# Evolution of apoptotic signalling pathways within lophotrochozoans

**DOI:** 10.1101/2023.12.11.571055

**Authors:** Helen R Horkan, Nikolay Popgeorgiev, Michel Vervoort, Eve Gazave, Gabriel Krasovec

**Author notes:** Authors for correspondence: GK, HRH. Deceased (Michel Vervoort). **Competing Interest Statement** The authors declare no competing interests. **Data Availability** All data needed to evaluate the conclusions in this study are present in the paper and the Supplementary Materials. Any requests can be addressed to the corresponding author GK and HRH. **Funding** GK was an Irish Research Council postdoctoral fellow (project GOIPD/2020/149) and is currently funded by the Fondation ARC pour la recherche sur le cancer (project ARCPOST-DOC2022070005318). HRH was a doctoral student in the Science Foundation Ireland Centre for Research Training in Genomic Data Science (grant no. 18/CRT/6214). EG is funded by Association pour la Recherche sur le Cancer (grant PJA 20191209482), and comité départemental de Paris de la Ligue Nationale Contre le Cancer (grant RS20/75-20). NP is funded by Institut Universitaire de France (IUF), Paris, France. **Author contributions** GK managed the project. GK and HRH wrote the manuscript. HRH, GK, MV, NP, and EG produced the data. All authors contributed to comment, discussion, and approved the manuscript.

## Abstract

Apoptosis is the main form of regulated cell death in metazoans. Apoptotic pathways are well characterised in nematode, fly and mammals, leading to a vision of the conservation of apoptotic pathways in metazoans. However, we recently showed that intrinsic apoptosis is in fact divergent among metazoans. In addition, extrinsic apoptosis is poorly studied in non-mammalian animals, making its evolution unclear. Consequently, our understanding of apoptotic signalling pathways evolution is a black-box which must be illuminated by extending research to new biological systems. Lophotrochozoans are a major clade of metazoans which, despite their considerable biological diversity and key phylogenetic position as sister group of ecdysozoans (*i.e.* fly, nematode), are poorly explored, especially regarding apoptosis mechanisms. Traditionally each apoptotic signalling pathway was considered to rely on a specific initiator Caspase, associated with an activator. To shed light on apoptosis evolution in animals, we explored the evolutionary history of initiator Caspases, Caspase activators and the BCL-2 family (which control mitochondrial apoptotic pathway) in lophotrochozoans using phylogenetic analysis and protein interaction predictions. We discovered a diversification of initiator Caspases in molluscs, annelids and brachiopods, and the loss of key extrinsic apoptosis components in platyhelminths, along with the emergence of a clade specific Caspase with an ankyrin pro-domain. Taken together, our data show a specific history of apoptotic actors’ evolution in lophotrochozoans, further demonstrating the appearance of distinct apoptotic signalling pathways during metazoan evolution.

**Significance statement:** Apoptosis, a form of programmed cell death, has been long studied in model organisms such as fly, mouse, and in humans. The restricted focus on these models has led to an overall view that the evolution of genes involved in apoptosis is highly conserved across all animals. The advent of next generation sequencing has led to a boom in the omics data available across the tree of life. Thanks to this, we explored the evolution of key genes involved in apoptosis in the clade Lophotrochozoa (*i.e.* molluscs, annelids, flatworms, brachiopods), one of the three large clades that make up bilaterian animals. We found a complex evolutionary history of apoptosis genes, with multiple losses, gains, divergences and redundancies, highlighting the value of exploring gene evolution and apoptotic mechanisms in Lophotrochozoans.

## INTRODUCTION

Apoptosis is a form of regulated cell death that sculpts the animal body during embryonic development and allows the removal of obsolete tissues or supernumerary cells (Jacobson et al. 1997). Present at the metazoan scale (Krasovec et al. 2019, 2021; AnvariFar et al. 2017; Ballarin et al. 2010; Kiss 2010; Jeffery & Gorički 2021; Vega Thurber & Epel 2007; Krasovec et al. 2022), apoptosis is defined by a conserved set of morphological features that depend on a multigenic family, the Caspases (Hengartner 2000; Cohen 1997). While Caspases are found in all animals, they have mainly been studied in nematode, fly and vertebrates, in which apoptotic signalling pathways are well described (Hengartner 2000; Fan et al. 2005; Steller 2008). Historically, two main apoptotic signalling pathways were described, currently known as extrinsic (formerly the death receptor pathway) and intrinsic apoptosis (formerly the mitochondrial pathway) (Galluzzi et al. 2018). In both cases, Caspases play a central role upstream (as initiator Caspases) and downstream (as executioner Caspases) of the apoptotic signalling pathways (Hengartner 2000; Kumar 2007; Fan et al. 2005). Importantly, each apoptotic pathway is characterised by a specific initiator Caspase, itself activated by a protein complex in which a specific activator is involved (Figure 1). While all Caspases are composed of the common P10 and P20 domains, initiator Caspases also contain an additional long pro-domain, either a CARD or DED domain, specific for intrinsic and extrinsic apoptosis, respectively.

**Figure 1:**
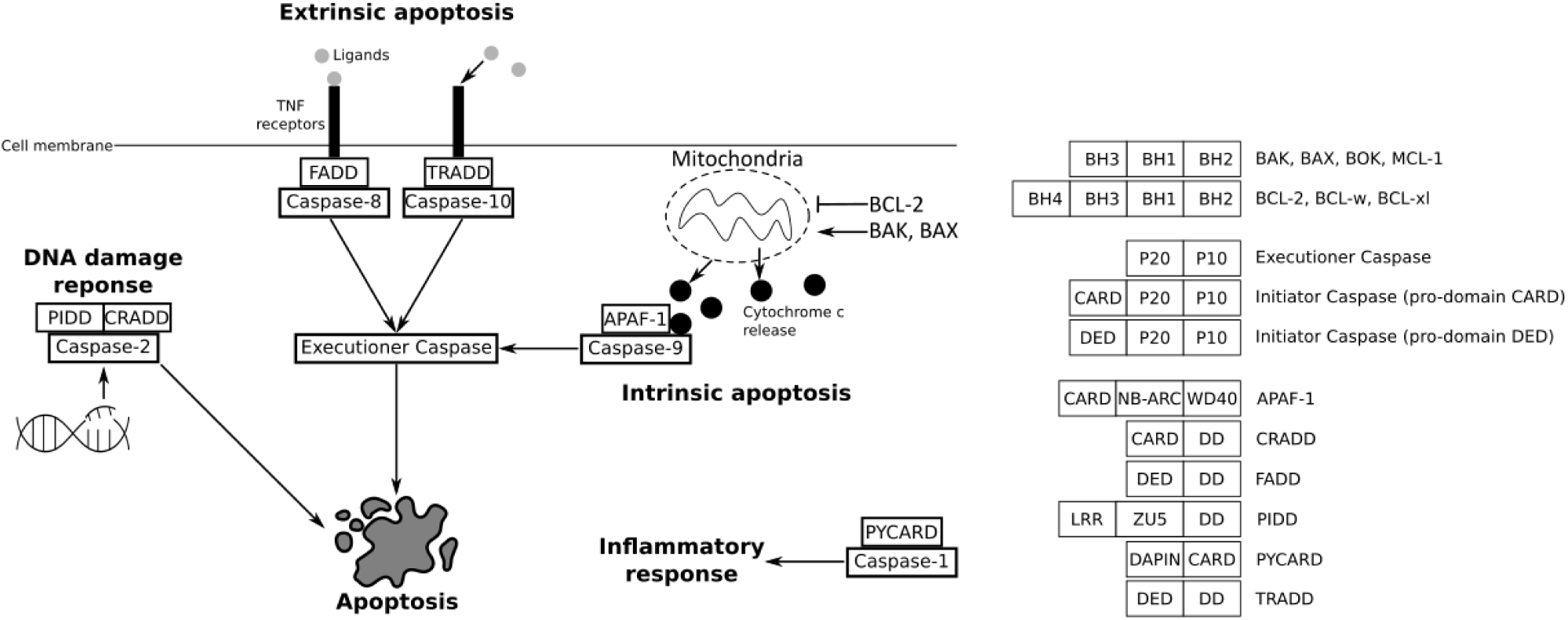
Main apoptotic pathways described in mammals. Initiator Caspases rely on a specific activator and are specific to each pathway. Extrinsic apoptosis depends on Caspase-8 or Caspase-10 and their activators, FADD and TRADD, respectively. Intrinsic apoptosis is controlled by the BCL-2 family (*i.e.* anti-apoptotic BCL-2, pro-apoptotic BAK or BAX) which can induce mitochondrial outer membrane permeabilisation, leading to cytochrome c release. Cytochrome c binds to initiator Caspase-9 and APAF-1 forming an activation complex, the apoptosome. Initiator Caspases induce activation of the common executioner Caspases downstream of the regulation pathways. Caspase-2, activated by CRADD, is involved in apoptosis induced by DNA damage. Caspase-1 is activated by PYCARD, leading to the inflammatory response. BCL-2 are characterised by the BH domains, with especially the BH4 for anti-apoptotic members. All Caspases have the P20 and P10 domains. In addition, initiator Caspases possess a long pro-domain, either a CARD or a DED. Activators APAF-1, CRADD, FADD, TRADD, PIDD, and PYCARD have a long domain which can be a CARD or a DED, in addition to a short death-domain (DD).

In mammals, intrinsic apoptosis is activated through mitochondrial outer membrane permeabilization (MOMP), a process controlled by BCL-2 family proteins (Figure 1) (Hengartner 2000). MOMP leads to the release of cytotoxic molecules such as cytochrome c into the cytosol. Cytochrome c forms the apoptosome complex with the specific adaptor protein APAF-1 (Apoptotic Protease Activating Factor 1) and initiator Caspase-9, leading to the activation of executioner Caspase-3 (Hengartner 2000; Galluzzi et al. 2018; Kalkavan & Green 2018). Key components of the intrinsic pathway are present in fly and nematode as well (Hengartner 2000; Steller 2008; Lettre & Hengartner 2006; Bender et al. 2012; Driscoll 1996). However, there are several fundamental mechanistic differences between these models and mammals. For instance, cytochrome c is crucial to trigger apoptosis in mammals, whereas it is dispensable in fly and nematode. Indeed, since BCL-2 members BAX and BAK have not been identified in *Drosophila melanogaster* and *Caenorhabditis elegans* genomes, MOMP does not occur. In *D. melanogaster* the APAF-1 homologue, DARK, oligomerizes into an eightfold apoptosome complex that does not require cytochrome c for assembly (Hengartner 2000; Steller 2008; Dorstyn et al. 2002). In addition, *D. melanogaster* apoptotic pathway depends on the removal of inhibition controlled by Grim, Reaper, and Hind (Steller 2008). These three proteins do not have homologues in other species. In *C. elegans*, the unique BCL-2 homolog CED-9 interacts with the APAF-1 homologue CED-4 (Hengartner 2000). When EGL-1 binds to CED-9, CED-4 is released, resulting in the formation of a homotetrameric complex that activates the Caspase CED-3 (Shi 2006). More recently, molecular characterization of the BCL-2 family in the placozoan *Trichoplax adhaerens* identified a *bona fide* BAX-like protein which performs MOMP *in vitro* (Popgeorgiev et al. 2020). Altogether, these findings suggest that although MOMP is ancient in metazoans, intrinsic apoptotic pathways are divergent and have evolved independently in most metazoan phyla. Interestingly, we have recently discovered that DRONC and CED-3, the initiator Caspases of intrinsic apoptosis in fly and nematode, respectively, are homologues (here orthologous, coming from speciation) to the vertebrate Caspase-2 but not to Caspase-9 (Krasovec et al. 2023). Caspase-9 is deuterostome specific, while Caspase-2 is ancestral and conserved among bilaterian animals. Interestingly, Caspase-2 in mammals is implicated in apoptosis induced by DNA damage, but not in the major intrinsic and extrinsic pathways (Figure 1) (Krumschnabel, Sohm, et al. 2009; Krumschnabel, Manzl, et al. 2009). Caspase-2 is activated by PIDD (P53-induced protein with a death domain) into the PIDDosome platform (Zhivotovsky & Orrenius 2005). Despite Caspase-2 being ancestral to bilaterians (Krasovec et al. 2023), the PIDD activator has not been investigated so far in non-mammals. In addition to Caspase-2 and -9, mammals possess other Caspases with a CARD pro-domain, the inflammatory Caspases, especially Caspase-1, which is able to cleave cytokines and thus trigger the inflammatory response (Sollberger et al. 2014). The activator PYCARD (PYD and CARD domain-containing protein) forms the inflammasome complex with Caspase-1, allowing its activation. Similar to PIDD, the presence of PYCARD in non-mammals has not yet been investigated (Figure 1).

Our knowledge on extrinsic apoptosis relies mainly on mammalian data due to its absence in fly and nematode. This pathway is activated by death receptors (*i.e.* TNFr – Tumor Necrosis Factor receptors) on cell membranes, themselves able to recruit activator proteins FADD (Fas-Associated protein with Death Domain) or TRADD (Tumor necrosis factor Receptor type 1-Associated Death Domain) into a DISC (Death-Inducing Signalling Complex), which activates Caspases-8 or 10, respectively (Figure 1) (Galluzzi et al. 2018; Fan et al. 2005). Despite this clear molecular cascade, extrinsic apoptosis is poorly described outside of mammals, except for some anecdotal studies, such as in cnidarians or amphioxus in which FADD can induce apoptosis in cultured cells (Steichele et al. 2021; Zhao et al. 2019; Yuan et al. 2010), or in the tunicate *Ciona*, where Caspase-8 is known to be responsible for apoptosis (Krasovec et al. 2024). Consequently, the evolution of extrinsic apoptosis is relatively unknown.

Taken together, it clearly appears that apoptotic signalling pathways’ evolution is a black box that needs to be revisited. Indeed, the lack of knowledge outside three key model species (mammals - deuterostome, fly and nematode - ecdysozoans) artificially creates biased inferences on apoptotic regulation pathways’ evolutionary history. Thus, the other major group of bilaterians, the lophotrochozoans, remains largely unexplored despite its tremendous diversity. This gap of knowledge creates a critical void and a comprehensive and global understanding of apoptotic signalling pathways evolution is definitively impossible to reach without exploring lophotrochozoan animals (*i.e.* molluscs, annelids, brachiopods, and platyhelminths).

In this study, we explored though comparative genomics the apoptotic network of four lineages of lophotrochozoans: molluscs, annelids, platyhelminths, and brachiopods. We conducted exhaustive reciprocal BLAST searches followed by phylogenetic analysis to identify apoptotic actors, especially Caspases, BCL-2 family proteins and activators such as FADD or APAF-1. We then analysed sequence evolution based on mutation rate and predicted initiator Caspase/activator interactions. Our results highlight an unexpected diversity and complexity of the apoptotic network within lophotrochozoans, characterised by the loss of the key intrinsic apoptosis activator APAF-1 in molluscs, annelids and brachiopods, in parallel with the loss of the extrinsic apoptosis actors in the platyhelminths, notably FADD and DED-Caspases. Brachiopods lost Caspase-2 despite being the conserved and ancestral initiator in bilaterians. Surprisingly, platyhelminths present Caspases with an ankyrin pro-domain, a unique feature never observed in vertebrates, fly or nematode. Annelids present several species-specific diversifications of initiator Caspases, characterised by unusual pro-domains, suggesting a variety of novel functions for these Caspases. Instead of a DED or a CARD pro-domain, we identified, among others, Caspases with a Z-binding, a kinase, or a PARP catalytic domain. The differential evolution between platyhelminths, molluscs, annelids and brachiopods suggests specific apoptotic pathways that have not yet been functionally described. Our work highlights the non-linear, divergent evolution of apoptotic pathways in animals and argues in favour of the study of lophotrochozoans as emerging biological models.

## RESULTS

### Evolutionary history of Caspases in lophotrochozoans

We investigated 60 genomes of lophotrochozoan species including 23 molluscs, 17 annelids, 1 brachiopod and 19 platyhelminths, to identify their Caspase (both initiators and executors) arsenals (Supplementary Table 1). We detected sequences of CARD-Caspases and DED-Caspases (initiators) as well as executioner Caspases. We also identified several Caspases presenting an ankyrin pro-domain (ANK-Caspases), an unexpected feature which has not previously been observed in either deuterostomes or ecdysozoans. To assess the main relationships between the different Caspases, we first conducted a phylogenetic analysis on all Caspases (both initiator and executioner Caspases) from a selection of 13 species representative of lophotrochozoan diversity (Figure 2, Supplementary Table 2). We chose this strategy of species sub-selection (while covering the whole diversity of Caspases types), as we know from our previous study on this gene family (Krasovec et al. 2023), that their short lengths combined with a high number of sequences leads to poorly resolved topologies. Consequently, this strategy allowed inclusion of all types of Caspases at the lophotrochozoan scale without over saturating the sequence number. The tree was rooted using a *Reticulomyxa filosa* (foraminifera) Caspase-like sequence, already successfully used for metazoan Caspases phylogenies (Krasovec et al. 2023; Kaushal et al. 2023). We first noticed a broad separation of the executioner Caspases *versus* all others. Indeed, they form a coherent group (PP=0.73), along with a clade composed of DED-caspases and ANK-caspases (plus a few additional unassigned sequences) as sister group. Most Caspases with DED pro- domains (DED-Caspases) form a strongly supported group (PP=0.99) composed of mollusc, annelid, and brachiopod sequences, but devoid of platyhelminths. ANK-Caspases form a maximally supported clade (PP=1.00) specifically restricted to platyhelminths, branching as a sister group to the *Haliotis diversicolor* sequence and close to DED-Caspases. The topology is characterised by a polyphyletic distribution of Caspases with a CARD pro-domain (CARD-Caspases), most of them being close to the root.

**Figure 2:**
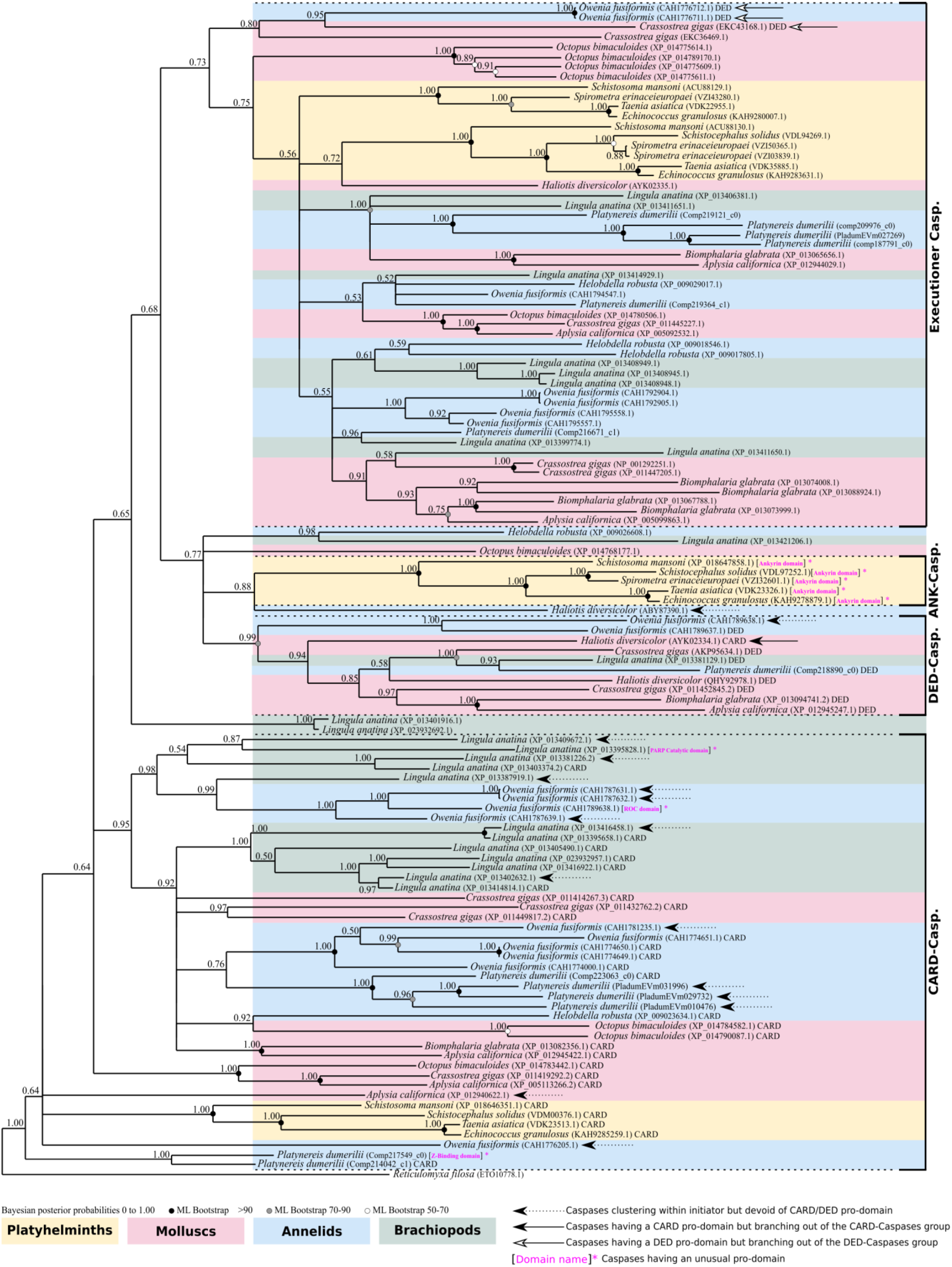
Topology of phylogenetic analysis of Caspases at the lophotrochozoan scale made by Bayesian inference method. Caspases with a CARD pro-domain (CARD-Caspases) present a paraphyletic distribution at the base of the tree. Among them, five sequences do not have a CARD domain (dotted black arrow). Caspases with a DED pro-domain (DED-Caspases) are restricted to molluscs and annelids, devoid of platyhelminth representatives. DED-Caspases form a monophyletic group, with only one from *Crassostrea gigas* out of the cluster (white arrow). Caspases with an ankyrin pro-domain (ANK-Caspases) form a monophyletic group specific to platyhelminths. Three CARD-Caspases (black arrows) are close to DED-Caspases and ANK-Caspases. Executioner Caspases are grouped into a maximally supported clade composed of sequences from molluscs, annelids, and flatworms. Outgroup is a Caspase-like of the foraminifera *Reticulomyxa filosa*. Node robustness correspond to posterior probabilities. Bootstrap correspond to robustness from maximum likelihood phylogeny which gave similar topology. The analysis includes 1 brachiopod, 5 platyhelminths, 3 annelids, and 4 molluscs. All caspases analysed have the common P20 and P10 domains. The presence of a classical CARD or DED pro-domain is indicated next to the sequence name. Caspases with a pro-domain other than CARD or DED are indicated in brackets and in magenta. Phylogeny was conducted on the full length of amino acid sequences containing the pro-domain and the common P20 plus P10 domains.

Due to their pivotal role, we next decided to focus more specifically on initiator Caspases (Supplementary Table 3). Due to the short length of Caspase sequences, this strategy allowed us to reduce the number of sequences per species and secondarily to increase the sampling to 50 species of lophotrochozoans to conduct a more in-depth analysis of this lineage. We reconstructed their phylogenetic relationships and identified three main groups, consistent with our previous analysis: ANK-Caspases, DED-Caspases, and CARD-Caspases (Caspase-2 and Caspase-Y) (Figure 3). ANK-Caspases and DED-Caspases are sister groups and together form a well-defined (PP=0.61) clade. The ANK-Caspases exhibit maximum support (PP=1.00) and are specifically restricted to parasitic platyhelminths. Indeed, both the free-living *Schmidtea mediterranea* and *Dugesia japonica* do not have ANK-Caspases. In contrast, the DED-Caspase group (PP=0.95) is composed of mollusc, annelid, and brachiopod sequences, together with two non-parasitic, early diverging platyhelminths, *Dugesia japonica* and *Schmidtea mediterranea*. Consequently, platyhelminths which have DED-Caspases do not have ANK-Caspases and *vice versa*. This suggests a specific Caspase evolution for parasitic platyhelminths.

**Figure 3:**
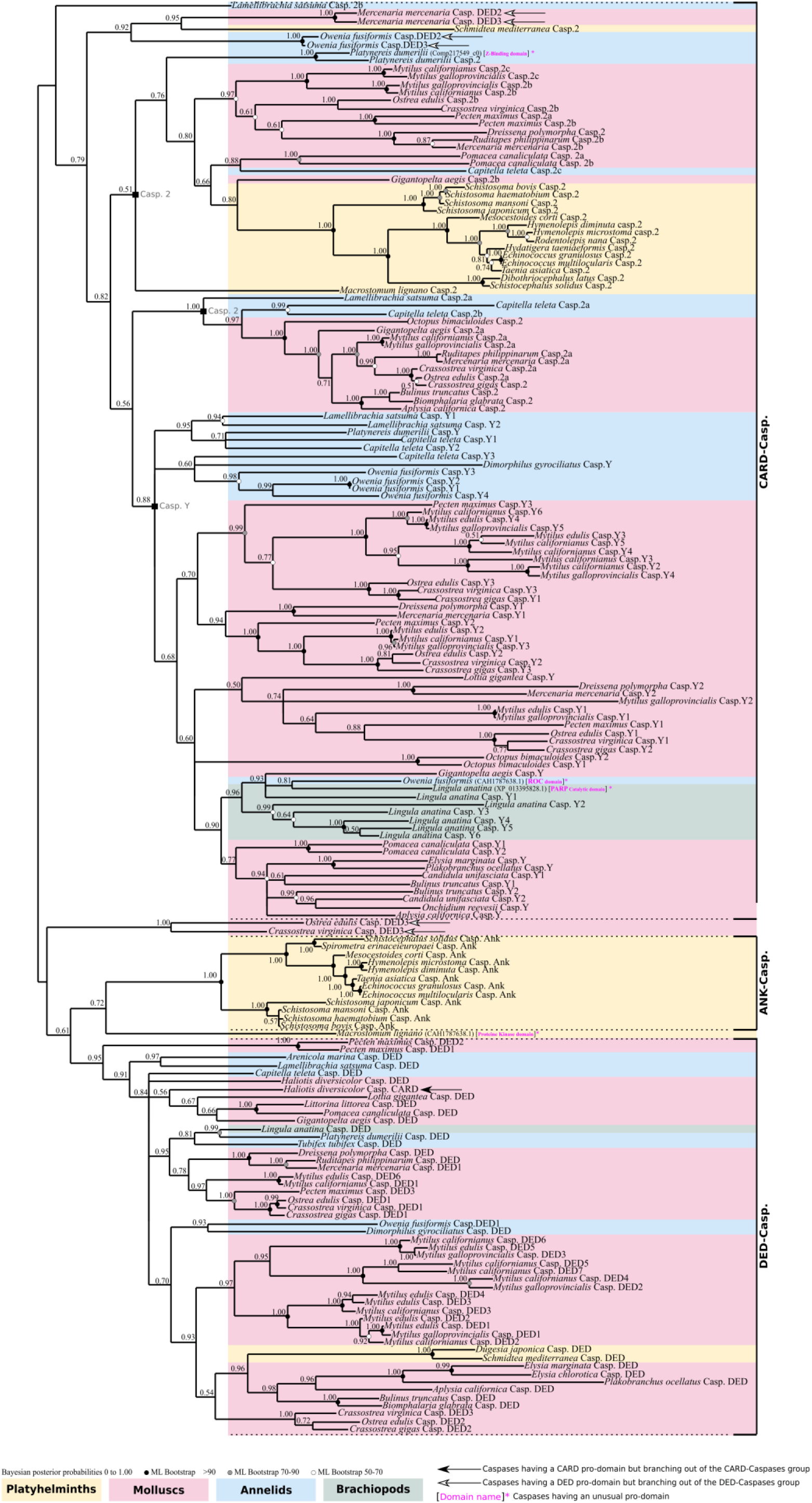
Unrooted phylogenetic analysis of initiator Caspases made by Bayesian inference method. Three distinct clades are present. The CARD-Caspase comprises the Caspase-Y specific to molluscs, annelids and brachiopods, devoid of platyhelminth sequences. The second major group of CARD-Caspases is the Caspase-2 which is split into two sub-groups and composed of representatives for each lophotrochozoan clade (molluscs, annelids, brachiopods, platyhelminths). DED-Caspases are monophyletic and composed of molluscs, annelids, brachiopod, and two non-parasitic platyhelminths. ANK-Caspases are monophyletic, forming a clade sister to the DED-Caspases and are restricted to parasitic platyhelminths. Several Caspases present unusual pro-domains (brackets, domain names in magenta). The sister group of *Platynereis dumerilii* Caspase-2 is one of them, characterised by a Z-repeat-binding pro-domain. A *Macrostomum lignano* Caspase with a protein kinase domain branches as sister to ANK-Caspases. Inside the Caspase-Y group, brachiopod *Lingula anatina* has a Caspase with a PARP catalytic domain, in addition to *Owenia fusiformis* which encodes one with a ROC pro-domain. Node robustness correspond to posterior probabilities. Bootstrap correspond to robustness from maximum likelihood phylogeny which gave similar topology. The analysis includes 1 brachiopod, 18 platyhelminths, 7 annelids, and 24 molluscs. Phylogeny was conducted on the full length of amino acid sequences containing the pro-domain and the common P20 plus P10 domains.

Both Caspase-2 and -Y possess a CARD pro-domain and consequently belong to the CARD-Caspase type. Caspase-Y is a strongly supported (PP=0.88) group which we already identified in a previous study and is composed of sequences from molluscs and annelids (Krasovec et al. 2023). Here, we show that Caspase-Y is present in brachiopods as well and, interestingly, that there are several duplications in this species harbouring 6 Caspase-Y paralogs. The Caspase-2 encompasses sequences from molluscs and annelids, as well as a strongly supported clade of platyhelminths sequences (PP=1.00). Most lophotrochozoan animals possess a Caspase-2, (with the exception of the brachiopod *Lingula anatina* due to a secondary loss), which is consistent with our previous work showing that this Caspase is conserved and ancestral in bilaterians.

In addition to the expected Caspase types, we discovered four specific Caspases with unexpected long pro-domains, which have never been identified before (Supplementary Figure 1). The platyhelminth *Macrostomum lignano* presents a Caspase with a protein kinase like pro-domain, positioned as sister to the ANK-Caspase group (Figure 3). We did not detect any other Caspases with an unexpected pro-domain in platyhelminths. In addition, we identified a sequence from the brachiopod *Lingula anatina* within the Caspase-Y clade that has a PARP catalytic and alpha-helical pro-domain instead of the CARD pro-domain. Similarly, annelids *Platynereis dumerilii* has a Caspase with a Z-binding pro-domain which is a paralogue to its Caspase-2 (Figure 3), and *Owenia fusiformis* encodes a Caspase with a ROC domain profile as a pro-domain.

To explore the history of Caspase pro-domains acquisition and potentially discover early emerging initiator Caspase types, we artificially constructed three topologies corresponding to three potential scenarios of evolution, firstly with ANK-Caspases as sister to [DED-Caspases + CARD-Caspases], secondly DED-Caspases as sister to [ANK-Caspases + CARD-Caspases], and thirdly CARD-Caspases as sister to [DED-Caspases + ANK-Caspases] (Figure 4). Next, we conducted topology tests to evaluate the most likely scenario (Nguyen et al. 2015; Kishino & Hasegawa 1989; Shimodaira 2002; Shimodaira & Hasegawa 1999). Interestingly, regardless of the statistical test used, there is no topology that appears to be the most likely, making it difficult to hypothesise on an ancestral type of Caspase, arguing instead for a complex evolutionary model, but also confirming the ancestral presence of both DED-Caspases and CARD-Caspases among lophotrochozoans (Figure 4).

**Figure 4:**
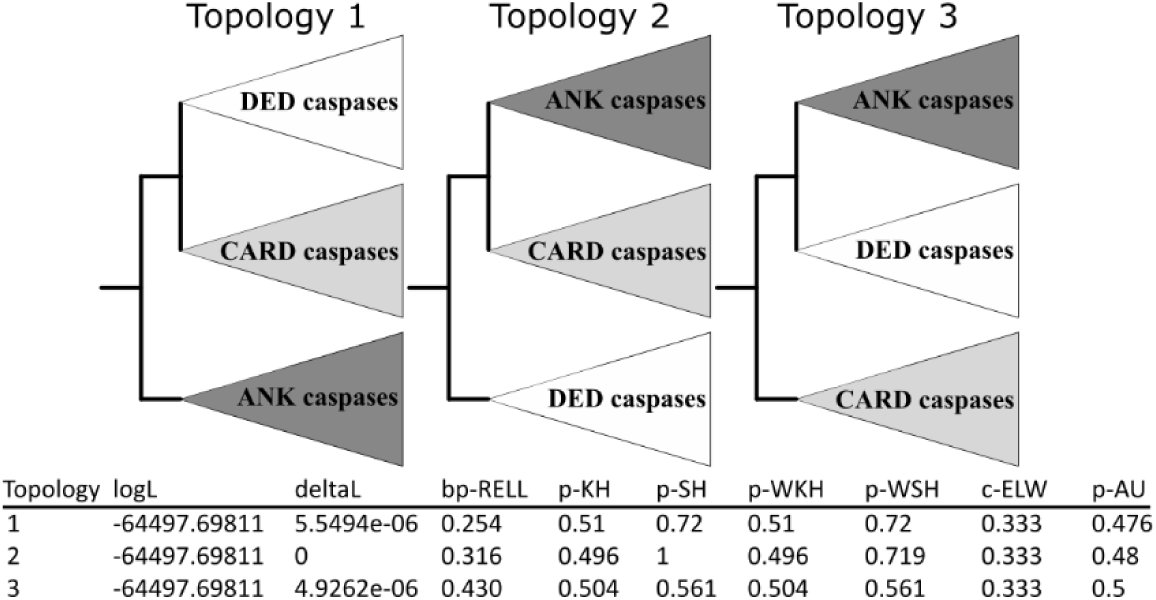
Topology tests conducted with IQ-TREE between initiator caspase evolution models. Three artificial topologies were built composed of three clades containing all DED-Caspases, CARD-Caspases, and ANK-Caspases, respectively. Topology tests do not allow identification of a single most-likely scenario, preventing the formation of a hypothesis on the emergence of an ancestral pro-domain within the Caspase family. Tests are noted as p-KH for Kishino-Hasegawa test p-value (Likelihood Method), p-SH for Shimodaira-Hasegawa test p-value, and p-AU for approximately unbiased test p-value. deltaL: logL difference from the maximal logl in the set. bp-RELL: bootstrap proportion using RELL method. c-ELW: Expected Likelihood Weight. p-WKH: p-value of weighted KH test; p-WSH: p-value of weighted SH test. A tree is rejected if its p-value < 0.05. Tests p-values do not allow to discriminate a more likelihood topology between the three scenarios.

Taken together, our topology indicates that Caspase-Y are a group of CARD-Caspases specific to [molluscs + annelids + brachiopods]. Brachiopods lost the ancestral Caspase-2. ANK-Caspases are specific to parasitic platyhelminths which have lost DED-Caspases. Conversely, free living platyhelminths do not have an ANK-Caspase and retain the DED-Caspase. The proximity between ANK-Caspases and DED-Caspases could suggest accumulation of divergence of the DED domain of ancestral platyhelminths during evolution.

### Diversity of Caspases in annelids

The presence of unusual Caspase pro-domains in some annelid species caught our attention. Therefore, we decided to perform an in-depth investigation of Caspases in this lineage, thanks to an increase in species sampling (Supplementary Table 4). First we conducted a phylogenetic analysis on a representative set of annelids comprising of 2 early branching species, 1 Errantia and 11 Sedentaria, according to recent annelid phylogenies (Martín-Durán et al. 2021; Lewin et al. 2024). The topology resulting from this analysis showed unresolved nodes, especially regarding CARD-Caspases, indicating that we were close to sequence saturation in terms of available phylogenetic information, as we previously experienced with Caspases (Supplementary Figure 2). This could be exacerbated by the overrepresentation of Sedentaria, a group from which most of the annelid sequenced genomes originates and encompassing species with a large number of caspases. Thus, we conducted a second phylogenetic analysis with modified sampling by removing the Sedentaria *Lamellibrachia satsuma* which encodes a large set of Caspases, and we added the Errantia *Alitta virens* (Figure 5). Globally, Caspases cluster in a coherent manner, similarly to our previous studies. The majority of executioner Caspases cluster together (PP=0.58). Most DED-Caspases are in a strongly supported cluster (PP=0.98), with the exception of two *Owenia fusiformis* DED-Caspases. CARD-Caspases are closely related, clustering in a strongly supported group (PP=0.83). CARD-Caspases are divided into the two types we identified previously: Caspase-2 and Caspase-Y.

**Figure 5:**
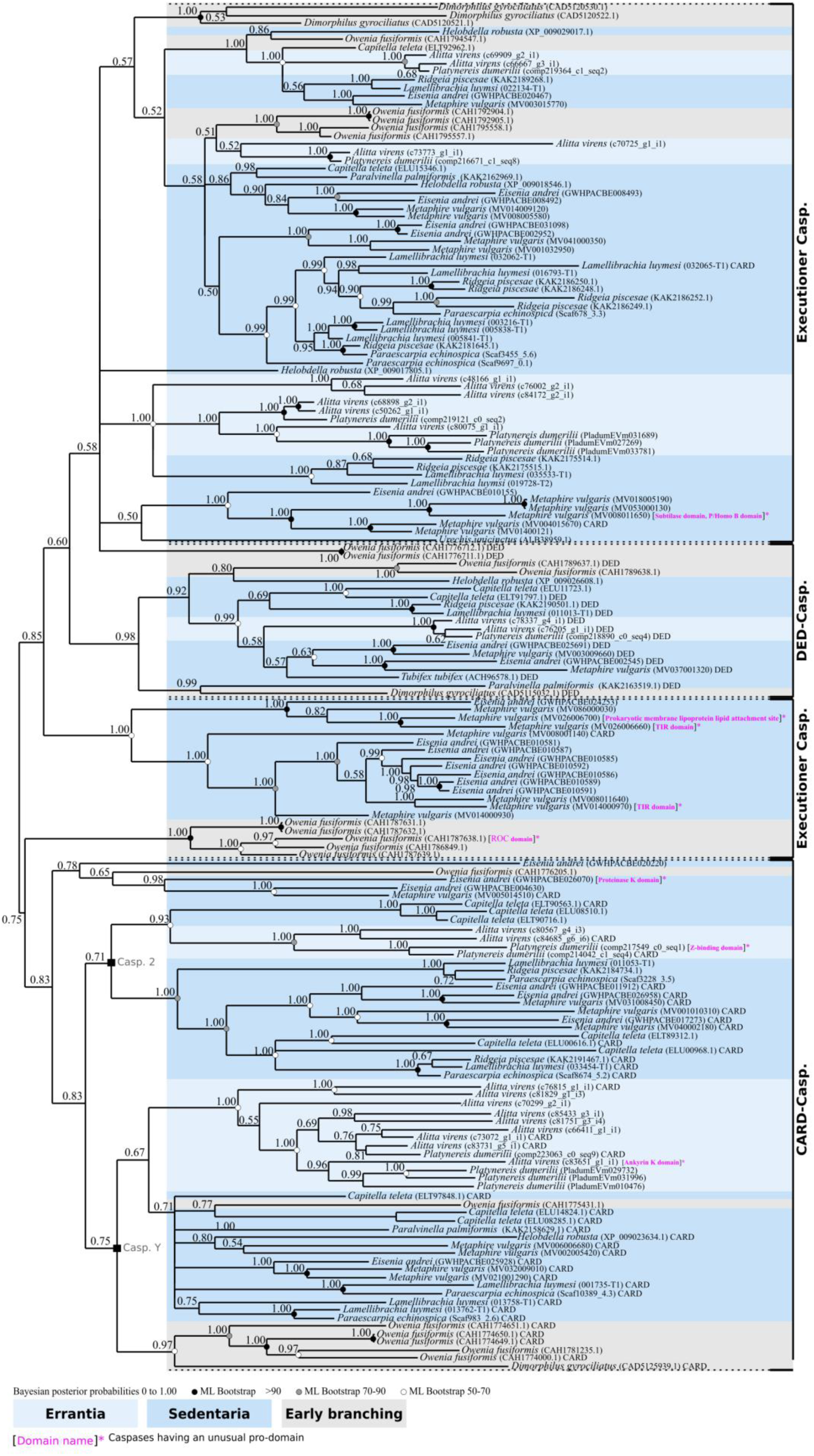
Topology of phylogenetic analysis of Caspases of annelids made by Bayesian inference method. Most executioner Caspases cluster together. Caspases with a DED pro-domain (DED-Caspases), except for two *Owenia fusiformis* sequences, form a monophyletic group. Caspases with a CARD pro-domain (CARD-Caspases) are split between Caspase-Y and Caspase-2. A few Caspases without long pro-domains branch inside executioner Caspases. Annelids cluster in a coherent way according to species phylogeny: the Errantia sequences form several paralogous groups, similarly to Sedentatia sequences. Early branching annelids are represented by the paleoannelid *Owenia fusiformis*, and Dinophiliformia *Dimorphilus gyrociliatus*. Globally, each species encodes DED-Caspases, CARD-Caspases, and executioner Caspases. In addition, several Caspases present unusual pro-domains which are: a Caspase with a protein Kinase domain (*Eisenia andrei*), with a prokaryotic membrane lipoprotein lipid attachment site profile (*Metaphire vulgaris*), a subtilase domain (*Metaphire vulgaris*), two Caspases having a TIR domain (*Metaphire vulgaris*), one with a Z-binding domain (*Metaphire vulgaris*), one with a ROC domain profile (*Owenia fusiformis*), and one with a ankyrin domain (*Alitta virens*). The presence of a classical CARD or DED pro-domain is indicated next to the sequence name. Caspases with a pro-domain other than CARD or DED are indicated in brackets and in magenta. Bootstrap values correspond to robustness from maximum likelihood phylogeny which gave a similar topology. The analysis includes 10 Errantia, 2 Sedentaria, and 2 early branching annelids. Phylogeny was conducted on the full length of amino acid sequences containing the pro-domain and the common P20 plus P10 domains.

Early branching annelid sequences (*Owenia fusiformis* and *Dimorphillus gyrocilliatus*) are distributed along the topology without branching at the base of groups composed of [Errantia + Sedentaria], possibly illustrating their uncertain phylogenetic position. Sequences from Errantia (*Platynereis dumerilii* and *Alitta virens*) are clustered within each Caspase group, indicating their orthologous relationship. Regarding Sedentaria species, most of them cluster together, especially Clitellata (such as *Lamellibrachia luymesi* or *Ridgeia piscesae*) which form several monophyletic sub-clades. Our sampling and the resulting phylogeny indicate significant variation in the Caspase family among annelid species. Some annelids have few Caspase members, such as *Tubifex tubifex* or *Helobdella robusta* with one and four Caspases, respectively. Conversely, other species present an expansion of the family with numerous paralogs such as *Metaphire vulgaris* which encodes 27 Caspases. This disparity could be due to genome duplication in this species, which has resulted in numerous paralogs for different genes (Jin et al. 2020).

As hypothesised, we discovered species-specific diversification of Caspases with peculiar pro-domains (Figure 5, Supplementary Figure 1). In addition to Caspases with a Z-binding and ROC pro-domains in *Platynereis dumerilii* and *Owenia fusiformis*, respectively (see above), we detected a protein kinase domain Caspase in *Eisenia andrei*. In *Metaphire vulgaris* we detected two TIR domain Caspases, a Caspase with a prokaryotic membrane lipoprotein lipid attachment site profile as the pro-domain, and a Caspase with a particularly long pro-domain comprising both a subtilase domain and a P/Homo B domain (Figure 5). *Alitta virens* encodes an ANK-caspase, similar to those found in platyhelminths, which is likely to be a convergent feature. Most of these particular Caspases are paralogous to sequences from the same species having a classical domain architecture, suggesting species-specific duplications. After the duplication event, the appearance of unexpected pro-domains may be the result of gene specific evolution. Taken globally and according to current knowledge, our results indicate that annelids may be the animal group with the greatest diversity and inter-species variation in the Caspase family.

### Losses and gains of activators is consistent with initiator Caspases inventory

Initiator Caspases are activated by specific platforms that depend on an activator which possesses a protein domain similar to their corresponding initiator Caspases pro-domain (Figure 1). Thus, the CARD domain of APAF-1 binds to the CARD pro-domain of Caspase-9 (mammals) or Caspase-2 (fruit flies and nematode) (Hengartner 2000; Bratton & Salvesen 2010, 1; Zou et al. 1997, 1). Mammalian Caspase-2 is activated by CARD domain containing PIDD (named CRADD, also known as RAIDD) (Lin et al. 2000; Tinel & Tschopp 2004; Duan & Dixit 1997) and Caspase-1 by the PYD and CARD domain-containing protein activator (PYCARD) (Sutterwala et al. 2006). Similarly, FADD and TRADD (containing DED domains) activate initiator Caspase-8 and 10 in the DISC complex, respectively (Fan et al. 2005; Pennarun et al. 2010).

We conducted an exhaustive inventory of Caspase activators within lophotrochozoan lineages (Figure 6A). Interestingly, we found a FADD homologue in the [molluscs + annelids + brachiopods] group, which is consistent with the presence of DED-Caspases in these taxa (Figure 6A1, Supplementary Table 5). However, FADD was not detected in platyhelminths. Our FADD phylogenetic analysis gave rise to a topology that is globally consistent with the species tree with molluscs grouped in a maximally supported monophyletic clade (PP=1.00) and split into gastropods and bivalves, in addition to the cephalopod. Annelid and brachiopod sequences branch first in a paraphyletic distribution, and appear as sister group to molluscs. The relative topology consistency between sequences and species suggests gene conservation in these groups. Conversely, APAF-1 was only detected in platyhelminths, but, consistently with our previous work (Krasovec et al. 2023), we did not identify any APAF-1 in [molluscs + annelids] which confirms the loss of this activator in these taxa (Figure 6A2, Supplementary Table 6). Here, we did not detect APAF-1 in the brachiopod *Lingula anatina* as well. The APAF-1 topology is congruent with the species tree and all clades are maximally supported (PP=1.00), with the monophyletic cestode platyhelminths branching with the sister group of trematodes. At the basis of the topology are the first branching triclad platyhelminths. Next, due to the absence of APAF-1, we investigated the potential activators for [molluscs + annelids + brachiopods] CARD-Caspases and discovered that these species possess a CRADD homologue, which is absent in platyhelminths (Figure 6A3, Supplementary Table 7). Phylogenetic analysis of the CRADD genes suggests a conservation of the gene, as the topology is in accordance with species relationships: molluscs are monophyletic (PP=0.56) and encompass gastropods (PP=0.83) and bivalves (PP=0.99), with the cephalopod as sister of the gastropods. Brachiopod and annelid sequences are distributed between the root of the tree and molluscs. Lastly, we were unable to detect any PIDD, PYCARD or TRADD (which activate Caspase-2, Caspase-1, and Caspase-10 in mammals, respectively) despite an exhaustive search in all available genomes/transcriptomes/proteomes of lophotrochozoan species from NCBI database. This indicates the potential vertebrate specificity of the PIDD and PYCARD activators, as it was proposed for TRADD (Zmasek & Godzik 2013).

**Figure 6:**
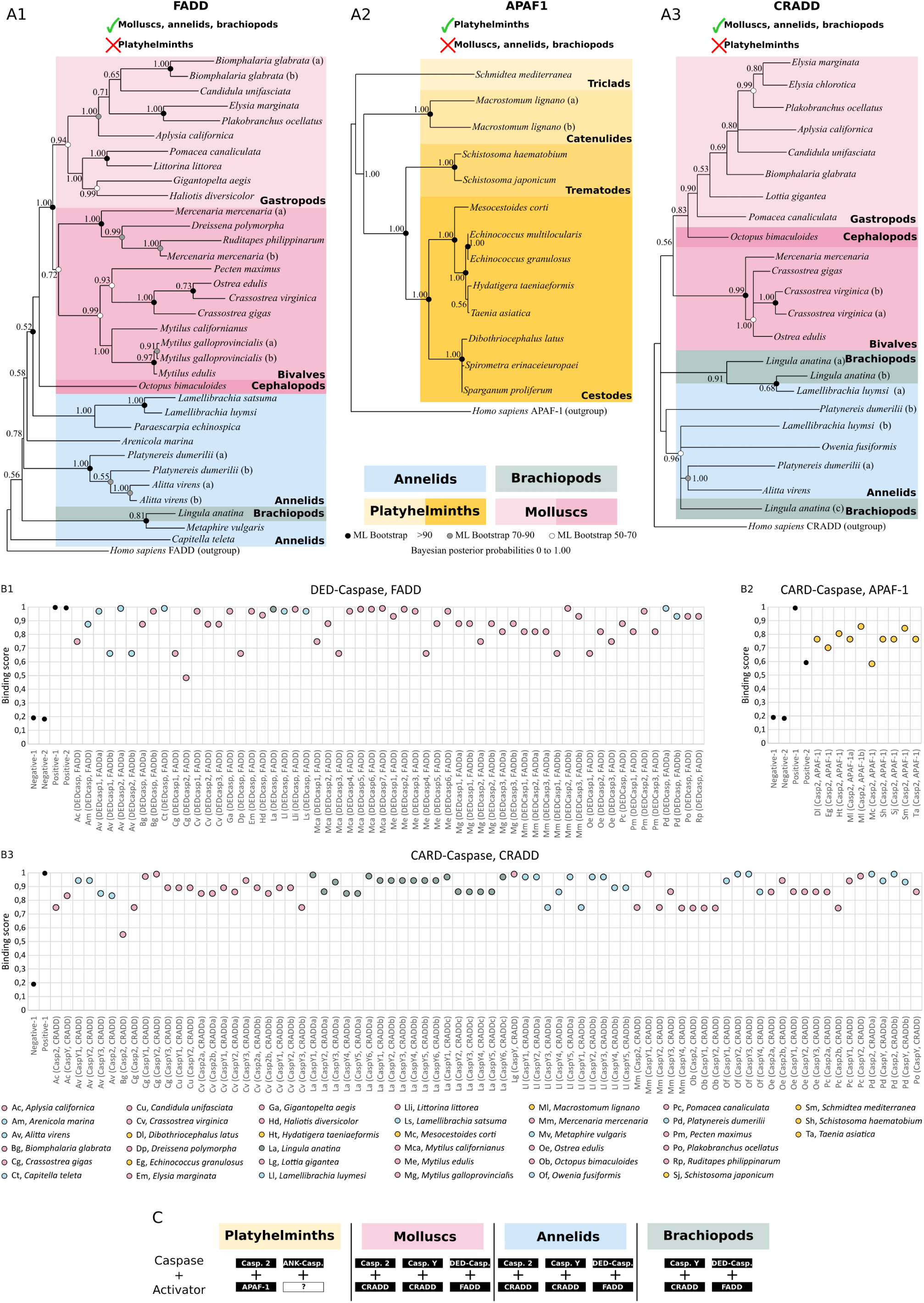
Analysis of initiator Caspase activators. **A**, phylogenetic analysis of initiator Caspase activators at the lophotrochozoan scale made by Bayesian inference method with human sequences as outgroup for FADD (A1), APAF-1 (A2), and CRADD (A3). FADD (containing a DED domain) presents a topology relatively consistent with species relationship. Molluscs are monophyletic and split between gastropods and bivalves, with also the presence of a cephalopod. Annelids and brachiopods branch early. The FADD analysis includes 1 brachiopod, 8 annelids, and 20 molluscs and was conducted on the full length of amino acid sequences containing the DED and Death domains. No FADD was detected in platyhelminths. APAF-1 is absent in molluscs, annelids and brachiopods but was detected in platyhelminths. The analysis includes 12 species and was conducted on the full length of amino acid sequences containing the CARD and the NB-ARC domains. The sequence topology is similar to platyhelminth species relationships, indicating a conservation of this gene within the group. CRADD in absent in platyhelminths but present in molluscs, annelids and brachiopods, and shows a topology relatively consistent with the species tree, similarly to FADD. The CRADD analysis includes 1 brachiopod, 4 annelids, and 13 molluscs and was conducted on the full length of amino acid sequences containing the CARD and the Death domains. Node robustness correspond to posterior probabilities. Bootstrap correspond to robustness from maximum likelihood phylogeny which gave similar topology. **B**, *in silico* test of binding affinities made with PSOPIA software between DED-Caspases with FADD (B1), CARD-Caspases with APAF-1 (B2), and CARD-Caspases with CRADD (B3). DED domains from FADD and DED-Caspases are predicted to bind, suggesting an activation of extrinsic apoptosis. CRADD can theoretically interact with all CARD-Caspases of these species including Caspase-2 and Caspase-Y. Predictions indicate that APAF-1 interacts with platyhelminths Caspase-2. **C**, theoretical couples of initiator Caspase/activator identified in lophotrochozoans resulting from analysis conducted in **A** and **B**

Next, we explored the theoretical initiator Caspase/activator binding interactions that could trigger their activation (Figure 6B). To do so, we evaluated interaction scores (S-score) using PSOPIA software, which is specifically designed to predict protein-protein interactions. First, we calculated the S-scores between DED-Caspases and FADD for all species where both proteins are present. Consistent with the literature, our results suggest that mollusc, brachiopod and annelid FADD could activate the corresponding DED-Caspases (Figure 6B1). We found the same result with the APAF-1 activator and CARD-Caspases from platyhelminth species (Figure 6B2). Interestingly, the S-scores for CRADD and CARD-Caspases of [molluscs + annelids + brachiopods] also indicate a high binding probability, with a global preference for Caspase-2 rather than Caspase-Y (Figure 6B3), except for a few species such as *Owenia fusiformis*.

Our results indicate a consistent relationship between the inventory of initiator Caspases and potential interactors, allowing the proposition of activation pairs (Figure 6C), according to specific loss within each group of lophotrochozoans that we focused on.

### BCL-2 family is conserved with non-linear evolution within lophotrochozoans

Finally, we focused on the BCL-2 family which are crucial regulators of MOMP (Chipuk et al. 2010). Within their primary structure, BCL-2 proteins possess one or more conserved sequences, called BCL-2 homology or BH motifs. Based on their number, they are divided into multi-motif and “BH3-only”. Multi-motif BCL-2 can be either pro-apoptotic including BAX, BAK and BOK which allow cytochrome c release following MOMP in mammals (Kalkavan & Green 2018; Banjara et al. 2020; Moldoveanu & Czabotar 2020) or anti-apoptotic such as BCL-2, BCL-XL and MCL-1. Notably, multi-motif pro- and anti-apoptotic BCL-2 members have conserved 3D structure, suggesting a common evolutionary origin. On the other hand, BH3-only proteins represent a polyphyletic group with pro-apoptotic activities (Aouacheria et al. 2013). Thus, BCL-2 family evolution is complex, with phylogenetic analyses usually producing unclear topologies (Krasovec et al. 2023). Importantly, the short length of BCL-2 proteins restricts the number of informative sites for the phylogenetic analyses. To circumvent this, we conducted phylogenetic analysis on the BCL-2 family by incorporating only 2 molluscs (*Biomphalaria glabrata* and *Crassostrea gigas*), 1 annelid (*Platynereis dumerilii*), 1 Brachiopod (*Lingula anatina*), and 4 platyhelminths (*Schistosoma mansoni*, *Taenia asiatica*, *Echinococcus granulosus*, and *Schmidtea mediterranea*) (Figure 7, Supplementary Table 8), to represent the diversity of lophotrochozoans without increasing the sequence number. Despite these precautions, the topology obtained by Bayesian inference is of low-resolution with numerous unresolved nodes, in line with what we previously observed (Krasovec et al. 2023). Conversely, the topology arising from maximum likelihood method allows the identification of BCL-2 family relationships (Figure 7) but with low node robustness. The topology suggests three main pro-apoptotic clades corresponding to BAK, BAX, and BOK. Molluscs, annelids and brachiopods possess all of them, in addition to numerous specific duplications (*i.e*. *Crassostrea gigas* BAX or *Lingula anatina* BAK). Conversely, platyhelminths only have the BAK pro-apoptotic, suggesting a loss of both BAX and BOK. Four anti-apoptotic clades are identified, one composed of the MCL1 restricted to molluscs and close relatives, and three BCL-2-like groups. We named them BCL-2 like as evolutionary relationships with BCL-2 are variable and difficult to establish. The relatively clear separation between pro- and anti-apoptotic genes clarified which species have a complete BCL-2 inventory (both functions represented). All species from our sampling have both pro- and anti-apoptotic BCL-2, but the number is variable. In particular, the three parasitic platyhelminths have only three BCL-2 representatives, while the diversity is high in molluscs and close relatives. Interestingly, there are no differences in BCL-2 family composition between parasitic and non-parasitic platyhelminths, whereas this is the case for the Caspases repertoire. Thus, the composition of the ancestral BCL-2 repertoire remains elusive.

**Figure 7:**
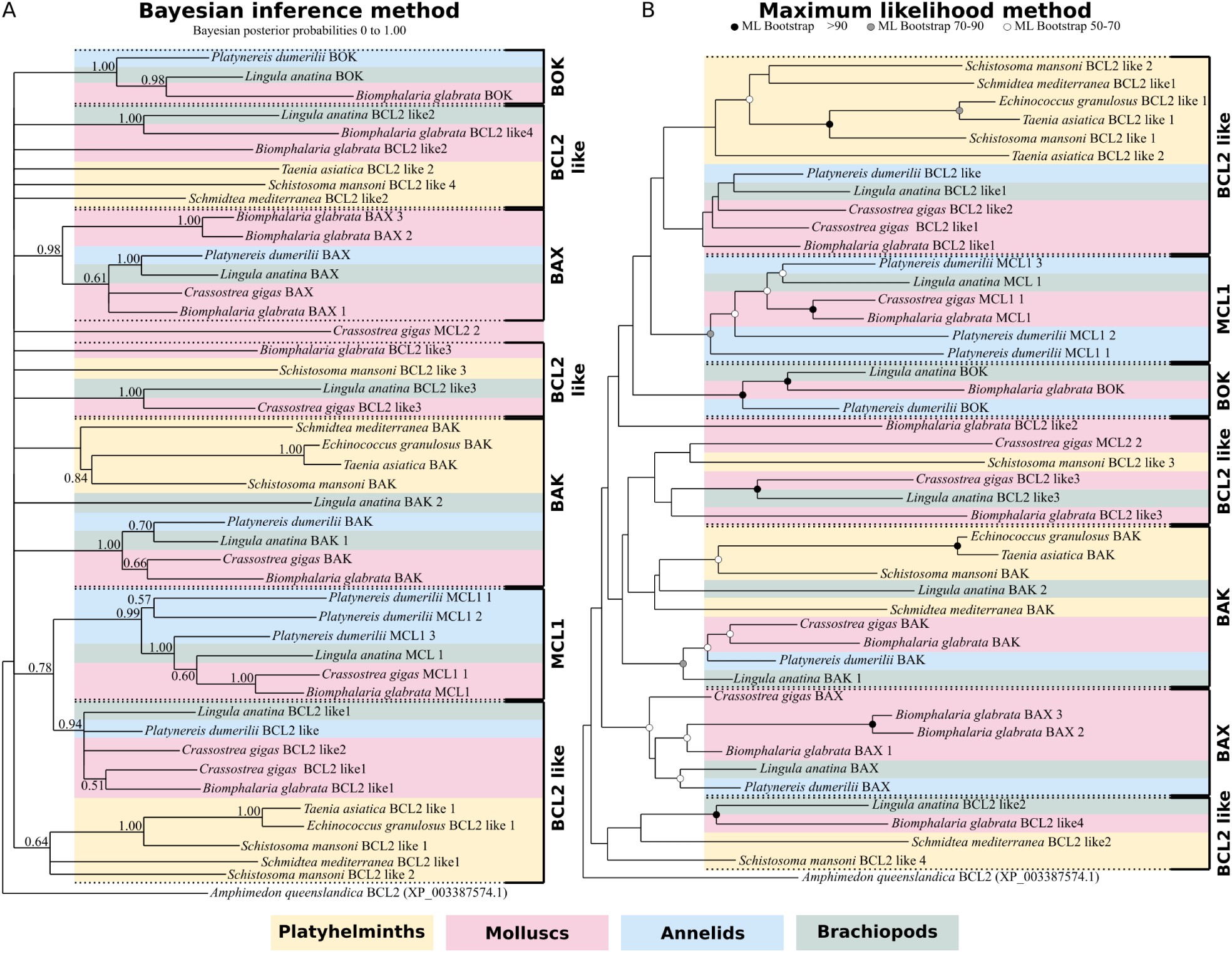
Topologies of the BCL2 family at the lophotrochozoan scale obtained by Bayesian inference (BI) and maximum likelihood (ML) methods. Node robustness corresponds to 500 bootstraps and posterior probabilities in ML and BI analyses, respectively. A, the BI tree exhibits numerous unresolved nodes, but the topology allows confirmation of the evolutionary relationships of several groups including BAX or MCL1, among others. **B**, the ML tree is well resolved, and sequences group into several monophyletic groups composed of either pro-apoptotic or anti-apoptotic BCL2 members. Interestingly, the platyhelminth network is very limited, composed of a unique proapoptotic (BAK), and a small number of BCL2 sequences, likely antiapoptotic. Conversely, molluscs, brachiopods and annelids have the three classical pro-apoptotic BAK, BAX, and BOK, and a large number of antiapoptotic genes including MCL1 and several BCL2 like genes. The analysis includes 1 brachiopod, 4 platyhelminths, 1 annelid, and 2 molluscs and was conducted on the full length of amino acid sequences containing the BH3, BH1 and BH2 domains.

## DISCUSSION

### Molluscs, annelids and brachiopods present specific apoptotic network architecture

Molluscs and their close relatives’, annelids and brachiopods, have an apoptotic network that is distinct from mammals and ecdysozoans. Molluscs possess a large diversity of BCL-2 which likely leads to MOMP (conversely to ecdysozoans), as cytochrome c release from mitochondria to cytoplasm has been observed in *Helix pomatia* and *Crassostrea gigas* (Pirger et al. 2009; Li et al. 2017). The loss of APAF-1, which is the shared component of intrinsic apoptosis, suggests a specific modality of CARD-Caspase activation. In contrast to ecdysozoans, Caspase-2 (and also Caspase-Y) in molluscs may be activated by CRADD similarly to what occurs in mammals, despite being more phylogenetically distant (Tinel & Tschopp 2004; Duan & Dixit 1997). This makes these animals the only ones currently known to have a potential intrinsic apoptosis which is not based on an apoptosome, but resulting potentially from Caspase-2 activation by CRADD and cytochrome c. The presence of only one CARD domain containing activator in these animals certainly induced an evolutionary constraint leading to the conservation of CRADD. The loss of this unique activator may lead to the inability to trigger apoptosis and could be deleterious for development and homeostasis. The dN/dS is a ratio usually used to evaluate selection pressure on genes during evolution (Ota & Nei 1994; Nei & Gojobori 1986). A ratio lower than 1 indicates a prevalence of synonymous mutations (without impact on protein sequence, and likely protein functions) resulting from a purifying selection. Conversely, a ratio higher than 1 corresponds to an accumulation of non-synonymous mutations which likely altered the amino acid composition of proteins, that may indicate positive selection. The ratio we found for lophotrochozoan CRADD less than 1 (dN/dS=0.91) confirms purifying selection on these genes (Supplementary Table 9).

Caspase-Y is restricted to the [molluscs + annelids + brachiopods], indicating a diversification of CARD-Caspases in this clade. Interestingly, CARD-Caspase diversification in mammals led to a functional specialisation between apoptosis and immunity. Caspase-1 is involved in the inflammatory response, while apoptosis is regulated by other CARD-Caspases such as Caspase-2 or Caspase-9 (Sollberger et al. 2014). In molluscs, it has been shown that Caspase-2 of *Crassostrea angulata* is expressed during the apoptotic event of larvae metamorphosis, suggesting a primordial role of this Caspase in apoptosis (Yang et al. 2015). Importantly, Caspases have been reported to be fundamental in immune response to various stress and inflammation in several mollusc species (Kiss 2010; Romero et al. 2015; Sokolova 2009; Sokolova et al. 2004). Therefore, we may hypothesise that the diversification of CARD-Caspase in molluscs and their close relatives’, annelids and brachiopods, may have induced separate functions as well. Caspase-2 could be involved in apoptosis, and Caspase-Y in immunity. Interestingly, we did not detect Caspase-2 in the brachiopod *Lingula anatina*, although this Caspase is known to be ancestral and highly conserved in bilaterians. This rather unique feature is only shared with the ambulacraria (hemichordates + echinoderms). This loss could be due to the diversification of Caspase-Y (6 paralogues in *Lingula*) which are consequently the only remaining CARD-Caspase in brachiopods. Indeed, the loss of one type of CARD-Caspase is often accompanied by the loss of the second one (Krasovec et al. 2023). Echinoderms are characterised by the loss of Caspase-2 and the duplication of Caspase-9, while it is the opposite for urochordates (Krasovec et al. 2023).

Molluscs, along with annelids and brachiopods, possess both FADD and DED-Caspases, two actors known to trigger extrinsic apoptosis in mammals (Figure 8). The theoretical binding affinity between both strongly suggests that extrinsic apoptosis could be activated with similar modality as that which is already known in mammals. However, the evolutionary relationships of DED-Caspases at the metazoan scale are still unclear, as the phylogenies performed so far are difficult to analyse and interpret (Sakamaki et al. 2014, 2015). Indeed, the topology of DED-Caspase sequences are significantly different to the species tree topology, and the branching of animal clades is not consistent with the current knowledge on metazoan relationships. Importantly, the presence of a potential extrinsic apoptosis in molluscs indicates that its absence in fly and nematode could be a secondary loss that does not reflect the commonality of non-deuterostome animals. Finally, DED-Caspases and FADD are the only two apoptotic actors identified here to present a positive selection pressure (dN/dS>1), a selection relaxation that could be due to the presence of both intrinsic and extrinsic apoptosis-like pathways, leading to functional redundancy (Supplementary Table 9).

**Figure 8:**
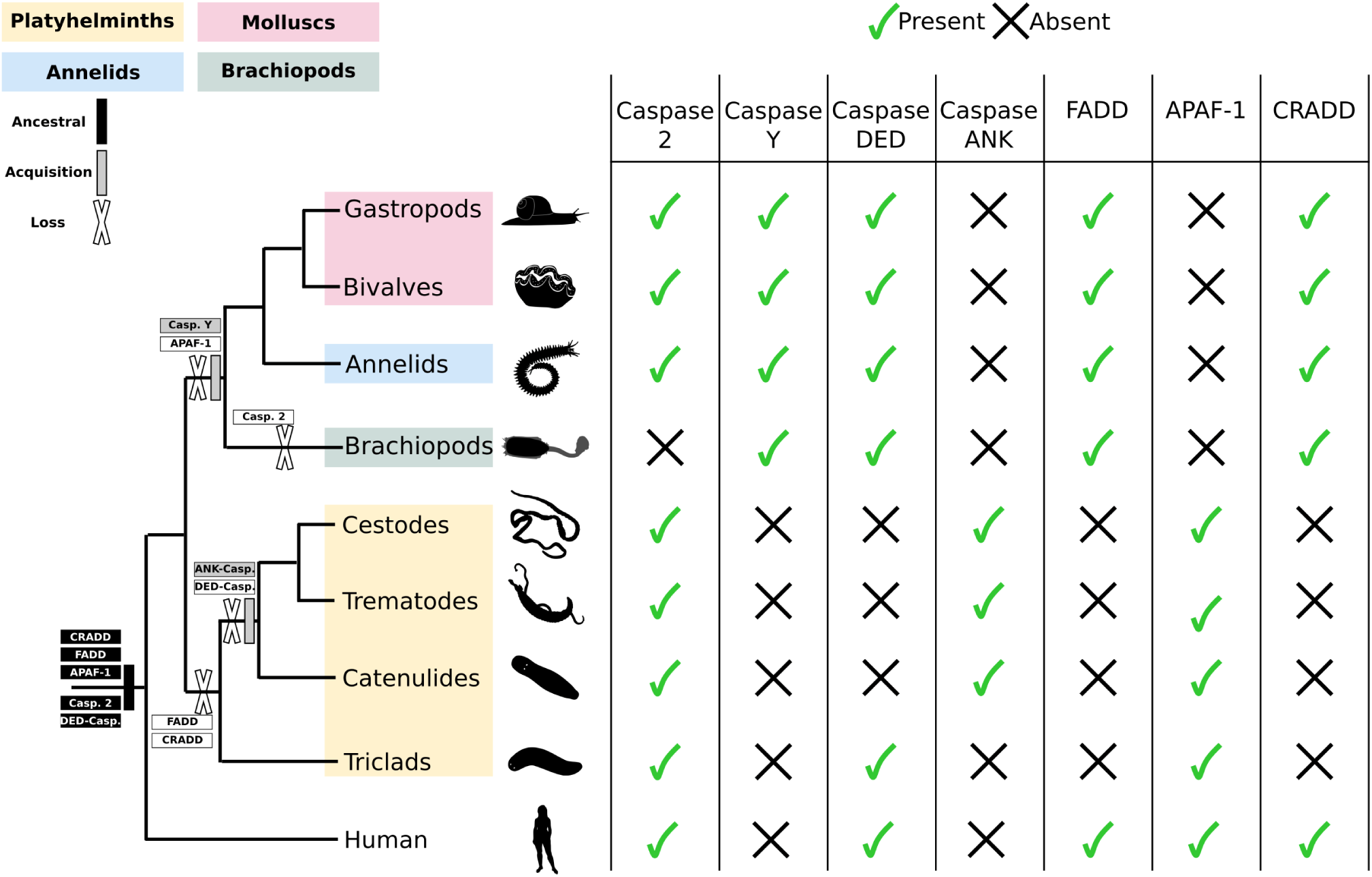
Summary of presence and absence of initiator Caspases and their potential activators in lophotrochozoans. Caspase-2, which is the shared and ancestral common initiator in animals, is detected in most lophotrochozoans but is lost in brachiopods. Caspase-Y is a specific acquisition of molluscs, annelids and brachiopods. DED-Caspases are lost in parasitic platyhelminths, which also acquired the specific ANK-Caspases. CARD-Caspases are likely to be activated by APAF-1 in platyhelminths and by CRADD in molluscs, annelids, and brachiopods. FADD, activator of DED-Caspase, is lost in all platyhelminths. This apoptotic network suggests a loss of extrinsic apoptosis in platyhelminths and a divergence in CARD-Caspase activation modality within lophotrochozoans.

Taken together, pioneering experimental works on lophotrochozoans (Yang et al. 2015; Pirger et al. 2004; Sokolova et al. 2004; Li et al. 2017) coupled with our study suggest that molluscs and close relatives present original apoptotic pathways which are different to what is known in mammals, fly and nematode.

### Annelids present a variety of species-specific initiator Caspases

Our study identifies annelids as the group with the most diverse Caspases repertoire known to date. We discovered a high diversity in this group, both in terms of Caspase number variation between species, but also regarding the presence of Caspases with unusual pro-domains. Among them is a Z-binding domain Caspase encoded by *Platynereis dumerilii*. Interestingly, Z-binding proteins are known to activate necroptosis in mammals, suggesting a rich diversity of cell death in this annelid (Maelfait et al. 2017). The ROC pro-domain, present in a Caspase from *Owenia fusiformis*, is known to be involved in protein GTP-binding, especially with kinase (Carlessi et al. 2011). This could allow the interaction with kinase proteins, such as death-associated protein kinase (known as DAPk) which is a mediator of cell death, cellular stress or autophagy regulation (Lin et al. 2010; Fujita & Yamashita 2014). The Subtilase and P/Homo B domains discovered in a *Metaphire vulgaris* Caspase are both normally found in proteins with protease activity (Siezen & Leunissen 1997; Rawlings & Barrett 1994; Anand et al. 2022; Zhou et al. 1998). Interestingly, subtilisin-like serine proteases are able to cleave various substrates: aspartic acid, serine and histidine. Consequently, this *Metaphire vulgaris* Caspase (MV008011650) seems able to cleave not only aspartate residues, which would make it a very unique Caspase in terms of catalytic activity.

### Platyhelminths conserve intrinsic apoptotic actors but have lost extrinsic apoptosis network

The history of DED-Caspase in platyhelminths is particularly interesting, as early branching platyhelminths possess one DED-Caspase while the parasitic species appear to have lost this gene. In addition, the DED-Caspase activator FADD has not been detected in platyhelminth genomes despite our investigations (Figure 8). The loss of one of these actors may lead to a lack of mobilisation of the other one, inducing a mutual relaxation of selection pressure leading to their parallel loss during evolution. Gene losses are common in animals with parasitic lifestyles due to genome and morphological simplification (Keeling 2004; Fairbairn 1970; Zarowiecki & Berriman 2015), which could explain the absence of some apoptotic actors in platyhelminths. This absence of extrinsic apoptosis in platyhelminths is shared with ecdysozoans, but is likely the result of convergent evolution, as extrinsic apoptotic actors are documented at a large evolutionary scale and are notably present in molluscs, annelids, and brachiopods (this study, (Sakamaki et al. 2014; Li et al. 2015; Lee et al. 2011)).

However, all parasitic species display an ANK-Caspase, a type of Caspase firstly identified in *Schistosoma* (Lee et al. 2014), which is unique to parasitic platyhelminths (with the exception of the *Alitta virens* annelid ANK-Caspase). Interestingly, ankyrin domains are known to be involved in protein-protein interaction and recognition (Mosavi et al. 2004). Furthermore, proteins with ankyrin domains are able to interact with Caspases (Flütsch et al. 2014; Schweizer et al. 2007). Consequently, ANK-Caspases may be initiators leading to the activation of upstream executioner Caspases. As ANK-Caspases are close to DED-Caspases, we may hypothesise that the acquisition of ankyrin pro-domains could be the result of a significant modification of the DED pro-domain of the ancestral DED-Caspases during evolution. Indeed, the recruitment of a new, but not yet identified, activator with an ankyrin domain was likely positively selected during the establishment of parasitism in these species. These specific caspases suggest a different pathway to the one currently known, which is of primary interest to investigate.

Platyhelminths present an apoptotic network composed of a unique CARD-Caspase (homologue to caspase-2) and the activator APAF-1, similarly to fly and nematode. In contrast to ecdysozoans, platyhelminth BCL-2 members seem able to induce MOMP and consequently the subsequent cytochrome c release, at least in *Schmidtea mediterranea* (Bender et al. 2012). The intrinsic apoptosis of platyhelminths appears to be one underlaid by CARD domain interactions, implicating the common APAF-1 activator, Caspase-2 (like fruit fly and nematode) and cytochrome c release (as in mammals). The resulting hypothetical apoptosome may share similarities with, or diverge from, apoptosomes from mammals and ecdysozoans, and may represent a new type of activation platform (APAF-1 + Caspase-2 + cytochrome c). The presence of a single pathway type likely induced purifying selection for conservation of these actors, since a deleterious modification or loss would lead to dysregulation of apoptosis (dN/dS<1) (Supplementary Table 9).

### Apoptotic signalling pathways evolve in a divergent way between metazoan clades

The current understanding of apoptotic signalling pathways evolution has benefited from decades of studies conducted mainly on nematode, fly and mammals (Hengartner 2000; Steller 2008; Lettre & Hengartner 2006; Ellis & Horvitz 1986). Ecdysozoans are characterised by the loss of numerous genes and the absence of extrinsic apoptosis, whereas mammals have an abundance of different apoptotic players; this has led to the false conclusion that apoptotic pathways are simple in “invertebrates”. In this study we discovered a large diversity of apoptotic network components in lophotrochozoans, along with clade-specific differences, indicating a complex evolution marked by multiple losses, acquisitions, and diversifications (Figure 8).

Phylogenetic evidence suggests that the central initiator of intrinsic apoptosis, Caspase-9, is deuterostome specific, while Caspase-2 is conserved among metazoans, and triggers apoptosis in fly and nematode (Krasovec et al. 2023). Consequently, it is not surprising that the only shared initiator Caspase that we have identified between all lophotrochozoan groups studied here is Caspase-2. Functional evidence shows that Caspase-2 in mammals, and its orthologues Dronc and Ced-3 in fly and nematode, respectively, are involved in a wide range of processes, including cell cycle regulation, tumour suppression, oxidative stress, ageing and cell death, and DNA repair (Tinel & Tschopp 2004; Krumschnabel, Sohm, et al. 2009, 2; Krumschnabel, Manzl, et al. 2009; Braga et al. 2008; Olsson et al. 2015). In addition, Caspase-2 has the ability to activate both intrinsic and extrinsic apoptosis, in addition to DNA damage pathway apoptosis in mammals (Braga et al. 2008; Olsson et al. 2009; Lavrik et al. 2006; Zhivotovsky & Orrenius 2005). This employment capacity is highlighted by the large number of proteins able to interact with Caspase-2, such as APAF-1 in ecdysozoans and CRADD or PIDD in mammals (Lin et al. 2000; Duan & Dixit 1997). This ancestral multifunctionality suggests that a mutation/loss of Caspase-2 would lead to destabilisation of various signalling cascades, increasing the likelihood of deleterious outcomes.

Our data suggest the existence of a possible apoptotic pathway in molluscs based on the Caspase-2/CRADD interaction, and another one in platyhelminths triggered by the Caspase-2/APAF-1 complex, in both cases relying on cytochrome c release. Cytochrome c release is likely an ancient trait of apoptosis regulation which is conserved in metazoans (Bender et al. 2012; Chipuk et al. 2010; Banjara et al. 2020), suggesting that intrinsic apoptosis in fly and nematode reflect an accumulation of derivative features concerning the implication of BCL-2 and mitochondria. The conservation of both anti- and pro-apoptotic BCL-2 in lophotrochozoans certainly results in their ability to play on mitochondria and apoptosis, but also on their multiple, ancestral and pleiotropic functions making them crucial in various molecular pathways (Popgeorgiev et al. 2019; Bonneau et al. 2013; Prudent et al. 2013; Popgeorgiev et al. 2020).

Finally, our data indicate that apoptotic signalling pathways have accumulated taxon-specific features during evolution, making these pathways divergent among metazoans and potentially convergent to the ultimate conserved trait: cell death executed by apoptosis regardless of the molecular pathways involved.

## MATERIAL AND METHODS

### Sequence dataset construction

We conducted BLAST searches for Caspases in a maximum of available lophotrochozoan genomes representing molluscs (highlighted in red along the figures), annelids (highlighted in blue), brachiopods (highlighted in dark green) and platyhelminths (highlighted in orange).

Putative lophotrochozoan genes were identified using tBLASTn and BLASTp searches using genes from human, fly, nematode, cnidarians, and oysters as queries. We extended the query in accordance to the results (*i.e.* newly identified Caspases were added to the query). Reciprocal BLAST was performed on all genes, including newly identified lophotrochozoan sequences. BLAST searching was conducted using NCBI databases for most species, and on PdumBase in addition to our recent transcriptome for annelid *Platynereis dumerilii* (Paré et al. 2023; Chou et al. 2018). After identification of genes in target species, proteins were analysed with ScanProsite (ExPaSy) (Gattiker et al. 2002) to verify the presence of specific domains which are the P20 and P10 for Caspases, CARD and Death Domains for CRADD, CARD and NB-ARC for APAF-1, DED and Death domains for FADD (see protein domains architecture, Figure 1). Sequences arising from BLAST search but devoid of the usual domains were not conserved to prevent false positives. Sampling was conducted on available lophotrochozoan genomes/transcriptomes, and for each species all Caspases and Caspase activators were explored.

Caspase family proteins are short (containing the large common P20 and the small P10 domains) with a high number of genes per species, which rapidly limits the relevance of the phylogenetic analyses (see Supplementary Figure 2 as example) (Krasovec et al. 2023). To reduce artefact branching and unreadable topologies, and to maximize phylogenetic diversity across lophotrochozoans, the dataset constructed for phylogenetic analysis with all types of Caspases was built with restricted number of representative lophotrochozoan species which recapitulate lophotrochozoan diversity. Secondly, analysis of Caspases with long pro-domains only was conducted with all species possible (for which unambiguous Caspase identifications were made). We provide here Supplementary Files 1 to 4 containing all Caspase sequences we found (including the executioner ones) for annelids, brachiopod, molluscs, and platyhelminths, respectively, which are present into the long pro- domains only phylogeny (topology in Figure 3). In case of redundancy (several accession numbers for one protein), we keep only one of them. The BCL2 phylogeny was also conducted on a representative species subset to overcome the same aforementioned artefacts which we already experienced.

FADD, CRADD, and APAF-1 analyses were conducted with all sequences identified at the lophotrochozoans scale which had the corresponding specific domains.

Multiple alignments of protein sequences were generated using MAFFT version 7 (Katoh & Standley 2013) with default parameters, and Clustal Omega (Sievers et al. 2011), to verify the congruence of the different alignments. All sequences were then manually checked in BioEdit 7.2 software to verify the presence of specific domains previously identified. Gblocks version 0.91b (Castresana 2000) was used to remove vacancies and blur sites. Final alignments comprise of 379, 214, 212, 560, 199, and 145 amino acids for the following alignments; Caspases with long pro-domain alignment, BCL-2 alignment, Apaf-1, CRADD alignment, and FADD alignment, respectively.

### Sampling strategy

Species sampling was conducted to include the highest diversity of lophotrochozoans possible in our analysis. When sampling we looked for caspases and potential activators in of a maximum range of genomes of lophotrochozoan species. Regarding the APAF-1, FADD, CRADD, and initiator Caspases sampling, we collected all sequences we detected. The PSOPIA score was calculated for all species were CARD-caspases plus APAF-1 and/or CRADD were identified, and same for DED-caspases plus FADD. Species sampled in one of our Caspases activator phylogenies but which is not included in the PSOPIA score analysis are species for which we did not detected initiator caspases candidates. Similarly, species represented in initiator Caspase analysis but not in APAF-1, CRADD, or FADD analyses, are species for which we did not detected caspase activator candidates. We maintained a maximum consistency between our analyses by keeping the same species. However, there are some differences resulting of species specificity and likely heterogeneity in genome properties. For example, we fund molluscs in which we did not detect any CRADD (*i.e. Gigantopelta aegis*), platyhelminths without APAF-1 *(i.e. Hymenolepis diminuta*), annelids and molluscs devoid of DED-Caspases (*i.e. Helobdella robusta* and *Candidula unifasciata*, respectively), or annelids without CARD-Caspases (*Tubifex tubifex*). Further details of this variability are available in Supplementary Table 1. This variability explains why some species are present in the Caspases analyses but not activator analyses, and conversely. Consequently, the differences in species sampling between our analyses are only the result of the diversity of lophotrochozoans in terms of the repertoire of apoptotic actors.

Regarding the analysis on all type of Caspases and BCL-2, we had to change the strategy to run the phylogenetic analysis in accordance with alignment length, in other word, the quantity of phylogenetic information. Caspase and BCL-2 proteins are short, which makes it challenging to conduct phylogenetic analyses with a large number of species, and consequently a large number of sequences (especially when including all of the multigene family), without sacrificing reliability. For these reasons, model animals and taxa were restricted, but animals with functional or well-studied genomic data were prioritised, in addition to be representative of the lophotrochozoans diversity.

### Phylogenetic analysis and sequence analysis

Phylogenetic analyses were carried out from the amino-acid alignment with Maximum-Likelihood (ML) method using PhyML 3.1 (Guindon et al. 2010). The best amino-acid evolution models to conduct analyses were determined using MEGA11 (Tamura et al. 2021) and appeared to be WAG for the Caspase alignments, Apaf-1 alignment, CRADD alignment, FADD alignment, and Cprev for the BCL-2 alignment. Node robustness was evaluated by 500 bootstrap replicates sampling.

Bayesian analyses were performed using MrBayes (v3.2.6) (Ronquist & Huelsenbeck 2003) under mixed model. For each analysis, one fourth of the topologies were discarded as burn-in values, while the remaining ones were used to calculate posterior probability. The run for the Caspase alignment was carried out for 3,000,000 generations with 15 randomly started simultaneous Markov chains (1 cold chain, 14 heated chains) and sampled every 100 generations. The runs for APAF-1, CRADD, and FADD alignments were carried out for 500,000 generations with 15 randomly started simultaneous Markov chains (1 cold chain, 14 heated chains) and sampled every 100 generations. For the BCL-2 3,000,000 generations with 15 randomly started simultaneous Markov chains (1 cold chain, 14 heated chains) was conducted and sampled every 100 generations. ML bootstrap values higher than 50% and Bayesian posterior probabilities are indicated on the Bayesian tree. The outgroup for the all Caspase phylogeny is *Reticulomyxa filosa* (ETO10778.1), a sequence recognizes to be sister of metazoan Caspases (Klim et al. 2018). For the BCL2 phylogeny, the outgroup used is a BCL-2-like from Porifera *Amphimedon queenslandica* (XP_003387574.1). Outgroups for APAF-1, CRADD, and FADD phylogenies are their human relatives.

Topology tests were evaluated using IQ-TREE (Nguyen et al. 2015). We conducted Kishino-Hasegawa test (Kishino & Hasegawa 1989), Shimodaira-Hasegawa test (Shimodaira & Hasegawa 1999), and approximately unbiased test (Shimodaira 2002).

The Dn/Ds ratio was calculated using SNAP v2.1.1 with default parameters on the ORFs previously codon-aligned by HIVAlign tool. SNAP v2.1.1 sort out raw Dn and Ds allowing next calculation of the ratio as presented in Supplementary Table 9.

### Protein-protein binding score prediction

Theoretical protein-protein binding scores were calculated using PSOPIA (Murakami & Mizuguchi 2014). Binding scores range from 0 (no affinity) to 1 (maximum binding probability estimation). For each category of measures (CARD-Caspase/APAF-1; CARD-Caspase/CRADD; FADD/DED-Caspase) we included positive controls (the relevant paired vertebrate proteins according to the literature) and negative controls (a Caspase plus a randomly selected protein which should not interact). Negative controls for DED-Caspase/FADD binding (Figure 6B1) are Negative-1: *Crassostrea gigas* FADD (NP_001295786.1) plus *Phodopus campbelli* insulin-like growth factor 2 (AFQ62592.1); Negative-2: *Mus musculus* Caspase-8 (AAH49955.1) plus *Hydra vulgaris* rhamnose-binding lectin (XP_002166914.1). Positive controls for DED-Caspase/FADD binding (Figure 6B1) are Positive-1: *Homo sapiens* Caspase-8 (AAD24962.1) plus FADD (CAG33019.1); Positive-2: *Mus musculus* Caspase-8 (AAH49955.1) plus FADD (AAA97876.1). Negative controls for CARD-Caspase/APAF-1 binding (Figure 6B2) are Negative-1: *Macrostomum lignano* CARD-Caspase (PAA59014.1) plus *Columba livia* Xrcc4 (PKK31261.1); Negative-2: *Drosophila melanogaster* Dronc (NP_524017.1) plus *Hydra vulgaris* rhamnose-binding lectin (XP_002166914.1). Positive controls for CARD-Caspase/APAF-1 binding (Figure 6B2) are Positive-1: *Homo sapiens* Caspase-9 (BAA82697.1) plus APAF-1 (NP_863651.1); Positive-2: *Caenorhabditis elegans* Ced-3 (AAG42045.1) plus Ced-4 (CAA48781.1). Negative control for CARD-Caspase/CRADD binding (Figure 6B3) is Negative-1: *Macrostomum lignano* CARD-Caspase (PAA59014.1) plus *Columba livia* Xrcc4 (PKK31261.1). Positive control for CARD-Caspase/CRADD binding (Figure 6B3) is Positive-1: *Homo sapiens* Caspase-2 (KAI4016188.1) plus CRADD (CAG28571.1).

### 3D protein models

3D structures were predicated using Phyre (Kelley et al. 2015) in normal mode and the best model was selected based on percentage alignment coverage and confidence score. The resultant PDB files were imported to Chimera X (Goddard et al. 2018) for model construction. Functional domains were detected using ScanProsite (ExPaSy).

## Acknowledgements

Acknowledgments

The authors acknowledge members of Uri Frank lab (University of Galway, Ireland) and Eve Gazave lab (Institut Jacques Monod, France).

## Notes

### Competing Interest Statement

The authors have declared no competing interest.

### Summary of Updates

Increased species sampling for phylogenetic analyses (caspases, caspase activators, BCL2). Addition of an analysis at the annelid scale. Addition of supplementary tables summarising species sampling. Addition of 3D models of caspase proteins. Addition of PSOPIA interaction scores (caspase/activator)

