## Supplemental Figures Tables Files for "Evolution of apoptotic signalling pathways within lophotrochozoans"

**Title**

† Deceased

* Authors for correspondence:

GK,

HRH,

**Affiliations**

^1^Centre for Chromosome Biology, School of Biological and Chemical Sciences, University of Galway, Galway, Ireland.

^2^Centre de Recherche en Cancérologie de Lyon, U1052 INSERM, UMR CNRS 5286, Centre Léon Bérard, Université Claude Bernard Lyon 1, Lyon, France.

^3^Institut Universitaire de France (IUF), Paris, France.

^4^Université Paris Cité, CNRS, Institut Jacques Monod, F-75013 Paris, France.

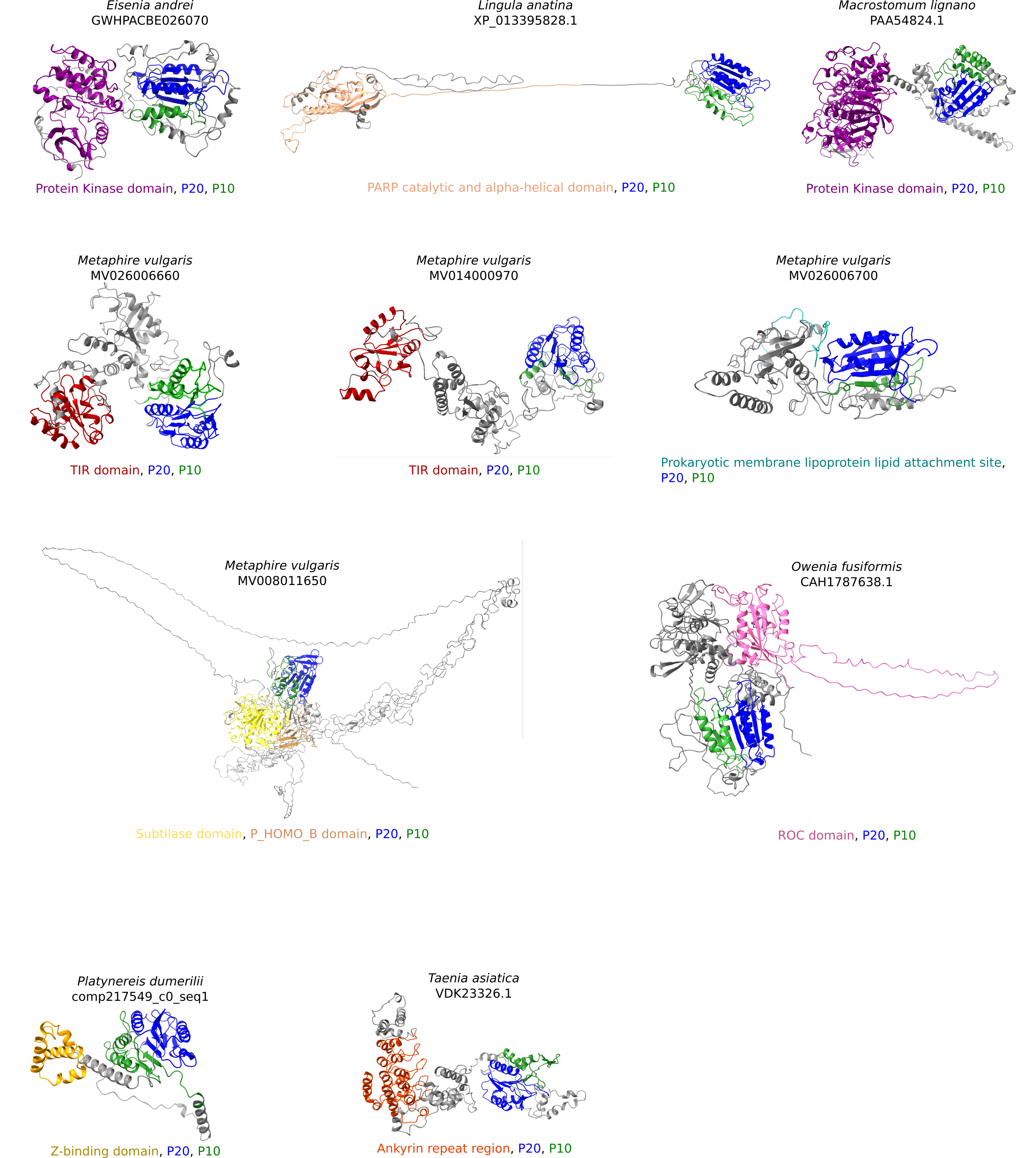

**Supplementary Figure 1:** **Specific structures of unusual initiator Caspases in lophotrochozoans.** Common domains of Caspases are the P20 (blue) and P10 (green). Other colours represent the pro-domains.

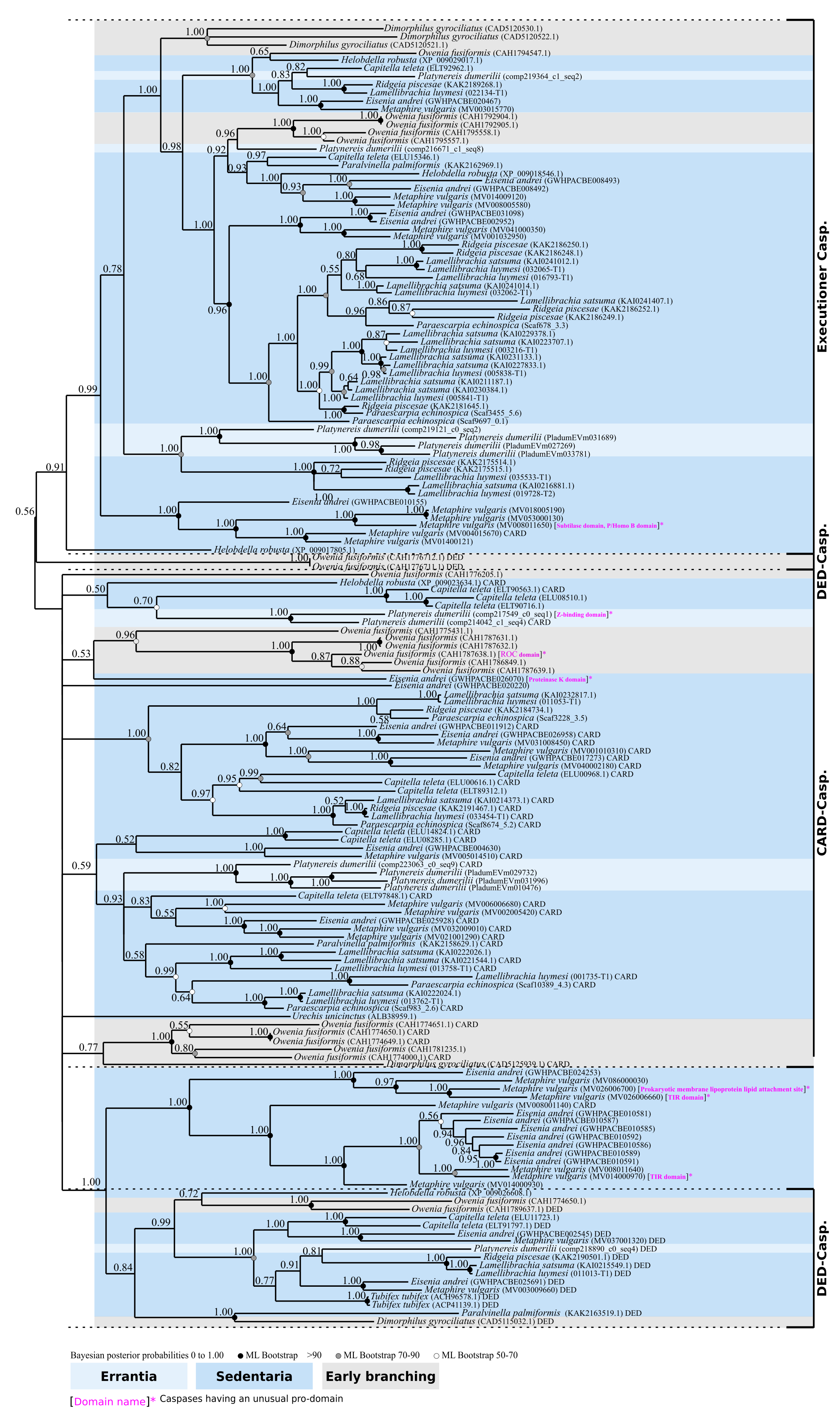

**Supplementary Figure 2:** **Topology of phylogenetic analysis of Caspases of annelids made by Bayesian inference method.** The analysis includes 11 Errantia, 1 Sedentaria, and two 2 branching annelids and was conducted on the full length of amino acid sequences containing the pro-domain and the common P20 plus P10 domains. Most executioner Caspases forms a clade. Caspases with a DED pro-domain (DED-Caspases), excepted two *Owenia fusiformis* sequences, form a monophyletic group. Caspases with a CARD pro-domain (CARD-Caspases) present a paraphyletic distribution due to several unresolved nodes. Unresolved nodes likely result of the large sequence number: Caspase are short, which makes it challenging to conduct phylogenetic analyses with a large number of sequences without sacrificing reliability. The presence of a classical CARD or DED pro-domain is indicated next to the sequence name. Caspases with a pro-domain other than CARD or DED are indicated in brackets and in magenta. Bootstrap correspond to robustness from maximum likelihood phylogeny which gave similar topology.

**Supplementary Table 1:** List of species used in our study. x: presence. N/A: absent, not detected.

| **Species** | **Executioner Caspases (no pro-domain)** | **DED-Caspase** | **ANK-Caspase** | **CARD-Caspase** | **FADD** | **CRADD** | **APAF-1** | **Phylum** |
| --- | --- | --- | --- | --- | --- | --- | --- | --- |
| Aplysia californica | x | x | N/A | x | x | x | N/A | **Molluscs** |
| Arenicola marina | N/A | x | N/A | N/A | x | N/A | N/A | **Annelids** |
| Alitta virens | x | x | x | x | x | x | N/A | **Annelids** |
| Biomphalaria glabrata | x | x | N/A | x | x | x | N/A | **Molluscs** |
| Bulinus truncatus | x | x | N/A | x | N/A | N/A | N/A | **Molluscs** |
| Candidula unifasciata | x | N/A | N/A | x | x | x | N/A | **Molluscs** |
| Capitella teleta | x | x | N/A | x | x | N/A | N/A | **Annelids** |
| Crassostrea gigas | x | x | N/A | x | x | x | N/A | **Molluscs** |
| Crassostrea virginica | x | x | N/A | x | x | x | N/A | **Molluscs** |
| Dibothriocephalus latus | N/A | N/A | N/A | x | N/A | N/A | x | **Platyhelminths** |
| Dimorphilus gyrociliatus | x | x | N/A | N/A | N/A | N/A | N/A | **Annelids** |
| Dreissena polymorpha | x | x | N/A | x | x | N/A | N/A | **Molluscs** |
| Dugesia japonica | x | x | N/A | N/A | N/A | N/A | N/A | **Platyhelminths** |
| Echinococcus granulosus | x | N/A | x | x | N/A | N/A | x | **Platyhelminths** |
| Echinococcus multilocularis | x | N/A | x | x | N/A | N/A | x | **Platyhelminths** |
| Eisenia andrei | x | x | N/A | x | N/A | N/A | N/A | **Annelids** |
| Elysia chlorotica | x | x | N/A | N/A | N/A | x | N/A | **Molluscs** |
| Elysia marginata | x | x | N/A | x | x | x | N/A | **Molluscs** |
| Gigantopelta aegis | x | x | N/A | x | x | N/A | N/A | **Molluscs** |
| Haliotis diversicolor | x | x | N/A | x | x | N/A | N/A | **Molluscs** |
| Helobdella robusta | x | N/A | N/A | x | N/A | N/A | N/A | **Annelids** |
| Hydatigera taeniaeformis | x | N/A | N/A | x | N/A | N/A | x | **Platyhelminths** |
| Hymenolepis diminuta | x | N/A | x | x | N/A | N/A | N/A | **Platyhelminths** |
| Hymenolepis microstoma | x | N/A | x | x | N/A | N/A | N/A | **Platyhelminths** |
| Lamellibrachia luymsi | x | x | N/A | x | x | x | N/A | **Annelids** |
| Lamellibrachia satsuma | x | x | N/A | x | x | N/A | N/A | **Annelids** |
| Lingula anatina | x | x | N/A | x | x | x | N/A | **Brachiopods** |
| Littorina littorea | N/A | x | N/A | N/A | x | x | N/A | **Molluscs** |
| Lottia gigantea | x | x | N/A | x | N/A | x | N/A | **Molluscs** |
| Macrostomum lignano | x | N/A | N/A | x | N/A | N/A | x | **Platyhelminths** |
| Mercenaria mercenaria | x | x | N/A | x | x | x | N/A | **Molluscs** |
| Mesocestoides corti | x | N/A | x | x | N/A | N/A | x | **Platyhelminths** |
| Metaphire vulgaris | x | x | N/A | x | x | N/A | N/A | **Annelids** |
| Mytilus californianus | x | x | N/A | x | x | N/A | N/A | **Molluscs** |
| Mytilus edulis | x | x | N/A | x | x | N/A | N/A | **Molluscs** |
| Mytilus galloprovincialis | x | x | N/A | x | x | N/A | N/A | **Molluscs** |
| Octopus bimaculoides | x | N/A | N/A | x | x | x | N/A | **Molluscs** |
| Onchidium reevesii | x | N/A | N/A | x | N/A | N/A | N/A | **Molluscs** |
| Ostrea edulis | x | x | N/A | x | x | x | N/A | **Annelids** |
| Owenia fusiformis | x | x | N/A | x | N/A | x | N/A | **Annelids** |
| Paraescarpia echinospica | x | N/A | N/A | x | x | N/A | N/A | **Annelids** |
| Paralvinella palmiformis | x | x | N/A | x | N/A | N/A | N/A | **Annelids** |
| Pecten maximus | x | x | N/A | x | x | N/A | N/A | **Molluscs** |
| Plakobranchus ocellatus | x | x | N/A | x | x | x | N/A | **Molluscs** |
| Platynereis dumerilii | x | x | N/A | x | x | x | N/A | **Annelids** |
| Pomacea canaliculata | x | x | N/A | x | x | x | N/A | **Molluscs** |
| Ridgeia piscesae | x | x | N/A | x | N/A | N/A | N/A | **Annelids** |
| Rodentolepis nana | x | N/A | N/A | x | N/A | N/A | N/A | **Platyhelminths** |
| Ruditapes philippinarum | x | x | N/A | x | x | N/A | N/A | **Molluscs** |
| Schistocephalus solidus | x | N/A | x | x | N/A | N/A | N/A | **Platyhelminths** |
| Schistosoma bovis | x | N/A | x | x | N/A | N/A | N/A | **Platyhelminths** |
| Schistosoma haematobium | x | N/A | x | x | N/A | N/A | x | **Platyhelminths** |
| Schistosoma japonicum | x | N/A | x | x | N/A | N/A | x | **Platyhelminths** |
| Schistosoma mansoni | x | N/A | x | x | N/A | N/A | N/A | **Platyhelminths** |
| Schmidtea mediterranea | x | x | N/A | x | N/A | N/A | x | **Platyhelminths** |
| Sparganum proliferum | N/A | N/A | N/A | N/A | N/A | N/A | x | **Platyhelminths** |
| Spirometra erinaceieuropaei | x | N/A | x | N/A | N/A | N/A | x | **Platyhelminths** |
| Taenia asiatica | x | N/A | x | x | N/A | N/A | x | **Platyhelminths** |
| Tubifex tubifex | N/A | x | N/A | N/A | N/A | N/A | N/A | **Annelids** |
| Urechis unicinctus | x | N/A | N/A | N/A | N/A | N/A | N/A | **Annelids** |

**Supplementary Table 2:** List of species and sequences of Caspases used to conduct phylogeny analysis of Figure 2.

| Species | Accession number | prodomain | Data Base |
| --- | --- | --- | --- |
| Aplysia californica | XP_005113266.1 | CARD | NCBI |
| Aplysia californica | XP_012945247.1 | DED | NCBI |
| Aplysia californica | XP_012945422.1 | CARD | NCBI |
| Aplysia californica | XP_005099863 |  | NCBI |
| Aplysia californica | XP_005092532 |  | NCBI |
| Aplysia californica | XP_012944029 |  | NCBI |
| Aplysia californica | XP_012940622 |  | NCBI |
| Biomphalaria glabrata | XP_013082356.1 | CARD | NCBI |
| Biomphalaria glabrata | XP_013094741.1 | DED | NCBI |
| Biomphalaria glabrata | XP_013088924 |  | NCBI |
| Biomphalaria glabrata | XP_013073999 |  | NCBI |
| Biomphalaria glabrata | XP_013074008 |  | NCBI |
| Biomphalaria glabrata | XP_013065656 |  | NCBI |
| Biomphalaria glabrata | XP_013067788 |  | NCBI |
| Biomphalaria glabrata | XP_013088924 |  | NCBI |
| Echinococcus granulosus | KAH9285259.1 | CARD | NCBI |
| Echinococcus granulosus | KAH9278879.1 | ANKYRIN | NCBI |
| Echinococcus granulosus | KAH9283631.1 |  | NCBI |
| Echinococcus granulosus | KAH9280007.1 |  | NCBI |
| Haliotis diversicolor | QHY92978.1 | DED | NCBI |
| Haliotis diversicolor | AYK02334.1 | CARD | NCBI |
| Haliotis diversicolor | AYK02335 |  | NCBI |
| Haliotis diversicolor | ABY87390 |  | NCBI |
| Helobdella robusta | XP_009017805.1 |  | NCBI |
| Helobdella robusta | XP_009018546.1 |  | NCBI |
| Helobdella robusta | XP_009026608.1 |  | NCBI |
| Helobdella robusta | XP_009029017.1 |  | NCBI |
| Helobdella robusta | XP_009023634.1 | CARD | NCBI |
| Lingula anatina | XP_013414814.1 | CARD | NCBI |
| Lingula anatina | XP_013402632.1 |  | NCBI |
| Lingula anatina | XP_013416922.1 | CARD | NCBI |
| Lingula anatina | XP_023932957.1 | CARD | NCBI |
| Lingula anatina | XP_013405490.1 | CARD | NCBI |
| Lingula anatina | XP_023932692.1 |  | NCBI |
| Lingula anatina | XP_013401916.1 |  | NCBI |
| Lingula anatina | XP_013414929.1 |  | NCBI |
| Lingula anatina | XP_013399774.1 |  | NCBI |
| Lingula anatina | XP_013408948.1 |  | NCBI |
| Lingula anatina | XP_013408945.1 |  | NCBI |
| Lingula anatina | XP_013408949.1 |  | NCBI |
| Lingula anatina | XP_013387919.1 |  | NCBI |
| Lingula anatina | XP_013381129.1 | DED | NCBI |
| Lingula anatina | XP_013411651.1 |  | NCBI |
| Lingula anatina | XP_013403374.2 | CARD | NCBI |
| Lingula anatina | XP_013381226.2 |  | NCBI |
| Lingula anatina | XP_013395828.1 | PARP | NCBI |
| Lingula anatina | XP_013409672.1 |  | NCBI |
| Lingula anatina | XP_013395658.1 |  | NCBI |
| Lingula anatina | XP_013416458.1 |  | NCBI |
| Lingula anatina | XP_013411650.1 |  | NCBI |
| Lingula anatina | XP_013406381.1 |  | NCBI |
| Lingula anatina | XP_013421206.1 |  | NCBI |
| Octopus bimaculoides | XP_014783442.1 | CARD | NCBI |
| Octopus bimaculoides | XP_014790087.1 | CARD | NCBI |
| Octopus bimaculoides | XP_014784582.1 | CARD | NCBI |
| Octopus bimaculoides | XP_014780506.1 |  | NCBI |
| Octopus bimaculoides | XP_014783598.1 |  | NCBI |
| Octopus bimaculoides | XP_014775611.1 |  | NCBI |
| Octopus bimaculoides | XP_014775614.1 |  | NCBI |
| Octopus bimaculoides | XP_014789170.1 |  | NCBI |
| Octopus bimaculoides | XP_014775609.1 |  | NCBI |
| Octopus bimaculoides | XP_014768177.1 |  | NCBI |
| Owenia fusiformis | CAH1774000.1 | CARD | NCBI |
| Owenia fusiformis | CAH1774649.1 | CARD | NCBI |
| Owenia fusiformis | CAH1787639.1 |  | NCBI |
| Owenia fusiformis | CAH1774650.1 | CARD | NCBI |
| Owenia fusiformis | CAH1787638.1 | ROC | NCBI |
| Owenia fusiformis | CAH1794547.1 |  | NCBI |
| Owenia fusiformis | CAH1795557.1 |  | NCBI |
| Owenia fusiformis | CAH1781235.1 |  | NCBI |
| Owenia fusiformis | CAH1774651.1 | CARD | NCBI |
| Owenia fusiformis | CAH1795558.1 |  | NCBI |
| Owenia fusiformis | CAH1792905.1 |  | NCBI |
| Owenia fusiformis | CAH1792904.1 |  | NCBI |
| Owenia fusiformis | CAH1787632.1 |  | NCBI |
| Owenia fusiformis | CAH1787631.1 |  | NCBI |
| Owenia fusiformis | CAH1776711.1 | DED | NCBI |
| Owenia fusiformis | CAH1789637.1 | DED | NCBI |
| Owenia fusiformis | CAH1776712.1 | DED | NCBI |
| Owenia fusiformis | CAH1789638.1 |  | NCBI |
| Owenia fusiformis | CAH1776205.1 |  | NCBI |
| Platynereis dumerilii | comp219364_c1_seq2 |  | PdumBase |
| Platynereis dumerilii | comp218890_c0_seq4 | DED | PdumBase |
| Platynereis dumerilii | comp216671_c1_seq8 |  | PdumBase |
| Platynereis dumerilii | comp217549_c0_seq1 | Z-BINDING DOMAIN | PdumBase |
| Platynereis dumerilii | comp223063_c0_seq9 | CARD | PdumBase |
| Platynereis dumerilii | comp219121_c0_seq2 |  | PdumBase |
| Platynereis dumerilii | comp214042_c1_seq4 | CARD | PdumBase |
| Platynereis dumerilii | PladumEVm029732 |  | PdumBase |
| Platynereis dumerilii | PladumEVm027269 |  | Paré et al, 2023 |
| Platynereis dumerilii | PladumEVm031996 |  | Paré et al, 2023 |
| Platynereis dumerilii | PladumEVm010476 |  | Paré et al, 2023 |
| Platynereis dumerilii | PladumEVm031689 |  | Paré et al, 2023 |
| Platynereis dumerilii | PladumEVm033781 |  | Paré et al, 2023 |
| Schistocephalus solidus | VDL94269.1 |  | NCBI |
| Schistocephalus solidus | VDM00376.1 | CARD | NCBI |
| Schistocephalus solidus | VDL97252.1 | ANKYRIN | NCBI |
| Schistosoma mansoni | ACU88130.1 |  | NCBI |
| Schistosoma mansoni | ACU88129.1 |  | NCBI |
| Schistosoma mansoni | XP_018646351.1 | CARD | NCBI |
| Schistosoma mansoni | XP_018647858.1 | ANKYRIN | NCBI |
| Spirometra erinaceieuropaei | VZI50365.1 |  | NCBI |
| Spirometra erinaceieuropaei | VZI03839.1 |  | NCBI |
| Spirometra erinaceieuropaei | VZI43280.1 |  | NCBI |
| Spirometra erinaceieuropaei | VZI32601.1 | ANKYRIN | NCBI |
| Taenia asiatica | VDK35885.1 |  | NCBI |
| Taenia asiatica | VDK22955.1 |  | NCBI |
| Taenia asiatica | VDK23513.1 | CARD | NCBI |
| Taenia asiatica | VDK23326.1 | ANKYRIN | NCBI |

**Supplementary Table 3:** List of species and sequences of Caspases used to conduct phylogeny analysis of Figure 3.

| Species | Accession number | Domain | Name | Data Base |
| --- | --- | --- | --- | --- |
| Aplysia californica | XP_005113266.2 | CARD | Casp.2 | NCBI |
| Aplysia californica | XP_012945247.1 | DED | Casp.DED | NCBI |
| Aplysia californica | XP_012945422.1 | CARD | Casp.Y | NCBI |
| Arenicola marina | UCK81519.1 | DED | Casp.DED | NCBI |
| Biomphalaria glabrata | XP_013082356.1 | CARD | Casp.2 | NCBI |
| Biomphalaria glabrata | XP_013094741.1 | DED | Casp.DED | NCBI |
| Bulinus truncatus | KAH9490831.1 | CARD | Casp.2 | NCBI |
| Bulinus truncatus | KAH9494957.1 | DED | Casp.DED | NCBI |
| Bulinus truncatus | KAH9513478.1 | CARD | Casp.Y1 | NCBI |
| Bulinus truncatus | KAH9499437.1 | CARD | Casp.Y2 | NCBI |
| Candidula unifasciata | CAG5125026.1 | CARD | Casp.Y1 | NCBI |
| Candidula unifasciata | CAG5131086.1 | CARD | Casp.Y2 | NCBI |
| Capitella teleta | ELU00968.1 | CARD | Casp.2a | NCBI |
| Capitella teleta | ELU00616.1 | CARD | Casp.2b | NCBI |
| Capitella teleta | ELT90563.1 | CARD | Casp.2c | NCBI |
| Capitella teleta | ELT91797.1 | DED | Casp.DED | NCBI |
| Capitella teleta | ELT97848.1 | CARD | Casp.Y1 | NCBI |
| Capitella teleta | ELU14824.1 | CARD | Casp.Y2 | NCBI |
| Capitella teleta | ELU08285.1 | CARD | Casp.Y3 | NCBI |
| Crassostrea gigas | XP_011419292.1 | CARD | Casp.2 | NCBI |
| Crassostrea gigas | AKP95634.1 | DED | Casp.DED1 | NCBI |
| Crassostrea gigas | XP_034301971.1 | DED | Casp.DED2 | NCBI |
| Crassostrea gigas | XP_011414267.1 | CARD | Casp.Y1 | NCBI |
| Crassostrea gigas | XP_011432762.1 | CARD | Casp.Y2 | NCBI |
| Crassostrea gigas | XP_011449817.1 | CARD | Casp.Y3 | NCBI |
| crassostrea virginica | XP_022338210.1 | CARD | Casp.2a | NCBI |
| crassostrea virginica | XP_022343382.1 | CARD | Casp.2b | NCBI |
| crassostrea virginica | XP_022321371.1 | DED | Casp.DED1 | NCBI |
| crassostrea virginica | XP_022325102.1 | DED | Casp.DED2 | NCBI |
| crassostrea virginica | XP_022294613.1 | DED | Casp.DED3 | NCBI |
| crassostrea virginica | XP_022286252.1 | CARD | Casp.Y1 | NCBI |
| crassostrea virginica | XP_022345548.1 | CARD | Casp.Y2 | NCBI |
| crassostrea virginica | XP_022343665.1 | CARD | Casp.Y3 | NCBI |
| Dibothriocephalus latus | VDN15196.1 | CARD | Casp.2 | NCBI |
| Dimorphilus gyrociliatus | CAD5115032.1 | DED | Casp.DED | NCBI |
| Dimorphilus gyrociliatus | CAD5125939.1 | CARD | Casp.Y | NCBI |
| Dreissena polymorpha | XP_052285027.1 | CARD | Casp.2 | NCBI |
| Dreissena polymorpha | XP_052235568.1 | DED | Casp.DED | NCBI |
| Dreissena polymorpha | XP_052225747.1 | CARD | Casp.Y1 | NCBI |
| Dreissena polymorpha | XP_052285285.1 | CARD | Casp.Y2 | NCBI |
| Dugesia japonica | BAP74679.1 | DED | Casp.DED | NCBI |
| Echinococcus granulosus | XP_024352296.1 | CARD | Casp.2 | NCBI |
| Echinococcus granulosus | KAH9278879.1 | ANK | Casp.Ank | NCBI |
| Echinococcus multilocularis | CDI97360.1 | CARD | Casp.2 | NCBI |
| Echinococcus multilocularis | CDS36952.1 | ANK | Casp.Ank | NCBI |
| Elysia chlorotica | RUS70675.1 | DED | Casp.DED | NCBI |
| Elysia marginata | GFR67360.1 | DED | Casp.DED | NCBI |
| Elysia marginata | GFS16061.1 | CARD | Casp.Y | NCBI |
| Gigantopelta aegis | XP_041377847.1 | CARD | Casp.2a | NCBI |
| Gigantopelta aegis | XP_041378168.1 | CARD | Casp.2b | NCBI |
| Gigantopelta aegis | XP_041374485.1 | DED | Casp.DED | NCBI |
| Gigantopelta aegis | XP_041378464.1 | CARD | Casp.Y1 | NCBI |
| Haliotis diversicolor | AYK02334.1 | CARD | Casp.CARD | NCBI |
| Haliotis diversicolor | QHY92978.1 | DED | Casp.DED | NCBI |
| Hydatigera taeniaeformis | VDM32914.1 | CARD | Casp.2 | NCBI |
| Hymenolepis diminuta | VDL19533.1 | CARD | Casp.2 | NCBI |
| Hymenolepis diminuta | VDL11723.1 | ANK | Casp.Ank | NCBI |
| Hymenolepis microstoma | CDS30656.1 | CARD | Casp.2 | NCBI |
| Hymenolepis microstoma | CDS29941.1 | ANK | Casp.Ank | NCBI |
| Lamellibrachia satsuma | KAI0214373.1 | CARD | Casp.2a | NCBI |
| Lamellibrachia satsuma | KAI0241012.1 | CARD | Casp.2b | NCBI |
| Lamellibrachia satsuma | KAI0215549.1 | DED | Casp.DED | NCBI |
| Lamellibrachia satsuma | KAI0221544.1 | CARD | Casp.Y1 | NCBI |
| Lamellibrachia satsuma | KAI0222024.1 | CARD | Casp.Y2 | NCBI |
| Lingula anatina | XP_013395828.1 | PARP | Casp, parp | NCBI |
| Lingula anatina | XP_013381129.1 | DED | Casp.DED | NCBI |
| Lingula anatina | XP_013403374.2 | CARD | Casp.Y1 | NCBI |
| Lingula anatina | XP_013395658.1 | CARD | Casp.Y2 | NCBI |
| Lingula anatina | XP_013405490.1 | CARD | Casp.Y3 | NCBI |
| Lingula anatina | XP_023932957.1 | CARD | Casp.Y4 | NCBI |
| Lingula anatina | XP_013416922.1 | CARD | Casp.Y5 | NCBI |
| Lingula anatina | XP_013414814.1 | CARD | Casp.Y6 | NCBI |
| Littorina littorea | QNL15268.1 | DED | Casp.DED | NCBI |
| Lottia gigantea | XP_009055665.1 | DED | Casp.DED | NCBI |
| Lottia gigantea | XP_009066758.1 | CARD | Casp.Y | NCBI |
| Macrostomum lignano | PAA59014.1 | CARD | Casp.2 | NCBI |
| Macrostomum lignano | PAA54824.1 | PROTEIN KINASE DOMAIN | Casp.PK | NCBI |
| Mercenaria mercenaria | XP_053379488.1 | CARD | Casp.2a | NCBI |
| Mercenaria mercenaria | XP_053378967.1 | CARD | Casp.2b | NCBI |
| Mercenaria mercenaria | XP_053385490.1 | DED | Casp.DED1 | NCBI |
| Mercenaria mercenaria | XP_053404642.1 | DED | Casp.DED2 | NCBI |
| Mercenaria mercenaria | XP_045184554.2 | DED | Casp.DED3 | NCBI |
| Mercenaria mercenaria | XP_045161198.2 | CARD | Casp.Y1 | NCBI |
| Mercenaria mercenaria | XP_045162709.1 | CARD | Casp.Y2 | NCBI |
| Mesocestoides corti | VDD79553.1 | CARD | Casp.2 | NCBI |
| Mesocestoides corti | VDD77147.1 | ANKYRIN | Casp.Ank | NCBI |
| Mytilus californianus | XP_052082663.1 | CARD | Casp.2a | NCBI |
| Mytilus californianus | XP_052081364.1 | CARD | Casp.2b | NCBI |
| Mytilus californianus | XP_052081366.1 | CARD | Casp.2c | NCBI |
| Mytilus californianus | ADB80146.1 | DED | Casp.DED1 | NCBI |
| Mytilus californianus | ADB80147.1 | DED | Casp.DED2 | NCBI |
| Mytilus californianus | XP_052079629.1 | DED | Casp.DED3 | NCBI |
| Mytilus californianus | XP_052067829.1 | DED | Casp.DED4 | NCBI |
| Mytilus californianus | XP_052081018.1 | DED | Casp.DED5 | NCBI |
| Mytilus californianus | XP_052081014.1 | DED | Casp.DED6 | NCBI |
| Mytilus californianus | XP_052081013.1 | DED | Casp.DED7 | NCBI |
| Mytilus californianus | XP_052102868 | CARD | Casp.Y1 | NCBI |
| Mytilus californianus | XP_052073552 | CARD | Casp.Y2 | NCBI |
| Mytilus californianus | XP_052073549 | CARD | Casp.Y3 | NCBI |
| Mytilus californianus | XP_052080984 | CARD | Casp.Y4 | NCBI |
| Mytilus californianus | XP_052080923 | CARD | Casp.Y5 | NCBI |
| Mytilus californianus | XP_052095498 | CARD | Casp.Y6 | NCBI |
| Mytilus edulis | CAG2226391 | DED | Casp.DED1 | NCBI |
| Mytilus edulis | CAG2226387 | DED | Casp.DED2 | NCBI |
| Mytilus edulis | CAG2237348 | DED | Casp.DED3 | NCBI |
| Mytilus edulis | CAG2231765 | DED | Casp.DED4 | NCBI |
| Mytilus edulis | CAG2233488 | DED | Casp.DED5 | NCBI |
| Mytilus edulis | CAG2224734 | DED | Casp.DED6 | NCBI |
| Mytilus edulis | CAG2203577.1 | CARD | Casp.Y1 | NCBI |
| Mytilus edulis | CAG2189547.1 | CARD | Casp.Y2 | NCBI |
| Mytilus edulis | CAG2231374.1 | CARD | Casp.Y3 | NCBI |
| Mytilus edulis | CAG2184393.1 | CARD | Casp.Y4 | NCBI |
| Mytilus galloprovincialis | VDI15888.1 | CARD | Casp.2a | NCBI |
| Mytilus galloprovincialis | VDI33080.1 | CARD | Casp.2b | NCBI |
| Mytilus galloprovincialis | VDI78201.1 | CARD | Casp.2c | NCBI |
| Mytilus galloprovincialis | AIS73193.1 | DED | Casp.DED1 | NCBI |
| Mytilus galloprovincialis | VDH91938 | DED | Casp.DED2 | NCBI |
| Mytilus galloprovincialis | VDH91689 | DED | Casp.DED3 | NCBI |
| Mytilus galloprovincialis | VDH97448 | CARD | Casp.Y1 | NCBI |
| Mytilus galloprovincialis | ADZ24778.1 | CARD | Casp.Y2 | NCBI |
| Mytilus galloprovincialis | VDI42160.1 | CARD | Casp.Y3 | NCBI |
| Mytilus galloprovincialis | VDI06563.1 | CARD | Casp.Y4 | NCBI |
| Mytilus galloprovincialis | VDI22398.1 | CARD | Casp.Y5 | NCBI |
| Octopus bimaculoides | XP_014783442.1 | CARD | Casp.2 | NCBI |
| Octopus bimaculoides | XP_014790087.1 | CARD | Casp.Y1 | NCBI |
| Octopus bimaculoides | XP_014784582.1 | CARD | Casp.Y2 | NCBI |
| Onchidium reevesii | QMS54626.1 | CARD | Casp.Y | NCBI |
| Ostrea edulis | XP_048751990.2 | CARD | Casp.2a | NCBI |
| Ostrea edulis | XP_048747578.2 | CARD | Casp.2b | NCBI |
| Ostrea edulis | XP_048776588.1 | DED | Casp.DED1 | NCBI |
| Ostrea edulis | XP_048757091.1 | DED | Casp.DED2 | NCBI |
| Ostrea edulis | XP_048769780.1 | DED | Casp.DED3 | NCBI |
| Ostrea edulis | XP_048770132.1 | CARD | Casp.Y1 | NCBI |
| Ostrea edulis | XP_048760265.2 | CARD | Casp.Y2 | NCBI |
| Ostrea edulis | XP_048742450.2 | CARD | Casp.Y3 | NCBI |
| Owenia fusiformis | CAH1787638.1 | ROC | Casp, roc | NCBI |
| Owenia fusiformis | CAH1789637.1 | DED | Casp.DED1 | NCBI |
| Owenia fusiformis | CAH1776712.1 | DED | Casp.DED2 | NCBI |
| Owenia fusiformis | CAH1776711.1 | DED | Casp.DED3 | NCBI |
| Owenia fusiformis | CAH1774649.1 | CARD | Casp.Y1 | NCBI |
| Owenia fusiformis | CAH1774000.1 | CARD | Casp.Y2 | NCBI |
| Owenia fusiformis | CAH1774650.1 | CARD | Casp.Y3 | NCBI |
| Owenia fusiformis | CAH1774651.1 | CARD | Casp.Y4 | NCBI |
| Pecten maximus | XP_033724734.1 | CARD | Casp.2a | NCBI |
| Pecten maximus | XP_033724726.1 | CARD | Casp.2b | NCBI |
| Pecten maximus | XP_033740501.1 | DED | Casp.DED1 | NCBI |
| Pecten maximus | XP_033740498.1 | DED | Casp.DED2 | NCBI |
| Pecten maximus | XP_033728057.1 | DED | Casp.DED3 | NCBI |
| Pecten maximus | XP_033724674.1 | CARD | Casp.Y1 | NCBI |
| Pecten maximus | XP_033724604.1 | CARD | Casp.Y2 | NCBI |
| Pecten maximus | XP_033742269.1 | CARD | Casp.Y3 | NCBI |
| Plakobranchus ocellatus | GFO26738.1 | DED | Casp.DED | NCBI |
| Plakobranchus ocellatus | GFO14050.1 | CARD | Casp.Y | NCBI |
| Platynereis dumerilii | comp214042_c1_seq4 | CARD | Casp.2 | PdumBase |
| Platynereis dumerilii | comp218890_c0_seq4 | DED | Casp.DED | PdumBase |
| Platynereis dumerilii | comp223063_c0_seq9 | CARD | Casp.Y | PdumBase |
| Platynereis dumerilii | comp217549_c0_seq1 | Z-BINDING DOMAIN | Casp.Z_bin. | PdumBase |
| Pomacea canaliculata | XP_025083090.1 | CARD | Casp.2a | NCBI |
| Pomacea canaliculata | XP_025081418.1 | CARD | Casp.2b | NCBI |
| Pomacea canaliculata | XP_025088866.1 | DED | Casp.DED | NCBI |
| Pomacea canaliculata | XP_025080583.1 | CARD | Casp.Y1 | NCBI |
| Pomacea canaliculata | XP_025080591.1 | CARD | Casp.Y2 | NCBI |
| Rodentolepis nana | VDO03831.1 | CARD | Casp.2 | NCBI |
| Ruditapes philippinarum | XP_060600760.1 | CARD | Casp.2a | NCBI |
| Ruditapes philippinarum | XP_060565700.1 | CARD | Casp.2b | NCBI |
| Ruditapes philippinarum | XP_060560660.1 | DED | Casp.DED | NCBI |
| Schistocephalus solidus | VDM00376.1 | CARD | Casp.2 | NCBI |
| Schistocephalus solidus | VDL97252.1 | ANK | Casp.Ank | NCBI |
| Schistosoma bovis | RTG87993.1 | CARD | Casp.2 | NCBI |
| Schistosoma bovis | RTG88383.1 | ANK | Casp.Ank | NCBI |
| Schistosoma haematobium | CAH8464935.1 | CARD | Casp.2 | NCBI |
| Schistosoma haematobium | XP_012801329.1 | ANK | Casp.Ank | NCBI |
| Schistosoma japonicum | TNN06772.1 | CARD | Casp.2 | NCBI |
| Schistosoma japonicum | TNN08539.1 | ANK | Casp.Ank | NCBI |
| Schistosoma mansoni | XP_018646351.1 | CARD | Casp.2 | NCBI |
| Schistosoma mansoni | XP_018647858.1 | ANK | Casp.Ank | NCBI |
| Schmidtea mediterranea | AEX93483.1 | CARD | Casp.2 | NCBI |
| Schmidtea mediterranea | AGY56134.1 | DED | Casp.DED | NCBI |
| Spirometra erinaceieuropaei | VZI32601.1 | ANK | Casp.Ank | NCBI |
| Taenia asiatica | VDK23513.1 | CARD | Casp.2 | NCBI |
| Taenia asiatica | VDK23326.1 | ANK | Casp.Ank | NCBI |
| Tubifex tubifex | ACP41139.1 | DED | Casp.DED | NCBI |

**Supplementary Table 4:** List of species and sequences of Caspases used to conduct phylogeny analysis of Figure 5.

| Species | Accession number | prodomain | Data Base |
| --- | --- | --- | --- |
| Platynereis dumerilii | comp219364_c1_seq2 |  | PdumBase |
| Platynereis dumerilii | comp218890_c0_seq4 | DED | PdumBase |
| Platynereis dumerilii | comp216671_c1_seq8 |  | PdumBase |
| Platynereis dumerilii | comp217549_c0_seq1 | Z-BINDING DOMAIN | PdumBase |
| Platynereis dumerilii | comp223063_c0_seq9 | CARD | PdumBase |
| Platynereis dumerilii | comp219121_c0_seq2 |  | PdumBase |
| Platynereis dumerilii | comp214042_c1_seq4 | CARD | PdumBase |
| Platynereis dumerilii | PladumEVm029732 |  | PdumBase |
| Platynereis dumerilii | PladumEVm027269 |  | Paré et al, 2023 |
| Platynereis dumerilii | PladumEVm031996 |  | Paré et al, 2023 |
| Platynereis dumerilii | PladumEVm010476 |  | Paré et al, 2023 |
| Platynereis dumerilii | PladumEVm031689 |  | Paré et al, 2023 |
| Platynereis dumerilii | PladumEVm033781 |  | Paré et al, 2023 |
| Helobdella robusta | XP_009017805.1 |  | NCBI |
| Helobdella robusta | XP_009018546.1 |  | NCBI |
| Helobdella robusta | XP_009026608.1 |  | NCBI |
| Helobdella robusta | XP_009029017.1 |  | NCBI |
| Helobdella robusta | XP_009023634.1 | CARD | NCBI |
| Owenia fusiformis | CAH1774000.1 | CARD | NCBI |
| Owenia fusiformis | CAH1774649.1 | CARD | NCBI |
| Owenia fusiformis | CAH1787639.1 |  | NCBI |
| Owenia fusiformis | CAH1774650.1 | CARD | NCBI |
| Owenia fusiformis | CAH1787638.1 | ROC | NCBI |
| Owenia fusiformis | CAH1794547.1 |  | NCBI |
| Owenia fusiformis | CAH1795557.1 |  | NCBI |
| Owenia fusiformis | CAH1781235.1 |  | NCBI |
| Owenia fusiformis | CAH1774651.1 | CARD | NCBI |
| Owenia fusiformis | CAH1795558.1 |  | NCBI |
| Owenia fusiformis | CAH1792905.1 |  | NCBI |
| Owenia fusiformis | CAH1792904.1 |  | NCBI |
| Owenia fusiformis | CAH1787632.1 |  | NCBI |
| Owenia fusiformis | CAH1787631.1 |  | NCBI |
| Owenia fusiformis | CAH1776711.1 | DED | NCBI |
| Owenia fusiformis | CAH1789637.1 | DED | NCBI |
| Owenia fusiformis | CAH1776712.1 | DED | NCBI |
| Owenia fusiformis | CAH1789638.1 |  | NCBI |
| Owenia fusiformis | CAH1776205.1 |  | NCBI |
| Owenia fusiformis | CAH1786849.1 |  | NCBI |
| Owenia fusiformis | CAH1775431.1 |  | NCBI |
| Tubifex tubifex | ALB38959.1 | DED | NCBI |
| Tubifex tubifex | ACP41139.1 | DED | NCBI |
| Capitella teleta | ELU00968.1 | CARD | NCBI |
| Capitella teleta | ELT91797.1 | DED | NCBI |
| Capitella teleta | ELU11723.1 |  | NCBI |
| Capitella teleta | ELT92962.1 |  | NCBI |
| Capitella teleta | ELU15346.1 |  | NCBI |
| Capitella teleta | ELU00616.1 | CARD | NCBI |
| Capitella teleta | ELU08285.1 | CARD | NCBI |
| Capitella teleta | ELT90716.1 |  | NCBI |
| Capitella teleta | ELT97848.1 | CARD | NCBI |
| Capitella teleta | ELT90563.1 | CARD | NCBI |
| Capitella teleta | ELT89312.1 |  | NCBI |
| Capitella teleta | ELU14824.1 | CARD | NCBI |
| Capitella teleta | ELU08510.1 |  | NCBI |
| Urechis unicinctus | ALB38959.1 |  | NCBI |
| Dimorphilus gyrociliatus | CAD5120521.1 |  | NCBI |
| Dimorphilus gyrociliatus | CAD5120522.1 |  | NCBI |
| Dimorphilus gyrociliatus | CAD5120530.1 |  | NCBI |
| Dimorphilus gyrociliatus | CAD5115032.1 | DED | NCBI |
| Dimorphilus gyrociliatus | CAD5125939.1 | CARD | NCBI |
| Paraescarpia echinospica | PE_Scaf983_2.6 | CARD | NCBI |
| Paraescarpia echinospica | PE_Scaf8674_5.2 | CARD | NCBI |
| Paraescarpia echinospica | PE_Scaf9697_0.12 |  | NCBI |
| Paraescarpia echinospica | PE_Scaf3455_5.6 |  | NCBI |
| Paraescarpia echinospica | PE_Scaf3228_3.5 |  | NCBI |
| Paraescarpia echinospica | PE_Scaf10389_4.3 | CARD | NCBI |
| Paraescarpia echinospica | PE_Scaf678_3.3 |  | NCBI |
| Metaphire vulgaris | MV032009010 | CARD | NCBI |
| Metaphire vulgaris | MV021001290 | CARD | NCBI |
| Metaphire vulgaris | MV002005420 | CARD | NCBI |
| Metaphire vulgaris | MV001032950 |  | NCBI |
| Metaphire vulgaris | MV041000350 |  | NCBI |
| Metaphire vulgaris | MV005014510 | CARD | NCBI |
| Metaphire vulgaris | MV037001320 | DED | NCBI |
| Metaphire vulgaris | MV003009660 | DED | NCBI |
| Metaphire vulgaris | MV040002180 | CARD | NCBI |
| Metaphire vulgaris | MV031008450 | CARD | NCBI |
| Metaphire vulgaris | MV003015770 |  | NCBI |
| Metaphire vulgaris | MV008005580 |  | NCBI |
| Metaphire vulgaris | MV014009120 |  | NCBI |
| Metaphire vulgaris | MV014000930 |  | NCBI |
| Metaphire vulgaris | MV001010310 | CARD | NCBI |
| Metaphire vulgaris | MV026006660 | TIR | NCBI |
| Metaphire vulgaris | MV014000970 | TIR | NCBI |
| Metaphire vulgaris | MV008011640 |  | NCBI |
| Metaphire vulgaris | MV026006700 | Protein kinase domain | NCBI |
| Metaphire vulgaris | MV01400121 |  | NCBI |
| Metaphire vulgaris | MV008011650 | Subtilase | NCBI |
| Metaphire vulgaris | MV004015670 | CARD | NCBI |
| Metaphire vulgaris | MV053000130 |  | NCBI |
| Metaphire vulgaris | MV008001140 | CARD | NCBI |
| Metaphire vulgaris | MV018005190 |  | NCBI |
| Metaphire vulgaris | MV086000030 |  | NCBI |
| Lamellibrachia luymsi | LLU\|FUN_033454-T1 | CARD | NCBI |
| Lamellibrachia luymsi | LLU\|FUN_013758-T1 | CARD | NCBI |
| Lamellibrachia luymsi | LLU\|FUN_013762-T1 | CARD | NCBI |
| Lamellibrachia luymsi | LLU\|FUN_005841-T1 |  | NCBI |
| Lamellibrachia luymsi | LLU\|FUN_011053-T1 |  | NCBI |
| Lamellibrachia luymsi | LLU\|FUN_005838-T1 |  | NCBI |
| Lamellibrachia luymsi | LLU\|FUN_032062-T1 |  | NCBI |
| Lamellibrachia luymsi | LLU\|FUN_003216-T1 |  | NCBI |
| Lamellibrachia luymsi | LLU\|FUN_011013-T1 | DED | NCBI |
| Lamellibrachia luymsi | LLU\|FUN_016793-T1 |  | NCBI |
| Lamellibrachia luymsi | LLU\|FUN_022134-T1 |  | NCBI |
| Lamellibrachia luymsi | LLU\|FUN_001735-T1 | CARD | NCBI |
| Lamellibrachia luymsi | LLU\|FUN_032065-T1 | CARD | NCBI |
| Lamellibrachia luymsi | LLU\|FUN_019728-T2 |  | NCBI |
| Lamellibrachia luymsi | LLU\|FUN_035533-T1 |  | NCBI |
| Eisenia andrei | GWHPACBE025928 | CARD | NCBI |
| Eisenia andrei | GWHPACBE011912 | CARD | NCBI |
| Eisenia andrei | GWHPACBE004630 |  | NCBI |
| Eisenia andrei | GWHPACBE002952 |  | NCBI |
| Eisenia andrei | GWHPACBE025691 | DED | NCBI |
| Eisenia andrei | GWHPACBE026958 | CARD | NCBI |
| Eisenia andrei | GWHPACBE017273 | CARD | NCBI |
| Eisenia andrei | GWHPACBE020467 |  | NCBI |
| Eisenia andrei | GWHPACBE002545 | DED | NCBI |
| Eisenia andrei | GWHPACBE024253 |  | NCBI |
| Eisenia andrei | GWHPACBE031098 |  | NCBI |
| Eisenia andrei | GWHPACBE008492 |  | NCBI |
| Eisenia andrei | GWHPACBE026070 | Proteinase K | NCBI |
| Eisenia andrei | GWHPACBE010591 |  | NCBI |
| Eisenia andrei | GWHPACBE010589 |  | NCBI |
| Eisenia andrei | GWHPACBE010581 |  | NCBI |
| Eisenia andrei | GWHPACBE010586 |  | NCBI |
| Eisenia andrei | GWHPACBE010587 |  | NCBI |
| Eisenia andrei | GWHPACBE010585 |  | NCBI |
| Eisenia andrei | GWHPACBE010592 |  | NCBI |
| Eisenia andrei | GWHPACBE008493 |  | NCBI |
| Eisenia andrei | GWHPACBE020220 |  | NCBI |
| Eisenia andrei | GWHPACBE010155 |  | NCBI |
| Ridgeia piscesae | KAK2191467.1 | CARD | NCBI |
| Ridgeia piscesae | KAK2181645.1 |  | NCBI |
| Ridgeia piscesae | KAK2186248.1 |  | NCBI |
| Ridgeia piscesae | KAK2186250.1 |  | NCBI |
| Ridgeia piscesae | KAK2184734.1 |  | NCBI |
| Ridgeia piscesae | KAK2189268.1 |  | NCBI |
| Ridgeia piscesae | KAK2186249.1 |  | NCBI |
| Ridgeia piscesae | KAK2186252.1 |  | NCBI |
| Ridgeia piscesae | KAK2190501.1 | DED | NCBI |
| Ridgeia piscesae | KAK2175515.1 |  | NCBI |
| Ridgeia piscesae | KAK2175514.1 |  | NCBI |
| Paralvinella palmiformis | KAK2162969.1 |  | NCBI |
| Paralvinella palmiformis | KAK2158629.1 | CARD | NCBI |
| Paralvinella palmiformis | KAK2163519.1 | DED | NCBI |
| Lamellibrachia satsuma | KAI0222024 | CARD | NCBI |
| Lamellibrachia satsuma | KAI0241014 |  | NCBI |
| Lamellibrachia satsuma | KAI0221544 | CARD | NCBI |
| Lamellibrachia satsuma | KAI0214373 | CARD | NCBI |
| Lamellibrachia satsuma | KAI0222026 |  | NCBI |
| Lamellibrachia satsuma | KAI0230384 |  | NCBI |
| Lamellibrachia satsuma | KAI0232817 |  | NCBI |
| Lamellibrachia satsuma | KAI0241012 | CARD | NCBI |
| Lamellibrachia satsuma | KAI0227833 |  | NCBI |
| Lamellibrachia satsuma | KAI0211187 |  | NCBI |
| Lamellibrachia satsuma | KAI0223707 |  | NCBI |
| Lamellibrachia satsuma | KAI0231133 |  | NCBI |
| Lamellibrachia satsuma | KAI0215549 | DED | NCBI |
| Lamellibrachia satsuma | KAI0229378 |  | NCBI |
| Lamellibrachia satsuma | KAI0241407 |  | NCBI |
| Lamellibrachia satsuma | KAI0216881 |  | NCBI |
| Arenicola marina | UCK81519.1 | DED | NCBI |
| Alitta virens | c73072_g1_i1 | CARD | NCBI |
| Alitta virens | c83731_g5_i6 | CARD | NCBI |
| Alitta virens | c73773_g1_i1 |  | NCBI |
| Alitta virens | c84685_g6_i6 | CARD | NCBI |
| Alitta virens | c83651_g1_i1 | ANKYRIN | NCBI |
| Alitta virens | c66411_g1_i1 |  | NCBI |
| Alitta virens | c80567_g4_i3 |  | NCBI |
| Alitta virens | c85433_g3_i1 |  | NCBI |
| Alitta virens | c81751_g3_i4 |  | NCBI |
| Alitta virens | c70299_g2_i1 |  | NCBI |
| Alitta virens | c69909_g2_i1 |  | NCBI |
| Alitta virens | c66667_g3_i1 |  | NCBI |
| Alitta virens | c70299_g2_i5 |  | NCBI |
| Alitta virens | c80567_g4_i2 |  | NCBI |
| Alitta virens | c78337_g4_i1 | DED | NCBI |
| Alitta virens | c81829_g1_i3 |  | NCBI |
| Alitta virens | c76815_g1_i1 | CARD | NCBI |
| Alitta virens | c50262_g1_i1 |  | NCBI |
| Alitta virens | c68898_g2_i2 |  | NCBI |
| Alitta virens | c80075_g1_i |  | NCBI |
| Alitta virens | c71693_g2_i1 |  | NCBI |
| Alitta virens | c84172_g2_i5 |  | NCBI |
| Alitta virens | c76002_g2_i1 |  | NCBI |
| Alitta virens | c76205_g1_i1 | DED | NCBI |
| Alitta virens | c48166_g1_i1 |  | NCBI |
| Alitta virens | c70725_g1_i1 |  | NCBI |
| Alitta virens | c120789_g1_i1 |  | NCBI |

**Supplementary Table 5:** List of species and sequences of FADD used to conduct phylogeny analysis of Figure 6.

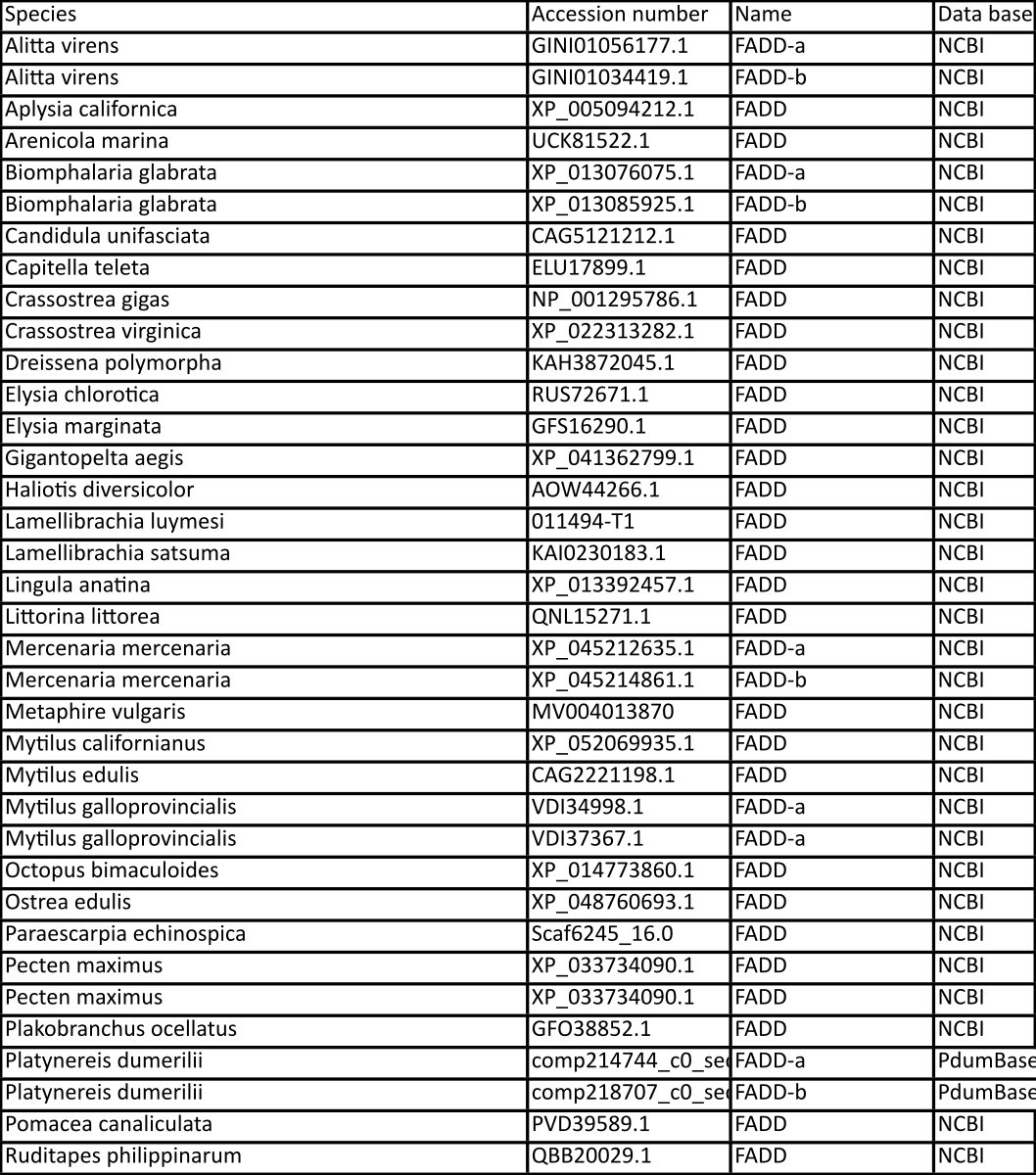

**Supplementary Table 6:** List of species and sequences of APAF-1 used to conduct phylogeny analysis of Figure 6.

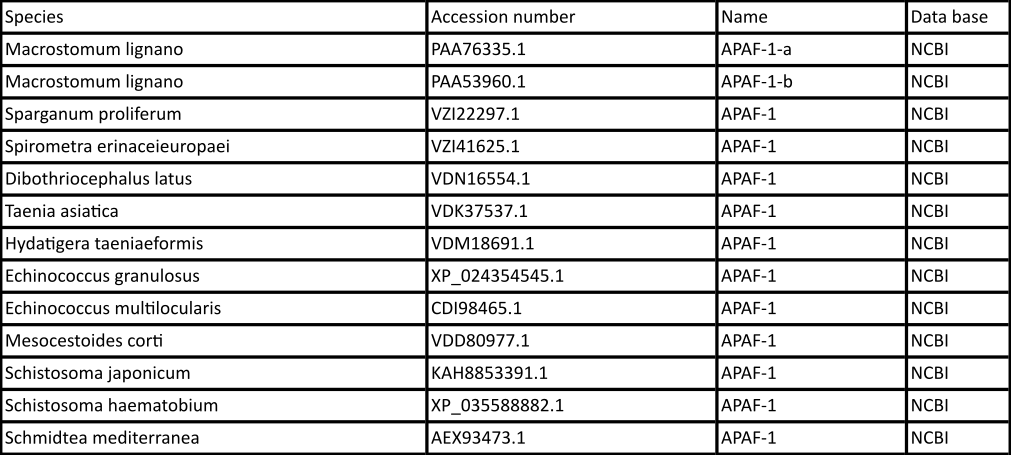

**Supplementary Table 7:** List of species and sequences of CRADD used to conduct phylogeny analysis of Figure 6.

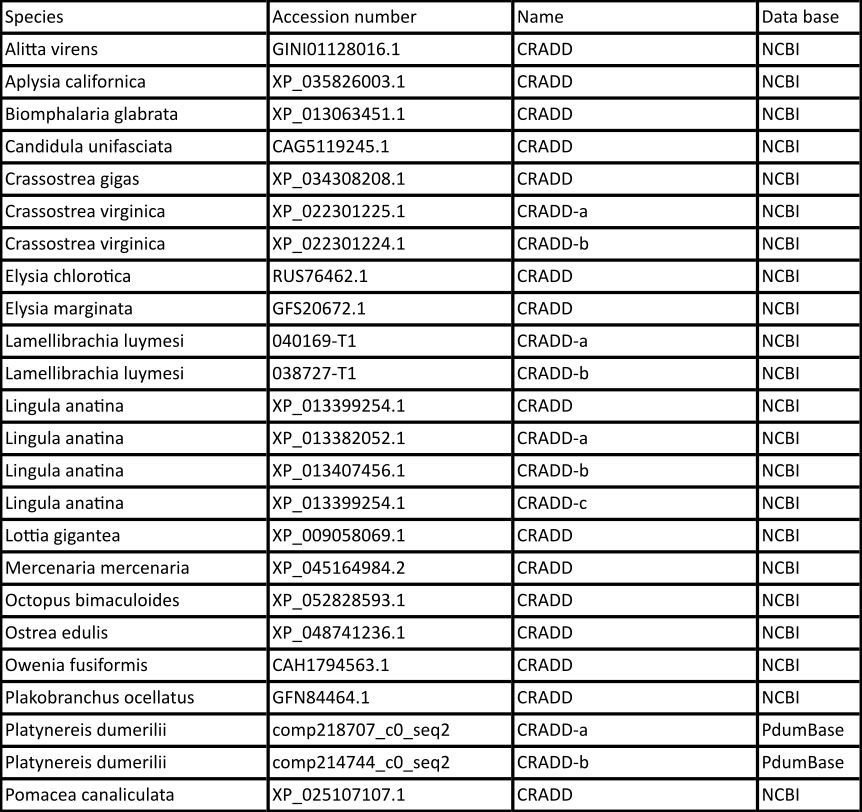

**Supplementary Table 8:** List of species and sequences of BCL-2 used to conduct phylogeny analysis of Figure 7.

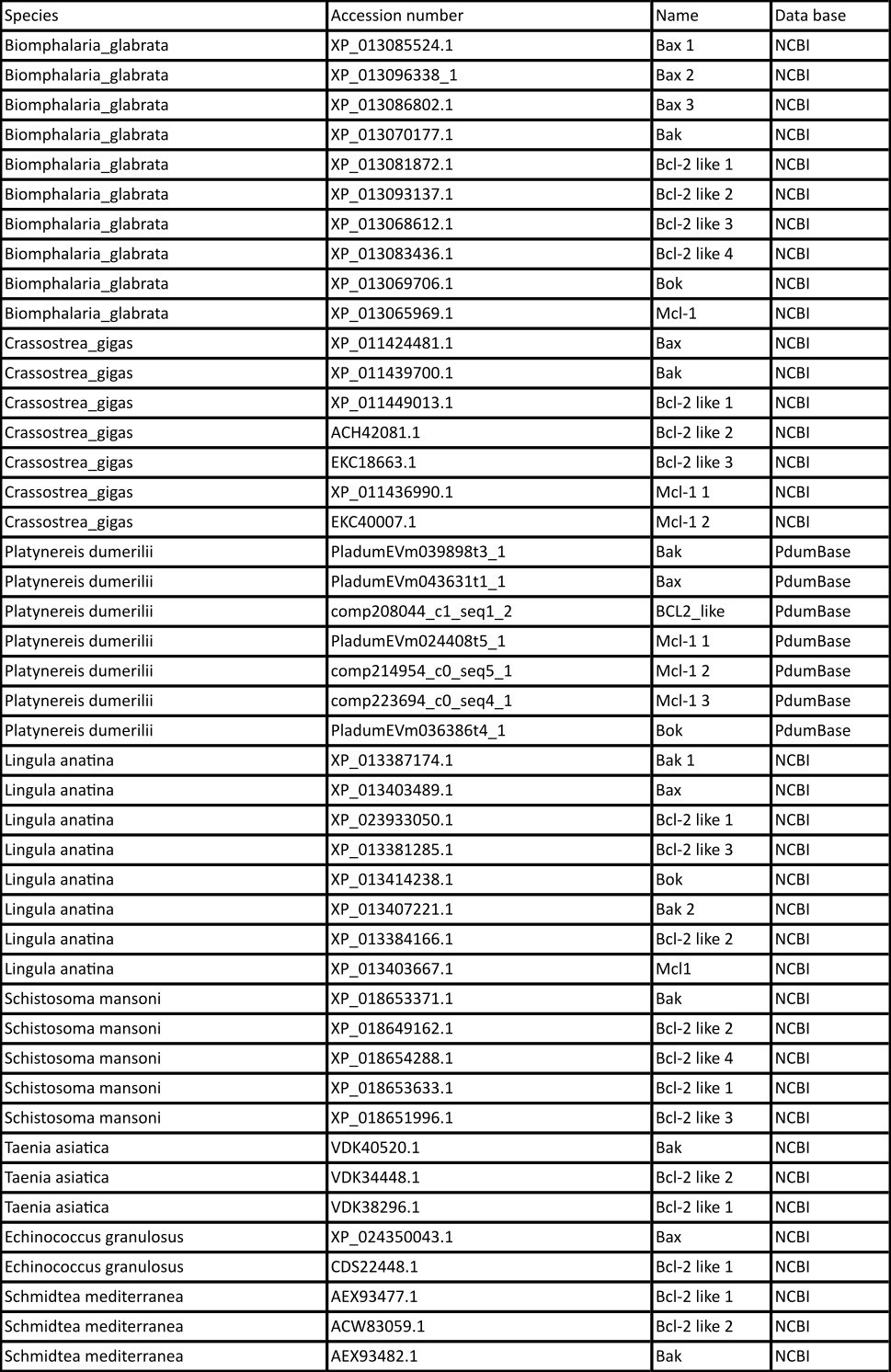

**Supplementary Table 9: dN/dS ratio calculated on the ORF using SNAP v2.1.1.** Apoptotic actors with CARD-domains have a *ratio* indicating a high selection pressure (dN/dS<1), in opposite to apoptotic actors with DED domains (FADD and DED-Caspases) characterised by a relaxation of selection pressure (dN/dS>1). dN/dS<1 are red, dN/dS>1 are blue.

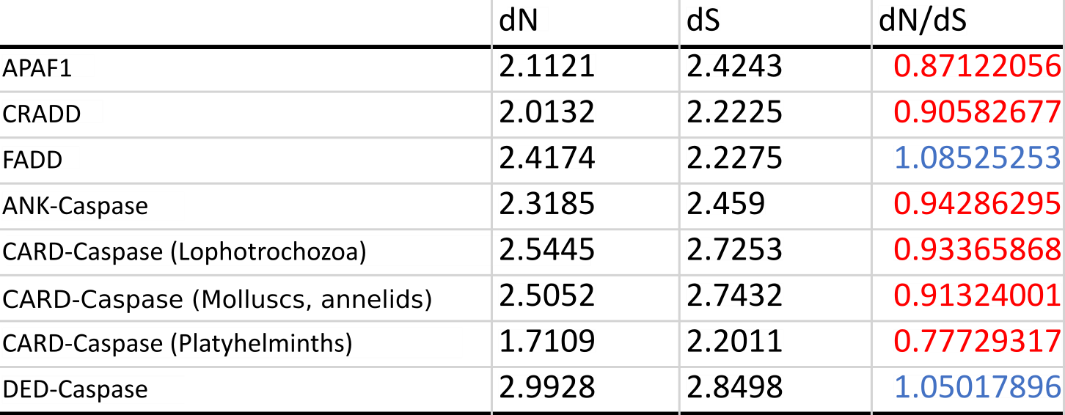

**Supplementary File 1: Caspase repertoire of annelid species which are included in analysis of Figure 3, Figure 5, and Supplementary Figure 2.**

>CAD5120521_1_DgyrCDS9086_Dimorphilus_gyrociliatusOK

MTDKTDAKPDFQQINKRSTDHEDARTYRAAGEDLRVSDEYPMTSSRRGRFIIINNVNFLG

RTGMNTREGSDRDAISLKMRFKALDFDVVVYKDQSTAQMEKIFGDAAANVDYNRRADCFG

CALLSHGDHGIIYGIDGFIEIEKLTEPFNGDKCKELAGKPKIFLIQACRGTKFDSGVEIA

DSEAVNPGADDIFKNRTFKIPALADFVFAYSTVPGYFSWRNSREGSWFVQAITTVFEKYC

DEKDLLWMLTRVNYEVAYKFESKTNQESMRQKKQIPCITSMLTKDLYFRRKY

>CAD5120522_1_DgyrCDS9087_Dimorphilus_gyrociliatusOK

MDSNDSKNKGSIGNTAIPNEEDPKQLIIDLCKLFYSIGWCTGTGGGISIRKGNEIFIAPS

GVQKEKIELEDIFVQDIEGKDIYLPPSEKKLKKSQCTPLFMNAYRLRNAGAIIHAHSAAA

VKVTLITPGYEFKISHIEMIKGLKRDSTGKKYNYHDTITVPIIENTPFECELEEVMARAM

KAYPDTCCVLVRRHGFYIWGSTWQEAKTMSECYEFLFEQAVFMKQMNIDLTASPVIPQAN

IRFDLSAKQDMNDKLFDKIKLTDCENSCKLSHVEYEMTSKPRGQFIIINNVNFKEVTKQE

TRVGSDSDSLKLQNLFESLGFEVKEFRDETVSGMRSILQDAAKRKSFNLKSDCFGCAILT

HGTLGAVYGTDNIIRINELAALVDGQNCPELIGKPKIFIIQTCRGLLYDDGARIECTDAV

SEGCGTFISRASEIAETYRELVFSDFIFAYSCFSDYVSWRNETGSVFIQAITTIFEEYSD

SMDLMKMLALVCEKVATYESNTKDPNLNGKKQIPSILSYMTKILKFKPGNY

>CAD5120530_1_DgyrCDS9095_Dimorphilus_gyrociliatusOK

MAGHQQEYFDVNRRRSETIEEEYGTSQDGRHFIIINNMEFKKNLRLPSREESEYDKINLK

MKLRACGFNVKVHNNKTTTEMKQILKDVSVKTQYKNPVSVFGCALLSYGQNETIYGTDGP

IEIQELIELLKGDVCSVLCGKPKLFIIQACPAIKIGVEETDGNTFPKKKKRNLSSLNEKN

IPTMADFVYAHSSIPGFMPWGISQNNDNAGSWFIQAIIFVFNKYRNSHDVLTMFTIDDDD

DGGGGDNCG

>CAD5115032_1_DgyrCDS4053_Dimorphilus_gyrociliatusDEDDED

MLESDCVGNPDSHQQHEVEWIHKLYKISAELKIRDINSLKFLLKDLVSSSKLEKTDRGSE

IMTLLIDNNKIDLTTGNMDCLQECLYYIGRRDLLRHLNISSRQVMDGLKTKRSTLSNYRL

LLHKLSEEIKSDDFKNLKSYVEMSGCTSRGKLDEAETSQQLLMLLEEEGLITENDIKFLK

AAFGEILSDCGYLEKLIEEYESINGQILGQSDFETDSGKKINIPSKLTLNIERESYTIDT

KPKGYCIIINNAKFLNDCLPVRNGAEEDIRNIQRVFTKLNYAIKLFRDLTASEMEKVLGN

IASENHSNFSCLVCFVLSHGTTGHVFGRDRRKLEIMKIPNFFGASKCPSLSGKPKLFFIQ

ACQTEGETALFVRRRNIESDGVEDNIVSDMADILIGYSTVPGLKSFRSRSEGSWYIKQLC

VQIARNYDKHDVLQILTNVNREIGKVNAPFENGYYKQTPAPYFTLTKAFFF

>CAD5125939_1_DgyrCDS14119_Dimorphilus_gyrociliatusP20P10CARD

MALRKWRNFLSSYVLRRNNNDKLDHRIVKESNSKEKREKNLSDFVNEEREPLIEGKFHLH

NSSNIDLEDCYNTKGQMKGRIVIVSNTHFEKKEILRDRTGSHWDCRNISEIFTTLGYDVE

IWNNCTKPEFEQKLRKEINSTTHEIANSFFFVVLSHGDENGFYLKDGNPNNNRSSGFYEF

SQIERKFSNENCVGLSGKPKIFIYQACQGNRKVSCSEDQRDEIDFFHHPESDILVAFSTV

KGYVSWRSPTHGERMEINEKSVITGNLVYLMQNLDADILLLATCQEKLILNEHDVEEIRN

CSTTLERNKLFLDKIQRRPVFERFLQVLRIVQKGFIAEKLQNYMNEHQLDGHFQNVDGN

>ALB38959_1_caspase3_Urechis_unicinctusOK

MKGVQEKNSIHDVELLEKVMRGLGFDVITHQDLTTSDIHKELAKYANDNEYNKNANCIGV

AAMSHSKKEGQFKAADMTISIEEILKPFKLNKKNEDLRNKPRFFFFQACRGTEHDTGHRI

EGIKTDNDSETSVPDSKVDRVLSVDRDDDQTEDDDRSQLLPVVSDMICAYSTYKGYTARL

GTDGSIFIQTLCSTLTKGAGEFDLVELLTIICNDVTQRDIKFKNKTTEVEMTQKQTPTIV

SSLTKKFYLKQR

>ACP41139_1_caspase_8_Tubifex_tubifexDEDDED

MSHNVESDACIAHLDYRKLLIDVHENLVSRDLAALKFLCSDIMPVAVLERVTSGLELLDL

LETRRYIVAADNVLFLAELLYYIGRKDLLRKLNVTDNEVRTSLTDKQHSHVKQHRVLLHT

ICQELTTKELLSLKFQCSAKITRAKLETIKTPQELILELERQRVITHDNFTYLLSLLRDI

HRQDLVLKVSQFAGLDKSTGKQTNLPVNTTPEYKRLESDDCSLSPTNRVVSLQPAQVQEQ

WIDKGEVDQSRLVPCYDTGMVPGTLQQPGLVFSEDTRVVPGYNEDSTAELHRRLAVGERY

VRQDDSLIMSQVPQNVELSRYLPVIIASGGSQPHASAGIDQLAIGQQLRLSTLHDVSIDD

ELPCYTMNRKPRGLCVIVNNEKFYKLDSDNDSKKMPDRTGTQIDCDHLQLVFESLQFKVE

ILNNLTHDELARGLLSAAGSDHSMYDCFVCCILSHGALGTVYGCDGKTVRVYDLMSYFKG

ASCPSLRGKPKIFFVQACQGTEKQQGFPADIETDSACAPSEVETIPNEADFLLGLATVPG

YVSYRSKTQGSWYITTLVSMLSRYANRYDLLSILTRVNERVSEAFTKEGNLKQVPAPMFT

LRKKLYFC

>ACH96578_1_caspase8_Tubifex_tubifexDEDDED

MSHNVESDACIAHLDYRKLLIDVHENLVSRDLAALKFLCSDIMPVAVLERVTSGLELLDL

LETRRYIVAADNVLFLAELLYYIGRKDLLRKLNVTDNEVRTSLTDKQHSHVKQHRVLLHT

ICQELTTKELLSLKFQCSAKITRAKLETIKTPQELILELERQRVITHDNFTYLLSLLRDI

HRQDLVLKVSQFAGLDKSTGKQTNLPVNTTPGYKRLESDDWSLSPTNQVVSLQPAQVQEQ

WIDKGEVDQSRLVPCYDTGMVPGTLQQPGLVFSEDTRVVPGTTEDSTPELYWRLAVGDRY

VSQEDSLIMSEVPQNVELSRYLPVIIASGGSQPHASAGIDQLAIGQQLRLSTLHDVSMDD

ELPCYTMNRKPRGLCVIVNNEKFYKLNSDNDSKKMPDRTGTQIDCEHLQLVFESLQFKVE

ILNNLTHDELARGLLSAAGSDHSMYDCFVCCILSHGALGTVYGCDGKTVRVYDLMSYFKG

ASCPSLRGKPKIFFVQACQGTEKQQGFPADIETDSACAPSEVETIPNEADFLLGLATVPG

YVSYRSKTQGSWYITTLVSMLSRYANRYDLLSILTRVNERVSEAFTKEGNLKQVPAPMFT

LRKKLYFC

>PE_Scaf983_2_6CARDOKParaescarpia_echinospica

MEERHREKLRCNRVAFIKDLIVNEDLLAHLEQNDILTQTMIEEIEVKPTKQKRASELLNV

LTKRGPGAFSVFVDALVCTNQTHLATLLESGIGAQAQPRQNEHAVPLRLPSQPMDDTEQG

MLAVKIHPVDEIDAAKKINEGDVYRMSSQPRGRAVIISNEHFSRGQPTRDGSIFDLINLE

ELFIQLGFDTVTHSDLTAKAIRDTLEEESRRQQPSANAFFLCILSHGAKGHIFGADGKTI

SIEQITTMFDGKNCPQLSGKPKLFIIQACQGEDSTRGVPQADGACESVDTDMAQRLRELR

LGDTADSMPSLPNVHEKADMVLAYSTVTGYKSWRLPESGSWYIRELVKIFANFAYKEHFM

DLLVKVQHQVSEKHTFFDEHKSSGLYVQMPETRTTLRKKLYLFPGFYG

>PE_Scaf8674_5_2CARDOKParaescarpia_echinospica

MLCKHHKEILERCRPELVEEIEPDLLFPELRKRKLVTSRQEDKIKRNGMRVRAEQVEDIL

DLIPRRGEHAFDGLCEALTDSDQERIVKAYLKPTVSVTTEQEPPTKKARKHDPTQNDEED

MDEGIDLHDGPISVHVNQSTEEFYKRNYKEAYPMYQSCRGLACIINVKRVVGTDDRTGTE

IDCDNLQQLWTQLDFQVFVYNDEDDLSVEGIRDKVIQFAAQPGHQHSQCCVLCVLSHGKN

GAIYGTDGKLLPVAELKYQLDAKRCPALKDKPKLIFLEACRGDLADTDVADCPTQADFLV

GFPSQPGYKAWRNSRKGSWYISAIVHVFMEFAATKNIDELMIKVNNIVSRKLSRTHLNED

NNKAQMPEVRHSLRHPLYLFPGITDDEGC

>PE_Scaf9697_0_12OKParaescarpia_echinospica

MLLLNFFNYVLFAFCRMGTRLDEDVYRMDHKKRGLFLLINNRCFDKKTEMKERTGTDVDA

ANLADVFKGLGFDVRSENNQSTTDMMKIFTEVAKQDHKNSDCFVCAILSHGDNGLVYGTD

GIVKIDWIASLFKGSNCPTLASKPKFFIIQACRGLEFDDGHDVPDGTGESEQRIYRIPEM

ADFLYAYSTVPGYFSWRNQVKGSWFIQAFCKVFKEEDTDKTDRNILTLLTRVNRIVAYDY

ESHSSLEGFNSKKQMPSIVSMLTKQLYL

>PE_Scaf3455_5_6OKParaescarpia_echinospica

MGDEPDANKVLTGSGGRAAMNAAECMDIDVDEDVYKMNLPKTGVCVIINNRYFEKSTGMS

DRSGTDVDAANLYQLFMTLGFDVRLEQNKKTTEMLRILNRVAKEDHSNNNCFVCAILSHG

DSGIVYGTDGVIQLDMLINPFKGKQCASLLGKPKVFIIQACRGNKLDHGVEVADAIGDQP

EQKIYRIPEEADFLYAYSTAPGYFSWRNSNKGSWFIQALCKVFKTDAEKNDILTLLTKVN

RLVAYDFQSNASREDMNRKKQIPSIVSMLTKQLYLPQKK

>PE_Scaf3228_3_5OKParaescarpia_echinospica

MAAEPDAVPSRSCPSVLLVSTDWREWDITSSMVTVNSIVAQCLTSTRSDILLNCTVVQFQ

DSIPAEVKAAADRANVRLIGCERPRRVSAPKKDWLTRFWEQHYPRLVNGHNFSHIIGYFP

QTAKTALSIRDRIRRRGAECTCVLFIEDSRREELTEESAKLINQANTVYTLGPLLFREYN

RHISRPHRMIFPVPYMGDHIATFHPHTPGTVSPIDIVTFYPENGIPGPDEEYQHKTLVLV

IKAISNVAKSYFGRCRTNNIEWHIYGVPDECSVQEQLKAVIREHSVFPELTWRLHKQPAD

CTRILQLASLCVIPWHSSYDPNALYPLLYGVPTLMCASSGVAEMFFNDVPRNDDGSTLLT

PMAVVNCSGDTGGDGWTAKIHSIISNPVDAGADFLELGRRIRESTALQEGMRAFSVATFG

GQDVQLSSNMAALGQRPVLSPSTSTGVQHHGHPGQIGRPVLGRRTTQTDISGPSPSPRPG

HPIPQGNGEPSPGPGPHVPLGDGRGNVKPWTDVNKYTRLVVDSENVYTMTASPRGYAFII

NINKVQGVNEERKGSEVDEHNFCNLFTQLNFLCITNKDLTRQGILDELNTVACSEALWHH

NCLVICLMSHGKEGVIFASDNREVAVTEILEIFSRCEALKGKPKLIFIQACRGKHTEKGE

TRADGPTDNAAARRPPVMSDMLVVYPSSLGYRSYRNTVHGSWLISKLMQVFSEHACEEHI

IELLTRVLREVSQMETSGRDVATGSMEQPEAHWALRKLLYFFPGM

>PE_Scaf10389_4_3CARDOKParaescarpia_echinospica

MEKQHEARIRNNKFELQKRLIVDELLLAHLQAGGVLTDGMLQDIEVKQTRSGKASVLLSL

LPKRGPTTFKVFVEALIKTGQDDLAKLLDPHQTLVSQNGKQPEKEIQTSTEVSNDPEDPP

ASGRVDKKEASASEKKETSGKESTVTPEKNETASFFNEGESPFITIEVKSPTVKVHLETK

FIRNGRAIIINNENLSEEQSRDGSLFDVTNTSELFKKLGFAVVTKPDLTLQKMKDVMQSE

SKNCDGIDGLVLCIMGRGRCGQVYGTDGKTIAIDQLTEHFNDVNCPALKGKPRIFITETC

WDGDKSPDHPQSPGVDDLDKGMQSLNISHPNQAWQENNTSDVHNNVMEVHCTVKGYKTMR

SDKQGSWFIRALIKTISEGSHEDDVVSLMKEVRRKWTETKITDMQITVTASEPSNVYFHP

AKET

>PE_Scaf678_3_3OKParaescarpia_echinospica

MDRADRGVFVIINNRHFHPSTRSKERSGTDFDAARLYELFMRLGFEVRLENNKTKTEMLM

ILHKAAKEDHTNRDCFMCAILTHGDVDRVYGTDGAISIDTLVSPFKGQQCSTLRGKPKVF

IIQACRGTMLDHGVEVADGKYEPEQKIYRIPQEADFLYCYSTAPDYVSWRDENNGSWFIY

ALCRVFLNDSSRLEIMAMLTKVSRLVAYAHESKTVCKDMAHKKQIPSIVSMLTKLLYLSP

K

>MV032009010_CARDp20p10Metaphire_vulgaris

MMTPTDQDNLLRNRLLLVRQTKVTEDLLSCLLEDYTLTVEMKQEIEAERTESKRVGVLLD

TLMRRGNRAFDRFIYALVQTDQDSLALSLDKDRAHPYIETRNKARQLNSNPQLNAIGYSQ

NSHWQGPFQYMMAQIMMSGAGHGVSSSVAGALVSHPTDELPNNSAVDANYEDIRVALNEI

NGPNARSIMDIQVLPGTNFHLANNHQQVYPISHLIRGRALIINNQKFRLSNMFRDGAEYD

RENLKELFQQLSFEVIVRENLTAQQMKELVKEESLNPAHHNADMFILFVLSHGCHGNIYG

TDVENVSIDKDITSCFSPENCPALMGKPKIFFIQACQGELRGRVYSPDSAEGDAISEIDP

KLIEKMKELQLGDKEGYVPEYFNRQDVVMSYATVRGYRAWRRTDLGSWFVRCLVYVFSRR

SHSEHLLDMLTEVNALLTQLKTSGVEGRYGQLAEKSDTLTKKLYFFPEKLEWQQQQRQVK

GRKAALMCYV

>MV021001290_CARDp20p10Metaphire_vulgaris

MMSQKHRDVLIKKRLKLVQGVKVTEDLLSCLMQDYTLTIDMKEEIEAERISSKKVGTLLD

NLMRREDTAFDRFVFALVETNQDGMAQLLDENSAEAYIATRNAIRRTNNHASDTPSAAQI

PMPATSSLEVQQLISMFMSAATTAAANQMGIALSSSSSSGSVVAQPHDGIGNEDTVDNKL

EEDRALAIINGHSATKEMKFTVRPATNLHLINHHQVYPMNYGLTRGRVLIINNHYFLDSQ

LASRHGAGYDRKNLKELFEKLSFTVEIHDNQSTDQMRTLFEAECTNPDHGKASMFILFIL

SHGSQGNVYGSDGAKLSIEDDITSCFTPEKAPLLAGKPKVFFIQACQGEIKGRLYGSDSH

ENEAVSEKDEEVTKQMGDLKIEDKKGFVPGYFKKSDVVMSYATVKGYLAWRHTDFGSWFV

RCLVHVFSQRAHSEHLLDMLTEVNSLLSKLKTEESCGQLAEKSDTLTKKLYLFPGADPRT

SCSVPK

>MV002005420_CARDp20p10Metaphire_vulgaris

ILQMDLKQKEVLMSNRLNLVRDLIVNDELLSILRTNNLITEELQEKVQANPTAQGRASCL

LDILPRRGKRAFSRFIKALNETQQKHLTTLLCSNEGALECTDLDFTDLACDYWKKALDVM

KNEIECPDTHLMKMKVLPMTEDFRSKLRLQSDVYRLDPERKKLAVIINNKYFSKSPEREG

SQNDIENFSYLLEQLDFTLIRRQDLDVVGIKQQLEDARMKLQSGLYDVFVCCIMSHGAKG

VVYGIDQNELNISNDILTGFSDQNCPEMVGKPKLFFIQACQTENGTDACPAVSDTGKAVW

SNSDLFLAVATAEGYKAWRHPELGSCFIRTIVYVFSYRACKDHINDLLAQ

>MV001032950_p20p10Metaphire_vulgaris

MDKSDAMPPPTMSQEKPRGSLSPDVNLHSTNGSLGATSSGAVDSGYESRESISSPEASEN

NVAASLGPVSSTHRLVDEAHCYRMDRRKRGKCIIFNNREFDASTAMNSRPGSDLDAKNMY

DIFQSVLQFEVTLLHNQTTMEMLQRVANVAAENHHDSDCFALVILSHGREELVYGRDGFV

KLSTLIEPLKGSGCPSLIGKPKLIFIQACRGEQFDEGVNVSDAKPDDERPTAEKAYKIPT

EADFLYAFSSAPGFYSFRNSTRGSWFIQAITEIFKKHGTELELMQLLTRVNRKVAYDFES

KSDNKEHSGKKQVPSIVSLLTKELYFSKKK

>MV006006680_CARDp20p10Metaphire_vulgaris

MEPLWKFRSHDGVRFKATDRLQLRRLNEDERGAGDDFDDCLAPCFTASIRPNRRSQFRFK

KEHEKATRDNWHFDKIALRIFLSLVVTSCSDHRKRLQSNTVTIVKNLVFSEEFVSVLMRD

GILTPEEKERIEAEKTRSDKSRYLVMDTLPRRSDAAYPVFIEALVESRQDSLAKEIDSSL

ASKFIEVRDKNDVEIQLQHPVKVSEASVDFVWKPYHLDESKTWYCKLFDSSSDIKDLGIE

LCKMTPDRKKNIEEMMGKRKVYRLIRHNDGFFGRAIIISNQYYTNMNPRLGSDIDCENLV

KLFEQLHFRTVVRKELTGKEMKKFLEEERNNEEHHQYSLFVLCILAHGNNNKVYGIDGND

GKENDGAVDISDITSWFSGVQCPRLAGKPKLFFIQACQGGKVMLSVSPKLRNSDGPSSTA

VASDYVRNAVTLHPDGYPDFTDSLPLNIPEAADLLVAYSTAPGYISWRHPEYGSWFIRCL

IYVILNNAYEEHVFDMLTEGFWSGLAKAPCTESIAEKGWWTQSRCRSMRFKLSTSIHPKR

PFSRKVDGYRHPSCVRPFA

>MV041000350_p20p10Metaphire_vulgaris

MSKLVQSSKEQGQQGTSVLSDVAQRPAQSLQTQQDVSVIQQSFGLPETLSNSVITSFAAA

SIASGGSGHHVRAIDPNHCYKMDHKKRGKCIIFNNKEFSKSTTMPARHGSDFDALSVREI

FRDLLQFDVDMHDNRTTIEMLQIVAKAAAEESNHDSDCLALVILSHGGEGGVVYGTNGII

KLEVLIEPLKGLGCPALIGKPKLIFIQACRGMQLDEGVNASDTKRNKTAATLDDAAYKIP

TEADFLYAFSCPPGYYSFRNAKQGSWFIQAIAKVFKKHGTKLEIMQLLTRVNQTVALEFE

SNSQNKEYTGKKQIPTIVSLLTKELYFTRNR

>MV005014510_CARDp20p10Metaphire_vulgaris

MATKDNWEDVIRTNYTDLLKMIDTDNGLHGEIRRLRIFSPIALKSFWDLKDKTERIDKML

RNLQRKDYESFNLFCEALKASDQGHVVKNHLSRVKDVSSRQVLVTDSVADVDASHNRGIS

HEMKACFQHHWNDLIAEIAFDEDLLGELEKYKIMTPIQLNKLRDKATEAERTEYMLNILE

RGSERQFESFRSALRSCNQHFIADDYLRPRSAQQRQAINRSSSLADPLPAGNPTSSAPPR

SAPVGREDVPPFQCQQQETQLNVGNHSVNIEELCVKTNEQVRLEGAYRCDSNPRGRAVII

NNEFFLRGHLTHREGSRNDVRLLKDLFQFLHFDPKVHENKTDEEIMKILKEESKDEGNSQ

CDMFVLFIMSHGICGEIFGMNGLTVKTEDILELFDGKNCRSLCNKPKLIFIQACQGEMKT

VGVAADEGSTTVTDSGIYVSVLKKTNSADDFDLSLPLRPLPVYQNDAVPSETVHEKADIA

LIYATVRDHVAWRRRDTGSLFVKYIDEVFRSRASKEDLMSMLTQVFERLHSHRDDNHHYK

QIGEKRDTLRRKVYLLP

>MV037001320_DEDp20p10Metaphire_vulgaris

MSPCSIAELLYLIHQNRLISKLQYTVEIVKQIVDEGKGCCNPCRVLLFRIAEHITSDRLD

DFRVFCPQISRAAMEKIKDPLDLFVALERLGLINVLENDLQFLREILSEVGFVEEVKHID

QFLRGQRNKSTNALSSSSSSSQSSHHQSPRNSHYNVRPNKNPVSLRPYYETTSSNNTNAN

ISPPEATEFNFPRQFPGEQPSLCTYSTIRMCTEDSSADGQTAGQATSANQCVNSDDFYPM

THDPRGICLIINNEQFHGENNLSRRKGSDRDCRLLEDVFKKELKFDVKVCRDLTDDEILE

ELEQTAARDHSQFDCFVCFILSHGEEGSVFGANGKAVQLKHVTGCFVPGRCRSLVDKPKL

FFIQACQGDSFQEACVIQTDCKGDDAEARPAPDGVAALKESIPSLGDFYLCLSTVPGYVS

FRSPALGSWFITALEKMLRLYAASEDLLRIMTRVNREVGRATDKKDGHICKQMPMSTQTL

RKDVKFL

>MV003009660_DEDDEDp20p10Metaphire_vulgaris

MEQFQNVLELFQELEKLGKIGEPNNFMMIAEMLLHIGRSDQLKTLNLTKEMIAGIVAESP

TLDPFRVLLFKVSEQLTSDETKMLIFIYGAKLPRSKKEIIKNPEELFLTLERHGVSLKSD

PNSITGMMKSINRTDLVKMIQKYYERIKLNRCSDDDKTIYCNQKLDSTLNVSLRDSSRFR

VERDPVNKRTAIAQRNDFGLRREMPSCEDNNATCHRSNIVALMEDEHFDLPCYPMTRTPR

GYCLIINNEVFSSVPGDQTSRAMPNREGTQRDAEMLEYLFRQKLNFKVIVRPNLTAEEIS

KKMADMAFNVDHSPYDCFVCCILSHGVLGSVYGVDGRTVEIKNLMNFYKGASCPSLLGKP

KLFFIQACQGKDKQPGFAISTDGPESIAAERVSDEVNLGETIPNEADFLIGYATVPGYVS

FRSKSQGSWYINTLVKMLDKYVERYDLLSILIKVNEEVSRAIAAMQDGIYKQSPMPMATL

RKRIFFR

>MV040002180_CARDp20p10Metaphire_vulgaris

MAMFNDQYWRRSMTGKWKDMISKNRPNLCKTIDTAEFIDHLESLKSTNGKYNQVSAILTA

LLKRPFEDFVKFCEALENSGQRHIIRNFLSSNQEIPEPSPPPPLSLQPSASTQADARPAE

VVDVDPLRGVWKDLIRDNYPSLVNDIDPDNGLMAHLISKKIISEWMKDIFTSKEGRSRRA

AAILDHLLLKRPYEDFLQFCDSLNATNQGHIVQRYLTLNRSRRLNQLDCQGSRTHTVDLK

RTASEVFDDESLTRLRELDWKAVFVETRSIMVDGVETSNGFINELFALGVFNSMSYEACE

EKETPYKQMEFVLDYMRRRPDEDFIKFCTALAKTNQRHIVETLLEYKIENRPLSIARQSS

WVDENAGGDRAGNRPVIAHPEEEEMPSAEEKPHDLGVGDVQHGMEVEETPTRASVICHPQ

DDSMSTSQAPEGATEPPQPHHEIPGYVGQNSDVAIAYSLLPNYVHSSQIHHGAPTGRENY

DDAGPAEIIRVNVKRCNEEFYQEYCNDSYKISGHYRGLCVIINNELVQGMNTRYGTRKDG

QLLADLFNGLEFEVKQFFNLTATGIQQEMERIAVDNDLRYHQCFVLFVLSHGQSLPDQSG

RRTDCIYGSDGNTIAVNDILLPLTDTRCPDLVGKPRLIFFQACRGEHSQTQSDSWSSRTE

TDARAEQTQSDTPLPRPLPPVYRDFLIEYVSYRNTELGSWYVNALVQIFMQKACNTSLSE

MLNMVSGKISKWSTEGNDLPNSRMAQSSNYESHLTKSFYFFPGIAR

>MV031008450_CARDp20p10Metaphire_vulgaris

MREELLEILSKNRPQLVENIDPEHGILNDLLAQNVFTRRQAKTIEVMPDPFRQVEAILDN

LRRRPDEDFNKFCNILCKHGQDEVVEKYLKPQKPASSEIDDDEKHSTGCKSQVLTSGSIQ

VRKSTREFYDQYKQSAYPMCNQFRGKCVIFNVETVRGMARREGTERDLQNLRSLFTTLHF

CVEAFTDPTAKDITDKMQKIATDPGLEQDQCLVLFVLSHGRVVTTMTRHGVASIEHIYGS

DGLPVKTEDILDPITNVRCPAMRGKPKLVFFQACREVFKVGSLETDAVATDAVCAAVATT

ISSTSSTSSSSSYAISAASASSSQPPISFKDFVVGYSTESGYLSFREVQSGSWYVDAIVR

VFSEFAHEMDLCAMLRKVNGIIADQWPTGSAVDNSLRQVATHIDSLTKPYLFFFPGIVTE

S

>MV003015770_p20p10Metaphire_vulgaris

MAVGTDVLVKSSSAVSTVLEQFDFSHPRRGICLLINNHHFNRGLTQQLDRDGTEVDAENV

QKTFQELGFEVLRYDDLTVTDMSINVREVAERDHSDVDCFACVILSHGDEGVIYGTNGII

KIDRLTYYFKGDVCASLRGKPKLFFIQACRGRHYDFGAEVESDATDSVESASVRKIPVEA

DFLMAYSVVPGYFSWRNNVDGSWFIQALCRVFREHGGTLELMSLMTLVNKIVAYEFASCT

DTEFTNNMKQVPCMASTLTKLTYFKPAKVGRLIQPPKGYAKSW

>MV008005580_p20p10Metaphire_vulgaris

MSSEDLSKDDDKDQIDGPFNTLPLDSSEFIYDMRHPKRGKFIIINNRTFQPATGMNERTG

TDADAYNLYNIFSKFGFEVSLFNNLTKDAMLQEMLAAAQEDHTDRDCFGVAVLSHGDADV

LYGTDNIIAIDNFVCPIKSCKSLVGKPKLFFFQACRGNELDSGMLLQSDLPDDNDSKMPS

LVIPMEADFLYAYSTVPGYFSWRNSSKGSWFVQALTQTLSKPESEKMEFVRIMTRVNHAV

SHQFESNASSPEMNKKKQVPSIVSMLTKDLYLGPKV

>MV014009120_p20p10Metaphire_vulgaris

MSDRVDNAGHQDHVSFGGGDDGDVDELDSPFKTLSLDSDEFVYNMKHPKRGKFVIINNRT

FQPQTGMNERSGSDVDAAGLYEIFGKFGFDVNLYHNQTTDQMLKLMIAAAGENHSDRDCF

GAAVLSHGDADVLYGTDGIIAIDNFVAPIKSCKTLVGKPKLFIFQACRGNEFDNGILQAD

AIGDGKPQGLVIPMEADFLYAYSTVPGYFSWRNSSKGSWFVQAFARILNRPETEKMDFVR

VLTRVSHAVAYQFESNASRPEMNRKKQVPSVVSMLTKDLYLGPKK

>MV014000930_p20p10Metaphire_vulgaris

MLSASFPLLGRFELYEGLDFYPMNSTPRGKCLIINIIHFTDKSLAKRHGSDKDTPVLKKL

FEDLRFDVEVHLDVTYETLETLLITMNRMDHSNYNAFVCCLLSHGKLGRIYTSDGRPVEI

LNVVQYFADKQCPSLKGKPKMFFIQACQSGYREVFQDQTSSAQAISIPHLSSDFADRCAI

EELHGKPCVECDAAPPIESTSSTDFTDRAVLVPSFPDFLMSYSTLPGSDAFRNIEFGSYY

VQALDEFLRKGDEIDRALKQVTTKVKKDLKERGMIDGMDLSEQLPFHLTSAMERLIYLLG

HNYC

>MV001010310_CARDCARDp20p10Metaphire_vulgaris

MNDTLCDRLKTLEANFEWRNVLQRNLFGLCKHIEPDNGLIDQLTGSEVLNRKAAESFRVK

STKYSRNQAILEFIQRRPFRDFLKFSDSLCKTEQGHIVDSFFYSGGIETLMQARHGASCP

NFQALDESDASPDPSCPKSSFDWRHHIKKHLTCLNEIIDPDHGLIPSLAHKQVIPEWIMD

KYLKQDGRCSRVQTILLELISKGDRHFQLFCDSLVETNQSHVVTDYLSPNSVECFRHSLR

RSLTCSAVDATGDHLATNWKTLLVKHRTDLTERITPNDDLVTYLEYSGVMNKHCAEHCRD

IEPEYKRNEFILNYLQTRPDKDFAAFCTALKHFDQKHVIDEYLQEENVSSAVQESTSAPV

TPVSHLEHGGDRNCEPAPQEESFQNENSQTENLQTPQEDAFVHLGGIGDGDRGSGLTTAI

QCASERETEQNLEPCFTEERESRSLDSAYSSQLDAEMAVTPDSSEVVEDICLSRPTPIRN

VIVLRSSDEFYNINRSSAKSYTMNKRVRGRCIIFNNHRFSRMKERTGTNVDAQLLVKLFK

RLLFSVEEVKDSKAADIISRLNAVSQDETLENEQCLAVFILSHGDYRNGKCIIYGVDEEH

FTLDDVLNMFTPSNCRQMIGKPKLIFIQACRGEQYDVTDVDGPFENEDQPVISSYPDFLI

GFPTQAGYRAFRDIDYGSWYINALTRVFMARAHNTSLNSMMHQVAGIASKWLSQTGARET

NNKTQFAIYTCSFTKDLFFFPGIVE

>MV026006660_TIRp20p10Metaphire_vulgaris

MEAPMDNSNAATEAVAEDMSKLQISPAPVLDDLMKWHLYIGYSQADREMVKELLAFMRNN

KLKCSVRSSLDVNEQSQTDNIKQGIVFSHKCLLYISPDYLKDPNSQQELKEAKRKFKLLG

SNSVFCIVQSNVNPTQFERYKQFQLIPLSVNENVPDCSRAVKNCLDNYHNKDAILKDYEL

KMDLQLGPKENIGVGVAYTYFFGFLQIVVPGFWEKFQETMANNGINDDNLVKKMLIVIRK

SINSPPRLNEPGVEQTKNLCIKQVFARAGKDRREYTLYVYKLAANDKYKNDFYFIADFPA

CLFALREAFDYELNTTMMKNDDVFEARNEFVLTVKALLSHQDNQAKFGNQWELLLWHDED

NRVELSEFLHQQIESKHLTELSRDITLLPVNPVDDRETFKVGRGVRDYAMYAEGEHYDLK

DSDRGYCVIISNYYFLDQRLDRDGGQAEIDKLEELFKALKFEVVTERNLQGQEMFSILKE

YRDKDHSNKDAFVCCLLSHGSKGRILGTGFTDDSISVVELMELFYEENCPGLKGKPKLFF

IGTCQVGDSVAKDEEPGTNLQSTSATPISSSAGIDMIPPTVDFIVSYACLPYSKANRHPK

EGGFYVQALCKYLKQDLEIDVALRRVNQEVRKRWNEIGAKQDPFHIIMPPQRFLFLKHL

>MV014000970_TIRp20p10Metaphire_vulgaris

MSLTDNGMRSLEGPFTHHVYIGFSPKDKSAVFELRAQLEHRGVICYLRPVTNDVKSSIQE

GIRVSQRCLLFLSPDYVEDSWYQFESEVVLEKAERFCRDTLVIMKSGSSVEVPKKLCGFK

EIVFDLKKIKDEKANARLADIITTEPNVQLAVLPKQNFGYGAALGYFYGFLRLTLPDFRR

RLVEETLKNEPGLLPTTVKKMLIIMPESCFCAVEMKSPGKIEFTDRFVRNREYYFAGEFA

TPLLTMFETYHFGLCHLTKDQLYAERDAFYFTLKAILSHPDNSECNDQFRLLYWSDPETK

EKNRLSLYDFLLPVLRDELALEKLSESKPGKVLSDGHASRTPVPAPMRQFSTFEPVNAPG

HNPSRIYEEIEVYPMNGRPRGMCLIINIVNFNKGHQHLTQRRGAERDTEILRNLFEDLQF

TVQVHVDVNFSTLEELLLTMHLMDHSKYDAFVCCILTHGKLGVVYTSDGKPVEILSLVDY

FSDRQCPTLKGKPKMFFIQACQRGDAEPANAFPSPPLSSDSSARPAIDPDKVQEAMTSEV

SLEADAMGAPHFSVVDDSLPVTFRHKSVLVPGFPDFLMSYSTLPGSISYRDTLAGSLYVQ

ELNDCLRKGSEIDRALKQVPTAIPPDVWHGSLDLLLDTWPEWHF

>MV008011640_p20p10Metaphire_vulgaris

MNRPTAIGHTEARGKLHRAPRRISSAPAVKQGEGRGIYWGGWSGRNDCQNNQIQLKASTR

NSSQEIPSVSLSCAIGKCVVVKSRSSEFIIMPLFENDIQQLEGPFSHHLYIGFSKNDVKD

VFRLRTQLEKRKVSCCLRSFKDEIKTSIRDGVRKSQKCLIFLSSDYLDDDWYTIEVEALL

EKVAQFCRDSLIIMKKSESSVIPEKLCGFRHILFHPDKLDDEKANAKLVDIITSEPNVQL

TVLPKQNIGYGAALGYFYGFLRLTLPDFKERLIKETLLNEPNLMAKAVKKMLIIMPESCF

CPVKMVSPGEIEFTEKFLVRAACRAGNIRRDYKSSLFRIKDKVENEDYYFVGELATPLLT

MFEMHYFDLCGLTKDQLYAERDAFYFTLKAILSHPENEECNNQFRLLYWSDPETGFTDDK

NRLNRQSLYQFLLPVVKEELRLERENERHPSKIGTPISRRLRSPSRQLSSFEPVEVSGPN

PARIYEEVEQYPMKKIQRGICLIINIINFDKGHVHLKPRRGSENDTKVLQRLFEDLHFKV

AVHTDLNASTLENLFLTTHLIDHSDYDAFVCCILTHGKLGVVYTSDGKAVEILSLVDFFS

DRQCPSLKGKPKLFFIQACQRGYAEPSEETGHSLPRDGTAGSRTESDDVQGGMTGLRLDE

DALRAPKIATSAEESVTATFKHKSVLVPGSPDFLMSYSTLPGSISYRDTETGSLYVQELN

DCLRKGQEIDRR

>MV026006700_PROKp20p10Metaphire_vulgaris

MRSPAAAGAGGTCPVQSSLSASVGQSTGGCNGVREKMLQEIRRSPVVKKPTDKKYIVEKM

LIIIPKTLLSPQKLTDKTNIRRTGLSIKQCFHRAGKENREYTLSIYELTDQKSGNVYYFV

TDIPSCLYAIREAYEFEQNSMSEDDADMTRKKFVETVETLLRHHENQSEFANQWVLLEWP

DDDKDNNRMELSEFVSQQIESNELAVSLDSAIPEGTRRFGYAHGISEYPMYSDRPRYALK

NFPRGYCVIISNYEFSNPNLNRQGAEVEIEKLEELFKLLKFEVVTERNLQGQEMFSILKE

YRDKDHSNKDAFVCCLLSHGSKGRILGTGLKDDSVSVVKLMELFYDDKCRELSGKPKLFF

IGACQVGEPVAEDKEPSTTLQSTSAAPVPPDAGPDVSSESSNTKTDAVPEELSANVMRDL

VPSAVDFIVSYACLPYTEALRDKDKGGLYVQALYKHLDRDLEIDIALRNVSQEVRDLWDE

YKNAGGTARPDPFHIIMPPQKLLFLNSI

>MV01400121_0p20p10Metaphire_vulgaris

MSANTGKIVKHKKKICLFEFVTDLLRITQHFCVPEGRTRGHCTVLVYLDTTDKLSARHAW

SSCANKHGIIGPEAWSKNRSDDRYSMGSPFRGLFIIINNEIFQDAPLRKGTEKDAVRLKA

TFEQLGFQVKVYDNQKCIEMLQIMAYAKDLSDDYDCVAVAILTHGSENKIYGVGEKDVCS

LDTVLAPLKDTKKGIPKLVLIQACRGENLDPGHVVQEDHDASSKAEDKAIRVPMESDFLY

HFSTSPGYFSFRNQTNGSVFIKTLCDVFDQHIKENVEIDVIKSMIKVNRRVAEDYESYVT

DPKSKALHKKKQMPSFVCMLRKELFFKKQT

>MV008011650_subtilasephomoBp20p10Metaphire_vulgaris

MGRCRRRWGSIGRWDGQGPGWLVAVIWIAFVRAGCAAVYLNKWAVEIAGGSDDADRIAAE

YGFKNLGQIGSLENYYLFEQRRLVKRSASPSKHHHFQLKKSPEVIWVEQQSVKRRFKREY

KVELDRWKPKLNDPVWPEMWYLNHGARGGHDMNVNGAWQLGYTGKGVVVTILDDGIEHSH

GDLRQNYDPQASFDINDNDIDPAPRYDKADENRHGTRCAGEVAAVANNSRCGVGVAFQAR

IGGVRMLDGDVVDAVEAQALSFNPQHIDIYSASWGPEDSGKLVDGPATLAKEAFIQGVLR

GRGGKGSIYVWASGNGGSMLDSCSCDGYANSIYTLSVSSTSEFGTKPWYLEECSSTLATT

YSSGENYKERKIVTTDLHDRCIDNHTGTSASAPLAAGMCALALEANPNLTWRDMQHIVLM

TSRSEPLRDAQWVVNGVGRPVSHRYGYGLMDGRAMVLLAEKWTTVPEQRVCTTDVLTVNQ

EIEGSGISVTFDTDGCKDSAGEINYLEHVQARISLVYSRRGDLQIHLVSPNGTRSTLLPR

RKYDYQPGEFRNWSFMSVFYWGEQPQGTWTLVLENVGSLPNSGTFENWQLIMYGTKEDPV

NLNDTTPKRPIVNVPAQGESMAGPLKTTSMRFRPGITNRQKPAKPGSRMTTKQATKEGKT

TPRRLLSPTAKVTPSTNLTQMRTLAPKANTTLRIQQGHQTSIVSSDGAVTSQSVNNDVTT

RMPHPGTATEEQAVNICDVECTADGCFGPGPENCFVCRHARYDNRCVSSCPRLTFYDAES

GSCHPCHFPCMECTGSFERNVQSCTSCGPDYLLSTEGNSTTCVKLCPVGFYPDFNMSRCF

PCVPNCLICVEDAINCLRCEDDFELVNSACTKACDESEYRDEDGSCQSCDVICETCKGSD

AMDCLQCAPGALHENDLCVANCSEGYYLDGDVMCEMCHPSCSTCFGPEDDECFSCFHPLK

LFGLSCVDECPTGHFEDNDETCHVCHDGCLACIGPRENDCLDCPPTKFYFDGRCADNCSE

GLFPVEVFNHVTEGMTNQCEVCHGTCRTCSGGSPVDCIRCFKGLVLSSGRCIGSCETRKY

KEGDQCLDCHDSCAACHGPSSDHCLECPSDRFLLGGQCLTGCPYGYYRMDGHCLFCSIEC

NECTDYGADNCVACSSGYVLHNGFCVVPFLEFGISCSGQQFYDAERQNCVPCHDSCRTCS

GSTERRCLSCLQPRVLHSQSGRCLPCCADVDSSQDVAQCCTCNHVFGICVDVWDNLRPEA

TRMSDVLTALPGKITTSAAISSSAASQVYPSSRGTAFLPSSEPFSVIKFPFSSRKPIMPT

ERKVQTTATSALDTPAADMFATKRVGITKADRGERTRYEFAFTFDPSLSRTTGFGRRDVT

RAAEESNLGFWSPWVIVVFVVVCLGSVCAVALTVVFCLVRAKAGDKMCWSKRHYHPVPVV

CNGTMQPVKSTGSQKVFEGDRISDAKMAYSKSYKAGEPFRGLLVIINNVDFSRAGMPNRG

GSERDVARLRNTFDQLGFETELHNDQTCGQMLREMTKVAHSPTLEKYDCLVVVILTHGDD

NNIIHGVGPNNDSYTIDRVLQPLKDIRKGRGQGIDKEMPKLVFVQACRGENLDRGHIVQT

DDSGVNNGGDCIRVPIEADFLYHYSTSPGYYSFRSSTKGSVFIQTICDVYDDHIAKEAEV

DVLKAQLEVNGKVAEDFTSYIPSDPAFNNMKQMPSFVCMLRKSLLLKKRTA

>MV004015670_CARDp20p10Metaphire_vulgaris

MTDVLETQRDTEADYLSMESQRDKLVEIMESEFGLPEKLLKMRALSHRQVDKIRYAVARE

EKNEMILDFILQSGKQQDLMKALSESFQTHLVNYINSNGVYRSEFGLDWPLTEREHEILN

DNEEQLTDILDTCELVPKLYSKQVINLKQRNFISSRTSDQARNEALLGMIRRLSRASYFT

LIQCLKDSHQMHIADILENGGVVLRVHATLETDNLDETETQIANEFTDLLMQRNASSEQA

IRAFNAAQLNIIAIKRGQSIVIYICCSTEDELDRVCKMIDNSEMKTFLQQLFDLLTHNIN

FTVLLVKIYKSDVNKAKQIFQRMKELKNKGTECDSAFASKEDSTPSISFGRNRPIIVAER

QQNMASPTQSSDRYNMGSPFRGLFIIINNEIFQNAPRRKGTEKDAVRLKATFEQLGFEVK

VFDNKKSIEMLQIMADANDLSYDYDCLGVAILTHGNENKIYGVGENDVCSLDTVLGPWKD

TKKEIPKLVLIQACCGNNLDAGHVVREDLDTSSKAEDKAIRVPLESDFLYHFSTSPGYYS

FRNSVIGSVFIQKLCDVFDEHIKANVEIDVVKSMIQVNRRVAEDYESYITNPKDKALHRK

KQMPSFVCMLRKELFFKKQT

>MV053000130_p20p10Metaphire_vulgaris

MAPRLNPSQKNYLNGFAQILRHGPAIICYVNGDGNHHLSLPHPMIRLKKFIEGDRISGVK

MACSKSYKTSETFRGLLFIINNVKFHKTSGMKNRDGSEKDVERLKKTFGQLGFEIKCEEN

RTCDQMISDVETVAFSDLAKYDCLIVVLLTHGEDNNIVCGVGPKVDQRKIDEILKPLKDA

CRGHDLDRGHNVQADDVVSNNKTDTDCIRVPIEADFLYHYSSSPGYYSFRTSKKGSVFIQ

AICDVYDECIKEGAEIDVLRSHLEVNGRVAVDFKSYRPNDPAFHNMKQMPSLVCTLRKSL

YLKKQTA

>MV008001140_CARDp20p10Metaphire_vulgaris

MCRVHLMLSTIIAIMAKGEAQRYQEVFVNDYVSMKCSPNGSIPIQWHHNSDLTYVYAGGR

LRDPYDSRFLSKKDNATGEYELIIPNATLQDAGWYYCSGEISGTRISSTQLTVLHVNPVC

SSNISPRGFVGPNTCDLRNDNMTFSCDVSFSGRAPYVIWRSANDSQTFCSSSCEVKSNRS

ICNLTFRADPLYDGSVFICDAITSSGRRFSCNSEIIKVWYLSADYENSEKRIRDSTRCAV

RSNLNVQYKWTLKQNSERIFLGANQTIRFQQPGIHQCEVEYSIRNATCSSLVKNVTVDAE

DKEQSNQLMAILIPIGIVVAALVLGGLILFLLKQRRKPPVDMSSPLTEKEKLAIDSNRGT

LLELIDTAGLFDLLKKNQVFDEEHLNKISSKSTSNEKNEALLKDLENGTLGSYWQTIKCL

HLSNQSHAAEILETGGFSVECLPADIDRNEPFSRDLYIGFSPEDKETVDALIKQLRNNGI

NCYLPSLDSDKHARIQNGVNASKKCLLLWPSKYVKEKWYEMEKTVVLRKVELFGRDTLII

VKGKTEQKVPKGLKNFKEIVLNKEQLDTGQIAELVTEIQKEPRGIMKFGLADGYFYGYLN

IILKDFRKRLTRDDSTEAPVLADCLPNTLKKLLIIFPESCICPSTMVTDGKVELHKVIKF

KVDRAGKQRDYANSLFKVKEGTEEYYFAGEFSASISTLYDAFKQKEESKEMTEDQLHDER

NVFFYSLKTILNQPKNAAFKDQVKLVYWSDPESDNPNKVDLHKCLMQVIRKEINSEQTLD

YPMYNRPRRGKCLIIRPTTIHSEQGDVTITEEKDHEILQKLFQDLYFEVEIYTNVDSNAL

NKALAETKATDHSSYDAFVCCILSYGKPGEIYTSDGNRMRILDIGEHFSDNQCPSLTGKP

KMFFIQTSQPGGSDPPKDNSSPSAFDFGVVIEPEKVRLLKDDANTSSPMSVDDSLLLPDF

PDFLMSYSCIPEDVVTRNDGSASCYINALDECLRQGNEMDLALNEVTKKVREELAAKAEF

RGKDLSKWLPFHRVSATRRRIYL

>MV018005190_p20p10Metaphire_vulgaris

MKNRDGSEKDVERLKKTFGQLGFEIKCEENRTCDQMISDVETVAFSDLAKYDCLIVVLLT

HGEDNNIVCGVGPKVDQRKIDEILKPLKDVKKAKAKAKEIQKEMPILVFVQACRGHDLDK

GHNVQADDVVSNNKTDTDCIRVPIEADFLYHYSSSPGYYSFRTSKKGSVFIQAICDVYDE

CIKEGAEIDVLRSHLEVNGRVAVDFKSYMPNDPAFHNMKQMPSLVCTLRKSLYLKKQTA

>MV086000030_p20p10Metaphire_vulgaris

MCLPLFSVIVDIRIVQTGEDGFTRQQNKDNCRKIIEKLNDVIEKRKQSCISVHRQLEDEG

IGLTGFSVDHEIVARFCCKRRSSLGRLEELLLSGQLRNMLDEDFRRLPSSGVALHGISDF

KLSAEREAFCKESDFERPRYALKHYPRGYCIIINNFEFPEALKKLFEGLKFKVDARENLT

QEQMKKCLTAYRDKDHSESDAFVCCLLSHGEKGRIAATDGQLFPVIELMELFYDTNCLGL

AGKPKLFFIGTCQGIGEDEKYERMQSTSTEPNPPHISPGTDSDGWLSSEAKKDPADLVPF

TDKEDKDKDRADIVRKLVPNPIXXXXXXXXXXXXXXXXXXXXXXXXXXXXXXXXXXGRPY

AQCEDRADIVRKDLVPSCVDFIVSYACLPETCAWHDSEKGGLYVQALCKHLNQDLEIDIS

LRNVSQEVRDLLNTYVKKGQIYARQDPFHIIMPPHKCLFFGSDNHDP

>Lamellibrachia_luymsi_033454T1CARDp20p10

MLCKRHKDILEKRRPELVDEIEPDSLFPELRKRKLVTGRQEDKIKRNGQRVRAEQVEDIL

DLIPRRGQHAFDGFCDALEASDQERIVKDYLKPNVPVEKEQDPSVIPAKRARKHEPTQNE

EEDMDEGIDINDGPISVRITRSTEDFYQEKYKEAYPMCQPCRGLVCIINVKRVIGTDDRA

GTEIDCDNLQQLWTQLGFQVYVYNDEDGLSAQGIRDKVTQFAALPGHQHAQCCVLCVLSH

GENGAIYGTDGQLLPVVELKYRLDARRCPALKDKPKLIFLEACRGDLADMDVADCPTHAD

FLVGFPSQPGYKAWRNCKKGSWYISAIVQVFMELAATKNIDELMTKVQG

>Lamellibrachia_luymsi_013758T1CARDp20p10

MDDAHIRKLRKNQVKLIKRLDVGNDLLAFLVGNHALTEEMVERIEVHPVGHRSRASDLLA

LLPKRGPKAFDIFIEALVETEQEYLAELLDEVLTAKLLKKKKTTTTVHSQETKEPGEVQL

VDKNFAENLLKKGGVYKMTAIPRGRAVIINNEMFQDGDIRKGTEYDQKKLKQLFEDLHFE

VVTYKDLSAKVIEETLTREKELQKPSANAFVLCILSHGKRDQITGTDRNEIPIEVITSMF

DGKSCKTLANKPKIFIFQACQGVTWTKGQDETDFVTDQMNKVDIDSPDAEPFFPKETKHE

KTDMVMAYATVPGYVAIRNIEVGSWFITDLCDTFAKHAWEDDVLHLLNKVCAKVAERRTS

EGYYQASQFVSSLTKDLFFFPGYSTNSTPVGSASVQATDT

>Lamellibrachia_luymsi_013762T1CARDp20p10

MEDRHRDKLRSNRVALIKDLIVNEDLLAQMEQNDIFTPSMVEEIEVKPTKQKRASELLNV

LTRRGPDAFGVFVDALVCTGQTHLAKLLEPGITGQGKENRNEYKPLRLPSQPMEEDDERG

IISVKVRDVDKNEAAKKLNEGGIYTMSSNPRGRAVIISNEHFQGACQPTRDGSIWDLINL

QELFRQLGFETVTYSDLTAKEIGDKLEEESSLQQPSSNAFFLCILSHGAKGLIFGADGRT

ISIEKITTMFDGNHCQALSGKPKVFIIQACQGEDSTQGVPQSDGSESVDAEMESRLRELR

IGDTPDSMPSLPNVHEKADMVLAYSTVT

>Lamellibrachia_luymsi_005841T1p20p10

MGDTPDARGVMGGGGAANAVESMDTECDEDVYRMDCSKRGVFVIINNRYFQKTTGMGERN

GTDVDAANLYQLFMGLGFDVRLEQNKTATDMLRILDRVAKEDHTNNDCFMCAILSHGDSG

LVYGTDGIVQLDNLVNPFKGRQCPTLAGKPKVFIIQACRGNQLDHGIEMADATGEPEQKI

YRIPEEADFLYAYSTAPGYFSWRNSNKGSWFIQALCKVFQKERGERDILTLLTKVNRMVA

YEFQSNASRPDMNRKKQIPSIVSMLTKQLYLRPK

>Lamellibrachia_luymsi_011053T1p20p10

MVTVNSIVAQCLISTCSDILVYSTVVQFQDSISSEVRAAAERAHVQLIGCEKPRREAVPK

KDWLTKCWEQHYPCLAESHNFTHVIGYFPQTAKTAIAIRDGLRGKGLDCTCVLLVVNSRR

KKPLEKSIKLINKADVVYSVGPLLFRECDRHVTAPHRMIFPVPHVIDVTQPSVLLSDIPG

TMPNKDIVTFYPENGIRGPGEEHQHKMLVSVIKAISEVAKSYFLRHNANVVEWHIYGVPD

KDREEQKAVIQEHSVFLEFKWMLHQQPADATKILELASLCVSPWQTSYDPNALYPLVYGV

PTLLCAGSGVAEILFSDFSKDDESSTLLTAMAVVNSSSDTEGDGWTARIQGIIYSPINMC

GSRQTVAGADFQELGQRIRVSAALQESMKTFSTAMCAGQVSPGPLTSVVVESAGAMATFT

AVSIEQRPAPSTMMSATTLQHVQHAQAMQAGHCGRPLLGRKMPLVASTGDLPRERPGPPM

GPGRNGPTVKPWTDVERYEQLVDSEGVYKMTASPRGFALIININKVTGGEERMGSEVDER

NLCNLFSQLDFKCITKKDLTRQEILDELKDLSKSEELRHHDCLVVCLMSHGKEGVIFASD

KKHVAVTEVCEIFSRCKPLYGKPKLIFIQACRGEHTDEGETRADGPTDNVAVTKPPTMSD

MLVVYPSSLGYKSYRNTAHGSWLISKLVPVFLEHACHEHIAELLTRVNREVSQMETLGRD

VPPGNVAQPEAHWALCKLLYFFPGK

>Lamellibrachia_luymsi_005838T1p20p10

MGDSPDARVATGDTPDARVATGDTPDARVVAGGGGAANAVESMDTGCDEEAYRMNCPKRG

VFVIINNRYFKKQTGMGERNGTDVDASKLYQLFMGLGFDVRLEENKTCTDMLRILDGVAR

EDHTNNDCFMCAILSHGDFGVVYGTDGIIQLDTLVNPFKGRQCPTLAGKAKVFIIQACRG

DLLDHGIDVVDGVKEREQEIHRIPEEADFLYAYSTAPGYFSWRNSSNGSWFIQALYKVFQ

RDRGERDILTLLTKVNCMVACEFESHVPTDESMHLKKQIPSIVSMLTKQLYLPPK

>Lamellibrachia_luymsi_032062T1p20p10

MDHIRLRIPEEGDGDAGRALPMSTGVRAFIGSRTSGRDGGASKREPYHLDYFRVARVKCC

WLNPQSEMAEHDVSCAETSLKNADTQEDDASAEKLNVKMRIQMFNRLAKEQNDETPLMKP

KIKNPVQQVKPPSTQATTPPTEETCRNCLPHEGTTSHATSHASPQSLTGNEQMPRTNQVE

TPPPPQNKSKREADTCDTPVLSGTPIMEGTRVMEDTPNTNMAMSGGSNGARGTDFKSDED

VYRMDHPKRGVFIIINNRRFLPSTHMNERAGTEVDAANLYKLFKGLGFDVRMEHNKTTTE

MLKIFDNVAREDHTDNDCFMCAILSHGDLGIVYGTDGIIKLDTLVNPFKGRRCPTLAGKP

KVFIIQACRGKKMDEGVEVADSPEGSSPKIYRIPEEADFLYAYSTAPGYYSWRNGVDGSW

FIQALYQVFHNDSHRLEIMALLTKVNRVVAYEFQSSTRNKQMSQKKEIPSIVSMLTKELY

LSPK

>Lamellibrachia_luymsi_003216T1p20p10

MGDTPDARVAMGVGGSADAVESMDTECDEEAYRMNRPKRGVFLIINNRDFDKKTLQKRRD

GTDDDAANLYQLFMGRGFDVRLEDNKTCTEMLTILDGVAKEDHTNNDCFMCAILSHGDFG

VVYGTDDIIQLDTLVNPFKGRQCATLAGKPKVFIIQACRGDLLDHGIEVADAIGETERKI

HRIPEEADFLYAYSTAPGYFSWRNSNKGSWFIQALYKVFQKDGGEHDILTLLTKVNRMVA

CKFESYVPTDESMHLKKQIPSIVSMLTKQLYLPPK

>Lamellibrachia_luymsi_011013T1DEDDEDp20p10

MPSRENMERDAIDENMYYRRTLKQVDDGLVSDNISAMKFLCRDFIPGAKLEQVSRGINLF

EVLEQKRLLGNNNTALLIKILQTIGRYDLAKLLEPCTGTSAEELSSYRVTLFKVCDDLTS

QEVDTMKFMCEGHLKKKLPNVKSCENLFVALEESGHVTAANTKVILDLLSDIHREDLHQL

FIPLQQPQRNSPQFRPILGSGAANSRTVSHIASNIQAEQELNIQFEQLDIEKGNLAMPPT

NLPLPARLLSPVEDMPGAQVTPSAGHSAQVRPPIVPQNASLDCYDMSADPRGICLIINNE

FFTCTDLRSPKLKQRRGTRVDCGKLTDIFEKLRFHVSVQSDKTALEIKELMIYFSTQVDH

TAFNCFICCILTHGKLGKVYGTDGATVDIMELTSSFKGAKCQSLVSKPKLFFIQACQGLD

KQTGVQVEMDAPESISPSLQETIPNEADFLVGFATVPGFVSYRSTIDGTWYISKLVEMLD

TFHTREHLLDILIKVNDEVSKVAKETNDTVYKQAPAPLASLRKKVFFC

>Lamellibrachia_luymsi_016793T1p20p10

MGCFASRSSAVLDKEAAPYSGGTHRKGSSHEGNTSYSTNKHESKVDAPVKEVTPVMEDTP

VMDDTPVMEDTHVMEDTPHTKIVISGGGEAGIGIIGTNAESDKDVYRMDHPKRGVFIIIN

NQHFLHLRERVGTDVDASHLYQLFMELGFDVRREQNKTAAEMLMILDKVAKEDHSNNDCF

MCAILSHGDTGFVSGTDHKVKIGQLVNPFKDDQCPTLCGKPKVFIIQACRGDKVDHGVEV

ADNIQEPEEKIYPIPEEPDFLYAYATPLSYYSWRNPEKGSWFIQALCQVFQKGNHELEIM

ALLTKVNHMVAHEFETNLPNNKNKPKQIPSIVSMLTKQLYLSPK

>Lamellibrachia_luymsi_022134T1p20p10

MEIITKVATEDGKFVERYQHDDIITIVATEDGKFVERYQHDDIITKVATEDGKLGTPTAQ

TASLELESYSNKHTRIGLCVIINNKNFNKLLTGQSNRDGTDVDAKALENVFMHLGFDVTR

YDDLSKTDMSINLRQVAEDDHSDADCFVCVILSHGEEGVVYGTDGKVPLDKLLGYFKGDI

CPSLAGKPKLFFIQACRGRRYDFGTVLGEDQVDALPVANTRKIPTEADFLIAYSVVPGHF

SWRNNVDGSWFVQALCRVFEQHGSSIDLLQMMTLVNKIVAYDFESCTDDNFTSQMKQVPC

IVSMLTKQVFFKPKY

>Lamellibrachia_luymsi_001735T1CARDp20

MNKQDESCLRKNKYALQTGLKVDEILLGHLNAKEVLSDGVRQDIESKPTRSAKASALLST

LPRCGPSAFKVFVEALDESGQKHLVELLGPPQTLPPRNEHSPAKMPTEVSSEPKENVASG

KSNKEQTAGSKTKETTEQPSTASETEETKQVLYYDDCKSSFINIELKDLTAKISDNPKKA

RLIKTGRGLVINNEQLTEKESRNGSTFDGINLCQLFKKLRFEMDVKPNLTVQQMKDAVEA

EKSLCNDVDGFLLCVISVGGCGQIRGVDGEKISIDELTANFVNCPALKGKPKIVIIETCQ

EGNEKCPNVEPQSSGVDELEGALKVVHISKPEPSNMIEVRCAIKGYTTARSECGSWFIRT

LIKSIIENAHQDDVVTLMKAVKHKWSNDGDMNMEITVSPDEQSSIYFHPQA

>Lamellibrachia_luymsi_032065T1CARDp20p10

MDPVHKDTLQKKHGLLVERLQPDLILDHLQLDGIFEEQDCDNVRSSRTEQEQSQKLLDKL

LLCGPRAYPCFLKALKAEHAKDLVKELDKQEAESRMAGHNLPGAETSLENADSQEGDAST

EELSVKMRIQMFNKLAKEQNGEKPSVKPKVKNSAQQVTTTQATEATSSGEKCHNGSSYGG

STTHSLTRPQPPLQLQVPEVPVQMPTSDNTSKRGVDTPDTPVMEDAPVMSYNVMEDTPVV

EGTQDVSTVLESDEDVYRMDHSKRVAREDHTNNDCFMCAILSHGGAGFVYGTDGIIQIGT

LVKPFRDEQCPTLCGKPKVFIIQACRGDNVDDGVEVADSIGDSSQKIYPMPVEPDFLFAY

STVPGYYSWRRGDTGSCFIQALCPVFYDDSNRLELMALLTKVNRMVACEFESHVPDNESM

HRKKQIPSIVSMLTKQLYLSPK

>Lamellibrachia_luymsi_019728T2p20p10

MEDHDSVKIHCGTAVIINNISYKDGDRVGTKEDENNLKTTFERLGLKVIIHQDVSTIELL

LKLQTVAEMDHTTKDCFVCAILSHGGQCVLTQGGEQRMCDIIQCNDGIISAGDLWNCFTD

DKAPTLKNKPRLFFIQACRGLGCPATVQLTGDGQSPKIMKSEKWEEEEKKLNYGFPAHLP

IYPEIMLAWATVPGYYSFRDVEGSPFIEYLCEGLRTYREDDLLTILTWVQNQVAIKYTAL

LEQGDNFVSFKQMPWVASMLTKNVYFRNAQA

>Lamellibrachia_luymsi_035533T1p20p10

LHKTTDIEASVDYSRLDHRRNYEETYTIRCGVAVIFNNVDYLEGPRRGTLKDEETLKTTL

ADLGLGVMVYRDVDTTTLLRQLQRVARMDHGNRDCFVCVILSHGGQSELTQEGEQRMCDI

IRCRNNIISARKLVGCLTDDVAPTLVNKPRLFFFQACRGEALPATVEIAEDGRQPRTVER

ESWDEEERRLNYELPADLPVYPDFMLAWSTVPGYFSFRLSSGSPFINHLCDGLRKHKEDD

LLAILTWVQSQVAIKYSSYAPSTVNDPSRRNIHNKKQMPWIASMLTRNVYFC

>GWHPACBE025928_CARD_P20P10

MMTQHHNSILTKQRLLMVRSLKVTNILLACLREDYTLTDEMQQEIEAEKTNSDKVGTLLD

ILVRRGDKAFDRLIRAIVETDQDVIALTLDKSLAQKYISVRDSCRMEVQHSSEPIDQSPG

LSGVQPTQWHNMMVPAGGPQPYPPYAMMMMPVMFPMQNMQHPNMPHSFGGMMQPLPAMNM

TSTSGALVAHPSDVIMEGGKYSVGMTKPGSRLTYSPDELEAVLEEINGGDVQEIMDIDLK

AATNLDDIDDLMIYRMQSLPRGHAVIINNRYFLNRMQNREGAQHDTENLRILFEKLSFTV

IVRENLTAKQIHDLLTTESQHQDHAKSDAFVLFLLSHGSQGNIYGIDGEKLSIEDEITTL

FDPEHCPALNGKPKIFFIQACQGNLRTPRFASDSDKFEPAEMDESLLEKLKKLEIVDKEG

FVPGFHKKSDVVMSYATVKGYIAWRRSDVGSWFVRCLVSVFSRRAHCKHLLDMLTEANFL

MSKLQTSKHEGNCVQLAEKTDTLIKELYFFPSSGSASKSEGSCEKKMN

>GWHPACBE011912_CARD_P20P10Eisenia_andrei

MLPERVGGMKDSYKQILTINRPALVEEIDSENGLLNQLESKKVLTRRQAVSISRNPDPFR

KNELILDALHKRPDDDFDKFCEALKATNQGHVVDAYLLNNATSSASSASSSPSSSPLSAG

TVPNEQLMRLGIDIRVKNELDKIAEKMSNLTLRQGGLIVDSAAVPALSQDPVNHENMDTE

EQAPVFNSPQLIDSEDIVVRRCDSNFYASAKDNSYPMNRRSRGKCLIINICHTNGQDPRE

GAEVDERILHKLFTELHYDVHVFKNLTARQIDNEMVRVAEDSSLVSDQSFVLFVMSHGSI

RYYDGVFTEVIFGSDGDPIATSQILEPLAKCPALTGKPKIAFFQACRGDNYDQVDCISDK

SSEDEDMTRPDACPPGANSLKDFVVGFPTQSGFRSFRNTRNGSWYINAIVRVFMEDACHM

DVSAMLRKVNGIVSKWLPKTNDPRMQQKGQIASISDNLTKPYLYLFPGIVEDN

>GWHPACBE004630_P20P10Eisenia_andrei

MATEENWEEVLRSNYTELLKNIDTASGLFGELKKTRIFTPIALKSFMDCKDTTEKIDRIL

RTLQKKDFESFKKFCTALENCAQEHIVRKHLSKVQEVTCRQVTTTDAAINEDSAHCRAIS

DGMCECFKQHWNDLIAYITFDDGLMSEVERNKIFTTLQINKLKDIDIEMKRTEQMLTWLE

RGSEKQFTDFCNALVTNNQLFIVREYLTPKTIPERRPPESSPPVILENPGQSQEIHKDPE

IRQAPEGSSGVRPLPNEVTSKDIFFSPCNKEFLDRLQAKVSKKVAYKCESKPRGRALIMN

NKFFTNLGTRLGSDVDTENLDALFKRLDFETVIHSDYDAQKMWEIIEKESQEEHHKDYDM

FVLFLMSHGIQGEIYGMDGKSLKTADIVNLFNGKNCPNLLSKPKLFFIQACQGNVKTVGV

TATDAVMMACDSGMPEQPQDVILEEMTRKLDLKDDFNMSIPIEPLPEVQSDAVADNKVHE

MADIAVIYATVEGHVAWRRVDTGSLFVHCLINVLIKRAHERELTGMITEAFSMLHELEMV

ENQGRYKQIGEKRDTLRKEVYILPL

>GWHPACBE002952_P20P10Eisenia_andrei

MDRTDAKPLQSSRSSSSSTKQKYGERKTEVSYPVQESLSRQESAYSSMGSSDGNFSLRSG

SEGDFQRSPKPANGPRSMGDSGFVSNASGASSLSSEHSLSDSLQSACSIAGPKVDTENYY

KMNHPRRGQCIIFNNTKFIKETGMDERKGSDVDAANVYRVFQKLGFEVELRNNVTTVEML

QHVIRIASMNHSDCDCLAMVLLSHGDDGIVYGTNGFMRFEELIAPLKGSGCKSLVGKPKL

FFVQACRGTILDEGVDVTDAVPDSDSKEEEKKYKIPVEADFLYAYSSPAGHYSFRNNTKG

SWFIQALCKIFEEHAGDKEIMQMLTRVNRCVALEFESRSRKKEHSGKKQVPSIVSMLTKE

LFFGKK

>GWHPACBE025691_DEDDED_P20P10Eisenia_andrei

MSQRVEVDAVSSNFFDYRSALLVIYNQLRSSEVQAMKFLCSDFISIAKMERIQNALDLFQ

ELEKLQRIGEPDNVHLLAELLYHVGRADQLRTLRLTKETVARAIQENSVLDRFRVMLFKL

SEQLTTKDTKELLFIYGKDLPRPKKESIEKPEELFQALEKHCVLTRNNPTCILDMMDSIG

RSDLRHLVREHFGSRAGRMPQMPNHSVAGETIRLQSDFDRPLNIGATQDLSRTVAESARN

KLIKDNQYAIWNDPQKSYSSKVCQPVTESDSAIFRQYSQGSRGMAEPSLGGTGLFRTQSC

EDNNAVSQRSAFVQLVEDDQLELPCYPMTRKPRGICLIINNEKFYKDPSDSASRKMPDRE

GTQKDAKLLQDLFSEKLKFQTMTHHDLTANEIKDKLLAAARKVDHSSYDCFVCCILSHGI

LGSIYGVDGRTVEIKDLTNFFKGASCKSLQGKPKIFFIQACQGKDKQIGYPVATDDGLGP

MSVPQQDGLEAEELPGETIPNEADFLIGYATVPGYVSFRSKSQGSWYINTLVKMLHKYGS

RYDLLSIMIKVNEEVSRAIAAMNDGSYKQSPMPMATLRKRIFFQ

>GWHPACBE026958_CARD_P20P10Eisenia_andrei

MRQELSDILSRNRPELVENINPEHGLLNDLMAMGVLTRRQAKSIEVLLDPYRQVESLLDM

LRRRPDEDFDKFCEALVITGQAEVVEVYLKPGKSTSSLGGSSSKQEQESNIGVATESLVN

PEFETIASTSGIRVRNTTSEFYNRFKNEAYPMCNAYRGKCYIFNVNAVYGMDRRQGSEED

LQNLKMLFQQLHFTTDTFTNPTAKDMIDKIIEIARDPDLDQDQCIVLFFLSHGRVTQMRT

NQGIVAVEYIYGSDGEQVATEAILEPLTNARCPAMRGKPKLLFFQACREYSKLDFEEASL

QVTHTDALPVVAESANKPASAPAAPAASSSSSSDQSPISLKDFVIGYSTESGYPSFRNTQ

SGSWYISAIANVFSQSAHEMDVCAMLRKVNGMITEQRPVGAAVDNRLSQVASQVDSLTKS

YLFFFPGIIEGCQRPR

>GWHPACBE017273_CARD_P20P10Eisenia_andrei

MTLLQDAGWRRDMESEWKEVISRNRPDLCKMIDTEGEFIDHLEGFKVFNNRTCQSFKALE

KSNQGHIIRDCLSLNPQATGTLTTDPVVVATESTQADARPEGSDEVRSNHPLRNIWRDLI

RDNYPALVAGINPDDGLFTHLVGKKVILDWMQDTFTTKTGRSNRVSAVLDHLIQKRPFED

FLKFCDALNETNQRHIVKKYFIRDRPSANQSDNSMVEVRPYVCDSTQDRSELLAKYPTLS

NLDWKATFAETRSKLVDMIDASNDFINLLRSLGIFNDISLEACQEREDIFQRSEFILDYM

RRRPDEDFLRFCVALVKSNQDHVITCLLNHRIERPPCESRSCPPQPVTRGGGDGGNETRV

KTVCQPVDDNKPMEEMEMEVNQQQHLDRGGLDVQPRLCEHQGFYVAHHQQEVCNNAMPQA

PQSIDLHPQRQQVICQPQDDDQGENLQSMDTQNPIAMQHPVANEAPYHEAAICGDLNELR

NCQIIEDDARGPVEIDNVYVTHCRRDFYERVCRHAYPMYRPHRGKCIIVNNRIVHGMNMR

NGTEQDGRLLHALFQQLHFRVEELNDASAECLILFILSHGQAIETPAGMVDCVYGCDGQP

VTVNEILEPLTDVNCPHLRNKPKLVFFQACRGDHFQEQTQTDAPPVFRDLLVGYPTQKNY

RSFRNTVLGSWYINAVVQVFMQRACDTSLSDMLSLVTGKVDAWSTEANNHQTGQTATYTS

HLTKSLYFFPGIVAEDDNR

>GWHPACBE020467_P20P10Eisenia_andrei

MIVKNGLEDCDSSTSLETLRHSITKSATIPGILDEYNFTHQRRGLCLIINNHHFNKQLTN

QNDRDGTEVDAANCESTFMRLGFDVTRRDDLTVTEMSIVVREAAEGDHKDADCFACVILS

HGDEGVVYGTNGTMRLDRLTANFKGDICPDLRGKPKLFFIQACRGRRYDFGTELESDTVD

SVDSVDSIVQRIPVEADFLMAFSVVPGHFSWRNNVDGSWFIQALCRVLQEHSETVELMHL

MTLVNKLVAYEFQSCTDSEFTSNMKQMPSMVSTLTKLAYFRCKY

>GWHPACBE002545_DEDDED__P20P10_RECA2Eisenia_andrei

MERLVTTDAQRDIFDFRRALKSIYDGLVQRDVDAMKFLCSDIIPLQKMEQMKTALDLFSE

LQTRRMIFPPDNVRLVAELLYCIGQHRLSRMLPYSCGDVRESVEGGRSQLHPCRILLHRI

CEDITQENFKTLKFFCTDIISKSAMEKICSPHALILALEKQGKIVVNKGNLSYLKGILND

AKLEELVAHVDQFVSRSLTGSQINQNVRSPDTCQSSASNNIVANLCVSPQGISAPNKPLS

QCIHGAGGHQNTDPRVPRVFPLRDVSEEQLTTGAEMQACQGSRATVDTHQHENGLPSYSM

TYHPRGICLIFNNMRFYKDENDPESKQQKVRMGSDRDCELLERLFGEKLDFKVEVYHDLS

DAEICNELFARASTDHSNFDCFVCFVLSHGVNGAIYGANGKLVQIKQITSYFRPSSCHSL

CGKPKLFFIQACQGTERPSVSDIVTDGPTIPEGGVGVMPKAETIPDEADFLVALATVPGY

VSYRSRRNGSWYITTLVKMLQELADRCDLLSILTRVNEELSRTNELVNGLVCTQLLARLC

RKPSLALLEPELFPSGIEPRTMIELYGEEGSGKTQYLVHLIANAILPPSLNGAIIGGLGV

SIILIDNDYSFNLLRLISILEVRISSASASPIKNSESSQENLTGTVPANAESTASVDRCS

PKKPRLSKKNDLCAPSMLRQSTENSTQKFQTSQTHNRELVNEMDIECCVKECLKRLYIVH

CNSSAQLFITLHTLETVLANRSEMCLLAIDSISSFFWLDKFNGTENNLSLIAEIIQKLAD

TYSLVVFATKSSAVYQRPHKKDATDNANHKDEHKEYLCKAWQKLVSSRIIFSKDTSGEID

LGNSQAFCASVCGKERILFTIDESGLRFLSS

>GWHPACBE024253_P20P10Eisenia_andrei

MLSEHCLVPERDDLMKWHLYIGYCQKDRQAVQSLYQSLHELVHTAGLTCYFRSSLDAVEL

SAELNVEEGINFSQRCLLYISPAYIKDCWSEYELHEADRKVKYFGRKSVVALIESSVDQC

HLDKFSQFTKITQPFHDELLTELVSEFELKPILSTFPKSNIGVGVAFTYFYGFLQMQLTD

IIKNLQEVLLGQHAVDWNCVVKKMLIIIPESVNSPPTLTEPEHGSLEQTNLTVRQFYARS

GREKREYSVPIYKLTSVDGQESYYFVGDTPACLYALREAHDRESAGLSKSELHQARNEFC

FTLKGLLNHPDNLVKCSNQWEMLFWPDTKENPNRRKLSEFLCEKIKAKRLTTDVVHMPID

GPVQATAETESYKPQKGIQCSTVYRDMEKYQLKRDNRGCCVIINNSTFSNSSLEERTGAE

VESDNLRELFTYLHFDVDLRKNLSAVDMKVCLESYQHRDHSSHDAFVCCIMSHGMHGSVC

AADGNQVTIMSLMELFYDDACPSLQGKPKLFFIGACQTEASGSSAVSETGRKLIANASPD

SVARPAVAQPVSNLLPSSYDFIVSYATLPKSVAYADRESGGLYVRALCKNLKEDYEIDVA

LKKVSSDVNTLLKDYSEKHDESYTVQKPFHICITQKSLFLCAEM

>GWHPACBE031098_P20P10Eisenia_andrei

MDRTDAKPLQSSRSSSSSSTKQKYGETKTEVSYPVQESLSRQESAYSSMGSSDFSLRSGS

HGDFQDSPKPANGPRSMGDSGFVSEASFFKFEHSLSDSLPSACSIAGRTSTGETSYKMNH

PGRGGQCIISTTPNSPKNTVASMNHSDCDCLAMVLLSHGDDGIVYGTNGFMRFEELIAPL

KGSGCKSLVGKPKLFFVQACRGTILDEGVDVTDAVPDSDSKEEEKKYKIPVEADFLYAYS

SPAGHYSFRNNSKGSWFIQALCKIFEEHAGDKEIMQMLTRVNRCVALEFESRSRKKEHSG

KKQVPSIVSMLTKELFFGKK

>GWHPACBE008492_P20P10Eisenia_andrei

MQTPAKNTNPEDGKDNIQRSQEVHELRYEMNHGKRGRFIIINNKIFGKDTGLEERRGTDV

DSENLQIIFGDILGFEVVVKNDLTRTEMLQLMIAESGKDHGDCDCFGVAVLTHGEEGKLF

GRDELISIESFVDPIKNCKSLAGKPKLFFFQACRGTKLDDGVEVPDESDSTDANTNVPSS

LKQKEILLPLEADFLYAYSTVPGYFAWRNGEHGSWFVQALTDKMKSLASADIDLVRILTR

VNHQVAYKFESNSKTMPKKKEVPSIVSMLTKDLYFPKK

>GWHPACBE026070_PK_P20P10Eisenia_andrei

MHNLVKEEPAVMLGSDGQSNEFLIESELDKIAAIKKVDVESHRLKSYPDSIGRGTGILGT

LSNPIHKNIVTLFHCEKHANKVYFVMEYCNGGNLQEYLKSKRKLDQRTLRCFLRQIAAGL

KVAHDMHIVHRDLKPENILLCFTGPKYPDPDYITLKIADFGIGRFLADDEMARTLCGTPQ

YTAPEILVGRPHRYKVDSWSVGVIAYECQVGSLPTDSLMAFKDVEIPTNITADFADLLNG

LLEKDENKRYDIKQLCNHPFLAANDVKTDFHVVETLSLSRRCRRCQRWSRDVVETDRVGL

LGVQNCFDDSLDFLDFKKDFHDVETDFREVKLPVPSAAHREWIKSEIKSECVYAVSGQAV

KGRVVVFSNRYFVKKTLSSRNGTEHDEKNLKNFFGTKLKFETDVYKDFTAEKMRQKLEEE

TMKRLTHFDVFIVFFLSHGIEGEIYGTDGRTLLTRQIVDLLDGERCPQLIGKPKLVFIQA

CRIVRSVPDVSVQTDGGALVQEMDNWTLDDRDPEVKANFDKLTREGVDPDSPPEPYHPQA

DMAIFFATVEGDPSYRQVTEGSFFVRYLLDVFTKKAHKEDVQSMVTEAPEILVRRPGRHD

HRVDLWSLGVIAYEFIVGSLPKDIIALNRYHASRGNTRTLKIMAATEEQYAEKRRLCPSG

NQQRRHDIAVDSCIILHQKH

>GWHPACBE010591_P20P10Eisenia_andrei

MDNEDVSIQSLEGPWISHVYIGFSPADRKTVFDLRLWLEKHGVICKLRETNDSIQESIRE

GVATSQKCLLFLTPDYMKDHWYKMETNAVIEKSQRFSRDVVIILKNQNQEVIPEILQEFK

AYSFKPEKLNDEKFVATILDAVIREPNVKLAVLPKQNIGYGSAYSYFYGFLRLVLPDFRK

RLVEETLKNDPEVLQKTVQKMLIIMPESCFCASELKKENHIEFTGKFVVRVAHRAGNQKR

DYKSSLYKLIDHERQQDYYFVGELATPLLTMYETYYFGLCGMTKDQLYAERDAFYSTLKA

ILSHPDNAECNDQFKLLYWSDPETDKPNQPDIYQFLLPVLRKEIMLERDNERNPSKVGKT

TNAHPSKPKPQLLKTFINVQVDGFNPARIYDEIGSYPMNRRPRGLCLIVNIANFEKGHRH

LSQRKGAERDGVRLQKLFKDLHFEVQVHSDLNAQTLELLFLTMHIADHTNYDAFVCCILT

HGKLGVIYTSDGKEVEILRLVDFFTDKNCPSLKGKPKMFFIQACQRGDDAESGSAGSSIS

QSSHSTSSDSAASRDTVEDTREGISSIQAEISNIIVTDVEPALAVDATSAPTVAVIDESL

PTTFKHKAVLVPGSPDFLMSYSTLPGSISYRDTQTGSLYIQALDDCLRKGEEIDRALKMV

TSSVMKQLRQRE

>GWHPACBE010589_P20P10Eisenia_andrei

MDNEDVSIQSLEGPWSSHVYIGFSPANRKTVFDLRLWLEKHGVICKLRETNDSIQESIRE

GVATSQKCLIFLTPDYEKDNWYKMETTSVIEKSQRISRDVVIILKTQDLEVIPEILREFP

SYPFKPEMLNDEKSFATMLDTVIREPNVKLTVLPKQNIGYGSAYGYFYGFLRLVLPDFRK

RLVEETLKNDPKVLQRTVQKMLIIMPESCFCASELKKENHIEFTGKFVVRVAHRAGNQKR

DYKSSLYKLIDHERQQDYYFVGELATPLLTMYETYYFGLCGMTKDQLYAERDAFYSTLKA

ILSHPDNAECNDQFKLLYWSDPETEKPNQPDIYQFLLPVLRKEIMLERDNERNLSKVGTT

TNAHPSKPKPQSLKTFITVQVDGFNPARIYDEIGSYPMNRRPRGLCLIVNIANFEKGHRH

LSQRKGAERDGVRLQKLFKDLHFEVKVHNDVNANTLESLFLTMHILDHTNYDAFVCCILT

HGKLGVIYTSDGKEIEILRLVDFFTDKNCPSLKGKPKMFFIQACQRGDDAESGSAGSSIS

QSSQSTSSDSAATCVTVEGTREGISSIQAEISNIAVTDVEPALAVDATSSPTVVVIDESL

PTTFKHKAVLVPGSPDFLMSYSTLPGSISYRDTQTGSLYIQALDDCLRKGEEIDRALKMV

TSSVMKQLRQREGHDRPQERFQLPFHLTSGMDRLIYL

>GWHPACBE010581_P20P10Eisenia_andrei

MIYKEVCCRDLEGPFSHHLYIGFSPANKKAVLDLCTKLQNNGVICYLRSTNDGVKESIRD

GVKTSQRCLLFLTPDYINDAWYEGEMLAVVKKSECFSRDMVIILKSADLEVIPEGLREFS

THLFKPEMLNDDTFMVPFMDMITREPKVDLSAGSRRIAGYESARNYFYGYLRLLLPDFKA

RLVDTLKNDPDDPAFVSKAVHKMLILMPESCFCPPELTLENKIEFTGKFIVRVAHRAGNQ

KRDYKTSLFKVIDNKEHQVYYFAGEFATPLLTMYDAQECGFGGMTKDQLYEERDGFYFVL

KAILSHPDNADCNDQFKLLYWSDPATGKPNQPDIYEFLLPVLRKEISLQNENEKNPSKIA

AIDNIHLLPSNRRPLSSFGTVQVHGSNPARAFEMLEYYPMDHFPRGLCLIINIINFDEDH

SDLEFRQGAEKDQELVEKLFNDLQFEVKVHRDLDESDLLTLFFETNATDHTNYDAFVCCI

MTHGKLGVIYTRDGKPVEILKLVDYFTDKNCPSLKGKPKMFFIQACQGADSEPETDAAGN

SLPLEYSIGDHLALPSSQRTDLESLSVAQGIDEDAVAEPAASTLDETFPLPLPRDAVLVP

GSPDFLMSYSTLPGSVAYRDIHTGSFYIQALDKCLRKGEEIDRALKMVSSNVREKLRQRG

ELDRTEERFQLPFHLTSGMDRLIYL

>GWHPACBE010586_P20P10Eisenia_andrei

MANEDVGSQSLEGPWSSHLYIGFSPANRKTVLDLRVWLEKHGVSCKPRATKDSIQESSTV

SIQESIRDGVTTCQKCLLLLTSDYVKDDWYKIETNAVIEKSQRFSRDVVIILKTQDLEVI

PNILEGFKTYPFKPEMLKDEKSFAKILDVIIQEPNVKLAVLPKQNIGYGAAYGYFYGFLR

LVLPDFKKRLLEETLQNDPDLVPKTIKKMLIIMPESCFCPAELKSEKIEFTGKFIVRFAH

RAGNQKRDYKSSLYKLIDVERDQDYYFVGELATPLLTMYETQTDGLCGMTKDQLYAERDA

FYSTLKAILSHPENTGCNDQFKLLYWSDPATKKANQLDLYDFLLPVLRKEIMLERENERN

PSKSGTTANVHPLLPNPRLLDTFITVDADGFNPARIYDEIGRYPMDRRPRGLCLIVNIAN

FDKGHRHLKARTGAERDGECLKKLFGDLHFEVQVHNDVNAQTLESLFLTMHIADHTNYNA

FVCCILTHGKLGVIFTSDGKEVEILKLVDFFTDNNCSSLKGKPKMFFIQACQRGDNEMET

GAAGNSMPHSSSTGDHSAQNTASDSSANVADRHTVEYTQEGISNIAVTEIEPILAADATC

APSVSTVDESLPRTFKHKSVLVPGSPDFLMSYSTLPGSISYRDTTTGSLYIQALNECLRK

GEEIDRALKMVTTSVREKLRQREGYGQGQQRFQLPFHLTSGMDQLIYL

>GWHPACBE010587_P20P10Eisenia_andrei

MANEDVSIRNLEGPYSSHLYIGFSPDNEKIILDLREKLRRHGVICYLRAMNDGIRESIRE

GVATSQKCLLYLTSEYVNDKWFQMETMAVVEKSERFSHDVVIILRGTNLVVFPDRLRPFK

TQSFKPLDDEQFFTKMLDTITKEPNVKLAVLPKQNIGYGSAYGYFYGFLRLVLPDFKKRL

VEETLKNDPDLVAKTVQKMLIIMPESCFCATDLKSENHIEFTGKFVVRVANRAGNQKRDY

KSSLYKLIDHERKQDYYFVGELATPLLTLYETHYFGLCGMTKDQLYTERDAFCFTLKAIL

NHTDNAWCNDQFKLLYWSDLATGKRDHLDLYKFLLPVLRTEISLERDTERNLSKVPQNRL

LLNTFENVPVDGSDPSRIYNEVDEYPMNRRPRGLCVIVNIANFEKGHPHLLPRKGAEKDA

ECLQKLFEYLHFTVQVHTDVNALTLQSLFLAMHIADHANYDAFVCCILTHGKLGVIYTSD

GKEVEILSLVDFFTDRNCPSLKGKPKMFFIQACQRGSAADMTDRDTVEDNRKGISNIEVG

NSNIAVTEVEQVLEVDATNDPTESHPNFRHKAVLVPGSPDILMSYSTLPESISYRDMQTG

SLYVQALDKCLRKGEEIDRALKMVSSNVREKLRQRGELDRTEELFQLPFHLTSGMDRLIY

L

>GWHPACBE010585_P20P10Eisenia_andrei

MANEDVGSQSLEGPWSSHLYIGLSPANSKTVVDLCLKLKEKRVIYKLRETNYSIQESIRE

GVATSQKCLLFLTPDYVKDDWYKIETNAVIEKSKRFSRDVVIILKTQDLKAIPEGLQDFA

QELFLPETLNDEKFLEKILNAIFREPRVKLAVLPKQNFGYGAAYGYFYGFLRLVLPAFRK

QLVEETLKNYPDLVSKTIHKMLIIMPESCNCAPELNSNAQIEFTGKYVVRVFHRAGNQNR

DYKSSLFKLIDHEKQQDYYFVGELATPLLTMYETYSFGLCGMTKDQLYAERDVFYFTLKA

ILSHPDNAECNDQFKLLYWSDPEAEKPNKLDLYKFLLPVLRKEIISERENERNPSKMGTT

TNVHSLKRQPPLLNTSISVPVDGSNPARIYDEIGRYPMDRRPRGLCLVVNIANFDEEHRH

LKQRKGAERDGECLQKLFGDLHFTVQVHNDLNALDLESLFLETHKVNHTNYDAFVCCILT

HGKLGVVYTSDGKPVKILSLVKFFTDKNCPSLKRKPKMFFIQACQRGDSAETCDAEGISS

IQAGISNNVVTAVEQAMELEATSIPTIAVLDDSLPSTLRHKAVLVPGSPDFLMSYSTLPG

SISYRGEQFGSLYIRALDKCLRKNMEIDRALKLVSSSVKNQLKNQDEYDKAEERFQLPFH

LTTGMDQFIYL

>GWHPACBE010592_P20P10Eisenia_andrei

MATEGVRIRSLEGPWSSHLYIGFSPAYWQTVRDLRSWLEKHGLICKPRSTNISIKESIRE

GVATSQKCLLFLTPDYEKDYWYTTETTSVLEKSERFSSDVVIILKNQDLEVIPNILRGFK

TYSFKPEMLNNEESFAKMLDVIIKEPDVKLAVLPKQNIGYGSAFGYFYGFLRLVLPNFKK

RLLEETLKNDPDLVPKTVHKMLIIMPESCFCPPELKSVNRIEFTGKFVVRVAHRAGNQNR

DYKSSLFKLIDHETQQEYYFVGELATPLLTMYESYYFNLCGMTRDQLYTERDAFYSTLKA

ILSHPDNVECNDQFKLLYWSDPATDKKNQLDLHKFLLPVLRKEIFEQRDPSKVETDNPVP

SSNRPLLDSFITVPVGGSNPARIYDEFGRYPLDRRPRGMCLIVNIANFDKGHRHLQHRKG

ADKDGERLKKLFGDLYFTVQLHNDVNAETLESLFLTMHILDHTNYDAFVCCILTHGKLGV

IYTSDGKEVEILGLVDFFTDRMCPSLKGKPKMFFIQACQRGDSETESGSTGSSMRHDGTT

GDHVAPQFSQNTSSDSAASGDTVEDTREGISSIQAGISNIAVPDVEKALDVDATSSPTLS

VLDESSLTTFRHKAVLVPGSPDFLMSYSTLPGSISYRDTQTGSLYIQALDECLRKGEEID

RALKMVTTSVMGRLRQREGHDPAQELFQLPFHLTSGMDRFIYL

>GWHPACBE008493_P20P10Eisenia_andrei

MTSTAGGSKGDSSRNQNVELKYRMDHPKRGRFIIFNNTTFNVRGKKLDARSGTAVDSRNL

KDVFEKVLHFEVEVHNDLTCGQMEDIMRSEATSDHSSCDCFGVAVLTYEETEGVLYGSDS

TITIERLLKPIKTCKSLYGKPKLCFIQACKPVEEKSKHHPDGSLQSDGLHERKKIRLPLE

ADFLYVYSTVPGIYSWSECDRGSWFIEALTDKLKTLAYEDIDLIRILTRVINEVANQLIS

SKKVKEKVPSITSMLTKDIYFTRKPDSSS

>GWHPACBE010155_P20P10Eisenia_andrei

MASREELLTTEDRYPVQSGGFRGKFVVISNETFLSQKLKKRDGTENDVKRLKETFSGLGF

DFTNNIYRDKKALDMWKIMIEVGKAGLEKYDCLAIAILTHGESDGRIHGAGAGDDLIQLD

SLFEPLKHGRKLKGKPILIFIQACRGENLDPGTLTQTDTPTRDNGSDDTPRFMKIPIEAD

FLYHFSTSPGYYSFRNANGSIFIQALCKVMAKYTEKKEEIDLVKALTRVNREVAEGYSSC

VPSDRSFNDMKQMPCILSMLRKDVYLHPK

>GWHPACBE020220_P20P10Eisenia_andrei

MEESQRRVLNDSLTFLKANLLTVELLDHLLEKGIISYELKQKINYEKTDGLKVAELICML

TYRGHQAFDVFLEACRATEQGHVADHLEAKLTAIKENQYSNTRTFVVEWLQTNIELKSSP

EAAGLSADELVCLANTAISTASERDNKDCRDLIVTELKKILCEVFSQPETAMNGNLLPLQ

LKQTTVKETTLETTGAKSPSETTCLPKSMRDWTVAHVCQWLERNGLDSDVIQQFQHERIT

GKALQYMNESDFKELSAVRLGDRITILEERNQYVTNPVSSYGEKQIVDVSLSSVEATSQE

KICIASSKAIDPAIGQMSCDKNSGAVPMELNFDPPPEKLEIAEKVLAIEEPVAEINTEDV

SENLRKSKEAKEVELSSSDSEEDFSASETKEDIKSKQSIQVSESKDVISTKPKEEARTSK

PSEAIDVSETEEDILTKFQKPSEVLRTFGNPPNSSFKYKKGAKLSCIETRSGGNLLQPIH

RFVMLEIDTIAGKMEFANEAVSFACACLNDRANGTIHFGIVPEASSGCTVHGQITGLVLK

KKIEKYADFLKKAFKNCFLEDQQDIIQECVRPAVFIEVLSETTEITEFIIEFDVIPEYNV

CKDKAFLARLCKGEKDAWEDYVIFQIVNGNVSRLTVSQQNNYMEYKEYVAAERKQQEDAA

RKKFEPRDELSKKLLHLLCDGEETFRGDQYRILVINKPEETMNSDTMKETFSFLADLEWK

VIFDFDAEAKICKFLQDEETAMKIIMSDEFSRHSESNVYKKDHLKQTQEDIRNSVQPSWI

FVNGYSPLNDESLSPRNWKMRKSDGFKEAVRFIGHQFPEGRALVVFLLLSKDTDVVLEAA

EEFITWFPDQWLCVAEEESIGKPWIEELKRRNCIDRDSENDRVVFGMPWSNVQDAVVSIT

SSRRKNCCEIPTSAGSFVVLKNKTINELSDLEILGSFECDNSDIRNDANELRLREAAQKQ

TFFQGNGVSWWNFWFPGQVCERDVLRRLRKAVKSSLDGIDDDVGKVILSHQPGAGGTTTA

KHLLWELRKSYRCAIVQTINDGQTANQILNLYQYEDSNPKPVLLLLDNPDEEKANLLVAD

VSENAKNLTRKGEDQAKCVCVFLICSRHSKVGSQYDPKQLVLKHVLTPKEQNWFKDMHKK

LLAEHKREPSIFTDPKLLISFNMLKENFNMDFIQRMVKSFLDNITDEKERRLLKYISLIN

AFDLECRALPTAAFDAIMIENPWAKKSHRWGKKAYSNSCTDIFYAASKARDDLLEISKDI

LKRRGHLPNGKPETRFAPLIQTIVNDETIEKASAILEEGLKLTSDPFIAQQLARLFITVQ

NWQQAIKYAKMATEQKPDNSYLWDTFGRIYEGQLLEQYNKLCVGSRSTSTEEFVPIIKIA

LNGMEMFTKVQHTSQHEKRLSVNQAGFFGELQMIVCLLDCLTYLFQDKAKLRHFILEREY

IPEECSSLACYEDTNYIAKLKELNVQVQIAVNRLEDELFQLREEAINEFRRNHVRYQSDS

LVRLKENLNSYFGEDDDNVPESLEEREKCDFRRRRIFRLAGFNLRSIFDLRSKSDDGHNR

LQRIQEMAKLNVLSEYVLVTDYQTLISVTLALRSMDPEYLCNFNEYDSLVKWSKKLYELR

DSSIVPRLEPYLFYVLFNWPRENSNVSPKVIESAIKNWKDAYYLKYPRQKDEGKPYRKRD

TTMFFLAKGQDMESIYTFFEEIGSRRNDTKVRSAFWRHPVTVKKLQRFSGVLSYGGNEVK

VQLQSGQGNKAHISIPPSFPIQDRQMWNKEVYFVIGLSWIGPKAFDITLDEPIITTELCS

EAVSSKTVFHRSGRSVVEFTEEEFRRKLKVINEKLNMIQNQRDKKEVNVGFDTGEDQLMH

EEDKLNKEKRLLIQMRQSIMESRIPEY

>KAK2191467_1_hypothetical_protein_NP493_53g19021_Ridgeia_piscesaeCARD

MFSKTHRGILEKHRPELVDEVDTDSLLPELRKRKLLTSRQEDKIKLNGQRVRAEQVEIIL

DLIPRRGKHAFDDLCEALIASDQERIVQDYLKPDILAKTEEEPPRKKARKQDATQNDEEE

DMDEGIDINDGPISVRVHQSTPEFYEKKFREAYPMCVPCRGLACIINVKHVVGTDDRAGT

DIDCENLQHLWVQLGFQVVVYNDEDDLTVQGIRDKVIQFAAEPGHQRAQCCVLCVLSHGE

NGAIYGTDGKLLPVAELKYHLDAKRCRALKDKPKLIFLEACRGDIADTDVADCPSHADFL

VGFPSQPGYKAWRNCKKGSWYISAIVQVFMELAATKNIDELMTKVNNIVSRKLSRSHKYE

DCNKVQMPEVRHSLRLPLYFFPGITDEDGC

>KAK2181645_1_hypothetical_protein_NP493_386g02006_Ridgeia_piscesaeOK

MEIDVDEDVYRMNHHKIGVCVIINNRYFDKTTGMSDRSGTDVDAANLYQLFMGLGFDVRL

EQNKNTTDMLRILSRVAQEDHTNNDCFVCAILSHGDSGIVYGTDGIIQLETLVDPFKGRR

CASLLGKPRVFIIQACRGNKLDHGVEVADAIGEEPEQKIYRIPEEADFLYAYSTAPGYFS

WRNSNKGSWFIQALCKVFKTDVEKHDILTLLTKVNRLVAYDFQSNASREDMNRKKQIPSI

VSMLTKQLYLQPKK

>KAK2186248_1_hypothetical_protein_NP493_207g00020_Ridgeia_piscesaeKO

MDHPKRGVFIIINNKKFHSSVDQSERSGTDVDADNLCQVFTGLGFDVRQENNNTTAEMLA

ILVKASKEDHSDNDCFMCAILSHGTDGNRVYGTDGEVDIRCLVSRFSEEKCPTLIGKPKV

FIIQACRGARMDPGVEVADALGDSAETVSRIPEEPDFLYAFSTVAGYYSWRNSTRGSWFI

EALCQVFRDNRTGKEIMALLTEVNRRVARNFESHTPDEDMNRKKEMPSIMSMLTKQLYLL

PK

>KAK2186250_1_hypothetical_protein_NP493_207g00025_Ridgeia_piscesaeOK

MNHAKRGVFVIINNKTFHPSTTLNERSGTDVDAAHLYEVFMALGFDVRLESNKTTAEMLM

ILVKASKEDHSDNDCFVCAILSHGTDGNRVYGTDGEVDIRYLVSPFSEEKCPTLIGKPKI

FIIQACRGARMDPGVEVADALGDSAETVSRIPEEPDFLYAFSTVAGYYSWRHGVNGSWFI

QAVCQVFRDIRAGQEIMALLTRVNRKVALDFESSVKKQELNGKKEMPSIMSMLTKQLYLL

PK

>KAK2184734_1_hypothetical_protein_NP493_255g04023_Ridgeia_piscesaeOK

MVTVNSTVAQCLTRHCSDILINCTVVEFQDSIPAEVRVAAEQAHVQLIGCERPRRGALPK

KDWLTHYWQQHYPHLVNGQNFSHIIGYFPETAKTALSIRDSIRRRGADCTCVLLVVDSHR

DELTEESAKLINQADAVYSLGPLLFREFDRHVSKPHRMMFPVPHMGDPITTLNLHTPGTM

QPKDIVTFYPENGIPRPDEEYQHKLLVLVVKAISNVAKSYFCQCHANNIEWHIYGVPEDH

RVREQLKAVIQEHSVFPELTWRLHKQPTKTIKVLQLAVLCVSPWQRSYNPNALYPLLCGV

PTLLCASSGVAEILFSDISRDDDGSALLTARAVVNCSGDSGDIGWRTKIQDIVNNPLDAS

TDFVEVGRRIRGSVALQESMKTFCVATFGGQVVQPPSEMTALGPVSTGALDSPMSASSQQ

PGQPGQVGRPLLGRRIPPADPPGSVPVSPGPLVSQDSDGPSPGPGPQVPMGRGRIPVKPW

TDVNRYNRLIVDSENIYKMTASPRGIALIVNINKIQGTKEERTGSEVDEHNLCNLFTQLN

FSCITKRDLSRQEILDELNKLASSQELWGHHCLVVCLMSHGKEGVITASDNRDVAVTEVL

NIFSRCEALKGKPKLMFIQACRGKHTEGEVKADGPTDNVAATKPPVMSDMLVVYPSSLGY

TSYRNTKQGSWLITNLVEVFSEHACDEHISELLTRVMRKVSHMETSGKDVATGHRAQPEA

LLALRKLLYFFPGM

>KAK2189268_1_hypothetical_protein_NP493_111g01008_Ridgeia_piscesaeOK

MAGHMERKEGETLAESRCPESGGEKEARKNKIMIGGLLVSAQTQAASLQLDSYSNRHPRL

GLCVIVNNKTFNKQLTGQSNRDGTDVDAKALENVFMHLGFDVRRYDDLNRTDMSISLRQV

AEEDHSDADCFVCVILSHGEEGVVYGTDGKVPLEKLLGYFKGANCPTLAGKPKLFFIQAC

RGRRYDFGTVIGEDQVDALPLANTRKIPTEADFLIAYSVVPGHFSWRNNVDGSWFVQALC

QVFEQHGHSAELLQMMTLVNKIVAYDFESCTDDDFTSHMKQVPCVVSMLTRQFFFKPKY

>KAK2186249_1_hypothetical_protein_NP493_207g00007_Ridgeia_piscesaeRH2_P20_P10

MLEPPTQPSSKPMPEPPTQPSSKPMPEPPTQPSSKPMPEPPTQPSSKPMPEPPTQPSSKP

MPEPPTQPSSKPMPEPPTQPSSKPMPEPPTQPSSKPMLEPPTQPSSKPMLEPPTQPSSKP

MLEPPTQPSSKPMLEPPTQPSSKPMLEPPTQPSSKPMPEPPTQPSSKPMPEPPTQPSSKP

MPQPPTQPPSKTTSEPPTQPPSKPTSEPPTQPPSKPTSEPPTQTPSKPTSEPPTQTPSKP

TPEPPTQTPSKATSEPPTQTPSKPTSKPPTQPPSKPRPQLPPKPRVHLGNTNALSQPSSK

SPSQPSLQPPSKPPTQSPSQSPSKPPTQSPSQPSPQPPSKPPTQSLSQSPSKPPTQSPSQ

PSPQPPSKPPTQSPSQPPSKPPTQSPSQPPPRSVPQSTSQPAPQPVLQSPSQSPPRSVPQ

SPSQPAPQPVLQSPSQSPPRSVPQSPSQPAPQPVPQSPSQPAPLPCAQIGHTKVMGQADT

RSGQVVSEERDVTEAVIGADTDFDDGVYTMEHAKRGVFLIINNRNFDALKERYGTDWDAA

NLYEVFTGLGFDVRMENNKKKREMREILCEASREDHTNHDCFMCAILSHGAGDCVYGTDG

AIPLDTLVSPFKSEECPTLIGKPKVFIIQACRGQKVDDGVEVADGEQEPKPKEKIYRIPQ

EQDFLYSYATVSGYVSWRDEVWGSWFIESLCRVFEDDSSRLDIMALLTKVNRLVAYEYES

STDTEELDQK

>KAK2186252_1_hypothetical_protein_NP493_207g00027_Ridgeia_piscesaeOK

MDYEDKRGVFIIINNSKFIQNTHLTDLPGYVIDAARLYELFSVLDFDVDMKQNQTTEEVK

TTPSNVARKNHTKYTCFMCAILSHGSNDVVYGTDGPIPIDTLVSPFKDEQCPTLRGKPKV

FIIQACRGVEMDHGVEVADAIAEPEENIYCIPQEADLLYAYATVNGYVSWVSEKDGSWFI

QSLCRVFGNSQLRSKLDVIAMLTKVSRLVAYEYETNNNNYRRSEPQEASTVSRFDANKAN

VSAIEITRLRRLAPYADLMSGVNCLRPSFKMFASREIQKLTGLIQSSGTNKHLTCHLQQT

SHDSDSLNP

>KAK2190501_1_hypothetical_protein_NP493_78g07001_Ridgeia_piscesaeDEDDED

MPSRENMERDAIDEKVYYRRTLKQIDDGLVSDNISAIKFLCRDFVPAAKLEPVSRGIHLF

EVLEQKCLLGNNNSALLIKVLQTIGRANLFKVCDELTSKEVDTMKFMCEGHMHRKLLNVN

TCEDLFVALEESGHLSADNTRVILDLLSDVQRQDLHSLFVPSQHLRHGTQQVRPILGIGA

ATSGAPTRIIRSYQAEDELTSHFQHMDIVKGNNAMPPPNTPLPACTGQPLSPAEDTSSAQ

VMPSAGQASHHPPPAAAVSPQVFPADTSLGSYDMSADPREKLQGIFEKLRFQVSVQIDQT

ASDIKELMIYFSTQVDHAPFNCFICCILTHGKLGKVYGTDGATVDIMELTSSFKGAKCKS

LVGKPKLFFIQACQGLEKQTGVAIESDAPESISQSLQETLPNESDFLVGFATVPGFVSYR

STTDGTWYISKLVEMIDTFHTSEHLLDILVKVNDEVSKVAKETGDRVYKQAPAPLVTLRK

KVFFR

>KAK2175515_1_hypothetical_protein_NP493_727g01061_Ridgeia_piscesaeOK

MYSIRCGTAVIFNNRQFTDGPRIGTEEDENNLKRTLEGLGLKVTVHHDVDTTTLLRELQK

VAKKDHSKKDCFVCVILSHGGHSELTEEAEQRVCDVIVCRNGIIAVRNLVNCLSDEEAPT

LLNKPRLFFIQTCRGSSLPATVESHGDGRPPKTVEPATWQDVEKSLAYKLPADLPVYPDF

LLAWSTVPGYFSFRRPSGSPFIHHLCDVLREYREYDLLTILTLVQSYVATKYSSFAPSTV

NDPKKTNIHNKKQMPWVASMLTRKVYFC

>KAK2175514_1_hypothetical_protein_NP493_727g01078_Ridgeia_piscesaeOK

MHGYESNKSLSCCVNVSSVPVAKEDHSQKDCFVCVILSHGGQNELTDEGERRVCDVIMCR

NDIIAVRELVDCLKDEKTLVDKPKLFFIQTCRGTSLPATVESHGDGRPPKTVEPARWKEE

ERNLAYKLPADLPVYPEFLLAWSTVPGYYAFRHADGSPFIEYLCEGLRQRTEHDLLTILT

WVQNQVAIKYTATLDLENGLRRCKQMPWVASMLTRNVHLQQ

>Pdum_Caspase_PladumEVm033781

MAGHNDDQQYDFTHAQRGVAITINMDFENGRCPPRPTAKLDEEHMQTVFERLQFEVRSYM

NLDKDEIIDILKEGVAPACYLSQLITKFAEEDHTNRDCFALAIGSHGCNEEEEDGNQTIM

TDTIEANDKSFSLREITEIFSEKNCSDLAGKPKLFFIQACRGDKMAENVTLDEPDFCLPP

VTKIHDIPLGRDFLLAYATIPGHLSMKRRGKGSWFIIALRIIILQNYLREDLLTMLGLVN

NYVVRNFYAKSAIQEYNNKRVIPWIASMLTKDVNFKQK

>Pdum_Caspase_PladumEVm031689

MEPKRRKLAASNESNEVGEEYDFSYSDRGVAIIINMEYESWREHHRPTAEDDEKHMDQVF

KKLGFKDVRCHKNLTKEKIMETLKEVAAEDHRDRACFALAIGSHGRSKVEQHENKKHIMD

FIDAYDQSFRLRDITKIFSEENCPSLAGKPKLFFTHACRDEVTGMGANAMLDNQPQTKVL

DIPMGRDFLLAYATLPGHRAGSRQGSTGLNGSWFFSGLHAIVRRHDDKDKDLLTMLGLVN

QYVVQNFQFRLNQGDKNVRNDERSEEKATTSRNLNLKNRRFIPWIASMLTKDVKFK

>Pdum_Caspase_PladumEVm010476

MHPSYKESGLAREEDLTNLVWRQLRGGATDWSESKGKIRITSESIARDVMAKLKIYTPLD

RMLSTRIMEAYMAKCDINKEWSTNFVGLLTFLIREGADPFCPLSPGGKPQKKTAFKVFLE

RFSPHLSDSLMPSVSTFLEEAFKKWETSETKESILYDVIRDLPSDIVKGESSHLSTIKGL

IVRNCLEIIKTNPGLCDHEDREGKTPLHKLMENILIDELWKRMPELVAGTDTLWDLMSGL

ITKKNVNIKDKDKVTPHAWILWWQESVTDYQIFNTKKKQNLVRKIEDIKKQFEENGACEE

DALQGLKNFRSKRQSGIFDGFKKLFRKNQGSKDDGHDRTKDQKNTSITTDSKTIATISSQ

KRTNHSEQASPKDEERDKNVPLGQEDLKKTNFAISMVTPISARGTILEDTECDQHLPITV

RPASPQLVKEVLNNPEDCYKMSAEPRGRVIILNNEIFDQSETAEVKLEGRDGSYFDVENL

RRLFEALRFEIVVWSNISAEEIKKKIKEESQSEFHEMADAFVFIILSHGKKGQIFGTDGK

SVHIDAHIVGAFDGRKCQQLIGKPKLFFIQACQGDQVSEGTMKLDSSAPLRQPSFSPPTD

EGPVHEKADVLIFYSTVPGYASWRNTEVGSWFIRDVVICINNYAHKLHVGDIMTKVNKQV

SEREVRFQGYKQISEYRSTLTKFLYFLPST

>Pdum_Caspase_PladumEVm031996

MASYQMDEADYASSGAGGQKTKGNGHIPINLQPSNAKLIETVLSNHEDYYKMSAEPRGRV

IILNNENFEQSDDEEDNLPGRHGSYFDVENLRNLFDALHFETVVWSNVTSYEIRQKIKEE

RKSDFHKNADAFIFVIMSHGTDEHIHGTNGRKVHIKKEIIDFFDGNNCPQLIGKPKLFFI

QACQGEKATTGAVSDSATPLDPKKAANILHEKADIVLFHSTVPGYMSWRNPGLGSWFIRE

LVLCINQYSHEWHFGDIVTKINKKLSEKRMTSKGFKQISKYESTLRKQLFFLH

>PdumCaspase_PladumEVm027269

MEPKHKRLKVAASNDVGEAYNFTHSKRGVAVTINMDFENGRKAPRPTAEEDELIMATLFE

NLEFEDVRCYKNLNKDEIIKKLKKVAAEDHTNRDCFALQIGSHGNNKEEAEEYQEIIMDI

IEANDESFRLRDIMQIFSEANCPSLAGKPKLFFIQACRGVKMAENVTLDEPEFSPPPMEE

PDVSPMDEADLSPQIMDEPDLSPIPITKIHDIPLGRDFLLANATIPGHRSVRRPGKGGWF

IIALKEIILQNYLREDLLTMLGLVNNYVVRNFYVDTKNEKNKKYKNKRVIPWIASMLTKD

VKFTPKFRNDDDNAGPRMPKQHPAPKTTLSGSPTGDNH

>Pdum_Caspase_PladumEVm029732

METQGEAEDLAAGASKDMDEPDLVTAQKNTWPHLPGDRKGDGRIAITLKSSSPKLVRKVL

DNPEDYYKMSAKPRGRVIILNNEIFDKSERAEDNLPGRDGSYFDVQNVKEMFAALHFETI

VWSNITAAEIRMKIKEERQSEFHKKADAFVFVIMSHGSMDNIYGTNGKRVHIKNEIIDAF

DGKKCPQLIGKPKLFLIQACQGDQATLGAVRPDSSVPVKGPNEDASDMLHEKADILIFYS

TVPGYISWRNKQLGSWFIRELVTCIKDYAHEFHVGDILTKVNKNMSEKKMTSKGFKQISK

YESTLRKQLFLLH

>Pdum_Caspase_Comp214042_c1

MARPKASAGMSKLHEKCIIRNHTYLIEQLKGLGKWGLFDKMIENEVMRKKWIDEIKEIKH

ASKQIGKLLDELQRSGPHAWDGFLHCLDASDFQEVSWRLRETEKLLEENMDTNAHQLNEY

FEKNHPPENVQSKLRKSMAPSKGPVYGSASPNEEEQYVMESRPRGLMLIINNEDFSRCGK

ENREGSKKDCDGLKDMFQRFGFLVEVENNQTKEQMLAIIKIYAEMANGGSYDSVIVAILS

HGDESVFYGVDGKAIMESELTGPFNGLNSENLRGKPKLFILQVCRGDKTRQSVSHDGPNK

KPPSNFLVQEGEAREEPNIPTTVDDEEEEMSRALDRLKITQNRHPDADMVLAYSTLPGHL

SMREDNNGSWFVQSLIEVFTEHAHDTEMLVLLRMVNKRMSQKMLKSAAQIPQPRFTTLKA

WYLNPVQD

>Pdum_Caspase_Comp219121_c0

MDEGACSSEEYDFSHPERGKAIVINNIKFEERTQQPFRKGAASDSDNLADTFALLGFDVK

VYTNLTTTEMLTVLQQAAREDHSKRDCFVCAISSHGNEDLQKKSRWSSKTETSDVIYGTD

GVLHLQLITDMFKEKKCPTLIKKPKLFFIQACRGDMLAENVEVTDGLMEDEVDWGGHHIF

VQNIPIYRDYFLAYSTPPGFYSFRRGIGSWFVISLCDVLQTCHKDTDLQTMLTKVNHRVA

YEFSVNTPRDRRTHGRHIVPWTASMLTKDLFFRPKTDEGKRIEFV

>Pdum_Caspase_Comp223063_c0

MEKDDRKTLQKNTVLLMRDLVVDKMFLAVLQQRGTLNATMLEDVDCHVKKTDRNSEFLNL

LPRRGPKAFNDFIVALMESGQDHLADALSPELAAQWRKANPQGPTALAPREPPGGYYPNY

GMPLMPQMPVPVAHPGSFVPSSQGDKERDSFIPINLQPSNPVFVHQVLNNPKEYYKMSAK

PRGRAIILNNESFDQSAKAETRLTEREGSYFDVVNLMELFKALNFDTMEWPNFTADRIRK

EIEKERQSDFHNGADAFILIILSHGTRGHIYGTDGRKVNIEKEIIDAFDGNRCPQLSGKP

KLFFIQACQGDQITMGTIKSDSGDAHDAVLDEKQDKLEDDKDAVPTNDSFVHEKADILIF

YSTVPGYKSWRNTELGSWFMRDLVICINDYAHLLHVGDIMTKVNNQLSERQVKCRGFKQI

SKYESSFRKLLYFLPKS

>PdumCaspase_Comp217549_c0

MAGIAVLESELISVLRENGTVSTKDIVRRLKSFTKKEINRQLHQYQKDGIVDKVSDIPPA

WRLKGKGGAANVTDHMDNEISRSGMDRLCEGKMKCLPPPAHPLPERYVMKSQPRGLMLVI

NNEDFDGCGKETRHGTNKDRDGLKDMFKEFGFQVECKNNRTKEEMLKDIKLYAQMANDNN

YDSVVIAILSHGDESIFYGVDGEAMMESELTSPFNGFNSEKLIGKPKLFILQVCRGDKKR

QSVSHDGPPQPPNNFLIKEGAAEDQPKISPEEDDNLLKELARLKITKNRHPDADMVLAYS

TLPGHLSMRDSENGSWFIQSIIEVFSKHAHNTEMLALLRMVNNLMSTKMHKWNAQIPQPR

FTTLKEWYLNPVQD

>PdumCaspase_Comp216671_c1

MDYEEVFQFQYNMNYQQRGMAVIINNRKFDSKTGMNERSGTDVDAAQLYQKFKLLDFNVH

MYNNQSCGSMLKIVTEAARQNHSDSDCFLLAILSHGDEGVVYGTDGIIQLDMLVNPFKGD

KCPSLVGKPKVFIIQACRGNKLDHGVEKTDADSEMEEMPKQRIPAEADIIMAFSTVPGYF

SWRNSTKGSWFVQALCKVLERYGREKDILWILTRTSYLVAYEFESNASKEFMNKKKQIPC

VMSMLTKELYFHPKRS

>PdumCaspase_Comp218890_c0

MEEKLNGLQSQIKDMIFGSSVDPEKSAPATDGGYRKVPRPTNSQQQRSPYLEFTTSILQG

IDEDLKKRDVEDLKFLCTELIAEARLEQCSRGLDVFDELLYQGYITRQDGSLLIELLVYV

GKMKKVRQLGIDPHRFRQSLMYGGKLPAYRVLLWKFCEEVGKDEFSTLKYFAKEYGVSKQ

ELESIQCMTALFKTMEDRGVLAPLDTNVIMDMAMCLKRQDLMDLVAQYQVNPGLPPMYSR

AKDPASSGMAPIPGTEKVHPLRNRTDASSVSPAATGRYGTTEPPPYSPHPMMEEESLSGA

GAVHNFYREKQMAMPNFPLPLPGQRPELDLPRYKMNSVPRGYAVIINNRDFYHVPSDPAS

RKFSDREGTEVDVRNLEYIWKKLKFDVKTYTNLSDHEIHRLMTEYGMCPENSQQKKDHSN

KDCFVCCILSHGTIGGVYGVNGKLVNIKDITNAFKPAMCPELTDKPKLFFIQACLGSDTQ

KGSSMDKIQQDSGGGISTVETIPNEADFLLGYSTTPGFVSYRSKEQGSWYITKLVQMLDE

YAHRYDLMSILVRVNEEVGKAVAQVENSEFKQSPAPLVTLRKQVVFD

>PdumCaspase_Comp219364_c1

MAEKSDQAEFKADQVDAFGFSYVSIKPLRLSGKSYKANPISLVQEEYNFTYPNRGTAVII

NNRNFDKERTGQSDRDGTDVDASALESVFIKMGFDVVRHDNLNRMDMMLTFRKLAATDHT

DSDCFVCAVLSHGEEGVLYATDGVVYIEKITRLFKPDECPTLAEKPKIFLFQACRGRFYD

YGHNVDDDQPDAALDPVSVRKIPVEADLLMAYSVCPGYYSWRNNVDGSWFVQAITEVFKE

HSDSMDIMDMMTMVNRKVAYDFASCTDREFTSDMKQMPCIVSTLTRKVYFRPKEEQSP

>KAK2162969_1_hypothetical_protein_LSH36_88g00021_Paralvinella_palmiformisOK

MAEGAGDKMPLEERTPAEGQDKLDSRLNSLALSSSVDDKYRYHMNHKNRGHFIIINNKNF

DSSTNMGNRNGTDEDAANLYQRFKELGFDVKSYNNLKKIQMLKVMTDASRWDHSECDCFG

CAMLSHGDEGVVYGTDGQIALDRLIQPFKGPQSYTLRGKPKIFIIQACRGDQLDHGVEVQ

SDSEVPIEEKVYRIPVEADFVYAYSTVAGYYSWRNSTKGSWFIQAMCKMLKQHAYNMNFV

QILTRVNYEVAYEFESNASKESMNKKKQVPSIVSMLTKDLIFNRK

>KAK2158629_1_hypothetical_protein_LSH36_166g00039_Paralvinella_palmiformisCARD

MALYQMSPEHKGKLQGNRVYLIENIEPSERLLSYLLQERVLTEDMYEEIDVERLRKRKVG

KLLDILPRRGPVAFQHFINGLMKDDQECIAECLDPDLVAQWNSVHPKLKPGSHNPYPGQQ

PLPVPQSHQSFQPKSFFESEESGFFNPEVKSVDVSDKNIIKLIRDTKKNYTMSSKPRGRA

IIINNSYFVNNSPRDGAAFDSQNICTHLEQLYFDVKLYADLTAMEMKQIFRDEGSQRDHS

DEDCFFAFILSHGARGYVYVPVDELGQAIEDLHIKDSDKQMPHMIHEKSDVVLAYSTVPG

YRSWRRTDLGSWFVRAIIKVFAAEAYQEDLLSLLVKVRETVSEYHTKHENHSFVQLTRTD

ETTRKKIYLFPGYYEKNRASFV

>KAK2163519_1_hypothetical_protein_LSH36_78g00047_Paralvinella_palmiformisDEDDED

MANILMDAIPDPRSYRFILKDFSDKLSKRDLGMIKFICADHVPASELEKLDNGFSVWKSL

EDKGYLGQNDYTLLEEILILLCKLGSEISQKNLEAVKYIFTEELSRCASEEATDIWKFFL

LLEKQDLIAPGHVANLGNIIETLAMPRLQKLVDEYKESDPGKNANIDSELGHIYSPEMNW

QRLDLGVTEVQVSDDELKFQPLSHPVQTSDTSVQPSGSADGRCQQMPHSCTDDREFRHFY

MPPNSNYGPSSSVQKYPQHHIRPILPHTWNFGTTCPEKNDRELGRHLDTKGPICPPSEDE

HAAGERRDVREVKQMRNFSKELSMPELPAYKMTACPRVDKMKRILTKYANDEERMKKSDA

FVVVILSHGKFGAVYGADGYELKIVKILQYFRPDKCPALLNKPKLFFVQACQGKDKQKAF

NKIHADDVTEVEDDLMPHDADFFIGYATIPGYVSYRHEDDGSWYISTLIEHLNKYAKK

>KAI0222024_1_Caspase2_Lamellibrachia_satsumaCARDP20P10

MHKSNNKILRMDDRHREKLRSNRVALIKDLIVNEDLLAQMEQNDIFTPSMVEEIEVKPTK

QKRASELLNVLTRRGPDAFRVFVDALVCTGQTHLAKLLEPGITGQGKESRNEYRPLRQPS

QPMEEDDERGILPVKVHPVDEIEAAKKLNEGGIYTMSSNPRGRAIIISNEHFQGACHPTR

DGSIWDLVNLQELFRQLGFETFTYSDLTAKEIGDKLEEESSLQQPSANAFFLCILSHGAK

GLIFGADGRTISIEKITTMFDGNHCQALSGKPKVFIIQACQGEDSTQGVPQSDGSESVDA

EMESRLRELRIGDTPDSMPSLPNVHEKADMVLAYSTVTGYKSWRLPESGSWYIRELVKTF

AHFACKEHFMDLLVKVQDQVSKKHTYFDRHTSSGLYVQMPEMRTTLRKKLYLFPGYYG

>KAI0241014_1_Caspase3_Lamellibrachia_satsumaP20P10

MAEHHKSCAETSLKNADTQEDDASAEKLSVKMHIQMFNRLAKEQNDETPLMKPKIKNPVQ

QVKPPSTQVTTPPSEETCRNCSTHEGTTSYTTSHASPRSLTGNEQMPKTNQAETSPPPQN

KSKREANSLDTPVLSGIPIMEDTRVMEDTRVMEDTQVMEDTRVMEDTRVMEDTPDTIMGM

SGGRGATNGARGTDFKSDEDVYRMDHPKRGVFIIINNRRFLPSTRMNERGGTDVDAANLY

QLFMGLGFDVRMEQNKTTTEMLKIFDNVAKEDHTNNDCFMCAILSHGDLGVVYGTDGIIQ

LDTLVNPFKGRRCPTLAGKPKVFIIQACRGTKMDEGVEVADSLEGSSPKIYRIPEEADFL

YAYSTAPGYYSWRNGVDGSWFIQALYRVFHNDSQRLEIMALLTKVNRVVAYEFQSSTRNK

QMNRKKEIPSIVSMLTKELYLSPK

>KAI0221544_1_Caspase2_Lamellibrachia_satsumaCARDP20P10

MDEADKKKLSNNLVRLTEDLEVGLALLTFLQEKQVLTSENKAKIEILPEKGLDRASALLS

LLPKRGPKAFRIFIEALVKYGHEHLAELVDKDLTAHLLEKKSMVTQETTPMVVYSQETKE

LAVRPVDKNEAEKLLKQKDIYDMSKNPRGRAVIISNFDFGDATSNRRGTDSDKKKLEELF

MKLFFDTIICDDLKAEQIRQTLKEESRNQKSSANAFVLCILSHGERDHIYGTDWELISIE

EITTMFDGKSCPQLINKPKIFIVQACRGDKVTQGVSPDKTDSDVGDLADKIEKMNVCDTQ

LKVHEKADMISAFSTVLDYASWRNPQDGSWFINDLVDTFVELAWRDDILHLLSEVRRRVS

RRITNDESIALTHTGDQLSKNLYFFPGYAKPS

>KAI0214373_1_Caspase2_Lamellibrachia_satsumaCARDP20P10

MLCKRHKDTLEKRRPELVDEIEPDSLFPELRKRKLVTSRQEDKIKRNGQRVRAEQVEDIL

DLIPRRGQHAFDRFCDALEASDQERIVKDYLKPNVPVEKEQDPSVIPAKRARKHEPTQNE

EEDMDEGIDINDGPISVRITRSTEDFYQEKYKEAYPMCQPCRGLVCIINVKRVIGTDDRA

GTEIDCDNLQQLWTQLGFQVYVYNDEDGLSAQGIRDKVTQFAALPGHQHAQCCVLCVLSH

GENGAIYGTDGQLLPVVELKYRLDARRCPALKDKPKLIFLEACRGDLADMDVADCPTHAD

FLVGFPSQPGYKAWRNCKKGSWYISAIVQVFMELAATKNIDELMTKVNNVVSRKLSRSQR

YEDNNKVQMPEVRHSLRLPLYFFPAITDDDGC

>KAI0222026_1_Cell_death_protein_3_Lamellibrachia_satsumaP20P10

MSNNPRGRAVIINNKYFYDEEHHRHGTDHDKEKLQELFRKLYFDTITHDNLTAEQIRETL

KEESRKQESSANAFVLCILSHGENDHILGTDEEMISIDDITAMFDGKSCPTLINKPKIFI

FQACRGKAKTSGVTLEQTDLDETDFEDKMKQMNIRDSTDAEPIPKRHEKADMIKAFATVP

DYVSWRHKLEGSWFINDLVDTFAECACNDDILSLLGEVTKRVATRKTSTGEYIGLTQMHV

QLLKKLYFFPGYSKPSQNHTHKRSDSIQATD

>KAI0230384_1_Caspase3_Lamellibrachia_satsumaP20P10

MGDTPDARRVMGGGGAANAVESMDTECDEDVYRMDCSKRGVFVIINNRYFQKNTGMGERN

GTDVDAANLYQLFMGLGFDVRLEQNKTATDMLRILDRVAKEDHTNNDCFMCAILSHGDSG

LVYGTDGIVQLDNLVNPFKGRQCPTLAGKPKVFIIQACRGNQLDHGTEVADATGETEQKI

YRIPEEADFLYAYSTAPGYFSWRNSNKGSWFIQALCKVFQKDCGERDILTLLTKVNRVVA

YEFQSNASRPDMNRKKQIPSIVSMLTKQLYLRPK

>KAI0232817_1_Caspase2_Lamellibrachia_satsumaP20P10

MTASPRGFALIININKVTGGEERMGSEVDERNLCNLFSQLDFKCITKKDLTRQEILDELK

DLSKSEELRHHDCLVVCLMSHGKEGVIFASDKKHVAVTEVCEIFSRCKPLYGKPKLMFIQ

ACRGEHTDEGETRADGPTDNMAVTKPPTMSDMLVVYPSSLGYKSYRNTVNGSWLISKLVP

VFLEHACHEHIAELLTRVNREVSQMETSGRDVPPGNVAQPEAHWALCKLLYFFPGK

>KAI0241012_1_Caspase3_Lamellibrachia_satsumaCARDP20P10

MRHTSTCERTYVLEIGNRQSGEESKQRICEKDARCCTKLDYFRVPRGVNRPRNMTMDPVH

KDTLQKKHGLLVERLQPDLILDHLQLDGIFEEQDCDNVRSSRTEQEQSEKLLDKLLLCGP

RAYPCFLKALKAEHAKDLVKELDKQEAESRMAGHNLPGAETSLENADSQEGDVSTEELSV

KMHIQMFNKLAKEQNGEKPSVKPKVKNSAQQVTTTQATAATSSGEKCCNGSSYGGYRSQP

RPKPPLQLQVPEVPVQMPTSDNTSKRGVDTPDTPVMEGAPVNSYNVMEDTPVVEDTQYGN

TVEESDEDVYRMDHPKRGVFVIINNSHFKGLDERSGTDADAANLCQLFTRLGFDVRQKDN

MTVAEMVKILNKVAREDHTNNDCFMCAILSHGGAGFVYGTDGIIQIGTLVNPFRDEQCPT

LCGKPKVFIIQACRGDNVDDGVEVADSIGDSSQKIYPMPEEPDFLFAYSTVPGYYSWRRG

DTGSCFIQALCPVFYDDSNRLELMALLTKVNRMVACEFESHVPDNESMHRKKQIPSIVSM

LTKQLYLSPK

>KAI0227833_1_Caspase3_Lamellibrachia_satsumaP20P10

MDTGCDEEAYRMNCPKRGVFVIINNRYFKKETCMDERNGTDVDASKLYQLFMGLGFDVRL

EENKTCTDMLRILDGVAREDHTNNDCFMCAILSHGDFGVVYGTDGIIKLDTLVNPFKGRQ

CPTLAGKAKVFIIQACRGDLLDHGIDVVDGVKEREQEIQRIPEEADFLYAYSTAPGYFSW

RNSSNGSWFIQALYKVFQRDRGERDILTLLTKVNCMVACEFESHVPTDESMHLKKQIPSI

VSMLTKQLYLPPK

>KAI0211187_1_Caspase3_Lamellibrachia_satsumaP20P10

MDCSKRGVFVIINNRYFQKNTGMGERNGTDVDAANLYQLFMGLGFDVRLEQNKTATDMLK

ILDGVAKEDHTNNDCFMCAILSHGDSGLVYGTDAVIQLDNLVNPFKGRQCPTLAGKPKVF

IIQACRGNQLDHGIEVADAIGEVEQKIYRIPEEADFLYAYSTAPGYFSWRNSNKGSWFIQ

ALYKVFQKECGERDILTMLTKVNRMVAYEFQSNASRPDMNRKKQIPSIVSMLTKQLYLRP

K

>KAI0223707_1_Caspase3_Lamellibrachia_satsumaP20P10

MGDTPDARVATGGGGVANAVESMDTGCDEEAYRMNRPKRGVFLIINNRDFDKKTLQKRRD

GTDEDAKNLHQLFKGRGFDVRLEHNKTCTEMLRILDGVAKEDHTNNDCFMCAILSHGDFG

VVYGTDDIIQLDILVNPFKGRQCATLTGKPKVFIIQACRGHQYDDGIEVADAIGETEQKI

YRIPEEADFLYAYSTAPGYFSWRNSGNGSWFIQALCKVFQKDCGERDILTLLTKVNRMVA

YEFQSNASRQDMNRKKQIPSIVSMLTKQLYLRRK

>KAI0231133_1_Caspase3_Lamellibrachia_satsumaP20P10

MGDTPDARVATGDTPDARVATGDTPDARVVAGGGGAANAVESMDTGCDEEAYRMNCPKRG

VFVIINNRYFKKETGMGERNGTDVDASKLYQLFMGLGFDVRLEENKTCTDMLRILDGVAR

EDHTNNDCFMCAILSHGDLGVVYGTDGIIQLDILVNPFKGRQCPTLAGKAKVFILQACRG

DLLDHGIDVVDGVKEREQEIHRIPEEADFLYAYSTAPGYFSWRNSNNGSWFIQALYKVFQ

KDGGERDILTLLTKVNCMVACEFESHVPTDESMHLKKQIPSIVSMLTKQLYLPPK

>KAI0215549_1_Caspase8_Lamellibrachia_satsumaDEDDEDP20P10

MGGSEGKTFGNPGLSGCSSPMGGPEGKKFGNPGLSGCSSPMGGPEGKKFGNPGLSGCSSP

MGGPEGKKFGNLWSTEQTPTSIVVLNKTSLLLSVCCLLSSLRPAEMPSRENMERDAIDEN

MYYRRTLKQVDDGLVTDNISAMKFLCRDFIPGAKLEQVSRGINLFETHYLLSSRVTLFKV

CDDLTSQEVDTMKFMCEGHLKKKLPNVKSCENLFVALEESGHVTAANTKVILDLLSDIHR

EDLHQLFIPLQQPQRNSPQFRPILGSGAANSRTVSHIASNIQAEQELNIQFEQLDIEKGN

LAMPPTNLPLPARLLSPVEDMPGAQVTPSAGHSAQVRPPIVPPNASLDCYDMSADPRGIC

LIINNEFFTCTDLRSPKLKQRRGTRVDCGKLTDIFEKLRFHVSVQSDKTALEIKKMMIYF

STQVDHTAFNCFICCILTHGKLGKVYGTDGATVDIMELTSSFKGAKCQSLVGKPKLFFIQ

ACQGLDKQTGVQVEMDAPESISPSLQETIPNEADFLVGFATVPGFVSYRSTIDGTWYISK

LVEMLDTFHTREHLLDILIKVNDEVSKVAKETNDAVYKQAPAPLASLRKKVFFC

>KAI0229378_1_Caspase3_Lamellibrachia_satsumaP20P10

MQRRATKQLPDRQWSTQEFLYNYKAALPGSRRAFQGKVEDLTTEATAYGQEEPKRMGDTP

DARVATGDTPDARVAMGGGGSADTVESMDTECDEEAYRMNRPKRGVFLIINNRDFDKKTL

QKRRDGTDDDAANLYQLFMGRGFDVRLEHNKTCTEMLRILDGVAREDHTNNDCFIQCPTL

AGKPKVFIIQACRGDLLDQGIEVADAIGETEQKTYRIPEEADFLYAYSTAPGYFSWRNSN

KGSWFIQALCKVFQKDRGERDILTLLTTVNRRVACEFESYVPTDESMHLKKQIPSIVSML

TKQLYLPPK

>KAI0241407_1_Caspase3_Lamellibrachia_satsumaP20P20P10

MKCILSEVAKEDHSKYDCFMCAILSHGGPEHVSGIDENVIIHDLVKPFKGKQCPTLLDKP

KVFIIQACRGKKVDLGVEIEDDDAMELSEEICHISDKPDFLVAYSSSPGFKAYRNVYEGS

WFIQRLSDVFEKDADKLDIMALLTKVSCTVAAYENYEKQIPSIVSTLTKRMYLSPNTDAP

IDMDAYRMGHPKRVVSLIINNCQCQTKGGSDHCGTDDDIAYLEKLFKGRGFNVKPTSKML

KILEEVAKEDHSNNDCFMCAIFTDGSAGGKVDIDNLVNPFKGQQCPSLINKPKIFIILAC

RCDKLNNCAVVTDDGSEQKALIPIEADFLYAYSAAPDGNGHNSTSRHAFLMAVLPLVKEC

RPHIEEVNYMTNSLSSHYRHRMIL

>KAI0216881_1_Caspase7_Lamellibrachia_satsumaP20P10

MCFREDHDSVKIHCGTAVIINNISYKDGDRVGTKEDENNLKATFENLGLEVIIHQDVSTI

ELLLKLQTVAEMDHTTKDCFVCAILSHGGQYVLTQGGEQRMCDIIQCNDGIISAGDLWNC

FTDDKAPTLKNKPRLFFIQACRGLGCPATVQLTGDGQSPKIMTSEKWEEEEKKLNYGFPA

HLPVYPEIMLAWATVPGYYSFRDKKGSPFIEYLCKGLHMYREDDLLTILTWVQNQVAIKY

AALLEQGDNLVNFKQMPWVASMLTKNVYFRNAQT

>ELU00968.1 hypothetical protein CAPTEDRAFT_194348 [Capitella teleta]CARDP20P10

MREARIVTEIVEENIKYGKTPYEQKRELLDYLATRPKSDYNKFIQCLADDNHRDAAKEFALPGFEPKPPQ

PEPSFGRSTQGQSAPYGIQQTPAEASGSIDAPDSLMDTTLVDGPIDVNVQMSSHRFYKDHKEKAYRMDRN

PRGPALIVNVKKIKDRKTRDGTDVDRDNLIKLWDQLHFRTKALNDEDGLEAEEIKQHIKTFVDGVQAEDQ

ACILCLLSHGNNFSILGTDNNEVLIDDIKKTFDENKHLEGKPKIIIIQSCRGRSIDLYETDAVSYRANSN

DFIVGYATKFGNSISTTASYIISFLFSIQTMKLFEPRAEACT

>ELT91797_1_hypothetical_protein_CAPTEDRAFT_220640_Capitella_teletaDEDDEDP20P10

MKAVTRVTECHTRSMRITMETDASVYRQDLLVLLKQLDECLGRNDLVRMKFLCADIIPKG

QMEQISRGIHLMEELIKRDRLTPGNLLLLNQLLYKIGRHDLLRTIGVNKTLVERTFGHEQ

GILQNRILLHDLAEQMTKEDLELMKFFCLDAKITRIPLNNTKNPQELFSLMEECGVIHAG

DLSFLREVFRRMQRNNLLAILGITQTDHVRPLCEGSDTSSSITDFSLHSTSSPPDSVVPL

RNPPMPPFSPQLRSQLPRYKMTSKPRGVCLIINNKIFTKDPNDLHSRQMSDRQGTDMDRA

KLAEVFKKLDFIIKVENDLSDQEIEHRLRKYGQHIDHSRYDCFVCCLLTHGAEGRVYGSN

GRSLKIKELMRYFKAQNAHSLRGKPKIFFINACQGKEKQLAVDLETDSPYQPLPLRSSGR

SRRAETQPLAFATTNPVETDAPGGLERETIPDEADFLLGYASVPGYVSFRSKQDGSWYIS

VLCHMLNKFAASEDLLSILTKVNEEIGRATANQDQKQCPAPLFTLRKKLFFFSENRGN

>ELU11723_1_hypothetical_protein_CAPTEDRAFT_112689_partial_Capitella_teletaP20P10

GVCLIINNEIFSKDPNDHFSRHMENRKGTSVDCQRLKNVFKNLKFIVKINDNLTDVEIVS

KLKKYGQRVDHRSYDCFVCCLLTHGKEGKIFGSNGKSVYIKTIMDFFTAESSKTLQGKPK

IFFVNACQGKEIQRVTQVETDSVGRNPNRETIPNGTDFLVGFATVPGFACFRSKADGSWY

IQCLCEVLTEFAQSEDLLGILTEVNKRVSKHTYDDLAQCPAPSYSLRKKMYFCDRTLIDR

TTKT

>ELT92962_1_hypothetical_protein_CAPTEDRAFT_224953_Capitella_teletaP20P10

MNWLSGSGLMTSSPSSLPALERYNFSHPNRGRCIIINQRNFHKALTGQNDREGTDVDAEA

CESAFLSLGFDVTRHDNLNQTELSMTMSQASKEDHSKNDCFVCVILSHGEDGVIYATNGT

TKIESLVSRFKGNDCPSLAGKPKLFFIQACRGRRFDFGAELPDKSVDELDAVIDDSPKVI

RIPTEADFLMAYSVVPGHFSWRNNVDGSWFIQALVRVIREHGDKMEIMSMMTLVNRIVAY

DFESCTDDDFTSNMKQVPSIVSMLTKQVFFEPKY

>ELU15346_1_hypothetical_protein_CAPTEDRAFT_139929_Capitella_teletaP20P10

MSHPKRGLFIIINNKRFHPKTQMGERTGTDSDAANLYSRFKDLGFDVNIYTDLKAEDMLK

VMIDAAKQDHSRSDCFGVAVLSHGDEGIVYGTDTIIKLETLLGPFKGEKCPSLIGKPKLF

FIQACRGTALDHGVLQHDAMAEENKIYRIPQEADCLYAYSTVSGYFSWRNSTKGSWFVQA

LCKALKQHAREKDLVWILTRVNREVAYEFQSNASKEFMNRKKQVPSIVSMLTKDVVFSSK

>ELU00616_1_hypothetical_protein_CAPTEDRAFT_225565_Capitella_teletaCARDP20P10

MMSMTKEHKHILTKYRVELVEEITLDHALWDQLRTRGVITQSVHRNIMHNNHEKEQIGLL

LDHLPRRPDTSFDSFCEALIATEQDHIVSQYLRQADQPECCQKRAINDSEDVSSSKKICL

QDDKMMEEDDLDLIDGPCDVYVTHCRSQFYENHKRGAYPMNKVWRGHALIINVQTVMGQP

PRKGTDIDRDKLEKLWHAMHFQTQIYNDEDGLGAEEIYEKVKAFVSKTTDEDQACVICLL

SHGAEDRIFGSDGDTIPVTRIIRLLEGKNAPQLAGKPKILIMQACRGKTEDSGVRYEDEP

DNAQLQPSGSDVIVCHPTLHGYMSWRNTQRGSWFISAIVQVLAKHAHDLDILKLLTRVNA

AVSLKESSQGQKKQIPEFSSCLRKDLFFFPGIRK

>ELU08285_1_hypothetical_protein_CAPTEDRAFT_220100_Capitella_teletaCARDP20P10

MDTYQKDCLKQNRPYLQNAIDLSNELLCVLSSKDVITDEHLEQLQKKKRYDTMTSAVLHL

LSEILPKRNPEAFHQFLEGLRCSGQGHVAETLENWLKKNKSNYEQASTSRATPTKPSTKR

MSGSEDQRDDSLSTIDHLLWDIENSPNIMKIKLQERKMSIGGDQTEPEECYEVTRKPRGV

AVIINNVNFKNKFETRDSSLYDVINMENLLRKLHFEVKKHKDKDAQGIIKCLEDERSNLP

ESDIFVAVIMSHGSKNCVYGTDGESVQIEEITNIFNGTKCPILKDTPKVFFIQACQGNDH

ISLSCFRIPRTCHQWKCESVGASQHDGSSVTEPQGDGTDLNLDETDAEPTETNEPEKAHQ

VVVSSTTPGDTAWRYSDLGSPFIRAIVAVFNRFAHLEHLLDMITMVTHVMLQWSKKHQQA

QICEKRSATFSKKLYFYP

>ELT90716_1_hypothetical_protein_CAPTEDRAFT_163805_Capitella_teletaP20P10

MESCKYRGHALVINNHQFKTDALKNREGWKADSENLQKLFNFLRFEQLELCNLNAEEMKE

AIANFARDEKSLNASSLAVVIMTHGDENSFFGVDGRTIHPYDDVVLNFHRDNAPDLIEKP

KLFIFQTCRGNREDPVQQVFVPQYESEALVAGATGGQPVQRKIVPLPNMLLVNATLPGYL

AYRHTTEGSWLISDFCRIVRSKAREDHVVDIITEVSRGLNDREVEAIQQIHPENLLSRKW

FLFPRERSPHDLRTPSEDRLEFVYIIPLVTKPAVQFNGHIVCRPLCNCF

>ELT97848_1_hypothetical_protein_CAPTEDRAFT_224852_Capitella_teletaCARDP20

MSQFSMSEAHRKALRKNRMLLVRDLILNEDFFSCIMENRIITEEMRESIQSVRHQSKKER

AAALLDILSRRGPQAFEEFIRACVFSDQENLAFALDAKMAAVYHAQLEELMNSQPAAPPI

ARPPPNIAPPANKVMPSSQTPAAGYTPPQINKPATTGSMTEIPKEMVENSIMKIPMIPIN

PNDPNIQKEQLNEAISYKMSQAKRGRMLIINNEYFVDPNFSNREGSQFDTVNICELFENL

HFDTTLKENVTAKGMKKLLVDEARMQFNSEAFVLFIMSHGAQGRVYGNDGKYLDIEQDIV

PVFDGKNCPDLINKPKIFFIQACQGDLKSAGLSSDNVDAARGDRYEMNACHDIEAKEVTE

ALEKLKIGDSKDSTPSFDRTVHERADMVLAYSTLPGRENWLKMDFESLFDAPKMIPMPGP

HKEYPR

>ELT90563_1_hypothetical_protein_CAPTEDRAFT_209613_Capitella_teletaCARDP20P10

MRVMLVILVAETKLRFSQLFACEGVVFTKVYLGVHGANLQARVEGQTSCADSIALIVDAC

VMRDQQCAVVQKCKTFLLKHLKPVNVFADELLSENVFSRVQLDAIERESSPVDQKQRLLN

ELMTSQNDDAFSGFLRALDLTGQRRSAEVLRSTERKLQAAVSLTAGDQPPPFPIPASDDS

QYRTESLSHDSVDTVPVETFPVPATSGHDRSASLDLRVQRNRSSAGHLFENIYDMESCTY

RGHALIINNYEFESDDLKNREGWQSDSDNLRDLFQFLEFEQLELCNLNAQGMKKAIARFA

QDEKSLNASSLAVVIMTHGNETSFFGVDGHKVHPYTDILLNFHRDYAPGLIEKPKLFIFQ

ACRGHRQDLAQEVFDPQDEPEDGPSVTGATGGQPVPRKTVPLPNMLIVNATVPGYVAWRN

IDDGSWLIKDFCRIVRSKAHEDHLLDIITEVSRQLNNRNFKAIQQIHPEHLLTHKWFLFP

PRL

>ELT89312_1_hypothetical_protein_CAPTEDRAFT_178431_Capitella_teletaP20P10

MSEDLRPHRRIINDRNDVFATSICFKGKLWSKMREARIVTEIVEENIKYGKTPYEQKREL

LDYLATRPKSDYNKFIQCLADDNQRDAAKEFALPGFEPKPPQPELSFGRSTQGQSAPYGI

QQTPAEASGSIDAPDSLMDTTLVDGPIDVNVQMSSHRFYKDHKEKAYRMDRNPRGPALIV

NVKKIKDRKTRDGTDVDRDNLIKLWDQLHFRTKALNDEDGLEAEEIKQHIKTFVNGVQAE

DQACILCLLSHGNNFSILGTDNNEVLIDDIKKTFDENKHLEGKPKIIIIQSCRGRSIDLY

ETDAVSYRANSNDFIVGYATKFDYEALRAEGGSVYIRAIVQVFMNHARNKDIPSMLREVN

RLVSAKKIDRVDGKHVKQSTEHHDQLRMDLYFFPGYIA

>ELU14824_1_hypothetical_protein_CAPTEDRAFT_227245_Capitella_teletaCARDP20P10

MDSSHKKCLLKNRPFLLKEISMSVDFLGHLSAESVITEEQQERLENIQRNITTKAAVLHF

LSQILPMRGPAAFNIFLKALRDSEQDHVADQLDDWLKKNEENSCMKINLQKTKMSTESTK

GCYKVTHETRGLAVIINNVDFRGKFDKRDSSVFDVINMAELFRKLHFEVKKHKNKDAQGI

IKCLEDERSNLPESDIFVAVIMSHGSKNCVYGTDGESVQIEDIIDIFNGTKCPILKTRLK

RVASESDESDGASQHDGPLVTELQRGVTDLNLDEPDAQPTESTENEKLHRIVVTSTALGD

TAWRQSDLGSPFIRAIVAVFNHFAHLEHLEDMISMVTDVMFQWGKKHSITQICEKTRATL

TKKLYFFP

>ELU08510_1_hypothetical_protein_CAPTEDRAFT_192905_Capitella_teletaP20P10

MSVAEVHSSGRSSFGLLEMKEAIANFARDEKSLNANSLAVVIMTHGGENSFFGVDGQIIR

PHDDVVLNFHRDKAPDLIEKPKLFIFQACRRNREDPEEQVPQYESEAMVAGATGGQPVQR

KNVPLPNMLIVNATLPGDLAYRHTTEGSWLISDFCRIVRSKAREDHVADIITEVSRVLND

REVEAIQQIHPENLLSRKWFLFPRERSPHELRTPSEDRLDFVYIIPLVTKPAVQFNGHIV

CRPLCNCF

>CAH1774000_1_unnamed_protein_product_Owenia_fusiformisCARD

MYTMEKEHKTLLQRKRVELLDINVGPKFMGYLFAVEILSRDMKEELEAIPEPTRRASRFL

DLIVRRGPKAFNELYEALIRTSQGDLADILKPDQAEKRQKDEDRRIEAMKASMPASPSHG

SSPLRRSGDSFRGQPIDTASEDEPLPEEWPLPKDSWSALAEINSMKEMRIQSQAWEDRYN

GKNPDYYPMNSSPKGRMIIINNKNFDKSKVKLSYREGTDNDATCLSQLFRKLGYSVQDSK

SWKDLTAKEIKEKLQKESQLESHNRHTSFICAILSHGAKGNRIYGVDGDTIAMKDIKEIF

QNCPALTGKPKIFIVQACQGSAKDTGVLDTSFNSSMSYDTDQGRQSPVQFSNRIKPAPCE

STDPSLKSPTESGDDSMDASYGEFGDSIDSVPTSMPTAQSCDSLKSLPGNSDILVANATT

DDMASWRIPDVGTWFIQAIIYVFARYAHNTEIGHLFRKVNHLVSKATTKKLPLQYKQQSC

IDETLRLQLYFYPGMGSENVL

>CAH1774649_1_unnamed_protein_product_Owenia_fusiformisCARD

MDPNHKKILIKNRVALIDLDVNEEFIGNLKANGVLSRNMVEDIEVEQRRSRKAGVFLDIL

ERRGENAFDEFYEALIKTEQGHLADILKPDMAASRKSEQLARIAGNKSALPQSNSTGRHF

QSSTVQSMNGQHIHYPSEDFNVRSSIPVEDEELPSEWPPKNWDPYKEIEQMKVMGHPSDW

PTKVHDRFVGNSKECYAMKGRKGRAVIINNQNFDDTSTLTNRSGSEQDAEILSKLFTKLG

YDVRKYDNLTACDMQKYLNNEAKQDYHNYCCSFVCAILSHGGEGNTIYGVDGDTMSVKDI

TLIFNNQNCEYLSGKPKIFIIQACQGNTKDTGALDRSDSSNNILESLSLDTTDGSISPLK

GGCESSHGRYYVPDGEDSVDGSYPGLDRCDSVPTDSDILVANSTTEGKTSWRVGGRGSWF

IQAVVYVFAKMAHNTDISKLFVKVNGLVSSATTKTNFKQASKVENTLRKDLYLYPGIGES

YC

>CAH1787639_1_unnamed_protein_product_Owenia_fusiformis

MSSPHSAGQWGFKILLSWKQRQPRSVNKRKVLADALREVYRDDLAGLVDGNFKPKSRSTT

TKAVIPQQEVTRQHTSSIVEPKTKYTARGVPPTGSGSDNPEDTDSEEGNSDDDSTKEFFE

KWRNHDGVYINSSTPRGRCLIINNFKFEPEMRNRYGTEHDLNSLKDLFTKFKYTVTVKTD

LTARDIKDTLNEESKQSMHDMYESFVLAILSHGTVDAVYGVDNETVPYKEIYNYFNGENC

KSLIGKPKIIIIQACQGKDEDGGVVPDGGEVKQDDNVDASSPSPSDLMDEKDYPDGAQKS

PTMADFFIATATTQDYVSWRNTQTGTWFIQALVHIYRKYAGKKDISAMSSKVNSLVASKT

STAGKKQIPDFRNRLRKKFYFFPGH

>CAH1774650_1_unnamed_protein_product_Owenia_fusiformisCARD

MDPNHKKILIKNRVALIDLDVNEEFIGNLKANGVLSRNMVEDIEVEQRRSRKAGVFLDIL

ERRGENAFDEFYEALIKTEQGHLADILKPDMAASRKSEQLARIAGNKSALPQSNSTGRHF

QSSTVQSMNGQHIHYPSEDFNVRSSIPVEDEELPSEWPPKNWDPYKEIEQMKVMGHPSDW

PTKVHDRFVGNSKECYAMKGRKGRAVIINNQNFDDTSTLTNRSGSEQDAEILSKLFTKLG

YDDMQKYLNNEAKQDYHNYCCSFVCAILSHGGEGNTIYGVDGDTMSVKDITLIFNNQNCE

YLSGKPKIFIIQACQGNTKDTGALDRSDSSNNILESLSLDTTDGSISPLKGGCESSHGRY

YVPDGEDSVDGSYPGLDRCDSVPTDSDILVANSTTEGKTSWRVGGRGSWFIQAVVYVFAK

MAHNTDISKLFVKVNGLVSSATTKTNFKQASKVENTLRKDLYLYPGIGESYC

>CAH1786849_1_unnamed_protein_product_Owenia_fusiformis

MEGEHTVPVGPPDYVENGDHQENQNNQGNPDNYENPENKENAKYKDNQEDNESDENQEEE

YGEVIQLETETIPELDDLNIPDQKTQEMNEIFKKFIYVQLKNDHQIQDIIPEGGGELQSS

RPGVEETLRVARHLATLGDDLGTRYDAQFTEMINAIPMSPQTGREAFIGIARQLFSDGVV

NWGRIVTLMAFSGRLSVHAARQGSAAMAHNISGYMTRFFVQEDIPKWVDDHGGWENAFNM

GSKEEVTFDVLLDRPEYLEAYQRAREEGTRHVYQARVMIVGQYGVGKTSLRKRLLDEGFS

KQYKITDGIQVDPSACSVDIKNSVIWKTKAKGFKAGEDTNDPLEVYESEVAKNLWHAFND

GTSVDGPVIPVSQGAKTTNGLKPKESRKGTDTIDNVITKEHVTPVNTDKPGTSSDYNNST

NNRETLRTDGDSSAPDYISKIPESIKNKFLSLTEDVTKQIRQLGGLNGDSSEDEKQLKLS

FWDFGGQHVFYTTHQTFLTNRAIYLLVVDLTKDLDEKVETFRQSRSGTKLDTTAPETVKD

YLDYWLNSIHTHAGKSSSGSESLSPPVIVVCTHKDAVSMEYHMKQPYMSRFKAWWQGAVL

YDTEQQSQHIENYFHKIDMNLQDKVYYRHVCPRYFAVDNQYNGDPELDCLKKMIVKLAED

QGFWGQEVPIKWLVLEKKLREFKLETPDRKARHFLEFDEVAEIGFKEGMDEDSVRACLGF

YHSVGDMIYFNEINLNDLVILDPQWLIDVFKSIITVPEFHRVVSGSGRNSRIYWEKLDKE

GLLEEELIDIVWEEQNTEISNHKKVLIEFMQKFDLICPLKNNPGYSQKKRQFFVPCLLPK

PEPSDIIKDKPLAGPTIFYQFTFLPNGLFHRLVANICLEKSWPLYGKNVFFDYARFEVGK

GAHVLTILAQDNHLELKIHTLSDDVTDKDNNFMCMRIRKDIEAMLKSTIQKYCPSAGYKV

SVRCRCPGVPEENQLIVPVSEMDIILSQNKLCELHHNTIKLQSYELWYQTVQIFKPVPTK

PKEAKPKPTRKKSSKKSKKQRPSDTDDELSASDSDSDGSNLEDTDSEDNEDDGELHVHKS

TQKFYKKWYNHKGAYCNISMPRGRCLIINNYTFKPKMSNRDGTEHDLKRLKQLFKIFKYT

VVVKTDLTAGDIRKVLIEERNRSIHAMYDSFVLVILSHGTVDAVYGVDKETVPYKEIYNF

FNGENCKNLVGKPKIIIIQACQGSKKDDGAVADGDEDENIDDIDSSSPPEPTIESASMDE

APTPLDEVDSPDGAQKSPTMADFFIATATTQDYVSWRNKETGTWFIQALVHIYRKYASRK

DINTMSTKINNVVASNTSMSGKKQIPSFNNQLTKDFYFFPGH

>CAH1787638_1_unnamed_protein_product_Owenia_fusiformis

MQFCNLSLSEVFTLDRTRRVLRYNQRYPEYLEAYQKAREEGTREVYHTRVMIVGQYGVGK

TSLKKRLLDEGFTKDYNSTDGIQVDPSACYIDISNSVIWKTKAKGFTLKEDTDNPQDVYD

SEVAKKLLHAVKDRTSGKHVRDSSTGATCTIPPDRIPDTIKSKFFSLEDGKNQEEKQLKL

SFWDFGGQHVFYTTHQTFLTDRAIYLLVVDLTKDLDEKVDTIRQSRRGPKPDSAAPEKVK

DYLDYWLNSIHTHAGKSSPETQSLSPPVIIVGTHKDAVTDQYIDDYFHKIVNSLHDKVYF

HHVHKTYIAVDNKSDDDHELNKLKETIVQLAEGQRFWGQEVPIKWLLLEKNLRGLKTKSQ

DREPVHFLKFEDVKAIGFKERMDEASVVACLQFYHSVGDMIFFNEIDLRNLVILDPQWLI

DVFKSIITVPEFHRDSRGRNIFGSIVWEKLDKDGVLQEEFIETVWKKLYADISIHSDVMI

ELMQRFDLICPFENNQECNQGKRLFFVPCLLPKPEPGDFIRDKPLVGGTLFYKFTFLPKG

LFHRLVAKICLKNIWPLYGAVFFDYARFEVGDGCCVLTLLAEDNHLELKIHQTLEERTDR

GDNLMGIKIREEIEVILMTTIKNYCPNVDPEISVRCICPNVEEEKRTLVPISEKDINRGH

KQCKSHHHSIKLDQYKLWYSTAHTAETKSANTKSGEKTTKQNRGKSAAGSSDPEGSNLED

TDSGENIDDEHIHVHTATAEFFEKWSKDEEIYTNASMPRGRCLIINNYAFEPKMRNRDGT

EHDVKRLKQLFERFEYTVVVKNDLTAGKIRETLTEESSRDHADSFVLVILSHGTVDAVYG

VDKEKVPYKEIYNFFNGENCKKLAGKPKIIIIQACQGKKKDSGVVTDGEEDENDDNVDVS

SIASFKSLDLSDDSTAHTDDEEEDSPGDTSPIMADFFIASATTQDYVSWRVTDSGTWFIR

ALVKIFRKHAGTKDILRMSTMINKLVATKTTRSGKKQMPTFNNQLTKTFYFFPGHGCKE

>CAH1794547_1_unnamed_protein_product_Owenia_fusiformis

MTTTSTDAYPIGGRETVLMTKPVAGLSEYNFSHKRRGKALIINNRNFDAMLTGQMDRDGT

DIDADAITARLTDLGFDVDVLHNLTTTKMSFALRNAAEEDHSDADCFVCVILSHGRDGRV

YGIDGTITIESLLRHFKGDECQTLAGKPKLFFIQACRGTKFDHGAEVTDAATDEGMDEID

AAPVVRIPSEADIMIAYSTVPGHFSWRNNVDGSWFIQALCRVLKEHSHDMDLMHMMTFVN

QIVAYEHASCTDDDFTSNMKQIPSIVSMLTKLVYFRRN

>CAH1795557_1_unnamed_protein_product_Owenia_fusiformis

MDERDSKPNSQDQNNTDALTQQVGQVNMGDTIDGRNTTQMMAGLRVSNNEDVNNEDRYNM

KHPRRGMAVIINNRSFQSSTGMNERNGTDVDASNIFQRLNDLGFEVIIKTNQTTRQMVDL

MSKAGKMNHSDADCFACVILSHGEEGVVYGTDGVVPIEKLTEPFKGNKCESLRGKPKMFF

IQACRGQKFDEGVETTDDDGEGLDETDANTVHRIPAEADFLMAYSVVPGYFSWRNSTRGS

WFIQALCSVLEKYGENPDYDILRIMTRVNRMVAYDFESRAAREFMNRKKQIPSIVSMLTK

EMYFTPRTG

>CAH1781235_1_unnamed_protein_product_Owenia_fusiformis

MSMETAHKRHLIKNRVKLLDLIVNEEFISYLRSTSTLSRAMVEDIEIGLDSRDGESLKGQ

IVQIVQQPSSPASSPLEEEFELPSEWPLKDWGYLSEIMNMKLCDQNTCRPMFNGGNNSYP

MNAPCKGRVIVINNEEFVKKANRKGTDRDANSLRELFLQMGFDVLPLKNWKNLTATEIRT

KLKHQSQQLKELYDCSSFICAILSHGTTGNMVIGVDEDEISVQEIKDIFCKCKALSGKPK

IFIVQACQGEERDEGCLDHSFERSTSSSSGGRAEPSESLDPSAISKPPVDGGDFVDGTPE

SMQSSPTSLPTYADTLVANATTEGMAAWRHSEDGSWFIQAIVYVFAKCAHRMEILDLFRQ

VNNLVAIAGTRTTKSEYKQASTSSSSFRRKLYLFPGMSGID

>CAH1774651_1_unnamed_protein_product_Owenia_fusiformisCARD

MDSKHKKVLNKYRVRLLQHLKISSSFISYLQAVEIMSRAMKEDIEIEQKRSKKAALFLDL

LERRGERAFDEFYNALIHTDQAELADLLKPGVAIERRAIQQQRVEEKSAQLAAMSYQLQL

NPAVVDQYQSSQSSQYQSSQNPKGQSQPNDIVVISNMNSMRGQPLGTSSDDGNCISITSL

VESNQSSCPIENDERPLDEWPADDWAPVTEIDKMRTSPDDWKTRISGGNKYYTMMKRGRA

IIINNAKFNGKMGNRPDSKEDVKNLTHLFSHLGYTVSPTKKWQNLTAENMRSQLRKESRH

NDHQDVSSFVCVILSHGNNGLIYDVDGAIITIEEIKTMFNSKNSPHLAGKPKIFIIQTCK

LTANGMDNESSIESNLSSMPSMSFDINDGDRKGLQGPNDVKEECDSVQVPDDSDMIVVSS

TTEGTDSFSSGRKGSWFIQAVVYVFVKHAHNTEIRELFTKVNQLMLKEGTKQARKQHASD

TPKLTSQIDHTLQKDLYLYPGLNSD

>CAH1795558_1_unnamed_protein_product_Owenia_fusiformis

MDERDSKPNSQDQNNTDALTQQVGQVNMGDTIDGRNGGSDERYNTSHPRRGVAIIINQRD

FDPDTGMGARMGTDRDADALEVRFYALGFEVVRYNSITATLMVSVLNKAGKMNHSDADCF

ACVILSHGEEGVVYGTDGVVPIEKLTEPFKGNKCESLRGKPKMFFIQACRGQKFDEGVET

TDDDGEGLDETDANTVHRIPAEADFLMAYSVVPGYFSWRNSTRGSWFIQALCSVLEKYGE

NPDYDILRIMTRVNRMVAYDFESRAAREFMNRKKQIPSIVSMLTKEMYFTPRTG

>CAH1775431_1_unnamed_protein_product_Owenia_fusiformis

MDLKRQILQFFESHPDESLQALDLKHLGQKKDINHQLYALQKEGVIEKVQEAYLPLWKLI

SQPMSHNQDGTLSHMGDTVTTNETTATKAQPSLATSDQLITSQMKYLDPSLKNLFDQAGI

SADQLQDKETASFIYDFIEQQGGLDAVRKETAPPPPTSRNRRAPPPPPPSRGLQRGLGAS

PPPAPSRSMDAPPQPPPLSQIRQGAKSKEMNNLDLNNLDPSLKSVFDQVGISAKDLQDKE

TATFIYDFIKQQGGLDAVRKETERGCSPATFKPVKSGMAARPPPPSRNRGAPLPPPPGRN

RAVPPPPPSSRGLQRSMDAPPPPASSKGMGASSRPLPPRQGTKLKEKSDPTNKTKFTSAP

DEECESALVPPESVNYNIPGYVFAIGETQCTNCSVAFSQYESKFTCDCRTIALCKKCSID

HTQHGTQRMFKENVIQKHRYDKLRTSITKNLLGKLKSDITCTSILKGSNVNKVLIDEEIF

GRSQSAAGLAIICGHKDILQYLLENYGTSLFKEALGTWQKSLGMLVIQNPQEDLACLSYI

DEFYPDLIRKDFGNQHTGNALHVACSDSVSVKASVSKSHLKQVNYLIAQEFDLNAKDSSG

NTPLHNACKITDPSIVETLLTHGANSSAENHEMQTPLFHLMASIEDKQPCHVMIGKNVRK

CIVLLRKAGASHKTSGFVSIDTGSACKRVALSDLKAYKRPGVKDAVTLRDLKIIEGINPN

SRDLKMDVTYNQVPEEQGTGSSKLACDIYPQSFEEPEKIQVSTENLEEDDTSSSHPLGKR

LAMITNQSPNELLTETIETNSTTPLSAGASKTDITGHIIDIPSHSTCVVSTETSDAIPVT

HPTTTTGEPITDISSYITGKPACPSTSNVSTEIMIASQNIPAAGGSAISSNIVPTTTAAD

TVASEMSSLTVTSSSSSQGKPVNAAEMDLAQMCLDFKQMHAELLEKELKETLGQRENVLL

SNYASKMSDINEMLKKLAVSKYEPILDVVIEKVKPDDLGMRQLSKNLDEVYKMTSKPRGR

ALIINNEDFSKTKLQNRDGSTIDRVRLETLLRGFDFEVETLLDKTAQEINDILEDERRRH

DHHYNDAFILVILSHGTNKPNSCVYGVDGAKVPINTITSYFEGKNCPDLKGKPKLFFMAA

CQGNTCHSATDYANAPSPDESQMSEPNIPSIPDTDDGPSDKPHKHTKADMLVACASVEDY

VSFRDEQLGTWFIYHLVKVFGEEAHQKDINKLLTKVKRDVANRKTPSQKHIMIPNIPYNT

LVKELYFFPGIVGTSGVTLPPDSNF

>CAH1792905_1_unnamed_protein_product_Owenia_fusiformis

MEALNLQGPKDEKYRYNMKHPKRGMAIIINNKSFDPETRQQERKGSDVDASNIRQQLHEL

GFKPEVKTNQTVKEMLGLMKKAGSMDHSQADCFACVILSHGNDNGEVYGKDATVSIDTLT

EPLKGNNCASLAGKPKLFFIQACRGSKLDEGIEVTDGHGGDDIDETNTNTTYRIPAEADF

LMAYSVVPGYFAWRNSREGSWFIQALCIELEEHGEDPDFDILKIMTRVNRRVAFDFQSNN

PRIPAMNQMKQIPSIVSMLTKELYFTPKDDD

>CAH1792904_1_unnamed_protein_product_Owenia_fusiformis

MADLVDGTPTSKTAARAQGDTQNVAGLEDNMEALNLQGPKDEKYRYNMKHPKRGMAIIIN

NKSFDPETRQQERKGSDVDASNIRQQLHELGFKPEVKTNQTVKEMLGLMKKAGSMDHSQA

DCFACVILSHGNDNGEVYGKDATVSIDTLTEPLKGNNCASLAGKPKLFFIQACRGSKLDE

GIEVTDGHGGDDIDETNTNTTYRIPAEADFLMAYSVVPGYFAWRNSREGSWFIQALCIEL

EEHGEDPDFDILKIMTRVNRRVAFDFQSNNPRIPAMNQMKQIPSIVSMLTKELYFTPKDD

D

>CAH1787632_1_unnamed_protein_product_Owenia_fusiformis

MQTKRIRRESAPYTNASSPTGRCLIINNFDFYPRMPNRYGTVHDVDRLKRVFTRFKYIVC

VKNDLTAGAIRKTLTEESTQDHADSFVLVILSHGAVDAVYGVDGKTVPYKEIYNYFNGEN

CKKLAGKPKIIIFQACQSSREDDGVTSHERSVDCSVQQVEADSSAASLPMADFLIGYATT

ENYSAQRVISAGTWFIQEMLNIFEKHAVEEDILTMSSKINALVADRSISTGEKQMPYFRS

SLRKKFYFNPPDIDAMNI

>CAH1787631_1_unnamed_protein_product_partial_Owenia_fusiformis

WYNTAHMGETGEMQTKRIRRESAPYTNASSPTGRCLIINNFDFYPRMPNRYGTVHDVDRL

KRVFTRFKYIVCVKNDLTAGAIRKTLTEESTQDHADSFVLVILSHGAVDAVYGVDGKTVP

YKEIYNYFNGENCKKLAGKPKIIIFQACQSSREDDGVTSHERSVDCSVQQVEADSSAASL

PMADFLIGYATTENYSAQRVISAGTWFIQEMLNIFEKHAVEEDILTMSSKINALVADRSI

STGEKQMPYFRSSLRKKFYFNPPDIDAMNI

>CAH1776711_1_unnamed_protein_product_Owenia_fusiformisDED

MAKRQRLLSLKDRKEGLHRMLIEDLSLHISKEGLDGLKSVTPELGEGRAEEITTFRKLCT

FLENRAFLGPDNYDRLKQMFESVGLYKLIERLDEAEKNISQLGVDTPSGSPTMISPTTPG

STPMPNSPHFDASHGLAATVNWGNNDDLAYPLNEGAYPGLALIINNFTSPNQASYRKGTI

EDEKRLVDLFQNTLKFRVRVERDVQRESLGDILTQCQAHLEHEDKFFKPTVEKYNCFVCV

ILSHGNKDGPLMVPTDNKKQKSDYVSVDEIVDHFKPDKVKSLKGKPKIFIIQACRGSNFQ

QAVDDESQQTPEDVTKQEEIDSDINPRVSTPVDADIVIQYSTTKDYVSLRNKQTGSWFIE

ALTDTIFERHKQEDLMSMMTRVNKTIAQKSGMVDEKITKQMPCIVSTLTGRVYFN

>CAH1789637_1_unnamed_protein_product_Owenia_fusiformisDEDDED

MVNMDQVPNHPPQSTFRYVKELTEDENKVLSTDVELDHIRAKNNLKMRKDLKKISDQLST

ENITDIKFLLRDLIPLGKLEQVKSGIHLYDALESQQLMRISFDEPNFELLVDLLYFIFRY

DLLLILGCEQHNIKARLKLRPSTCRISEYRLMLYKIIINLQEQEKRAMIGEARPWFNHLP

KEQFECIKSLQMLFIQMEKIRYLAEDKVDKLQWLLHKIGSMDLVEDLKKYKRNHYCGASY

GEGNIEEASPYAGDINNTYNVNPQALDLHVKHQKYAMTVPLTDPKGPPSRPSEESEPEFE

KSSSKDLLQATDPYSFPPPQSPGLSDQVKMVAIEDKGNKLLNQTVNKYEASENIPSINHE

PIVPASIDELSDDEQMDDSDSHRSDFDDSSDSLASPINIARPPGTRDNMNASLDNRTISN

AQDLTQFMLPGDSTNVNKHDKLLPNRNDNNLEEVQYLEFGKDKQWIPKSQRIANLATTQS

ASPVSPSPKAPEPHPSPSLVSGSGGNKVGLNLVSELHGAKSSKGKDDIHVSERRGHVSQA

GQVQGQGKLLVGAYQSASPVNSQDLTINNLETDSEPEVVSQMLESYTIESKPKGLCLIFN

NYKFDKLPGGQNSQVQALEDRNGTDVDLEKLKKLFTQLGFQVVVEENLTSCDMLNKSVAY

SRTRKFSEFHEKSDAFVWCILSHGIRDAVFGTDGKEVNINSLTYFFKGSNCPELIGKPKM

LFVQACRGIHESKAVVPLKKSEQFRTDGPTATTPDEADWVIGQATSQGHMSFRSESHGTW

YITKLCENLNRYHGRKYDLLQILTKVNQDLAKGEADFEDKYGNSYIGKQQPVPYFTINKN

IYLTCINSKCKRCS

>CAH1776712_1_unnamed_protein_product_Owenia_fusiformisDED

MAKRQRLLSLKDRKEGLHRMLIEDLSLHISKEGLDGLKSVTPELGEGRAEEITTFRKLCT

FLENRAFLGPDNYDRLKQMFESVGLYKLIERLDEAEKNISQLGVDTPSGSPTMISPTTPG

STPMPNSPHFDASHGLAATGTIEDEKRLVDLFQNTLKFRVRVERDVQRESLGDILTQCQA

HLEHEDKFFKPTVEKYNCFVCVILSHGNKDGPLMVPTDNKKQKSDYVSVDEIVDHFKPDK

VKSLKGKPKIFIIQACRGSNFQQAVDDESQQTPEDVTKQEEIDSDINPRVSTPVDADIVI

QYSTTKDYVSLRNKQTGSWFIEALTDTIFERHKQEDLMSMMTRVNKTIAQKSGMVDEKIT

KQMPCIVSTLTGRVYFN

>CAH1789638_1_unnamed_protein_product_Owenia_fusiformis

MKVSMRDTGDNIEVNNYEASENNAEDLSQSSEIPLVPLSVSALETEVDSESGQLQDITSD

EESNFDVSNSLSPQIFKPPLNTTTPSKSEYIQNLENKQAEENVQDLSQILKNEIHGAAFD

DGSQFGQSTEEFLGNDDAVVCVPGKISEPVVKSLHEKECEPMEETNSELGIENREFLPRP

NNGQYAICTPENVNYAQDLTTYASNHDSDVRSHTIGCMANNYHDEKNRSDWVPKKNKPKR

NRASTTGKPLLQSAVPVEHQSNLIYGYPENGFNYSKEASVKIEKNNVLNQAGEFHSDIRL

HDAEKNEAGFIQTAHPVFNQPQDLTIHRIPDDVKMKEDLFQTASPVILSQDLTMSSPEIE

VEQNPNTTMLESYVIESEPKGLCVIFNNHVFHHLQEDKDSVKLTNRKGSNEDKERLIKVF

SQLGFEIHLKENLTANEMLNRANYYARSYKEFHKKSDAFVLCILSHGEGDCVFGSDGKKV

PLHDNTEGCLAYYFKGANCPALIGKPKMMFIQACRGPQEGIAVKPHSYQERQTLKPCSHA

IQTDGDIPSSQGTTRIQTLPDEADWVIGQATSIGHASFRNEETGSWYITKLCSNLHRYHG

RKYDLLKILTRVNLDLSKAEADCTDKNKKTFIAKQQPAPYFTTNKNIYLTCTNTKCQRCF

NQPNGKT

>CAH1776205_1_unnamed_protein_product_Owenia_fusiformis

MAKSSQKTVMRQKGTWERWGSVQRARRGEVIDEPERVKQNGHKVAKEKATLESLQFLNPN

LSKPFVNNRYKKKYMIIPDNMKLSRSRLRQLKRCDTVVLKNSQWMDLVDFYTNRFFRIMR

LVGSALSVLALAGNVLSLRTNNWSTQNGLEKVVLTFMGFTAVFDLIAVGLCTRGIFQENM

FEIIYNFSSAAELQIICTFGNHWNSQYLADLKMNPDDITTPDDSPRDLRVKCARSGSLAS

ISSQDEEVENMRQLPLAERKEQILRKLCDLNNQLRNINKKSNLVLLSDAINMLQENGSNE

EVTSLESLIEIWQDRSAEANKELCSHIETYIFEIHEIYGIFNSLKTHIIPLVDVHFNKNN

IEIDEHKVWAQKSWLALREEIKILENTSQKGEIESSQLSSKDARKYKFKDKQVSLPRNDP

VLYLRSYMIEQLTSILMYLHQWVLFHNNAANLTLLRTDRTSINQAIEHFELETKLLEKEI

GDLKETLKQRPNIESDDAILEALHANPEFAEEIGQMAAKMKELKQEHVKDNDELKKKQKR

KDDIEKLFKGVKKPAMKKELLDLEKSIKTLTARVTNFEQRLNNHDNVKEQLKYKYLETFN

KMEEDIKVKTARIEAINEQIMEPKEKLEATVKNIGEIMFQLTRCQKNWMDIGAKNWHSLT

ECREVTPCISRTTPEALRARWVSYYSIMNPRGVALIISNEPTKLAKGERPGGRHDVNTSR

NLFKNLGFDVRVRENLTSNEMTGVMKSLASEYNSKSTSYSLLTIVILTQGTNGKLLDKDG

KSVSLPSLLECFSTTNCPALNGRPKLCIIQRCQWSDNINVENIDDVDNTTINQPEETINS

TKETLSKNADFAIVEHQLNNWDNLRDEEGGSLFVQIMAEVVAAKAFYLDFSAMFSEVVRL

LRARHTPVFPTILSTLTKSLYFFPGDSTRPSP

>XP_009017805Helobdella_robusta

MDSGYQSNKDNDSKYNYAMTHDTRGIALIIGNESFKDSNVADRSGNSKDLANLKALFIMLKFSVRPHTNL

TVKKMKELIEEVASEDHSKYDCFVLVLMSHGGQEFIYGVDGVINLKELLLPLNGNKCKTLIGKPKLFFIQ

ACRGTEIDEGVEYKAKFETDADEVEERIMKIPKEADYLYSYACADGYVSFRNANGAWYVQALCEVFQNYS

HKYDILTMLTLVSCKIKDFESNMTNAAYNKKKQIPTFISMLTKRLYFC

>XP_009018546Helobdella_robusta

MTNKERGTFLMINNKMFNSKTGMTERTGTDVDADALCQLFQKLGFKIRRYDDRTVAQMIDVIKAATKEDH

TNNDCFALALLSHGDDGIIYGIDDHIPLEFLLAPVKKCSTLAGKPKLFFIQACRGTKLDPGTKVESDSGD

QQTIKIPIEADYLYAYSTVPGYFSWRNSLKGSWFVQSLAATYDDVLRQQQQPVDLLKIFTRVNKKVAFEF

QSNSSSLNMDKMKQVPYIASTLTKDLYLYPKK

>XP_009026608Helobdella_robusta

ETAGICLIINNEKFDGELKLTERRGSDIDRDQIKTFFQKLKFKVIVYENLTANVMQEKLNETSKQEVKES

HDCFVCFILSHGEMGYVVYGTDCQEVPLLSLTTPFHKGHCPQLENKPKLFFVQACQGNQYNRGSINLADQ

LEEDSAKKFKVMEEGDFLLSFASLQGYASMRSPTTGTWFIASMISVFHEHYQEYDKFIICFLGVFLLKI

>XP_009029017Helobdella_robusta

MANSICENLKLEEYNFKSYSIIGKCIIINNEHFHEKTGQSNREGTEIDARNADNAFRRLGFEVTLYNDLS

WYEMRNAIQEAAGDDYGNYACLVVVILSHGREGKVYATDGTLQIDSEVVDKFKGENCQSLAGKPKLFFIQ

ACRGSKFDHGVEMRDYVDGVSVNEVRIISSEADIMIAHSSTAGFYSWRNNVDGSWFIESLCNVLNSDLVN

DVDLVSLMTLVNKLVSDNYQSKNTDEKFSSNKKQIPCVLSTLRKLVFLRPRD

>XP_009023634Helobdella_robustaCARD

MRQIGYLLDLLPRRGPKAYEGFIQALQSTGQTHAIEILEPDPKPLWGDEQAKAIAGKISNYCSVSVFMQT

TFVDGELELINNDYLMKKSKYSTLNYALQCIYPVNKNYKNRVVIININFDNGNKARRDGSEKDFENLKEL

FGLMKYDVIGKTELSKKEMDEFIVEERSYLNTSNESHNVWIMIIMSHGTIGGCYDADGRLHAFSDIVNNF

DGVNCPGLKGVPKIFIFQCCLGNKKSYPCGTLFNKCESASADYESSCTTAIPPINDRHDKADVIRLFSTV

EEYVSIRNPELGCIYISAFIYVLSNMCRHENLQTMILRVSFHSTLVQIIK

>ALB38959.1 caspase-3 [Urechis unicinctus]OK

MKGVQEKNSIHDVELLEKVMRGLGFDVITHQDLTTSDIHKELAKYANDNEYNKNANCIGVAAMSHSKKEGQFKAADMTIS

IEEILKPFKLNKKNEDLRNKPRFFFFQACRGTEHDTGHRIEGIKTDNDSETSVPDSKVDRVLSVDRDDDQTEDDDRSQLL

PVVSDMICAYSTYKGYTARLGTDGSIFIQTLCSTLTKGAGEFDLVELLTIICNDVTQRDIKFKNKTTEVEMTQKQTPTIV

SSLTKKFYLKQR

>c73072_g1_i1CARDOK_Alitta_virens

MNETDRNRLRSNTVLLTRDLTIDKMFLAVLQQRKTINDTMKEDIECHRKR

TDKNAEFLGLLPRRGPNAFNDFIIALMETSQDHLADALSPELAAKWRTAN

PNGPSVPRGREEEECRIGGEGEFANARPHPGAFVPPAAGDEERDSFIPIN

LQPSNPVFVRQVLNNPTDYYRMAAKPRGRVIILNNETFDQKAEKKLSERD

GSYFDVVNLTQLFDALGFKTELWSNLQSEKIRKQIDEERRSDFHNTADAF

IFIILSHGTKGAVYGTDGKPVLIEEEIIGPFDGNNCPQLAGKPKVFLIQA

CQGDKITMGIIKADGGDADDAVIEAKKEELEDQKDAVPMGESLIHEKADV

LVFYSTVPGYKSWRNTELGSWFMRDIVININQYAHCLHIYDILTKVNNQL

SGRQVKYKGFRQISKFDSTLRKSLFFLPKS

>c83731_g5_i6CARDOK_Alitta_virens

MNQDDRNKLRSNTVLLTRDLHIDTMFLAVLKQRGTLNDTMIEDIECNRKK

TDKNAEFLNILPRRGPRAFNDFIIALMETSQDHLADALSPELAAQWRKAN

PKGPTPMARDNDSGYGQGYGQGYGQGMERLHHATAHPGAFIPPAEGDRER

DSFIAINIQQTNRLFVQQVLNNPKDYYKMSAKPRGRVIILNNEIFDSAEK

KLPEREGSLFDVVNLTELFKSLEFETKHWSNCTADEIRGLVKEERKSPFH

IDADAFIFIVLSHGTRGVVYGTDGGKVQIEREIIDSFDGNNCPQLAGKPK

LFFIQACQGDQITIGAIKSDSDDTGDAESDTKNDEMEDQKDAVPTSGSFI

HEKADVLIFYSTVPGYKSWRNTELGSWFMRDLVININQYAHMLHIGDILT

KVNNQLSERQVKYRGFKQISKYESTLRKWLYFLPK

>c73773_g1_i1OK_Alitta_virens

MDQSDAVPEAVPDLISSVECGASGGASAASAQGSGDQPDARSLNIFGGSS

QRQSTTEASKPVEKMDYEEVFQFEYNMNYQQRGMAVIINNRKFDSKTGMN

ERSGTDVDAAQLYQKFKMLEFNVHMYNNQTCGSMLKIVTEAAKQNHRDSD

CFLLAILSHGDEGVIYGTDGIIQLDMLVNPFKGDKCPSLIGKPKIFIIQA

CRGNKLDHGVEQADADSDMEEMPRQRIPAEADIIMAFSTVPGYFSWRNST

KGSWFIQALCKVMERFGKEKEMLWILTRTSYLVAYEFESNASKEYMNKKK

QIPCIMSMLTRELYFRPKKI

>c84685_g6_i6CARDOK_Alitta_virens

MASVEAGMKKMHRDCLRLNLTYLTRETLLDALCDSLKENGVMKEDWIDDI

LSYNRPPKDTCHDLLEKLKCSGPKAWEGFLTCLDETDQKHIAWRLRETEN

CYRENPTITARELKEYFVTNPAPSELLPKPEKRASLHSHSSMVHINSEFY

DQPHPNAKQNEDNCYRMNSEPRGFMLIINNVKFTDGQTERLGSDQDRDRL

ISMFEGFGFVVNSKDNLDKKKMEQQIQIFAKTANDFKDIDSIIVAVLSHG

VDGVVYGIDGSPLKVDDLIGPFNGIRSPNLIGKPKLFILQVCRGRDYRMS

VPSTDGPKSEPKAGPSNSIMVQGGESQDKPAIPTSVDEELSDAMAQWHLD

LKEAPRYVHPDNDMLLAYSTLPGQLSFRDDKKGSWYVQALCNVFQDNAQG

VHVLDMLVRTNREILRFGGESYVQCPMPMVGLTKDWFLNPL

>c83651_g1_i1ANK_OK_Alitta_virens

MDTQPSGVMELKRACMEGDLPKVKELLNRQNVNHKVPHMLGQTTSLLHLA

ALCGHQKVVGHLLKFGADMSVRDHTGDTALHYAAQKGHTQVVRLLVMKGA

DVFLEGNHQKTALHAAAQSDHIDTLDYLLDVGQINVVALDKDNRTPLHRA

SMNNHSRSVKRLIENGSNVNAKDASQNTPLHLAVKGGKLKNEIDTVQVLL

RNGADRTAINIKKKRALDVAHPRLKLIIALSQPSLDGDKECIYININIKP

ADPELVRRVLNNPEDCYKMSAKPRGRAIILNNEIFGKSAESKLSDREGSS

VDVVNLSNLFDNLGFDTKRWINLSADEVKREINNERQSAFHKNADAFVFV

VLSHGCEGAIYGTDNQTVEITRDIIDMFDGNNCPQLIAKPKLFFIQSCQG

KQVTPGVFHSSTINATPSRNREEEADDDNFPVRNSVHEKADVLVYSAAVP

GYKSWRDTDTGSWFIRDLVENINRHAHTLHLSDILVKVNNQTSKRESMYK

RYKQISKLESTLTKLVFFLPET

>c66411_g1_i1OK_Alitta_virens

MDAKPRGRAIILNIENFKRVQKDSPKLTNRKGTYCDRENLRNVFTALGFE

TIVWNNFSAEKIKKKIDEERRLDFHKTANAFFFFVLSHGRKGIVYGSDGK

PVLIEEMLGLFDGKNCPQLVDKPKLFFFQACQGDKTTKGVAKSEERDEKK

ESVPLDNSTVSVLAIFNEDQKEAVPLDSSFIHEKADFLVCYSTVSGYSSR

RNKKLGSWFIRFLVMNINEYAHSHHICDILTKVNNQLHEREKEGCKHISQ

YTSTLRKSLFFLPKS

>c80567_g4_i3OK_Alitta_virens

MAHKRAGMSHKHRECLRMNYTFLLKEIMVDDLCDSLREDGVVKEDWIDDI

RSRVMMMSMKDRVGDLLEKVKCSGPKAWDGLLSSLENTNQKRIAWRLRES

ERLYEENQSIRPQELKTHFENTPTPLECLPTTPEDTAMRPSLSSVASINS

AYFNDLQRESNNDHYKMESNPRGCVTIINNYEFEEQTDLKERLGSKKDRI

RIKTMFENFGFAVQYWDNLKAEHMKSVLKNCASPEDDVDCVVIAILSHGN

GDSVYGVDGILLKYEEIIQPFKANNSTKLKGKPKLFILQACRGRDYRMGV

STDGPGGNDSKAGPSLNLQLDPDPGDEQEEDLVQQVANWNLNSEEMKQNI

HPDMDMVLAYSTLPDHLSHRDDIKGSWFVQALCNVFTERAKTTDVVNMLA

FTNGEISKMTRNESVAQCPQPTSTLRKIWNLNPT

>c85433_g3_i1OK_Alitta_virens

MNETDREKLRSHADLLNANLVVDYSFISVLQEMGIWYDFRDRSRSSDNEA

KSKKFLLILPNLGPHAFSAFIFTLVKTNQEVLANILCPESVEWWKTNNSF

IQKSSAEASSSAESFTFETGDTVVSLDTFRMKSVILQPSDSEEVEYMASS

IEINLQAADPKLVQEVVSNPKFFYRMCAKPRGKAIILNNEVFYDNDKHEE

RYGSSFDVANLTKLFDQLDFKVELWQNLTAKEIENKVDEERKSEFHADAN

AFVFIVLSHGDDTLIYGTDGKIIQTQKCIIDKFDGKNCPKLIDKAKIFFI

QACRGRIVTNGVLVADDAFIKIESGGPAILHEKADMLVFHSTVPGYKSWR

DLDLGSWFMRDLVININKYAHVFHLVDILTLVNNRISERQERYRGFVQIS

TFESRLRKFLFLLPKSNDTDLNKV

>c81751_g3_i4OK_Alitta_virens

MLMNPKKFYRMSAIPRGRAIILNNETFEDGRRFSKRNGSYYDVANLSDLF

TGLEFQIDIWSNCTAQQIKAKVEDERREQFHSRVDAFVFISMSHGGLGVI

YGTDGIKVHIERDIIDKFDGENCSQLIGKPKLFFIQSCRGDEATVGSDAA

SDSVQDEGSDSTVTDGPPVIPSRIHEKADVLAFYATVPGYVSYRNKHVGS

WFVRDLIVSINEHAHDQDIGHIATQVNNKMNERKSDGSGQCQTSKLEHTL

TKFLYLLPKYADS

>c70299_g2_i1OK_Alitta_virens

MIERFSQMSTKKPDEKAVQQACLRLNTSWLNQEIQMEDLCLKLLENGVLE

EEDSSSILEKKRGAKLVELLKILKKSNKSGTWDGLLRSLEKNKKDVALRL

KVTACIYEAYSAVTSEDLEKYFNTYPQRTGPDTVDLTSQGRNITINLQKT

TLEVLQKVRNSEEYYEMSAKPRGRAIILNNEEFDKGKPRDGSSLDVDNLK

RLFEALEFEVVIWNNCTAEKIKENVKTECKHKAHENADAFVFIVLSHGQT

GRITGTDDYPVNITRDIIDWFDGDSCPQLVGKPKVFFIQACQGSRSSTGV

LPDSTETRSQPENICSDANENFTHEKADILVSYSTVPGFKAWRDPQQGSW

FMRDLVINLNMYAHQLHLADILVKVRRQVSKRSNRQGFIQVPKDESTLTK

HLHFLPRT

>c69909_g2_i1OK_Alitta_virens

MAGKGEPGYSGQAEFKADQVDAFGYSYVSIKPPRLSGKSYKANPISLVQE

EYNFTYPKRGTAVIINNRNFNQEKTGQSDREGTDVDAAALESVFIKMGFD

VTRHDNLNRIDMMMTFRRLAAADHSDADCFVCAILTHGEEGVLYATDGVV

IIEKITRLFKPDECPTLAEKPKIFLFQACRGRFYDYGHDISDDQPDAALD

SITVRKIPVEADVLMAYSVCPGFYSWRNNVDGSWFVQAITEVFKEHSDSM

DLLDMMTMVNRKVAYDFASCTDREFTSDMKQMPCMVSTLTRKVYFHPKEE

TPKEP

>c66667_g3_i1OK_Alitta_virens

MAGKTEQESAEFKSDQVDAFGFSYVSIKPKRLSGKSYKANPISIVQEEYN

FTYKQRGTAVIINNRNFDYEKTGQNDRDGTDVDASALESVFMKMGFDVVR

HDNLNRMDMMLTFRRLAATDHSDNDCFVCAILSHGEEGVLYATDGVVYIE

KITRLFKPDECPTLAEKPKIFMFQACRGRLYDYGHNVDDEPDAAFDPVSI

RKIPVEADVMMAYSVCPGYYSWRNNVDGSWFVQAISEVFREHSDSMDILD

MMTLVNRKVAYDFASCTDREFTSD

>c70299_g2_i5OK_Alitta_virens

MSAKPRGRAIILNNEEFDKGKPRDGSSLDVDNLKRLFEALEFEVVIWNNC

TAEKIKENVKTECKHKAHENADAFVFIVLSHGQTGRITGTDDYPVNITRD

IIDWFDGDSCPQLVGKPKVFFIQACQGSRSSTGVLPDSTETRSQPENICS

DANENFTHEKADILVSYSTVPGFKAWRDPQQGSWFMRDLVINLNMYAHQL

HLADILVKVRRQVSKRSNRQGFIQVPKDESTLTKHLHFLPRT

>c80567_g4_i2OK_Alitta_virens

MMMSMKDRVGDLLEKVKCSGPKAWDGLLSSLENTNQKRIAWRLRESERLY

EENQSIRPQELKTHFENTPTPLECLPTTPEDTAMRPSLSSVASINSAYFN

DLQRESNNDHYKMESNPRGCVTIINNYEFEEQTDLKERLGSKKDRIRIKT

MFENFGFAVQYWDNLKAEHMKSVLKNCASPEDDVDCVVIAILSHGNGDSV

YGVDGILLKYEEIIQPFKANNSTKLKGKPKLFILQACRGRDYRMGVSTDG

PGGNDSKAGPSLNLQLDPDPGDEQEEDLVQQVANWNLNSEEMKQNIHPDM

DMVLAYSTLPDHLSMIDDFKGGWFVQALCNVFTERAKTTDVLNMLSFTNG

EISKMTRNESVAQCPQPKHTLRKIWHL

>c78337_g4_i1DEDDEDOK_Alitta_virens

MGDKSNGLSNLQNRIKAHIFGNSVESSVDKVPPVTGPTSCDGGGYRKGPS

SNDSSYLEFTAPVLQSIDEDLKKRDVEDLKFLCTELIAEAKLEKCARGLD

IFDELFYQGYITRQDGSLLIELLVYVGKMKKVRQLGIDPHRFRQSLLMYG

GKLPAYRVLLWKFCEEVGKDEFSTLKYLTKEFGISKQELEGAESMTGLFK

TLEERGILAPLDTVVLIDMAMCLKRQDLIDLVAQYQANPGLPPQYNRSKG

GNSGGIANPSLNIPHRPGMNFPFDRTDAGSSLATGPHNTAEPPPYSPHPS

WEEESLSGAGAVHNFYREKQMAMPNFNQPLPIPRPNQQVLDMPQYKMDSN

PRGYAVIINNRDFYHVPSDPASKKFSDREGTEVDVRNLEYIWKKLKFDVK

TYTNLSDHEIYRLMVEYGLTPENPNGKRKDHTNKDCFVCCILSHGTIGGV

YGVNGKLIQIKDITNNFKPSICTGLVDKPKLFFIQACLGTDKQKGAGLGN

IQEDSGEGFGVVETIPNEADFLLGYSTTPGFVSYRSKDQGSWYITKLVQM

LEEYAHKYDLMSILVRVNEEVGKAVAQVESSEYKQSPAPLVTLRKQVVFH

>c81829_g1_i3OK_Alitta_virens

MYYKMLSCRAIILNNEKFERKEHKDRKGSSVDLNNTRRLFEKYGYDVDVW

QDLTADKIKENIKKENENLKSKKFDGFVFVIMSHGSRGYILGIDGKRVNI

ENDITEPFDGKNCEVLIGKPKIFLIQACQGRRLSEGVKYCNEPDDEGEAL

EEPDCWVLTHEKADMIEFHSTYPGFKALRNTIYGSHFIRSIVRNVYEHGH

EEDFYDIMTKVKCDVAALVIPFIDKPEKGIEIPMKLKQMPSSQNDTLMRK

FYFPLFRN

>c76815_g1_i1CARDOK_Alitta_virens

MENTFSMSKADRHTLDVNLPLLTKDLMIDKMFLATLKQRGTLNDERIEDI

EYYSKRQDMNAAFLKIIRSRGPKAFESFIFALRETNQDDLAQALTGSVQA

TVPNKQATGVEGGGAEGGPGFPMHEKFSQMKLETKTPGEAPTVDERDASE

RLSSIPDIPEEPDIPIDLKPINPELLKNPKKYYQMSRCRAIIMINEKFEG

GKNREGSAVDKRNLMRLFDALGYDCEDWDDCESRDIERKISEERKRLESC

ETDGLIFVACSHGKQDFITGTDGKRVHIQQQIIDQFDGKNCSSLISKPKI

FIFQCCRGGGITEGVEFVPGNEADSFEDDDFSVTVHEKSDIFEIHSTYDG

SKSYRKSDSGSHFIRCLVKNIVKYAHEEDLASIIPWITQDLEAIDIRAPI

TDESGIGRFKNVKQIPVYKSVPKKKFYFLQLKHEKNS

>c50262_g1_i1OK_Alitta_virens

MDNRDWEDEYDFSHAERGKAIIINNIQFEDYTDQPYRKGAASDSENLGDT

FALLGFDVKVYTNLTTTEMLSVLQQAAKEDHSNKDCFVCAISSHGSEDLQ

KKSRWSSKTETSDVIFGTDGVVHLKLITEMFTEKKCPSLARKPKLFFIQA

CRGDMLAENVEVADGVMEDEIDWGGYHVVTQDIPIYRDYLLAYSAPPGFF

SFRRGIGSWFVISLCNVLQTCHKDNDLLTMLSKVNHQVAVEFSVNTPRDK

RTHGRHIVPWITSMLTKDLYFRPKTDEGKRIEFV

>c68898_g2_i2OK_Alitta_virens

MDNEAKDDEYDFSHAERGKAIIINNIQFEDFTRQPYRKGATSDSENLGDT

FALLGFDVKVYTNLTTTEMLSVLQQAAKEDHTNRDCFVCAISSHGSDDFQ

KKSRWSSKTETSDVIYGTDGVVHLKLITEMFTEKKCQSLAKKPKMFFIQA

CRGDMLAENVEVTDGVMEDEVDWGGYHVVVQDMPIYRDYLLAYSAPPGFF

SFRRGIGSWFVISLCNVLQTCHKNTDLLTMLSKVNHQVAVEFSVNTPKDR

RTHGRHIVPWIASMLTKDLYFRPKT

>c80075_g1_i1OK_Alitta_virens

MDPENEYHFNHQERGVAIIVNIMQFDNRQIEPRPHAAVDSVTLEETFKML

GCEVRSFVDLRRAEIINELINVATEDHSGRDCFIFAISTHGDEQFKSEEL

RQLRLDVLAARDMEINLRDITEIFTEKNCPSLKGKPKLFFIQSCRGEKVA

KNVEIEMEDEEMEDEDSVDFGWPRTQVVQNIPIMRDCLLAWSAPPGFFSF

KRKSHGSWFFTAMRRVFKERHHNTDLLTMLVEVNHSVAREFESNSPRSGN

LHQRHIIPWIASMLTKDLYFRPKTP

>c71693_g2_i1OK_Alitta_virens

MLVNPFKGDRCPSLIGKPKIFLIQACRGNKLDHGVEKTDADSEMEEVPRQ

RIPAEADIVMAFSTVPGYFSWRNSTKGSWFVQALCKVLERHGKEREILWI

LTRTSYLVAYEFESNASKEYMNQKKQIPCIMSMLTKELYFRPKKN

>c84172_g2_i5OK_Alitta_virens

MLLRYLDELVTPWFLVVIQDLRMSSEEYGYKFNNLRGMAVIIVNETFQGM

DMRQGARNDLKMATKVFGSLGFEVKAYLDQTASQMLERIKTAVTEKDIMN

LDCFVCVISSHGDPIQQTINEKSTWTDWLWGIDQPIEVGTLLKPTTIAEC

PQLANKPKLFFIQACRGGRASYVVNVDEGKEYPPVVSPDWTVEAVVQNIP

IQKDFLIAWAVQPGFLAFHRPSGSWFIKGLCKILGQYAEKCEELLKKHQK

EKISKPLARYMHENLEIYDIVTLMTMVNGYVGKGKRFISNMPNDDYLDYK

KAMPWVAHQLIQEVHLYKLLIPKYKLSSS

>c76002_g2_i1_Alitta_virens

MAEDQYVFKNKGLAVIIVNENFQEDDLVSRSCATKDLEMSSKTFEDLGFE

VRLYHNQTATQMLYRIIEASKDAKKMDCFVCVISSHGDSEDWIWGIDEPI

KLDTLIKPVTREECRDLANKPKLFFIQACRGGRTSNAIRIEAGKPMVEGK

EPDKHPHVVPNMPIQKDFLLAWSVQPGFVAFQRPSGSWFIKGLCKILDDY

LNEYKQLKQEHQSKTTEETLEDFLNENLEMYDLLTLLTMVNRYVAIDFQT

NIPKDPAFHKKKAMPWIAHQLVKDIYLHKLLLLNEQTA

>c76205_g1_i1DEDDEDOKP20_Alitta_virens

MGDKSSGGSDLHNRIKANIFGSSVDREPEVTPTACDGGGYRKCPPPAPSN

YLEFTAPVLQGIDEDLKKRDVEDLKFLCSELISETQLEKCTRGLDIFGEL

QYQGYISRQDGALLIELLVYIGKLKKVRQLGMDPYKFRQSLLMYGGKLPA

YRVLLWKFCEEVGKDEFSTLKYFAKEFGVSKQELESAESMTSLFKTMEER

GILAPLETEVMMDMAMCLKRQDLVDLVAQYQANPGLPPTYTRADNLYNGG

LTGPPPAAGGGKPLPYYEITDAGSPLINSPFNSNQPPPYSPQPLSEEETL

SGAGAVHNFYREKQNLMMMMPDYPLPTPRTVPQVVDLPQYKMDSIPRGYA

VIINNRDFYHVPSDPASKKFLDREGTEVDVRNLEYIWKKLKFDVKTYSNL

TDSEIYRLMVEYGMTPENPKGNRKDHSNKDCFVCCILSHGTIGGIYGVNG

KLVPIKDVTNTFKPGMCPGLIDKPKLFF

>c48166_g1_i1OK_Alitta_virens

MSSDEEEYEIVKRRAMAVIIVNENFVQPEDKREGAKKDLEMTSETFKRLG

FEVKDYFDQTATQMLHCIQKALQDKDIKQLDCFVCVISSHGYLETEDMGD

KHVRNDWICGTDTEIKLDTLLKPATLEECPGLENKPKLFFIQGCRGGRAS

YQVQVQPGTGGGKGKPPIVDQWAVGATVQNIPLQKDFLLAWAVQPGFIAF

HRRSGSWFIKGICKVLNAYADKGEELLQKHQQEKISKPVDRYMHEHMQEY

DIVTLMTMINGYVATGKEFVVNKSGDPYLHAKKMMPWMANQLTKNFHLYE

LFLPK

>c70725_g1_i1OK_Alitta_virens

MENMAQPYKMTNKSRGIALILSMRTTQQGQRDYAKFDEAKLFTAFSEIGF

TVRRECDLRKQDIITTLKNVSKENHKDHDCFVCCMLSCGDYGVIEGTDGP

VQVDEFLEFFKGDVCSDLAGKPKIFIFQACNPILYKKETTDSETSTDAAI

NTYTIPSEADFLMVWSLVPGILTDAEKDKGCAFITSFCKRLEAGRGKDEI

NRILTRALLDTNNQLEKCAPMTFKKANYRLPCVINMLTKPLYLFSDI

>c120789_g1_i1OK_Alitta_virens

MFTEKKCPSLARKPKLFFIQACRGDMLAENVEVADGVMEDEIDWGGYHVV

TQDIPIYRDYLLAYSAPPGFFSFRRGIGSWFVISLCNVLQTCHKDNDLLT

MLSKVNHQVAVEFSVNTPRDKRTHG

>UCK81519.1 caspase-8-like protein [Arenicola marina]DEDDED

MEPHRQFNASSVANTGHDAVPESRPFRVMLKTISDALSKYDIAALKFLCSEIIPSSKLESLHNGLGLLGQ

LEDRRYITPLQCQFLAELLFRIGRRDLVKRAGFNLAAVKTTIENGTSQHLTPYRVLLHTLSEQLSTSDVE

TLKFMHRDKLDKLTLEKTVDSRQWLTALEQRALIAPDRLAFLDQCFRDVGGDRHAALIQRYNEELVLHSM

CRSDLSELPTSNNTSLTNSFPYGDVARERALPSVTSAVATSGEELVLTSTCRSDLSALPTSNNTSLTNSF

PYGDVARERALASVTSAVATTRESPESRPRGHQERRDVPPCVDNQSGGVISVIPGECPMTPDMFHHLPAP

MLVDPGLGVPRMPGPDVCDRDDPDNPCYNMTATPRGICLIINNEQFQSDPNDRDSKSLSERTGSTADQEN

LENAFGALRFSVHVERNLTAEDIQRLMTYLAHRVDHARYECLVCCILTHGNTGLVYGSDGKHVTIRDLLN

PFKGSSCSSLLGKPKVFFIQACQGKIKQEAANPLVQTDTPGGAGDDLELPNEADFLLGYSTVPGYVSFRH

AEYGTWFIWFLCQNLTKYAHKYHLLDILTMVNRAVARERFTQDDSDHVYKQMPAPFSTLRKRLFLA

**Supplementary File 2: Caspase repertoire of brachiopod *Lingula anatina*.**

>XP_013414814_1LingulaCARD

MDKFHRDQIRKNRLALIQDMEPGPVADYLVQEGVLTDAMLEDIEAARTRRDKVSKILDDI

PRRGPGAFEHFYNGLIETEQPHLADLIWPDRLSQTRAKSAGRRSLIAQVSEDRDDEDDDD

KLPEKFPTADIADFSVIDKLIDVKDPELKALLTNPKESTYKNFSQPKGRAVIINNVLFGG

DKDRKGSEKDATQLQRLFEKLNYKVVLHSNLTAEKMLGMLKEESKNSDHARHHSFILIIL

SHGANDVVYGTDWKKLSIPVITDIFNGKNCPFLVDKPKLFFIQACQGENYDKGSGDTQAD

GGDLAEYFSRLELAGRQEERTDSSDVSVPTMADILVAMATTPDYVSWRNESLGTWFIIAL

VYAISKYAHKYDISQILTKVNYLVSKGETIKGKYRQVSQWRSSLRKKFYFFPGLHQS

>XP_013402632_1Lingula

MLKKESKNSDHAARHSFILIILSHGTNDAVYGTDMKKLSIPAITDLFNGKNCPFLVDKPK

LFFIQACQGKNFDKGSGDTKADGDLAEYFSRLEMAGRPKERTDSSDVSVPTMADILVAMA

TTPDYVSWRNESLGTWFIIALVYAISKYAHKDDISQILTKVNNLVSKGETAREKFRQVSQ

WRSSLRKGFYFFPGLHQS

>XP_013416922_1LingulaCARD

HYSVGLQSHLLKYVVIAVRFNNPIKTLNFVFQAERTRRNKVAKILDVVPRRGPKAFEHFY

YALIVTGQGHLADLIWPERVTQTHAETASRKFVITHEQASTNQDDEDDDNLPEKFPTPDI

ADTTVIENLKDVQDPKLRALLTNPKEDTYQNFSKPKGRAVIINNVLFGGDKDRKGSEKDA

TQLQRLFEKLNYKVVTHSDLTAEKMLVTLKEESKLSDHEAYHSFVLIILSHGAKGVVYGT

DWNELSLPVIIELFNGTNCPFLVNKPKLFFIQACQGENADPSSEDTKADGDFFGYLNNLE

LARREEQTDGSGVSVRARADILVAMATVPNYVSWHKESLGTWFIIALVYIISRFAHKYEI

CKLLTKVNHLVSKAETKDKCNQVARFSTSLRKDFYFFPGLHQS

>XP_023932957_1LingulaCARDp20

MEPGPVADYLYQEGVLTYPVFERIKETRSRSEKVDTILDVVPRSGPEAFEHFYNALIVAG

QQFLADLIWPDRVSQTHAETASRRSMHAQASQKRDDEDDDKLPENFPTPEIADFSVINSL

KDVQDPELRDLLTNPKEPKGRAVIINNVKFDGDERRVGSDKDATRLELLFEKLKYNVVVH

SNLTAEEMRGKLKEESENLENAAHHSFILIILSHGANGAVYGTDWKELSIPAITDIFNGN

NCSFLVHKPKLFFIQACQGANVAKKSEDTEADFKFLEYLSKLEEVGNAVETDDSDVSVRA

MADILIARTTVPGIGI

>XP_013405490_1LingulaCARD

MADEVDRDRIRRHRVALTKNMVEPKEVLVYLNQWGVFTNTMKESTEELGSKREMVNKILD

DTPKRGKHAVRMLYDALLMTEQTELADLMYPELKEDRLAGKLNVTGQKSEVVQSQSAASA

FRRQMSNPPEEGDSEEEELPPDFPTEETVKPDIYKDLKEVTSRRIKNFATKPNKAYTNTS

YPRGIALIINNVKFSDTKLSQRTGSDLDAQKLEKLFHKLDYNVQTKTNLSAAAIKSTLQE

VASRQEHRSHHSFILVILSHGAKGKIYGEDGNHVEIEEILKQFNGINCPLLIDKPKLFFI

QACQGEKTDGPGTQTKKENPDGERDAIEEITMQLENSRIQERPDAVNASVPTMADILLAM

ATTADYVSWRNDCKGTWFIQALAYVISKYGHRDSIASLMTKVNNLVAKAKTNQGQFKQVS

ECRSSLRKDFYFFPGLRNPQHAFGGGNPVNHQTQGQGQEHATQGQGHFQSQGHVTAGVNM

ATGEGSTVGSRYILTALQPGGDPTNVQSQTVPDERSKFNNQAGTDGYYGGEQYGPSYGPQ

GYGTNQHGSFHNSQPYGPLYPAQQQGPQFISQQFGPHFGSQQYGPHVQPPRYGYSYSTPQ

YGTHFDAPKYGGDYQHVGYIGNPPSGNPPIYYEQHRSTPDPGVPHVGPSLSGGYVPSVYG

PARTGHDNLAHPVEETEAKYVFSG

>XP_023932692_1Lingula

MTSKPRGYALIINNRYFHGSQLTERLGTDVDRDNLQNVFKALNYDVTVFDNMKGREMENA

IRGFSIRDSHRHVDSTIVVLLSHGSENVIYGTDCVPVEIDKLIKFFDGAHCPALRGKPKL

FVIQACRGKNFNRGFETTDGETAPSGRDAGPVPATRTSLNVVGPSMLEQLQEMHLRDTEN

LRHKLPEGADMLVAYATVPGFVSWRNSERGSWFIQELCTVLYKYAHKEDLLSMMTVVNNK

VATAYESACNKKQMPAPVTTLRKKWFFYPRM

>XP_013401916_1Lingula

MTSKPRGQALIINNRYFHGSQLTERLGTDVDRDNLQNVFKALGYNVTVFDNMKGMEMENA

IRGFSMRDSHRHVDSTIVVLLSHGSENVIYGTDCVPVEIDKLIKFFDGAHCPALRGKPKL

FVIQACRGKSLNPGFETTDGETAPSGDAGPVPATRTRQNVVGPSMLERLQAMNVRETENL

RQKLPEGADMLVAYATVPGFVSWRNSERGSWFIQELCTVLYEYAHKEDLLSMMTVVNNKV

ATAYESACNKKQMPAPVTTLRKKWFFYPRM

>XP_013414929_1Lingula

MESAEKPATAMNTVDTSGGPFGVPAGEPKVEVSEPTRRALEGNLEYNFKHVRRGMAIIVN

NRNFEEKTEQGERIGTDVDAGALANLFMDLGFEVTRYDNLSTMEMMSVMRQAAQEDHSDA

DCFACAFLSHGIEGRIYGTNGLISIDLLLSFFKGDKCLSLAGKPKLFFIQACRGKKNDEG

VDIPLDETDSATEPKVRRIPTEADYMVAYSVAPGYYSWRNSVDGSWFVQGLVQVLKEFHQ

EMELLQMMTLVNKVVAEEFASDTGYEYNSGMKQIPTIVSRLTKFVYFRPKYLDYS

>XP_013399774_1Lingula

MDDSEPDLVDANLAHAAGGYPGSQRARLPEEIVENEYNFNHPNRGYAIIINNRKFEHHTE

MPERNGTDLDAAALYQRLSALDFKVELVNNLRREEMRSKLEEVAKRNHSDNDCVVVAILT

HGEEGVVYGVDGWIELEKLVAPFKGDKCLTLVGKPKIFFVQACRGNKLDHGVEVCDAEVV

KDEPDAATVFRVPSEADFLMAYSVVPGYFSWRNSTNGSWFIQSLCKVLEKFGHTLDIMTM

MTRVNRMVAFDYESNANKEYMSKKKQIPCITSMLTKDLYFRPKLPGRH

>XP_013408948_1Lingula

MDDTEDGHGDITDAPLSRRGLRVAPEAGRPQQVSSSGDHDMEYPMTYENRGLAVIFNNRN

FASHTGMAERRGTDQDANGIENRLRALGFDVHRFDNVTCYRMSMTMADVGRRNHSQNDCV

ALVILSHGEEGKVYGTDGIIEVDQLLAPLKGDRCLTLAGKPKLIFIQACRGTKLDGGVEV

ADAEMWEDGRTHVHRIPAEADFLIAYSVVPGFFSWRNSMNGSWFMQALIQVLGDYGQQLD

ILSLMTIVNRKVAYEFESNAASKAMSKKKQVPCITSMLTKLVYFRPKH

>XP_013408945_1Lingula

MASSAGIHSLLAIRLKKEITEDQLELIKHRFTSKIGAGRIENMDTLDLLNELNNRMIISK

GKYNKLRQVFKEVGLEKLNRHLDKAEQRIQNIKSGKANGKNEVKEDLSSSESDEGACGGS

EEQTEESWSGDEILRVAPEAGRPQQVGSSGDHDMEYPMTYENRGLAVIFNNCNFASHTGM

AERRGTDQDANGIENRLRALGFDVNRFDNVTCSRMSMTMAEVGRRNHSQNDCVALVILSH

GEEGKVYGTDGIIEVDQLLAPLKGDRCLTLAGKPKLIFIQACRGTKLDGGVEVADAEMWE

DEPDGRTHVHRIPAEADFLIAYSVVPGFFSWRNSMNGSWFMQALIQVLGDCGQQLDILSL

MTIVNKKVAYEFESNAASKAMSKKKQVLCITSMLTKLVYFRPKH

>XP_013408949_1Lingula

MEDTKDEMILGDHDMKYKMTHKKRGYAIVINNQVFQPQTDKEERRGTKQDAEGIKNRFEA

LGFVVEPYENLTCSQMLRTMGEVGRRNHSKNDCLALVVMSHGDEGIVYGTDGIVELDILL

APLKGDKCPTLAGKPKLIFIQACRGTKLDGGVEVADEAGLDELDGRNHVHKIPAEADFLI

AYSVVPGFFSWRNPTNGSWFMQALIQVLGDYGQQLDILSLMTIVNRKVAYEFESNAASKA

MSKKKQVPCITSMLTKLVYFWPKQ

>XP_013387919_1Lingula

MYSCGNLRNRTGTDADSETLQNLLYQLQYRVQVHHDLSSEQIHRLLERESKRDVHKKYDS

FILIILTHGTKEGMCGTDGNLVPYDVIKQYFNNANCEGLRGKPKIVIIQSCLGKFMDEGV

PLKQLDALPASLPHIEEKWQQISVADGRPALEENKEKDSLLCVTEERRDADLQRGEPLDE

KLPTMADFLVAYATTPDYVAMRDTKTGTWFIQGIASVFAKYACTEDAVSLFGKVNAHIAK

RQTRGLGIKQVSSFESTLLKKVYFFPGR

>XP_013381129_1LingulaDEDDED

MIETDAVQPAQASYFRMLLKKVDEQLVSRDLNAIKFLCRDVLPHSKLEKINRGLALFTEL

EYKGLLSPNNLKLLEECLFRIQRADLLKQMGFSLRDMKNKIEQHGEQLSPFRVLLHDIAE

DVSKSEVESMKFIAQAHVPRYKTENAKDLQTLFVALEQQNLLTPDDLDVLVEMVQCIDRK

DLLTKIDSYKTLSRHGNVIDLKRREPVVVPPKVPMMPKRPEPVLRPNPPPPGRSQQTPQN

TDNNRRQLVGQGDTASEPNIAVHDQEATEARDQFYNPKAGHADAQHVAQRLERITIRDDH

MDLPSYKMDSNPRGMCFIINNEKFYKIEGDQASKKMPDRTGTDVDCGNLHYVFSKLKFTV

KVYKDLTDQEITRTMVDAAYNIDHSTYDCFVCCILSHGQLGSVYGVNGRLVLIKDITNCF

KANLCTSLAGKPKLFFIQACQGKDKQEGSPLEMDSAMPGRQQVEEDAPGEQETIPNEADF

LMGYATIPGFVSYRSRTHGSWYITKLVEMLDRYGERYDLVSIMTRVNEEVGKANAELRDG

RYKQSPAPMFTLRKRIFFT

>XP_013411651_1Lingula

MDQTNWEYDFSNKSRGVAIIINNETFENDAEKNRDGSRRDEKSLYQAFKTLGFKDIRVLR

NLTASAMYHTLFKASRENHLDSDCFACAIFSHGGELQGPPIVDASGDSIVSMHGMVYGTD

NRPIHIKLLLELFRDNKCPTLRGKPKIFLLQACRGYKLDDGVEIEEICDQDYDVCDAPAG

TIDVPDDADMSNPAIQRDIPIFNNFITVYGTPPGFFAFRKKKGSWFIQSLCSVIEKYGST

MGFQHMLTRTAHVMATEYESYNPNKEHLSGKKSVCSVVSKLHKEVYFRPKTIWH

>XP_013403374_2LingulaCARD

MVSEVLLSELMACNLITDYDIEKIKSIPTRYDKAGHILDVVKTRGPEAFEKFQEVLRNTE

QRHLADLLDRPSGKVFYTAVPENQDRLAAMCPEIEYKNNSEPRGRALIINNKRFSVDWLP

ERDDTEVDVAALRKLFGYLHYETDVISNLTADEMMEEIQQESRRDDHKNYDSFILFLLSH

GTEGAVYGTDGYPLPYKYIKYILNNKNCLGLLGKPKMIFIQACQGKILDSGTLLDGQTVQ

QPQQAPPAANLDDKVPDAPLEIPNLPSEPTMSDILEVHATSPGYVSWRRTGKGTWFVQSI

VDIFHREAHKYDVLQMLVNVSAALSQMQTVKGNKQALEIVTSLRKKVFFNITPDAEFQTM

FTELKSLLLFLSPLYTSDYYDKLLIVPLHVYAAAFPENQDRRAAMCPEIEYKNNSEPRGR

ALIINNKRFSVDWLPERDGTEVDVAALRKLFGYLHYETDVISNLTADIF

>XP_013381226_2Lingula

MNLAKKSEETPIITIEKLMEMAQRVGLPDGELQEKFIPYHHRQGDILHFKIDGLENVVII

DPLWLADVFSAIMTPASIHSGCSRGHFLKDELERGVLKEEVFDEYLKEFKMAEKKTEIIR

LMLHFDLMLEISQSVENPLPGHGKRRFIVPSMLVSNNSANETNPQKYLLIAFPHILFPTG

LFHRLIIRLLRKYKAQQYKNNIPVVGFDSASFRLADKQSVLHLAIENKAMKLFITNATGL

PPKTHVYSEATKTVENELTVLRDTYCPRMHFGYSIRLKNDGPNIPINALDVQTDDSICVY

VENSEVSVDLRPYRVWFSAAPENQDRRAAKCPENEYKNDSEPRGRALIINNKIFSVDWLP

ERDGTEVDVVALRKLFRYLHYETDVISNLTADEMKEEIEQESQRDDHKIYDSFILFLLSH

GTEGAVYGTDGYPLPFKYIKAILNNRNCLGLSGKPKMIFIQACQGKILDSGTLLDRQTVQ

QPQQAPPAANLGDEVPDAPHEIPKPSNDPTMSDFLEVHATSPDYVSWRHTGKGTWFVQSI

VDTFYREAHKYDVLQMLVNVNYAISQKQTIKGNKQAPEIVTSLRKKVFFNITPGEEESVQ

DQN

>XP_013395828_1LingulaPARPcatalyticP20P10

MATGGARPKTNSIWGNALRWFTGIQNEVTERQDRRYGREHRANGDYTAQDFLGASTSSYD

LLGGLPDTLSSQLNISRRKKSKYEKFTESLSNPSQSACFTMGVSAEIFCAVTELVQGTWQ

SKKTGRGRDTRNLEHTFIVVKRVEAIKNEGLLERYKLTKRQHRLRQTQGRLKPYYYVDPT

MKKPMTVETESPVLDTLNEDINEQYLFHGTKSANIPSICAMGPDFRIARKGMFGRAIYLS

ESATKADQYADEMDSRRTEDLQILICRVLLGNAYYCEEQQTLRRPPCVCGEKICRDHHET

FDSVIGDTRKLFREFLLFTADQCYPEYIVTYDRIDQFQARAVPQLPVTSLCLRAADPALF

MEEAYKDDVYQNCGLPRGRAVIVNNINFQHHEKRDGSEMDCKALVELMEQMSYTVIVHND

LTAQGIFDVMETESKQPSHWTSDSFIVFILSHGMSNAVYGTDGYSLPYSMMYMHFTTDSC

PALAGKPKMFFIQACQGTKKDSRVAGESVPLTMRQDGGQSDSARARCNESESKFGANMCD

IAVATATTPDYVSWRNKENGSWFINAIVRVFSEEARSKHFLELLTSVNSWVASMKTQGQG

QYQQVSSMETMSFTKRFYFWPRLCIVNGELQELD

>XP_013409672_1Lingula

MSTNREEKVKVWPCTVELFEKKAYDAKRVYKNYTEWRGRAVIINNVSFDNEFSLDRKGTE

VDVEYLKSLFRQLHYLVLKHDNLTAEAIDQLATREARTDHSAYSSFIMVILSHGGENTVI

GTDGYHIGYDQILAHFEASRCPTLNGKPKMFFISACQGIQKAPLFNTEQAPPRPQDRETP

EKADMILVTSTTRGDKSWRSTERGTFFIQCLVKSSRKEHVTPNYLIY

>XP_013395658_1LingulaCARD

MEKTDRERIRRNRTAFIKDMMPDDVIEYLYSMGIFTDSTKEDVQAYRTRTDKVRKILDTV

PMRGKQAIHVFYDALVETEQGHLADLLFPERKEERLAKEAARAPPTSTPCPANTQPVNTQ

PRVTQPVQPEEEDVQLPELPCDAGQSLVFVDSLKFDLSDSVLRLQEKYSQVYKNASANKG

RALIILNIKMEGHDILRTNLTRLLKKLGYKVDHKKNLKAEKIKEVLRNEAQNGEHGDAES

FICIFIGYGKGGILTGKDGKTISLSEVTCFFNGQNCSSLVNKPKLFFAQTCVDNDTDKTE

GETREVMEQLTADLQAATLRDGGEGTVATASVPTMADIFVAASTEYTSSTEASPNFWFLE

ALVYVFCTFAHSDSLNSMLNKINSILSKKRAQLDYQSTSLLKEFYFFPKRPVTSDSDSCQ

QNDAVESATEGRSSLAQASCGDDTAQLVLG

>XP_013416458_1Lingula

MRGKQAIHVFYDALVETEQGHLADLLFPERKEERLAKEAARAPPTSTPCPANTQPVNTQP

RVTKPMQPEEEDVQLPELPCDAGHSLVFVDSLKFDLSDSVLRLQEKYSQVYKNVNANKGR

ALIILNIKMEGHDILRTNLTRLLKKLGYKVDHKKNLKAEKIKEVLRTEAQNVEHGDAESF

ICIFIGYGKGGILTGKDGKTICLSEVTCFFNGQNCASLVNKPKLFFAQTCVDYETNKTEG

ETREVMEQLTADLQAASLRDGGEGKVATDAVPTMADIFVAASTEYTSSTETSPNFWFLEA

LVYVFCTFAHSDSLNSMLTKINRILSRKSAQLDYQSSSLRKEFYFFPKRPVTSDSDSCQQ

NDAVESATGGRSLLAQASCGDDTAQLVLG

>XP_013411650_1Lingula

MDDTETDTPPSRGGFVDLGKDKRPDKQVAHVRAGPENEYPMDYRYMGKAIIFNIMEFKGS

GIKRKDVDAAKLYGKLEDLKFQVSLKNNFSAEEIKDCLARVAKENHQDMACFMCIILTRG

REGLILGKDRSIPVEELLQPFKGNKCPSLAGKPKIFLIQADQDITFEKESADFFPLGTKY

KPEETKKYRIPAEADFLVATSVIPGKWSWDITLPTASHFKKYKNASCFIHHLCEMLEKES

EQEEESERLDLLTLFMRVSQLATCDVQQALGDTHKALFEEYDLIPCVTSTLTKDIIFPIS

D

>XP_013406381_1Lingula

MSAGGGDSTCDVQLHYDFKHGKRGKAIIFNHMEYDPKAKTKERLASICDANMLVETFKKL

GFAEDEIHIYHGIHVQPGSESLIAKANGSEKDTSVAYEERQKAFQAVLKQKKILPENIKF

YKGSKLTQLTSSETLKIIEEAAKDEKYNEDADCIVFAVLSHGKEGPDMSDDSVLQDLIRC

YDDRLHIEKFMQPLRDDTCLSLKGKPKLFFIQACRGTGRDDGVLVKEDAHSGKVSAGPPR

TITSTPVEMRNIPAYNNFLTAYSTPPGFSSSKGSRGSYFIRSLCSKLNEHYNSGMDLIRI

LTRASSHMAKEYSPTRGEGYPALLKKKQVPVIVSKLTKLVVFTAN

>XP_013421206_1Lingula

MGPLQRFISFAKVMSVFPSTLHFAVGLTSMLCSALSSWSFIKQVGIVEEMRNLFFSLFRL

QLFAIYVVIAIFIFKWATELDRIFGAKTNVFILQRKSFKKFCNRDSCYIMAKLVGVSARN

TWPYLKFAAHLESIIALMAWCICSYRYFPISMSLDDLPVMLAENFSVHISVGICFLDSFD

YYQNFFSTILIKGKSILVLTRNFQRVTLSRIPFKLFCFLQEFYFYRPFFDPIFIKRKIRH

LVRFHRKKKASRKGKKRRGAVNRNDGKAMLVNKVKPDVAKLPRQPPKKVPGKTTGIGPNS

QIERSKFDKNKNSSGETNMSTVLTNSSAPVLKYPIMESCVPEIQEDENSEASPHVETPPY

VYPRLGLQVTGNRRTRTCSETSTASSTESAYSITSEGQYRGITLIINNCRYFPATKMHNR

YGTMKDEKHIEDVFSKLKYKVIKFRDYTCSDLKHRLRQLGSKMEAHYDSFVCFVLSHGED

GYFYTVDGERIFLADLISYFYSKNCPQLENKPRLFVSQMCLGGAELNDGGSADNTGKEAV

SKRTKCRSAIMKDAEGEKDCKKSLSTVPAAQEADFLLAFATLPGCAAKRYAATGSPFIKT

FLGCVKKYYNRKGRTRKCVTDILKLTGKRLKKKENQVHCSFLSINKAFYLTIPAMN

**Supplementary File 3: Caspase repertoire of mollusc species which are included in analysis of Figure 3.**

>XP_005113266.2 cell death protein 3, partial [Aplysia californica]CARD

MEKEQKEILITRRVELVENLNMDGGLFTQLMAKKILNQRTVNNIKMKRTPEEQAEELLNY

LPKKDCFERFCEALEADDQEEIVTKYLKVPSKTSNTPAVTAEQPQNSANTVASPTAAPSR

NSSLPAASVETIDAAIQPPSVQPVNSQPTISVIRIKEPDNNHLCIEYQNTSASPQKKMRL

EEITNGSHGYEEDSTRYVNKHFIAQESDRSLSCNNSLSVVPVLNPQSVAHNPNVSDPVRY

MAAYAQQTSQGPASAQLTITGQASLPLVPPPSMSSASPMTNSPAHIPQPFRPDLSTVQPF

PQLSRQSDPLTRFPSNEAGHQYSHPHQYLSQHGLLPPSQKHASVPAATSGQPPPVADYNS

LTHAADAVDTMTIRIHGREISQVDGSTPPQNGQGSELSSVTLNAAAGPAINLSAPPLPMS

SLPAAAASPSPTLTPTLTAPSPVAVPAQQNISIPSNIPSDMEHMDLPDGCVNVKVEHSSR

QFFINNYKKSYPLHKIPRGLFLIINVDEVEGKPPRKGTNFDRDNLMNLLCQLHFEPVVYN

DSDGLSAQEIVIKLQDFARRKEHKRADACVICLLSHGEEGYIFGTDGKRIPLDEIFLLFG

NTNCKELMGKPKVFIIQACRGGILDKGVRMDEPDGGTLPNMPVAQLPTMSDMLIGCPTMA

GYYAWRNRARGSWYIEAIVQVFMKYAKCEDICVMLNRVNQLVSRKVSECNQKEMNQMSQM

SEYKSSLRQPHFYLFPGI

>XP_012945247.1 PREDICTED: caspase-8-like [Aplysia californica]DED

MDLEYRKRLAYIDDGLHKTDCKIFKFLCEPLIGRSVKDDSPMSIFEALEGRRMLSASNMT

VISEALQIIQRYDLLEDIRTDSATTYHGPMVQTSMICPFRATLAEISENLPDEDLNRLRF

IYNKVPKKTKIGDGIDLFKFMMQRGDLKPSSLTALTDIFSSIERTDLVNSLQSFQDQHVQ

RGQYMWQARQALTQAQEETYDDRPPTVESMYDEPENASTGEPVVTRAELLPPICTSGATS

SWETNCNVLGLNTPHSPMTSLNFPQPGPGECSEQLKPSGYQNSVLRLLSEIQHSGNYNNS

ANLNSAQQSSSETGIRFEGEHVTESTVPVTGSSEPLLESSQEADPPLNELGVSLLNIDDA

AVGATASTGVPEGYKPVGNYKMEADPRGLCVIVDIENFTHDPHDPQAKKLNPREGSKYDR

EALTQTFSRLLFKVEVVENPNDRDLLKYLEQVSKRDHSYFDCFVACILTHGGNGFVYGSN

GRRVYLEQVTSYFRGTNCPSLLEKPKLFFLGCCQGTSTQQAVELLCDIEADGPGRNMRFL

PDEADFLIGYATVQDYVSYRSRSQGTWYITELTKILDKYAEIDDLNSILLRVNAAVAEYA

HEKTETNGNTKLLKQVPAPQTTLRFALKFL

>XP_005099863.1 PREDICTED: caspase-3-like [Aplysia californica]

MDMDTVDALGEPTESDVVDARGQPGGGSRVGAAAQKESDFVADMYNMKHKNRGIAVIFNN

KVFHRRLQMGDRQGTDVDCENMYNLLSTMGFDDIRTFHDSTVKEMRQEMQQVSKEDHTQN

DCFVCVILSHGKEGYVFGQDDKVPIDELVLPFKGNNCPSLGGKPKLFFIQACRGTDLDSG

VEVSDAAPDEEESMEEEVVMRRIPTEADFLMAYSVVPGYFAWRNSSRGSWFVQAIYDVFS

EHWEKMDVLSMMTRVNKKVAYDFPSNASKEFMNRKKQIPCITSMLTKDVYFKG

>XP_005092532.1 PREDICTED: caspase-3-like [Aplysia californica]

MSSHQAELPYILRSVSGERRDVRPGAPSEPNEYNFNFKKRGLFLLINNRHFTDDTGQPPR

EGSDVDAERLEERFQDLGFEVRRYDDVTCAKMTSLLYDAAASDHSDHDCFGCAVLSHGRE

GLVYAKDGHVPLDLLVLPFKGDRCPSLIGKPKLFFIQACQSERSGNYLDTSCSSRGRSPV

SPSSSRRSPVSPPSLAGDEDSPELSGIIHMRKIPIEADCLFCYSTVPGYYSWRNNQDGSW

FIQALCIVLENYSTKMELMHMLTHVNRIVAYEFASCTDKEFSENMKTMPCILSMLTRYVY

FRPKKQDIR

>XP_012940622.1 PREDICTED: caspase-6-like isoform X1 [Aplysia californica]CARD

MGDGLTYSRLIAGVLNQHQIEIAKTVDYEELRPYLVQRNVFTTLDLQVAEAKKPYFYTII

HLCEKAKTLDRKGFEDFIDAIAEDHYWLADKIRADYDEKLSASKVNQVKTSCPPGSTSSG

MRLPGLSALQQKAPPHSRSQSANLAARAHDTQAMKPALVSVPRGGGGGSLTSSPQSRPPV

SGSALEVCKEFKGASPRDPDMCLQDPEAERLVRASASSPRSHHSMGGISPVQRGGNNKYA

EKRRKNPNSVYAMDSHPRGFVLIINNEKFDTLPERHWSESDRTKLFHMFKFLGFQVSLEE

NCNTAKMKELLQLFARFDRLKTASTMAVVILTHGGSNDLLFGTDGKVVHGTPIADTYITK

FDLIQAFKPESCPSMNGKPKLFIIQACRGVEDDRPASAQHLSWPSQITTDGADTPSANSV

GAAAVPHRAAASDMCFVHSSTIGYKAYRQKNVGSPFVDTLTKMVTDWARDLKFQDIIQEV

QKSMAQQPLPYDNYTLPELSTHLMQKWYLNPPDV

>XP_012940623.1 PREDICTED: caspase-6-like isoform X2 [Aplysia californica]

MGDGLTYSRLIAGVLNQHQIEIAKTVDYEELRPYLVQRNVFTTLDLQVAEAKKPYFYTII

HLCEKAKTLDRKGFEDFIDAIAEDHYWLADKIRADYDEKLSASKVNVKTSCPPGSTSSGM

RLPGLSALQQKAPPHSRSQSANLAARAHDTQAMKPALVSVPRGGGGGSLTSSPQSRPPVS

GSALEVCKEFKGASPRDPDMCLQDPEAERLVRASASSPRSHHSMGGISPVQRGGNNKYAE

KRRKNPNSVYAMDSHPRGFVLIINNEKFDTLPERHWSESDRTKLFHMFKFLGFQVSLEEN

CNTAKMKELLQLFARFDRLKTASTMAVVILTHGGSNDLLFGTDGKVVHGTPIADTYITKF

DLIQAFKPESCPSMNGKPKLFIIQACRGVEDDRPASAQHLSWPSQITTDGADTPSANSVG

AAAVPHRAAASDMCFVHSSTIGYKAYRQKNVGSPFVDTLTKMVTDWARDLKFQDIIQEVQ

KSMAQQPLPYDNYTLPELSTHLMQKWYLNPPDV

>XP_012944029.1 PREDICTED: caspase-7-like [Aplysia californica]

MAANMEELVSGTQALTVKDESDLDLDSVEGFRKSLKKKISQFACGLGNKEDDENRYSVRS

SQSSSSSSSRDFAENRTAFVTAELKTPAVPVDPKADVYDFTCEKRGLAVIINNEDFSQCQ

DFDDRPGSTYDESSLYHSFQRLGFNVLLHRDLPGWKMVEVLKAAASSYDHRNADCFACAI

LSHGDQTWTEREWDRLGTKTRLDLLFGVDGRAVATKFIVELFSDNYCPGLEGKPRLFFLQ

ACRGSKLDHGQELPVIRPRYTPGIRPRVREMGDGPDRASGGIDLPDISEINSPTSSSVDP

PEFQRKLVDYTDASSDGDAGPSEDSGQLGNTAERTVADDVDAGWGFTREVPVSPTPLYKD

CLVMYATPPGYFAWRRSTGAWFVQSLCTVLNSGMVHRMSLVNLLSRVCGLVAHNFQSQNP

SRPSMHEKKAVPVVESMLVKDVIFTRKQ

>XP_012945247.1 PREDICTED: caspase-8-like [Aplysia californica]DEDDED

MDLEYRKRLAYIDDGLHKTDCKIFKFLCEPLIGRSVKDDSPMSIFEALEGRRMLSASNMT

VISEALQIIQRYDLLEDIRTDSATTYHGPMVQTSMICPFRATLAEISENLPDEDLNRLRF

IYNKVPKKTKIGDGIDLFKFMMQRGDLKPSSLTALTDIFSSIERTDLVNSLQSFQDQHVQ

RGQYMWQARQALTQAQEETYDDRPPTVESMYDEPENASTGEPVVTRAELLPPICTSGATS

SWETNCNVLGLNTPHSPMTSLNFPQPGPGECSEQLKPSGYQNSVLRLLSEIQHSGNYNNS

ANLNSAQQSSSETGIRFEGEHVTESTVPVTGSSEPLLESSQEADPPLNELGVSLLNIDDA

AVGATASTGVPEGYKPVGNYKMEADPRGLCVIVDIENFTHDPHDPQAKKLNPREGSKYDR

EALTQTFSRLLFKVEVVENPNDRDLLKYLEQVSKRDHSYFDCFVACILTHGGNGFVYGSN

GRRVYLEQVTSYFRGTNCPSLLEKPKLFFLGCCQGTSTQQAVELLCDIEADGPGRNMRFL

PDEADFLIGYATVQDYVSYRSRSQGTWYITELTKILDKYAEIDDLNSILLRVNAAVAEYA

HEKTETNGNTKLLKQVPAPQTTLRFALKFL

>XP_012945422.1 PREDICTED: caspase-9-like [Aplysia californica]CARD

MEQHHKELIQTNYVKLSNLLSADVDSVARELWSQKILTDGMLDKVVSNNSDRKKAYALLD

VITRRGPHAFQKLYECSLAVGLEDVADVLRPDLAPHAPPRAPAPPPKARSTPGETSAPTI

AAPESSDEEDLPDVWPPAESPNLMKLKVKKATDLEDPDRRKKWLRHYRKSLTPKGQEGVY

EMKHRKRGSLILINNEKFDHMSDRDGTQKDRLALDSLFRRMNFNFIVHSNKTSQVLLEVL

EEEARKDYSCHDCVVVILLSHGHKGVVFGTDGKIENGKPLNCVEIEKITKLFRQAPSLYG

KPKLFFVQACQGNAKDSGVAPGQAAETESGDSPGKDDKPSPGKPAPSPGPTGPSSGLDTQ

QDSAAGQAGEKSGPPTPKVTGASTAGKTSGPGKDAVDSCDDEGRDPEAEGAVGGTQAEGE

GGESPLDFLSEEEGDAAGEKLPGAGDVFIGMATPPDFLSWRRPDFGTWFIQALVEVFAKW

AYKLELDQLMTKVNQLVTKAATKRGSYKQVGEKRGTLTKSFYFFPGLYEDMEDL

>XP_013088924.1 PREDICTED: caspase-3-like [Biomphalaria glabrata]

MEEARAHRYSMKNPKKGIVLIFNNTNFHSSSLPTRHGSDADEESMRKLFRVMDFEADDIR

IRRDVSVNEMKRELAEVSRLDHTDYNCLVCFILSHGEDGYVHGTDGKMAVTDITSFFQGN

RCPTLNGKPKLFFFQACRGTKFDEGVDYVDSCVPDQVQENLGFAIPQYEPDFLIAYSSVQ

GYFSWRNNAKGSYFIQAIHEVFSNNWGXXFLPQNVHSMYTVIFLPQNVQNDLDSNLVAMT

QLTPVISFCRLNSPQ

>XP_013073999.1 PREDICTED: caspase-3-like [Biomphalaria glabrata]

MSDTTDAKLESFQAGGDQSNVPESLVTTDETDSIGGYRSSGRIASMPESDFLSQHYAVKD

RIGRAVIINNKTFGQRTGMDKRPGTDTDAVSMQTVLTTIGFQVERFDDLTADEIKNKLQQ

VASDDHSSSSSFLCVILTHGEEGYVFGTDDRVSLDDLVAPFKGDKCLSLAGKPKIFLIQA

CRAPEAGYEGDDGMEEEMDYDKSIRRIPTEADFLIALSVVPGCLVWNKSPKANSWFIKAI

HDVFSRNWNKLDLMTMMTRVNKIVADEFETKNKAEFDKRKKQIPCITSMLVKDFYFFKR

>XP_013094562.1 PREDICTED: caspase-3-like [Biomphalaria glabrata]

MKSEKRGKVLVINNAKFETLTERDGSGQDEESIYETFRELKFEVQVLHDKTHEEIESILE

TESKSDFSEFDCFILFVLSHGHKDVVYGTDGYFDEITKQPKNCVQITYLKSKLCTNKSLI

GKPKMFFIQACQGNKKDSGLFINPADQESNLNLPQSESSMSQESDIDLSKNPALSDSGTE

NTSIANSGQSVEDQTKHIMESVLPTEAHDAYGPKLPSQAEVFVALATTAGYLSWRNTMWG

TWFIRAITFVFAKYAYKYDLNKLMTMVNSLVAKAETIKGKYKQVAEKRDTFTKTLYFFPG

LPENSIDHQQTHYNN

>XP_013074008.1 PREDICTED: caspase-3-like [Biomphalaria glabrata]

MQRTVAMPMDSPDAFPSYRRRTQENTESENVALEHQNLIMTEEELLSEKYKMDYPKRGTA

IIINNSNFKRETQKPKRYGSAKDEVCMHELLRKLAFQDIRVKTDRTCAQMKGDLKAVAEV

NHSSYSCFVCVILSYGEEGYVYGTDDKISVDELFEPFKGNNCQQLAGKPKLFIIQACCAT

DVQYREEADAAREKEEEEEDIALIKIPSEADFLTVYSEVPGHFGWESDTKRSWFIQSLSE

AFLDEWENKDVLTILTRVLQKMDIHLETLKGSSNKSIQIPCITSMLTKSWFFK

>XP_013065656.1 PREDICTED: caspase-3-like [Biomphalaria glabrata]

MGSFQDDVSPIMKSCKTKDELDGFRKNIKNKLKRFFCSPSSLSPTSPSTETLEFVPVKTA

LNAAGPSTLNQIKKQDDDDAYSFTSSKRGLAVIINNEVFHEPKQFEKRSGTERDAAALEE

TLTGIGFDVLMENNLTSTQMTCVLTSAAIDYEHADADCFVCVVMSHGDVMWFERGGTFNK

KMERNDVIFGVDGLPVKTKFIIDTFNNWNCPSLKDKPRLFFFQACRGEKLDHGTEIPLLK

DLQISLRTTSSSSVSCGSNNGYLLTNDSLVMKNGKDETDSKPETYNGTKPNQQDILPQFS

SPQLPVPPPSLYEDCLVMYATPPGYASWRREDGSWFIQSLCHVLNSPQVGSGKVSLLRAL

TQVSGLVALKYETYSEDDPSYHEKKQQPVIYSMLIKDVFLKRRVPMGARTINYGVSEKVF

NGCNEELRSSVYV

>XP_013094741.1 PREDICTED: caspase-8-like [Biomphalaria glabrata]DED

MSYFRDGQYRETLALIDSEMHEDDIKNLKFLSEPFIPKKLQKKFTSALELFEALEKSHQL

SENKVEVLVELLHIIGRVDLAMKLLEDAHLLGHHVAHGKYVNPFYVLLFKLSEELNIQEL

QSLSFLYGKVPKREKIVFGIDLFSIMLQHVNLSPTKLNDLESYLANINRNDLVQEIRDFK

SAHEYNILYGDKNKDYQCMMDETMSSPVLNYNSNDTNATHHSDASVIASEEEFEVRPVPE

SQTSQSNQSHESQESQQSYRKEILVCEDENLEVGMENNLALSLNLKHFTQDLSSGPLSVN

DILFEEELEDVTEYYRMKARPRGICLIIDNQRFTFDETDLEATHLGNRMGSEKDCEALKY

TFEEKLHFQVDIRQNLKDFEIAAAVRDVSSLDHSQYDCFVCCILTHGGEGFVFGSNGRKY

SIKEIINCFSAQKCPTLAGKPKLFFLQACRGKRHQAATVVEMDAPEMIPADEDLLPPEAD

FLLGYATVADAVSYRSRSTGTFYISELTKALNTLADKYDLLRILTRVNYEVSRLVYSEGQ

GEQRIVYKQIPAPQYTLRKVVKFL

>XP_013082356.1 PREDICTED: uncharacterized protein LOC106067681 [Biomphalaria glabrata]CARD

MDKAQKDIITANRVELVENITVKDGLYSQLLQKKVLSQRSINRIKGSGVDEQQVEELLNE

LMKRDCFELFCQALIDDTQVDVVKKYLKPPASQTVFTPTPAASEPSPIQTTNQQLVGSLQ

PPTNQQPTGPQQPSSVITNKQLTPSFMNTNENASLPLQPSMVSTDCASGMKRANPSPPQS

ATICIRESDNEHLIVQYVPPCKAARQEEPSQLNNVQGEVPETFEDDETRYHLKHFIADKP

IPMSPYIMNNPQILSNPNIIDPQRYITAFLPPAGRLQGVEHLAARFLSLPTNSVTVIPEG

RNMMDTQSIRKLLQPQNPLLQTQIINNKESSFKRIDQESFPRFNVTVGQLGVLSASKTTV

QETLGPGFNTDDAVDNISMYSISTPTLSTKLSEQPRLNDKTDITRVMTEFNEDKFHLPAS

FEEDMSSMDLPDGCVNVKVEATSKQFFLQNFKKSYPMKQIPRGLALIINVDEVEGKPPRK

GTNFDRDNLCSLLTQLHFNIMCYNDQDGLTALDIVNKLKTFSRLKDHETSDACFICLLSH

GEEGYIFGTDGKRIPLEEIFMLFGNTNCKGLIGKPKIFIIQACRGGMYSQVFKNSF

>XP_013067788.1 PREDICTED: caspase-3-like [Biomphalaria glabrata]

MSTMEARDVTDALPSIRTRASTFPSVAPESDYNSDRYRMNYKNMGMAVIINNKNFHPRTG

MNTRTGTDVDAANMKDLLKTIGFENVVTLNDLTAFQMIDHMRKVALLDHSENSCFVCVIL

THGEEGYVFGTDDKVQVEQMVAPFKGHKCLSLAGKPKIFLIQACRGTELDPGVEVPDGME

IEMEEEVRRIPTEADFLMAYSVVPGYFAWRNSTRGSWFIQAVYDVIKTNWKTMDLLTMMT

RVNKKVAYDFQSNANKEHMNNKKQIPCITSMLTKDVFFYQ

>XP_013088924.1 PREDICTED: caspase-3-like [Biomphalaria glabrata]

MEEARAHRYSMKNPKKGIVLIFNNTNFHSSSLPTRHGSDADEESMRKLFRVMDFEADDIR

IRRDVSVNEMKRELAEVSRLDHTDYNCLVCFILSHGEDGYVHGTDGKMAVTDITSFFQGN

RCPTLNGKPKLFFFQACRGTKFDEGVDYVDSCVPDQVQENLGFAIPQYEPDFLIAYSSVQ

GYFSWRNNAKGSYFIQAIHEVFSNNWGXXFLPQNVHSMYTVIFLPQNVQNDLDSNLVAMT

QLTPVISFCRLNSPQ

>KAH9490831_1 KAH9490831.1 Caspase-2 [Bulinus truncatus]CARD

MFAMYSNVYSKIAIVSETCKGLAKCETVNHSNSCRRSVKRTMDKTQKEIITANRVELVEN

ITVKDGLYSQLIQKKVLSQRTVNRIKGAGVDEQQVEELLNELMKRDCFDLFCQALIDDTQ

VDVVKKYLKPPPVQTASPTPVAASDPTSVQTSGQQPAQPQLEPLVTTDCASGMKRVSASP

PPTAVLCIRENDNEHLILQYAQPPKKIARQEDPVIFENDETRYHLKHYVSEDVILEPTHI

LNNSQTLSNPNIIDPQRYAAAFVSSPARLKGAEQSELTLGITEAKKVMDSQSIRNLLRPH

NTFHISNITQLASNNKNRFMRIGQESSSPNSKISIGQMGVLSSQTSRGVQETLGPGSNCD

SVDNVSTLSYVIPMQQTQLSNIERLLLKDNQASEITRVMPDFFSDQCQVLAGFEEDMSSI

DLPDGCVNVKVEHSSKQFFINNVKKSYPMQKIPRGLALIINVDEVEGKPPRKGTNFDRDN

LCSLLTQLHFNILVYNDKDGLTALDIVNKLKAFSKLKEHAMCDACFICLLSHGEEGYIFG

TDGKRIPLEEIFMLFGNTNCKGLIGKPKIFIIQACRGGYLDKGVQLDEPDGGPVASKEGT

QVPSMSDMLICYPTQTGYYAWRNRARGSWYIEAIVQVFMRFAKNEDICAMLNRVNYLVSR

KVSECNQKEMNSMSQMSEYKSTLRRPHLFFFPGIGVSVNQYVKYVVRGPRNMSSDGPEIC

HQTAQKYVIRRPRNMSSTAQKYVVRRSRNVSSDGPEICHQTAQKYIIRRPRNMSTEKCVI

RRP

>KAH9499438_1 KAH9499438.1 Caspase-2 [Bulinus truncatus]

MWEDIIEENKGASKRSRKLLQILLSRGPNAFQKLYESVISEELFDAADTLMPENMPHWKS

DTPDDVTYLADFIKNPNDTMVLHNPLSKQNGELDVLDRLQIKDIKDLLNKDFERGAKKYD

TGSFSSQTQVKELETGPSQTQVKELETGPSQTQVKELETGPSQTQVKELETGPSQTQVKE

LETGPSQTQVKELETGPSQTQVKELETGPSQAQVKELETGPSQTQVKELETGPSQAQVKE

LETGPSQTQVKELETGPSQTQVKELETGPSQTQVKELETGPFSSQMISKNTLSTSTQNPA

ELSEQNQIHRKENVTHLKSPVPTVDLTPAQPLVNTSENLLTELHVLPKSPQVEEDWITKK

NRSIKLVDMNSEMAQKFKQLMCNPDGSIYVMKNKLRGNAVIIDNIHFDHKTPREGSEKDT

ESLENLFSTLSFNARIHTDLTSRAMLQVLEDECQMIEEKHECFILAILSHGTNGFIYGTD

GDKALENAISVERIRDIVCKVKNLINKPKLFFIQACRGDVKDPGLEEADNGKVPTHADCV

FSMSTTSAMASWRDRNHGTWFIQAVAKIFSEHAHNCDINKLMTHVNNLVSQRETSERVKQ

VSEFSNTLRKTLYFFPGLNEENIATMVS

>KAH9513478_1 KAH9513478.1 Caspase-7 [Bulinus truncatus]CARD

MEQKDKDIIRRNYSDLASAIKPLIQTVTKDLFLKNILTERMYEEIIEAHRTATERAEALL

DRIPRRGPTAFNELYNSVLKCELYEAANILKPDINHWQLQITEPTEGHNSEKAVNVSNDM

EMDNGDDDDDIVLPDRWPLDAPTTEPIVQNVVENSNIFKIYKKSLGSQRHYKMKNSPRGR

ALIINNLKFQSLSDRQGSELDESGLISLFKQLDFDVVVENDKTCAEMKQILKNESERDHS

KFNCFVLVILSHGFKGGVFGTDGKVERESHYNYISLKGIKDMFDVTKTLIDKPKLFFIQA

CRGVNKNEGRIKPVEVSDGDDLTLAEELFTEVNETLESLIPTEEADDSTKVPHHADFYIA

MSTTSDMASIRNTLVGTWFMQALIYVFANYASRCNIHELMQLVNQLVSKAESKKGLKQVA

EMRHTLTKELYFFPGLTHS

>KAH9499437_1 KAH9499437.1 Cell death protein 3 [Bulinus truncatus]CARD

MQLAENIMYRIEIICKNLWSAKVISKHMVEDIMEGQNTDRKRAMALLDLLPSRGPYAFDL

FFKLLEENELFEAAYILKPEQRTIHANLSISFAAQPSERLQQHNVLGNVEQIDDELPDVW

PSAEALGSIEVRKITKNDTEIKKLYEYSLQPKGQTEFYTMKSTKKGKVLVINNEKFEELP

SRNGSSKDRSSIDKVFQSMNFDVIVKNNLTSQKMKDLLDNESKVDFSGIDCYVLIVLSHG

HDGVIFGTDGRTDEEANKSINCVEIKYVRDKICSNISLIGKPKIFFIQACQGKNKDFGLY

VSPADRQLVGSSARNETSDNQDGDIDLSKYDTKVDDLREKHLETQNEDATVDQSKIIEDV

VPVEATDAVGPKLPSQADIFVALATTAGYLSWRNTALGTWFIQAITYIFAKYAYKYDLNK

LMTMVNCLVAKAETIDECFMRVFSNNCLSCMKPSCQENLLNFS

>KAH9518589_1 KAH9518589.1 Caspase-7 [Bulinus truncatus]

MSAETANMDRGDQTDAFGSSKSTASIVAALDSDYISDKYKMNYKHLGTALIINNKNFHPR

TGMNTRTGTDVDAVNMSGLFKTMGFENVITLHDLTVVQMRDQLRKVASLDHSDSSCFMCV

ILSHGEEGYVFGTDEKISIDQLVFPFKGHRCKSLAGKPKIFFIQACRGTELDPGVEILEE

METEMDEEVRRIPTEADFLMAYSVVPGFFAWRNSTRGSWFIQAVYDIFIDNWKSMDLLTM

MTRVNKKVAYDFQSNASKEHMNNKKQIPCITSMLTKDVYFYRD

>KAH9518588_1 KAH9518588.1 Caspase-7 [Bulinus truncatus]

MLSETKDEPDAKPEKRDQFSAANEVDFVDAKRSAEKTPTMTESEFLSQHYVVKDFIGHAV

VINNKKFHQKTGMSDRIGSDIDAQKMKSLLTNIGFTVETYDNLKSFEMKDQLQNAASIDH

TNSSCFVCVILSHGEEGYVFAHDDRVALDELVSPFKGDKCPSLVGKPKIFLIQACRDPEI

DYECADSAGMEPESDPGQMRRIPTEADFLIALSVVPGCLAWKNSTQKGSWFIQAIFDVFS

KNWTKLDLMTMMTRVNRIVADEFEKKAEIEFNRRKKQIPCITSMLTKDFYFYRR

>KAH9500511_1 KAH9500511.1 Caspase-3 [Bulinus truncatus]

MSDFSSQRYTSKNPRKGIVLIINNKRFDSEDLPERNGSDEDVANLQRLFRIMDFESEDVM

VRNNVSVNSMRQELANISKLNHRDYNCFICFILSHGDDGFVYGTDRRLAVSDITRYFQDI

QCPTLIGKPKLFFIQACRGTKFDDGVDYVDSAAVDQGEEEVDLVVPAYEPDLFIAYSAVK

GYFSWRHSTNGSYFIQAVHQIFSRYWGELDLHRLMTRVHWVMANEFESSSSIEAMEGKKQ

MPCYVSTLTKDFYF

>KAH9518585_1 KAH9518585.1 Caspase-3 [Bulinus truncatus]

MSNNLDYIDAIGDSGNSSAQPNRRVYSSLTEDEFFSDTYKMDYANRGIAVIFNNYDFIQE

TNLPKRTGSEKDEDSIYYLMNTLGFEQIKIKRNRTCPQMEADLKEVADEDHSTNSCFACI

ILSYGEEGYVYGTDNKIALESLFDPFKGNNCLTLVGKPKIFIIQACPMRIMLYREESDSG

VMEEEFEELEEEVGKIPTEADFLVVYSDVPGSYGWSKKENSWFIQSICEAFETYWSKKDV

LTIMTAVLRKMEEHFEKSKRQALPIPCITSMLTKSLYFKQ

>KAH9505269_1 KAH9505269.1 Caspase-3 [Bulinus truncatus]

MPSCQPTPCSVVSRPHAQLSADPMPSSLECGARCGERRDVRPGAPSEPNEYNFNFKKRGI

FIIINNKHFDEETGLLAREGSDVDAERLEERFQDLGFEVRRYDDVMSAKMTSLLHDVASL

DHSDYDCFGCAILSHGREGVVYARDGGVSLDLLVLPFKADKCPSLIGKPKLFFIQACESR

RHGMYLDKPGRTRTTRSSSPASPSSSRRSPVSPSLTGDEDTLDMSPSVVLHMRKIPIEAD

CLFCYSTVPGYYSWRNNQDGSWFIQALCIVLENYGNKMELMHMLTHVNRIVAYEFASCSD

KEFTDSVKQMPCIVSMLTRYIYFRPKKQDIR

>KAH9513084_1 KAH9513084.1 Caspase-3 [Bulinus truncatus]

MVIEDFPAKTMEYEKMDGSYLNELDGASKFVKSLLPGLKCLANKPQSPTDSVNEFMKGET

AFITKQRCESGVNVCSPTKEKFLRFETAIINKPCAEPEVKACTTTEEKFMKVESAVVTQP

HTEPGVKVCSTTEDDDNTYTFTHGRRGLAVIINNVDFSKAGGFGDRAGSDVDAANLRESL

TELGFDILCLRNVSAVQMVNQLLLAARKYEHDDADCFVCVVLSHGEESYTQKSDINNLPH

HRTERKDLLFGVDGKSVPTKYVVEMFNDDQCPGLKDKPRLFFFQACRGGKVDDGATVNVV

PNVVATSCVHHHHRAEHDSHSSNTDSTQYLPLQMAGLQMCQPTNLPQESVGSIDSVKAVE

PVLVNPYPDTLYKDCLLMHATPPGHLSWRDPVEGSWFIQSLCQVLNCNELKSGRMTLSKA

LTLVCRLVAFGYESFNPTDRSYHRKKEEPVIYSMLVKDVVMKTKDKQSMFFT

>KAH9498393_1 KAH9498393.1 Caspase-7 [Bulinus truncatus]

MFGWIQGFKNKKKSEMNESLLEKIVDCDISLSLSWNTMDDENRYKFNHPKRGLAVVINNE

EFGNIKNLGDRHGSSQDAKSLFDMFQQLGFQVLHCKNLTAPKIVVLLKAVAMHYDHINSD

CFACAVLSHGDETHIRRDNSTSLRKDIIYGVDGETVSTEVITKFFDDEQCQTLIGKPRLF

FFQACRGDTLDGGVDINISSSSVIHTSVSMPEIITARTMTDSSSVDPPNYHIEHIDDTEA

KKDPPNTASAVIDSMEDYIEKDTIVDPPPLYKDCLIMYATPPGHFAWRRGSGAWFIQSLC

QVLESDAVKFGNIDLQKALVHVARKVTFDYESNMPDNKKFHRKKESPVLYSMLVKDIYFN

NRSEESMNM

>KAH9498390_1 KAH9498390.1 Caspase-7 [Bulinus truncatus]

MSTDVKKNETSCFSGDEGFVLEDDIKKEFEETDDDIDGFRKYIKNKIETVTYFIRSFNKD

ETCGISHNPISASSQDFKETIQTNKVSEENSKSDNSDIYDFTHHHRGFAVIINNEKFTDK

TNEYQDRPGSSEDAKLLVKTFKHLGFHVLLLENLTALKVIETLHSASRDYKHSLADCFLC

VILSHGHQITKVFQDRNSALQEDVICGVDGKVVSTKKILDYFTDSNCPDLKGKPRIFFFQ

ACRGIKVDDGVEIKICGKKKKWRVPSSIPDTADISCQTSAKIRDAESSFSTYCEMEDELD

AKSEEKYVAEKVHHKIQSVMNPAQLHKDFLVMYGTPPGYYSWRRKTGTWFIQSLCSVLNK

RNIQQTPLMQALVQVSGLVAQKYESSVPSDPNMDKKKEMPVIQSMLVKDIFFTKKEKANI

GNIK

>KAH9498392_1 KAH9498392.1 Caspase-7 [Bulinus truncatus]

MIFGSRVKMEGGIQGLVIVFSACMVQVLAFGNYACVGIYNVYLLEEFDHDNVGVSLISSI

HFALLLGCGPFISFLMTKVSYRTLSILGATLVFIGILGTHLIIHLPSMYLFFGVFAGLGG

CFIYLPSHVLSGLYYEKYRSLSTGVATTGTGLGAIIMPVIVGNLIEEYTWKGDEGFVLED

DIKKEFVETDDDIDGFRKYIKNKIETVTYFIRSFNKDETCEISHNPISSSSQDFKETIQT

NKVSEENSKSDVYDFTHHHRGFAVIINNEKFRDKTTEYQDRPGSSEDAKLLVKTFKHLGF

HVLLLENLTALKVIETLHSVSRDYKHSVADCFLCVILSHGHQITKVFQDKNSALQEDVIC

GVDGKVVSTRKILDYFTDSNCPNLKGKPRMFFFQACRGIKVDDGVEIKICGKKKKWRLFM

NPAQLHKDFLVMYGTPPGYYSWRRKTGTWFIQSLCSVLNKRNIQQTPLMQALVQVSGLVA

QKYESSVPSDPNMDKKKEMPVIQSMLVKDIFFTKKEKANIGNTK

>KAH9494957_1 KAH9494957.1 Caspase-8 [Bulinus truncatus]DEDDED

MMSSFQYRETLAEIDSELHADDIKNLKFLSEPYIPKKFHTKFTSGLEIFEALEKCKKLSP

NNVHLLVEILTIIGRVDLALKLKEKANLADESMIYKEHVNPFYLLLFNLSEELNIQELQS

LSFLYGKVPKRENINIGIDLFSIMLQHVNLSPTKLDSLESMLANINRNDLVQEIKDFANR

FVKQGKYVWSNTEVSSMEIYGSDESIQNTNQDKEQQFRESPVYDAMKENYLYEDVMDQDP

VLESVQSSENETVSLKHINIENRQELEDVTEFSKLTTISTENEQFSLRNINVEFQQEPED

VAEFYKMTANPRGLCLIIDNQIFTYNPGDLEATQLGNRVGSEKDCDALRETFQDKLHFQM

QIEKNKTDFEIAEIIKRVSLLDHRKFDCFVCCILTHGGEGFVFGTNGRKFSIKEIIKTFS

AQSCPSLAGKPKLFFLQSCRGKRHQLAFPTSAMLEIDAPQVEPDDDDLLPPEADFLLGYA

TVADAVSYRSRSHGTFYVNELTKALNSYADRFDLLRILTKVNNEVSKRVYTEGSGEERIV

YKQIPAPQYTLRKVLKFL

>KAH9510360_1 KAH9510360.1 hypothetical protein Btru_042890 [Bulinus truncatus]

MAAEFLREHQFIVDVMSACTHDVSMEVRFEYLRPLLVDIFPRMTLSRDEGHGRASCEIMI

NICDSAKRLSKDDFVKFVDAIHKDHPTSAQLIRNKYKEISGKDLPSFPSQPQRVPVVQEV

TPVHCHADIQARSNHRVQASNILAQAVTIPAQVTKPQLQFTDKALPDYKLQQLPLIGKNT

FTTVYLMESTPRGRALIINNMIFDVFSERKGSDVDVKLFRKMFDKLKFETRVEENKKAWE

MDQIFRQFAKDSQLKNVSAISIVILTHGGANDCYFGVDGKIFKEHGKNKPVNCITKVDLA

KIFSSRNCFHMKGKPKLFLIQACRGEDENESYTQNDQQQGNHIHDDAPKSEHREEVSTLE

ADDGENEKPRADTADMCFVHSSSIGYKAYRNILLGSPFVIELTTCISNADHTTDFQEIVR

KMQRIFSDKKLENSMMTLPDFSTQLVKRWYIKPPPNY

>CAG5125257.1 unnamed protein product, partial [Candidula unifasciata]

DAVDARSNDAMDAGAETSARLQASAQARSSDNLSDCYKMNHEKRGMAVIINNRNFHPSTR

MGERNGTDVDAEAMYRLLDVLGFEVIDKHDNKTAAQMRDILCKASKVDHTDSDCFVCVIL

SHGEEGHIYGTDQPIAIDELVRPFKGHNCSSLAGKPKLFFIQACRGSELDHGVTVADAAG

EDDVEIRRIPTEADFLMAYSVVPGFYSWRNSAKGSWFIQALFDVFTKNWKTLDVLTMMTR

VNKKVAYNFESQSSTEGWSKKKQIPCVTSMLTKDVYFKK

>CAG5121219.1 unnamed protein product, partial [Candidula unifasciata]

GDRRDVRPGAPSEPNQYNFNFKKRGIFILINNRHFEEETGQQPREGSDVDAERLEERFQD

LGFEVRRFDDVTCAKLTSLLYDVASYDHSDHDCFACAVLSHGREGLVYARDGQVTLDMLV

LPFKGDRCPSLIGKPKLFFIQACQTKRQGIYLDPPSRQSKPSRSPSPVSPSSSKQHSLGS

QSLTGDDDASESSGIVHMRKIPIEADCLFCYSTVPGYYSWRNNQDGSWFIQALCIVLENY

GTKMELLHMLTHVNRIVAYEFASCSDREFGESMKQMPCIVSMLTRYVYFRPKKQDIR

>CAG5115060.1 unnamed protein product, partial [Candidula unifasciata]

RRDVRPGAPSEPNEYNFNFKKRGLFILINNRHFEEEVGQHPREGSDVDAERLEERFQDLG

FEVRRYDDVTCSKMTSLLYDAASQDHSEHDCFGCAILSYGREGMVYARDGHVSLDLLVLP

FKGDRCPSLIGKPKLFFIQACQCRRHSIYLDPTSRQGKVSRSPSPVSPSSSRRSPMSPSF

TGDDDPVDACGLVHMRKIPVEADCLFCVSTVPGYYSWRNNQDGSWFIQALCIVLENYSTK

MELMHMLTHVNRIVAYEFASCSDREFGESMKQMPCIVSMLTRYVYFRPKKQDIR

>CAG5125026.1 unnamed protein product [Candidula unifasciata]CARD

MDQKEFVRRNYIKLTEAIAPNVEAVVVWLFSNGVISRHMKDNVLESNKTQHKKTWALLDL

IVTRGPRALDYLYNAVLEEQLYEAADILKPEKGPHMPVVVLHPYVSSPAVQQTPTMEYDS

PDEGLPDTWPSEEAIKACTTVRMVKETHRRMINLFKETLTPRGQEMRYLMSAKPRGRVLV

INNEEFTHLDNRVGSFEDRKSLETLFKALDFDVEVAKNQTKERMWELLVKESEKEYHKKA

QCFVLVILSHGHTGVVFGTDGRLRNGHPENFLEIKNITDLFCKSENLQGKPKLFFIQACQ

GANKNEGHPVGQADSTVPQKSAAADGEVPISSFAASSAPVAGVGDTKEQEEIIKEAKATL

QEVISTTQSDAAGDKRPSGADVFVAVATIPGFLSYRNTQIGTWFIQAISYVFAKYAYKYD

LNKLMTWVNNLVGRAETGKERYKQICEKKDSFQKEFYFFPGLVDDK

>CAG5131086.1 unnamed protein product [Candidula unifasciata]CARD

MEQRHRDIIRTNYCLLAEQIKTSVSDVCACLYQSRIISERMRENILETNGTSRQKAEALL

DLIVKRGHAAFEMLYTAILESELYEAADILNPIRAPHRNLGTAMSQTSDLDAPPSREEEI

AAALSMNVTPPIREEDVAAEADVLPMNETPPYREENTAVHAGVLMANELPAQSASSSSIS

SSSAIDNIPKSWPGLGFVKKEEIKVQIVQPDSPMYEEYLKSLTPQGQRFYYTLQHKVRGR

VLIVNNETFKTNHGTKKENEKVIYCSDREGSSYDKDNLEVLFTQLGFLVQTVNNQTKEDM

EKIFKEECEKNHREYDCFILIILTHGHKGIVFGSDGYSVIKANNELEHHNYIRVKDIKNM

FCSIKSLRGKPKLFFIQACRGDAYDKGVDDSTTGASLESGDTDMDIQLENANDTISELEE

MEESDGPHDKLPTAADYLMALSTVSDYVSWRNPNHGSWFVQAIVYTFWKNAHLYDLNRLL

TMANDYVSQAETAEGMKQVAEKKDTLRKSFCFFPGLHPGLPSSPLN

>CAG5136339.1 unnamed protein product [Candidula unifasciata]

MSANDMVDALGVGDTDEPDARGQPRNSLPRMSESAFSSLLYKMDHPNKGVALIINNTTFH

PRLFIRGHIGDSKRSLQEPSADAELMVDLLKTLGFKDVMFYKDQTAADMKSILQKVASMD

HTDSDCFVCVILTHGDDGYVCATDDKIPVDELLQPFKGHLCKSLAAKPKLFLIQSSKGFM

PEKGERSFESAGEESNEEETVRRIPTEADFLMAYSMIPAWKNISKPSWFIQSIHDVFEQN

WQTMDLLTMITRVNKKVAQAFELNANNVYENQNRQIPYITSMLTKDLFFTTK

>CAG5120412.1 unnamed protein product [Candidula unifasciata]

MKIMDRFTHDIASEVDFQYLNPHLRAIFPRIMLQRAEGRDRASYDIMTTICESARTLCMG

NFLNFLAAIRIDYEHVFNLINTYYREETGHNLPNYENAQNFSQSEPILLQNSPSLAQNAS

FPPQNAPLPQHNASIPPQNAPLPQHNASIPPQNAFLPQHNASIPPLNASRIFQNAEPLEP

NMSTAGYMDVLNSNELGTPETFHSSYNVSGSPAETTEPNISSQLRKYNKLVIENPNSLYT

MESTPRGHCLIINNELFQNLPPRKGSSKDLNNLDMMFNFLGFQVVSYENKTATGMKILLE

EFANSPALGQTSALAVVILSHGGPSDIIYGEDGALIDNSPVAGTFITKLQLQKVFSGTNC

PSMNGKPKLFILQACRGEESEDGQENNYCRDDAPSLDHIREPETVSDDPLPEHRAETADM

CFLHSSSLGYKAYRSMSQGSPFIQLFTERIFQHAHEKSLQSIVMMIQNEFSTKAITTTKM

TMPEYSSSLTKKWYFNPPHGRRSMYNFGTHE

>CAG5135344.1 unnamed protein product [Candidula unifasciata]

MDEETVVKLESEIEEESQAEYVSENKDSIDGFRQFVKEQLHNMCGKKGSPESSPQPSSSA

SSSSQDFLENRTAFIPSETGQEDSEDIYDFIHADRGLALIINNEDFSRSKNFGSRPGSSY

DEVSLYRTFKKLGFRVLVKRNLTARQMVEVLKLVKDEYDHANADCFVCCILTHGDETISR

EANEKRGLMRQDLLFGVDGEAIPTRTVVSFFNDKYCPKLKGKPRLFFLQACRGSKYDHGI

DIQVSGSTHTTRTVGGKNRDGIDKGSPVDQTDARTITRKNSFSSSVDPPEYPKNTLDSTD

AKPEANVARGNLSLPAAGASGDNADRSPAREVPACPTPLYKDCLVMYATPPGHFAWRRKN

GAWFVQSLCDVLNADHIDQISLMKALVRVSYQVAINYQSENPYRPDMHQKKETPVIESML

VKDVYFTKRK

>CAG5119597.1 unnamed protein product, partial [Candidula unifasciata]

ATSAASFLDVTKPSKVAREPDIDLTDVYDFSHRQRGLAVIINNEDFSLSRLNNVRGPLDD

RPESSYDALALSRTFEKLGFDVLLKNDLEDKKMVVLLESAKDDYDHANADCFVCAILTHG

EETFVSTGNNLPEVRLDMLSGANGKSVATSTLVNMFSDDACPKLKGKPRLFFIQACRGEE

CDNGARIQLIKHTATTSESDRQSHDGTAHGKTQTETERSHDGEASSNCLPRAQGLQTGTT

DANPDYSENSIEEDLQNINMNATASGGVDMIDVPGAAVQRVRHVRLCPTPLYKDSLIMYA

TTPGHYAWRRDSGAWFMQTLCKVLDNPDIGSKSLLSALVNVAGIVTRDFESSTPNRLSTH

QKKETPVIESMLVKDVFFTPK

>CAG5136338.1 unnamed protein product [Candidula unifasciata]

MDDSEMDGPSYEKSKVCQETASQAKIEEEESYCFSHPRRGVAVIILYSQDFKGGERPGWK

IDKENLQRTFSHLGFRVDIYQDLNKQDLMVHLQKVSREDHSDCDCFVLAVSSHGQECHVG

GVREDTILTIDTSIKTREVVDMFTNKACPSLTGKPRLFFIQACRGQKLDHGHEILPEKST

SNDDTLDASPVDSDLTDVIPMGFIHHRYLHTMQQNTAVANLMSRLRHVTITKQGRQVLQE

GALPKQVVSPAPCHEDFLVMYATPPGYFAFRRQNEGSWFVNSLTKVLYASDGTRNLTKEL

TRVIRLVAENFFSKNKDKDLHGKKQTPVLYSMLCKDIYMGPKKPLQDGR

>Octopus_bimaculoides_(XP_014780506.1)

MGSGDKKSLRPGAPSDPLEYNFNNKRRGIFIIINNKHFDESTGQHSREGTDIDAERLEER

FQALGFEVQRYDNITRTKMTGLMYDMASKDHSDVDCFGVAILSHGREGAVYATDGSVPLN

ILVLPFKGDRAPSLVGKPKLFFIQACRGTKMDDGVEVVDGSSFTEDDSVDAVASADVIMR

TVPTEADFLFAYSTTPGYYSWRNNEEGSWFIQALCIILENHGKEMELMHMLTLVNHIVAY

DFESCTNEDFTSKKKQMPCIVSMLTKYVYFRPKKQEII

>Octopus_bimaculoides_(XP_014783598.1)

MKYNNMNEKDKETLIAKRVELVENIMMEELLTHLIAKGVVTKRNAEYINAINVNSKKVEY

FLDLLPKRGPDAFQIFCEALNTTGQEFIVSHYLKEDNKTNDLSFDCIPTAKDNEPSNQLL

LKDNDNSIIRIKYSSTSPLKKCNSTPTKSSPKSPEVRHLLKHVISEPTPSQFSLNGNQTG

ITRCDRIDNSMSPAIEEPWPKKRKVCNEQDIDILKIKPATQEFYYKNHSQSYKMDGIPRG

LALVININKIAGMAPREGSSIDHRRMSYLLKQLQFTVISYDNENLTAENILIKVREFKDL

PEHKNADATVVCLFSHGTNGYIFGTDGEKVPLNDVFSLFDVDECPDLRKKPKIFIMQSCR

GDNVDLRRVPTLKVSAEDFADNVPVQKTEPTMSDSLFCYPTQEGYRAWRNYEEGSWFIQA

LVDVFAKHAWEKDICAMLNMVNSMVSQRTSLNPSTGLNGAKQMSEFKSTLRKPHLYFYPG

IGRKQN

>Octopus_bimaculoides_(XP_014783442.1)caspase2

MKYNNVRITSNFNKTMNEKDKETLIAKRVELVENIMMEELLTHLIAKGVVTKRNAEYINA

INVNSKKVEYFLDLLPKRGPDAFQIFCEALNTTGQEFIVSHYLKEDNKTNDLSFDCIPTA

KDNEPSNQLLLKDNDNSIIRIKYSSTSPLKKCNSTPTKSSPKSPEVRHLLKHVISEPTPS

QFSLNGNQTGITRCDRIDNSMSPAIEEPWPKKRKVCNEQDIDILKIKPATQEFYYKNHSQ

SYKMDGIPRGLALVININKIAGMAPREGSSIDHRRMSYLLKQLQFTVISYDNENLTAENI

LIKVREFKDLPEHKNADATVVCLFSHGTNGYIFGTDGEKVPLNDVFSLFDVDECPDLRKK

PKIFIMQSCRGDNVDLRRVPTLKVSAEDFADNVPVQKTEPTMSDSLFCYPTQEGYRAWRN

YEEGSWFIQALVDVFAKHAWEKDICAMLNMVNSMVSQRTSLNPSTGLNGAKQMSEFKSTL

RKPHLYFYPGIGRKQN

>Octopus_bimaculoides_(XP_014790087.1)caspaseY

MNTEQVHHISKQQFMQIRVDFVRHVDDIESFLDEALSNGLLIEGHRNEIMSQRNPDDQIR

KLHDYIFKKLPYDSDKLMSALKKSKHIKIFDLLDEQTAYPMKFKPHGRVILINNVKFDDE

ETYKERHGSEKDVEGITKLFTDFNFDVHPHPNKTAKEMKIIIEEATSKSTSGEDCFVMFL

MSHGIIGNIVGTDGKELSYSTINTILKESSQLKDKPKLIYINACQAKSEKEDVKQYFDVA

DLHVTFATVPESLAYRSSKRGSLFIESLLTVYKNNKEKCGISSLSFEINAQVAEKNDKIS

KDQVSSNYSTLKREVILQATD

>Octopus_bimaculoides_(XP_014775611.1)

MTTTDAQCVSEYTTDDRFHQSTGEDLMKCFNKYDMTHKKRGVAYIFNNENFQHSELKTRL

GSSKDAEDFKKALIKLGFHKDDINLYTDATAEEMCGALKQFGKKNSHPDLDCFICAILSH

GDSDDLIYGYDDVVKLDKLLSYLRPGCCPSLTGVPKLFFVQASRGSKIDFGIEKTDADLL

RNRERAPKIPVMADMLVVYSSCNEYPSFGNKEGGSWFMQALSKILMKYGNEYEIMKLLTA

VSNYVASLGFQSSDNSEYNGNKQMPQIMSTLIKELKFQSKN

>Octopus_bimaculoides_(XP_014775614.1)

MTNPEVESEHLTAIKHKDALFKSKSTTGACFPPSGEEDLGKYFAKYDMTHKKPGVAYIFN

NENFQDPLLNTRYGSSKDVHDFERALIKLGFHEDDINVYTDSTAEEMFDVFQMFNEENLD

IDCFICVILSHGSSDDIIYGYDGVIELDRLLSCLRPDRCPSLTGIPKLIFIEASRGNKLD

YGVLKSDAAVIEPDKNSSRIPIMADMLIVYSSFAGYQSFRDRQNGSWFMRGLSKILIEFG

TKYEIMKLLTAVSNYVASLEAPSWYNSEFIGCKQMPCIGSSLTKQLKFVRKNKSD

>Octopus_bimaculoides_(XP_014789170.1)

MDAVNVDGCTGAKFHLPTEEDLMKCFDKYDMTHKRRGVAYIFNNENFKDPKLETRHGSSK

DTEDLKTALMSLGFREDEINVYTDPTVKEMLRALEDFNEDNPDIKGNIDCFICAILSHGG

DDEIIYGYDGKIQLDELLSCLRPDRCPYLTGIPKLFFIQACRGSKIDVGVEKNDACGFGF

EEKPPRIPIMADMLVFHSSYKEYPSIRNKDKGSWFMQTLSKVLKKYGAEYEILKLLTAVC

NHVATLEYQSSKYPEYNSCKQMPQIMSTLRKELKF

>Octopus_bimaculoides_(XP_014775609.1)

MTTTDARCVSVTGDQNRRPIVEDLMNCFHKYDMTHKKRGVAYIFNNENFEDPNLKPRHGS

SKDTEDFKNALMKLGFEEDDINLYTDATIQEMLDALDLFGKENSDIDIDCFICAILSHGG

SDDIIHGFDGPVELEKVLSCLRPDRCKLLTGIPKLFFIQACRGTTPDMGIEKNDADYIDH

EERSPKIPIMADMLVVYSSCNKYPSFRDEKDGSWFMQALSKILTEYGTQCEILKLLTAVS

NRVAALGFRSSANPEYNGCKQMPHIMSTLIKELKFEQKN

>Octopus_bimaculoides_(XP_014784582.1)

MNAEQLYYISKNQFQKLRVDFVKYLDDIDSFLDEALANGLLTENNIDGIMSERDSNNQKR

RLHNILHKKLPDGSREFVSALKKSEHNEILKLLDEQTVYPMKFKTHGRVILINNGKFDDE

EKYQERFGSEKDVEGITKLFTDFNFDVHLYSNKTAEEMEDIILKEAAESTAREDCFVMFL

MSHGVLGNIVGIDGEKLPYSSINKTLKKSAHLKNKPKLIYVNTCEINPTEKDIKKYFNVP

HLYVTFATVPENLAFRSDESGSVFIKSLLTVYERNKEECDISSLSLDINSQVSKESNXIS

KDQVSSCYSTLKKKVILRATD

>GFO38322.1 caspase-3-like [Plakobranchus ocellatus]

MSDHVDAKGDYDQTDARGETRYDKHRSMPPKVSELDFVSDHYNMKHDERGLALIINNKKF

HPRLGLSERTGSDVDAYKMHMLLGTLGFEKIQQHDNVKANDMKDLLYQAAKKDHTDSDCF

ACVILSHGEEGYVFGTDNKISVDDLVLPFKGDRCTSLAGKPKLFFIQACRGQELDRGTTV

ADAIGDQEAEEEIVVRSIPTEADFLMAYSVVPGFFSWRNSARGSWFIQAIYDVFTEHWQD

LDLLTMMTRVNKKVAYDFESNASEEYMRRKKQIPCITSMLTKDVFFK

>GFO14050.1 caspase [Plakobranchus ocellatus]CARD

MDQVYKDLLKKNHMSLKRSISKGVIPVLEYLESTKIISEDMKERVECENKTPGDQASALL

YLIKTRGRNAFQALYDALLHAEIYEAADVLKPDQGPHGPSYQALAPPGRSSSIGQGECDE

LSEDEPLPDYWPLAEQECDTFVLVKTIQDCTKSKQKLQKLYDHSLSPAGQDDYYSMLHKK

RGFCLILSNQNFKHLEERKGTDRDVTHLNHLFTQLGYDTYVKKDQTSKGMKEILRKQVLD

KNHTNYDCFILVILTHGEKGVVFGTYGRLDEKSQKPEDCLEVLDIRKEVCSVPSLLHKPK

LIFIQACRGKQRDEGQSLNSGKKIKSDGDIGGQDAFSGGGESPNDPADSGGNQDIAAPNV

TPEVLPRPSEETEQGPLSSDFDLKARKKPVAPPTKPEDELDACGEIVPSLADSFLAMSTT

EGYLSWRNTKSGTWFIQAIVYVFSRFAYKLDINKLMEKVTRLVSKGEEEKKGYKQICEWT

PTMRKTFYFFPGQSKID

>GFN83778.1 caspase-2 [Plakobranchus ocellatus]

MTPHGKIERVIWSPATSLTTSPLFEILSHFPFQLNRTPEEQVEELLNELPKRDCFEDFCA

ALIADGQEHIVKKYLKAAPPTSAKSQPSAASASSPCSSASRQPSHAGPSETTDSSASFMH

SAVKRSHPPTLSSTATICIREEDKQHLIISFTDEGASPEKRKKALEAGTASVESALSGPV

AYKPDETRYTMKHFVCQESESDEHTSLVPISEANTLIFNNPQNIMNNPNISDPFRYAAAF

PANQQNMTSAAVNNSPIAQFCVAKSASLSSQPNAKDHSSILSSLSAGIQYAAAPSSSSPL

KDPSSLVNKQENAHIRSLLQPQQGEALLNAKPAMAAESGKFIRLEPGVDVSQFQDPQQYL

SAAGLLGRSKVFNSGGSSRSPSLNQNDAADTAFLSGPAKNLHLNLAKQALSTSTISDFQD

SLDSGLLDVTSENLSKFAVGAAISPSPLNYISSGSNNGMNDSVCDLDQIDLPDGCVNVKV

EHCAKQFFCANHKKSYPMRRIPRGFALIINVDEVEGKPPRKGTNFDRDNLSNLLCQLHFH

VIVYNDKDGLSAQEMVNKLKDFAKLREHADMDACFVCLLSHGEEGFIFGTDGKRIPLEEI

FMLFGNTNCRGLIGKPKVFIIQACRGGKSKWRVFSFKGVLPSRHVVGMSQWEDEVEARWR

SS

>GFN96839.1 caspase [Plakobranchus ocellatus]

MNHQGGEDISFHHSQFRGLAVIIINTGQRRGSDRDRDLLQEMFSYLGFQIHVLLNQNSKD

LSQELKRVSNEQLSDIDCFAVAISTHGIEAQTREKVMEDVIETADGKIKTKVILEMFADH

KCPQLINKPRLFFIQACRGEQLDSGCPMILQKVARPEQAERPEDQADVDTEGNPPPEVQN

RNIHKEGARALATASADDEDENVPVRPPVCFKDFIVMNATPPGYFAFRNPINGSRFIRSL

REGVCKYAHSMSLFRTLTKVSGLVAKEESYTPSTPHLHRKKQVPVISSMLTKDIYFKPQQ

AKYD

>GFO02264.1 caspase-3-like [Plakobranchus ocellatus]

MSTQGEDVEDLYFHHSQSKGLAVLIINMGENDQRRGADKDGDLLKEMFSHLGFQTLVFQN

KSSNDLLRELKNVSMEQLSYIDCFAVAISTHGREAQTKEKVMEDVIVTADGKIKTKVILE

MFADHKCPQLINKPRLFFIQACRGEQMDSGCPMILQKVAMPEQAERAEDQADVGTESHPP

PKVQNRNVQVSGAGVQEEGARALATASADDEDEDAPVRPPVCFKDFIVMNATPPGYFAFR

NPSKGSLFLRCLKEGICKYAHSMSLFRTLTQVSGLVAKEESYTPSTPHLDRKKQVPVICS

MLTKDIYFRPQ

>GFO26738.1 caspase-8, partial [Plakobranchus ocellatus]DEDDED

MDKAKYRELLQDLHSELADDDIKCLKFLAQPLIKKRQYEGIKDGLGFFEALEKCGMLSAN

NTMFLCQLLETMRRRDLISKITGEDVSWDAVGGDLELVCPFRKMLFDLSLEIPDENLVQM

HYLCGNVPKRTPIKDPLDLFMFMIRERCLAPDKLGFLKKVFIDVSRKDLYEKVEAFEGDV

HPNECDYRNVLIQTAGPVYQEAPFQSSFTSSHTRGQQLENLKEDPMSHQESGFSMQERMY

DIPYQTASIESNPNSSRQPLPDHNMEDSDFQIPDLNSSESSSSSYIPFTADISVGEAAAR

ITAEETGAIRYPVHQLDKSECYKMDAEPRGLCLIIDNQTFMPDHMAECHARNLAYRGGSE

VDRDNLRVTFQHLKFRVEEVINATDQKIRDTLMAYAAHDHGAFDCFVLCILSHGVSNAFY

GSNGGKVMEHELFDMFSAFNCPTLSTKPKLFFFAYCRGQLHRPDNLRTTFQHLKFRVEEL

IDVTDRKIRDTLTAYAARDHGAFDCFVLCILSHGVSNAFYGSNGGKVMEQELFDMFSAFN

CPTLSTKPKLFFFAYCRGQLHRP

>GFN78499.1 caspase-3-like [Plakobranchus ocellatus]

MEHHKAGGDEDGDVYDFNYLKRGLAVIINNEDFSPSSDFDDRPGSSYDASALYHSFAHLG

FDVLLYSNLTAWKMVEVLRAAARDYDHENADCFVCSVLTHGDQTWSNREYDRMGLTVRQD

LLFGTDGKAVATRTVVELFNDTASPGLRGKPRLFFLQACRGNKLDEGQEMHVVTPIRGIR

HRHGKDMPDAADGGGDGTLEAGKETIHTEDTGTAVEPAQRQTSLSSSVDPPGYPKIEDKV

DAQPAPDDEDDENNHHQRNTGLTEEESQSTPSSSQKASFTDAYGEGQASNVDHFSGFLGK

VQDVPVSPAPLYKDCLVMYATPPGYFAWRRKNGAWFIQSLCKVLDSRHLGNLSLVRALVQ

VNGFVARNYQSENPSKPSMHAKKEMPVIESMLVKDIFLTQKRRF

>GFO38603.1 caspase-3-like [Plakobranchus ocellatus]

MDSQSCIKDFRANFRPRLYQLKLCPALSDDYHLVSEVLNPLIHFISHHIIFANIKDWLVP

IFTTQDLEHAEASGKGSIDIMRNICEHAKRMMLPEFVQFGEAIVEENSHVAEKLMERVDE

LLATSKVSADESVQNGAFSLQPRTNSSSKFSSVESANHFNSYGSASAGHSSTFADHRTSG

RSGSGFQDSFSPLALCSNHTTKRNYTIQSSSCTNSTHAGCTALPSGQELSRIVRHSGQHP

SEVYDMSTTTRGLALIINNEDFENFSKRSGSSLDVTNLYNMFHYIGFKVTIQRNKTAEEM

KTLLRQFSQHKLLGTVSALALVILTHGGSNDCLFGTDGTFSRCGPVSGTFITKMDIIHMF

KGSICRSMEGKPKLFIVQACRGEEENSAQCYQTAAKRAGLPTKHNVRLRSDAPSVTPTSV

VNPSTTSSASSTYQHRHDSPGSIKSADTADMCFVYSSSYGYKAYRTRSAGSPFVQVFSST

VRRLAYEKEFRDIVQEIQRYFNTEPLPNNMMTLPDFSLQLMRPWYLNPPKWANSA

>XP_025081735_1 XP_025081735.1 caspase-2-like isoform X1 [Pomacea canaliculata]CARD

MQFDFHRSIRENRNVTTESRAPMGFEIAEHIQKYQNHQQQRMEKVHRDLLVTHRTELVEN

IDVTNGLFSELVQRKVLNQRMVRNIKNNKDIDTQIEELLDILPRRGPDAFHLFCESLIAS

DQQHIVDSILKVNLAEAPTEKTASSCTVCTVHSSPHVSSMATDTSCLTATTSLSITSATS

AALQQPSSPQKRGLKDEKKAPVESAEDLASETESDCSPRKIRSVKSDSPSVICIKDGDDK

SVFRICHSKIKEKPSELISQKQLDKNRFTTDEMRYVMKHFVTVDSQEERRQEVNRNLAET

LLHDMTRDYFNAFQSQPRLYPKNPIQASDNHRYVSSLRESLSRCDSADTMDRHFPQKAGA

TCHQTADKDETVSVLMPPQTTQALHLSEKNMDLTDGSVCVRVEHCNRQFYMENKEKAYRM

LGMPRGQALIINVNEVHDKPPRRGTDIDRDNMHNLLTQLHFNVIVYNDCDDLTAQGLVNK

LRKFAALPAHHTSDACIICLLSHGEEGFIFGTDGKRVPLDEIFVLFGNRHCPALLSKPKI

FFIQACRGGALDHGVKLDLDETDGPSIQQSHLQLPSVSDMLICYPTLEGYYAWRNRERGS

WYIQAVVQIFMRHAKCEDICAMLNRVNELVSRKVSRCPQADMDRMSQMSEFKSSLRRPYL

FFFPGIGSPHIDNV

>XP_025080583_1 XP_025080583.1 caspase-2-like isoform X1 [Pomacea canaliculata]CARD

MDPEHKHRLQIAWAQLLETVGNVIGVVDGLFAKKVITRAMKERIMQGHTVDSDKLRALMD

LLPKRGPEAFDAFYDVLIEQGEFEAADVLKPELRDKHKALSPPHPAVGIQETAVSLIIDD

DLPEKWPDPSVHDPVRTKVVPVTEEGMRQNFANALAGSKSIFPLWRKTRGIFLLIDNNNF

KDARTAGLDLGDRDGTNFDRLALDMLFRQLDFDVIVEKDCKAQEIADAVQREAGKSHLNF

DCFVCAILTHGERDIIYGVDGNKLLLDEIIAFVDGEKCPTLIDKPKIFFIQACQGKKKDK

GAGMVEQSSAAGDKPADDSISNIISRVINLRMQTEQHDAELEKRTGTKSDIFLAMATVPD

FVSWRNTNHGTWFIQAVVFIFSTLCHKYDLLSLMTKVNNLVSRAETKEERYKQVSEFKSS

LTKTFYFFPGLTKEVSSQLMTSSSDGRQTAHVHPQQPQQPEDTSSSVMYG

>XP_025080591_1 XP_025080591.1 caspase-3-like [Pomacea canaliculata]CARD

MMEKEHKDRLQRAWLPLLNTVGDVIRVIDGLFTKNIITRLMKERIVQGNAVHSDKLRALM

DLLPRRGPEAFDAFYDVLIEQEEFDAADYLKPELKGQHKTRNDDLPKKWPDRDVHDPDHI

KVSLVADKEMIQNFENAYSDEVSIYPMCRAKRGIFLLIDNQNFYAARAAGIKLADRDGTE

FDQTALHSLFEALDFKVEIKDNCSAKQIADALQEVARKSHRDFDCFACAILSHGEQGVVF

GVDGDKILIKDIIGAMDGVRCPTLIGKPKLFFVQACQGNKYDKGVSVVKLGDAAGDMPTG

DSVSTGDGVSTGADVSTVADHMKNMTMQDAVQHKEELDVMDLYNLVDNVLNKQCDNVSNG

KTGTKCDIFLAMATSPDYVSIRNTKFGSWFIQAVAYVFSKFAHKNELLHLMIKVNRLVSR

AETREGGYKQVSGFNSKLTKELYFFPGLKKQVPSPRPETSIEPQAQN

>XP_025101998_1 XP_025101998.1 LOW QUALITY PROTEIN: caspase-3-like [Pomacea canaliculata]

MADDNVDARPEGDQGDALASPGAPAVLPKGDLPVISKTDFVSDSYNMNYPRRGKAVIFNN

KHFLPKRAMGERTGTDVDARNLYFMFTEMGFDTDIKHDLKAHEMESYMVKVSKEDHSDAD

CFACAILSHGEEGYVYGQDARIEIEKLVGPFKGNRCKSLAAKPKLFFIQACRGTNLDDGV

XVADALGTEGRTEVEIRRIPTEADFLMAYSVVPGYFAWRNSSRGSWFVQSLVEVLSKNWK

RMDLMTMMTRVCRKVAYEFESNATREYMNRKKQVPCITSMLTRDVYFTPKAK

>XP_025101960_1 XP_025101960.1 caspase-7-like [Pomacea canaliculata]

MISCNVTMAADLEDVSNSNTGYLKETEDLEDDEPEGVKDFLKKKYKQVKKAFRRSGKKER

PLNMATTDDKEDGSRTEVSPIVEDPDEYNFSHSRRGLALIIDNENFTSASRLPSRQGSDK

DLTSLGDMFNRLGFEVEGYKDLKGEDMWKVLTEAAGWKWHKDSDCLAVAILSHGGEMKWP

QLWNHRKEILRQDFIFGVDGQYLLTSSIMDCFNDDNCPALKGKPRLFFLQACRGENVDPG

QDIAVVSDKPEGLHMGLNTPHGTYGANGSVSGDTIEQSEADTRPTQGQITHDTGSGDAES

QGQRNDANEEERDETNKDRPAVSLVDIYNDFLVMYATLPGYYAWRRPTGSWFVQSLCTVF

GTYRPRMSLIQALTRVTGMVTTLYESVNKKNPHESGMKQTPVIQTRLLKDVYFT

>XP_025101913_1 XP_025101913.1 caspase-7-like [Pomacea canaliculata]

MGDTGLSDEAASARVSNHGILPSSNIENAILKEQASKSTFCDKDDQNSCNTATVLSGLDN

LSVCYTSNIVGEPDQHISNVSGLGNQPSCNTGTYPGLHDQPSCNVPMIANLSDVSSSDTD

DFEDTEELDIDEPEGVKDSLRTSTNRFKKALRRSAKRERPQNVASINDEENRSVIDEAAP

AVEDPDVYNFSHRRRGLAVIIDNENFTSTSGMPSRPGSHEDLKSLRNMFNRLKFDLRCYK

NLKGADMWKVLTKAAAWKWHKDSDCLAVAILSHGDEIKWPQHWNHRKEIVRQDIIYGVDG

QCVLTSSIMDCFNDDNCPALKGKPRLFFLQACRGMKKDRGQNVAVVSYEHGGLQTDIQST

WGGCGFDGGDTVEQSQTDARPEQGQASRNTGCGYAESQGHRNDASEDETDGSTPAVSPAP

IYNDFLVMYATVPGYFAWRRPTGSWFVQSLCAVFEFYNPRMSLIQALTRATYYVTTCYES

ENKKNPCESGMKQTPVIQTKLLKDVYFT

>XP_025101988_1 XP_025101988.1 caspase-3-like isoform X2 [Pomacea canaliculata]

MLKSLSSSGDEQQTLHADYDESEEYKFDHKRRGLALVIINEKFRGHAPREGSQIDASRLS

DTFSLLGFEVRLVKDCTLEELRCILKGIAEEDHTDADCFVLAISSHGDEIIQKLPKDKKQ

VRQDVVFCTDFFMQTRDLVFAFSDSQCPTLTGKPRLFFIQACRGSQLDPGQDMEVFQSTL

DEVDGFPDAKVIVSPAPLFKDCLIMYATPPGYFAFRRPKDGSWFIQALCSVLENCDGTKT

ILQLLTRVTQKVTQEFMSRTPLNYETDCKKQTPCLYSMLTKDLLLKAAHQSGQILVWS

>XP_025101968_1 XP_025101968.1 caspase-3-like isoform X1 [Pomacea canaliculata]

MSNSHEILETKKSSDESDVLSITGAFRSMLKSLSSSGDEQQTLHADYDESEEYKFDHKRR

GLALVIINEKFRGHAPREGSQIDASRLSDTFSLLGFEVRLVKDCTLEELRCILKGIAEED

HTDADCFVLAISSHGDEIIQKLPKDKKQVRQDVVFCTDFFMQTRDLVFAFSDSQCPTLTG

KPRLFFIQACRGSQLDPGQDMEVFQSTLDEVDGFPDAKVIVSPAPLFKDCLIMYATPPGY

FAFRRPKDGSWFIQALCSVLENCDGTKTILQLLTRVTQKVTQEFMSRTPLNYETDCKKQT

PCLYSMLTKDLLLKAAHQSGQILVWS

>XP_025088866_1 XP_025088866.1 LOW QUALITY PROTEIN: caspase-10-like [Pomacea canaliculata]DEDDED

MGDTLQEDAIVLPRPELREAVMIVDNELHDDDVKTLKFLCQNLVPGNKVNRVKTAHELMD

LLESRNLLSSVNYDLIPDLLQCIGRIDIVESLGYDEEEVLTNRDRGLCQINPFYILLCTI

SNDLSDEDVKKMAFLYGKIPRSQALSSGLDVFTTMIHHKTAGPDNIQALKDIFMSIQRND

LVTKLENYSASGGASVFKDLKKIFLQSRTNITSFRFSLPVEANSSPRNTNIVCDPEETTV

TQLTPGNVGGEPVAPPSHCPAAKIADAARADNQAEDRQALQDALDIEIVPEKIKEVLAKE

IPDEYWVEISNHLCVPLRETGHSPKGDLNINIQTLGLDSWEVSMQESVYSHNEASSAASA

KYMSVETVSQSASEIGQITNNVQRMSLQSQNPVPIYKMTSMPRGICLIINNRNFYKDGQR

RSKEMPVREGTEIDKEKLMRTFGNLGFIVQPHDDVTDTGMLQVISQMALEDHTSYDCFVC

CILTHGLQGELYGVNGITVPIKDLTGPMRAQSCPTLSGKPKLFFLQACQGRDKQPGYPDI

QTDAPIATDSPQEIIPNEADFLLGYATVPGYVSYRSKTRGSWYITKLTELLSRHAYEHDI

LDILTLVNYEVGRGDAKMEDGMFKQSPAPMYSLRKKLLFTQLSN

>XP_025083090_1 XP_025083090.1 caspase-2-like [Pomacea canaliculata]CARD

MDERMRETLRVSRVFLLDNVVMKDLMDYMIEQDIFSDAMRERIMAGQTNREQIRKMLDEL

ARRPHGCFEKFIVALRKSSQEHCADFVQSIYSKNILSQDPSIHATMSSPIQASSDVETSS

SSQKPSYSLQYRGNNPEVTNYLIAGPCAALPDIIMEEPESFKAPVQESTFRQDLSSKDEY

PMDTKPRGLALVINVEKFANMSRREGSVTDMNQLHDLLNYLGIKACYKTNCTADEIHEYL

KAFSKHEELKNVSCLVVAVLSHGGKDGIIYGSDGNTLKDDDIRSYFSITECPLMENKPKL

ILIQACRGDQEDRVSQPTSISGSSLKRELGPSLDSNASNSSQRRADLSDICLIHSTVSGY

VSYRHIRDGSPFIGVFCQQMKLHAANEHLTDIMIKVNNVLATSELQCEVRTLASSTNTLT

RKWYFRPSDGFAPFLSQ

>XP_025081418_1 XP_025081418.1 caspase-3-like isoform X1 [Pomacea canaliculata]CARD

MALTSLCFDTKMKAVLRKYRVELVSSLQVDDLLDHLIRDHVFECDDYIKAEKTRQDKVRK

LLEDLQKRPASDFWKFIQAMISTQQSHLAEKLIQEYKHLGGVVPLVTSSQHEHPQQPTER

GRRNGSPYHEWSDGRVVTSPCDATTSDSEDHRQSRQQNTNRTVYEQKKAEYPLNIYKMES

TPRGLALVIDNESFDPGVGDRHGTVEDAEALRNLLIDLDFEIQMKRNCSAEEMRNTLQNF

ALSLSGREIDACIVAVLSHGRRNEEILGTDGQVVTTSEMRHYFGNQLCPTMAGKPKLFII

QACKGDMEDTLCGQRRHVQGESLADVEKEVFHQPAVPRPLPDNADICFLYATVEGFVAYR

NVFIRKLCEVFRANADKFHVKEIATLLNSEMASLSLDGRLTMSEDRGSLTKFWYLNPPSS

V

>XP_025096263_1 XP_025096263.1 caspase-9-like [Pomacea canaliculata]

MGSVQSRDNVDILVHGCGAGKLIRKYSQILWTNSRESNCLKIKVGTDTRQAHILGFLPTW

APNDVHFNVRQKTNRGLIFTSLKEKVVVLLFDFNDRMTLEETSKYFSAWYYNLGKHADRV

LLVGVNGGLEACDRRLATCNREPVTEAKLRTVMNSLRLHSLQTLKVDLDKPFDVKSVFVK

AGGLFYMNFVNQEPLIAEPLEDEEPKPEPPIDAVLHVHLGKYKLNSMPTGLCLVINNINF

NAASLDPRRGSEKDYSNIERVFSNLGFLVQEERDLTASQIDFLLQQTAHDYHSDLSCLVI

FFLSHGRGSDTIHGVDGHVVRLHDVMRVFDGENCPQMAGKPKLFFFQSCRGGVNMRGVQP

AVDSGVEDNADDDGEEEEERRYTIPVMRDTLIGFSTEANHVAYRCPENGSWYIQTLTETL

ERLCEDYDIEDILKITTEKVSLRSTPEGYKMCPNYRTNLAKRLRFRRSPSVAVRGKYSTL

AVQGR

>XP_060600760_1 XP_060600760.1 uncharacterized protein LOC132754183 [Ruditapes philippinarum]CARD

MEKRHKQILKDKRPDLVESLNMIGGLFTQLISRDILDQRMVRTIKNGKTEDEMAEGLLDF

LPKSNQGEKTFDLFCEALIADSQSSIVRLYLKPDSKKETEEKNVVQESNGANAVEDTLSQ

VSGSQVTGLGQTYVNIGEVVEYRDGERSVQSPGHVQTVAIREELQTQMYRRSIEPRDLHH

SQSSPNLPSNSEHDAMFFYQNTLRNPHAVYSVLNDRCSNRSDKERQLFNMGLPLASVSQI

MNDYYIDPLLVSDSGGIGLPKNVSSKDISQNPYAQKPNVFNRSPYTEINPASSGLVLGMS

PQKSFAFQTRDKNYPPVLQERHFLNASKEYMERVQGVDRSSEQELENIYCRQNEERFSDK

IKQNYERYNENSSSRRNLTPQVEEIHPSPSRYSEARSMVEIDRHIETNNSCDGSLQRSMM

SFPRHVTLLPTAGHPDLHGKDSLRGDNGQSPRLLRNDSVDTMHKRPLTEEIETGAKRLRE

DVDTCSLPNLIRPERTPTLPVLEARDDQIVKIPVLEEPRKQPSMPAVVVQPDQNLEKNQR

SSLLGAMDDPEIDLTDGPINVSVELSTRKFYLEHYKQAYAMRRIPRGKALIMNVNEVVGK

SARRGTDIDRDNLHNLLCQMHFDVTVYNDADGLTAMEMVQKLQDFSQHSSHVSGDCTVVC

ILSHGEEGYIFGADGKKFELDAVFSLFDNTCCPSLRGKPKLFIIQACRGGALDRGVQYYP

DEHDGSDARVRQVPSMSDMLICYPTQNGYYAWRNRDRGSWYVEALVQIFMKYARCEDVCT

MLNRVNLLVSKKVSHCPNIDMDQMSQMSEYKSTLRMPHLYFFPGIGGCS

>XP_060596494_1 XP_060596494.1 caspase-3-like [Ruditapes philippinarum] 1463 bp

MDPATPQDDTDAKGFIGSLLKTFVCGGNVGARSKAFMDIGDIENEEEIEEAYKRVAPAFS

RTELLSDMYKMDFPRRGLAVVINNKTFSSEMQKKGYGVRTGTDIDATRMCSRLKMLGFEI

ERYHDATCKEIKMAFSNAAAKDHRDADCFVGVILSHGEKDKIYGVDGPIETNDIFQYFKG

FACETLAGKPKIFFIQACRGHQLDAGANLNVADAQRFDYEAEPEEEIRIPNEADFIVSYS

TVPGYYSWRNSANGSWYVQALVLLLDKYGTTMEIKKLLTQLNKCVAYCDRYMSNTDDPKM

HGMKQIPSFTSMLTKDLYFKPKK

>XP_060596493_1 XP_060596493.1 caspase-7-like [Ruditapes philippinarum] 1463 bp

MGVCGSTAQTGTGDGPIDKRDDVDGPVSKLNDRKEDKEKENVDKNSNQTNQHVSVKNTGN

TSDTSGNDERDVGLVPVGYFNQNSFTGLDIIDLDDNEEYLQAIQKAYQRPSPRLTRLDLL

ADVYKMDFPRRGKAIVINNEKFKDSPQRKGTWIDYSKICNRLQRLGFQVEGHNDVTCNQI

MAAFVNTANEDHSDADCFVGVFLSHGESNEIHGIDGVMKLSTIFELFHASRCKGLAGKPK

LFFIQSCRGEAFDDGVHLNVADAQGYVQQEPEVELTIPDEADFLISYSTVDGYYSWRNAK

NGSWYIQALVRVLDECGTLLEIQKLLTRVHKLVANEFESKERSRKRRKQIPSFISRLTKD

LYFNPK

>XP_060591462_1 XP_060591462.1 caspase-3-like [Ruditapes philippinarum] 1463 bp

MSGQGEQSQDVVDAKPSDNTDAKPSRGASAGKFPVTTGDGEPIMSAEEFFADEYKMDYPQ

RGRAIIINNREFDRGLGLGERTGTDQDASSLQMRFMDMGFEVDLYNNLVAGEMQKVLVKA

AADDNYNRISDCFACAILSHGEEGVVFGTDAKIEVKQLLEPFKGNNCKGLLGKPKLFFIQ

ACQGMQFDSGVNVNAVDAKGQMELERQITVHKIPSEADFLIAYSVVPGFYSWRNSANGSW

FVQALSDVLAKNWRDMDLLTIMTRVNKKVAYEFESKTGKDFMNQKKQIPCITSMLTKEVI

FKPKK

>XP_060580422_1 XP_060580422.1 caspase-3-like [Ruditapes philippinarum] 1463 bp

MDEIADDINTLVLKADETDAHVDTGGGTFDGSYTNENSVSIGGAQDVAHGEQTTQHDAKP

LDIEENVDDLEARGDDHGAASTDARMLGCLGSTDSAEKVNTILSVDPKLDVYDMNYKYRG

HALLINNEKFSIQLRRQGLGNRDGSSVDDDSMKKRLSELGFAVHAYPDLSAYSMKSVCSK

MAAKDYSECDCFICVILSHGEEGVVYGTDKPVEITELTKYFKGDKCPSLVGKPKLFIIQA

CRGYQTDAGVNVNVADAKGIGEQGKPAMVKLPVEADFLFAYSTIPGYYSWRNGANGSWFI

QALSEALKDFGKTMEIRKLLTLVNKKVAYNFQSFNPENKDFHKMKQIPCITSMLTKDLYF

STVKKKKLFF

>XP_060573657_1 XP_060573657.1 caspase-3-like [Ruditapes philippinarum] 1463 bp

MDREYPSSTELSVEWDGSQASLRNEGLSLHGSNQSLQMSSASPRLSPISNDSTSPYLLRK

VSADKDKHHVRPGAQSEPNEYNFNYRRRGVFLIFNNKHFLESTGQLEREGTDVDAERLEE

RFQDTGFEVRRYDDVTYARMTKLMQDIASLDHSDCDCFGCAVLSHGREGYIYATDKLVPL

EMLTQPFKGDKCPSLVGKPKLFFIQACTDKRTEHKRASPSRFVNGAQSTHIASRKVPVEA

DFLFSYSTVPGYYSWRNHQEGTWYIQALCIVIENYGNKMELMQMLTQVNRMVAYEFESCS

DEEFTEDLKQMPSIVSMLTRYIYFRPKKPDIRG

>XP_060560660_1 XP_060560660.1 caspase-8-like [Ruditapes philippinarum]DEDDED

MSATVLPNLESLHTEDPPGDEDTEDVQCDSLRLFSQEPSITAITDLHQRLIDIDEEVGDD

ELEGLKFLCRDHIPVSKLEKSQTALEVFDIMKKMVLIESNNVDYLLECLGRVHRLDLVRK

LGFQPDAVRNHLQNFKKLLPFRTLLMDIIEDLDDKETEQIRFYLRGKIRKKILNDKRSLL

QLFVFMEQEGLLSHDNLDLLTKVMESVRRGDLVDKIRIYQGLEPLQRQGPGLNPTVPSQQ

SQPTVPSGQPSNIARSVSAGTISFNSEENETTEVPDRRNIPQHLLDQIVTDLTPQVEDLG

IELGFTNHQLVQYQRQIPEPRERTLKLIQEWETNVQDKGKIVPALCYTLTKLGCKKSAED

LKRWTNSLNEPTVRPGSTEYQPTGPPGDGDNVCVSLREAQLDSVRQQVALSRLQQREPDV

RQAPPLPQGPAFANVPELRQVLPQTQVPVPGQVLPPTQVPTPGNMTEFRQMNGHQQILSN

VIPAQNDVNIINGGIAVYKQGDNYLVNAGISRELAPQMVPLENFIKGQGQSQQIQGVGNS

RQMALPVVHQPSRQIPELPVYPMNRRPRGWCVIINNKEFHVIQGDNTSKEMPARNGTDKD

AESLQHTFSKLGFIVRRRDNLTETEMMTYLIDIAHNVDHTDFDCFVCCILTHGVLNYLYG

SNGKLVSIKDLTETFQTNRCQTLAGKPKLFFLQACQGRDKMLGGSELERDMPRLESDQGD

REMIPNEADFLLGYATVPGYVSFRSRNHGSWYIRKLCELLEKYANKYDLMSILVEVNREV

AGANAAMDGGLYKQIPAPLVTLRKQLYFK

>XP_060598890_1 XP_060598890.1 caspase-3-like [Ruditapes philippinarum] 1463 bp

MKFNNPKLDRDGSDKDEARLFKVFTGLKFNVKSEKDKSKTQMLSILEDYAKMDHNQMDCF

VCCILSHGKEDEVVCCDTECATIEKITSFFRATNCKKLAGKPKLFFFQACQGSKRMGGWQ

VDGDGNRTFKEGSIPDEADFLLGFSTTSGYVSFRKKEGSAYIQTLCEQLQKHAHQYDLLT

IMTEVHRIVAEEVFEEENHKQMPTTLNRLRKQLYFNVKQLKD

>XP_060573497_1 XP_060573497.1 caspase-3-like [Ruditapes philippinarum] 1463 bp

MDNVDEARLATSQGGSDENCAKGKINKDRSEAELFGMNEYLPKKERRLVKYAEADSPPQE

AKETSETTPLRERLVKVFSRLKFNVDSVKDTSGTEMYSILKKYAEMDHTQMDCFVCCILS

HGKKDEVVCCDNECVNIENITSYFRATDCRSLAGKPKIFFFQACQGSKRMGGWQTDGGHD

CDSERTFKEGSIPDEADFLLGFSTTSGYVSFRQGGKGSVYIQTLCEQLQNHAHQYDLLTI

MTEVHRKVAKEVIQENTKTDHHKQMPATLNRLRKQFYFNVKQLKD

>XP_060593237_1 XP_060593237.1 caspase-8-like [Ruditapes philippinarum] 1463 bp

MDTILKEYANKDHTDMDCFVCCILSHGNKDGVVCCDKNCVSIETITGYFRASCCPSLAGK

PKLFFFQACRGIDHMGGWQTDGPIALDGEKNLKVGSLPDEADFLLGFATTSGYVAFRHNI

NGSIYIQTLCTLLEKYASE

>XP_060586583_1 XP_060586583.1 caspase-7-like [Ruditapes philippinarum] 1463 bp

MKKSCRGRCLILSNQTFYGPLEKDKEGVMRCILPTRHGTVKDAEDLKKLFTQLHFDVITR

TELSAQDIHSELLREASHSDHSQAECFVCVICSHGTKDGIYGVDGKTISLENVTKYFDGD

HCKHLQAKPKLFFIQACQGERADRPGSNVIDGGQDLSNLLERTHIGDQTDAAKETFNDTL

NPNTVPTKTDMLIACATHPGYVSLRNTQYGSWFIQAVIYVFQKYACKEDIFSLLTLVNSL

VARGRTALNQEGPRDVVQASLFTASFTKKFLFFPGVAADSEPR

>XP_060579252_1 XP_060579252.1 caspase-7-like [Ruditapes philippinarum] 1463 bp

MMMEEDNNTARVDTEVTDALKFETIKEKITSPVKKLLRLGNEHVLWDESDEYDFNNANRG

TAIIFNMEQVKTDSGELESRIGSSIDAERVRETFKSLGFSISYYENHTKEQVMDILTKEA

KRDHSDEDCFACIILTHGSSYYRVNSKEQYHTREDMIMAADTTMLTSDILKLFRDSNCKT

LQGKPKLFFIQACRGDTFDKGAFLETRDEVDAKPENSTEHIPCPIYKDFLVMYATPPGYY

AFRHPINGSWYIQALCRALSSAGVRNKTLLQLLTSVINIVSTEHQAQAPQYLGNKQTPCF

MSMLVKDVYFRLKEQKMSLPI

>XP_060579255_1 XP_060579255.1 caspase-3-like [Ruditapes philippinarum] 1463 bp

MEQGANEVHADDETEFPGSKTVSRIINSVTRSFQHIWGSDIEPPVWDETDEYDFNHPCRG

TMIVFNNEHFKGHSSRPGSSVDADNLYKTFTKLGFSVTCHHNQTRDQMIHILKKVAKSDH

SKADCFACAILSHGDEVHVVDDNDPGRHVREDLVYATDQIILTREIVNMFTDKKCKTLQG

KPKLFFMQACRGSTLDSGVNLSVMEDEVDAIVAETIEVTPSPIYKDFLLMNATPPGYYAF

RRPDTGSWFVRALCEILSHPNAGNQSLNQLLTSVIRLVCQQYQSQSTNLKISGKKQTPCF

MSMLVKDVYFRPTGRSLDKADQK

>XP_060565700_1 XP_060565700.1 caspase-9-like [Ruditapes philippinarum]CARD

MEAAQRQAIRKCRVRLVNDIQIEYMFDSIESHGLFTPIMLECIKAKESRPEKVRTFLDYL

QRRGPGAFDQFIQCCRESEQADVASYVLQEYQTIKDNTAQNSSNNSKGVSMANNIYLTQE

TQRSSQGQMETQGSSEMSVMETQGSSQISTSVMETQGPSQCVPSSSASVPQGINEVDRRE

EEMQEDNEGDAVMDEQPSYQGSESRIPGSTSRGTCSASSIPQPSLSVGHLSSSVGTVGYT

GNLTEEYKMDTDPRGFLLIINNRDFQRGLSTRDGTDIDCGKLRKLFLSLGFGVDVRQNLT

ALEMKQTLHMLSQHSQLDQVDSLAIALLTHGTEDVLYGVDAQHITVHEAFMPFTAEKCPA

LHHKPKLFIINACRGDAEDKGSLTFRHVDTTSTSQDISAVPSQSLACNFEDQYGTNTVSY

DVQSIPPVPRNAGRIPNMKDFLVAFSTIPGHVSWRHLNDGSFFIQGFVKVFEEYAHKSDV

MSMLVKVNNEVSKNINHQGVQIPAPQVMLTKVWHLNPKTKDFI

>XP_060574475_1 XP_060574475.1 caspase-7-like [Ruditapes philippinarum] 1463 bp

MALDPIIARHKLLLTTIKDEFDTDSSEYASFIEHFKGYVPRGQLAAKKELLGKFDALASR

GHLGPGKYDVLKKISEDSGNNNITTLIEKAEEDISVLRKSSDSNGQQGSSTAPPRSRPTG

RDATNDGPSSKRARTELDNPDPLYRRHGNHGYWDNTGKKGLLLILNFKENRAGTDLDVDR

LKDFFGPARLNFEILDPRQGELHDLSTSQVEAALESVQNKLNDKYHKPYYCFVCVILSHG

DEVGIKTRGDGRKSIDEIQKSFNNKNIPNFAGRPKLFFIQACRGHRDQTGQYVDDSDDAQ

EDENNANNGKDEADDLIHIPTDADTLTAFATTLGHRSFRRSGIGSWFITECMNVFTRDHI

QDHVEEMLITVRENVALNWVKDDGKTKQMPCVTSTLTKRLMFTKSE

>XP_060589701_1 XP_060589701.1 caspase-3-like [Ruditapes philippinarum] 1463 bp

MAATALHNKLLTDISDELVKDSSEYDNFVFHMTEFVKEGLLSSKKTILGKFGAMKTKGKL

DVGEYECIKKVAESSENIDIYNLIEKREAEIIEALEREKTGQKPDKSVSTKRKGTEVSSA

KKRKLEEETDQSDPLYGRDSRGRGFYDTAGKRGYLLILNNTHRRLGTGDDVSNLQYFFGD

ILRYRVDTISNDLSRADVIRALQSARDNIRSGATHEYFPFICAILSHGDENGLSFTDGAI

KVNEISKIFKDKIEEHNLRRNKLSGRPKLFFIESCRGGKAQGLHEVDDADDIVPQRNVDM

ADEIIYIPSDANTLLAYSSTKGYASYRDRSRPDTAHTYFIKALIDVFTEYYTTDHVEDML

ISVKERVAIDNASLYDTEKYTYTKQMPCTQSTLTKRLYLTPYNGQ

>XP_060598297_1 XP_060598297.1 caspase-1-like [Ruditapes philippinarum] 1463 bp

MSQVKDGCENSNVEDDGSDKESHENISSLEKAEKTFDELTLTENTELGRKENQEVEKEEL

CDEGDGLLKPLRKLGSAIIKKIGSKTIEIEPYTTTQENWTYSAISDERRKDDGETAKSEE

DIPKNDEKTVDITESDHYNFTHQKRGYLLLIINEKFYNQATRTGAHLDLIKMKEIAKKFG

FEILNNGAETNMTQSETLNWLNKAQTMDHSECDCFMFMISTHGLEQRNPRMGGGFDHALV

CADDKLIFTSSIVEMFNETNCPSLKGKPKIFIVQACRGNMVDHGSDLKISIDGGKFSSNV

KDRKAKSDQVSHEPSDVTDTKGAPDSWHKENLRGIETPYLKCHIDMLIMYAIPPGMLAWR

NTSAGSWMINYMHEILMDYDMEAPKCFLKDLTKISRKMARRSTDTPSMPEFHEKKAVPVI

EHMLTRDILFSLKH

>XP_060598298_1 XP_060598298.1 caspase-3-like [Ruditapes philippinarum] 1463 bp

MESHESVVCKNDVTDAFPDEVQDKRPGKVLTNTRTIQSRSIPSLQGKEKGYMLVIVNETC

GIHGEREGAQKDFQNIKEIGEHLGFQILNKDSYMNLTKDETSEWLKIASEADHTACERFM

IVVSSHGNEETNEERGKVKEHAFYCADSKPYFTNTMIEMFNKCDTLEGKPKIFIMQACRG

QNPDAGVEVPTTSVMSSGTKKDESDTTTHARDDIPDTKPGLDSWNEEIPYFKSYNDMLVV

FATSPGFYAWRNKLKGSWLIHYLHGALMTNRTPSNFLQVLTEVAGRMSERELTDNQLKCV

PVICHMLRSPIPI

>XP_060573228_1 XP_060573228.1 caspase-like [Ruditapes philippinarum] 1463 bp

MDDSEEYDMNHLFRGHAIIIANEKFAGDTNRPGARQDANKMQALFQSLDFLVTTYMNVTA

ADMMDIIRKGSKSFANLDADCFVLVISSHGLEKPVFDSSLQGGSYMYLPKDSTVWRHALI

GSDNKLIFVDEILKIFDDQENKDSYLKGKPKLFFLQACRSGHLSYKNGILGSYDTGTNVK

VSISDTNLTNIQPSDAGYHDEDFIELVNSVAGIKLSQEAENATDGMPVVGRVTYDYLKSP

VVETDPVTPVTPRVMKQSAYDIVPVPCPDDCLIMYPTVPGKYAYRINQTGSYMLSYMFHH

AHRNKLLRGCDILQYLTSVNNAMASTEIYFDPSAITVVPDEIKMLQESLPFKVTACIVHR

LTKTLRFQPRSTNSVWQELNARRTF

>XP_060588760_1 XP_060588760.1 uncharacterized protein LOC132744161 [Ruditapes philippinarum] 1463 bp

MMSGYKFDHKFRGTALIIGNEFKGQGLNERKGCLKDVEMAQELFKRLDFNVTSFKNLSAV

DMKKKMEIASKFTYNNDCDCFVLVLSTHGHEAKEIDLKKSNEATVWRHQVLAEDNVPIYV

DEIIDIFNDENSVGLKGKPKLFFIQACKSRGEIGSADTYDTGVDVNVVQGTKNPSVRDPN

TLRTDPRDSANNLYAIDLTDAEDMYKDWEKDPDALQLGAISRSSDQATDQDAEGEVTEID

TGGSGDSGQLPPSPFDIVPLNCPDDFLIMYPVMSGKIAFRKPKTGSDLIKFLKEKLNEVF

NGCNLLQYLTHVSHEIAGNKHKELKQILKTNLKKQKEARRTRESFDPLETFRKHLKGTIK

YLSSEENHEKLVTIIGEEHRSQLKGDPSVAEPYLLGVIDNRLTMQQMTEILKTILPFKMT

ACIVHRLTGVVYFKPKTKKSMMGILWKKSLPKETIVKKK

>XP_060580285_1 XP_060580285.1 caspase-6-like [Ruditapes philippinarum] 1463 bp

MATFSCGYSDNINQEVHKQGYAVFIVNINKNGDGGGFEFEERKGAREDLSNLQTVFESIG

FEWINNKCYWHTSLPVGRWDHRKGYRTERVTGNHTEPVPDRKECCIRCWIQALDHSRSKC

FLLVISSHGQELTGKETEIIFSDGHSVPLTEILDALSDKNCPSLRGKPRIIILQACRSNF

EDKEKVTKMQDTGVYVNVVLDQHQAVKASGATASSNVSDQQSDLVVDYTFQHTALRNMPD

NFMLVYPVVSGQRSQRSENYGSWLIYGLKEMYNKYYVGKPVNFFHLLTLVSGYIARERET

IVYKHKEGTNEIMLDEHGKQIRDEEQSGRKNAVNSFHRLCEPLIFYPKTKELPKMDEPTH

LLGNIDEPNFFGKKSKKGKKTLGQLLSEDK

>XP_060580283_1 XP_060580283.1 cell death protein 3-like isoform X3 [Ruditapes philippinarum] 1463 bp

MAEGGMPVGRNKPGYAALVVNIESVRSDLNDFRPRPSAREDITNLLHIFNKLGLTCINES

SSDHTNLTVGQWDHNRASNHTNSDVENCGACLRCKIRSKDHSESRCFFLAMSSHGREITE

TEVMFSDEESVKLTEIFDALSDENCPSLKGIPRIILILACRGNFKDPERNQARQDRGVPI

NISNKPKGKEDHKEPARSVRRTNSQTPLTSGASSAQECYFHHASLSQLPNDFLLVYPSVS

GQLTHSNETDGSWLVNAMKDAINEYLGTVPPPTSLDFFHLITAAAGRVAMIESHVLETDN

NGQKVGRNGKRKLDTKDEEKSGFKNAVCSHHRLSEPLDFDLNHKQDSCIPKPTFF

>XP_048751990_2 XP_048751990.2 uncharacterized protein LOC125663707 [Ostrea edulis]CARD 1461 bp

MEKSKKDILIKNRVGLVENINIEDGLLTELLSRQVLTQRTVKSIQARKTTFDQTEELLDK

ISLKSNGFEALSEALIANGQEDIVNLYLKAAQEDNETVKSEAKIESVISQESQTQTVVQE

PTRPQSSPPRLPPPGILLSHPTTKAVQLFQESDCSSARVIAQAPPEGVHTSQLVHLIHQP

PTDHVFHPIADQSVQSIQASSSAHDSSIICVSGNPSQAYDFSNAVFLDRISKNDKSFKEN

EPRHNMKHYIDDPNYQVLEGSTPKTVSTHIPLVFNNNIHPMPKEVVYVPSIKPVPLQRED

TLDAMTKPFGNHEENCESPAKRQRKEDKIEITVFESNQQNPNDPVMNAAAVQPVKLQGEL

DDPEIELTDGPVTVRVDHTTRQFYITNYKKSYAMRRLPRGKALIINVNEVEGKPPRRGTD

IDRDNLYHLLTQLHFDVQVFNDSDGLTAKDIAQKLEIFAADPAHRHGDASFVCLLSHGEE

GFIFGTDGRKLQLDSVFKLFDNSNCPTLVGKPKIFVIQACRGGALDSGVPFTDEHDGSSA

AFKQLPTQSDMIICFPTQAGYYAWRNRERGSWYIEALVQIFMKYAKSEDICTMLHRVNLL

VSRKISRCPQIEMDSMSQMSEYKSTLRMPHLFFYPGIGCG

>XP_048761694_1 XP_048761694.1 caspase-3-like [Ostrea edulis] 1461 bp

MEEAMSPENFTENANLDVADAKPISETDAFGFGQGTFKTHSTKPIATTIQTTSEFYSTRY

KMDYANRGKAVIINNKKFNPNTGLNDRNGTDVDASALCCRLSELDFDVDLFHNLKADEIK

QQLHKAAESDHSNSDCFACAILSHGEEGYVWGIDRMVPINDLLEPFKGNKCLSLAGKPKI

FLIQACRGTKFDVGVDMNVADAKGFMDIEPEFSLQKIPSEADFLIAYSVVPGYYSWRNST

NGSWFVQALSEVLMKYGQTVDLLTMMTRVNQIVANKFQSNTSHAEMNAKKQIPCITSMLT

KEVFFTSK

>XP_048763304_1 XP_048763304.1 caspase-3-like isoform X1 [Ostrea edulis] 1461 bp

MMTLCKSLESLTSSRQSLATSVENQESPYLIRKVSGDRRSVRPGAPSEPNEYNFNYHKRG

LFIIINNKNFHPSTGKESRDGTDADAERLEERFQDLGFDVRRYNDVSSSKLLQLMNDAAQ

LDHSDSDCFGCAILSYGIEGRIYATDGMLPLDVLIVPFKGDKCPMLVGKPKLFFLQSCRC

SNLEQSVEDTVSYKSFLSEESERSFSGMRRIPVEADFLFFYSTVPGFYSWRNHQEGSWFI

QALCIVLENYGSKMELLHMLTQVNRMVTYEYESCSDEEFTDDVKQMPCIVSMLTRYVYFR

LKKPDIRD

>XP_048776588_1 XP_048776588.1 CASP8 and FADD-like apoptosis regulator [Ostrea edulis]DEDDED 1461 bp

MNEIMNDIETDRPDLIPEEDAVPQKGFDYGKFLVTVDNGLDSNDVENLKFLLRGIVTDSK

LQKAQRGLDLFSELQNLGMIDIENGDLELLEECLYRIHRKDLLKKIARDTAMVEQRLRRL

TEELQNKKVTHIESTKMLSPFRVLLFEIGEDIDDEEFGSVLFALTSLVPKGQLSKMKSIF

DLFLFLEKKDEMSHVKVDVLLKVLKVVDRADLIPKVNKYLSNQNRRSSPTQYNGPSQMEV

EPPQVPQEPLNHPPPQLPPVIPGSSAMSNSPLSDGLLINLSDAISKSPDWKDIPPYLGLE

DRAILETVNDDQIMPSLRIFKMLQHWQRDIMQNDETCRVTLIGVLQKFRVTRGLQILGQT

VEGDEHIPDPNPTDLQSLYQPPPGQIPQSLRHPAPEPLPDIQSLRPNMSEMETQTSIMIL

PHGNLPIREQTEENLYVSNAEARVQLEQRQPPPSENNYSQSERSIRGNALDENLAQAMRR

TNLDDEMDLPSYRMDRKPRGIAVIINNMKFFKVPNDPESKDMPNRTGTDLDAEKLMYIFQ

KLHFHVRRYDNLEDYAMNRRLVEVSFEDHSEYDCFVCCILSHGVLQNIYGTNGRLVRISE

MTGIFRSGKCPSLAGKPKLFFIQACQGREKQGGIDIEMDSDLELDHGRNSEMIPDEADFV

LGCATVPGYVSYRSRSQGSWFVNKLVEMLDKYAYRYDLLSILVKVNEEVGRANARLEGGV

YKQIPAPLVTLRKKLFFR

>XP_048757091_1 XP_048757091.1 caspase-8-like [Ostrea edulis]DEDDED 1461 bp

MPQSGDFRATLLDIDAHLDSKDLEILQFLCKDIVPFSVLEKCKRCLDVFANLEQKGRLKD

EDSKFLQECFHYMQRKDLIRKLGGDPQDLERKIKTDGPSMLSEFRILLYEISEDCGKDEF

ESMKFYSRTTLQLTKKMLDKVTSCLSLFNVLEREELLTPIQVKTMEQLIETTGDEELLDY

LYQYKNRNLVCSPSEELGAIPGMKLDLKINSLPNTSPMVPQEHQEAPQNLTAQSEGANPQ

MAAYPSLTSMNPPAQDAMDTFMSRQLSDGQSPQEDHDFNDSLNRFYKMTAKPRGYCVIIS

NKIFHGKSDLKERVGTGVDEDKLENTFGSHLNFIVKTYRNCTSDEIMEVCRKYSGKDHGN

FDSIVFCILSHGTTGAVYGSDCRSVYIRDITSCFTASQCQSLASKPKLFFFQACQGLVHQ

SGLNNMGTTRNVNSGLQTDTLYESPRPQEPPNEETTSPPPNMIPDESDFLLGYATVPGYV

SYRSRSTGSFYIKTLVEKLKKYTLGPRSHDLVSILTEVNNAVSELHYSDSQGALYKQVPA

PQFTLRRKVHFDKC

>XP_048769780_1 XP_048769780.1 caspase-8-like isoform X1 [Ostrea edulis]DED 1461 bp

MSDNINRRCILQEIHDKMSHDDFRCLSYLSKDVIKNRKRMELRSSLDLFLALEEKTAINC

TSQNFSLLEEMFRLMGRKDVISVLTRCKQDAVQAEKTKTSMTDKRILLFNIGEETGEKEL

NYLKESLGKSGVRKRELDHVTDVWDGLDLLEERIRETDHLSFLRNTLGQVEEIKVLLDDF

IQKDEVSEGATGCQSDKFPVQEEGDMEITNGEEAKAYSNMQTGMSMDGFPSGFSSSRWSG

LGVYDKGIHPGRCVIINNERFDDESRLPTRTGTNIDRDRLHDMFTMFGFNISMYNNQSRD

EVMNVLECVQKEDHQDNGALVVCFLSHGSLNTISGSCGQDIGISNMTSLFRADNCPSMSG

KPKFFILQACQGRETQNVWHRGQTTIRAEFDDPRTPSLPEEEVTHESPSWGQLDIAASEA

DFLVAQSTMPGYISLRDEKEGSFFIQSLVRHLKEHSRRTTGPTSIKLGTQYPWMQGFTCV

QMKGHTLFKGEIITKMQKLSGNLKIFARTTVLEKLKFT

>XP_048756175_2 XP_048756175.2 caspase-3-like isoform X1 [Ostrea edulis] 1461 bp

MDDLEVDCCPVQHGPVSVDYTTMEIKRNSRLDARVVQAPNYMRIGDQSMQFHPRQYHKPT

YHSQQSLEIDGDVMSMEEDSPHLVNDDSRFLSEGVKPYIEPASLSFGPYNSGIKAKTNTS

RRPQEVPYIDIPSSVFHDTEYKMDHRKRGIALIINNMNYADVRQRPEREESIHDASAMEN

LLEHLMFTPVTQRMGHKNMVVNDVSSKRLEELIQDVTNPALNDFSDCDCFLMVVLSYGID

GSIMCCEGERDKLIPVEEVVAQFKADKCPSLAMKPKLFFMQLCPAFTYNIADSVGRSGPT

SMEPEVRKIPTDSDILVQISTLPGTYGWYEDDNALSWYIQGLKDTLQKNCYDPLRQGQRS

REINDILLHVNRLYISKLREQGICPSRHYSAPAITSTLTKDLYFLPKGIEAT

>XP_048760265_2 XP_048760265.2 caspase-7-like isoform X2 [Ostrea edulis]CARD 1461 bp

MEDADRRALQLNRTFLVKNIANPDDVADGLFTFEIFTEGMKDEVEVEKIKEKKMRKLLDI

LPRRGPQAFKLFLKVLMDTHNDHVVKQLTSGSSTTYTPPYHQQDEEEDLPQTWPDPTSLE

GKLTVKLCSSHNTRFQNGRYYKMGRNPRGRVLVINNKIFFGPLEEDESGKRRVTLATREG

TEQDERKIFEVFEQMNFIVDVYNDKKGEDIKDILRQEVKKDFHRTADCFVLVIMTHGTSG

SIYGTDGKTVSIEEIKKQFNGENFPAMADKPKVLLIQACRGDERDEGVRESTHINQEIKK

AEEMVKDLTLNSETDSHTEIVTPTMSHMVVATASTLGMVSFRNKMYGSWFLQAVAFIFAR

YSYKDDILQMLTKVNELVSRGRAKDSSQKKEFVAISEFSSTLSKSLYFCPGLVSEEIQQN

RIQQASS

>XP_048742450_2 XP_048742450.2 caspase-6-like isoform X1 [Ostrea edulis]CARD 1461 bp

MAGDQWKDAIRRNYSFFKNNIADPLDVVEYLYGETDPPILNENLKDQIKVEGTTSKKVGK

MLDILMRRGPRVPQFLYQAFVETENPDCAEKLAPYLLLQEKEALKKDPSEWPPPKKEHRA

MEKEPVTYLRPNDPCCCLQYNNTDEVYSLRKECRGVVFIINNMQFNGKMSIRHGTMSDRN

NLKELFQALHFEVIVENDLTKQEIEDKTRLISQSDKVKDSDCFIFIILTHGGDEGVFGVD

EKSVSVSTLTEMFEPNNCPGLNEKPKIFLIQACRGEKKEMVDPVGGSGNVDPVGGSGKSH

ADGDGRKEHVDTVDGPNVKSEPNISATKTVHSKSDFLIAYSTPEGSLSWRKTDAGSWFIQ

AIVWIFKYEAHLKDLVEMLGKVNNLVSKGHADHGGTHYVTISNYQSSLRKKLYFFPGIYG

EKRVHPNIR

>XP_048770132_1 XP_048770132.1 caspase-6-like [Ostrea edulis]CARD 1461 bp

MDERQRGILRKSRSIFTRDLVDVDTVCDRLYQKEVLSEGMKNEILSENTRENKVRKLMDI

IPKRGKRAFRAFFDVLEETFQHNLSDILCPGGQMDDLETKYNALHGFQSKRATDKPLLQR

SMSAEVRREPVESISEFQEQHKTCEFPLECKDFQKLAIVEEDFVLPSNWPSEQEHAQSLN

FKICKSRSSPASLREEWSRMYPMTRRPLGRLVIINNFQFDGEVDDERMKRSQQDTTSLDV

LFKQLHFQTKQHSNLTLEGLSSLLEEERTRDHTSIDCYVMVFMSRGDGERIYGIDGYRID

IDTIISTFDFEHCSSLRRKPKIFIFQACDDVAAKSKLLGAEKKNYEILTSSTTSEHMRTD

MFVVNAVQDVAVSYHGIFGSRFIQSFVYLMRNLAFNEDFEEIVRKINHLNSVSEPEIIKF

ASSFSRRLFFNP

>XP_048747578_2 XP_048747578.2 caspase-2-like isoform X1 [Ostrea edulis]CARD 1461 bp

MEQIHKDALRRTRVALVKDLDVDVICDEVLTHDIFTQLMMEYIMAERTRIDKVRRLLDDL

VRRGPNAYRAFIQVLNATGYKFLADKILQNEDIVRREKEPNTSFPVQATNEIFSEIHPSS

ISSGFTTSNLNNNARNSLIGPVNNNMQQNPVSLGSSESNLSSNLSSVSQSCSEPNSKTGV

GGLTVHEMASTLSTPSESTDDDTLDLLSGSLNPTSMDVDDDVDSRTEELLLRDTGNEHGH

PSISEGYPVQPTDTSHTAYNFDPPFTMMSFYEQSKSLDKCYMMESTPRGIVLIINNKDFG

QNHKERTGTEYDCQKLTELFRNLSFQVLEKENLTAMEMKEELKAFTQNRQLQAVDCLVLA

VLSHGNSDKIICGIDSEGIDIYEHVFPLFSAKNCRMMRGKPKMFIFNACRGVKNGESFDT

FRPQTSRTTIPEKDQLQRDGPNIEPSDSQVLDMQDLLLCFSTFPGYTSYRDPNQGSWFIQ

EFVSVFQEQCSTEHVMDMLTEINRRVSQKKHSDQAAEMCVQMPFPSSSLTKKWYINPL

>XP_048776071_2 XP_048776071.2 caspase-3-like [Ostrea edulis] 1461 bp

MDNNKMTSTKIPTEARDLHNQLVIDLHDHINSKVDEHDFEKLLHRLKLKLPPMALLGCKN

VIDVIEKLQKKGYVRPGKYEEFKKLIECIDIGWVDVIDDRAEKIIKIIDDANKSQKIASP

MEDQRKLDMKRKLEDVEQEAPSQDPDGPTHASKKRKSRSEYSKSDEQVGLAIILNFTKGR

LGSVVDFKRIQKFFRSTLKFECHIEEDLSKEALQEFLEETANMFNTKRTAAKYYCVLVFI

MSHGNEEGILMDDGKHISLDDIVAYFKNDKLKYIVGKPKCFFVQACRGSSPQGAVDLSDD

KEEMMIDEVVGKVSIPTDADLLIAYATTPGYYAYRRPTIGSWFFHNVIEIFSAHYTEEHV

EDMLIDVKEALAYDPEWQTNEGYKQMPCTWSTLTKRFFLKSSVE

>XP_048742245_2 XP_048742245.2 caspase-2-like [Ostrea edulis]PS50137	DS_RBD	Double stranded RNA-binding domain (dsRBD) 1461 bp

MSERSDSLLKTPKSSFKQLDEIAQKTRVTILSQIESHPEGFLATLTLGDVKYTGIGSNKT

SAKDKAAAEALEEWKDKATPEKKVEKTPATRINELCQQNIEIKQRYEELEKDGDDFVCKL

VMTLDNRENMEEHIGRGRSKKEAKQNAASCGLQKSHILQPFTGGPGIPTEGRCYECKPKQ

TGYVLVVYFSRNRDWADIDVNNIKGFMESKLKFECDVVKDPTKGELTEVLESTSRHLNDH

ANAYYCFTFFLMGHGSQFGVRTADKGKKKMTITVEEILNYFKNDKVSNFTGKPKVFFLQC

CRGELHQDTVVQPDSDDEDSDKLRVPADGDILIAHATTEGYKAYRYKDFGSIFIYNCIKT

FETNYTTTHLEEMLIDVKAVIALDPRWRRTKEHSQMPCHWSTLTKKFMLIPADR

>XP_048728167_2 XP_048728167.2 caspase-7-like [Ostrea edulis] 1461 bp

MSAPRSANNDRISVSCSELYEMLTDQSVEISSSRRFLEDQEERDFGGSRHWKGESSENLH

HLPKIKHNRASPDISEFIVGNRSKSFPFMKDKPISSPTTMRKLDARTLEKELLTQFKNRR

LNPHSHVQMGMNKSPRQRLQKSFYPSAKQLAVQSLEQFQQNADTATTTTADDRFAVDVIE

SSSNMHTTVLSEDCILDGTPSKTSTLPSRDYFDGLSNPRWRSLNDAADSGFYSMTRKPRG

YAVIINNYEFYKEMIAFTSSGPQTLRVIQKGLRRDGSIEDTDKLSRFFRDELHFIVRGHA

NVSSYDMIKILNDVSSADHDEFDAFVCCIASHGRRGTLFGSDCREIKVEDVTGLFNGNHC

VSLSGKPKCFFIQGCRGNHISQKGVLYGQRNESSQDEHYIPLTCDFLIGYSLIKECTEFH

ARQSGSLLYKTLVKYFRKYCKRYDLINIMVRVKKALAKLQENSVGAEDIFECNFRAQLVF

>XP_056001291_1 XP_056001291.1 uncharacterized protein LOC125664819 [Ostrea edulis] 1461 bp

MNERMEIFKLAQSYKIKWELFLDSKVHEVLTAHAQSLGCELDLLILPLFVSVASSMTSGT

KIQVTLSWIERPRLCGVVAARRGERKTTMLNYIKSCCRQVKCHGGDTDDTESVPGIPTID

EAFTENIPETCKGDRLFVLDDLDIVLKNVTEMPDIWNLRTLQRLFDSRCSDETDDAIKEN

VNVIAAAQGVVISDIVNSPDPGRLLDRVIVYCARDSEDMVISTKTPDSLGKLMQRIYDLH

VNQNVTYTFSEEATQEFQRMSGIVNAIKKQCHNDELKRGCCNHVMSQMARLSAISCALTS

SIKETKADFTINKTTAQSMFVVSLYLLRQKCLLLHGSDSTTYSELFHSILNSETELLPTA

DTSPRESEDDVIEITLDRNNEPVARIIPKREDNMEISETSQATDVRMWTMDNKDYTVNQN

GRETSSVYYPAVQNLPQEVATQQKRQRLMNHPEGPNTEHCQQYEHNHPTHSEDNKQIHFS

NTLQPRGKFRPPPPPLKLRPHQLHVNRRFIPAPKRLTDRTQSKTCVFTCSDSLFVHICCA

KIRKCLLTKGLSITSSYACQYRLFPPVPLTQRTSSPRTTHPSWAAVKFFERLEALGYGEV

TVFRSQSVKFRKKALEEMSEEARNILKYQIGITEEEYRSAMTKEEDIQGPVQEYNYVFTV

HQNGDIETEPMQHVETEQNVQHSAICEILTTCYRIRHLYVICLYKRTVRYCLIEIRPRKI

RISSSGRAFFRNIEYRINISTFLIMQERNMENHNDLDNTTEVQHATSRVQFSPVILENSV

SLWSDTEPNRNQNDCVYRFGHNKRGYAILIMNNEFDNEDENLPGVQSDIKNMSELFNSLE

FDVIELINLTNDQLITELTGIQDAVTNDCDCFVCVISTHGGEDPGYYIGDTEKLVTKTEH

YILNSKGAVWTRYIMDMFGDEQCRALRGKPRLFFIQACRGGEVDKGIDLLEDDSDDIEDD

SKNHAVFQIPCYNDSLVMFSSTSGKYAWSEDFEGGWLLSALQSVFPSSTDTDLLSLLTRV

CGEVARREAYIPDNERAHRSKSAACVYHKLSKDIYFPKSNKKD

>XP_048760572_2 XP_048760572.2 uncharacterized protein LOC125669811 [Ostrea edulis] 1461 bp

MDLEQQALQGNDEWERFEKSETKPDGFKAFKKTLKDFIGLKSKKKGGHGVPYQTQMDDDE

RSHVISDTKKESQTASNTPEVPQKLQNNELSWSYPESSNPRSQQSHNEIAEDNTYSFRHE

KRGFAVLVINSEFDTQSDRENAVWDIYYMRKMFSEMGFDVKILQNLTSNDLLSKLKRIQQ

QITMESGCFACVISSHGMEGRAKVSGLKGREPFVRTEHFVYTKDGVVRTSELLELFNDCN

CRNLKGKPRMFFIQACRSRLDVQKDDEVDMGVEIPVRLVHSTTNKDTTSDAGGIYDDYTD

RQSEDFFVASLDPQYENIHVYGLSRKYTGNYIQTGNRYSINTRSPEAREPPRRTTTYKVQ

KPETKPENIEVFQIPCFNDFLVMFSSAAGTIAWSDSGKGGWLMYTLYHVFNDVTYTDKDL

LTTLTGVCGMMATEMQTYCPSNLRNDKAKSAACVYHRLSKDIYLQPSSYV

>XP_056017527_1 XP_056017527.1 uncharacterized protein LOC125669447 isoform X2 [Ostrea edulis] 1461 bp

MAQFFKRFLHKELQRHVKIRNDVADDREYDFRHEVRGIAIVISNGVFSPDTGSVTEPRSY

AEEELCKMREMFSGLGFIVMLFKDLTARQMHDVIQTVTSQEMDFFLEKSDAFACVLASHG

NEIQEDSPADPAVKVFHHIIYGTDRAIRTHLLVDLLHDNRCKKLQGKPKLFFIQACRSRF

NDPNRFDSGVKILTSNKRRIPEVGHDKKESISINSTTSFADTKSPISDGLCGDKIKDGGR

KSDGSESSIEVIPCLEVLHEVTHDVSEDPSIAQRGIKWIKKLVFRKVDRIARNDVHDEDT

MTTVMTPCHNDFLVMYSTSQGKASFGREEKGGWMIHVMHRIMVQHVEFIQKDKENCLDFL

SVMTNVMRHIGFYFETDTGNPNTSGYKAAPTLQHKLSKDLIFRNKYVS

>XP_048734212_1 XP_048734212.1 caspase-3-like isoform X1 [Ostrea edulis] 1461 bp

MWSTLDADFVQGSDRESSGDRREVHETELGPLFVSYIDSAVVPTEDCNIVDNEEKNEFVV

VDNNKPGLALLVGFEYFNNGEQSRPGVKKDLENMYELFHTHLGFEVQVKKDLTAEECMEW

IREVTSKTELLQRISCVVMVISSHGSEDEVKSGIVNPDFRRQEVKFYQHRFSTKDGSIQT

SEIIDKFDDGKCRALKGKPKVFFIQACRSRTCENVSAYDVVDGGVSIRIQPEGSSRPHPS

TQHVLSSQNRPVMQNEEAHSHMDAKGVKCSNIRAGNKIFTEMSGCVRAPMEEEVTLAPPP

CLHNCMIVMASPPGKAAWSDTFQGGWLITCLYKVYMEHLKKGHKIRMIRSITQINRLMAL

SMHSVTDDPQLNSMKCVPVMTHMLERDIVLVPKNLQ

>XP_056010880_1 XP_056010880.1 uncharacterized protein LOC130051874 [Ostrea edulis] 1461 bp

MSGKVGLLYVKNKNTVQLLSDEITTLLGAENIVDSGVEEQAIKSGELTPVVKKVGKNLSN

VNSLIVITDNAFWDRFWSEKSNKVHNKLEELLCDDDGDGVTLVLIVGCNIDQGLIETLVP

SSSHAHVISEEDLKKYLSKERKPPSVCHMVVSIKENNSESIKELSKQIVEKYGNQSLEDE

DVCKDKVDTPTQTPTLTFPVTPQWPTTKERVVVQTAGPNDDASENPDIDFSVPSMKSDDV

DNNETRYDISRPGLCLYIGVLEFDPEDSKVKLAGRKSSMLEFEKMKDSFECLGCFVKRKS

NPTDTEALKFLQQGMEWCEKLKPSYFVLIISTHGQEVPQVNKEDKSAKLHPSGIKEKGQP

KKEMVHQLFFNNGKSLLTRDIVKMFDDCPSLADKPCFFFIQACRSRFGIDGHENFDPGVM

VDVFTPRSQIEAAQITPDTEDASGNPKLLDKTDAKRSTAKLTEEDKKARLQQIEKWNKNA

KIATLYTHILEYKNWLSNQTGASQTRSQDTSEETAKLAEALTVEGFTFQKVQEILETCKG

YEVDILDFFFPEMIISDPAPCTNDSAIMFASAPGKEAYNRGNKGGWLIINMEEQLKESLK

DNPENVDLLAELTKVSGRMAYKYETDTDSSESSGHKSVPCLYHKLCKDVIICPQELARKK

KLLENHDNSEYSEFFKWA

>XP_048765195_2 XP_048765195.2 uncharacterized protein LOC125673006 isoform X1 [Ostrea edulis] 1461 bp

MEVDSEEYTYKFEDKRRYQGLALVIANFTKGPHKRNGAEQDLKYMKKTFKRLGFKVFYEE

NVTTPEFLSLLNHYSKSDTLKNYDCFAFAISSHGLEREERKGGSLVHHHVIQMFDDEFID

TSQILDYFSPKKCKIMKGKPKLFFLQACRIPESKASKDHLDAGIGFDHGSRTHPHHGGRE

LEKGTAYRDKTDYRESVEVSNEEVSEGDVSDTDVEDDPDLVIEKPPSSTTLGGQDTGDTL

EVLDQRQTEIITNLKRLQNQTWQLGRVYTATGTNQIKESNYQANQIKESKYQATEGEDEQ

KIPLQESKKTANESGRMEGVGLHLPTESRGSGDELDPPRGVRLDQRAALNITVVPCHNDM

LIMFASPQGLYAIRSKDHGSYMLKILYESVERYYGHGKLRNNKTNFLEVLRDVTAKMSAK

TFFGSKEYTIVPCIVHQLSKDIVFRQGQMMVDSKFSLKLPSSLASIIR

>QNL15268.1 caspase-8 [Littorina littorea]

MTSQDRMQTDAQAQDQPDFMSAVMKIDNELHEEHVESMKFLCQGLVVSSKLKRVETAHEL

VSLLQSSNNVKQNDLFLLADLLQYIGRIDVLEEIGFEEKEVIFRRRQQGSSIKPFYVLLF

QIAEELPMEDVKKAAFHYGKVPRSQTILSGLDLFTVMVQRQSIRPENVGLLKTIFDSLER

KDLVELIERYSATRGDSVMRDLEDNFVRTMSFRGFNYGNMFSGGRQIPPGSGFGPSLDDF

DVRDAPLPAPSQASEIGNTSMNTEPPSEGRMPPSAPPESLPYSIGSHGQVPEVPERNVAP

SSHTPANNSHAQSKPPDGYDERIIPWSVLEILSQKIHAKYWPDVTEFLGVHIDTVRWDST

KTNRQMNIVLLNRWLAQTETQEADLLESRVKLIRALNYADCKLLAEQVAKLLNINYNNVA

DESMSSVGSGSHLNAVSSSASAAPVTTDSMSQHEIMEQEVDPAVGAARVEVTLVGRLPAV

SPIPYYNMSNSPRGICIIINNRNFFRDRPEAKLMVAREGTEVDKMKLEKTFRGLNFLVSS

HDDMTDSQMIQLMNEIAHMDHSHFDCFVCCILSHGVQGSLYGTNGMTVPVKDMTGPFRAQ

ACPGLAGKPKLFFMQACQGKEKQTGHLNDIQMDSNLEEDAPSGELIPNEADFLLGYATVP

GFVSYRSKSRGSWYVTKLAEMLQKYGYDHDILDILTLVNFEVGKGDALIEGGVFKQSPAP

MYSLRKKLIFSRLR

>AYK02335.1 caspase-3 [Haliotis diversicolor]

MTHRPSQKVSAERRQSNVFNEERSLAMEHIYPMKHKRKNIVYIINNFIFGHGERKGSVVD

EESLTKTFTNLSFDVQSFQNKTRADMVNLLENVSRDIHKDTDCFVCVILTHGEEDMVHGT

DGPVEVTKLLAPFKGDRCPALAGKPKLFFIQACRGKQFDDGADALSVKNDGPEDDEDEMD

VKDETVTFRIPRNADFLVAYASVRGFYAWRNEKDGSWFIQALCKVLEEHGQTLDLMTMLT

RVNHTVAFDYSSKGKIKEKKQIPCITSMLTKDVFFRKT

>QHY92978.1 caspase 8 [Haliotis diversicolor] DED DED

MSMDVEADAVAERGFHTMLLKIDNDLHQSDLQALKFLVKGNLHKNKLTKMSTGHELFAAL

ETSKLIEASNRWLLVEMFDKIGRLDLIDYIDDEITEDLKRTHLPTGEISDYRLMLFDTAQ

DLTVEDTQKLTFLLHTLPKAKLGKVSSGLDLITLMEEAKIIAPDHLDKLRQYLVTLTRHD

LVRKIDLYTERRHQRVCVTATGASMGHDTGGDVRMTPVPMAHPGPPRHTADNGVPESIRL

QISKDLCQHKYMENLATLWTHLGYKEDFVNRLQSGFQGLTNGERIQKMIECAPSLQSADY

SQALMRLINAMRFSQMFQLSEQLVQQFKAHLPREMVYDQGTGVLQRFGDENLNQYVSTGE

ATETAQPGTDQSLNSINIHDKQSTIPTYKMNANPRGICVIINNFEFWPGSEGAPDLKNRT

GTHIDAAKLSDTFSRLHFDIVHYNDLSADAMTQRLRTIGQEMVKAEHDCFVCCILSHGDT

AGKVYGSDSKVVEVNKLKAPFRPDCCPNLARKPKLFFLQACQGKEMQEGVHVQSDSDLKT

DGPTVMVANEADFLVGYATTPGYVSYRSKSKGSWYINKLTMMLDKHAHTEDLLSILTKVN

NEVGMELAPLGGKQSPAPQFTLTKRLIFRKLE

>AYK02334.1 caspase-8 [Haliotis diversicolor] CARD

MIKSIHNVEERFKLYAQKCAKLLQLGQCNPKKVQRDISCTENYIMQVFREREQFIIGKED

KRHNSHRDTLRKCYHHIISDLELVNSSVLDRLLEDDVMSVSDKDSIIMKGTKKSQNEEFY

RYIYGKVTREDFHNSLLPSLKKDHGHLAEILSKELVTLDESTDDVQCGTCEALKIVQVKR

LATSLFKAGVIDNAVHADLRSDSITNSQKWKELLEHANVSHVHIIDAMEKEYPDLYHDFR

RRNITTLKCFCGAPDTLSVVSDFVSPYRRDVIANSTTGAGDVAALDDLQSFLSLPIQDTE

SAYPMNARPRGICVIINNRDFQESQDDAGGRYRGQHGLGKREGTDVDAKRLEYIFSKLEF

TIRPFTNLTDTQMVEELQCVASTIDHKPFDAFVCCILSHGVKDRVYGTNCRDVKVSDLTA

PFRASACPSLAGKPKMFFIQAAQGNIRMQGLNVEDGVELPVPCERTPVDASVVRDDPPQE

KPDKQKWRSKGSPDERNIIPDESDFLLGYATSPGYVSYRSKRHGSWYIIRLAEVLEKYGD

RLDLLSCLMMVNDEVAKGNAMTKDGVFKQVPAPMFTLRKKVNLTSDPMWFRSLSTINE

>ABY87390.3 caspase [Haliotis diversicolor]

MAEGLKIEIEAGDGKSNNLVMAAVVTGGVDNKGKGQPKSEGIKSDLSATKIKMKGPGKDS

QHRDVKMLGIVEGGMACQDVSEDTDGRGQGQEARYKMDSEPRGLCYIFSNKTFEDKPKQT

RIGTEKDRDALTEVWKKLHFEVESFDDFKAEGMETRIKELAKMDHTVYDCVVVIFLTHGR

DGAVEGSDNEWLDVSTITSYFMPSVCPSLASKPKLFFFQSCRGKEIQKAVPVQGDDGGSD

ADATETIVQANKMVAPEADFFFGFATVAGMQALRNPETGSWFIQSLVQALQASTKSDLDS

IMTTVRGKVNNRECETVDGKFLRQTAEVRTTLLKKLYF

>QLI42792.1 caspase-3-like protein [Onchidium reevesii]

MAGRGDFTDTPHGGVQDSTDAHGVHSTMGQPSSVPAATQSKSDFVSDRYKMDHPKRGIAV

IINNKDFLPQTGMGSRTGTDVDAESMHDLLNEMSFEKIIPKKNVSVKDMKELFWDVSKYD

HSRSDCFVCVILTHGEEGYVYGTDGKIPIDDLVYPFKGHNCKTLAGKPKLFFIQACRGQE

LDGGVDVSDAIGEEEAMEEEVVIRRIPTEADFLMAYSVVPGYFAWRNSSRGSWFIQAIVS

VFQANWKTMDVLTMMTRVNKKVAYDFQSNASSQSMNRKKQIPCITSMLTKDVYFTEK

>QMS54626.1 caspase-9 like protein [Onchidium reevesii]CARD

MEPRDRELIQKNYMSLVEIIKPRVDEVCGELLGMGIISSGMKEDITEVPFPSERKAMMLL

DLLMTRGNHAVEALYKAGLQQQLYGLADILRPENQPHGLIHTTGLQDSTPTASAADDSPD

EPLPDTWPDQAAIESIGTLRKVDKKQSRMLKLYKESLEVPGKGMIYRMKHRERGRCVVFN

NYTFDHLQPRDGSNQDRKGVEKLFQALGFDVEVYNNKTCKKMWELMEEEAKADHSQADCF

VMFIFSHGHSGVVFGTDGKKDEKEVLQNAIGIKRIRELFGNLDSLGGKPKLFFIQACQGD

LKDKGQEVHDGGTSPMKASTSPDLPETTLSLQQQKRLNEENIKDIKLTAKNLEEDQSDAM

GEKRPNQGDIFIAMATIPDHLSWRNTVHGSWFIQAVVYVFGKYAHKHNLDRLMNMVKNLV

AKSETRIKQYKQIGEIISTLNKDFYFFPGLCEKEPNC

>XP_011419292.2 caspase Dronc [Crassostrea gigas]CARD

MMDKAKKNILIKNRVELVENINIEDGLLTELLSRQVLTQRTAKSIQAKKTTFDQTEELLD

KIALKSNGFEALSEALIANGQEDIVDLYLKPPSEESEKLKRDEKSESVISQESQTVGPEP

TRPQSSPPRLPAPGPQVSQPITKAVQLLTHTESASPRVLSQGQQESQGVPTRQLQPALHH

PPPDHVFHPIQTHSVQSVPVQPSSVDSSIICVSGNPGQPYNFSNAVFLDRVSKSDVSFKE

NQQRHNLKHYINDPNYQVPPEVAPTTVSQHIPVMFNNNVHPVPKEVVYIPSYRPASLQRE

DTLDAMTKPLGNHEANCESPAKRQRTEEKIEITVFESNQPSQNEPPMMTQSSLQPVRIQG

ELDDPEIDLTDGPVTVRVDNTTRQFYITNYKKSYAMRRLPRGKALIINVNEVEGKPPRRG

TDIDRDNLYHLLRQLHFDVQVYNDKDGLTAKEIAQKLEMFAADPAHRQGDSSFVCLLSHG

EEGFIFGTDGRKLQLDSVFKLFDNSNCPTLVGKPKIFVIQACRGGALDSGVPFLDEHDGS

TGSKQLPTQSDMIICFPTQAGYYAWRNRERGSWYIEALVQIFMKYAKSEDICTMLHRVNL

LVSRKVSRCPQIEMDSMSQMSEYKSSLRMPHLFFYPGIGCV

>XP_011449817.2 caspase-2-like isoform X1 [Crassostrea gigas]CARD

MEDADRRVLQVNRTFLVKNIANPDDVANELFSNEIFTEGMKDEVEVEKLKEKKVRKLLDI

LPRRGPEAFQIFIKVLQDTNNDHVALKLKSGISPQNNSHHRQEDGEDDLPQTWPDPLSLE

GRLTVKPLTCPASRFQKTGCYKMGRRPRGRVLIINNKIFFGPLEEDENGKKRVTLATREG

TEQDERKIQEVFEQMFFVVDVHMDKKGEEIKALLKEEVKKDFHHKADCFVLVIMTHGTSG

AIYGVDGKTVSIEEIKMLLNGENFPAMADKPKVLLIQACRGETKDEGVKETAADIEKEIK

STGEILKDLSLNTVSDSKSESVTATMSHMVVAMASTLGMVSYRNKIYGSWFLTAVAYVFA

KYSHTNDIHQMLTKVNQLVSRGRAKDNKQNKEFVAISESSYTLSKDLYFCPGYFIEEVPQ

SRIQQASF

>XP_011414267.3 caspase-2-like isoform X1 [Crassostrea gigas]CARD

MANEDEWKNSIRRNYSYFKNNIADSLDVAEYLFGETNPPIITDNQKEQIKAENTTSKKVG

KMLDILMKRGPRVPQCLFDAFMETHNEECAKKLAPYLLAQEKHALKREPSEWPPLHEEHF

DMMKEPVTYLRPNDPCCFLQYNNTDEVYSMRRECRGLVYLINNEQFHTLSQRDGTKFDRD

NLKKLFTDLHFEVIVENNLTAEEIRKTTESISKCDKMKDSDCFIFIILTHGDEKGVCGID

GISVPVTTLTEMFEPNNCPEMNEKPKIFLIQACRGDKREMVCSSGGPGENKSQGDGGGDR

VDGMNAPDINVEREPNIYATQTVHSKSDILVAYSTPEGSLSWRKTDAGSWFMQAIVWIFK

FESHVKDLVEMLGKVNNVVSKGKARDVGSHFVTVSKYESSLRKKLFFFPGIYGDKRVHPK

CI

>AKP95634.1 caspase8-2 [Crassostrea gigas]DED

MNDIETDNTIWTDAVSEKGYDYGEFLGCVDSGLDTSDVDNLKFLLRGVLPDSKIQRAERG

LDLFTELQNAGVIDIDNADLELLEECLYRIHRKDLLKKMARDTNLVEQRLKRLQADLQKK

KISYMNSAKFLSPFRVLLFEIGEDIDDEEFKSVMFTLTSVVPKGQLSKMKSIFDLFLFLE

KKDEMSHTKVDVLLKVLKVIDRADLIPKVNKYISLQAKENNYSQNSQPTLMEVEPPQVPP

TGAQNHPPPQALGAGGGIRLASHAPLSKGILIRLADVIAKCPNWKDISPFLGLEDSSITA

VINDDHLRVSLRILNMLEYWQMNVMQNDETCRVTMIGVLQKFRITEGLQILGETVEEDDH

IPDPTPTNLQSMYRPPVGDIPEPIRHPAPEPLPTIQSLRPITSTSEIETQTSMMILPNSS

QPVRDENEENLSVSNTEARVQPVEREPQPSENNYVQSERSTRDDRVDETLAVAMRRTHLD

DEMDLPSYRMDRQPRGIAVIINNMKFFKVANDHESKEMPDRTGTDLDADKLTYIFQKLGF

LVRRYDNLEDVAMYRRLVDVSLEDHSNYDCFVCCILSHGVLQNIYGTNGRLVRISELTGI

FRSRKCTGLAGKPKLFFIQACQGREKQMGIDIEMDSDIEADNGRNSEMIPDEADFVLGCA

TVPGYVSYRSRSQGSWFVNKLVEMLDKFAYRYDLLSILVKVNEEVGRANARLEGGVFKQI

PAPLVTLRKKLFFK

>XP_034301971.1 caspase-8 [Crassostrea gigas]DED

MKMTHLGDFKAILLDIDAHLDSKDLELLKFLCKDVVSFSVLGKCQRCLDVFANLEQKGKL

KEENSKYLQECFHYMQRKDLIRKLSGDPQQLENRIRANGPNYLTEFRILLFDIAEDCGKD

ELEKMKFYSRTTLHLPKKMVDKVKSCLDLFNVLEKEEKITNIDVNTLEQLIKTTGDEELL

ESVYQYKNRNLVSSPTDEMNVEPGMKLDLHINHFSNTTAAVTPQAHQGPEHNLAVQTEGS

NPQMGAYPSLTSMNPPAQDAMDTIMPRQQSPAQSSEENLDSHDDMFHYYKMTSKPRGYCV

IINNQIFHGNLGERKGTDIDEEKLKEVFGNHLEFKVEVYRNATSEEIMRVCQEYSRKNHR

QFDCLVFCILSHGSTGAVYGTDSNSVRIKDITACFTASRCPSLAEKPKLFFFQACQGLSH

QSGLQTIGTTRNVDPNTQMDATQEPPRSEVNPNEDRTSSAHKMIPDESDFLLGYATVPGY

VSYRSRSQGSFYINILAEKLKKYALGPRGHDLVSILVSVNAAVSELHYSNEDGHMFKQVP

APQFTLRRRVQFDKC

>XP_011432762.2 caspase-8 isoform X1 [Crassostrea gigas]CARD

MDDRERSILRRSRSLFIRDLVDVVTVCDRLYQREVLSEGMKNEILSETTRENQIRTLMDI

IPKRGQKAYGAFYDVLEETYQYNLASILYPGGRPNTTPYPPKAAVEPRQYVGLPKTPLHN

WLTGAKREPMESIGEFQAHRKPVESLDDDLVVPSTELHPEHHKSLKEKDFRKLRTTEEDF

VFPYDWPSEQNHAQSMDFQIRLSQRPQGDSKEDELAVYRMKEATRGRLVIINNFLFGDEV

DKERMARSQQDATSLDVLFKQLHFQTKQHSNLTLKNLSSVLEGERTRDHASLECFVMVIM

SRGDGERLYGVDGYKIDIDTIISMFDSEHCSYLRGKPKVFIFQTCSDAPKTGTAEFQKKS

KTPTSSSSLTLEDSGDLRADMFVVKATKDVEVSYNGVFGSRFIQSLVYLLRNLVCEEDFE

EIVKKINHLNVVSEPEIVEYTSSFTKKLFFNP

>XP_011414268_2 XP_011414268.2 caspase-6 isoform X1 [Crassostrea gigas]

MANEEEWKDSIRRNYSFFKDNIADPHDVVQYLFGQTNPPIITHDYLDEIKAERTPSKQVE

KMLDILLIRGPMVPQCLFDAFVETHNEECAKKLAPYLLAQEKHALKKTTSEWPPLHEEHF

DMKKVPVTYLKPNDPCCFLQYNNTDEVYSMRRKRRGLVYLINNEKFDDLPRRYGTEFDRD

DLAKLFTDLYFEVIVHDNLTAEEIRKNTESISKCDKIKDCDCFIFIILTHGDEKGVCGID

GESVPVSTLTEMFEPNNCPELNEKPKVFLIQACRGDKKQKVCPSGGPGENGSQGDGSGDH

VDGMNASDGGWELGISATQTVHSKSDFLVAYSTPEGSLSWRKMDAGSWFMQAIVWVFKSE

SHVKDLVEMLGKVNDIVSKGKAKSGWGHYVTVSKYESSLRKKLFFFPGIYGDKKVHTDCI

>XP_011447134_1 XP_011447134.1 caspase-7-like isoform X1 [Crassostrea gigas]

MEEAMSPECVSNTDVPDAKPVSETDAFPFGKSKSQRKERGKPVATTIQTATEFFSHKYKM

DYPNRGKAIIINNKKFNPNTGLNERNGTDQDASALCCRLSELDFDVDLFHNLKAEEIKVT

LQKAAELDHRDNDCFTCAILSHGEDGFIWGIDRMIPINDLMEPFKGNKCLSLAGKPKIFF

IQACRGTKFDDGVDMNVADAKGFMDVEPQFSLQKIPSEADFLIAYSVVPGYYSWRNSTNG

SWFVQALSEVLMKHGQTVDLLTMMTRVNQIVANKFQSNTSHSDMNEKKQIPCVTSMLTKE

VYFTSK

>XP_034311529_1 XP_034311529.1 caspase-7 [Crassostrea gigas]

MEEAMNQECVSNTDVPEAKPVSGTYASQFEKSKAQSDIPVATSKQTSTEFFSHKYKMDYP

NRGKAIIINNKTFNAGTGLNVREGMDQDASALYNCLSKLKFDVDLKHNRTADEIKATLQK

AAKLDHRDNDCFICAILSHGGDGYIWGTDRMIPINDLIEPFKGNECLSLAGKPKIFFIQA

CHDDMNVGDGKEVEPQFSRLKIPSEADFLIACSVVPGYYSWRNERNKAKCPSFVQALSDV

LEKHGQKMDLLTMMTHVNQIVGKKFQPDTSHPDMNEKKQIPLVTSMLTKEVYFTIK

>XP_011445227_1 XP_011445227.1 caspase-3 isoform X2 [Crassostrea gigas]

MSSPRRSPVNVDHGSPYLLRKASGDRRSVRPGAPSEPNEYNFNYHKRGLFIIINNKNFHP

STGKESREGTDVDAERLEERFQDLGFDVRRYNDVSSSKLLQLMNEASQLDHSDSDCFGCA

ILSYGIEGRVYATDGMLPLDVLIVPFKGDKCPMLVGKPKLFFLQSCRCPNLEQSVEDSVS

YKSFLSEDSERSFSGMRKIPVEADFLFFYSTVPGFYSWRNHQEGSWFIQALCIVLENYGS

KMELLHMLTQVNRMVTYEFESCSDEHFTDEVKQMPCIVSMLTRYVYFRPKKPDIRD

>XP_034334496_1 XP_034334496.1 caspase-3-like isoform X1 [Crassostrea gigas]

MGRKDLVSLIRLRVGEGPLEHSDGWTFVSERRKTLYSVGEEIGQDELKKLKALLTVELRV

GKRTMDTVKDVWDCLDIMEKRLLDSELFSFLKTVYTTDERLLEIIDDFAKRSASAEATGG

QSIEETRSQEVAEHVHQRTNLDTENETTSQLQSQLSAYEIGKQAGICIIINNEFFSDRNK

HPPRCGTDVDGDSLRELFTMFGYEIGLHENKTCQEIMHLLDEVRKKDHKDNGALIVCILS

HGYLNTVTGACSEDLGIDEIMIPFRADCCPSLSGKPKLFIFQICQGQRTQPIVFHPTNAV

VKAEGDYNMTQDKAEVTDSPTLPRGPFLAEADFLIAFATMRGYASYRNENGSYFIQSLVD

LLRKRSPSEDVVSILVEVNHDVSQKIGKTEEGRYVAQIPESIIRLTKKFFLHPMNSKA

>XP_034334509_1 XP_034334509.1 caspase-8-like [Crassostrea gigas]

MHLLDEVRKKDHKDNGALIVCILSHGYLNTVTGACIEDLGIDEIMILFRADCCPSLSGKP

KLFIFQICQGQKTQPIVFHPTNAVVKAESDYNMTQDKAEVTDSPTLPRGPFLAEADFLIA

FATMRGYASYRNGNGSYFIQSLVDLLRKRSPSEDVVSILVKVTHDVSQKIGKTEEGQYVA

QIPESIIRLTKKFFLHPMNSKA

>XP_034310204_1 XP_034310204.1 caspase-7 [Crassostrea gigas]

MGDIEVDCPNVPYNFGRSNIDAQLVSGFGETEQFQRNRFENSYMFVDNDNVSNDIDMVSE

DHSEILSGIKDSSSRTQKPEKQDAGHQSGTKAIYPSSTNRDKVSAPISMPKSVFFGTDYK

MDHRKIGRALVINNMRYADRRQRPERDDSIKDASAMTNLLETLSFEPVTRRSGGQKEILI

NDVSSSQLMDIIKDATDPALNDYSDCDCFVMVVLSYGIDGAVMCCEGETDKLIPVEQIIS

QFQADKCPSLAMKPKLFFMQLCPAFTYNIADCVKDTGSSSSRLQPEIQSIPTEADILCQI

STIPGIYGWYEDNSALSWYIEGLKEALQKHLYNPLRDGRRSGEIFDVIFHVNRLYLGKLE

EQGIRPSQSYSVPHTTSTLTKSLHFLPKNRTM

>XP_034337084_1 XP_034337084.1 caspase-3-like [Crassostrea gigas]

MFLTFSLSHILGTHLGHNLGKFREHACRSARINSEPLPAGLFDWERETSYPVQSPVNESR

SYSVELDSMSSKYSLQSIEQRYTMDSDPRGAVLIINNRRFANNHKERLGTEQDCRNLKLL

FEELGFHVYVKEDLRSFEILRDCKDFAKDPILAKVDCMVLAVLSHGTADNVICGADAKMI

NVMDEICPIFSPLRCPYLSGKPKMYIFNACRGDSFYLPNEVGGYRSQVLENQDILFCYST

FPGYASYRDPDRGSWFVQEFVEVFREQANEEHVIDMLTEVNKRVSQKNHTDNSLEASCVQ

IPCPATSLTKKWYLNPLGAISLGH

>XP_011449627_2 XP_011449627.2 caspase-3 [Crassostrea gigas]

MATKKIPQEARELHNELIADLHDNINSKVDEHDFEKLIHRLKLKLSPIDLKACHNLIDAT

DKLQKRGHLEPGNYGELIKLVDCIDKNWVNLIREKGETIIEIINKANEPQPSVSSEESER

AKKRRLESDQDGQRSNENTDGQTQILKSSTKKERKKSVYPKTNDQKGLALILNFTKGRPG

ANADFCGINDFFEKTLNFECHVETDPPKEEVECFLEETACLFNNKRTAAKYYCLLVFIMS

HGNEDGIRTNDGKTISLDDIVSYFKNDKLKYFAGKPKCFFIQACRGSSPQGVVDLPDDKE

EVMIDQVVGKISIPTDADLLLAYATTPGYFAYRRPNVGSWFFCNLLEIFKEHYEEEHVED

MLIDVKERLAYDEEWQTNEGFKQMPCTWSTLTKRFFLAPQQLAE

>XP_034336797_1 XP_034336797.1 caspase-3-like isoform X1 [Crassostrea gigas]

MTGRMFRMLCSQPGDNDVGSVEPRVENTAQSRQQSSTGAPPISPQTSNEPIVGRDTDSSY

SFTHSKRGYAVLIMNHEFANEDENLPGVKHDIDSMSRLFTSMDFEVRQLINLTGDQLVSD

LWDIQTQISSDCDCFVCVISTHGGEDPGFYANSPQEPGYTKTEHYVLNNEGALWTRHIMD

MFNDQHCNALRGKPRLFFVQACRGNDVDKGVYLNENTENFESNKDQSEDISLNQDVFQIP

CFNDFLVMFSSTSGKYAWSEDLEGGWLLSAIDQVFSSGSAGDDLLSLLTQVCGEVASREA

FIPNNERAHCSKSTACVYHKLSKDIYFPSSKTKYSDVDC

>XP_019930379_2 XP_019930379.2 caspase-1-like isoform X2 [Crassostrea gigas]

MDQEDQRREFCENSEILAPESEQDGYKAIKKTLSVLIGRKSKNRKLKEKPHVQSQRDEDS

GAFTSNYEYQRESHPMQETSEPAWVYPQSSFPPRDPAPHPTSADTYRFCHETRGFAVLII

NSEFDSQAKRENAVWDEDYMREMFKELNFDVCVLRNLASKELLKEMKAIRDAVSVESDCF

ACVISSHGMEGKVSTVHPEVKKPFIRIEHFTYTRDGVVRTNELLELFNDRNCRHLKGKPK

MFFIQACRSRLDVQRADEVDMGVEVHLVETKATTSAGSPARFFTRGDVVGNYDDYSGRRP

EEFRRNFTAATLDPLYKNYQVHGLSRQYPGYYHVHAQASLQRQEPARTGTYKFKPTEPEP

ESEPIEVFQIPCYNDYLVMFSSAAGTIAWSDSGKGGWLMYCLYHVLHDVTNTDDDLLTTL

TRVCGKMARDMETYCPSTPHYDKAKSAACVYHRLSKDIYLQPVSHL

>XP_034304251_1 XP_034304251.1 caspase-1 [Crassostrea gigas]

MDREDQGREFCEKFKNLAPESEPDGYKAIKRTLVDPLVHHVTQPPFPPLGRFCHETRGFA

VLIINSEFDSQSKRRSAVNDEHYMNKMFEELNFDVHVLRNLSSKELEDKMKAIRDYVTAE

SDCFACVISSHGMEGQVRTSTPGVREPFIRTEHSINTRDGVVSTNKLLELFSDRNCRPLK

GKPKMFFIQACRGRIDVERSDEIDMGVEVLLVETPASTTSRGSSARFFTRGDMALTSSSN

TGDMATTSPVNTEDVVPTNSITMGDVVAKYDDYSDRRRDDFNEQRRYQVNVHASLQRQEP

ACTSTYTFEPTQPEPEPIEVFQIPCYNDYLVMFSSAAGTTAWSDSDKGGWLMHCLYQVFN

DVTYTDDDLLTILTCVCGKMARDMETNCPNSPHYDKAKSAACVYHRLSKDIYLQPKEIKR

ERKR

>XP_034332982_1 XP_034332982.1 caspase-3 isoform X2 [Crassostrea gigas]

MEMAEFKVGNRSKSFPFKREMPMFMSSSRKLDTKSLEKELLSRFKSRRLHPHYHVQMGIN

RSPRQRLEKSFYPSSRQSVQSIDRVYHNDVTLPSITCAADDRIGADVIDTFDVHRHKKHD

RGHNDKHRHSLPNLKPQSQFNGYSSQGVNFEKQFTSPRLRNSDDASDSRFYTMNKKPRGI

AIIINNYEFYKEMIAITNSGPQTLRVIQKGLRRDGSIEDTDKLGCFFRNELHFIVKGYAN

VPARDMVKILSDVSDDDHAQFDAFVCCIASHGRKGTLFGSDCREIKVHEITGLFTGSRCV

SLSGKPKCFFIQGCCGDHFNIKDVLSGPRNIPREHSYIPLTCDFLIAYSMVRNCTEYPAR

QSESLLYKTLLRYIRKYSHRYDLSSIMVRVKKALAKLQESTIGVDSVFQCNFRAQLSF

>XP_034331081_1 XP_034331081.1 uncharacterized protein LOC105339898 isoform X4 [Crassostrea gigas]

MDADKDTYQFHPERTHQGLALVIANFTEGEDRRGGADSDLKYMKKTFKRLGFKVDCHENV

KGSELVSILNKCSQHMLTSYDCFVFAISSHGMEFVQKDKMGNSVQQHVIKMFDGEYIYTK

DILDYFTDRKCKALKNKPKLFFIQACRIPETKSTHDQYDAGVGFDSGISTHNKKGGQDQR

TTRDKTDFGGSSGGTETDSDEESDVSDTDVEEDESSLSDSDIEPTAKGQGVCYWQKLEKR

QTEILKKLKKMEKGQTEILKKLKRVENGQTKILKRQKRMENKVWHLGREYMVAANQNKES

NALAAANQNEEEPSPQTTTNQNEEPSPRTTANQNKEPSPQTTANQNKEPCLQAGGNGDSN

SHSAADENENASGVTADLEDDIDPAKRRRLYPLESGGSGDDIDPARGRKIYPPAPLEFTL

VPCHNDMLIMFASPQGHYAFRNKTDGSYMLQLLYKCVEKYYANGGLAQNQTNFLDVLRDV

TTQMSEMKFMGSKEYTIVPCIVHKLRKDVIFTQGKSMGSTKFTFKLSEIISTLGMSFFTW

SNKN

>XP_034313845_1 XP_034313845.1 uncharacterized protein LOC117684706 [Crassostrea gigas]

MSSNADEVKFLTGKDLERGKLNFETSFLQELEKCKKLTRNNVNDNREYNFRHAVRGIAIV

ISNEIFDDGSKMRPRTYAKIELSKMNEMFAGLGYLVMSFKDLTATQMYEVIRTATSSKMD

FFLNNSDSFACVLASHGSEVLEKSVANPKVDCKVYHHVIYGTDRTIKTHDLFDLLAESRC

KALQGKPKLFFVQACRSRSNQGPSRYSKGVNVVEPISKNCSNPRQGCDQTESLSKDSAPQ

STEVVDGSVDQVKRRRYTLDQSDNAHDSNPKSASDTATTDESEHNFEEDAYRGNEASQHT

ELTNELMDWSSSDDVIPCLDVDIIRDENKDLVEEPSFIEIGKNWVRKGFFKRRNLFARPV

QDVMTTVATPCHNDFLIMYATSEGKVAYGREENGGWLVNAMHEEMVKHDPKNCIDFLSVM

TSTMYNIAYHYETNTGEPSTSGFKTAPTLQHKLSHDLIFRNK

>XP_011439359_2 XP_011439359.2 uncharacterized protein LOC105336650 [Crassostrea gigas]

MSSNADKVKFFTGEQLERGELSFERSLHRELEKCKKLTRNNENDDREYNFRHAVRGIAIV

ISNEIFKSNDGSKMQPRKYAEIELRKMNEMFAGLGYLVMSFKDLTAQQMYEVIRTATSSK

MDFFLKNSDSFACVLASHGSEVLEESVANPKVDCKVYHHVIYGTDRAIRTHLLLDLLDEN

RCKALQGKPKLFFVQACRSRVDPGAERFDRGADIPTDHSLKEHSKPKRGRDKTESMSDDS

ASQPTEVADGSVDQVDKRCRYSLDQSNNAHDSNPMSAGDMATNDESEDNFEEGVYRENEA

SQHTELTTELMDWSSSDDVIPCLDVDIIRDKNKDLFEEPSFIERGKNWVRKGFFKKRDLF

ARPVQDVMTTVATPCHNDFLIMYATSEGKVAYGREENGGWLVNAMHEVMVKHIVHIRENP

MNCIDFLSVMTFTMYNIAIHYETNTGEPSTSGFKTAPTLQHKLSYDLIFRNKYS

>XP_053379488MercenariaCARD

MEKRHKELLILQRANLVEDLNMRSGLFTQLLCRKIIDQRMVRKIKNGNTEDEQAEELLDY

LPKSDQGAKTFDLFCEALIADSQSNVVKSYLKPENKKESDENKTVQEATESIGDTSSQAG

GQTCVHVAEGVEYRDGDRSVASPSHVPNIAIREEFQSQMYRRSIEPRDLHHSLSSPNLPN

SSEHEARFFYQNSLRNPNAVYTVLNDRCINKMEKEKQLFNMGLPVASISQIMNDYHMQNN

NDPLLVRDSGSKILPKCASSKDFSQNAFAQPPPVFGYSPHKEINVSSMNSSFGPSPQKSF

AFHTRDQTYSPVLQESNFLSAKKGYGIGTIKERELEIENIYSSSKQNEKLFNEKIGSCYE

GYDESMRHKVSAPQVSEIHSSPSRFSDARNVMEMDRYIEVNNSSDGSQHRSMISFPRHVT

MLPVSHHPDLRSEESFRGDDSRSPRLMRNDSVDTMQKRPLSEEVETGAKRFREDVDTCSI

PNLVKPEKLPTLRVLEAREQQTINIPVIEEPRKQHSMPAVVVETDNHLEKSQRSSLLGAM

DDPEIDLTDGPINVNVELSTRQFYMTHYKQAYAMRRIPRGKALIMNVNEVVGKSARRGTD

IDRDNLHNLLCQMHFDVTVYNDTDGLTAMEMVQKLQDFSQHTGHASGDCAVVCILSHGEE

GYIFGADGKKFELDAVFSLFDNTCCPHLRGKPKLFIIQACRGGALDRGVQYYPDEHDGSD

ARVRQVPSMSDMLICYPTQNGYYAWRNRDRGSWYVEALVQIFMKYARCEDVCTMLNRVNL

LVSKKVSHCPNIDMDQMSQMSEYKSTLRMPHLYFFPGIGGCS

>XP_045161198MercenariaCARD

MMEQVHRKLLQRNRTTLIDNISNPIDVVEKLFEKNLFTEAMKQEVEAERTKNGQIRKILD

TVPKRGPTAFNIFYVSLLETANDTAADILKPELQGTHPHHGGAGERGYAIPQPGVELPQD

WPNVAAMTMEIKVLPCNVRSDLVRHNWISPNVYSMKKRCRGRCLILSNQTFYGPLEKDSK

GVARCILPSRHGTVKDRDDLKVLFKQLHFDVITRNELTVQDFHSELLREAAHSDHNQAEC

FVCVICSHGTKTGIYGVDGGVIPLENVTKYFDGEHCKGLQSKPKLFFIQACQGETADRPG

SNVVDGGQDLSNLLEKVHIADQTDAAKETSEQLDPNSVPTKTDMLVACATHPGYVSLRNT

KYGSWFIQAVIYVFQKYACKEDIFGLLILVNSLVARGRTALDEKGPRDVMQASLFTASLT

KKFLFFPGVVADSET

>XP_053378967MercenariaCARD

MEPAQRAAIRKCRVRLVSDLMIEHMFDSIESRSLFTPIMLECIKAKESRPEKVRTFLDYL

QRRGPDAFDQFIQCLRETEHDYVASYVLEEYLSMKNNTAQNNLNNSRGGTVPMSNNIYLT

QETQGASQGEMETQGGQMLNSVMETQGPSQTSSAIMETQGSSLGQLSSSVTVSQGINEVD

RREEEMQEDNEGDALMSEQSSVQGNESSVPGSTAQGTCSAPQPSLPIGQFSSQVTSMGPA

GYTGNFKEEYKMNTDPRGLLLIINNRDFQGVLTTRDGTDVDCDKLKTLFMSLGFGVDVRH

NLRAHEIKETLNILSRHQELSQVDCLAVALLTHGSEDYLYGVDANYFNVNEAFLPFTAEK

CPALHHKPKFFIINACRGDAEDRGSLSHRDIDTASTSHDVSNVPPQPLGCNFEGHYETSS

ITYDVQKMPPEPRSVGSRIPNMKDFLVAFSTIPGHVSWRHLNDGSFFIQGFVKVFQEYAH

KTDVLSMLVKVNNEVSKNIKHQGVQIPAPQVMLTKTWYLNPVMRQSI

>XP_053385490MercenariaDEDDED

MSTTVLPSQASLQQEVACVNDDTEDVQCDSLRVDNNRTGLSDLHQCLIDIDEEVGEEELE

GFKFLCRDHISVSKLDRAQTCLEIFDIMQKMVLIDVNNVDFLLECLGRVHRFDLVRKLGF

QPDAVRDHLPNFKKLYPFRTLLMDITEDLDDQETEKIRFFLRGKIRKKLLNEKRSLLQLF

VFMEQEGLLAPSNLQLLTKTMESVRRGDLVDKIREYQGLPPLKRHEPGLPRNLNPTVPSQ

HQVNVGPNEHVNRAFSQEPHQQREGLVQSGSAGNLPGQDQSSQIERSISAGTLSVPSGEL

EVSSRATLPEHLILQIAEDLKEQVNDMGIELLFSNAQLAQYEREIPDTKQRTVRILQDWQ

TNLDDKGRITPTLLYVLNKLGLKNASQAVKRWIDSINTDQAIPQQNMAGFEPSGPPGDSE

NVCVSMRHAQNVAEQIPPQQHLENVQVVENLQQLHISPNQLAENRCLYVNGEAVYRVGEQ

IVINPGVSRQLQPMVIPGQGEGQILPGQQGQGTAVGNPARVSGPVLQGQGRQSEGLTAPP

PLKQYRMNRDPRGWCVIINNEKFHVDRGDRESKEMPPRNGTERDAASLEHMFNKLGFIVQ

RHNNLSNTQMMNVLIDAAHNVDHTCFDCFVCCILTHGVLGHLYGSNGVLISIQDLTSTFQ

ANRCPTLAGKPKLFFLQACQGRDKMEGGAIQQDGPTPIPDKNNDISTDQGDSKEMIPNEA

DFLLGYATVPGYVSFRSRNHGSWYIRKLCELLEKYSNSYDLMSILVEVNREVANANASMD

GGLYKQIPAPLVTLRKQLFFGKLSQPPRV

>XP_053404642MercenariaDED

MALDPVTTRHKLLLTTISDELGSQGDEYDSFVEHFKGYVPRGLLQSKKTLLAKFDALASK

GHLGPGKYDVLKKICENSGNIDILELVQKAEADINEILNTGTERNSRQTTTDTGQPALRQ

APLQSPPVQETVNGTAGPSAKKARKEVDSSDPLYYMDREEHGYYDNSGKTGILLILNSYE

DRKGTEHDVEKLKGFFGGKGLRFEVFDPRENATQDLTTEEVDEVLRDVRSRLNGPLATHY

YCFICVIMSHGNEDGIKTKNGIISVENIRQFYNNKNMPKFAGRPKVFFIQACRGSAQQDS

QEIDDADDEKRPEAGQNDTEDESDEIIHVPTDADTLLAYATTLGYKSFRRRGIGSWFITE

CINVFEKYHKKDHLEDMLITVRENVALRWTKDDGAKQMPCVWSTLTKRLKLTENNTS

>XP_053406687_1 XP_053406687.1 caspase-3-like [Mercenaria mercenaria]

MASDSGLREDTTDAKGFLGSFVGKLMCGTVRSKAFMDIKDIEEEDIQEAYKKVPPAFKRK

ELLSDMYKMDFPRRGIAVVINNKNFSPEMEKKGYGVRTGTDVDATRICSRLKMLGFEVDR

YHDATCKQMKMAFSGAAAKDHSDADCFVGVILSHGEKDKVFGTDGPVDTNDIFQYFKGYS

CKSLAGKPKIFFIQACRGHQLDEGVNMNVADAKPFQYEAKAEEEEEIRIPNEADFIVSYS

TVPGYYSWRNSANGSWYVQALVHILDKYGTTMELQKLLTQLNKMVAYTDKFMSNTLQTNP

SMHGMKQIPSFTSMLTKDLYFVPKK

>XP_045188502_2 XP_045188502.2 caspase-7-like [Mercenaria mercenaria]

MGVCGSTGRTGTGNGPARTYDTVDGPVGETDNHEGRKNTGTANGNSNIRNHTNVTHIENT

ELSSGKDEKDVGLVLAGQLNPRSFANIDIIDLDDNEEYLQAIQQAYRRRRPKLTRKDLLA

DVYKMDFPRRGKAVVINNEKFKDSPQRKGTNVDMTRICNRLERLGFEVERHNDVTCNQIM

SAFVNSANEDHNDADCFAGVFLSHGGSNEIHGTDGILKLNTIFELFQASRCTSLAGKPKL

FFVQSCRGEEFDDGINVNVADAQGMNALPEEEVEVTIPDDADFLISYSTVNGYYSWRNAR

QGSWYIQALVRVLDECGTLLEIRKLLTRVNKLVAFEFESKDRRQSRKKQIPSFSSKLTKD

LYFTPK

>XP_045214182_1 XP_045214182.1 caspase-3-like isoform X1 [Mercenaria mercenaria]

MSEQGEKSNDVVDAQPTDTTDAKPMSGSSSAGIGVPKFQAVGGDGEPVISAEDFFADEYK

MDYPHRGKAIIINNRTFDRQLGLGERTGTDQDAAALNMRFMEMGFDVDIYHNLVVRDMAI

VLGKAAGDDSYNRSSDCFACAILSHGEEGIVFGTDAKIEVKVLLEPFKGNNCKGLLGKPK

LFFIQACQGMQFDSGVGVNTSDAKGQMEMEREITVYKIPSEADFLIAYSVVPGYYSWRNS

ANGSWFVQALSDVLAKHWHDMDLLTIMTRVNKKVAYEFESKTGKEFMNQKKQIPCITSML

TKEVLFKPKK

>XP_053406744_1 XP_053406744.1 caspase-3-like isoform X1 [Mercenaria mercenaria]

MFNATMDQVTDNMSTLGVKEDVTDASVDTGGTCSVSQGKENVNVDAGVAVIGEHEVAREE

QMTQHDAEPLDIEGSIDVLEARGDQGAASTDARVLGCLGATDSGQKVSSVPTVSQTLDAY

DMNHKYRGHALLINNEKFHIKLRQQGLGNRDGSSVDERSMKKKLNDLGFSVTGYPDLTAN

QMKSVCSKMAAKDHSECDCFVCVILSHGEEGVVYGTDKIVDIKELTQKFKGDVCPSLVGK

PKIFIIQACRGYQTDPGVNVNVADAKGKAEAEPAMVRLPVEADFLFAYSTVPGYYSWRNG

MNGSWFIQALAKALEDFGKTMELRKLLTLVNKKVAQNFQSFNPENKDFHKMKQIPCITSM

LTKDVYFSTTKRKKVM

>XP_045212901_1 XP_045212901.1 caspase-3-like isoform X1 [Mercenaria mercenaria]

MDREYPSSTELSVEWDGSQASLRNEGVSLHGSSQSLQMSSASPRLSPISNDSSSPYLLRK

VSADKDKHHVRPGAPSEPNEYNFNYRRRGVFLIFNNKHFLESTGQLEREGTDVDAERLEE

RFQDTGFEVRRYDDVTYARMTKLMQEIASSDHSDCDCFGCAVLSHGREGYIYATDKLVPL

EMLTLPFKGDKCPTLVGKPKLFFIQACTDRRTSIFGEKRASPSRFGINGSPSPHIASRRV

PVEADFLFSYSTVPGYYSWRNNQDGTWYIQALCIVIENYGSKMELMQMLTQVNRMVAYEF

ESCSDEEFTEDIKQMPSIVSMLTRYVHFRPKKPDIRG

>XP_045213875_2 XP_045213875.2 caspase-3-like [Mercenaria mercenaria]

MEPGTEEIHADDETEFPGSKTVSRIINTVTRSFQHFWSSELEPPVWDEADEYDFTHPCRG

TMIIFNNEHFKGHSSRPGSSVDAENLYQTFTKLGFNVSCHHNQTRDQMINVLKKASKSDH

SNSDCFVCAILSHGDEVHVVDDNNPGRHVREDLVYATDQIVLTRELVNLFRDKKCKSLAG

KPKLFFMQACRGSTLDSGVDLNVLVDEVDACLADTVEVTPCPIYKDFLMMYATPPGYYAF

RRPDTGSWFVRALCEVLSQPNAGNRTLNQLLTSVIKLFSQQYQSKSANLKISGKKQTPCF

MSMLVKDLYFKPKEGSTPV

>XP_053393150_1 XP_053393150.1 caspase-3-like [Mercenaria mercenaria]

MEPDTDNNTVKSDAEVTDAANLGALKEAIIRPLKKLLHLSNDPVSWDESDEYKFTHPCRG

IVVIFNMERVKGENVDSELREGSSIDVDSLKETFKNLGFNICYKENQTKKQVMETLAKES

KRDHSNEDCFACVILTHGGSYYRVHKTESSHKREDMIMAADTPLLTSDLLKLFHDNECKT

LKGKPKLFFVQACRGDKFDKGVYLQVEDEVDAKPADDSMEFSPCPIYKDFLIMYATPPGF

YAFRHPEKGSWFIQALCKTLSSPGVRNRSLHQLLTSVIHCVCVEYEAQALQFLGYKQTPC

FVSMLVKDVYFRPKIKSIPI

>XP_053389765_1 XP_053389765.1 caspase-3-like isoform X1 [Mercenaria mercenaria]

MYKHSEIQMNAQSSNQTPFSVIERRGAQDIRTENYTYKPSKRNSQFSFPEYRMDRIPRGY

CVIINNRHFYKENGDHSWPMPDRNGTEKDAEYLRYTFSRLHFIVRQQDNLSDTEMMKFLC

NIGQDDDNERYDCFVCVILTHGVLNHIYGSNGKLVAIKDVTAIFQPSRCKGLLGKPKIFF

IQACQGREKMTGAVECRDDEAELETDTVVSNRTDATRCVQSLFADEFVKSDYAESSLTRT

SITSDITKNHITSTDIQCDGDRNSDISIDSGHRENIPDEADFLIGYATVSGYVSFRSRNH

GSWYIRKLCEILDSHADRCDLLSILTEVNRQVATANAPMDGSLYKQVPAPYSTLRKLVFF

RFQEC

>XP_053386794_1 XP_053386794.1 caspase-3-like [Mercenaria mercenaria]

MSMSLNSGGRENHEDVVETSGLEITDKEADLNIPVNEMKGLSLTVNDDIEEQTDEGDGLL

KKLGTLACVVGKKKVKGKEEIEPFSTNQESWIYSNLASDIAEAEAENVSSNITGTHIKHS

PRQKQNIDSTESDCYKFTYPKRGYLLLVINEKFYNQASRKGADLDLLKMKKVAKKIGFQT

LNKGAETNLTKAETMNWLLKAQDTDHSTSDCFMFMISTHGLEQQNPRKGGLVDHALVCAD

DQLIFTSTIIEMFSETNCPSLKGKPKIFIIQACRGTSVDHGSDMTISNHKGKPDSTVRDR

KAKHDNPMHLQSDVSDTKGGQDSWVMNTPRDIDTPCLKCDADMLVMYAIPPGMLAWRNTA

DGSWMIHYLYEILMAYDLKAPKSFLNILTKVSARMALRSTDTPSIPEIHEKKAIPVIEHM

LTRDILFSLKL

>XP_053380576_1 XP_053380576.1 caspase-like [Mercenaria mercenaria]

MPFKRKDEVKCESERKRAMTEKHTDEVDTGFGASGTPPDQKEIETSNPLLKHLVFSFLDA

ENTRDLNKSETLSLLEKAKNTDYSTCDCFLFAISTHGLEQKNATKGGLVEHALVCADDEL

IFTSTIIETFNDKNCPGLKGKPKIFIIQACRGKNVDPGFDISVSMDNGGSSITVKDKKDN

FTPVPSDMSDTKAEQDTWITTASSLKCYVDMLVMYAIPPGMFAWRNTDEGSWMIHYLHEV

LLDEYKGSKSPISFLAVLTGVSARMSRRTTNTSKPEWNEKKAVPVIEHMLTRKILFSKKI

>XP_045211958_2 XP_045211958.2 caspase-7-like isoform X2 [Mercenaria mercenaria]

MADPLLAEQDEEILIEDQQDSILKTLRKYLGPDRKTFGKEDIVTNHQNTQSYSNRGSSAR

RETDGEIESIAPADHYKGFDITLSEQNEMVFTNTSNMFTYSSETNILEKGPATKAVNIAE

QQTTAQEHMDISDIDNQYQFNHKRRGHMVLFVNNQILNDDDHRKGADEDFRKMEEIRTKF

GFELLNSDMTTNLSKLAMQMVLRKAQQIDHSDCDCFAIMVSTHGQKLSSPRNRTGDIYGL

SCADGQIMELSEIVKMFDDAHCPTLEGKPKLFFIQACRGNDSGGKSDTADTKARYESQSR

TSVYESPSRTPRYESPSRTPRYESLSRTPRYESPSRIPRYESPSRTTRYESHLKTGRCES

PSRKVKYESTSKATRFESPSRTARFKSPSSSIVATLSDDDTPYLNCCDNDMLVMFAAPPG

MVAWRSEANGSWMIHHLHAVLMEYDFRKPKNLVIVLTKVCRLISQCETHVQDPKFHKKKV

VSVIESKLTKDILF

>XP_053382662_1 XP_053382662.1 caspase-3-like isoform X1 [Mercenaria mercenaria]

MADISESELAKIQSQGEEITLELTDHEDTQESNRRWKTNLEKHPWLRDPPECLKVKERTE

NMGKRIKQLEEDDEIYMMKEGGRKMCLIIGNTKCKSYEELMGCAKDNAYLSALFKQLNFE

VTVKNNLTAKEMTAFLNEKKENLKDVECFVCAITSHGNHTGILGTDEEVTSFLDIMKMFD

RDNCRELSNKPKLFFFDVCRGRHEDVPKNPTVVNESTDAMIPPIKSDILVAHATSHDYVS

FCHPEYGSWFCRSFVKVVSENACRYNILDMLLKVNRKVGEFQSAKGSLQSAQFDSSMRKQ

FFFSQPVIPERLQIKMLSNLREKRGTKRKPDSKIFEVSIIFILRKLKTLITHKLYGIMYA

>XP_053378767_1 XP_053378767.1 caspase-6-like [Mercenaria mercenaria]

MTQLLQEQKEEENLKDVECFVCAITSHGNSKGILGVDEEATTVLDIINMFDRDNCHELSD

KPKLFLFDMCRGGKQEDVPKHATDVTDSTDAMFPPMKSDIFVAYSTSHDYMSFCHPNFGS

WFCRSFVKVVSENAYRYSISDMMLMVNRMVGTFNSKNGSRQSAQFNCSMRKHFFFSPQTS

GEVKKNAVDEQKMSQPIHEVTSQQMS

>XP_053379902_1 XP_053379902.1 cell death protein 3-like [Mercenaria mercenaria]

MERTEDMEIRIEQLEEAKLIYKMTEGGRKMCLIIANSKFSRQARLEACKLDCRYLTALFT

GLNFDVEGKENLTAEGIKAFLEDKKRHLDVDLECFACVITSHGDPRGILGIDDAVIEFQN

ILDMFDRDHCPELANKPKLFLFDVCRRSGVDLVQLKAKKIVNYTMIPPTKSDILVAHATT

NDYVSFCDRNYGSRFCRSFVKVVSENACRYSILDMLLMVNEMVGEFTSGGVLQSVQFDCS

MRKQFFFPLPVIRQAKAIAAHETKTSLEEDRDVKQTGL

>XP_045201993_2 XP_045201993.2 caspase-3-like [Mercenaria mercenaria]

MAVSLHIKLLTDISDMMETDSPEYINFVYNMTQFVPKGLLNSKPNLLGKFAAMKSKGKLD

IGKYDNLKKVAKDSGNNDVDELIQEREQEIIEALASENLGKASGQSTLTTNCGQSVCQTG

SQKDRGQSSSVTPSASSRSGQIKSGADKVDINDKNVKKRKHEEETDQSDPLYGVDVQGRG

KYDTDGKKGFLLILNYERSGLGNDVRNLESFFGNDLGFDVKTVSGNLTASDIWTGLNETR

EKLNGDTKRTYYAFICVILSHGNKVGIAFKDGETISVDDIRKHFINSDQTMNNFTGRPKL

FFIQSCRGDDVQGLHQVDEADDHEKEDIIDESDEIFNVPTDADTLIAYANTQGYATLGDS

LDSSTQFTFFINALLMVFKKFYRRDHVEEMLISVRESLALGEKKYFHRQCKYSKQMPCTW

STLTKRLYLTAFQTAMEQN

>XP_052225747_1 XP_052225747.1 caspase-2-like [Dreissena polymorpha]CARD

MQQEHRNLLVKNRTLIIENMANPLDVTDMLYEKNIFTDAMKQDVEAQRTKIEQVRKILDL

VPKRGPKAFEVFCHTMVETHNYSIAAALGVQGQGHVIAQPGADLPEEWPNEKAMSAPVIV

EPCNVHSDLVRQNWHNPNVYPMRKAIRGKCLIISNKIFTKSEIDSEGVFRNVLRHRYGTE

RDVEKLTELFSQFHFEVIVKEDLQRDAMSRVLFQFSADKSLMDSDCFVCIICSHGTREGI

YGTDGEIIPIEEVTSHFDGEHCPQLQEKPKLFFIQACQGEKYDRTEEDQGPDISKVVKDL

KNIRLEDGANRTDAEGDRSPHTATATDAVGEMQPINTRTRVSIKQDILVACATHPDFVSF

RNELYGSWFIQAVVYVFQKYAHKESLVEMLTMVNKVVTTGRTGTATQCSKYETSLTKKFL

FFPGIIENAGS

>XP_052285027_1 XP_052285027.1 caspase-3-like [Dreissena polymorpha]CARD

MDPAQKTALRKSRVHLVNHLEMRHLYDYILKDELLTPIMVETIQAKECRPDQVRAFLDIL

PRRGPTAYNIFLKILTESGQNALANHIKQEYDRETGNASGLSELSLTPESSSDNEDMEVE

GVEVFKGTSLITAVQETGSPVANLPDPQSIVSLTSQGSSASLGPGYTGNYEEEYRMESSP

RGFLLIINNRTFRGDLEERVGTDVDCANLKCLFLSLGFGVELRTNLTAEEIIANLEVLSR

HEQLPSVDCLVVAILTHGSDDYLYGVDEMFVDVNHLYEKLSATQCPALIHKPKFFIVNAC

RGDAEDKGILATPSIGRCSVGVDVNKIPDTAQAVRRIANMQDFLIAYSTIPRHVSWRHKT

EGSFFVQGFVKVFKEWASTMDVVSMLVKVNHHVSRIDEYAGIQIPSPQVMLTKKWYLNPP

NAAYSS

>XP_052285285_1 [Dreissena polymorpha]CARD

MEQKHRDIIRQNRTRLVNDITDLASILDDLMTDKIITDSMSQDILAKKTPNEKTRKLLDI

LAQRGPKAFRRFCEALMNTQQDHIVEILDPEFYRNSVRTAKPVSSDATSAGGDTSLARPK

DEHPKSSIPQPVSVPIVPGKQKDERDKHIPNEFTSKGPDTKNEKPKKDDLPTASPSKTKP

VQPIPSEVTSETVARKNKNSNRPMSSSSEPSSLPNPNIFPGQAEGELPETAKDWPKELNL

RSNTEFKVKKLTGNIEDLDKYSLEGKRTCIVVANQHYTERYKTRLVDIDRLRIKNVLKEL

NFNDITIHTNKSSQQLLSCLEEERQKISKETSVLMLIVTSYGANGQIFCVDGDKVSLQQI

VDIFSVENCPLLKGKPKVVLIHAGGFLLEDGSSPSDTHCNEIRRSLSHLSLGTPTSKSKG

FFSETVSDCRPQTLLDDSQTVPTAPTPESKIVLSPSGILDETLNVYDNSDFLVAIATGPD

EDHPHYCSLFLLATLYVLCRYSHQHHFLDMLNKVNRLKKDASGQLVEICKCKSSLTKNLY

>XP_052235568_1 XP_052235568.1 caspase-8-like [Dreissena polymorpha]DEDDED

MIETMAVELTTKDKLEKFKFSHEIQTDTLGNNDDGSVPALTSGEFHQRLININDEMGSDD

IEALKFLCRDYLPASKLDKVESALEIFEYMKKAGMIDLNNVDFLLESFARIHRYDLINKL

GFSSDDVRKVIPTFKQVPDFRVMLMDLADDVDKTEIEKVRFLLKSSIKKKMLNEQQSILQ

LFVLMEQEGLLGPTKLSLLQTVFTSLKRPELENRIREYEGAPPSQVQPPGMPPSPSGQSE

SRENIQPLRDFQREMRQGDPLRAPINMQQSNNQYTPQRPGLANMAPPEHPNHSFDRSQSA

GTITVSSSAGDLAMGQENPRDIPEALFLQIVEDVKDRFEELLTELGCAREFFHYTQTYPS

PQKRMQEFLQNWKEDKRVGEVVSTLGYALQRLGCKQDAMALRDWAESVQLQWQASTASAV

PASPDLTDLLRDQKYMYNPVGPPELNEFVSLREAKPEIGTSESQTRHGGVAPPVLPPVTG

NIPVINASQMPPRVQEQIRNSGVLQSRREPNQTAEGIREMGHIPGPQQQVAMATNPDQGQ

RSAQVATRGDGTTRLLQQFAATNLNVKNPSGELDAYKMDSNPRGWCVIINNQDFYTDPSD

RDAQAMPPRMGTLKDAENLTYTFKKLGFVVQRYDNLTADEMMRTLVDASHNVDHSKFDCF

VCCILTHGVLNHLYGANSRLISLKELTSCFQTNRCPTLAGKPKLFFLQACQGRDKMEAGE

VPERDAPEPFNDVEVDTLPREMLPNEADFLLGYATVPGYVSFRSRNQGSWYIRKLCEYLG

KYANKHHLLDILTEVNREVAKADANLDGGRYKQVPAPTYTLRKRLYLRPIQ

>XP_052269676_1 XP_052269676.1 caspase-7-like [Dreissena polymorpha]

MDGLEGIDDPDAKHFFSKYLGGIRRAVSGAPRWQDVKLEEYVNVENDANYKRKPMFTKAD

KEKDEYKMNFPRRGKAVIVNNKNFTDNMQKLGYGVRSGTDIDHTRITNRLRRLGFDLDHH

HNATCQVMKDVFAKVAKEDHSDVDCFVAVLLSHGEEGKICGTDGIIELEEIFQYFKGAHC

PTLAGKPKVFFIQACRGKEFDSGTNVNVADAKRVVESDLEKSEVMKIPSEADFLLSFSTV

PGYYSWRNSTNGSWYIQALCCILDQYGTVLEIQKLLTKVAKLVAYAERYMSNTEDPAQNR

KKQMPSFTSMLTKDLYFHPKNGST

>XP_052269677_1 XP_052269677.1 caspase-7-like isoform X1 [Dreissena polymorpha]

MDASEGTGDSVVRDNSYTLRLLKHKDTEKDEYKMNFPRRGKAVIINNKNFTDDMQKRGYG

VRSGTDIDHTLITAQLQRLGFDLVHHHNATCRVMKDEFAKVAKEDHFDADCFVTVLLSHG

EEGKICGTDDIIDLEEIFQYFKGAHCPTLAGKPKLFFIQACRGKEFESGTNVDVVDAKHV

VESDLEISEVIRIPSEADFLLSFSTVPGYYSWRNSTNGSWYIQSLCCILDQYGTVLEIQK

LLTKVAKLVAYTEQRYMSNTADPAQNRKKQMPSFTSMLTKDLIFHPKDECT

>XP_052263052_1 XP_052263052.1 caspase-3-like [Dreissena polymorpha]

MSSRDQTDASPMEGDKTDAKGGDQTGALPPLPSASGGFGDKLGDKFQNIMTFEEFTADEY

KMDHLNRGLAIVINNEVFSKELGLGARTGTYQDAVALNERFEDMGFDVQMHENVPVKEMV

QILTKAAKDHNYNMNSDCFVCAILTHGEEGYVFGVDAKIETKQLLEPFKGNNCGSLVGKP

KIFIIQACQGSQFDEGVAVDVADAKAKPAREVQVYKIPSEADFLIAYSVVPGYFSWRNST

NGSWFIQALSRVLGLHWQDMDLLTIMTRVNKIVAYEFESRTTKEYMNQKKQIPCITSMLT

KEVRFKPKKK

>KAH3748363_1 KAH3748363.1 hypothetical protein DPMN_182807, partial [Dreissena polymorpha]

QGQRSAQVATRGDGTTRLLQQFAATNLNVKNPSGELDAYKMDSNPRGWCVIINNQDFYTD

PSDRDAQAMPPRMGTLKDAENLTYTFKKLGFVVQRYDNLTADEMMRTLVDASHNVDHSKF

DCFVCCILTHGVLNHLYGANSRLISLKELTSCFQTNRCPTLAGKPKLFFLQACQGRDKME

AGEVPERDAPEPFNDVEVDTLPREMLPNEADFLLGYATVPGYVSFRSRNQGSWYIRKLCE

YLGKYANKHHLLDILTEVNREVAKADANLDGGRYKQVPAPTFNSVKIAQGLINIKHGIYF

LVAAGLWKLDVLRGESSTINMFTAACQQFVKSIGKTTLHATPSLDEKDNVKMFQVVLKSK

RRRFIFWEKSVFRPSEFQLTDMMVDRKPLNVKMTTTDLVQEFNRTSKLSLKGKIGAEICK

EIIDVELDANDTITLDIKFGSVKKQELGEQDMIDELSDRHINMEHPYVRQIRKDKRNSLC

LISGVALLPDGGTIHRVTDEDLEAIAKAAPLKANGSAGVEYVKDRDIRLAKDTAVAYNVQ

ELDIDTNTGAIQLVLTGGKTGGGFRACVGYDEVDAGTGDVSVDFQLDHAREVFAYFLDLD

PKTRSEVNKNLLKLMEYPRDVDVIADLLNNIEAGQEVNNSVSDLAKRVVSPREVWQYVLE

QAGFSVDDDSLVIPKSKERVSMFGEFFDAATVVEESLLKLLTSLSASNAENMLLVLKSGI

QRAPVTVKNIVAEVLSDKTSHKMLEHLKYHLDELGVIAPPAEMPFCIDALYWIVFALYGK

>XP_052220239_1 XP_052220239.1 uncharacterized protein LOC127837318 [Dreissena polymorpha]

MAANTHLIGDKETNNWFKAYIGLNVTKEGLTNFLCAELQKVHTVVGKSCGKCPIENLIQC

PTNPYCNNRKRNYCPFHKSQKPQPCPICDNVKDNIILHHRYGKPSWRNTRAEQWALNDWE

IGKCYLPPDGYSSISSVQELDYNGVISIVLNCLHFQTCLSPLCLSPPPPDKQCPLEKVRQ

IGRDVRHTSDCKVTDADLQDYFKTLSTLLADPQYLQHDPSAIIACTRLSDLQNDRMSFEE

LGKLLKEVNQTFEEAKKTGERFSEMAERTLSKHLQKLDATNQFGLIRLKKMTILEQLCLY

SKKDREHRLEKCIERMQQAANKTAQADYERDVAAEEDEGNYTNEPSNIVASDPLFKLDLN

KAVRNAEEDTVQSDVQRSNNQTQAGISLVASDASIKDIVKKARNVIKFRGDSAPDDTMTT

ATEQMKTIRIDEQPTLPRYEMTHIPRGVCIIINNQEFNRNDLKRRDGSDKDREELEETFR

WLGFDVKCHNNLSRGEMILELITVAHNTDHTNYDCFVCCILTHGEHDTLYGSDGQPIHID

ELISNFNVERCPSLAGKPKLFFLQACQGDKNMVGGVQPQVDSSKASEVKTLPIFGDFLIG

HCTYPGYLSYRYLTGSIYIKTLCEQLRKLAHSCDILTILTMVNLEVSKWEGTYENNRVKQ

MPAPRYMLRKLLFLTPSESKT

>XP_052224167_1 XP_052224167.1 caspase-3-like [Dreissena polymorpha]

MYNNVCLQSESAVERRIAVRQLTIKSNKSASWFVDIRKLFWLYELGDPEDLLESPTEKEH

WKRTVNRTIDNIWLQQTLAESKTMKILQNLNMNMVKPRKPHPLLQQQAVSTYDANRQLLK

LKFMCGKYILQSDRASFSKNKVDDTCRVCYKSPETLQHMVLECSGLSAVRDPILRDIESE

IEKSFPCLWKSIVKTLKKKKKKEMSFDDTGLCIIINIQEFPGTKMQPRVGSEKDRDNLKK

TFTWLDFDVKCYDNLSGRAMIRMLEYVAKKIDHTNYDCFVCCILTHGELNHLCGSDGEPI

PLKKLTNNFNIDSCPGLAGKPKLFLLQACQDEPLVEGEKLQVDSSKASEVETLPNHSDFL

IGYCTAPGHPAYRDSTGSIYIKTLCEQLIKRAHSSDILRILTMVNLEVSKWVDTIENKRV

KQMPAPQSMLRKLLFLKPSESKT

>XP_052215323_1 XP_052215323.1 caspase-7-like [Dreissena polymorpha]

MAEIPDTVRERHVKLKTDIVAEFVTGSDEYNAFVEAFKAQGIPKGKLGNAKSLFDKFTLL

ESSGKLAPGNYDALINIASVSNCLDIVTLIRTADADIKSMLPDCPKDISGPSTSNGTRTK

KRAAEDPPCRLEEPKKNRVSSLETDEEDPIRTRDLKGRSCYDYKDRRGYLLIINYTNGRN

AADHDVKALKSFFRDEVHFEVDVNDKDLSLAELNEYLEAAKNKLNGVLGKNIYCFICAIM

GHGNQNGIATTDDKFKSAEDIRMEFRNDTIPNYTGRPKIFLLQACRGSVRQDGVEIDEDD

DKAENSEAEIDEDDIISKVPLDADTLLAFSTTPGYKSFRKTGGTSFRGSWFIQSCVQVFQ

EYYKSDHLEDMLITVRERVAELERPDDGKKQMPCVWSTLTRRLYLTKS

>XP_052286801_1 XP_052286801.1 caspase-14-like isoform X5 [Dreissena polymorpha]

MDEQTPKITETGTSKECDKQPETETETSPLTCAGESSFGPRDFEKTVFIHPAILSKGFGE

SIDAAYPCDGTGFVLLINYSHNRPGSEKDVNKLTGFFRDELGYELEVAQDNMTRHQLRNC

LENARTKIDTRFYCFICIVMAHGDETGIMTSDDKVQPYNEILETFDNKSMQNFAGRPKLF

FINACRGNRLQTSVPLVNLDHTEADAKPFTPPKLVAETADIFAVYSTVSGFASIRDTEEG

SVFIQSCVDVFKKYYDRKTVIEMTTAIQRKVQDRESCQVSEVKTTLTKQLYLKNVKWSSL

DQFAQLELKEI

>XP_052284842_1 XP_052284842.1 caspase-14-like [Dreissena polymorpha]

MGCVNSSNSSDQLGIIHNNDFDDNLQTNPSILSSLISCFEAQNHDEDATPTEHAIPIEPV

SQPETQLQMLAIAPLERGSHQQRRYAQFIHPAVLERGLVESIDVAYPTSGKGFVLIINYS

HSRQGSEKDVTKLVGFFQHELQHSVLLGNNNMTREELRAFFEDVKKSIDARFYCFICIVM

AHGNEKGIITSDGKVQSYNEIIETFDNRNMSAFAGRPKLFLINACRGDRRPWTMPFDFES

ITMDATPVHPIELVAETSDIYAVYSTVRGFVSTRDTQEGSIFIQTCVDVFQKYYKTRNVC

DMTTSIRRKVQHNHMCQVSEAKTTLTKQFYLMKSQ

>XP_052217267_1 XP_052217267.1 caspase-7-like isoform X2 [Dreissena polymorpha]

MAEYMSGSDGQTDVKQSSKIARAGEVTGTYGYLLILNQRKDREGSEKDVKNLLSFFEKIL

FFETDVIDQNTISETEMDDRINKARNKLNADPNISCFVCVIMAHGALGTFYVKGGPKTVE

SVRQAFCNESLPKFTGRPKVFLLQCCRGDKENKYTTVKDSDLKERMISVPNDADTLIAFA

TTPEHVAYREPYDPEEQKEQKQERQRQGSHFLNICIDVFKDQYEKKHLEEMLVRVREKVN

ATPFTVLENDEHILYSQMACVWSSLTKVLYLKKKLPDFFYSCR

>KAH3785603_1 KAH3785603.1 hypothetical protein DPMN_163696 [Dreissena polymorpha]

MTALLNERKRYATNGQGFLLIINYTYDREGCEYDKKKLINFFGRLNYRIAFFQNLQKRQL

LDGLRALPDILDRGEYYCFVCVIMGHGNKKGIMTADTNELKIIGIKKILDIFDNRRLKTF

AGKYMHLNELQYKLDDAGKKENSNANASTCTKDATNTSESNEMEILPPSGSDMVTFYSTV

KGFVSKRYPDGGTVFIQNVLRTFEQHHCLLDVETMMKMASRRIQRTHRQVPLFCTTLSRK

FHFIYCPRW

>XP_052271649_1 XP_052271649.1 caspase-1-like [Dreissena polymorpha]

MSCNVHNRGTAVIIVNKNFNRLNKRKWASKDYNNLMTIFYELGFRVQLVWNASAERMRRI

VREAASLCDNKNSECFVLAISSHGTETCKPGAGPNGANLYEHAVNGSDDQLVFLFELTQI

LEKETSLRGKPKICIIQACRNTHHDSKSQKDMLDTGIAIGTLGDDESDSKKKSSVSESEY

SDIKQFFKYDSDDESHSSSVAETDGPEHLAATGISDSVPIDCPEDFLIVHPVQPHKMSFR

GSKSGSFFVRSLFDHRKILEKESLLTYLTEVAGSIANKNIKFPSIDKKDLYKMCITLTHR

LKSDVTFHNNTFTRVIPTETYMVHPNNVVMSE

>XP_052268241_1 XP_052268241.1 uncharacterized protein LOC127869624 [Dreissena polymorpha]

MCDQNTSVDSTCYDGESGDLQKSMETASEVANKQHTALTKQCGALNKVTHSMFESQHKDL

EEAHIETESESVDDPDGISKFFKSLLCGTVKPKNVKENEESYQPKVKQTEENSLQTELLY

QENTYVHKASIPKLIPNITDAGNQYAFSYPVRGYMVIIVNETFKHQTTRKGSDLDWQYLH

QLARKLGFIVYNKHDTHNVSRSQTLKILLEMSRADHSGCDMFMFAISTHGMMQPNPRAKG

QTDHALICADDKMIFTSSITQIFNDENCPSLKNKPKIFLIQACRGEGSDSGSDVTILCPS

VKVHKTQRQFLGNAQCAGDGSQQIKDDSDAMGKSDSETTSSRNVFMGTSYGARPNSFVNL

RAIRQPHKTEPKETPYPVVDTPSLRCDNDQLVFYAIPQGQMAWRSEGDGSWMLYYLHTIL

NGRGCGQPLNLLKTLTKVSAQMSMRTSNSPSNLKLHEKKAVSIIEHKLTRDIILTPKQYH

TQHEHIGKYRQWAILRIFYLWINASSYNRVMEVKDDERKECCRVEQATGKTVCHTRDCEQ

ADLFNELVTLENSGEHHLLVIENESRQTGINNVETENSDKNCGTEATSDITMQTTRLSTD

SSLKNISSSKNIQIKTADFLSVINTIDQENKIHFPATENRKAIPDEKHIAENRNEICMTI

HTTTLNISSDFEKDDSDEYRKDGCDVVDGLGKLVKQKVKTLIVCASSKGMTVDDDSLPFQ

KANKEKVDENGDNLWVNSPLYGKETEINTCTTVVRESEEKVSRVSVGQEECGPFTVASTV

FTSELTETTKETCHFFLPSQLDETKTQSAQTVQISNTKCHGNGVLLVDPYYFNRGHRGYM

IIIVNQTFNHQAFRDGAHKDVEYLYNIAYKLRLTVYKGKNTNMSFTETKQLLEDVRCLDH

SDSDMLALGISTHGLEQPNPNAKGKSDHALVCSDDRMIFTSTITEMFSDDNCPSLKDKPK

LFIIQACRGEEHDSGADLSVAKKIALDRTCGGNAETCWQPCLDDSDVHFAKDSWATGPDN

QGQPVQLHDDLLATTGNESFALPMVPQFTSTQLVREQLGSSPHLKPFPFVDTPSLKCENN

QLVFYAIPPGMFAWRNTEDGSWMLYYLNNLIEMCDDRRPINMLKLLLKVNAVMARRTTNV

PSDKNLDKKKAISVVEHKLTRDLIFPPLVK

>XP_052277922_1 XP_052277922.1 caspase-3-like [Dreissena polymorpha]

MGWNSRTQKLKERVTTPLCAQTTGLIFTSTITEMFNDENCPTLKDKPKLFLIQACRGEEH

DSGADLSVAKEAALDRTCGGNTETCCQPCLDDSDACFAKDSWAGGHENQGQPVNLHDDLP

ATTGNEPTALPMVHQFTPTKLVREPLGLSPQLKPFPYVDTPSLKCENDQLVFYAIPPGML

AWRNTKEGSWMLYYLNNLIEKCDVRLPINMLKLLLKVNANMSLRSTDAPSDKNLNKKKAI

SVVEHKLTRDLIFPPFVK

>XP_052262360_1 XP_052262360.1 uncharacterized protein LOC127866063 [Dreissena polymorpha]

MTVDEDNRPFEKATKENAHEHGAVTSVKRPMYDKDIEIKTRVTAVLEKVEMVSQCNVSIK

ESCNQASTVFTSELTEDKTEKCRFSLPSKGEEKVNSFEQISQKSNAICTKNDILLVDPYD

FSRSHQGYMVIIVNQTFKHQAFRDGAFKDIEYLYNIAYKLGLNVYKGQNTNLSLSETLQL

LENVRTLDHSHCNMLAVAFSTHGLEQPNPKAKGKSDHALVCADDRLIFTSTVTEIFNDDN

CPSLKDKPKLFIIQACRGEEHDNGADLWVTKKTGVDRTCGGYADNCWQTSLETPDVHFTN

DSWAGGPDNQGQSAHLLDDHTITPGNESSALPMVTQIQSAQLLRAPLRSSPKLKPFPLVD

TPSLKCENDQLIFYAIPPGMFAWRNTEDGSWMLYYLNNLIEMCDERRPINMLKLLLKVNA

EMALRTTNVPSDKNLDRKKAISVVEHKLTRDLIFPSRVK

>XP_052269873_1 XP_052269873.1 uncharacterized protein LOC127871176 [Dreissena polymorpha]

MNCDRYFSVARYGTNEYKKFRDDVIACVEQFDGPCLPLQANRFIVKPEIISGPTEVIILM

FDEENEQTSFQKVVVRHADKELPATRVLTNVWAFQTGDLPSGDNRISVICDGQDIGSEIL

KVCEVDLKKAINAITVAMDIRPLTSLATLLQSAEEDVDEELAKLLSSKPSSIVETLLHMR

NNDGGGGKSLLTICKSIHLNESFVRINQENEVNSSDDYHELQSPLSRNSSVSRFYTASLY

RGTRDIGVGHSQGMESGVDQKLTKSSQTGRISNTKCPEQDVLLVDPYDFSRDHRGYMIII

VNETFKRQAFRHGADKDEAYLSNIAYKLRLTVYKGKTTNMSFTETKQLLEDVMCLDHSDS

NMLVVAFSTHGLERPNPKAEGKCDHALVCSDDRWIFTSTITEMFNDDNCPSLKNKPKLFI

IQACRGEEHDSGADLLVSKKAAPDLTCGGNAETQRFLDDSDVHFAMDSWARGPVHLHDDF

PAAAENESSAIHMVPQFTSTQLVDEPLGSSPQFKPFPFVDTPSLKCENDQLVFYAIPPGM

LAWRNTEKGSWMLYYLNKLVKKCDGRRPVNILKLLLKVNAKMALRTTNAPYDTNLDKKKA

ISVVEHKLTRDLIFPPLGDDQ

>RUS90572.1 hypothetical protein EGW08_001660 [Elysia chlorotica]

MSSDTVDAQGLPAESNDHDEQDARGEPRRDLHKSAPPLMSEADFVSDRYNMKHPERGMAL

IINNKTFHPRTGMGQRTGTDVDASKMNDLFTTLGFEKVRPLDDVTVADMREALVLAAKED

HTDRDCFVCVILSHGEEGYVYGTDDKIPIDELVMPFKGHKCPSLAGKPKLFFIQACRGQE

LDRGATVADGEGDEMEEEVIVRRIPSEADFLMAYSVVPGFFAWRNSSRGSWFIQAVYDVI

MKHWNTLDLLTMMTRVNKKVAYDFESNASAEYMRRKKQIPCITSMLTKDVFFK

>RUS70675.1 hypothetical protein EGW08_021565 [Elysia chlorotica]DEDDED

MTAFKMTDQEEYRILLQDLHSELPDEDIKGMKFLAQPLIKKRLYESIKDGLGLFGALEDC

TMLSRFNLVYLSQLLETVGRMDLQARITEEIECDAISGEESLICPFRKLLFNLHKEIEDH

DLMRMRYLHGTIPKRVTVDDPLDLFGFMIRELSLAPHKLDNLRKILSEISRQDLVEEIMF

FEDTVNPDQCNYSSFSKETSGPKFEPASLESSSFPSSLQNQEPSSQQEWGPSYQDTEHQQ

SGFFQQESLGSSSQATGVSSWAHSSGDSIEELKAHNVTGASGEPATATGPGLSTATITTT

TTTAPTAPVAPAATVGGFVCRPIPPQAEQQDVRKTLTLEDIEMFEDAPAVGTGSSEATGA

TGNINTESKQMANAALDLQERYKMSAEPRGLCLIIDVQKFSKPHLRFRYGSQVDKARLHK

TFTRLKFVVTQMDNPTAAQIDDCLLQISQLDHSAYDCLVICVLSHGGPHYFYGSDGVSFD

ENRLFGLFSAKACPSLANKPKLFFFSYCRGLQEQEGVPVVETDCPTVVPNPVMVPDLSDF

LFGYATPQHFVSYRNRSNGTFYIEVLTDVLDKDADRLELQHLLLRVNRLVSNKPTTTTHR

QIPKPEHTLKRQLYFL

>RUS82552.1 hypothetical protein EGW08_009682 [Elysia chlorotica]

MQSTNSIDLPYILRSVSGERGDVRPGAQSEPNEYNFNYKKRGLFLLINNRHFDEEIGQQD

REGSDVDAERLEERFQDLGFEVRRYDDVTCSKMSSLLYNAASLDHSDHDCFGCAILSNGR

EGVVYARDGQVPLDLLVMPFKGDRCPSLVGKPKLFFIQACQGRRNSRFLEPSSVGRAVSR

SSSPGSPSPSSHRRSPVSPSFNAEDDMPEPGSHVHMRRIPVEADCLFCYSTVPGAYSWRS

SQDGSWFIQALCIVLENYGTKMELSHMLTHVNRIVAYEFASCTEKGFGEHTKQLPCTASM

LTRYVYFRPKKQDIR

>RUS82190.1 hypothetical protein EGW08_010072, partial [Elysia chlorotica]

ISRVLHLTNGTKTVDLEKSAFVPEYRPSQLFVHPNNPLDLDLGFRFRKKRIKAGSSKMAQ

EYSSLQEDDRYPFRHGRGLAVIIIDGSSIGQGGREGSEKDSSMLQHMFTRLGFKLEVFEN

YNSQALLESLSRVSSEDFSEIDCLAVAISTHGGEGRTDAQTTEDFISTSGTKIKTSVILQ

MFIDEKCPTLVGKPRLFFLQACRGYLLDSGQEVQISSALERTVNLHTGTAQGPTRPMTRQ

DGAGSPVEMGIPLQEVKIRPVPCYKDFLVMYATPPGYFAFRNPDNGSRFINCLYQVMSNV

DPSLNLTRCLTRVVAAVAMETSYCPVNAPSRHNKKQTPVIYSMLTKDLFLLNRK

>RUS77802.1 hypothetical protein EGW08_014441 [Elysia chlorotica]

MTSLHEELSQKVNVTDGEENLDTVEGFRKDVKKKISKFLCGVRDATEESRRAPEVAPSPS

SSLSSTDFAQIRTNFVRMEPHGAGGEEDSDVYDFNFARRGLAIIINNEDFSSSSDFDDRP

GSSYDASALFHSFTHLGFNVILYENLAAWKMVEVLRAGARDYDHVNADCFACAILTHGDQ

TWSDREYDRMGTTVRQDLLFGTDGKAVSTRTVVDLFNDGASPGLRGKPRLFFLQACRGNK

LDDGQDVHVVTPVPKMIRRRGSRDVPDATDNPADGEDTLEGRGGRNSGAEVSAPLAERQT

SLSSSVDPPGFPKTKDMVDACMASDDEDENNETTSHQQGNVGASGGQGQTQNSNTSYLDA

GGEGPSEISSVDIGMGRWRRVREVPVSPAPLYKDCLVMYATPPGHFAWRRKNGAWFIQSL

CKVLDSRHLGNLSLVRALVQVGGYVARHYQSENPSRPSMHAKKEMPVIESMLVKDVFLKK

KIT

>GFS12612.1 caspase-3-like [Elysia marginata]

MSSDYVDASGEAGEGNDQPDARGEPRRDLHRPAPPLMTESDFVSDKYNMEHAERGMALII

NNRTFHPKTGMGERTGTDVDAQKMNELLTTLGFVKIRPLDDVSVQEIKNALYQAAKEDHT

DRDCFVCVILSHGEEGYVYGTDDKVPIDELVLPFKGHNCPSLAGKPKLFFIQACRGQELD

RGTTVTDAGGGGEEMEEEVIVRRIPSEADFLMAYSVVPGYFAWRNSARGSWFIQAIFDVF

MKYWKSLDLLTMMTRVNKKVAYDFESNASAEYMRRKKQIPCITSMLTKDVFFMQNKQ

>GFS16061.1 caspase [Elysia marginata]CARD

MDRADQELLRRNHMPLKKAIAPGVLNVLEDLESCGIITLDMKERTEHESRVPGDQASKLL

YLIRTRGKYAFQALYDCTVSAGLFDAADILKPEKAPHGPSYVGRAEPLPSSVPDEFSEDE

DLPDCWPPQKLEELDVRVKVKSFQDSKVAERRKKFYEFACSINGQKTHYPVQHKELSQCL

IISNEKFDNEDLENRDGTVRDVTALNSLFYGLHFDVTQKKDLKSEDMKKIITDKIVKYDH

SKSDSCVVVILSHGADGVVFGTDGRYDENTGKPLNGVEVEWIRQSMCAVPSLLKKPKLIF

IQACRGKKKDKGIPANSSQQETGSNQSSTSKEKTGGGTTDSAGQNPSGQSESGDSPNSNG

GGCQDSGASSNPREETPDSRDQDIDIRSRKNQICKERQAEDNVDAMGEPVPSASDVFLAW

STTPGYVSYRNTKQGTWFIQALVWVFSRFAYKHNINDLMTKVHNLVSRGEVEKTGYKQTA

ESTTRLLRTFYFFPKLKKEENL

>GFR67360.1 caspase-8 [Elysia marginata]DED

MSSSVAQWLARRTHDPAVTVDGGFGDWTDWTECSVTCGGGVRSRQRFCNNPAPQYGGAQC

TPNIYVEEQMCNTGACSPNGYAHGVAHGGTCAGQRGLFDCKNGIQCVSILEKCDCMPHCA

DASDEDVEYAGCLIGMEECLNSADNDDADDDDGDVDEDDDDDKMTMMMTTTTMLMTTTMG

MMYMMLMMMIIMKSITMIEQSYYRVLLQDLHSELLDEDVKCMKFLAQPLIQKRLYQNIQD

GLGLFGALEDCGMLSNTNLSYLSELFEVVGRIDLQARISEECEVSSDAVSGSEPLVCPFR

RLLYKLHREIPDTDLSRMRYLHGNVPKRETVTDPLDLFGFMIRERSLTPENLNTLKNILS

EVSRQDLVEEISHFEVTVRPNECNYLFKKGTGPSFQEASVERNTFSSSSSKDSNSHPEYG

MSYQDSDQQMPGRVFQQESVESENNTSLVTGLSSYTYAGQGIEDLKAAHFGGDAAATGMD

SIAAAAPPSLAPFLTEARLQSAAPGAMGNVCMNPAPRFNGSSLDRYKMDAEPRGLCVIMD

VQSFTETRLKFRAGSEYDRSSLRSCFERLKFHVEEMVNPSASNLYAYLKDISEMDHTAYD

CLVICILSHGGSKYFYGSDCREVYEDTVFDLFSAKACPTLASKPKLFFFSYCRGQEGVPL

IGTDGPSVAIDTDGSSVADLSDFLYGYATPQNYVSYRHKSKGSFYIQVLTQVLNEEAESL

ELQHILLRVNREVSNMPTGTAFRQMPEPKHTLQKQLYFL

>GFS10900.1 caspase [Elysia marginata]

MKTILRKFSQLRSLATVSSLAVAVLTHGGANDCLFGKDGQICRGQPVEGTFITKMDIIHI

FKGSVCRAMKDKPKLFILQACRGEEENPVGEGETGASSGAAGLSNERPPHNFRQDAGPEN

VAPTSSIQTRSRSPLVGRTGAVADTSDMCFVHSSSYGYKAYRTRNLGSPFIAIFTEKVRQ

LARYEEFKEIIRKVQKHFNKGPLSNNKMTLPDFSVQLVNQWFLNPPRWANTL

>XP_041377847_1 XP_041377847.1 caspase-2-like [Gigantopelta aegis]CARD

MDPQHKNILITQRANLVENLNIRDGLYTQLITRGVLNTRMVRTIQHNHSPDQQAEELLTI

IPTRGPDAFSKFCDALIADDQQHVVDLYLKIQSKTSGDKKKSEQCSDKDGDGTGVKKRPR

SPSPTIICFKDGTSNEVVLKYLSQGLPAKKSHSFSSCESSFSEFPGEVRLNMKHYVSSPV

SGFVDPFIPVALKKHSESPSTTHVGFLPQAYTERVSFTRCDTIDTMHKNTNVPMSSHSTV

SPSIEPDPVLFDETEDLNIDLTDGPVCVQVAMSSRQFYLTNHKKSYPMKKLPRGKALIVN

VNEVLGKPPRHGTDIDRDNLCNLLKQLHFDTIVYNDSDGLTAEDLRSKLKKFSGLPEHQE

SDGCVICLLSHGDEGFIFGTDGKKIPMDEIFEMYDNSRCRGLLGKPKVFIIQACRGGAFD

KGVSFMQDAGEIDGRPPVRPETSRQIPTMSDTVICYPTQHGYYAWRNCERGSWYIEAIVQ

VFMRHAKNEDICTMLNRVNRVVSSKVSRCPQIEMDRMCQMSEYKSTLTKPHLFFFPGIGC

P

>XP_041364768_1 XP_041364768.1 caspase-3-like [Gigantopelta aegis]

MADNSDQVDAAAGGKGQVDETDARGTRSGAGDIGQTEYGFATGELPVEKESSLLSDEYKM

NYPNRGLAVIINNRDFASHTNMGTRTGTDQDKININNRLRVLDFTTEIYDNMSINDMKML

MKKAGKLDHTRNDCFACVILSHGEEGTVYGTDGPVEIKDLVEPFKGHNCPSLAGKPKLFF

IQACRGQSFDKGVTVCDALGGKGGSLPMEPVTYRIPTEADFLMAYSVVPGYFSWRNSTHG

SWFIRALSEVLKEHADTLDLMTMMTRVNKKVAYEFESNAAKEFMNQKQIPCITSMLTKDV

YFRPKPH

>XP_041362615_1 XP_041362615.1 caspase-3-like isoform X1 [Gigantopelta aegis]

MDFKFFKKWRKYKPHIQYSDIPDTEQEFARRSADTKNVRPGAPSEPNEYNFNFKRRGLFL

IINNKNFDSSTGQMVREGTDVDAERLEERFQDMGFEVKRYDDVTRAKMSALMYDAASYDH

SNADCFGCAILSHGVEGRIFATDGYMSLELLVVPFKGDRCPSLIGKPKLFFLQACRGSKM

DYGVEVSDSTSASSKDEDETDSSNTVTIRKIPIEADFLFAYSTVPGYYSWRNNQDGSWFI

QALCIVLENYATKMELLHMLTHVNRIVAYEFASCSDKEFNDRMKQMPCVVSMLTRYVYFR

HKKQDIR

>XP_041374485_1 XP_041374485.1 caspase-8-like isoform X1 [Gigantopelta aegis]

MMSRGMQMDAEDEGGADFRADLLSIDGDLHVEDINTLKFLIQDFAPSKKLETINTGHELF

DYLQMSGRLNEQNKWVLAEMLKRILRLDLLDTFKVDEKDLDQYLAKFGSHFTRYRLLLLR

LAVDIGVDDLKRLLFCSTDIPRSKREKIATAMDFFTFFEQSKKLSPDNVDLLEDSLHHIG

RCDLSQVVTDYKQSSLTNYSNNFPTGAPGNYVCGTKPAYQSAKFNPPASQAPVTGLPYKP

AGPPARPAPVTGQQAEFSKPPALSHTGQAVPDPGSRSQPYDEQTPEQDAQDTLMAQQSDR

IDVDPEPEPVPYYKMDAVPRGICIIINNRDFKQDPDDPDSKALPSREGTDVDAKNLQDTF

QRLQFIVKCHTNLTDKGMTDLASQVSLEDHSAFDCFVCCILSHGVEGKLYGTNGRLLSIT

DLTGPFKSQTCSSLAGKPKLFFIQACQGTEEQCGQEIQTDSPYELVDVDAPPLTKVIPNE

SDFLFGYATVVGFVSYRSKTQGSWYISKLTEMLNKYAKKEDILSLLLLVNNEVGTEYAMR

SRGVCKQSPAPVYTLRKKLIFR

>XP_041374777_1 XP_041374777.1 caspase-8-like [Gigantopelta aegis]

MLKNNVEEDSTTPGILSRFVNVSKRYLGKAAVAVAEEAAQVVREKFGEPSSRDQDYYKME

ATPRATEPAQVVRETLGEPSSHDQNYYTMEATPRATEPAQVVRETLGEPSSHDQDYYKMD

ANPRGMCIIINNRDFYYDPSDSGCRNLGQRNGKRKLEETFEKLGFFVSTHENLTDKEMKD

TAETVGKMDHDTFDCLAFCVLSHGKEGEIYGSKGRAVNMSELTGPFKSQRCPSLAGKPKL

FFIQACQGRRTQGGRRVQTDSQPFQDENVTRGNNVIPEDVDFLIAYATVPGCVSYRSTQK

GSWFIYHLTKILNE

>XP_041354906_1 XP_041354906.1 caspase-3-like [Gigantopelta aegis]

MYSKLHLHLETMAKTKVDVFGSSSELKRLEHEEKITFKEYKYRCYYAVKGYHVCGVYWDV

CIWLTSFPSELQEMRDFRNDSTFCTYSMDIITLESSSWIKAIIRKTGRDGFFSLVGKHLN

DVLVLISESETKDKKQVAPTEQTLSIETPDEEQVTSKSGSSTEHQHEDPSSTFGEKKTST

AQEVGDTTTKQMKACSNSTSSLSTANKKADSILTPGGNEPGIQEELVEMPTKPMKPSSYK

MTSTPRGICMIISNEDFYSLTDPTKLEKRHGALQDVEKLSKMFTRLQFKVVLFRNLRDNE

VVMTAKHMSLNDHSHFDCFVCCILSHGTQGSVYGVNHVAVPISDVLSQFRSRACHTLAGK

PKLFFIQACQGLEPQIGVAVTDTPSVPQPAKLAADGTDFLVAYSTEPGYVAFRHRKSGSY

FIDALVTNLEGLPDFSVHDVMAKVCDDVSQLNINNGFKQVPQFTSTLTKQLVLARGSLAV

QSHCSGETGVQQEDATSPTLIHTFNDDVSSHQPKHTKKVTKTSVCVIS

>XP_041378642_1 XP_041378642.1 caspase-3-like isoform X1 [Gigantopelta aegis]

MSQRPLEDSGGRSHTNIINCNNVIVAGTIQCANVVVGGAQGVRVSPKTAHKDLTDEYYKM

DANPKGLCLIIQNENFSGGDIRKGAHKDLERMENLFKKLEFDVKVSPDRTANEMKDDMAT

EARHFKADFDCLVCIIMSHGEDGLLFGTDLEYVKVNDLSSYFQPDRCKNLIDKPKIFFIQ

ACRGEKWQVGRTKYDSTITSDVKAEKDSPGEVELDTVGAQSQLPAGIPNEKDFMFGYATV

PGYGAVRDENSGCWYIQALADVFEKYADSEDLNSMMTRVNAIVNEQRITKKGSAPVVQTP

VYHNALTKKVVFY

>XP_041378266_1 XP_041378266.1 caspase-3-like [Gigantopelta aegis]

MSRRLREASGGRSHINITNCSNVMVVGNIQCDSLVLGSHGVRVSHKTKTTAPKDLTDEYY

KMDSHPKGLCLIIQNENFPGGDTRKGAHKDLERLEKIFKNLEFDVRVSCDLTADKMKDVM

AKEAREFKADFDCLVCIIMSHGKDGQLLGTDNFYVKVNDLSSYFQPDRCKNLIDKPKIFF

IQACRGENYQIGCTSYDSTITSDVKLDKDLPEEVDLAAVHAQSQLPAGIPNEKDFMFGYA

TVPGYRALRDENSGCWYIQALADVFEKYADSEDLNSMMTRVNAIVNKQMTTKRCGPVVQT

PVYHNALTKKVIFY

>XP_041355471_1 XP_041355471.1 caspase-3-like [Gigantopelta aegis]

MSNTRAVIRNRLEIVSRQIDLTELNNQLEERRKKRMKTCYPVEIDIYLDYSTWESEEEED

MEEEENARHIWLCRVYDENPYRCRLSDERSRFVIQFGTLMELISMVRKIAFPNASFTQNR

PLQIVDTCKEFPVYSMTQKRRGICLIINNVDYADPSKVRSGSQKDCKDLKATFRKILFKV

IVLKNLTDSNIISAIKKMSTRDHSQYDSFVCCILAHGGRGTVTGVNDKPVHLEKITSNFR

VCDCPSLAGKPKVFIIQACQGSREQRGVELHQSDSISDDTPLMTLADGADFFMAYATEPG

FKSYRNPRTGSFFINSLCKNINQYKSHELSQIMRKVCAEIAHITINDDMKQVPWVSSTLT

KDLVLWPVYTNKTVT

>XP_041370746_1 XP_041370746.1 caspase-3-like [Gigantopelta aegis]

MDPAMINNLPLQIGSDSVDVSGLAKQLDQIKVHELNEISKDSVRAGSVQGEYGDIIIIIR

DNIRWQILETVNCFKCCVLSLNNGSYVCLLSYNNTSFVVRFSNLKDLINIFEKVAYPHLP

LAKNHPLIISNCSSKVPIYTMTRFPRGIAMIINNVVYNAPGLEKREGSGVDVKNLKDMFR

ISGYMTVVQFNDLTDSEMMIEVKKMAAGDHREYDCFICCILAHGGRGTVKGVNGKDVQLQ

TIFSQFRGPNCPSLVGKPKLFFIQACQGSQEQRNVVAPRYQSDVIRDETPVHMAAEGEDF

FIACSTEPGFVSYRHPEKGSVFITSLSKNVLELCDHSLSDIMHTVCSDIAKANEDSERKQ

VPWYSSTLTKTCVLRPLYVNST

>XP_041366892_1 XP_041366892.1 caspase-3-like [Gigantopelta aegis]

MLNNKFFSCLFAIQIAVDMYANKLYLFLLVFDVISIMLSMRFSPIGHECKQYKMDARPKG

MCVIINNKDFTSRRMSKWNGTDEDVRKLKSFFKQCEFEVKIFENQIQQDMLKELKQISLS

DHRAYDCFVCFILSHGQHGSVFGTDGRPVSVAEIELFFSARQCSTLAKKPKLFFIQACQG

GAPAQGVNVDKVDHEGSYECDVTCEFPVHEHADFLTVFATTPEHFAYRDPTNGSPFISTL

INVMDRCRGWSLDDIFNKVRDDMDKCVGQKQIPETKSTLTKLVVLT

>XP_041356116_1 XP_041356116.1 proline-rich protein 36-like [Gigantopelta aegis]

MAAALEEPPAELPRGPPISMAAALEEPSAELPRGPPISMAAALEEPSAELPRGAPISMAA

ALEEPPAELPREPPISMAAAFEEPPAELPREPPISMDAALEEPSAELPHGPPISMAAALE

EPSAELPRGPPISMAAALEEPPAELPRGPPISMAAALEEPPAELPRGPPISMAAALEEPS

AELPRGPPISMAAALEEPPAELPRGPPISMAAALEEPPAELPRGPPISMAAALEEPSAEL

PHGPPISMAAALEEPPAELPRGPISMAAALEEPSAELPHGPPISMAAALEEPPAELPRGP

PISMAAALEEPPAELPRGPPISMAAAPEEPTAELPRGAPISMAPALEEPPQRCILNQLTL

PPVVDESDQYKMDARPKGMCVIINNRDFTWPGLSRRDGTDEDVAKLDQFFTRSHFKVKVF

ENQTDSDMLEELKRISSLNHQPYDCFVCFILSHGERESVYGTNGRLVSVSEIQSFFSARQ

CRSLGKKPKLFFIQACQGREAAQGVNVEDTDHGGSWEYDVTVENPVHEHADFLTVFATAP

GFIAYRNATSGTQFISTLIDMMDKYHGLSITDTVTKVTNVMDMGAGLKQIPMFKSTLRKR

VVLN

>XP_041353231_1 XP_041353231.1 caspase-3-like [Gigantopelta aegis]

MERNIMIYVVGPEGVGKSTLVKSFTKGKVFRKRVEGFRCWATSIKTRDTDDEQIDIYEFR

GFKGNVSAGPSLTFWTVHVLIVMCTQHSLSESCKNFKFDFWRRRNTQVHLVQNIRSSEIR

DSRTLETEGKHFGLTNTYQLRVSSKGEVRQLFKKIIGDLHQGHPDFPRDSQHVSITLDDS

GYRSKASTAENQRPVSNNILQKYEMGVKPRGTCIIINNEDFGSLMEKRTGSDKDKERLRE

LFSDLDFDVHIFTNQTRSDILDILKRESTKDHSAHSCFVCCILSHGAQEVVFGTDGNAVD

ISELVAFYEGDQCRGLLGKPKIFIIQACQGNKQRPGVETLNGSDGGLRTDVSVCKTMPVD

ADFLIACATTSGSLAFRHQRDGSHYISTLVEKIRQNLAKSLTDVLTMVNSSMAQQDYEFE

GQRSKQIGCYKSTLRKQVVFGGIGQTR

>XP_041367451_1 XP_041367451.1 caspase-3-like [Gigantopelta aegis]

MMEIIFYMSRLFPQIALNETLIYYIGLIISIFFLNLYFECVSAINEHSTMQEQPNDGDRD

IRTRSDKYYDMEANPKGICLIINNECFTNSPTRKGSAKDVERLKCLFEKFGFVVEIKKNL

KSSDLLEELKSVSVVGVKPEHNCFVCFISSHGVGKSIIGTDEELVSISKIKEFFSASQCR

ALVGKPKMFFVQACRNDQKKPVSLWRSDVTGEPQSDFSDRHTDSDFLLAFATTPGSDAFR

DEAHGSLFIYHLVRILDENSHEDLLSVLTQLNDIFSVPAKLGVKQMSKFESTLTKKVVLG

KRSCN

>XP_041364823_1 XP_041364823.1 caspase-3-like [Gigantopelta aegis]

MAERLVDEYDFSFQTRGVALIVNNQRFEEHDPQLPTRAGSRHDTENLKLIFQKLGFEDIR

IKIDLTAREMTDWLESVAEEDHTYHDCLACALLSHGNETFTRLTDRVSQRHDVISGTDDD

ITTNHLIQIFNDANCPTLAGKPRFFIMQACRGEQLDDAVEVAVDEVDEVDGMMVEPAYTT

TPVPCFKDSIVVCASAPGYASVRQPRGGSVFIRNLSDVILNYDVTQLELQKLLTRVRQQV

AVGFESKSSRDPSRHMKKQMPCVYSMLTKDVYFRNKSNVTTQCA

>XP_041364822_1 XP_041364822.1 caspase-3-like [Gigantopelta aegis]

MADDSADMHVAALGRKFIGKVASYIKPNTHEASSQAVVMDLADEYDFSHRSRGIAVIITN

SKFEATTQRAGAENDTQLFKEAFKHLGFTDIRVAQDLTASEMKWALESIAKDKNVLQNSD

CIAIAFSSHGDEQMYIRHDNASVREDTIFGTDVPIRTVELMEIFSDFQCPELIGKPRLVF

VQACRGQKLDRGVEVVKDMDVTDSIMDVTVSPAPCFKDFLVIYATPPGHFAFRRPAEGSW

FVKCLADIMKLADVEKMSLTKILTRVVSKVALEYQSGHSDPRLNAKKQSPCFYSMLTKDV

YFRAKHQHKEEPMIVD

>XP_041364306_1 XP_041364306.1 uncharacterized protein LOC121379720 [Gigantopelta aegis]

MGSRPGSNCDAKALEKMFKRLGFTIKRYNNLEAKQMVTKLKKVAANRSFDHRKADCFACA

ILSHGNHVDTKSDVFEKRWDDVIFGVDGVLVSTKYLLDLFSDDACPELAGKPRLFFIQAC

RGEKTDSGVKKIVLKDKVSVGEGDNNSAEDSKSDGVGDQGDSTVETDVVGSRPLQHLPDH

DKQLEQMVATNGSKPASDSDTNHLTEVEDDRLTDGDNFDSDSSSESSKTEDQTDTIGTKG

NASELGAKLEEKKKRKVVTLVPAPLFKDFLVMYATTPGHFAWRRKTGTWFIKSLTEVMKV

NRNKNLVQTLTKVSRKVALDYESNIPENKKMHAKKQVPCIMSMLTKDIYFTQKT

>XP_041378464_1 XP_041378464.1 caspase-9-like isoform X1 [Gigantopelta aegis]CARD

MDDQHREILRRNRSAIIREIHDIGTIVYALEENGVITNNMLEDIMAESTPTDKKREMLNI

LSKRGPKAFDKFYEILDNNKEGSAADILKPELSEQRKAEEERKLAAVGRAKACVPLDDDD

DYDLPETWPDMNRLSEDVTVKVCEKSGQMYERFIRSNTDSGQTYKHQSKPRGLALVINNE

KFELPKKNGSNLADRHGTNIDQSTVSQLLEQLGYTVCIRKDLTAEKMLSTVNEFSQLDHS

LLDSFILFILSHGEKQGICGVDGKLVQKEEIIGAFNGKDCPKLSGKPKLFFFQACQGHKF

HHGAPIASAEPVSSSSGSDPDNMEHTLKKMKDLKLNKTSSDESRDDTVADIKVETSVPAG

GDVFIASATIPDYLSYRDEKKGTWFIQAIVYIFKNFAHKEDISKLMTKASLIQLSRPIWS

MKVTPTDDVPLSHLVTNGVTLMEYARANDSLVTCAPMSDDSIVEEVLAARTPVVAEPEPE

SDTENDDTADLPPPSLTSATDTLELLTRFVDSRDSNGNPVHAIGDLTNFLSSVAFMPCCN

ARQTLLTDFVN

>XP_041378168_1 XP_041378168.1 caspase-3-like [Gigantopelta aegis]CARD

MDDIQTEAIRRSRVFLVKTLILDVLWDYLLEKGIFNDEMKERIQCASTPSESKRRLISDL

LRRPREAYEDFLTCLDLADHKHCADKIRENEYQVRREQGCLSGVPDNTPHNEVPALSSYC

PPQVNLALTQEATSDVDMSDRSNTTPYICSPTPDTDCNPSDGISAYALNRRVSSNKVYEM

ESKPRGYVYMINNHQFDNFDARNGTQIDCRKLEKLFSQLGYFVSTDRDLKAEEMKRRLKE

FAKFPENLKVDSLIVVILSHGEQDDIIFGTDGESTNRGFKKYLTIKHLKSWFGNRRCPEL

EGKPKLFLIQACRGQSEDIGSASASVGSSVQRDVVDFPADTTADIMNESSLPEQSDMIFA

HSSVPGYVSYRDPEQGSVFINALVNVFTEESHREHLADMLTMVNKKIVDEFHTPEGLKCL

SSPVNQLTKKWFFNLDQAPEEWNKP

>XP_009060090.1 hypothetical protein LOTGIDRAFT_106792 [Lottia gigantea]

METEEAFYSEEYAMNKPNRGVAIIINNVSFHPSTQMGERTGSDKDHKALKQRFKELEFDV

REFKNLKRDDMESLMKKVAGESHDNNDCFMCAMLSHGEEGTIYGFDGPIEIQRLIEPFKG

HNCRSLAGKPKIFFIQACRGSKLDAGVEQQDSKGSRESETIEYKVQRIPTEADFLIAYSV

VPGYFSWRNSANGSWFVQAISEVFRKYGQKLDLMTLMTRVNKLVAYNFESNASKEYMTNK

KQIPCITSMLTKDVYFRSKN

>XP_009060089.1 hypothetical protein LOTGIDRAFT_209997 [Lottia gigantea]

MSDKGDGSGDNVNAVVPDNANVGAISLGQGECSVRGSDTGATGGRLSEEYAMKTENRGIA

IIINNVLFDPETEMDERKGGEVDQEALKQRFTELEFDVREYRDLTVAKMEKLMIKFAGES

HKNNDCFMCAVLSHGDKEYVYGADGVSIRTEDLIKPFNGDNCPDLAGKPKIFIIQACRGS

MVDNGVEIYSDDNGGRETFEIAQIIPIEADFLKAFSVVEGYVAYRNTEKGSWFVQAISEV

FRKYGQREDLITLMTEVNRLVAYEFELKAGEKYVKQMPCLTSTLTKKVYFKPKTGYPAN

>XP_009065630.1 hypothetical protein LOTGIDRAFT_133150, partial [Lottia gigantea]

LKDHENSDACFICLLSHGEEGYIYGTDGEMLALNQIFDLFNNNKCPLLKGKPKIFIVQAC

RGGSFDRGVSIDETDGGPAIPMIKQLPTLSDMIIGYPTQAGFYAWRNRERGSWYIEAIVK

VFIKHAKHDDICTLFNRVSIHYTLPLFDIFTRTVLASTGQDFSM

>XP_009060261.1 hypothetical protein LOTGIDRAFT_105903, partial [Lottia gigantea]

NFNYKNRGIFIIINNKHFDSSTGQDTREGTELDAERLEERFQDLGFEVRRYDDVKRTKMT

GLMYDIASQDHSNNDCFGCAILSHGKDGKIYATDGILPLELMVLPIKGDRCPSLIGKPKL

FFIQACRGINLDYGVDMSDASTTSSKDEDYTDAASQTVCIRKIPSEADFLFSYSTVPGYY

SWRNNQEGSWFIQALCIVLENYGFKMELLHMLTHVNRIVAYEFESCTDKEFSDRTKQMPS

VVSMLTRYVYFRPKKQDIR

>XP_009066726.1 hypothetical protein LOTGIDRAFT_134425 [Lottia gigantea]

MHEILQIYQMNSNPRGHCVVINNEIFHKSTELDDRDGTEKDVISLEKLFTQLHFLVDVRN

NKTAQEIVDIVQEYTDMDHTNYDCFVMVILSHGSHGNVYGVDGKEVDISKIVFSFDSIYC

QSLHGKPKLFFIQACQGITQKLDELDLQDIEQVDEALHNTARRADMFISMATSPGYVSWR

NINIGTWFIQAVVYAFKYWAHSCELVNIMTKVGTKHGG

>XP_009062283.1 hypothetical protein LOTGIDRAFT_210703 [Lottia gigantea]

MASLYKMNHPRRGKAVIINNVNFNTQARLNRRKGSRQDSDSLADCLRGLNFTAEKHEDKT

TGEISKIFEKVAAEDHSDADCLVVSLMSHGGKEGISGTDFNRRNFMQLQDERDGNYRSAG

VLELSDIIDPFKGDKCPTLVGKPKIFILNACRGDDMDYGVEGPIDQVDSVTSRYTRVQRL

PIDADIFVLFSCTEGYTTLRDEISGSYFVQDLCRVFLEYGKTLHLNELVLKVKEMVAARQ

TVVQGSAHLEFQMPVVLDTLRKLVYFDHKTSITRQPSADITRMIRLFE

>XP_009066755.1 hypothetical protein LOTGIDRAFT_134426, partial [Lottia gigantea]

YQMNSNNRGHCVVINNEIFEQQSRRYGTEKDVESLKELFKQLNFQITVHDNKTAQEIHNL

IKHYKDMDHTNYNCFVMMVLSHGSHGNVYGVDGKEVGIEQIVISFDSRNCKSLCDKPKLF

FIQACQGGRYTKNDDEDSSEISLPTKTRALTFLFYISINYISGFVSWRDKNFGTWFIQAV

VYAFKYWAHSCELAVLMTRVGTN

>XP_009059943.1 hypothetical protein LOTGIDRAFT_73381, partial [Lottia gigantea]

SYDFNHEKRGCCVIINNDKFTDMYDRPGSSCDVKNMETEFRNLGFDVKIYNNLTASYIDK

KLRKVSMKYDHEKADCFVCIILSHGEEHFDLKRERNDVIFGSDGHHISIAEIMERFSDPY

CHGLLGKPRLFFIQACRGVKTEQGVDITIKSIKKVEVPVHPQKPIDRKKLIYSFSCILGY

YAFRRESGTWFIKSLTYVLQNSLRSKTFQQMLTHVSYEMAMNYKSKNKKNLNIDGCVTIP

CIMSMLTKDIYF

>XP_009059944.1 hypothetical protein LOTGIDRAFT_74313, partial [Lottia gigantea]

NFNHTKRGIAVVVVNEKFEGHSKRAGAELDRELLSYCFRSIGFDVRVFTNLNGFQLVTEM

EKIALEMTRDPDFDCFVFAMSSHVEEDIIHATDGYIPTERILELFSDENCKGLIGKPRLF

FLQACRGNLFDIGTTVIDEVDSASDKMVITRIPCFKDYLVMCATPPGFYAFRRPSEGSWF

IIELCRALQKAGSNQPNILKHLRYVSKCVGRNFESKSEQSLYNAKKQSPVIYSMLVKDIL

FN

>XP_009060095.1 hypothetical protein LOTGIDRAFT_154127 [Lottia gigantea]

MCENSIRILDQKDCKSPLCVEGDRYCFHRKNIGKALIFNQSVFHHLSSRPGSKVDRENIE

RSLTHLGFQVDTYVNLSTEDVKSVLDAESARDYSLENSFMCVFMSHGSEGVIHCEDGPLD

ISTILQYFDAKNCPSLFGKPKLFIVNACRGDKMDPGVDVVDSIEDSLTFPIPFTLLDEDF

ILVYSVVSGYASARDFRTGSCFISALSETLLNHGKSMDFVKLMTRVNAKIAFEFERKGGK

KQMPCVYSMLKKDLHF

>XP_009055665.1 hypothetical protein LOTGIDRAFT_53303, partial [Lottia gigantea]DEDDED

NYRQKLVQLDSELDSDNVKCLKYLCFDVIPASKLEKITQGTDLFELLEQKGFLGNGNYFF

LAECLSKIGRRDLSEVALDHNQLITTLFEFVLFRSLWFDISEDLDAEEVCKCLYYYGDVP

KSKRGGIKSGIDFLNFLEKKKKIGPDNMDYLLDVLKTIERNDLVQMIQKYTERISGDFIA

GPPPKSIGPQSYSMTSPLGNQNKINDLIDALMKSEMYRMDRYPRGMIIIIIIIDFGLKLL

MGPFLFIADTLDRTFTELGFQVYVHNDLKAVEMVQILTRYSQEDHELFDCFVCVILSHGT

RDYIYGANCEPVPIRTLADPLKNLSCPGLCGKPKLFFIQACQGQDKQHDVVRVDDLDEDA

PPNGEVIPNEQDFLIGMATIPGFVSYRSKTSGSWFIIKLCEALQKFAKYWDILHILTYVN

SEVGKANADIDGVRYKQSPGPLYTLRKRLLF

>XP_009057120.1 hypothetical protein LOTGIDRAFT_121371, partial [Lottia gigantea]

FKNLGFIVKLHNNLKGDEIIQEMKSYAREDHRQYDCFVCVITSHGIRDEILGSDGVSVPI

RSLTDPFKAIPCPDLCGKPKLFFIDACQGHERQQGESFGKASPVDPVVEVIPNEADFLMA

YATVPGFVSYLDNRTGSFFIQTLCKALEEYAERLDILHILTMVNSAVGEFNITIDDSKCK

QSPGPLFTLRKLLFLKPPK

>XP_009066758.1 hypothetical protein LOTGIDRAFT_236948 [Lottia gigantea]CARD

MAMTREKREVIQKNFTKLTENIEDIDRVVSNLLSAGVITTNMEEEIMAKTTKRNKIRELL

NLLPKRGQVAFDKFYTALIDSEENVAAEFIEPHVINRSLVGSRLENQVTNPENNRDELPA

VFPPDQEEEVVVVPFSDADSVMTANYHKSLEVNNDEVYKNTKGLYLSAIVYKNTKGLYLS

AIVYKNAKGLYLSVIVYKNAKGLYLSAIVYKNTKGLYLSAIVYKNTEGLCLSVIVYKNTK

GRCLIICNENFDDVTRLCDLDWEKDVGNLMTLFEKLRFSVDIIKNCTVQDLKKRLDRDKQ

KDMKDVDCLVVFVLTYGENGQQVYGTDGTSMTIQSIVDYFYPTSCPGLSTKPKLFFIQTV

PRDLEGMDSTDGTPEQGEISLKKGVNKIQIGQSETSDSDVEARNMPSRGDTYLCVSATSD

KVKPDESDSGSMFILALVYAYKNYSYSNELGKLTLKMNALIKKGEIKNGRPVTIGSSVVS

SLAKSLYFFPKNCP

>XP_009060096.1 hypothetical protein LOTGIDRAFT_228989 [Lottia gigantea]

MAEERIESFDETNWMVSDEEGDVDSSDSEFEDREGKVVDISVCEEVKQKLKEHYSENSHH

EYEHYNGKNLMIVIQNKDFHESRQRHGSTADVENLKKTFKRLGFIVKTFKNLSGEKMDVK

LRKARQKFNHKDASCFACVILSHGNQYETGDYERKDSVLGVDEQAISIEYVMKIFNNDNC

PDLMGKPRLFFIQACRGKGYDKGAETLVPVEDSSNSNVEQISSELEETYIEATSGDYSPP

VTTFTYPTPCYSDSMVMFASPPGYAAYARKGGSIFVSCLCDEFQIYNRKHNFFQILSRVA

NQMALDFETSNVVEDTDYEKCKIVPCIHSCLTRDVKFQILDD

>XP_009060091.1 hypothetical protein LOTGIDRAFT_228982 [Lottia gigantea]

MNEDRDFDCFVLAISSHGEHLEKTVGDKTIVSDVIHCTDGYITTQNILQIFSDENCPGLT

GKPRFCFIQACRGRQLDPGVLINNHNIDINYEYDKFGYHNKPFTNHIVTTPFPCMTDFLV

MYATPPGYYAFRRMKEGSWFVLALAKVLRSACRDKCDINKLLRQVVKEVSYGYQSESNNY

ATDMKKQAPVIYSMLTKTVYMCPK

>XP_022345548_1 XP_022345548.1 caspase-2-like [Crassostrea virginica]CARD

MEDADRRTLQLNRVFLVKNIANPDDIAEGLYSREIFTEGMKDEIEVEKLREKKVRKLLDV

IPRRGPDAFVAFLDVLLETKNEHIANHLKSGISVQNKSNHQQDEPEEDLPQTWPDPLSLE

GPLTVKVITTQNRRPKPSWCYKMGKKPRGRVLVINNKIFFGPLEEDQFGRKRVILPTREG

TEQDQRKIEEVFKQLNFTVDVFQDKKGEDIRDILQQEVKRDFHREAECFVLVIMTHGTHG

AIYGVDGKTIPIEEIKRQINGENFPAMADKPKVLLIQACRGEERDEGVRENEDINQKIKE

AEKNLRDLSIHTESDSKRESTIPTMGHMVVAMASTLGMVSFRNRIYGSWFLQAVAYIFAK

YSYKDDIHQMLIKVNELVSRGRAKDSTVKKEFVAVSEFNSTLSKNLYFFPGVVTEEIKSI

GIQQASD

>XP_022343665_1 XP_022343665.1 caspase-6-like isoform X1 [Crassostrea virginica]CARD

MAREDAWKDCIRRNYSFFKNNIADPLDVAEYLFGETTPPIITDNQKEMVKAERTTSKKVG

KMLDILMKRGPRVPQHLYRAFVETDNPDCADKIAPYLIQLEKQALGKDPSEWPPLRDEHK

SMIKDHVTYLKPNDPCCYLQYANDNEVYSMRRERRGLVFLINNEQFYELQRRDGTRFDRD

YLKSLFIDLHFEVEVENDLTAKEITEKTRFISRSDKVKDSDCFIFIILTHGDEKGVCGID

GISVPVTTLTEMFEPNNCPELNEKPKIFLIQACRGDKKEMVIPMEQSGDAPSQPDGGGDQ

IDASNTPDINVDREPNISATATVHSKADFLIAYSTPEGSLSWRKTDAGSWFMQAIVWIFK

YESHLKDLVEMLGKVNNVVSKGKATDGQTHFVTVSNFNSSLRKKLYFFPGIYANKAVHPS

CI

>XP_022321371_1 XP_022321371.1 caspase-8-like [Crassostrea virginica]DEDDED

MNDIETDNTELVTEDAAPQQGFDFGQFLVALDSRLDSSDVENLKFLLRGFIPDSRLQKAD

RGLDLFTELQNTGVIDPDENDLELLEECLYRIHRKDLLKTIARNTEYVDRRLRKLQEDIK

RNVTHLNSSRFLSPFRVLLFEIGEDIDDEEFKSIRFTLTSLVPKGQLSKMKSIFDLFLFL

EKKDEMSHTDVSVLQRVLKVIDRADLIPKVNQYISAQSPSTKSKSQNDGHSQMEVEPTQV

PTIDRPPQPTGMPNPPLYEGLLISLADAISKCAEWKDILPFLGLKDSPIAQTINDEQVMP

SLRVLRMLQYWQTNIMQNDQSCRVTLVGVLQKFRMTQGLQILGQTVEEDDHIPNPTPVDL

RSLYQPPDGDIPSSIRHPAPEPLPNIQALRPVTSEIETQTSMLILPNSNQPIRDRNEENI

SVSNTEARVGVSERQPHLSENNHVYTERDMRDARIDENLSRAMRRTHIDDEMDLPSYKMD

RKPRGIAVIINNMKFFKVPDDPESKEMPDRTGTDLDAEKLTYIFQKLNFYVRRFDNLEDF

KMNRRLVEVSFEDHSQYDCFVCCILSHGVLQNIYGTNGRLVRIAELTGIFRSGKCPSLAG

KPKLFFIQACQGREKQGGIDIEMDSDLEVDHGRNSEMIPDEADFVLGCATVPGYVSYRSR

SQGSWFVNKLVEMLDKYAGRYDLLSILVKVNEEVGRANARLEGGVYKQIPAPLVTLRKKL

FFR

>XP_022325102_1 XP_022325102.1 caspase-8-like [Crassostrea virginica]DEDDED

MSQSGDFRELLLDIDAHLDSKDLELLKFLCRDVVPFSVLEKCKRSLDIFTNLEQRGKLKE

GKSIYLQECFHYMKRKDLIRKLGGDPQKLEDRIRYEGPAHLFPFRVLLFEIAEDCGKEEL

EKMKFYSKTTLHLPKKMTEKVKNCLDLFSALEKEELITNIQVRTLEQLITITEDEELMDY

LYEYKNKNLVSSPTDEMNAPLEIKQDLGINHCQDTPAVATQGQQQPAMNLPFQSERTSSP

RISYSSLNSMTPPAQDAGESLLSRQQFPQTPQVTPEPSDDFINFYKMTSKPRGYCLIINN

QKFHGEIKFVERKGTDIDEEKLKNVFGKHLHFKMDVTKDATSQQIKNICRDFSKKDHSQF

DCIAVCVLSHGTTGAVYGSDCKKVNIQEITSCFTASRCPTLANKPKLFFFQACQGLTHQS

GLQRNGGSTRVVNGGIQTDITPEPPREEGEEVIDDVATPQSMIPDESDFLLGYATVPGYV

SYRSRSAGSFYISILAEKLEKHALGSRSQDLMSILTEVNNAVSELHYRDPQGVMYKQVPA

PQFTLRRKLQFDKC

>XP_022316669_1 XP_022316669.1 caspase-2-like [Crassostrea virginica]

MEEAMCIESPQNQDVADAKPVSETDAFPFGSGKSETHSPKPVATTIQTTSEFYSHVYKMN

YPNRGKAIIINNKTFDRNTGLNERRGTDLDSSALNCRLSELDFDVDLHHDLKAEEIKNIL

KKAADLDHKDNDCFVCAILSHGEDGFVWGKDKKIPINDLMEPFKGNKCVSLAGKPKIFFI

QACRGTKFDEGVGMNLADAKNVGDTNMDVDPQFSLQKIPSESDFLIAYSVVPGYYSWRNT

ISGSWFVQALSEVLMKHAQNLDLLTMMTRVNQIVANKFQSNTSNPNMNEKKQIPCVTSML

TKEVYFKSKFMEQPMSPESPQNHDVADAKPVSETDAFPFGSGKSKPHSTKPVASTIQTTS

EFYSHVYKMNYPNRGKAIIINNKTFNPNTGLNERNGTDQDASALCCRLSELDFDVDLHHN

LKAEEIKNTLKKAADSDHRDNDCFACAILSHGEDGFLWGIDRMIPINDLMEPFKGNKCLS

LAGKPKIFFIQACRGTKFDEGVDMNVADAKGFMDVEPQFSLQKIPSESDFLIAYSVVPGY

YSWRNSTNGSWFVQALSEVLMKHAQTLDLLTMMTRVNQIVANKFQSNTSNPDMSEKKQIP

CVTSMLTKEVYFKSK

>XP_022313378_1 XP_022313378.1 caspase-3-like isoform X1 [Crassostrea virginica]

MMTLCKSLESLTSSRQSLAASVENQESPYLIRKVSGDRRSVRAGAPSEPNEYNFNYHKRG

LFIIINNKNFHPSTGKESRDGTDVDAERLEERFQDLGFDVRRYNDVSSSKLLQLMNDAHL

IMSSIDILTAAQLDHSDSDCFGCAILSYGIEGRVYATDGMLPLDVLIVPFKGDKCPMLVG

KPKLFILQSCRCSNLEQSVEDTVSYKSFLSEDSERSFSGMRKIPVEADFLFFYSTVPGFY

SWRNHQEGSWFIQALCIVIENYGSKMELLHMLTQVNRMVTYEFESCSDEEFTDDVKQMPC

IVSMLTRYVYFRPKKPDIRD

>XP_022292891_1 XP_022292891.1 caspase-3-like [Crassostrea virginica]

MLYKVGEKTGKLQLNALRAILLSQGVEDEIQDESMGIWKCLDLLEDCLAEEEFFSLLKVV

YKNDAQCLQLLTDFLNGSVTAGAIGGQIDKYPIQEVGEEDHPDGNPLTEKIRESELLPYD

LPLRDRFSMIPDEIKMGSTKPMSSQLLVYDRGKHPGRCVIFNNKTFVKEATHPTRDGTDK

DKDLLEQMFKTYGYEVFVFNDKSCPEILQILETFQRRDHQDYGSFIVCTLSHGDLDVISG

SCGGSIDINLMTSLFRADNCPSLAGKPKLFIYQACQGKSFQNVWHRTDNIIIANGDTHLT

NEKPDMTATVPDFYAMCFAPVEADFLIAHSTMPGYVSLRNEDGSFFIQSLVKNIKELSPR

EDLVSILTEVNRDVSEKIGKTSDERLASQIPEARVRLTRKFYLTQVKSNG

>XP_022294791_1 XP_022294791.1 caspase-8-like [Crassostrea virginica]

MEESFNCFEENNLNHCQDVTTKLLFLAKDVIPEAVREKMKQGQVRSALETNVVSLWIDGK

DPPFGFIKEWLLMEGEELFNIRDNDMFDDHCHVDSVGISEKRKMLYKVGEKTEKHQLNAL

RAILLSLGVEDKDQDESMGIWKCLDLLEDCLEEEEFFSLLKVVYKKDVQSSQLLTDFLNG

SFTTGAFRGQIDKYPIQEVGDEDHRYPDAYPQTDKIRESELLSNVLPLRDMFSMSPDEIK

MGSTKPQSSQLLVYDRGKHPELLEQMFKTYGYEVFVFNDKSCPEILQILETFQSSDHQDY

GSFIVCTLSHGDLDVISGCCGGSIDINSMTSLFRADNCPSLAGKPKIFIYQACQGKCFQN

VWHRTDNITLANSDTHLTNGKQDMTATVPDFYPMCFGPAEADFLIAHSTMPGYVSLRNED

GSFFIQSLVKNIKELSPR

>XP_022294613_1 XP_022294613.1 caspase-6-like [Crassostrea virginica]DED

MESRNVLIEIHNEMGKDDLKMFSYLTKDYISNRERKKIDSALDLFAALEKRSVITCVEGK

EFRVGFLIDCFTLMRRRNLVRKLRPQSILGQQSTMESVLNEKRKLLFTVGEETGGSELDA

LKTQLRVSSMEVGKRALDQIVDVWDCLDLMEERMEDGELISLLKRVYVDDDKLPGLLGDY

TRGLVSAGATGGQVDKVPVQVEGEEEEKQALPPQTEEREEPQGLTFNPPPSGSSTMTQDG

APSGFSIPQESRLGVYDRGRRPGCCIIFNNEKFVNESTHPPRVGTNEDRDQLVYVFGTLF

GFEVSIFDNKSREEIMEILTNFQTRDHKDHGAFIVCTLSHGDHTGVSGACGQTIPIDSMT

SLFRADSCPTLAGKPKIFIYQACQGQGTQNIWHRTHDAIKAEGDTHLMNGVPDVTSPELP

QLLEPLVTPAEADFLIAQSTMPGYISLRSGTGSFFIQSFVKHVKQHSPREDVVSILVEVN

RDVSQKVGQIEGGQLAVQIPEPKVRLTKKLYLFPVESEV

>XP_022338210_1 XP_022338210.1 cell death protein 3-like [Crassostrea virginica]CARD

MMDKAKKNILIKNRVGLVENINIEDGLLTQIASQQILTQRTIKSIEAKKTSFEKVEELLD

KISLKINGFEALRDALIANEQEDIVDLYLKLPTEEVEKPQKEETVISQESQTVSQDPSRP

QSSPPRLPAQGAQVWHSPAKTVQLQSHTESSTARVISLGQPDSHGIPIHGTVSNVVQPVP

SSSPPDHVFHPIQEPSVHSVPPSSLVSSLICVSGSASKPYDFSNAVFLDRGSKSSTSFKE

NEQRFNLKHFINDPNYQVPLEDAPKTVPWSNPTMINNNINAGHKEVVYVQSYRPVSLQRQ

DTTDAMTKPLGNHEGNCESPAKRQRTQEKIEITTFEPNQNSLNDSMSSQAALQPVKLQGE

LDDPEIDLTDGPVTVRVDHSTRQFYISNYQKSYAMRRLPRGKALIINVNEVEGKPPRRGT

DIDRDNLYHLLTQLHFDVRVYNDRDGLTAKEIAQKLEVFAADQDHRHGDSSFVCLLSHGE

EGFIFGTDGRKLQLDSVFKLFDNSNCPTLLGKPKIFVIQACRGGALDSGVPFLDEHDGSV

GSKQLPTQSDMIICFPTQAGYYAWRNRERGSWYIEALVQIFMKYAKSEDICTMLHRVNLL

VSRKVSRCPQIEMDSMSQMSEYKSTLRMPHLFFYPGIGSI

>XP_022286252_1 XP_022286252.1 caspase-6-like [Crassostrea virginica]CARD

MEERERTVLRRSRSVFTRDLVDVHTVCDRLYQREVLSEGMKSEILSETTRENQVRKLLDI

IPKRGQKAYWALFDVLEETFQHDLSSILGPGGRISPHKLLDARSDVYTRSPDVPRLNSAM

ARQSLAESVHKYRSQPMESNWEAPSPPAPVKEVLDPTGTCGLCTETEREDPKQSNLKGEE

KEEGFVLPEDWPSEQDLAQSINFEVRLSRKSRPRLVETEPDCYPMRGKARGRLVIINNFQ

FGQEVDVERMNRSQQDATSLDVLFKQLHFQTKQHSNLTLKKLSSVLEEERTRDQASLECF

VMVVMSRGDGEHIYGVDGYRINTDSIISMFDCEHCSNLRGKPKIFIFQACNDATKSTDNF

EKNYKKASMSSSFTLSDKLRSDMFILKATKDVEVRYGGAFGSRFIQSFVYLMRNLACEED

FQEIVTKINPLNATNEPEIVQYASSLTKRLFFNP

>XP_022343382_1 XP_022343382.1 caspase-2-like [Crassostrea virginica]CARD

MDQIHKDALRRTRVALVKDLNVDHISDDLLAKDIFTPLMMEYIMSERTRIDKVRRLLDDL

VRRGPNAYKSFLEVLNDTRHDFLAEKILRNEEIIRREKDPHSSIPVQEVNPGISNSVQHL

SSVSLGSSSTPLESSILNSRNSELQSYHPPSSGLIVSEMESADSNEDDILNLDPSQPLNP

TSMDIDDDDSRLSPDDLQSGPESQPVQLSDDNKTQYNLGPSSSFSTYDSFYMKDKCYKMD

SDPRGVVLIINNSKFEKDYGERQGTEHDCRKLKELFEGFKFKVFVKENLRAIEMMREFRE

FSKNPWLEKVDCMVLAVLSHGQKDNVICGIDGRGIDVLQEIVPIFSSKECPSLAGKPKMY

ICNACRGGKNGQALEIRCSQSIQRDELQRDGPNSFSENKIQIQETRDILLCFSTFPGYAS

YRDPDQGTWFIQEFVNVFAEQSHDEHVLDMMTAINKRVSQKQHRDHIDEAAWVQMPFPAS

SLIEKWFLNPPPLAGATGFSH

>XP_022320316_1 XP_022320316.1 caspase-3-like [Crassostrea virginica]

MAAKKIPSKAKDLHNKLVADLHDHINAKVDEHDFEKLKHRLKLKLTPMDLKACNNIIELT

DKLQKRGHVQPGKYEEFVNLIECIDTNWVDIIREKGEEISEIIDKANEPQLSLPVKESEK

SSESKKRKMEEENPDPVQCKDGPSTFQNTPSAKRKQKSFYRKTGKQKGLALILNFRKGRH

GTQRDFENINDFFQNTLNFECHVENDPSKQDLEDFLQETAGLFNNKRKAEEYYCLLVFIM

SHGNKDGILTDDDKTISLDDIVSYFKNDKLKYFAGKPKCFFVQACRGNTPQSTVDVSDDK

TEVMLDEVVGHISIPTDADLLLTYATTPGYYAYRRPAIGSWFFHNLIDVLKRNYQEEHLE

EMLIDVKEILAHDAEWQTNEGYKQMPCTWSTLTKRFSLVP

>XP_022342791_1 XP_022342791.1 caspase-7-like [Crassostrea virginica]

MDSEEQEKDFIEELEKLALESEPDGYNIFKKTLKAFTARKSKHRKDKDESISKSQTDEFS

PDRTPKHGHPREPHQLQEISESASGYRRPNDPALPDDDRYLFQHKKRGFAVLIINSKFDY

QPERPNADLDRYSMSKMFQQLDFTVILLENLTSSTLLEKMIDIQQRITPETDCFACVISS

HGMEGHAKVRGQGKNDPYVRTEHFICTNDGVVQTTELLDLFNDRHCRNLRGKPRMFFIQA

CRSRVDVQRDDEVDMGVEVPVQFVTPPASSGSSRDTLRPPTTGDVGVNYDEYQFRKPAAC

STYPIYENFRFNIKANNHVGGQNVQAPVKEEERKEPPTRTTTYKDPNPEPEPEPEKIEIF

QIPCFNDYLVMFSSAAGTTAWSDPGKGGWLMYCLYHVFHDVTYTEDDLLKILTKVCGKMA

ISMETHCPKTRFDKAKSAACVYHRLFKDIYLQPN

>XP_022323042_1 XP_022323042.1 uncharacterized protein LOC111124471 isoform X1 [Crassostrea virginica]

MTSSEIGLLYVSNVKQLADKVCTCLDQCKCKHHSIQENDIIPSGVPKLTKGAEKHLKGVG

ALIVLADNKHWENFWTKDKRMRNILEDLMYQDDDDEDNKPVSFCLFVGCNLDQGRPIKKE

EHETYFSEDFSVQVSYKLVSVDMSEQSITAVARGIVEHITNQVLPTRMLGPVGRQDTIDS

PTSPPCTPIQNIEPKLYQPVDTAGSDNYVTVSEGEDPVAGGEDPEENCYNLRTRPGLCLH

ICLLEFDPEDVNKTLEQRHSSSVEMEKMQSTFERLGCFVKSVRNPTDEEALEFLQEGKKF

CEELKPSFLVLIISTHGQESTYTIRNAKEHPSGVAASRREREHQLFFHNGKTLQTREIVK

RFEPPSCKSLADKPRLFFIQACRSIFDDENHENFDRGVMIQLQSSATREKSIEVPRQKHE

RLQDTADAKGRISEPSEQEMNERVQEVLKWSEEELETKRQILQYIKWLSESNVAKEENPD

GRKKLSDALTKKRFSQPKRDKILAASEGYEQDILDHFFNEAIVSEPAPCGNDTLIMFASA

PGKEAYNRAKQGGWLITNLEAQIKETMNKNPERVDLLAELTNVSHRMAYEFETETNRQDD

SGHKSVPCLYHKFCKDFIVWPEEMKRVKELCEKDDKWMDFFKMKP

>XP_022323040_1 XP_022323040.1 uncharacterized protein LOC111124470 isoform X1 [Crassostrea virginica]

MTSNKIGLLYVSNVMQLADKVCTSLDQRKCRHHSIQENDIKPSQVTKLTKGAGEHLKGVG

ALIVLADNKHWEDFWTKKNKMQNKLEELMDQDDDDEDNKPVSFCLFVGCNLDQGRPIQKE

EHEAYFSENTSKVSYKLVSVDMSEQSITAVARDIVEHLNNQVLPTRMLRPFQSQDTIDSP

TNPPCTPIENIKPEWFQPVNAAESDNDTIYRVTVSEGEDPVVGGEDPEENCYNLRTRPGL

CLYIGLLEFDPEDDKVELEKRHASSAELKKMQSSFERLGCFMKSVTNPTKEQALEFLQKG

KKFCEELKPSFLVLIISTHGQEVTKRIRNADSAKEHPSGVAASRHEREHQLFFHDGKALS

TRDIVKLFEPPSCKSLADKPRLFFIQACRSRFNDKNHENFDPGVMIPVQISASGDNALEG

SGQKHERKRPLLDTADAKQRIGKPSEQEQNKRVQEVLQWSEEELETKMQILQYMKWLSKS

DVAKEENPDGKKNLSDALTNNRFSQPQREEILASSEGYEQDIFNHFFHEAIVSEPASCGN

DALIMFASAPGKEAYNRAKQGGWLITNLEAQIKETMDENPERVNLLAELTNVSHRMAYDF

ETETNRQDDSGHKSVPCLYHKFCKDFIVWPEELKRAKELCEKDDKWKAFFNMKP

>XP_022333891_1 XP_022333891.1 caspase-3-like [Crassostrea virginica]

MSSRLDAGPDSNSEISGERHVATETELTSRRGVVNTYQSVEQSEIQRQEFFDGPTVIPTQ

DCDLGEGEFVVVDQHKPGLALLLGFETFSNGEQSRPGVVRDLQNMYQLFHTYLGFEVQTM

KDLTGAECEEWIRKVTSDNDLLERISCVVMVISSHGSEEEIQKEKTPGMEFRQKEIKYYQ

HRISTRDGSIETSKIVNMFDNPKCRALRGKPKVFFIQACRSRTDEAKSVYDFVDSGVPIR

IQSEDGYEAQPSKGPTLSTHHNPVVQCESDESRVDAKGIRAENIHFRGKRYMDLLQPAHS

TMEDQIVVAPPPCLHNSVIVMASPPGKAAWSESIEGGWLIMGLYQVLMQYVRDRHKIRLL

QALTRVNYFIALTLQSNTNDPSLNAMKCMPVMTHTLEKDIVIVPKNLQRIDTPPDLQNTV

IQP

>XP_048751990_2 XP_048751990.2 uncharacterized protein LOC125663707 [Ostrea edulis]CARD

MEKSKKDILIKNRVGLVENINIEDGLLTELLSRQVLTQRTVKSIQARKTTFDQTEELLDK

ISLKSNGFEALSEALIANGQEDIVNLYLKAAQEDNETVKSEAKIESVISQESQTQTVVQE

PTRPQSSPPRLPPPGILLSHPTTKAVQLFQESDCSSARVIAQAPPEGVHTSQLVHLIHQP

PTDHVFHPIADQSVQSIQASSSAHDSSIICVSGNPSQAYDFSNAVFLDRISKNDKSFKEN

EPRHNMKHYIDDPNYQVLEGSTPKTVSTHIPLVFNNNIHPMPKEVVYVPSIKPVPLQRED

TLDAMTKPFGNHEENCESPAKRQRKEDKIEITVFESNQQNPNDPVMNAAAVQPVKLQGEL

DDPEIELTDGPVTVRVDHTTRQFYITNYKKSYAMRRLPRGKALIINVNEVEGKPPRRGTD

IDRDNLYHLLTQLHFDVQVFNDSDGLTAKDIAQKLEIFAADPAHRHGDASFVCLLSHGEE

GFIFGTDGRKLQLDSVFKLFDNSNCPTLVGKPKIFVIQACRGGALDSGVPFTDEHDGSSA

AFKQLPTQSDMIICFPTQAGYYAWRNRERGSWYIEALVQIFMKYAKSEDICTMLHRVNLL

VSRKISRCPQIEMDSMSQMSEYKSTLRMPHLFFYPGIGCG

>XP_048761694_1 XP_048761694.1 caspase-3-like [Ostrea edulis]

MEEAMSPENFTENANLDVADAKPISETDAFGFGQGTFKTHSTKPIATTIQTTSEFYSTRY

KMDYANRGKAVIINNKKFNPNTGLNDRNGTDVDASALCCRLSELDFDVDLFHNLKADEIK

QQLHKAAESDHSNSDCFACAILSHGEEGYVWGIDRMVPINDLLEPFKGNKCLSLAGKPKI

FLIQACRGTKFDVGVDMNVADAKGFMDIEPEFSLQKIPSEADFLIAYSVVPGYYSWRNST

NGSWFVQALSEVLMKYGQTVDLLTMMTRVNQIVANKFQSNTSHAEMNAKKQIPCITSMLT

KEVFFTSK

>XP_048763304_1 XP_048763304.1 caspase-3-like isoform X1 [Ostrea edulis]

MMTLCKSLESLTSSRQSLATSVENQESPYLIRKVSGDRRSVRPGAPSEPNEYNFNYHKRG

LFIIINNKNFHPSTGKESRDGTDADAERLEERFQDLGFDVRRYNDVSSSKLLQLMNDAAQ

LDHSDSDCFGCAILSYGIEGRIYATDGMLPLDVLIVPFKGDKCPMLVGKPKLFFLQSCRC

SNLEQSVEDTVSYKSFLSEESERSFSGMRRIPVEADFLFFYSTVPGFYSWRNHQEGSWFI

QALCIVLENYGSKMELLHMLTQVNRMVTYEYESCSDEEFTDDVKQMPCIVSMLTRYVYFR

LKKPDIRD

>XP_048776588_1 XP_048776588.1 CASP8 and FADD-like apoptosis regulator [Ostrea edulis]DEDDED

MNEIMNDIETDRPDLIPEEDAVPQKGFDYGKFLVTVDNGLDSNDVENLKFLLRGIVTDSK

LQKAQRGLDLFSELQNLGMIDIENGDLELLEECLYRIHRKDLLKKIARDTAMVEQRLRRL

TEELQNKKVTHIESTKMLSPFRVLLFEIGEDIDDEEFGSVLFALTSLVPKGQLSKMKSIF

DLFLFLEKKDEMSHVKVDVLLKVLKVVDRADLIPKVNKYLSNQNRRSSPTQYNGPSQMEV

EPPQVPQEPLNHPPPQLPPVIPGSSAMSNSPLSDGLLINLSDAISKSPDWKDIPPYLGLE

DRAILETVNDDQIMPSLRIFKMLQHWQRDIMQNDETCRVTLIGVLQKFRVTRGLQILGQT

VEGDEHIPDPNPTDLQSLYQPPPGQIPQSLRHPAPEPLPDIQSLRPNMSEMETQTSIMIL

PHGNLPIREQTEENLYVSNAEARVQLEQRQPPPSENNYSQSERSIRGNALDENLAQAMRR

TNLDDEMDLPSYRMDRKPRGIAVIINNMKFFKVPNDPESKDMPNRTGTDLDAEKLMYIFQ

KLHFHVRRYDNLEDYAMNRRLVEVSFEDHSEYDCFVCCILSHGVLQNIYGTNGRLVRISE

MTGIFRSGKCPSLAGKPKLFFIQACQGREKQGGIDIEMDSDLELDHGRNSEMIPDEADFV

LGCATVPGYVSYRSRSQGSWFVNKLVEMLDKYAYRYDLLSILVKVNEEVGRANARLEGGV

YKQIPAPLVTLRKKLFFR

>XP_048757091_1 XP_048757091.1 caspase-8-like [Ostrea edulis]DEDDED

MPQSGDFRATLLDIDAHLDSKDLEILQFLCKDIVPFSVLEKCKRCLDVFANLEQKGRLKD

EDSKFLQECFHYMQRKDLIRKLGGDPQDLERKIKTDGPSMLSEFRILLYEISEDCGKDEF

ESMKFYSRTTLQLTKKMLDKVTSCLSLFNVLEREELLTPIQVKTMEQLIETTGDEELLDY

LYQYKNRNLVCSPSEELGAIPGMKLDLKINSLPNTSPMVPQEHQEAPQNLTAQSEGANPQ

MAAYPSLTSMNPPAQDAMDTFMSRQLSDGQSPQEDHDFNDSLNRFYKMTAKPRGYCVIIS

NKIFHGKSDLKERVGTGVDEDKLENTFGSHLNFIVKTYRNCTSDEIMEVCRKYSGKDHGN

FDSIVFCILSHGTTGAVYGSDCRSVYIRDITSCFTASQCQSLASKPKLFFFQACQGLVHQ

SGLNNMGTTRNVNSGLQTDTLYESPRPQEPPNEETTSPPPNMIPDESDFLLGYATVPGYV

SYRSRSTGSFYIKTLVEKLKKYTLGPRSHDLVSILTEVNNAVSELHYSDSQGALYKQVPA

PQFTLRRKVHFDKC

>XP_048769780_1 XP_048769780.1 caspase-8-like isoform X1 [Ostrea edulis]DED

MSDNINRRCILQEIHDKMSHDDFRCLSYLSKDVIKNRKRMELRSSLDLFLALEEKTAINC

TSQNFSLLEEMFRLMGRKDVISVLTRCKQDAVQAEKTKTSMTDKRILLFNIGEETGEKEL

NYLKESLGKSGVRKRELDHVTDVWDGLDLLEERIRETDHLSFLRNTLGQVEEIKVLLDDF

IQKDEVSEGATGCQSDKFPVQEEGDMEITNGEEAKAYSNMQTGMSMDGFPSGFSSSRWSG

LGVYDKGIHPGRCVIINNERFDDESRLPTRTGTNIDRDRLHDMFTMFGFNISMYNNQSRD

EVMNVLECVQKEDHQDNGALVVCFLSHGSLNTISGSCGQDIGISNMTSLFRADNCPSMSG

KPKFFILQACQGRETQNVWHRGQTTIRAEFDDPRTPSLPEEEVTHESPSWGQLDIAASEA

DFLVAQSTMPGYISLRDEKEGSFFIQSLVRHLKEHSRRTTGPTSIKLGTQYPWMQGFTCV

QMKGHTLFKGEIITKMQKLSGNLKIFARTTVLEKLKFT

>XP_048756175_2 XP_048756175.2 caspase-3-like isoform X1 [Ostrea edulis]

MDDLEVDCCPVQHGPVSVDYTTMEIKRNSRLDARVVQAPNYMRIGDQSMQFHPRQYHKPT

YHSQQSLEIDGDVMSMEEDSPHLVNDDSRFLSEGVKPYIEPASLSFGPYNSGIKAKTNTS

RRPQEVPYIDIPSSVFHDTEYKMDHRKRGIALIINNMNYADVRQRPEREESIHDASAMEN

LLEHLMFTPVTQRMGHKNMVVNDVSSKRLEELIQDVTNPALNDFSDCDCFLMVVLSYGID

GSIMCCEGERDKLIPVEEVVAQFKADKCPSLAMKPKLFFMQLCPAFTYNIADSVGRSGPT

SMEPEVRKIPTDSDILVQISTLPGTYGWYEDDNALSWYIQGLKDTLQKNCYDPLRQGQRS

REINDILLHVNRLYISKLREQGICPSRHYSAPAITSTLTKDLYFLPKGIEAT

>XP_048760265_2 XP_048760265.2 caspase-7-like isoform X2 [Ostrea edulis]CARD

MEDADRRALQLNRTFLVKNIANPDDVADGLFTFEIFTEGMKDEVEVEKIKEKKMRKLLDI

LPRRGPQAFKLFLKVLMDTHNDHVVKQLTSGSSTTYTPPYHQQDEEEDLPQTWPDPTSLE

GKLTVKLCSSHNTRFQNGRYYKMGRNPRGRVLVINNKIFFGPLEEDESGKRRVTLATREG

TEQDERKIFEVFEQMNFIVDVYNDKKGEDIKDILRQEVKKDFHRTADCFVLVIMTHGTSG

SIYGTDGKTVSIEEIKKQFNGENFPAMADKPKVLLIQACRGDERDEGVRESTHINQEIKK

AEEMVKDLTLNSETDSHTEIVTPTMSHMVVATASTLGMVSFRNKMYGSWFLQAVAFIFAR

YSYKDDILQMLTKVNELVSRGRAKDSSQKKEFVAISEFSSTLSKSLYFCPGLVSEEIQQN

RIQQASS

>XP_048742450_2 XP_048742450.2 caspase-6-like isoform X1 [Ostrea edulis]CARD

MAGDQWKDAIRRNYSFFKNNIADPLDVVEYLYGETDPPILNENLKDQIKVEGTTSKKVGK

MLDILMRRGPRVPQFLYQAFVETENPDCAEKLAPYLLLQEKEALKKDPSEWPPPKKEHRA

MEKEPVTYLRPNDPCCCLQYNNTDEVYSLRKECRGVVFIINNMQFNGKMSIRHGTMSDRN

NLKELFQALHFEVIVENDLTKQEIEDKTRLISQSDKVKDSDCFIFIILTHGGDEGVFGVD

EKSVSVSTLTEMFEPNNCPGLNEKPKIFLIQACRGEKKEMVDPVGGSGNVDPVGGSGKSH

ADGDGRKEHVDTVDGPNVKSEPNISATKTVHSKSDFLIAYSTPEGSLSWRKTDAGSWFIQ

AIVWIFKYEAHLKDLVEMLGKVNNLVSKGHADHGGTHYVTISNYQSSLRKKLYFFPGIYG

EKRVHPNIR

>XP_048770132_1 XP_048770132.1 caspase-6-like [Ostrea edulis]CARD

MDERQRGILRKSRSIFTRDLVDVDTVCDRLYQKEVLSEGMKNEILSENTRENKVRKLMDI

IPKRGKRAFRAFFDVLEETFQHNLSDILCPGGQMDDLETKYNALHGFQSKRATDKPLLQR

SMSAEVRREPVESISEFQEQHKTCEFPLECKDFQKLAIVEEDFVLPSNWPSEQEHAQSLN

FKICKSRSSPASLREEWSRMYPMTRRPLGRLVIINNFQFDGEVDDERMKRSQQDTTSLDV

LFKQLHFQTKQHSNLTLEGLSSLLEEERTRDHTSIDCYVMVFMSRGDGERIYGIDGYRID

IDTIISTFDFEHCSSLRRKPKIFIFQACDDVAAKSKLLGAEKKNYEILTSSTTSEHMRTD

MFVVNAVQDVAVSYHGIFGSRFIQSFVYLMRNLAFNEDFEEIVRKINHLNSVSEPEIIKF

ASSFSRRLFFNP

>XP_048747578_2 XP_048747578.2 caspase-2-like isoform X1 [Ostrea edulis]CARD

MEQIHKDALRRTRVALVKDLDVDVICDEVLTHDIFTQLMMEYIMAERTRIDKVRRLLDDL

VRRGPNAYRAFIQVLNATGYKFLADKILQNEDIVRREKEPNTSFPVQATNEIFSEIHPSS

ISSGFTTSNLNNNARNSLIGPVNNNMQQNPVSLGSSESNLSSNLSSVSQSCSEPNSKTGV

GGLTVHEMASTLSTPSESTDDDTLDLLSGSLNPTSMDVDDDVDSRTEELLLRDTGNEHGH

PSISEGYPVQPTDTSHTAYNFDPPFTMMSFYEQSKSLDKCYMMESTPRGIVLIINNKDFG

QNHKERTGTEYDCQKLTELFRNLSFQVLEKENLTAMEMKEELKAFTQNRQLQAVDCLVLA

VLSHGNSDKIICGIDSEGIDIYEHVFPLFSAKNCRMMRGKPKMFIFNACRGVKNGESFDT

FRPQTSRTTIPEKDQLQRDGPNIEPSDSQVLDMQDLLLCFSTFPGYTSYRDPNQGSWFIQ

EFVSVFQEQCSTEHVMDMLTEINRRVSQKKHSDQAAEMCVQMPFPSSSLTKKWYINPL

>XP_048776071_2 XP_048776071.2 caspase-3-like [Ostrea edulis]

MDNNKMTSTKIPTEARDLHNQLVIDLHDHINSKVDEHDFEKLLHRLKLKLPPMALLGCKN

VIDVIEKLQKKGYVRPGKYEEFKKLIECIDIGWVDVIDDRAEKIIKIIDDANKSQKIASP

MEDQRKLDMKRKLEDVEQEAPSQDPDGPTHASKKRKSRSEYSKSDEQVGLAIILNFTKGR

LGSVVDFKRIQKFFRSTLKFECHIEEDLSKEALQEFLEETANMFNTKRTAAKYYCVLVFI

MSHGNEEGILMDDGKHISLDDIVAYFKNDKLKYIVGKPKCFFVQACRGSSPQGAVDLSDD

KEEMMIDEVVGKVSIPTDADLLIAYATTPGYYAYRRPTIGSWFFHNVIEIFSAHYTEEHV

EDMLIDVKEALAYDPEWQTNEGYKQMPCTWSTLTKRFFLKSSVE

>XP_048728167_2 XP_048728167.2 caspase-7-like [Ostrea edulis]

MSAPRSANNDRISVSCSELYEMLTDQSVEISSSRRFLEDQEERDFGGSRHWKGESSENLH

HLPKIKHNRASPDISEFIVGNRSKSFPFMKDKPISSPTTMRKLDARTLEKELLTQFKNRR

LNPHSHVQMGMNKSPRQRLQKSFYPSAKQLAVQSLEQFQQNADTATTTTADDRFAVDVIE

SSSNMHTTVLSEDCILDGTPSKTSTLPSRDYFDGLSNPRWRSLNDAADSGFYSMTRKPRG

YAVIINNYEFYKEMIAFTSSGPQTLRVIQKGLRRDGSIEDTDKLSRFFRDELHFIVRGHA

NVSSYDMIKILNDVSSADHDEFDAFVCCIASHGRRGTLFGSDCREIKVEDVTGLFNGNHC

VSLSGKPKCFFIQGCRGNHISQKGVLYGQRNESSQDEHYIPLTCDFLIGYSLIKECTEFH

ARQSGSLLYKTLVKYFRKYCKRYDLINIMVRVKKALAKLQENSVGAEDIFECNFRAQLVF

>XP_056001291_1 XP_056001291.1 uncharacterized protein LOC125664819 [Ostrea edulis]

MNERMEIFKLAQSYKIKWELFLDSKVHEVLTAHAQSLGCELDLLILPLFVSVASSMTSGT

KIQVTLSWIERPRLCGVVAARRGERKTTMLNYIKSCCRQVKCHGGDTDDTESVPGIPTID

EAFTENIPETCKGDRLFVLDDLDIVLKNVTEMPDIWNLRTLQRLFDSRCSDETDDAIKEN

VNVIAAAQGVVISDIVNSPDPGRLLDRVIVYCARDSEDMVISTKTPDSLGKLMQRIYDLH

VNQNVTYTFSEEATQEFQRMSGIVNAIKKQCHNDELKRGCCNHVMSQMARLSAISCALTS

SIKETKADFTINKTTAQSMFVVSLYLLRQKCLLLHGSDSTTYSELFHSILNSETELLPTA

DTSPRESEDDVIEITLDRNNEPVARIIPKREDNMEISETSQATDVRMWTMDNKDYTVNQN

GRETSSVYYPAVQNLPQEVATQQKRQRLMNHPEGPNTEHCQQYEHNHPTHSEDNKQIHFS

NTLQPRGKFRPPPPPLKLRPHQLHVNRRFIPAPKRLTDRTQSKTCVFTCSDSLFVHICCA

KIRKCLLTKGLSITSSYACQYRLFPPVPLTQRTSSPRTTHPSWAAVKFFERLEALGYGEV

TVFRSQSVKFRKKALEEMSEEARNILKYQIGITEEEYRSAMTKEEDIQGPVQEYNYVFTV

HQNGDIETEPMQHVETEQNVQHSAICEILTTCYRIRHLYVICLYKRTVRYCLIEIRPRKI

RISSSGRAFFRNIEYRINISTFLIMQERNMENHNDLDNTTEVQHATSRVQFSPVILENSV

SLWSDTEPNRNQNDCVYRFGHNKRGYAILIMNNEFDNEDENLPGVQSDIKNMSELFNSLE

FDVIELINLTNDQLITELTGIQDAVTNDCDCFVCVISTHGGEDPGYYIGDTEKLVTKTEH

YILNSKGAVWTRYIMDMFGDEQCRALRGKPRLFFIQACRGGEVDKGIDLLEDDSDDIEDD

SKNHAVFQIPCYNDSLVMFSSTSGKYAWSEDFEGGWLLSALQSVFPSSTDTDLLSLLTRV

CGEVARREAYIPDNERAHRSKSAACVYHKLSKDIYFPKSNKKD

>XP_048760572_2 XP_048760572.2 uncharacterized protein LOC125669811 [Ostrea edulis]

MDLEQQALQGNDEWERFEKSETKPDGFKAFKKTLKDFIGLKSKKKGGHGVPYQTQMDDDE

RSHVISDTKKESQTASNTPEVPQKLQNNELSWSYPESSNPRSQQSHNEIAEDNTYSFRHE

KRGFAVLVINSEFDTQSDRENAVWDIYYMRKMFSEMGFDVKILQNLTSNDLLSKLKRIQQ

QITMESGCFACVISSHGMEGRAKVSGLKGREPFVRTEHFVYTKDGVVRTSELLELFNDCN

CRNLKGKPRMFFIQACRSRLDVQKDDEVDMGVEIPVRLVHSTTNKDTTSDAGGIYDDYTD

RQSEDFFVASLDPQYENIHVYGLSRKYTGNYIQTGNRYSINTRSPEAREPPRRTTTYKVQ

KPETKPENIEVFQIPCFNDFLVMFSSAAGTIAWSDSGKGGWLMYTLYHVFNDVTYTDKDL

LTTLTGVCGMMATEMQTYCPSNLRNDKAKSAACVYHRLSKDIYLQPSSYV

>XP_056017527_1 XP_056017527.1 uncharacterized protein LOC125669447 isoform X2 [Ostrea edulis]

MAQFFKRFLHKELQRHVKIRNDVADDREYDFRHEVRGIAIVISNGVFSPDTGSVTEPRSY

AEEELCKMREMFSGLGFIVMLFKDLTARQMHDVIQTVTSQEMDFFLEKSDAFACVLASHG

NEIQEDSPADPAVKVFHHIIYGTDRAIRTHLLVDLLHDNRCKKLQGKPKLFFIQACRSRF

NDPNRFDSGVKILTSNKRRIPEVGHDKKESISINSTTSFADTKSPISDGLCGDKIKDGGR

KSDGSESSIEVIPCLEVLHEVTHDVSEDPSIAQRGIKWIKKLVFRKVDRIARNDVHDEDT

MTTVMTPCHNDFLVMYSTSQGKASFGREEKGGWMIHVMHRIMVQHVEFIQKDKENCLDFL

SVMTNVMRHIGFYFETDTGNPNTSGYKAAPTLQHKLSKDLIFRNKYVS

>XP_048734212_1 XP_048734212.1 caspase-3-like isoform X1 [Ostrea edulis]

MWSTLDADFVQGSDRESSGDRREVHETELGPLFVSYIDSAVVPTEDCNIVDNEEKNEFVV

VDNNKPGLALLVGFEYFNNGEQSRPGVKKDLENMYELFHTHLGFEVQVKKDLTAEECMEW

IREVTSKTELLQRISCVVMVISSHGSEDEVKSGIVNPDFRRQEVKFYQHRFSTKDGSIQT

SEIIDKFDDGKCRALKGKPKVFFIQACRSRTCENVSAYDVVDGGVSIRIQPEGSSRPHPS

TQHVLSSQNRPVMQNEEAHSHMDAKGVKCSNIRAGNKIFTEMSGCVRAPMEEEVTLAPPP

CLHNCMIVMASPPGKAAWSDTFQGGWLITCLYKVYMEHLKKGHKIRMIRSITQINRLMAL

SMHSVTDDPQLNSMKCVPVMTHMLERDIVLVPKNLQ

>XP_056010880_1 XP_056010880.1 uncharacterized protein LOC130051874 [Ostrea edulis]

MSGKVGLLYVKNKNTVQLLSDEITTLLGAENIVDSGVEEQAIKSGELTPVVKKVGKNLSN

VNSLIVITDNAFWDRFWSEKSNKVHNKLEELLCDDDGDGVTLVLIVGCNIDQGLIETLVP

SSSHAHVISEEDLKKYLSKERKPPSVCHMVVSIKENNSESIKELSKQIVEKYGNQSLEDE

DVCKDKVDTPTQTPTLTFPVTPQWPTTKERVVVQTAGPNDDASENPDIDFSVPSMKSDDV

DNNETRYDISRPGLCLYIGVLEFDPEDSKVKLAGRKSSMLEFEKMKDSFECLGCFVKRKS

NPTDTEALKFLQQGMEWCEKLKPSYFVLIISTHGQEVPQVNKEDKSAKLHPSGIKEKGQP

KKEMVHQLFFNNGKSLLTRDIVKMFDDCPSLADKPCFFFIQACRSRFGIDGHENFDPGVM

VDVFTPRSQIEAAQITPDTEDASGNPKLLDKTDAKRSTAKLTEEDKKARLQQIEKWNKNA

KIATLYTHILEYKNWLSNQTGASQTRSQDTSEETAKLAEALTVEGFTFQKVQEILETCKG

YEVDILDFFFPEMIISDPAPCTNDSAIMFASAPGKEAYNRGNKGGWLIINMEEQLKESLK

DNPENVDLLAELTKVSGRMAYKYETDTDSSESSGHKSVPCLYHKLCKDVIICPQELARKK

KLLENHDNSEYSEFFKWA

>XP_048765195_2 XP_048765195.2 uncharacterized protein LOC125673006 isoform X1 [Ostrea edulis]

MEVDSEEYTYKFEDKRRYQGLALVIANFTKGPHKRNGAEQDLKYMKKTFKRLGFKVFYEE

NVTTPEFLSLLNHYSKSDTLKNYDCFAFAISSHGLEREERKGGSLVHHHVIQMFDDEFID

TSQILDYFSPKKCKIMKGKPKLFFLQACRIPESKASKDHLDAGIGFDHGSRTHPHHGGRE

LEKGTAYRDKTDYRESVEVSNEEVSEGDVSDTDVEDDPDLVIEKPPSSTTLGGQDTGDTL

EVLDQRQTEIITNLKRLQNQTWQLGRVYTATGTNQIKESNYQANQIKESKYQATEGEDEQ

KIPLQESKKTANESGRMEGVGLHLPTESRGSGDELDPPRGVRLDQRAALNITVVPCHNDM

LIMFASPQGLYAIRSKDHGSYMLKILYESVERYYGHGKLRNNKTNFLEVLRDVTAKMSAK

TFFGSKEYTIVPCIVHQLSKDIVFRQGQMMVDSKFSLKLPSSLASIIR

>CAG2189547_1 CAG2189547.1 CASP2 [Mytilus edulis]CARD

MNDEDRRKLQTSRSFLVKNIENIEDINEELYTREVFTDGMKADVEVQDTKEKKVRKFLDI

LPRRGPEAFDIFLDVLNQTKNKHVYDRLMTGSPGTTYQRSYIAQPVNLPTIGGDNQLPET

WPDERSILEASQCKIKPHDPSLMQHAVSGRAYQMKRNPRGRAVIINNKIFHGPVQKIDGK

NRVMLAMREGTDEDVQGLHKLFTDLHFHVKIHKDKNAQKIVDILRDEANDVGFHRNADCF

ILIILTHGKQNAIYGSDGKTVEIRDIKDMFNARNFPCMENKPKLFFIQACRGEEYDGGRN

KTVPDASSTDCAMSGNVVNLALERTRLDFDQISINNANDTDATVEKSVATDSHRLVAMAT

TMDMVSFRNRILGSWFMQAVIYVFRKYSHNEDILSMLSDVNHLVSLGRAKDRENRDAVAV

SEFESTLSKRFYFFPQQCETVPIQASTS

>CAG2184393_1 CAG2184393.1 unnamed protein product [Mytilus edulis]CARD

MAVDPKKALIKNRVAIVRDLDNGDDVADYLYANKLFTEDMRDSVLQEPEKEKKNRLILDT

LGRRGQKAAKGLHDAFLDTENSELAKLLAPYIRVIESEENRYRPKEWPPNSEEQDKMQKD

SVRRIKDLKSPSYIHEYTDENVYEIRNKTRGKVFIINNVEFQTKLSTRQGSEKDAENMKS

LFTELHFDVVQKDNLTGQDMLKFLKEERNKVDWNAMDCVILVVMSHGHTTGIFGTDGQEI

ELKKITELFESSQCPGLNEKPRLVFVQACRGENKESDHIGTGVQVSGNDSVTEGVQAMSL

GKDQIDAVAQMPNEDLKEKKTVHSAADFLIAYATPEGTPAYRHIHVGSWFLNAIVWTFKY

HAHQEEIQHMLLKVNELVSQGRTMAGNLTVSEIKNNLRKKFYFFPGVYDDPPKFIDG

>CAG2231370_1 CAG2231370.1 CASP2 [Mytilus edulis]

MERHGVLEIKDIKSPLMHDYNRDDVYKLQGETRGKVFIINNTFSYSKEKKREGSDVDVKN

LTDLFQQLHFEVVTQTDLPAEGMLRFLEEERDKIKASKEIECIVLILMSHGEGAEIYGND

NKMIELTKIIEVFDSDHCPSLDEKPRLVFVQACRIDVQHTADELDRLRVKLEETKVQDKP

GDQLDSAPFQIEKPAHVKHPSADFLVVYATPAGTLSYRNVQTGSWFLCAVVWIFKYHAKH

EELQHLLIRVNRLVAKGKGPVRFGQKLTVSEVKSNLRKKFYFFPGVYGDPPKLFEK

>CAG2231374_1 CAG2231374.1 CASP2 [Mytilus edulis]

MGSNNYEWKNAFKKNRVMIVKGLANPVEVADQLYRMNIFTEEMRDEITVKPKTRLILDTL

DRRGQKGVAALHRAFLDSKNRDLAEVLSPYVRAVERIENMTEPKEWPPKIEENSTMMDDH

VIKIMDKKSPWYIHEYGKEDVYKIQNDCRGKVFVINNVDFHGQLLPREASNKDADNLTKL

FTELHFTVIPKTNLTAQEMLSFLRSEAKELVDEDRAECVILIIMSHGVGTKVFGVDCLPV

EIKALTDCFSTANCQVLQGKPRLVFVQACRSVETASIEKTLHLVQSGKQVEMECIVKIDK

PIKQVFWEMNKAGAKYLITETDNQDKYSLQSSKILQTLIVKKVVESDQGKYRCKVIHDEK

VSSVSNFTELKLDVKDGARDVIDSVPSIQEKENCVLPTSTCSDHPSADFLIAYATPEGNC

TTCMI

>CAG2199917_1 CAG2199917.1 unnamed protein product [Mytilus edulis]

MFETIKQTNELEQKTRKLLHTLNYRGKQGVAALHQAFINTDNKGTCRKTCTFTSKKSRKE

KTNRTKIRKSSLEKCNVLQHHMVNRFRMYLVQIITAWPPNVKEENIMLADPLIKITDPKK

SLVYSRIWKRIRVQDTGKCRGKVFIINNVDFGNTKENREASNKDATNLSKLFKDLHFDVT

RKDNLTGHKINATELKLNVQDDTGTVTQQISQVTMGATGGCDETDSEPFQQQTTGDSSLK

STFLDHPLADFLVAYATPEGSKAWRHQHCWFWFMNAIVWTFKYHAHEEELHHLLIRVNRL

VAKGVTWSNNMTVAEVRSNLRKKFYFFPGIHGDLPETF

>CAG2198417_1 CAG2198417.1 CASP7 [Mytilus edulis]

MADKNDTTDAKNMEDTVDTFPFSFGKKSVPVNIHMSREEFEADIYPMNRKNRGFAVIINN

KTFDQGVGLGARNGTDVDASALANRFAELGFDVELEDDVTTEEMMQIMKKYAYMDHSEND

CFACAILSHGEEGIVYGKNGKVEIKKLLEPFKGNLCETLAGKPKIFFIQACRGTEFDEGV

EMNVADAKGEMDVDRQISTHKIPAEADFLVAYSVVPGYYAWRNSIDGSWFVQALSTVLEK

YGNQLDLVRILTRVNKLVAYKFQSNTSNPYMSRKKQIPCITSMLTKELYFFPKQ

>CAG2237348_1 CAG2237348.1 CASP8 [Mytilus edulis]DED

MANLRLRQILVEICSDLGSDDLEKLKFLLRDVTGSAKLEMATMAIDIFSAIERNKHVALH

DGSYLAECFSLIGRIDLVRKLELDPRRVEVGMETDPRLFPFRILLYKISEDLSDKEFEQL

KYHSTSINVLKDMLKALHRGDLLELVVEYQAKYPHLIDGENNTDSININGATQTLESPVV

DQYPSLPSRPQNGGPQTTEPSPMHVYEMNAFPRGFCIIINNNNFYPQSGFGCRVGSDEDA

RMLDETFCRLGFIVKIYNDFTVKDMAAVMHGLSRLDHSKYNAVVVCILSHGGEHVVYGVD

GKEIAIRYLTSFFRSSKCKSLANKPKLFFIQACQGNERQTGINQTDLSIQADGPSDLEEG

QSFPETDAEDVGRSNNSKIIPDESDFLIGYATVTGYTSFRSRTRGSFYIRKLTENLNQYA

NSQDLISILTRVNHDVSTENIPAGNNYEISKQVPMPQFTLRKQLFFRPENLPHDRTMPQS

AAGLSIISQSIQAVRTIAQPTDATKCKTNNIDYINNSNLVKLTGCTKIIGNINVVEATFE

GDPYLNIPALHPYQLDVFKSVKEITGVLVVQGKHKDFKDLSFLGNLTTIYGRGAKRYQGA

SLSVAYSSIEALNLSSLKRIRNGNVVIAFNDRLCYANTVKFTNLFRRKEQLATVVKNRSK

LECDLTRKRCSTVCGTNGCWGPRRENCVGNITENNSIEDSFNIVDML

>CAG2231765_1 CAG2231765.1 CASP8 [Mytilus edulis]DEDDED

MANLRLRQILVEICSDLGSDDLEKLKFLLRDVTGSAKLEMATMAIDIFSAIERNKHVALH

DGSYLAECFCLIGRIDLVRKLELDPRRVEVGMETDPRLLPFRILLYKISEDLSDKEFEQL

KYHSSKALGIKRGQLQKVKHVFDLFVLLEKNLLISPASINVLKDMLKALNRGDLLELVVE

YQAKYPHLIDGENNTDSLNINGATQTLESPVVDQYPSLPSRPQNGGPQTSEQSPMHVYEM

NAFPRGFCIIINNNNFYPQSGFGCRVGSDEDAKMLDETFNKLGFIVRIQKDLTAKDMAAV

MVDLSRLDHSKYNAVVVCILSHGGEHVVFGVDGVLIPIRDLTMYFRSSKCRSLADKPKLF

FIQACQGNVRQRGINQTDLSIQPDGPSDLEESIPETDVVTVPTNNSKIIPDESDFLIGFA

TVTGYTSFRSRTRGSFYVRKLTENLIEYANSQDLISILTRVNHDVSTENIPAGNNYEISK

QVPMPQFTLRKQLFFRPENLPRDGTVPQSAEGLRIISQSIEAVRTIAQPTDAPSQSGGK

>CAG2226391_1 CAG2226391.1 caspase-8 [Mytilus edulis]DEDDED

MALTCLTKDILVDICSDLDSDDMDNLKFLMKDVAGLHKLMPATNALDLFSAIENNKHVAF

HNGRFLAECFELMGRTDLVGRLGLNPTQIEKEMGQHNDSVRPFRILLYKISKDLASDDVE

LLASFSLRVFKLTQQEKATIKSALDLFVLLEKRGIIEIDHTNLLKEMVKSLENRGDLVDL

VDKYQGPLYEMTSKPRGICVIINNKKFIKMKAREGTDVDQARLEATFKKLGFKIAVYRDL

TVKNMADVMKDISIMDHSKFNALVVCVLSHGGENFIYRMDDRGGKTGNSRRQLVHGDGDR

LIQGNFVYGVDDKPIPIRYLTMYFQSSKCKTLANKPKLFFIQACQGEDMEPGVLLSSDLQ

MDGLSVDDSIDLDVAPVSENTEEKDAAPHASDNQKVIPDESDFLLGYATVFGYVSYRSRS

QGSFYVKSLTENLDNHADTLDLQALMVKVNHDVSEENIGEQAKKQVPMTQITLRRLLFFF

PDNLK

>CAG2226387_1 CAG2226387.1 caspase-8, partial [Mytilus edulis]DED

YKTMASTCLTKDILVDICSDLDSDDMANLKFLMKDVAGLHKLMPATIALDLFSAIENNKN

VALHNGRFLAECFELMGRTDLVRRIGLNPTQVEQQMGQTNYSVRPFRILLYKISEDLASD

DVELLASFCLRVFKLTRQEKATIKSAFDLFVLLEKRGIIEVVHTNPLKEMMRSLENRGDL

VDLVDDYQVRCGGPSVPFYEMTSTPRGICVIINNNKFIKMKSREGTDVDQARLEATFKKL

GFMIKVYRDLTVKDMAGVMKDISIMDHSKYNALVVCVLSHGGENFIYRMDDKGGKRGNSR

RRQLVHGDGDRPIQGNFVYGVDDKPIPIRYLTMYFQSSKCKTLANKPKLFFIQACQGEDM

QTGVLLSSDLQMDGPSVDESIDMDVAPVFENAEETDAAPHASDNQKVIPDESDFLLGYAT

VFGYVSYRSRSQGSFYVRSLTENLDKHADTLDLQALMVKVNHDVSEENIDIVKGEQAKKQ

VPMTQITLRKLLFFSPDNLK

>CAG2224734_1 CAG2224734.1 CASP8 [Mytilus edulis]DED

MIEFESLQNHNNNNQTDLTRGLITIDGELESDDINSLKFLCQDHIAQKMLEKIKDGIQIF

NELERLGKIDTSQGQVDYLLESLCKCHRIDLIRKMGFDQIQVKNRVESGRSNFSPFRLLL

FEIAEEMETADLKTALFTLSSVIPRSKHDKIKSICDVFIQLERSGDLSPLNVDVLYKLLK

MMDRGDLMTKVRAYQGLNPLINQIIAQPNRQDDIDSAMPFYKMNARPRGLAIIINNQHFH

KIRNDQSSKEMPARTGTEKDAEKLDYIWRKLGFEVCLFNDVDSTRMNQIMVETGFLDHSN

YDCLVVSILTHGVLGHVYGADGRIVRISQLTSVFSGRRCPSLAGKPKLFFIQACQGREKQ

EGHAIETDAGGSMPEQDLNVDRPRELIPDEADFLLGYATVPGYVSYRSRSQGSWYINKLS

QNLDKYAYQHDLLSILIKVNEEVGRATANIDNGAYKQIPAPMFTLRKRIIFR

>CAG2253531_1 CAG2253531.1 CASP8 [Mytilus edulis]

MRDRFPGQLGNRKQELRRLANGKVKDCVVCSYRSNGRRKRSRLQCGRCLREKLDYIWRKL

NFEVRLFNDVDSSGMCIIMVEKAFQDHSNYDCLVVSILTHGVLGHVYGADGRIVRISQLT

SVFSGRRCPSLAGKPKLFFIQACQGREKQEGGHAVETDGKGFKPIPDSSEERQYLNVDRP

RELIPDEADFLLGYATVPGYVSYRSRSQGSWYINKLSQNLDKYAYQHDLLSILIKVNEEV

GRATANIDNGVYKQIPAPMFTLRKRIIFR

>CAG2233488_1 CAG2233488.1 CASP8 [Mytilus edulis]DED

MVDRGTVVDVDNELEAFDVQSIKFLVKNFIHHVQLKKCFSLLDVFTALEVVKHITENNWK

EFLSECLFMIGKRNIIHILGLNSSEIEERVQRKEGFLIPFRTALYNIAEDLDSTEIEKLK

QEAINMVPNIIPALRKVTSMYDFLDVLEKRLLISHHSSDIFLVMLERINRSDLGIFIKDF

SGMCVIINNKTFSKESKLLPRRGTEFDEERLSQTFTKLKFKTCIYRDLSAEEIVEKITEL

AEVDHSKYGACVVCILSHGNETAVFGSYGHSVGINHLTSLLSPRNCQSLTGKPKLLFIQA

CRGIRDQTIQNNNKGQIEEEQHDIGDTNSDIQYIPPPSDPPLGEPRELDEINSDIQYQPP

PSDPPLGEPRELGDINSDIQYLPPPPDTALRGPHDIADINGDIQCLPPQPDPPLGEPHSL

QGEFDFLLGYATTPGYVSYRDCKQGTVYVRTLTEELMAYSDRLDLLSILTIVTNKLSDMN

IDGGAGKVVNLKINLPVIVSNKGIGKVPTELDQYCVCKNPVSCLHDIIMAVDPHKAQIQR

HLSIHFGISFTKGITHTDVLNELKTLVDQNEIKTIQITEKDCIVTVNDQNTKHTLLTRGI

NLKNRFIKMVDVEQQVTNVTIKDAPCEMSDITICGHISKFGDVVHGSLRRGLIKDTNIET

GTRYVQLTNCVPIIPTATKFGRFSVRVFADNDRTECPYCTKTDHTSYRCTNKPTRQNQAA

RKCYRCNSIAHKIKDCPHSEKLCYRCGKEGHIQRDCQTDDQEKYGDYVHDIQEGRDANKD

DSCTEDNNTEEVINELNKTTENVIDLNKTTDVWTNDISKLRSDTEQITANLTTCIDTVKT

KYPNARIGLSSILPRIGKSVSIQALNTAARSVNSFTQKLCVKSQVLKYIDLWDEFAPNKT

PYRALFDLNDPSGVHISMSGTELVENIFEDFINSECEGEYQTPSKKRNRSSSSTPGSSEK

QVTKMPKSY

>CAG2246135_1 CAG2246135.1 CASP8 [Mytilus edulis]

MNAIPIGIAVIINNEIFDGEFEDRTGTHIDAEKLNKLWTDLKFEVDYHYNLTAAQMIQTA

CDIARLDHTKYDSLVFCVLTHGANGVLYGKDENFVHTQQIIDLYLSSRCPTLAGKPKLFF

IQACQGTNTQEGLKLSTDTTNIYPDNYHREIIPNEADFLTRSHSNNDKSEQRCKGEIISR

WQQHLHTDSYIQLNIMKNFEAQVLIADKSNGLKTTLKIWSLHIKKVFIKSNRVIEFLRSN

I

>CAG2192887_1 CAG2192887.1 CASP7 [Mytilus edulis]

MKEEFYNMNHARRGDCVIFNIKHVGTEEVRAGSEQDVQRLTESFSFLGFEVIEKVDLTKK

EIIREIHSIADKNHYDNDCFVCVILSHGREGYVHTTDGEVSLSTLFDPIKNCKSLLGKPK

LFFIQACRGNTIDFGNLVQDGSLRGSETLQMLSSESDVLIAYATTPDYVSWRDKKMGSIY

IEVLCELLRLEGTETKLTELLQKVSNYTAKCFAVSRLCQVPCYTSSLSKSLVFNLKKRVV

MVMYNDGTYSNVSNTNAQINATHSSA

>CAG2199194_1 CAG2199194.1 CASP3 [Mytilus edulis]

MNRESSGLCIIFNMETVGNTSRYGTDIDVERLKDSFGRLGFDVEVKHNPSRRDINNYMND

VKSVNHNRKDSFACVLLSHGRDGHIRSSANEDVPLFEIIEPVKTCESLVGKPKLFFVQAC

RGTRTDSGMLTDAIEDINSDNLRKLSSHSDVLVSYATIEDFVAYRNGHDGSIYIKSLCYL

LDNEGFSTNMTDILNKVNRITVNAFDKSSIEKKTSYYRNGQVVKEKYYYNDRTRRVVNVE

NTHTENHEETSSDSGQEYVI

>CAG2211990_1 CAG2211990.1 unnamed protein product [Mytilus edulis]

MASPQLFKFLSRFSDLYVRLSIHKEEEIYRYNTRLVNNYYCHSHIIFSIIKEEMDSRKDV

TDASPIIGDCDHQTLEKQKLIMTNGTHSVSNSGPGTISKSHLDQERYSISAGIAVVINNI

EFKTELNLPDRKGSDVDASSLFQRFQELGFDSDLLNDASNEDMMEKFNEIKDDKENLKKT

GCLIVALLTHGNEDSVYMTDMSVKIKTVMDLFNATNCPELNLKPKIFIFQACRGGGLGRG

VNMKLIRDDPNKADAAGEYSEYDEKFGETIRIPNEADFLAVYSTSSGYGSFRNTEKGSPF

VRQLSEELLKMKKEDDFYRILTRVNKRVGMGYKPNFPNSGDVTQMPCFISHLTKELDQFD

EAQEPRQFILYFVRSQTAEDMEKFDVLDAVAGNKGETLTKTQITPIANETESNSGQVKGD

KKSLKTTDCLIVVLLTHGTEDSVYMTDGVVKIKNVMEYFNATNCPELHLKPKVFILQACR

GSDLGGGVEMKVIKNRDPNQADAMASYVEYDEEFGETIRIPNEADFLAVYSTSTGYGSFR

NTETGSPFVRHLSDELMKMTKGEDFYKVLTRVNKNVGMRYKPNHSSKGVVTQMPCFISHL

TKDLVFKQKV

>CAG2211993_1 CAG2211993.1 unnamed protein product [Mytilus edulis]

MNEMASQDTLDAKPDLIEETSTKLPVTKGTEPKTNEGTIYQSQLDQERYPIKSGIAVVIN

NTEFDARLGLSDRSGSDVDASSLFQRFMELGFESDLMNDASRDDLVEKLQEIINDKENLK

KTDCLIVALLTHGNEDSVYMTDRSIKMKSIMNCFNAENCPELILKPKIFIFQCSRKPDSR

QGEHITVQDTLKNKVTVPDAVDTVRIPNESDFLEVFSTSIGDRSFHNTESSSPFLRHLIE

ELHNIQEDDDIYKVLTKIRSKNPPSNDCNSEVIVVHSLVLQHDTSKEMADVLDAQRGKPE

DTSTSTIIHGPESNSGQGFVSKAHLDQEKYLISSGIAVVINNTEFHNNLKLGDRTGSDVD

AASLFQRFQELGFDSDLLNDATKVDMEDKFNEIKNDKKSLENAGCLIVALLTHGNEDSVF

MTDESVKIKNVMDYFNAENCPELIMKPKIFIVQACRGTGLGSGVNRTVVRRDPNQADAAA

EYHEYDKTHGEVIRIPNEADFLAVYSTSPGYGSFRNTEQGSPFVRYLSEELLKMKKEDDF

YKVLTRVNKDVGLGYKPNYAHSDIITQMPCFISHLTKDLYLKQKAAEDMENKNMTDGVPY

KKETVITSELNCNTGQGKISKAHLNQERYQISCGIAVVINNVEFEPNLNLGDRFGSEKDA

LTLFQSFKQLGFKSERLNNVTENEMTEKFNKIKEDKKSLAETDCLVIVLMTHGTEESVFM

TDVSIKTKNIMNYFNAENCPELQLKPKIFILQACRGTGLGGGVEMEVAYGELVRIPNEAD

FLAVYSSSSGYKSFRNTDTGSPFVDHLSRELLGMTHDDNFYNVLTRVNKNVGIGYKPSLG

MSGIVTQMPCFSSHLTKELFFNMKEMANRGKKDAHDEDPSSFVTEKERNTGQGSINQRHL

DQKRYSIKSGIAIVINNTEFDARLKLADRKTSDADAASLYQRFLELGFESDILTDVTKDD

LEEKFREITKDEKQLKETGCLIVAVLSHGDEDSVYMTDKPVKFKAVMDYFNAQNCPELIL

QPKIFIFQCSRRSASRQGEDLTFTQMKMNIPCGLETVRIPNEADFLAVYSTSLGDRSFDS

TEDPSPFVHQLWTELQKMKKEDDIYTVLTKVNNKVMRFEPANIDSKFIRQMPYFVSHLTK

DLFLKQIN

>CAG2211991_1 CAG2211991.1 CASP3 [Mytilus edulis]

MDDILQNRRNEKSVPVLMQEIEGSDRDAASLFQSFQELDFETDLLNDASYNDMEEKFEEL

KNDREQLEKIGCLLIAILTHGDEDTVYMTDRSIKIKCYACRGQGIDRGVNVKVVRDPNQA

DGVAEYYDYNKSYGVPMRIPNEADFLAVYSASHGYGSYRITSKGTPFVNHLTEELRKMKK

GEDFYKILTRVNRKVGIEYQPKIPRSGVTCRQQVFIKQETVNTELV

>CAG2211973_1 CAG2211973.1 CASP7 [Mytilus edulis]

MSGKSSLDASLIPDSWTGRPGDPRGQRPRQVPVPIPYPHRVPVAPITAVITKEEFDSKYY

QIKSGLVLVLNNKEFHVQGEDLGDRTGTDIDASTISQRFSELGFEYELETDLPKSATVDW

FERVKRQLIKKPVDCFALVILSHGTENGVYATDQFMRTEDIVETFNATNCPPLRFKPKIV

IIQSCRGTTLSRGINVGLQDPNQADAIADTREYNEWLMGIPTTVTIRLNYEADFLFIYVS

VPGAGAFRNAAEGTPFIRHLSKAIEEMEVNEDFYSVLTSVNKMVGMTYKPNIGEAEYNNV

VQMPCFVSHLTKSLRLKQSG

>CAG2187150_1 CAG2187150.1 unnamed protein product [Mytilus edulis]

MSHGNNEIILGTDGGSKNIIDLITLFNSNNCTSMTGKPKMFIVNACRGDSRDKIFGKTQP

VDVHYTAHTDGPISRKPANSDFLIAYSTFPGYVSYRDDQDGSPFVKCLVEVFTEMAHLVH

VSDMLTEVNNRMAESHNQIPNPTSTLRLKWHLM

>CAG2254645_1 CAG2254645.1 CASP8 [Mytilus edulis]

MDGGGTDWNTALLENLSQNCLITLGTRSLSKAIFDQRHRGFCTPIKDRRSRSVYVFVFFI

LSHGGPRMIYGVDGLPVQISKLTEEFTASSCKTLAGKPKVFFIQACQGKKRTAYSKASPR

CLRVVRSDIVPMPPVQDEPLDDMMHPNPSDFLLGFSTAPGYISYRDSRSGSFYIQSLCRQ

IQENVKGTCSLEFKLTYILMFEVKLNLYWCVVNPIYKTDRT

>CAG2204675_1 CAG2204675.1 CASP8 [Mytilus edulis]

MLTQKKDSSMAASNKTTKGQVKEDGKLKKDKKYIVQGLNGGPQEDETPQKITKGKSGKTE

SIELKLGTTELPLEDAQLVLDQYKKLKTDDRFVNTSGFLTELFRKLQPLEKGQSTRLEVD

MTVRQFQDYSEGFKYKMDAKPRGYALIISNGTFYQIPGDPLSRELPKRPGADTDAENLET

VLTGLGFRVVRKDNLKKWEMEHEIKTIARADHKAFDCFLLILSSHGNIGKLYGSDGQQVL

MSQQLKLFKQEACKDLKGKPKLCFFQACPVGEEDVADTAALPMEGKNKGEKSIPDDKPKF

LINDVDFLVGYSSLPGSVSYRSEQSSSFYLNSLYKNLKDKGEAYDLVTILEDVNRELTAK

KAKYRAGDVNAVSTFVSCLRGPIFFQ

>CAG2204673_1 CAG2204673.1 unnamed protein product [Mytilus edulis]

MKPSQQKNDSTREDVNVKTDQRRKKLANPSANQIEFLMEVYDSVLFGKNKKKETKTAQVV

ELHLGTTVLPLKDARGVLSLIEQLKTDQKYSATLGFMSDLFRKLNTLEDGQFSRIEIDMT

VRQYQDYTDGFQYRLTVKPVGYCLLITNSEFYQVPGCPISKKLPNRPGVEGDAKRLEGLM

STIGYKVLRKDNLKKWEIEREMKAMAEADHKKFDCFVCIISSHGTANKVYGSDGYDLMLG

PQLKRFTVDACPDLDGKPKIFLFQACNIETEDKMQKTLQPLINETDFLYGFSTLPGNLAY

NPKDGKKTGSYYLYSLYKHLSDKYKACDLVAILEDMNRELAEKKSELRPGDLGSVTMFVD

CLRGHLFFQ

>CAG2204672_1 CAG2204672.1 CASP8 [Mytilus edulis]

MASKRNIQKNTTMIKPSPSKNEPQLSKSQQRISKNEPSKSVQTLEKSGIMQNNNYGTGDE

KQKPGQKQNIGQTSKMPNTMSKFQVIPSKNDGRNGKAINNISKSLPTGQLKSIQGLQNVV

PPGQTSKTKNDLSKSQIISSKNKQMDTTTNNLVKSMPAAQLNNLKGRQDLDKKSQQQRLN

RPQPDGQVSKNQTGQKSTTKTESIQAQGQKSGMLPANMQKSQQQQNRPNNSQTSYQKGLP

PLKQQNTVNKGQQNQDKAGNLVSNKMTSQPNTQKSVNQQSRVQQNTQNLAKQSNTFPKVS

QNAENQQNNSQKNQLRQQKLGNQANNLQKTQTNTPKPANKPNKPNGQMKGNPADIVKTSI

ESGQKPGKQQNDLKKGMNPLKPANTGKSIQNYGNTGRSNQNAVDAGKSAQNTKRSSQNVT

IGGKPDKTASNTGRSSQTAGNTGKSQNTGKRGPGDKDKKKKKQLERRNNPVRPQIIEYPA

GVTTIPLPELQVLLDEIEEMRGDRNYSALVEFLSQLLQKVQNLETGNYSRTEIELFVRQF

QDYNEGFHYRMDSKSRGHALILTNYEFYQVPGDPWSYDFKDRPGAEIDADKMEDLLLEMG

YLVTRRDNLKHWEMEWEIKLMTETDHRKFDSFMCIISSHGNVNQVYGCDGLRVILNKQIR

RFRSGKCPSLTGKPKLFFLQSCQAEEEDQTFRSESMVPTDAKPKPGELEKSRFLINDVDF

LVGYSTLPGYLSYRHEQTGSWYLNALYENLKENSKQYDIVTILEDVNFDVSCKKDQYNYL

DSNQVSMFSSTLRGPLFFQ

>CAG2230213_1 CAG2230213.1 CASP3 [Mytilus edulis]

MSETADEKPCYYVPCLIKIESRCDIYETFNVTDCTCKKSAWLCFKFRFKPPHIMNHLIAS

LSRKYEIAEVVVPNQKKKSVALFGGIAVFELQKKTHLRKLLVMTFPNFIQVQVLEFGKNS

IIKRGLYKDIGDFVTDELKKIIHTRFKMSTVEFEKKIECGHTKPELVTGSNELSEEQITE

YLCGTCKTPHTFDGEWSEIESKALSDKYEKKPGQRGLVLIVNFMFHGVTTERKGSDVDCK

NLTNFFEEELKYQVECTTRDMTKLELQDYFQEIARKYLTVNSKDYHSFICVIMSHGNKDG

ILTTNGYITVEEIKKPFKNKTGGKFNGKPKVFLIQACRGKEEQEKVVVSDCFETEIHTGH

VEGYDEFPGNRQRRSSVEMSVFQPGVVGKPSGLSEVFSTDAHSYYSEHVPTDADIIVAYA

TTPGYISIRNIYEGAFFIRAFLSVAKNKYNEEHIEEILRNVRRELATNPEYAPQSGSLAG

KCQMGMLETTSTKKFFL

>CAG2211594_1 CAG2211594.1 unnamed protein product [Mytilus edulis]

MVNISKYNNDIFQFHGETSERKGSDVDCKNLTNFFKEELKYQVECTTRDMTKLELQDYFQ

EIARTYLTEKSKDYHSFICVIMSHGNKGGIRTTDGLITVEEIKKPFKNKTGGKFNGKPKL

FLIQACRGKEEQEKVVVSDCFGTKIHTGHVGRYDEFPGNRQRRSSVETSVIKPGILDQPS

ASSEVFYTDAHSYSSKNVPTDADIIVAYASTPGYISIRNIYEGAFFIRAFLSVAKNKYNE

EHIEEILRNVRRELATNPEYAPQSGSLAGKCQMGQVRDYIYEEIFLVMTVQ

>CAG2232363_1 CAG2232363.1 unnamed protein product [Mytilus edulis]

MYNKKDVSKLELQDYFQEMSRKHLTENSKDYHSLICVIMSHGNENGIETKDEYISLEEIK

KPFKNKHGGQFNGKPKVFLIQTCRGKGNQEKVEVSDLPSNDVQTGIPGEPSSSPEMFSTD

ESSFSKENVLTDADIIVAYATIPGYTSYRSEVNGTSFIRAFLSVAKNKYKDEHIEEILTN

VKRELSSNPKYVPQQLRSAGKMSDGNCTFYIKKEFFCNVRLINLIYEAMAV

>CAG2247236_1 CAG2247236.1 CASP7 [Mytilus edulis]

MVIQLSTTDQMPETFRTKGKQITLMYRLCMHQPRCELNMIQTITRWHTWFICRSDNCSDV

LDLNLGTGKPKIFLIQACRGIEEQEHVEEDDLPLLFKSDTLENLTTDARGYSTEKIPADA

DFIVAYATTPGYISIRDISKGAYFIRAFLTVAKKRYKDEHIEEILIHVKRELASNPDYVP

LTGSLAGTCQMGMVESTSRKKLYL

>CAG2192724_1 CAG2192724.1 CASP7 [Mytilus edulis]

MFQFLHTYDKLAGQRGLLLVINFKFAGQNERTASEEDVRKLHTFFSELDYDIVCQSDMTT

NELKRSLEKIRYEYLTRQSHKYHCFICVIMSHGSELGIATQDGTIKVDDLVRPFKMIMHS

VSLVRIQQRIGKPKIFIIQACRGGRSQVKEQYIEPNIINEADSTVCARLPTDADIIKIFA

TTPGYYSYRKVDSTSWRGAWFIQAFIAVARELPKSHIQEILTAVTHELAINPEYEIDGEK

CQLPMFEAALSKLFYLKGKLRKKIKN

>CAG2215809_1 CAG2215809.1 unnamed protein product [Mytilus edulis]

MAESEKRQYIVGVDFGTTYTGYAYATVKDLKTDSFTYCTAEWKGTNLWSKKAPTSIMMDE

NKQCVAFGYEADSQYFNLLAADEHTDYYYFTQFKQVLYNQILARTTKISDITGKEMKAID

VFSQAIQFVRRHIQRQVKQTNAIDLKDEELYFVLTVPSTKGDAAETFIKDAAKQAGLQDD

NFCIAEEPIVSSMFCLGLRTTRIQSRFDATVKSGMKYMVIDLGGAYTDITVYQIQLNSTI

QEVVPQGGGPWGGTSVDDKFITLLEDIFGETVMTRFKTEQLEDYIDLCRQFESRKRMVKI

NQENDIDISMPGSFISLLKQIRKMSIGEAAKQIHLDDDEIRCSGQKLFLPAKCFKRLFEQ

TLASLITSVGKILSNTKLSDLKTMIMVGGFSECELVMHAVKAEFGGNSMKIITPAEAGLA

VVKGAVLYGHLKLTATTKIPELEHFTTSARPPVSCIKDNFDDRFANLTTSEEYTKKSRSA

RFTPGSKKDVTDLKCFFEKLNYKVLHNEDDMTIDELKQYLEEIRNNRLKKDSHKYHSFYC

VIMSHGNETGIATTDGLIPPEEIVKFFRNNAIAEFIGKPKVFFFDMCRGSETHQKFMQIE

SESGPADMANARPSTTFLPNECDILVAYATTPGYQSFRKIRGNWTGSWFIQAFLAVAEGN

LQMHLEDILKIVRRELSNNPTYELGGKKCMMPNSETKTTKNIYL

>CAG2236484_1 CAG2236484.1 unnamed protein product [Mytilus edulis]DSRBD

MADKLEVLTKAPKSPYKQLEELVSKTLRKIKLQCDISQIENEFVAKLILGDEVYSGCGTN

KTTAKDNAANDALMNTKYHSETKIDKKAVDKLIAICQNMQINHTYEETYQEEDEFVCKLI

ITHENDGIVKDEEHVGRAKSRKDAKQDAASRALIKSTIIPKYSAKDMASADGSYACIGNQ

TGYVLIIIFRTDRDWADKDEENIIELMSGILKFEYSVVADPIKAELVKILENISSNLNEN

YKSYYSFMVFLMAHGSQFGVVTADKGKKKITISVDEIMEFFKNDKIPNFVGKPKMFFIQS

CRGNSIQETFVQPDNDDDETLERISAPTDADIFIAYATTEGHKAFRHKHFGSMFIYECIK

KFKEQYTKTHLEEMMIDVKAALAIGSADKTKRHVQMPCTWSTLTKRLFLVPTES

>CAG2212589_1 CAG2212589.1 CASP7 [Mytilus edulis]

MSHGGEEGIWTTDEQKIKMNDITAQFKNTEFRDFIDKPKIFIVQACRGKPVQELTGHVAD

DEMELEDDFIDSVSVPKDADILIAYATTDGHLSVRHRKDGSWFINALVQVLTETYEKRHF

EEMLVGVRHMLAKWRGRRKDPEDSGKKQILCQMPCSWTTLTKLFYLKASGDC

>CAG2218971_1 CAG2218971.1 CASP7 [Mytilus edulis]

MYLNYSTQLFNYVDKPKIFIVQACRGKPVQELTGHVADDEMELEDDFIDSVSVPKDADIL

IAYATTDGWYLAQILKISFMKGHRETGSWFINALVQVLTETYETRHFEEMLVGVRHMLAR

DPKWRGLRSDPENSEKKQILCQMPCSWTTLTKLFYLKAAGDC

>CAG2221876_1 CAG2221876.1 unnamed protein product [Mytilus edulis]

MPVPKEALDKHAYLTTQLSNYIDSSIDKDQYEQLLNRLKSVLTPKDRNNAKTIMHILESM

EKKSFIAPGKYADLKKILNEFDERIARDVIEPIEEEINDILAKNSKSTKIQPELRGGRDI

AVASSYTDEQPPNKKRKTSQIAIPDSAKYVYANSGQKGLMLILNFSRDREGSTDDVEKLK

SFFEDKFKFEIETLSDLKKEVLQKELTEFRDEYLNSSRHAKKYYCFVCVIMSHGNEKGIM

TDDRQPITMDEITSYFKNKETPHFIGKPKIFLVQACRGSATQPVVTPDDWPEDTMSVPHD

ADILIAYATTPGYKCFRTKYSDIGAWFIHVLLEEIEESYESDHLEDMLISVRHRLAIDEE

WRRSDFCQMPCTWTTLTKRLFFR

>CAG2230422_1 CAG2230422.1 CASP2 [Mytilus edulis]

MPIQSANVKFSHLAKRPRLAENGIIKILSNEDPKSKRIPPMTLVSAQSKEVLSSEVETET

IDGPRSHGEEGYIFGTDGRKIFLDNIMSLFENGHCKDLRGKPKIFIIQACRGEALDSGVV

FVDRHDESQRTTFQLRTKSDILISVPAHTGVQHSGKMYGEEHDGSPCKPYQLPTMSDMLI

CFPTQTGHYAWRNKDRGSWYIEALVQVFMKFARTEDVCTMLNRVNSVISRKLSNCQQIEM

NQMSQMSEYKCTLRMPHFFFFPGIGNE

>CAG2230419_1 CAG2230419.1 CASP3 [Mytilus edulis]

MASKLFCTGVLIALLLQSRLIHSLTGDFTCLGRPKSRPASGRVIEPSSNKRTKAFDAFCD

SLKEDLQEHVVDLYLTPKEEEKSETETTNSCSSRLVNGEPSHAETTSFMRIHSHVTVPST

TTSLSYENANLAISGANYTSAVQPRPHSSPPRLINPPDNHHQFFVQYSSNSRDPNQNFSK

EELKGEARYKFQHYISDPYYNGLNNDKPRDISHESFQKINSLPHHWDSSPRKFAGNTHIL

IRNDSVGPRKEGYIFGTDGRKIFLDSIMSLFDNSHCKNLRGKPKIFIIQACRGGALDSGV

MYGDEHDGAPWYYAWRNRERGSWYIEALVQVFMKKVLWCNKYDKRTDMSTSQLFIKMKEG

RKKELFNVIWILVIVGIPVIVGIPVVGILVIVGIPVIVGIPVIVGVPVIVGVSVIRIPVI

IRIPAIDGIAAIVGVSVYVLPSNKVSECDFMISIWQ

>CAG2209875_1 CAG2209875.1 CASP8 [Mytilus edulis]

MQCTEDTIVTTNACLRDQIRCKPLVDFMSSQTATQIRLSTSELVLECTDTFKRSEGNEEK

TVNFESFPISSTLPDWLIGSNRALLFHLYQRGIMKYELMRLRTFETAVSYRAIRMALGTE

YTNKLITEDTTREMFREIEMTVHNYTRLAIIINNELFNGDMENRTGTQKDGDKLKELWNN

LKFKVMYHINKTAADILQIASDIAQLDHTTYDCVVFCIMTHGGYGVLYGTDCVPLEIQKI

VNFFEVAIVQRQQLSQDSSDIDADTRKDIIADEADILICFSTVSGSVSYRCPQNGTWFIN

SLVKNLRLYSKSYDLQTILVIVNNDVGRNHSQKAAISTNKFLCRSVLYVNLYFLVDDCFI

>CAG2203577_1 CAG2203577.1 CASP2 [Mytilus edulis]CARD

MYITYGVLTEGMMSEITVSTVYITYGVLTEGMMYEITVSTMYITHGVLTEGMMSEITVST

MYITYGVLTEGMMSEITVSTMYITYGVLTEGMMSEITVSTVYITYGVLTEGMMSEITVST

MYITYGVLTESMMSEITSEHTREQQARSLLDILPRRGQTAYDYFHQALVDTHNQELADVL

KPEFASRPELSKAKTLEETSQRYDKTTLQHAASVGSPIIDPQSLFSGSIDIHSTDTVPPF

TPETVEPMKIKMIKCNRQQIISNNPDCYSMDGNPRGKLIIINIMSFDNHGIPSRDKQAHD

LSATSLSLLFKQLKFKIDIKTDINKKELLDLITAEQQSRETYDCFAMVILSHGLGDLIFM

KDGEPLKMQNIMDQFDTENCPKLKGKPKLFFIHACPSDSGEQDNTEMSNIFPLIHEDHHT

LTKADIFLARATKYDPIVSNTESKGFWFIQAFVHIMKYRGHHDHLLDIMTKVNGLAREDD

QLVESMNTLTRSLYFFPD

>CAG2194586_1 CAG2194586.1 CASP7 [Mytilus edulis]

MTAADLTLPSNRTDHFIFPAHVTMDFSDGNSYNFDQKERGIALLINNRNFKTANDRKGDE

LDQQNMKKLFDDLGFKTCSYEDQTADGMRGIATEIAQDSIHVDSDCFVCVLASHGREKQT

KTDTHGPYLEKEQEIMGVDGNEIKTSEFVGIFSDNRCEGLRGKPKFFFIQACRIAGVLKM

DEGYPLRPLESLKLINEKPANKLMDSTEEVPMVTMAPCPEDTLIMFASVSGNLAIRNPRD

GSWMVQELYKCIVKHQKELERMNFLDILTDVLARLSKKETSAKQFLKQFSPGCFIHCLTK

DVFMKEKVREC

>CAG2194587_1 CAG2194587.1 CASP7 [Mytilus edulis]

MDSSDSITYNFGQAERGIALLIHNKKFESKDGTRNGDELDRRHLKTIFKELGFKILSFED

QKADEMLKIATDIAKESVHEKSDCFVCVVSSHGNEEQTKSDTHQSTLEREQEIMGVDGKG

IKTSDLVDIFNDNNCKGLSGKPKFFFIQACRVSGIRKMDEGYPKQNMTPMKGSHKQDKVD

ATTKGNFLHVSNNIIPHHRRAFDHEKDSADAIYEEIDTSDTDDDEDLTYLKAKYQKNTVI

MFAALSGNYAVRTSGKGGWMLNELYKCLAKYAKDGRKLERINFLEILTEVLAGISEKTFN

PPKDSKEYHLNGHFSPACFTHCLTKEVYFKKKSSLRKIKQVISKAKKKVTEMLF

>CAG2207841_1 CAG2207841.1 unnamed protein product [Mytilus edulis]

MDTKIKECAYNFGYAKRGIALLIHNEEYDPNSDFNDRLGDNIDFKHMLEIFKELGFDVRP

YRNLTSTDIIEKTEEVSKMKDLHKESDCFVCVIAAHGEEMVIQKDEGTKEHVIFGTNGKP

VRTSEIVGMFDERNCKELADKPKFFFIQACRTGRVIGNRLDIGHDVSKVDQMVRDEIFKT

TKQNSNNQIIDVECPADTLLMFASLSNNSAVRTPSTGGWLLASLYNQIKEYIKNGKLWSI

EFTQVLNSTLQEMATKFYKPSDETSKWYGAFSPGCFTHTLSADVLFNKKL

>CAG2207843_1 CAG2207843.1 unnamed protein product [Mytilus edulis]

MPMDVSNKDFAYDFNNERRGIALLIHNEVFDKFSGYKDRPGDSGDFIRMQEIFTKLGFEV

DAKRNLTGRKIFDTIQTVSQSKIHEKSDCFVCVIASHGEEMQIETKNNVIIKEHVIVGTD

GQCVKTRDIVEMFNEDNCEGLKDKPKFFFIQACRTSNTTVLGMDSGHILTITERKQPPGG

VDDVEEMHVDSKITDNVDANYISTNKDNLDTRHMSTADTYLSRRQLVHQVTDVECPEDTL

LMFASLSNNFAVRNSSSGGWLLTSLYSQIKEHIETGKIGSTEFTQILNGALKEMTTKNFK

PSDTTSRLYGAFSPGCFTHCLSRDVLFLEKQYR

>CAG2194585_1 CAG2194585.1 CASP7 [Mytilus edulis]

MEGCYNFNQKERGIALLIHNEYFLQSSGYQDRRGDDRDYRRMKKIFKRMGFKVCSFRDQT

AKEMLRIADEVAGMEKEHRESDCFVCVVASHGVEAQKDTRVHSLSHDREHMIVGVDGKEM

STSDFVEKFDGENCEGLQGKPKFFFIQACRIGRYRGSSEGMDKGLEVGITTEENMEKKKK

KKTKGKHMVDGSVQERATSEKQDSEDETVDESDDSDEDEEDIETSDTDDDEDFAELRAKL

QIDADLSKESHPKQRLESEKRIQRKNREEVDAKGIMPDEDSVNVSVGSSFSKIDVPVNAK

TNQQIVSLDKTKPSAHKDELDDKKLVRKKDSKLHTEEVDTKGYKPSTRRVDYVVPMVTVV

PCPEDTLIMFASTSGNFAVRSPAAGSWLLNKLYQQLKVHTKDLEKFQKIDFLNILTEILT

CMSLETYSPGEKSNTKI

>CAG2194584_1 CAG2194584.1 CASP7 [Mytilus edulis]

MEGCYNFNQKERGIALLIHNEYFLQSSGYQDRRGDDRDYRRMKKIFKRMGFKVCSFRDQT

AKEMLRIADEVAGMEKEHRESDCFVCVVASHGVEAQKDTRVHSLSHDREHMIVGVDGKEM

STSDFVEKFDGENCEGLQGKPKFFFIQACRIGRYRGSSEGMDKGLEVGITTEENMEKKKK

KKTKGKHMVDGSVQERATSEKQDSEDETVDESDDSDEDEEDIETSDTDDDEDFAELRAKL

QKVDADLSKESHPKQRLESEKRIQRKNREEVDAKGIMPDEDSVNVSVGSSFSKIDVPVNA

KTNQQIVSLDKTKPSAHKDELDDKKLVRKKDSKLHTEEVDTKGYKPSTRRVDYVVPMVTV

VPCPEDTLIMFASTSGNFAVRSPAAGSWLLNKLYQQLKVHTKDLEKFQKIDFLNILTEIL

TCMSLETYSPGEKSNTKI

>CAG2191994_1 CAG2191994.1 CASP7 [Mytilus edulis]

MNQHLIARRKKIMQLIDRFKKTTRLSQSQTDQAKRQQLPSVKNDLFNDIYDFNHPKRGVA

IFIINSSFHNQSKRPFAGEDIKKMRELFELLDFDVVLLENKTSVDLWKSLQDINLKVEEN

CDCLACVISSHGAEISITSSIDTSTQPRHHVLFTADGIIPTDRILQMFDDDHCPNLQGKP

RMFFIQACRCILNTENDTESEIDEGFAFDLVKEQVGDDMIPVKVVKKKRSFFRRRQKETL

ADTKGRPEEEEINLTNCKSIEPVDFCLIPCYEDYLVMFSSSSEKIAWSDSGKGGWLTFCL

YEVFKHLLDTEYSVSLLHILTDVCGKMAKEMETRTPSIPSYDRSKSAATIYHMLTKDIFL

KPMVHKLVPLPGPVLESSV

>CAG2232222_1 CAG2232222.1 CASP7 [Mytilus edulis]

MPSQTNKTKNKEKPHEKNNVCSHIYDFKFEHSNRRVAILIINSFKISEQPERPFADEDKT

KMKKLFELLDFEVIVLENRKSGDLLEDLQAINRQIENNCDCLACVISSHGAEIPIASSSD

TSPQPRHHVLYTEDNIIPTDTILQLFNDENCPNLQGKPRMFFIQACRSIFDIAPKSEVDE

GYTIDLVQDQVDIDKVPEKTKETLADTHGRHEEEENNLSNCKPIEPVDFCLIPCYEDYLV

MFSSSSGKLGWSNKGKGGWLMYCLFTVFEELLDTNFDNFLQILSRVCGKMATDLETSHDN

KMYNETKSAAVIYHMLTKDLYLRPKSMMESET

>CAG2225971_1 CAG2225971.1 CASP7 [Mytilus edulis]

MTLTVQNQRPRRFAEHDETNMKHLFKSLNFDVIVLKNKETDRLSQDLQEIKSQIDENCDC

LACVISSHGSEYPVMSSSNTSPQLRHHEVSTADVRIPTDTILQFFDDDHCPNLQGKPRMF

FIQACRSTPNSCEENRVDKGFLINSVQEQVDMKDKVIDTQVVEFFDISCYEDYLVMFTCS

SGKLGWSNKSGGWLMQCLHSVFNELLDTNYDDFLQILTRVCWKMARDLETHTDSKVNDRT

KSAAVIYHMLTKEIHLRPKVKTEPKL

>CAG2191993_1 CAG2191993.1 unnamed protein product [Mytilus edulis]

MSTDSSDSDDTEDTQFTEIDSSPDGIKLRQKILTVIKRIRINRKSGTRLRNGKTVKGRQI

EAETNSTEDQEFQPVLESNDYNLSWRKPGLAVFIINSRFEHQPYRPFAETDCHYMVKLFK

HLGFEVRLLKDLKGQDLWRELIAIQTSITTEYDCFVCVISSHGAELPLSRFWDSQRKLRE

HVIYTRDGSIPTDKILNIFNDEGCKNLQGKPRIFFIQACRCVMGADEDLDVDKGVDIEVT

RTRIHEQDYPVFHFKRRSKYNFDNTKDEVKKASKSDENENCDFILDETLLTTSKADTRGR

PEDHHRKVSSGRFHSAEPASFCFIPCHEDFLVMFSSASERIAWSDSGKGGWLMYCIYKVF

KGLLNAEFKTDLLYILVGVCDKMARGLAAATPSDPLRNRTKSAATIYHMLTKDIYLQPKH

VMKQSVCSGQFEGK

>CAG2203760_1 CAG2203760.1 unnamed protein product [Mytilus edulis]

MKGACIVEPERISETKKDSPMTSTSSGGTNSFYGLSSRKVEHTFHTNATQSSSALENSSF

YQFTGSKMGYAIFIINSEFQSEKERQYAVKDRDNMITLFQQLGFEIKLLENKSKKELLQN

LAGKPKMFFIQACRCLKDSPETSHYDYGFDIPSKHAKISCHYDEKSSMDQESVFQIPCSE

DFLVMFTCSPGKSGMTDQRGGWLIHCLYCVFKTLLDSNYKEHFLQILTKVCGKMAQDLKV

NCSSERLFHETKSAAVIYHMLTKDLYLKPKKFGCVMRQTSKKPTYV

>CAG2249235_1 CAG2249235.1 CASP7 [Mytilus edulis]

MEDRKADNIDRKDEFPDLDCKYETYSSMDGLGKNIKKFFGISKSDSSQTYTATVIPAGVT

EDFPHQSTITEVKTPKDSRREAGNRNEGTEESSTTVVYKDTGKAMKASNSQTSQSLQATG

GRGANLQEQSPTLSVNTQYIISPDENNKKPGLALIVTGQKFAIGNGDFRNGAEKDLRNMF

EIFYNLNFEIRWEPFADKTTEDLKNALIKCAEDPTLEQKKCLVIIITSHGGEVKLGRPYR

PENIPHDINVYGHAITTYNGMIPTNEILEIFDVDRCPYMEGKPKLLFIQACRSRNNMKTV

DKGHDIQVAQKQDIKQMRDFFNRSTMDPRHQEDDTNIIYNTVTMKDMLLTPCYQDFMVAF

ASTEGHIAWSEDSGGVLLTAMYNVFKKKIDNKQDIDLLPTMTEVCSKMMDYESNFTDQRY

DRSKAVMCLLHMLGEEIIFKPKY

>XP_052082663_1 XP_052082663.1 caspase-2-like isoform X1 [Mytilus californianus]CARD

MDPKHKELLIRKRVELVENLNVSDGLFTQLIARKIFNQRMVRTIQHNKSPDQQAEELLNI

LPTRGPNAFDAFCDSLKEDLQEHVVDLYLKPKEEEKSEAETTNSCSRLVNGEPSHVESAS

FVRIHSHVTVPSTTTSHTHENVNLAISGANYTSAVQPRPHSSPPRLINPPDNHHQLFVHY

SSNSRDPNQNFSEEELKGEARYKFQHYISDPFYTGLNNVKPRDISHEAFQKINSLPHQWE

SSPRKFVGNSHILIRNDSVDAMTKNGKEYPDNMEQSAKRPRLAEDEIITIPVYEDLKPKR

TPPIVLVSTQSKEIIDPELETGKRLMVPGGEHDDPSLDLTDGPVMVRVESSTRQFYIEHY

KKSYSMIRIPRGKALIVNVNEVEGKPPRRGTNIDRDNLYHLFSQLHFDVTIYNDSDGLTA

KEIAIKLQEFSELPDHKHADACVVCILSHGEEGYIFGTDGRKIFLNSIMSLFDNGHCINL

RGKPKIFIIQACRGGALDSGVMYGDEHDGAPCKPYQLPTMSDMLVCFPTQTGYYAWRNRE

RGSWYIEALVQVFMKFARTEDVCTMLNRVNSVVSRKISRCPQIEMDQMSQMSEYKSTLRM

PHFFFFPGIGSG

>XP_052067946_1 XP_052067946.1 caspase-3-like [Mytilus californianus]

MADKNDTTDAKNMEDTVDTFPFSFGKKSVPVNIHMSREEFEADIYPMNRKNRGYAVIINN

KTFDSGVGLGARNGTDVDASALANRFAELGFDVELEDDVTTEEMLQIMKKYAYMDHSEND

CFACAILSHGEEGIVYGKNGKIEIKKLLEPFKGNLCETLAGKPKVFFIQACRGTEFDEGV

EMNVADAKGEMDVDRQISTHKIPAEADFLVAYSVVPGYYAWRNSIDGSWFVQALSTVLEK

YGSQLDLVRVLTRVNKLVAYKFQSNTSNPYMSRKKQIPCITSMLTKELYFYPKQ

>XP_052068823_1 XP_052068823.1 caspase-3-like isoform X1 [Mytilus californianus]

MVTLCQSTESILSRLSPTSVDKPESPYLLRKASGERRSLRPGAPSEPNAYNYNYPKRGLF

IIINNKNFHPSTGQRCREGTGADAERLEERFQDLGFEVRRYDDATASSMTKIMVEAANYD

HSDCDCFGCAILSHGIEGRIYATDGMLSLDVLTGPFRGDRCSSLVGKPKLFFVQACRTAT

MNHSFDSMSDKSGLSEEYEPALSSIRKVPVEADFLFAYSTVPGHYSWRNNQDGSWFIQAL

CIVIENYGTKMELVHMLTQVNRIVAYEFESCKDEEFTDKMKMMPCIVSMLTRYVYFRPKK

PDIRD

>XP_052079629_1 XP_052079629.1 caspase-7-like [Mytilus californianus]DEDDED

MANLRLRHILVEICSDLGSDDLEKLKFLLRDVTGSAKLEKATMAIDIFSAIENNKHVALH

DGSYLAECFLLIGRKDLVRKLELDPRRVEVGMETDPRLLPLRILLYKISEDLSDKEFEQL

KYHSSKALGIKRGQLQKVKHVFDLFVLLEKNLLISPTSINVLKDMLKALHRGDLLELVVE

YQAKHPYLIDGESNADSTDRNGATQPLESGPPVVDQNPSLPISHPDIANGTPQNPTASPV

LVYEMNAFPRGLCIIINNNKFYSESGFGSRGGSDEDARMLDETFWKLGFIVKVYKDLTAK

DMNAVMHGLSKLNHSKYNAVVVCILSHGGEHVVYGVDGNEITIRYLTSYFRSSRCKSLAD

KPKLFFIQACQGNERQTGINQTDLSIQADGPSDLEEGQSFPETDAEVTSNNSKIIPDESD

FLIGYATVTGYTSFRSRTRGSFYIRKLTDNLNEYANSQDLISILTRVNHDVSTENIPAGN

NCEISKQVPMPQFTLRKQLFFRPENLPQDRTMPQSADVSRIISQSIEAVRTIAQPTDASS

QSSEVTNRSDNLRPITQPSDAN

>ADB80147_1 ADB80147.1 caspase-8 [Mytilus californianus]DEDDED

MALTCLTKDILVDICSDLDSDDMDNLKFLMKDVAGLHKLMPATIALDLFSAIENNKTVAL

LNGRFLAECFELMGRTDLVRRIGLNPTQVEKEMGQTNYSVRPFRILLYKISEDLASDGVE

LLASFSLRVFKLTRQEKESIKSAFDLFVILEKRGIIEVAHTNPLKEMVKSLENRGDLVDL

VDEYQVIVPDNSGHTGSRTIPDYPDGGVSLHSTSHLAQSRRPQNINIGPSVPYYEMTSNP

INIGPSVPYYEMTSNPRGICVIINNKKFKKMRAREGTDVDQVRLEATFKKLGFTVRVFKD

LTVKDMAGVMKDISIMDHSKFNALVVCVLSHGGENFIYRMDDRKEKTGNRQRLVQGDGDR

PIQGNFVYGVDDKPIPIRYLTIYFQSSKCKTLANKPKLFFIQACQGEDMQTGVLLSSDFQ

KDGPSVDESIDLDVAPVSENAEETDAAPHASDNQKVIPDESDFLLGYATVFGYVSYRSRS

QGSFYVKSLTENLDKHADTLDLQALMVKVNHDVSENNIELVKGEQAKKQVPMTQITLRKQ

LFFSPDNLK

>ADB80147Mytilus_californianusDED

PRGICVIINNKKFKAREGTDVDQVRLEATFKKLGFTVRVFKDLTVKDMAGVMKDISIMDH

SKFNALVVCVLSHGGENFVYGVDDKPIPIRYLTIYFQSSKCKTLANKPKLFFIQACQGED

MQTGVLIPDESDFLLGYATVFGYVSYRSRSQGSFYVKSLTENLDKHADTLDLQALMVKVN

HDVSENNIEQKKQVPMTQITLRKQLFF

>XP_052067837_1 XP_052067837.1 caspase-1-like [Mytilus californianus]

MIDPVEIDEAVTNINTAVTGAVTAIIQLSITESKLVGLSEVIARREDWTQLPSALQIDYD

RISQIMQANPTNELIVHGFLKHWFIEQRNKLGTSPSADGLLNAFQQLGFTDAIEEYARLS

HESEYEFEQESENLDARSERSGNQEENFNVSSGEARVSQGNEDTDFRAATPPINQSNSGA

QVGPSLNPLINQSIRQPNQQPNRQDDVIIRQMEAVAIDDSAMPFYKMNARPRGLAIIINN

QHFHKIRNDQSSKEMPARAGTEKDAEKLNYIWQKLDFEVRLFNDVDSTKMNQIMVETAFE

DHSTYDCLVVSILTHGVLGHVYGADGRIVRISQLTSCFSGRRCPSLAGKPKLFFIQACQG

REKQEGHAIETDAGGSMEERQDLNVDRPRELIPDEADFLLGYATVPGYVSYRSRSQGSWY

INKLSQNLDKYAYQHDLLSILIKVNEEVGRATANIDNGAYKQIPAPMFTLRKRIIFR

>ADB80146_1 ADB80146.1 caspase-8-like protein [Mytilus californianus]DEDDED

MIEFESFQYHNNINQTDLTKGLITIDGELESDDIKSLKFLCQDHIAQKTLEKIEDGIQIF

NELERLGKIDTSRGEVDYLLESLCRCHRIDLIRKMGFNQIQVKNRVESGRSNFSPFRLLL

FEIAEEMETADLKTALFTLSSVIPRSKHDKIKSICDVFIQLERSGDLSPVNVEVLYKLLK

MMDRGDLITKVRAYQGAQPTGGNTELGYGPVEEGQIPRTIEIAMEPEYQGEGHIRPAYTV

EQPRPTPATEHSVQFTGNQIPLHLGQFQNSTASVIPSQQEIVDIRGTRSNKNGRLDDNVD

SNMAKRLKVLDRERDRLVPSQQEDPVFLYSQQEQPSTGNYPSVIGIRNQPTAEPSPVNRT

GISSEVPVNHLVPRNDVVTEEQSPGDSGLTAGTVSLSDRSSQDISSGEVQLSITESKLVG

ISEVIARREDWMRLASALMIDYDRISEINRANPTNLLRVHGFLRHWFIEQKNKFGTSPNA

DGLLNAFRQLGFTDAIEEFVRLSLEPESENEQGSENLDARSEGSVYQEENFNVSSGEARV

SQGNEDTDFRAATPPINLSNSGAQVGPSLNPLINQSIRQPNRQDDVIVRQMEAVAIDDSA

MPFYKMNARPRGLAIIMNNQHFHKIRNDPQSKEMPARTGTEKDAEKLDYIWRKLDFEVRS

YNDVDSTRMTQIMVETAFEDHSKYDCLVVSILTHGVLGHVYGADGRIVRISQLTSCFSGR

RCPSLAGKPKLFFIQACQGREKQEGHVIETDGKGSMPVSDSSEERQDLNVDRLRELIPDE

ADFLLGYATVPGYVSYRSRSQGSWYINKLSQNLDKYAYQHDLLSILIKVNEEVGRATANI

DNGAYKQIPAPMFTLRKRIIFR

>ADB80146Mytilus_californianusDED

PRGLAIIMNNQHFHKARTGTEKDAEKLDYIWRKLDFEVRSYNDVDSTRMTQIMVETAFED

HSKYDCLVVSILTHGVLGHVYGADGRIVRISQLTSCFSGRRCPSLAGKPKLFFIQACQGR

EKQEGHVIPDEADFLLGYATVPGYVSYRSRSQGSWYINKLSQNLDKYAYQHDLLSILIKV

NEEVGRATAGAYKQIPAPMFTLRKRIIF

>XP_052081128_1 XP_052081128.1 caspase-8-like [Mytilus californianus]

MVILLNYFFRGFDPLINQRIRQPNRQDDVIVRQMEALAIDDSALPFYKMNAIPRGLAIII

NNEHFHKIKNDQSSKELPTRTGTEKDEEKLDYIWRKLNFEVQLYNDVDSTRMNQIMVETA

CQDHSMYDCLVVSILTHGTLGHVYGADGRIVRISQLTSCVSGRRCQSLAGKPKLFFIQAC

QGTENQEGHAIETDADILRPDERHNLNVDRPRELIPDEADFLLGYATVPGYVSYRSRSQG

SWYINKLTVNLDKYANKHDLLSILIKVNEEVGRATATNDNGHYKQIPAPMITLRKRIIFR

>XP_052081014_1 XP_052081014.1 caspase-3-like isoform X1 [Mytilus californianus]DEDDED

MVDRGTVVDVDNELVSFDVQSIKFLVKNLIHHVQLKQCSSLLDVFTALEIVKHITENNWK

EFLSECLFMIGKRNIIHILGLNSSQIEERIHRKEGFLIPFRTALYKIAEDLDSTEIEKLK

QEAIDMIPNIIPALRKVTSMYDFLDVLEKRMLISHHSSDILLVMLERINRSDLGIFIKEF

SGGFYHMTRPYSGMCVIINNKTFSDASKLPARRGTEFDEERLSATFTKLKFKTCIHRDLS

AEEIVAKITEVSKLDHSKYGACVVCILSHGSETAVFGSYGHSVGINHLTSLLSSRNCQSL

TGKPKLLFIQACRGTRDQTIENNNDEQKGEEQHDTGDINSDIQYLPPPPDPPLGETHNLQ

GEFDFLLGYATTPGYVSYRDCKQGTFYVRTLADELMAYSDRLDLLSILTIVTNKLSDMNI

DGGAGKVHKQCPMPQFSLTKKLYFFPENFI

>XP_052081013_1 XP_052081013.1 caspase-3-like [Mytilus californianus]DEDDED

MACASDEPMRLIDACNDLDSHDIASFKYLCQDDIGLNNLENCIEVTDILDLLSNKESLLH

YVGHRLHLMEKPCLIKGLGLELEAIKTAVEGNNRSTITPYRKVLFNIAQDLGSKDVTSMI

EKAKDCLQNFTPAVRKIEKNGKSGKIFNFFNVLEEKDHLSSSNISFLYQLLFDISRDDLA

KLLKPPKDFNLKDFYPVNQRQRGKCLIINNTKFEDDLLDRKGSNFDAEELSKTFHQLNYQ

IMTCGDLRAEEILDYLNTVSKLDHSDGSSLVVCILSHGSLRTVCGVDGLHIPLRNLTEKF

TASNCKSLAGKPKLFFIQACQGKKEQILPHHRKQKYSQESELPGTRQVESARNDIIVMDT

DENETVADVIPEESDFLLAFSTAPGWSSYRDPDKGSYYIQKLCEQIQKECMRHHLLDILT

DVNNRISKCDIPIKDGQIVKTVPSHFSSLTRKLYFFPENIR

>XP_052081018_1 XP_052081018.1 caspase-3-like isoform X1 [Mytilus californianus]DEDDED

MGCASDETMRLLDVCKDLDSHDIASFKYLCQDDIDLKRLDTSHETTDIFQLLSHKESLIE

YVGHRLCLMDRPCLVKELGLDLEAVKSAVGDVTTSTITQYRKALFNVAQDLSMEDVIAMI

QKSRDCLKNFSPAVRQIEKIGKSGKVFEFLNIIQEKKHITSNNVSFLYQLLYDINRGDLV

HILDSIKGYDLNGFYPFDQCYPGKCIIINNCKFENDLLERKGSHLDADELSKTFRQLNYK

IERHEDLKSHQILEVTDTLSKQDHSRYSSVVVCILSHGGPRSVYGVDGFPVPVRNLTEKF

TGSNCKSLAGKPKLFFVQACQGKKEQIVQRKAAPRPAAKARNDVIAVEIDEDDTVEEVIP

DESDFLLAFSTAPGCSSYRDPNNGSYFIQALCDQIQKDCIRNHLLDILTDVNKRVSKYDI

PIEDDQTVKTVPSYYSSLTHKLHFFPVNTR

>XP_052067829_1 XP_052067829.1 caspase-3-like isoform X1 [Mytilus californianus]DEDDED

MDGGGTDYWKTALLEVSDDLDSRDALSVRYLVQDEIGNPDFEKKRDAIDILNQLLHLKDQ

ELVKHVGECLYLMEKFTLIRKLGLNKEELRNQVENNDTPTISPFRKALFNITKELSTDDV

DKMIRLSKDFLQNLSIRRLERIDESARVFELFNILEQRKEISSEDSSFLTKLFKEMNRKD

LDHHLDTLTNGGFKPVGIYPVNTESPGFCIILNNEKFEDSEKFHDRIGSQADVAFITRLF

ENLRYKIIVKTNLRSREIIDFVHRLKAEDHGVDTSFVFFILSHGGPRMIYGVDGVPVQLR

KLTDEFTASACKSLAGKPKLFFIQACQGEKEQLMPRRTSPRRHRGVRSDVVLVSPDQDET

LDDVVIPNASDFFLGFSTAPGYISYRDPSSGSFYIQSLCRQIQEKCKSHHLLDILTEVNS

QVSKQNILLDGGQRAKTVPTQETSLTAKFYFYPETSKYTV

>XP_052095186_1 XP_052095186.1 caspase-3-like isoform X1 [Mytilus californianus]

MKKGSSMAATNKTNKGQVKEDGKLKKDKKYIIQGLNGGPREDEAPQKITKGKSGKTESIE

LKLGTTELPLEDAQLVLDQYKKLKTDDRFVNTSGFLTELFRKLQPLEKGQSTRLEVDMTV

RQFQDYSEGFKYKMDAKPRGYALIISNGTFYQIPGDPLSRELPKRPGADTDAENLETVLT

GLGFRVVRKDNLKKWEMEHEIKTIARADHKAFDCFLLILSSHGNLGKLYGSDGQQVMMSQ

QLKLFKTEACKDLKGKPKLCFFQACPVGEEDVLVVADTAALAMEGTNKGKKNISDEKPKF

LINDVDFLVGYSSLPGSVSYRSEQSSSFYLNSLYKNLKDKGEAYDLVTILEDVNRELTAK

KAKYRAGDVNAVSTFVSCLRGPIFFQ

>XP_052095188_1 XP_052095188.1 caspase-3-like [Mytilus californianus]

MTVRQYQDYTDGFQYRLTVKPVGYCLLITNSEFYQVPGCPISKKMPNRPGVEGDAKRLEG

LMSTIGYKVIRKDNLKKWEIEREMKAMAEADQKKFDCFVCIISSHGTANKVYGSDGYDLM

LGPQLKRFSVDTCPDLQGKPKIFLFQACNIETEDKMQKTLQPLINETDFLYGFSTLPGHL

AYNPKDGEKTGSYYLYSLYKHLSDKYKACDLVAILEDMNRELAEKKSELRPNDLGSVTMF

VDCLRGHLFFQ

>XP_052059080_1 XP_052059080.1 caspase-3-like [Mytilus californianus]

MGDELDARRGEIVDTSTESTIPYGNESNVGKGFVSKAHLEQDRYPISSGIAVVVNNTEFD

DNLNLGDRKGSDVDAASLFQRFQELGFDSDLLNDATKVDMEEKFNDIKNDKISLKEAGCL

IVALLTHGNENSVYMTDESVKIKNVMDYFNAENCPELIMKPKIFIFQACRGQGLGRGVNR

RVIRDQNQADAGAIYTDYDKIHGEVIRIPNEADFLAVYSTSTGYGSFRNTEQGSPFVRYL

SEELLKMKKEDDFYKVLTRVNKDVGLGYKPNYAHSDIITQMPCFISHLTKDLYLKQKD

>XP_052059022_1 XP_052059022.1 caspase-3-like [Mytilus californianus]

MVQLQSQQQYCCHSDIIFSIIKEEMGSRKDVKDSAPFSDDDETIKKEAPIMTNGTHSTNS

GPGTISKKHLDQERYPVSAGIAVVINNIEFNENLNLPDRKGSDVDASSLFQRFQELGFDS

DLLNDASNEDMMEKFNEIKKDKKNLENTDCLIVALLTHGNEDHVFMTDMSVKIKTIMDLF

NAESCKELILKPKIFIFQACRGSGLGHGVNMKVTRDPNQADAGAVYTEYDKTFGETIRIP

NEADFLAVYSTSAGYGSFRNTEKGSPFVRHMSEELLKMKKDDDFYKILTRVNKRVGMGYK

PNYPNSDNITQMPCFISHLTKDLYLKQMN

>XP_052059064_1 XP_052059064.1 caspase-3-like [Mytilus californianus]

MEKTQDTLDAVSGNKGETLTKTTITNGTESNSGQGKIAEEHLNQEEYTISSGIAVVINNI

TFDTRLKLGDRVGSDLDASSLFQRFMELGFDSDLLNDATKNDMDMKFDEIIKDKESLKTT

DCLIVALLTHGNEDTVYMTDTSIKIKYVMDFFNATKCPELVLKPKIFILQACRGSDLGGG

VERQVIKNKDPNQADASASYVEYDEKFGETIRIPNEADFLAVYSTSTGYGSFRNTQTGSP

FVRHLSDELREMTKGENFYKVLTRVNRKVGMGYKPNHPSSGVVTQMPCFISHLTKDLVFK

QKD

>XP_052059089_1 XP_052059089.1 caspase-3-like [Mytilus californianus]

MNEIANQDTLDAKPVLIEETPTKLPATNGTERKRNEGIIYESHLNQESYTIKSGIAVVIN

NTEFDARLCLSDRSGSDVDASSLFQRFMELGFESDLMNDASKDDLVEKLQEIMKDKKQLS

KTDCLIVALLTHGNEDSVYMTDTCVKMKAVMNYFNAENCPELILKPKIFIFQCSRRSDSR

RGEDIIVEDTLKSKASIPNDADTVRIPNESDFLEVYSTSIGDRSFHNTESSSPFLRHLIE

ELHNIKVDDDIYRVLTKVNSKVMTFEPGHTESKDMRQMPYFVSHLTKDLYLKQVN

>XP_052059131_1 XP_052059131.1 caspase-3-like [Mytilus californianus]

MAERNTKEGIIYKSHLDQESYTIRSGIAVVINNTEFDARLGLSDRPGSDVDASSLFQRFM

ELGFESDLMNDASKEDLVEKLQEIMKDKNQLSKTDCLIIALLTHGNEDSVYMTDTCVKMK

AVMNYFNAENCPELMLKPKIFIFQCSRRPDSRQGEDIIVADTFKNKVSIPNDADTVRIPN

ESDFLEVYSTSIGDRSFLNTESSSPFLRHLIEELHNIKEDDDIYRVLTKVNSKVMTFEPG

HTESKDMRQMPYFVSHLTKDLYLKQIN

>XP_052058990_1 XP_052058990.1 uncharacterized protein LOC127699369 isoform X1 [Mytilus californianus]

MACRDMMDAAPGKEEKVINSFSESNTGQGNISTAHLNQERYQISCGIAVVINNVEFEANL

NLSDRIGSDEDALTLFQSFKQLGFKSERLDNATKDEMEKKFNEIKEDKKSLEETDCLIIV

MMTHGTEESVFMTDMSIKTKNVMDYFNAENCKELQMKPKIFILQACRGTGLGGGLEMEVV

TDKEIYKIEPNQADAGGFYREYDKTHGDPIRIPNEADFLAVYSTSVGYTSFRTTDMVSPF

VHHLSEELLNISKEDNFYKVLTRVNKKVGLEYKPSPKMSGIVTQMPCFSSHLTKELFFNL

KEMANRGKLDANDQDTLTTSSITKETGRNTGQGIINKSHLDKTSYTIRTGRAVVINNTEF

DSKLGVYDRRGGDGDAASLLQRFMELGFESELLSDATKDDLEEKFQEIKGDKEELQKTDC

LIVAVLSHGDEDSVYMTDTPVKIKAVMDYFNSQNCPELIQQPKIFIFQCSRRAALRYGGN

FKVEEEKKENYIPYGLETVRIPNEADFLAVYSTSLGDRSYGSTEGSSPFLCQLMKELQEM

KKDDDIYTVLTKVNNKVMRFEPGSIDSKDIRQMPYFVSHLTKDLYLKQIN

>XP_052094026_1 XP_052094026.1 caspase-3-like [Mytilus californianus]

MLKFKHVTYLFYLSISLSFCHIYFPEEVIDCWSAENQDKLDACLYKGEGNTKAVPNTYET

ESNSGQAKISRSHLKQEKYSISSGIAVVINNETFERDLRLPDRIGSSSDAASLYQSFQEL

GFEHELLSDASNSEMKEKFEEIKNDKEQLEKTGCLIVALLTHGDEDTVYMTDRSIKITTV

MSFFSAVNCPELALKPKIFIFQTSRGEGIGRGVNVTVVRDKPRDPNQADGIAEYYDYNKS

HGVQMRIPNEADFLAVYSTSHGYGSYRSTEKGTPFVNHLTEEFRNMKEGEDFYKILTRVN

RKVGIGYQPNIPRRDVTQMPCFISHLTKKLVLRQ

>XP_052073944_1 XP_052073944.1 caspase-3-like [Mytilus californianus]

MQVLILVTGDKKEQAKRRLSFKAVDCFVLVILSHGTDNGVYATDKLIRTGDIVEAFNATH

CPPLRFKPKILITQSCRGTHLSGGINVDVFDPNRDDDSCYPSEYPEWIGIPITKTIRFPN

EADFLFVYAAVSGSGAFRNVAESTPFIRHLSKAIEDMEVNEDFYHVLHKVHTTMGVT

>XP_052081364_1 XP_052081364.1 caspase-2-like [Mytilus californianus]CARD

MDKIHRKALTQCRSKIVSDLSPDNEFWDKIAEKEIFTPLMLEYIQAKYPRPEKVRQLLDD

LLRRGPDAYWKFLYCLDNSGHRHLADVIRQKEQLILNPNYNPIIQEARFTGNIRGTSTSF

QPTAPNNQGDIQLPHSVSAQTINERNTAGQFPHSQSNPNITSSGTPVTSPSDESASSVTT

SSSSTTSYPSTQLLSNTSVSGQNSSNNDISMENEHDGLTSQLPGTSSQYDNRQQTSSLES

TPETSSHQATERTKKYKMESTPRGFVLIINNIHFRNHKHRPGAEQDRDKLKAMFEDFGFE

VDVKEDQTAMQVEELLSQFSLLQDLYRVDSLIVVLMSHGNDEFILGSDCQRLNVINLIKK

FNADYCPAMMGKPKMFIVNACRGDDVDCNPILGSTQPCYDKTETDSGPTSLDRKQPANSD

LLIAYSTFPGYVSFRDEEQGSWFIQCLEEVFRESASDEHVMDMLTQVNDKVASKSESMMG

YNQTPAPSTTLRYKWYLNPQTV

>XP_052081366_1 XP_052081366.1 caspase-2-like [Mytilus californianus]CARD

MDEIHKKALNQCRVSLVKNIIFESEFCDLIKQKEIFTDGMLEEIQANTSRRKRVGQLLDD

LRRRGPDAYNKFLYCLENSGHRNLADAVKEKEKQIRNENQQPVLRETIHGAQHLQHSISA

QPLLSPDPLSNMNSSQSAPNLASSGSIVASPSEDLQDITSRKEVQGNSNFQRETGYNEDI

RMDVADHINPGFSNPLPGAESSRSSSSGGLGSYNITTDPRGIVLIINNINFVSNNENRHG

AEIDQEKLMEMFRDFGFEIRVHIDKTAQEIEEILKYFSKDEQLLEVDSLIVVLMSHGNNE

IILGTDGECKNIIDLITLFNSKNCTSMTGKPKMFIVNACRGDGRDKIFGKTQPVDVHDTV

HTDGRGLSKQPANCDLLIAYSTFPGYVSFRDDKNGSPFVKCLVEVFREMAHLVHVLDMLT

EVNNRMADSHNQIPNPTSTLRLKWHLM

>XP_052068814_1 XP_052068814.1 caspase-3-like [Mytilus californianus]

MGKHLSKSSTKQKDEAADSSSQENDDHGFYPQYPNVYHSHVFGTQRYSVNQQTLGRYTSV

GNLSTTPTTSDYVSEIGRNPSSRASVCVNTERPPNKQNKDFETDARPSMRRSQSCNQIPD

FYSMNEEYYSMNHPRRGDCVIFNIKHVRNDDIRSGTEQDGQKLKKTFSSLGFEVIEKVDV

TKKEFKKEIKTFADRNHDSDDCFVCVILSHGREGYVNTTDGELSLTAIMDPIKKCKSLLG

KPKLFFIQACRGNKIDFGNLAQDGSQREGVTVQKLSSESDVLIAYSTTSDYVAWRNTEMG

SIYIEILCEILRLEGAETKLTELLQKVNNCTSKYFDDHQCCQVPCFTSSLRKSLLFSRRK

QVVMIMYNDGTYENVPNPIASINDTIL

>XP_052061917_1 XP_052061917.1 cell death protein 3-like [Mytilus californianus]

MGMYISQRSDDRGHVKQELSQDVYGYPTPKKNISRSDSNITSDSDVYIRPCDIREDVDRK

HTTDKFQVNKDRWLLNPDGYGYMMPQATTQRSDSNVTSKSGVYNIPYDKKEEIYRETPTS

TFQFNEEMFTSNPGSLSVNGHLQANKHAAKIIYKDPQLRRHNSYHYMDIRQARQGPLIHH

NSQRSTGKQEMVMTKQKMSDMQTVKSEGFDNSKMHRSVSMNSFIDTSRGIRTKHPLQADH

KQSKEGHSNSSVTTFRMSQSFRQETQNRKYSAPSQQHRSFKRENSQRREYIDQFERDAGL

ETGRTEMWNIENLYVPTEELYVMDGDRSGLCVIFNMETVGTTSRFGTEIDVNGLKKSFGN

LGFDLEVIQNPTRRAIKDYMNSVKSLSHHRKDSFVCVILSHGRDGHIRSSDNEDVPLLEI

IEPVKSCESLIGKPKLFFVQACRGSKTDNGVWTDAAEDISDDNLRKLSLHSNLLVSYATI

EDFVAYRNGKDGSIYIKILCYLLDKEGFTTKLTDILKKVNRMTLDAFGDSSVCQVPSFTS

TLRKSVLFKQKQEKKARSTFK

>XP_052102868_1 XP_052102868.1 caspase-7-like [Mytilus californianus]CARD

MNDEDRRKLQTSRSFLVKNIENIEDINEELYTREVFTDGMKADVEVQDTKEKKVRKFLDI

LPRRGPDAFEIFLDVLNQTKNKHVYDRLITGSSGATYQRSYIAQPINLPTIGGDNQLPET

WPDERSILEASQCKIKPHDPSLMQHTVSGRAYQMKRNPRGRAVIINNKIFHGPVQKIDGK

NRVMLAMREGTDEDVQGLHKLFTDLHFHVKIHKDKNAQKIVDILQEEANDVAFHRNADCF

ILIILTHGKQNAIYGSDGKTVEIRDIKDMFNARNFHCMENKPKLFFIQACRGEEYDGGRN

KTVPDASSTDCAMSGNVVNLALERTRLDFERISVEKANDTDATVEKSIATDSHRLVAMAT

TMDMVSFRNRILGSWFMQAVIYVFRKCSHNEDILSMLSDVNHLVSLGRAKDRENRDAVAV

SEFESTLSKRFYFFPHQCETVPIQASTS

>XP_052095498_1 XP_052095498.1 caspase-3-like [Mytilus californianus]CARD

MAVDPRKALIKNRVAIVRDLDNGDDVADYLYTVKLFTEDMRDTVLQEPEKEKKNRLILDT

LGRRGQKAAKGLHDAFLETENNELAKLLAPYIRVIESEENRFSPKEWPPINEEQNKMQKD

SVRRITDLKSPSYIHEYTDKQVYEIRNKTRGKVFIINNVKFQVNKLSHRHGSEVDAENIK

SLFTELHFDVVQKDNLTGQEMLKFLKEERDKVNWNTMECLILVIMSHGETTGIFGTDGQI

VELKKITEMFESSQCSGLNEKPRLVFVQACRGENTESDHTGTGIQVSDKDGVSEGIQAIS

LETEQTDAVAQAHNEDSTKRTVHSAADFLIAYATPEGTPAYRNIHVGSWFLNAIVWTFKY

HAHQEEIQHMLLKVNELVSQGRTMAGNLTVSEVKSNLRKKFYFFPGVYDDPPKFIDS

>XP_052073549_1 XP_052073549.1 caspase-3-like [Mytilus californianus]CARD

MDDMEWKKAFRVNRVAIVTGIANPNAVADCLFSSGIFTDVMRDAVQQEPETDCKSRLILD

ILDKRGSRSVKCLYTAFLQTSNGDLADLLEPYAKNIESTENLNDPKAWPPAIDEQAAMTK

HRVFKIKDKESPLLHDYSKIDVYKMHGEKRGKVFMINNTFSHTKHPETQKREGTSVDVQN

LTELFQQLHFEVVLKTDLNAEEMVKCLEKERDSITDWNKIECVVLILMSHGKGEYIFGDD

NIPVKLTDITAVFDSYHCKGLDEKPRLVFVQACREVETPGEQLSKVCLKMEKQKLQSHTP

GGQDQVDAEPSKDEQISQVKHPAADFLIVYATPAGTLSSSSIDTGSWFLNAVVWTFKYHA

KHEELHHLLLRVNRLVAKGKDPDVLVVGKKLTVSEVKSNLRKKFYFFPGVYGDSPHLFKD

>XP_052073552_1 XP_052073552.1 caspase-3-like isoform X1 [Mytilus californianus]CARD

MDWKKAFAKNRVEIVRNLANPNDVAGCLFSDRVFTEEMRDGVQQELETEQKNRMILDTLH

RRGQRSVKSLYNAFQETGNDHLADLLLPYAKIIEQKENFNDPKEWPPEKSEQTAMEEHGV

FKIKDLKSPLLHDYNKEDVYRLCWEKRGKVFIMNNKFSNSTWERHGSDVDVKNLTDLFQE

LHFEVVTKTDTPAKEMFEFLIKERNKIENWKDIECIVLILMSHGTSGYIYGSDRIPIKLS

DLTYVFDSEHCIGLDDKPRLVFVQACRGVTDQGKIAELCVKVENQKIQPKSGDELDSETF

HDEKPTQVKHPSADFFIVYATPEGTSAYRHIESGSWFLNAVVWTFKYHAKREELQHLLIR

VNRLVAKGKGSALYEQKLTVSEVKSNLRKKFYFFPGVYGESPQFFNE

>XP_052080984_1 XP_052080984.1 caspase-3-like [Mytilus californianus]CARD

MSCDWKTAYKKNRVKIVGGLANPDAVAGLLFQSEIFTDEMHEIVKQIDENEQKTRKILDL

LNYRGKNGLAALHQAFLDTSNKELAKGLAPYVLAVEEGDNKTEPKEWPPSVKQVEVMLND

PVIKIKDTKSPLFIHEYGQEYVYTIQSELRGKVFIINNVNFKGPMQDRPASIKDAENLTK

LFEELHFTVLRKDDLTSQDMKSFLLEEAKQLDKEKCAECAILIIMSHGSGTKIYGVDCWP

VELKSLTDCFSSENCESLHGKPRLVFVQACRSYDKSDVTEKFQKMTTEPRDEMDAEPSKL

EVKENRSPKCTFPDHPSADFLIAYATPEGTKAWLHNDVGSWFMNAIVWTFKYHAHEEELH

HLLIRVNRLVGKGSTLKKEMTVAEVKSNLRKKFFFFPGIHSKIPTLF

>XP_052080923_1 XP_052080923.1 caspase-2-like isoform X1 [Mytilus californianus]CARD

MTAENFDWKNAFRKNRVKIVKGLANPDEVGGHLYQMKIFNDEMYDTVKQIEETEKKTRKI

LDLLNYRGKKGVAALHEAFLDSNNEDLAKDLAPYVMEVEKRDNKTEPLKWPPSKLENEDM

LKDQVIKLKDPKSPWCIHDYGKDHAYTIQNECRGKVFIINNVNFNGPLQNRVPSNIDAEN

LTKLFIELHFTVLQENDLTAQGMQSFLRSEAKQLEKEENCAECVILIIMSHGVGTKVYGV

DCLPVEIKELTDCFSTANCESLHGKPRLVFIQACRSVETASINKTVHLVKPGGQVEIECI

VKSCKSIQNVYWEWDKNKVKKEDFYRITEWNSSNKYCLQQMKNVRSLFVKNVSENDKGKY

RCIVKHDDGSFSVSNFTDLQLDIKDDSKNELPEEFEKLTTKGRDEMDSVPSVQETQDEIL

PKCTFLDHPYADFLIAYATPEGTSAWLNNEVGSWFMNAIVWTFKYHAHEEELHHLLIRVN

RLVGKGCTWSNDMTVAEVKSNLRKKLYFFPGIYGNPPELFKEK

>XP_052061832_1 XP_052061832.1 caspase-7-like [Mytilus californianus]

MHSYKPKSDAKRRQPRSPKTLNATPITLLDGRSSPRGTGNHLSRRLSPLHISSSRNLPRP

PDIIIHSPNNERTEISKERSAAKSGRRKRINSSSRRRKRSIDTNRSPRLDNDFNLLDDSA

SLRRLSVSDNESNYSSESESRFSSVNSLLSGGRAFADQLTKCISVLDRRGQYYFMNHNRR

GYFAIFNFQMCGDKIRLGSENDTKRLQTCFTKLGFQTEILQNCTTAEMDRELKRFMALDQ

TDCDCFGMAILSHGENGRISTRDGSITLENVIRKVKKCESLLGKPKLIFVQSCRLDNNLT

TDAAIVQSIPKLAIESDLLVSYATTKGNVAYRSPESGSPYIQILTEILEAKGEKESLFDL

LTEVNAKLQEMIQSLGASCEQVSVFETTLKKKLQFDKIVAVYTKYSDGRIEYENKLEENI

QDQTSVKSTQDLRN

>XP_052076503_1 XP_052076503.1 caspase-3-like [Mytilus californianus]

MPKVDLIPFEDCDLLANADRLGPRIIKAGNRTLFSSKEVKSIKEVNVLFPEDKGLGASFT

HEDRIRPTNPTPRRNRVCNLQITNFTHDKKKLTNSEINFHGTSILDRDKETFRSEESQWL

NKFDKYEKKGLLLIINFMFSGEEERTGSDEDVGNLCTFFTKLKYDIDCRVDMTTEELRTC

LEDVRHNYLSRNSSKYHCFICVIMSHGSKKGIETTDGFIEIDEIVDYFKNDKMKKFVGKP

KMFIFQACRGEDIKQIQNNQASHRMVQHDSGSRLVKLPKEADILKIYATTPGNLCYRTTG

SKSWNGSWLIHAFLDMAKKMHSKHHIDEIMTAVKHQLSTNPEYVKDGETCQMALSENSLT

KLFYL

>XP_052096691_1 XP_052096691.1 uncharacterized protein LOC127731865 [Mytilus californianus]ROC

MEYLQSVMSEESSGVTAGQGMEGDMVPTEIKLMADKRSIDLYLKLLESGSEKKRDIRLVI

VGKKGAGKTSLIKRLFSEENTDVTSTNGIEIHTIKCKAMSDDGIWNKLDGTNEETEIHAR

LLKQYKETIEASVEEGAHEAVKENTLTTILYKRIEESKESETVTQQPKIRKHHVTTESQV

EPPVTSQQRNQVLKQAKMDIEAILKSNVDLHDKEEYATLLLWDFAGDEEFYHTHQTFLSQ

DAIYLVVTKLTEADDIQAQDLFQLWMDSIHCYSRLEEDKSSSADSKTSDDLDPPVVIVGT

WKDAVTSDVEEIDDACRENIFKFTENMSEDQLRHIRQEYFISNTEDDPSVFQKIREDILN

LARKMKKWNIDYPLKFIQLETRLQEKKKELPIISFQEIQNIATDTPKPLNDEELRLFLEF

HHEIRALVYFKDLPSYIILDTQWLSDAFRYIVTAKKFRAISIKNRKKWDELYSRGKLHSE

VLEDIFKKRDNMFFKHKDHILNVMERFDIIIRPNMSETADEKPCYYVPCLIKKELHCDIY

ATFNVTDDTCKKSTWLCFKFRFKPPHLMNHLIASLSRKYEIAEVIVPNKIKKSVALFGGI

AVFELQKKTYLRKLLVMTFPNFIQVQVLEFGKDSFIKRGLYKDIGDFVTEELEKIIHTRF

KMSNVEFEKKFECGHTKPELVTGSNEFSEEQITEYLCGTCKTPHTFDGEWSEIESKALSD

NYEKKPGQRGLALIVNFMFHGGTNERKGSDVDCKNLTDFFKEELKYQVECTTRDMTKLEL

QDYFQEIARTYLAVNSKDYHSFICVIMSHGNEIGILTTDELITVEEIKKPFKNKPGGKFN

GKPKVFLIQACRGIEKQEKVVVSDCFETEIHTGHVGRYDEFPGRSSVETSVIKQGILDKP

SGSSEVFSTDAHSYSSENVPTDADIIVAYATTPGYISIRNIYEGAYFIRAFLSVAKNKYK

EEHIEEILRNVRGELATNQEYVPPSGSLEGKCQMGMLETTSTKKFFL

>XP_052084453_1 XP_052084453.1 caspase-14-like isoform X1 [Mytilus californianus]

MDAKEEAKDLHAKLTSDLNEQFGLFESDRYEQFVNRLKAKLSPKASKEAKNIMDILEALE

RNGTIGPGNYKYLKSVIRDYDVRVVADLIEPTEREIQKILSRGQGTSASGSTCYNNEGQY

SVDGKLEKKRKLEEEIPDDNRYVKERGQKGYMLILNVISKKRSGSENDIIALESFFRGQL

GFTVNVKEDVTKDELLQTLTETKRTLTGSEAKKYYCFICVVMSHGEEDGIKTIDGQKIKM

DDITAKFKNTEFKDFIDKPKIFIVQTCRGKPVQPLAEHVADDEMELEDDFIGSVSVPKDA

DILLAYATTDGHLSFRHKETGSWFIDALVKVLKEKYENQHFEEMLVSVRHMLARDPKWRG

LRSDPKHSEQLQILCQMPCSWTTLTKLFYLKAAWDC

>XP_052093697_1 XP_052093697.1 caspase-3-like [Mytilus californianus]

MPAIPKEAFDKHAYLTTELSNYIGSSIDNDQYQRLLNRLKSVLTPKDRQNANTIMDILES

MEKKSYIAPGKYADLKKILNEFDERIVRDVIEPIEEEISAILEKNSTNTALPPGQSDSSN

KEAVASSSTDKQPPKKKRKISEIAIPDSAKYVYENSGQKGLMLILNFSRDREGSTDDVEK

LTTFFGGKFDFEIENQPDLKKEDLQKELTGFREEYLNSPRHSKKYYCFVCVIMSHGNEKG

IMTDDKQAISIDEITSYFKNKETPHFIGKPKIFIVQACRGSSIQPVITPDDWPEDKMSVP

HDADILIAYATTPGFKAFRTKYSDEGAWFIHVLLQEIEESYESDHLEDMLISVRYKLAID

EEWRRHEMCQMPCTWTTLTKRLFFR

>XP_052092435_1 XP_052092435.1 caspase-3-like [Mytilus californianus]DSRBDP20P10

MPEEKLLATSQCINHTYEETYKEEDEFVCRLIITHENNGIVKDEEHVGRAKSKKDAKHDA

ASRALRKSNILLNSSAKDLACADGSYACIGNQTGYVLIINFRKDRDWADKDEENIIELMS

GILKFEYSVVADPTKAELVNILENISSNLNGNYKSYYCFMVFLMAHGSQFGVITADKGKK

KITISVDEIMELFKNDKIPNFVGKPKMFFIQSCRGKSIQDTFVQPDDDDDDTVERISAPT

DADILIAYATTEGYKAFRHKNFGSIFIYECIKKFKEQYSKTHLEEMMIDVKAALALGSVD

KTKKHVQMPCTWSTLTKRFFFVPTES

>XP_052097427_1 XP_052097427.1 caspase-3-like isoform X1 [Mytilus californianus]CARD

MDERSRTILRKGRTILIKGLCDIPGICDKLFAYGVLSEGMMSEITCEHTREQQARALLDI

LPRRGQSAYDFFHQALVETHQQELADVLKPEFASRPELSKAKTVEEAPQRYDRTTLHHAA

SVGSPITDPQSLFLVGSIDIHSTDTVPPFTPETVEPMKIKMQKCNRQLIQPDCYSMDSKP

RGKLIIINIMSFDNHGIPTRDKQAHDLSASSLTLLFQQLRFKVVLKTNINTMELLSLIKA

EQTSAETYDCFAMVILSHGLGDLIFMKDGEPLKMQNIIDQFDTENCPKLKGKPKLFFIHA

CPSDCGEPENTDMSNIFPRIHEDHHTLTKADIFLARATKYDPIVSNTESKGFWFIQAFVH

IMKYHAHHNHLLDIMTKVNGLAQEDDQLVDSMNTLTRPLYFFPD

>XP_052068642_1 XP_052068642.1 uncharacterized protein LOC127707919 [Mytilus californianus]

MEGSYDFNQKERGIALLIHNEYFLKSSGYQNRPGDDRDYKRMKKIFKTMGFKVCSFRDKT

KREILEIADEVAGMEEKHRGSDCFVCVVASHGLESQKDTDVHSRSHEREHVIVGVDGKEM

STSDFVEKFDGENCEGLQGKPKFFFIQACRIGRYRGGDEGMDKGLEIGITKEENMKKKKK

TKKKGKHMVDGSVQEPDTSEKQDSKDETFDESDDSVDDEEDVVTSDTDDDEDFAEFRAKF

QEIDADLSKESHPKQRLELEKRLRRKKIEELDAKGLMPDEDSVKVSVGSGFSKIDVPANA

KTDEQIVSVDKTKPSAYEESSLQLDDKKLVRKNDSKLSTEEVDAKGYKPSERRVEYVVPM

VTVVPCPDDTLIMFASTSGNFAVRSPAAGSWLLNKLYQQLKVYTKDPGSFQNINFLNILT

EVLTCMSLETYSPGEKSHTKYLEGRFSPGCITHCLTEQVFFTEKPSKRKFSFLKLFKR

>XP_052068645_1 XP_052068645.1 caspase-6-like isoform X1 [Mytilus californianus]

MTLHRRIKTMDSSDSIIYNFDKAERGIALLIHNKNFESEELGTRNGDELDRRHLKKIFKD

LGFKILSFEDQKADKMLKIAADIAKEGVHEKSDCFVCVVSSHGNEEQTKSDTHQSSLERE

QEIMGVDGKGIKTSDLVEIFSDDNCKGLRGKPKFFFIQACRISGDRKMDEGYPEQNMMLK

TEIHKQDKTDATVIDDPVNHEKESADEIYEEIDTSDTDDDDELTYLKDKYQKKEKQIHTA

SSLTNPTVLQSEGTHREEIDAKGSKPDEDVPMITMVPCPENTLIMFAALSGNYAVRSSVK

GGWMLNELYKCLAKYAKDGNKLERINFLEILTEVLAGISEHTFNPPEDSHEYFLKGHFSP

GCFTHCLNKEVYFKKKSSLRKIKQVIFDALNKVTEMFF

>XP_052092692_1 XP_052092692.1 caspase-7-like [Mytilus californianus]

MDKMDKECAYDFSHEKRGIALLIHNEEYGPESGFKNRPGDNEDFKHMLEIFKELEFDVHP

YRNLTSTEISKRAKEVSEMSKIHEKSDCFACVIASHGEEIELETEKNIHIKEHVIVGIDG

QVVKTRDIVEMFNADNCKELKDKPKFFFIQACRTSNATVLGMDSGHTVTITEQKQPQGGI

VDGIENMQADSQIIDKVDANCISTNANSLCENNLDTRHLSSAEQQVVHQVTDVECPDDTL

LMFASLSRNYAVRNPTNGSWLLTSLYSQIKEHIDTGKICSTEFMQILNGVLKEMTTKNFK

PSDTTSRLYGAFSPGCYTHCLSRDVLFLEKQYK

>XP_052092690_1 XP_052092690.1 caspase-7-like isoform X1 [Mytilus californianus]

MDVVNKEFTYDFNNEKRGIALLIHNEIFDENCDYNDRPGDNGDFNRMLEIFTKLGFDVKP

YRNLTATQIRKTMEKVSEMSKIHEKSDCFACVIASHGEEMELETENNIHINEHVIVGTDG

QVVKTRDIVEMFNADNCVELKDKPKFFFIQACRTSNATVLGMDSGHTVTITERKQPQGGI

VDGIENMQVDSQIIDKVDANCISTNANSLCENNLDTRHLSSAERTLSRQQLVHQVTDVEC

PDDTLLMFASLSSNYAVRNPTNGGWLLTSLYSQIKEHIDTGKICSTEFTQILNGALKEMT

TKNFKPSEITSRLYGAFSPGCLTHCLSRDVLFLEK

>XP_052092684_1 XP_052092684.1 caspase-3-like [Mytilus californianus]

MEVSNKGFTYDFNNERRGIALLIHNEVFDKYSGYKDRPGDSGDFIRMQEIFTKLGFDVDA

KRNLTGRQIFSTIQKVSESKFHEKSDCFVCVIASHGEEMQIETKNNVIIKEHVIVGTDGQ

CVKTSDIVEMFNEDYCEGLKDKPKLFFIQACRTSNATVLGMDSGHTLTITERKQPQGGIV

DGVEDMQVDSKITDNVDSSYIPTHENQKSNNENSQCEDNLETRHLLSAETYLSRRQLVHQ

VTDVECPEDTLLMFASLSNNFAVRNSSAGGWLLTSLYSQIKEYIETGKICSTEFTQILNG

ALKEMTTKNFKPSDITSRLYGAFSPGCFTHCLSRDVFFLEKQYK

>XP_052106425_1 XP_052106425.1 caspase-1-like [Mytilus californianus]

MYNQYRFKQTTGQIPSQTDKVKSNKEPSVKNYLFNDIYDFNHSNKRVAILIINSSFSEYS

KRPFASEDYTKMKTMFELMDFDVKVLVNKKSSDLRKELQTISRQMEESCDCLACVISSHG

AEIPMESSNDTSAQPRHHVLFTADGIVPTDRILQEFNDDNCPNLQGKPRMFFIQACRSIF

NPDNDPESEIDKGYTIDLLQELADNDIVPDKMNEKVTNAQIDAVDFCDIPCYEDYLVMFS

SSSGKLGWSDKRGGWLMYCLFTVFEKLLDTDFDDFLQILSAVCGKMARDLETSHPKKVYN

RTKSAAVIYHMLTKDLYLRPKSIKETET

>XP_052058715_1 XP_052058715.1 caspase-3-like [Mytilus californianus]

MATSESGSKNDESSPDSVKKKIIHLIDRFKQTTRLTPSQTDKAKSQQLPSVKNNLFNDIY

DFNHPKRGVAIFIINSSFHEQPKRPFAGEDSKRMKELFELLDFDVVLLENKRSVDLWKSL

QDINRKVEENCDCLACVISSHGAEIPITSSSDTSPQPRHHVLFTADGIIPTDRILQLFDD

DHCPNLQGKPRMFFIQACRSVFNTENDTESEIDEGFTINLVQEQVGDGRIPVKIVKKKRT

FLKKGQKGQETLADTHGRSEEEEINLSNCKQIAPVDFCFIPCYEDYLVMFSSSSEKIAWS

DCGKGGWLTFCLYEVFKHLLDTEYSVSLLHILTDVCGKMANEMETRTSSIPAYDKSKSAA

TIYHMLTKDIFLKPMVHQLVPLPGPVLESSV

>XP_052058711_1 XP_052058711.1 caspase-6-like [Mytilus californianus]

MSTDSSDSDDIEDTEFTEIDSSPDGIKLRKKILTVIKRIRINRKSGTRLGNGKIIKERQK

VAEKNSTEDKEYQLCVPAPVLYSSDDIYKYNMSRRKPGLAVFIINSRFEHQPYRPFARTD

CRYMVKLFKHLGFEIHLLEDLKGQELWRELIAIQRSITTEYDCFVCVISSHGAELPLSRL

LDSQRKLREHVIYTMNGTIPTDEILKIFNNEGCKNLQGKPRIFFIQACRCVMGADEEFDV

DKGVHIEVTQTRIHEEDYPVFHIKRRNEYNDDDTKNEVKNEHCDFVLDEKLPTTSKQDTR

GRPKDYDSKVSSTRFHSAEPASFCFIPCHEDFLVMFSSASERTAWSDNGKGGWLLYCLYK

VFKGLLNSEFKTDLLHILVGVCDKMARELAAGTPSDPIRNRTKSAAAIYHMLTKDIYLQP

KHATKQSFCSRQFEEK

>XP_052098329_1 XP_052098329.1 caspase-7-like isoform X1 [Mytilus californianus]

MNISIGKSNSISHISKKKKNMQSESQFDDSSIMKLVRRMKQAFIVKPESISGTNEDYIRS

SIPSGEFNDNYGFSSRQRELNAHLNTTRFSSDTRNSNSYQFTGRKIGYAIFIINCKFDAQ

SERPFADLDCKNMKKLFEQLGFEIKMLENNSSEELLQNLKVIRNSIGRDHDCMTCIISSH

GCEIPTRNNERQHVIFTKDSTVLTEDILHNFGDDECKNLVGKPKMFFIQACRCVGGVSKT

SDIDKGFDIQDKSAKCNAVSSTENMNEESVFQIPCDEDFLVMFTCSSGRLGISDPSKGGW

LMYCLFHVFEKLLDTNYEQDFLQILTRVCGKMARELVAKSESVKEYDKTKSAAVIYHMLT

KDLYLKPK

>VDI15888_1 VDI15888.1 caspase 2 [Mytilus galloprovincialis]CARD

MESKHKELLIRNRVELVENLNVSDGLFTQLIARKIFNQRMVRTIQHNRSPDQQAEELLNL

LPTRGPKAFDAFCDSLKEDLQEHVVDLYLTPKEEEKSETETTNSCSSRLFNGEPSHAETT

SFMRIHSHVTVPSTTTSLSYENANLAISGANYTSAVQPRPHSSPPRLINPPDNHHQFFVQ

YSSNSRDPNQNFSKEELKGEARYKFQHYISDPYYNGLNNDKPRDISHESFQKINSLPHHW

DSSPRKFAGNTHILIRNDSVDAMTKNGKEYPDNMEQSAKRPRLAEDEIITIPVFEDPKPK

RTPPIVLVSTQSKEIIDPELETGKRLMVPGGEHDDPSLDLTDGPVMVRVESSTRQFYIEH

YKKSYSMIRIPRGKALIVNVNEVEGKPPRRGTNIDRDNLYHLFSQLHFDVTIYNDSDGLT

AKEIAIKLQEFSELPDHQNADACVVCILSHGEEGYIFGTDGRKIFLDSIMSLFDNSHCKN

LRGKPKIFIIQACRGGALDSGVMYGDEHDGAPCKPYQLPTMSDMLICFPTQTGYYAWRNR

ERGSWYIEALVQVFMKFARTEDVCTMLNRVNSVVSRKISRCPQIEMDQMSQMSEYKSTLR

MPHFFFFPGIGSG

>VDI74424_1 VDI74424.1 Hypothetical predicted protein [Mytilus galloprovincialis]

MDSRKDVTDASPIIGDGDHQTLEKQKLSMTNGTHSVSNSGPGTISKSHLDQERYSISAGI

AVVINNIEFKTELNLPDRKGSDVDASSLFQRFQELGFDSDLLNDASNDDMMEKFNEIKDD

KENLKKTGCLIVALLTHGNEDSVYMTDMSVKIKTVMDLFNATNCPELNLKPKIFIFQACR

GGGLGRGVNMKLIRDDPNKADAAGEYSEYDEKFGETIRIPNEADFLAVYSTSSGYGSFRN

TEKGSPFVRQLSEELLKMKKEDDFYRILTRVNKRVGMGYKPNFPNSGDVTQMPCFISHLT

KELDQFDEGK

>ADZ24781_1 ADZ24781.1 caspase 3/7-2 [Mytilus galloprovincialis]

MSKSDQRDSVPVTPEEQTSLKQGLMTNGIKQPERLLKKEYVCNSGPGTISKSHLNQDTYP

INAGIAVVINNIEFKRDLGLPDRKGSDVDASSLFQRLQELGFDSDLLNDASIEDMNEKFK

EIKDDKENLKKTGCLIVALLTHGNEDSIYMTDVSVKIKTIMDEFNATNCPELILKPKVFI

FQACRGGGLGRGVNRKVIRDPNQADAGAEYDAYDEIYGDTIRIPNEADFLAVYSTSAGYG

SFRNTEKGSPFVRQLSEELLKMKKDDDFYRILTRVNKRVGMGYKPNFPNSGDVTQMPCFI

SHLTKELYLKQMN

>VDI74419_1 VDI74419.1 caspase 3 [Mytilus galloprovincialis]

MNSAMKLCGDLFAISAVKNSFSLWPRLCILTLNKQGITQLWMLFHDADYSVVVFIRLRSC

GGKSSLLAHLDHEKYLISSGIAVVINNTEFHNNLKLGDRTGSDVDAASLFQRFQELGFDS

DLLNDATKDDMEDKFNEIKNDKKSLENAGCLIVALLTHGNEDSVFMTDESVKIKNVMDYF

NAENCPELIMKPKIFIVQACRGTGLGSGVNRTVVRRDPNQADAAAEYHEYDKTHGEVIRI

PNEADFLAVYSTSPGYGSFRNTEQGSPFVRYLSEELLKMKKEDDFYKVLTRVNKDVGLGY

KPNYAHSDIITQMPCFISHLTKDLYLKQKG

>VDI74422_1 VDI74422.1 Hypothetical predicted protein [Mytilus galloprovincialis]

MANQDEVDASHYKGEENTKSVPITSEIESNSGSANIAKVHLKQANYAISSVTAVIIDNET

FESDLNLPDRKGSNVDSSSLFQSFQELGFETDLLNDATYDDMEEKFKDIKNDSEQLEKTG

CLIVAILTHGEEDTVYLTDKSIKIKKVMSFFNAENCPELILKPKIFIFQACRGKEIGSGV

NVTVVRDKPRDPNQADCIAEHYDYNKSYGVPMRIPNEADFLAVYSSSPGYGSYRSTEKGT

PFVNHLTEELRRMKEGEDFYKILTRVNRKVGIGYQPTIPRSDVTQMPCFISHLTKKLVLT

KINA

>VDI40415_1 VDI40415.1 Hypothetical predicted protein [Mytilus galloprovincialis]

MESNYGQAKISKSHLKQEKYSISSGIAIVINNETFERSLKLPDRIGSSSDAASLFQSFQE

LGFENALLTDASFEEMKEKFEEIKNDKEQLEKTGCLVVALLTHGDEENVYMSDRSIKLTT

VMSFFDTVNCPELVLKPKIFIFQTSRGEGIGRGVNVTVVRDPNQADGVAEYHEYNKSYGV

PMRIPNEADFLAVYSTSHGYRSFRSTDQGTPFIKHLTEELRNMKEGEDFYKILTRVNRKV

GIGCVIQWEKVLRNAAVLALPPSETSSLLALPPPATVKESISIVLIMQE

>OPL33846_1 OPL33846.1 hypothetical protein AM593_03819, partial [Mytilus galloprovincialis]

MMDDVPYKTEKVITNTSNCNTGQGKISTSHLNQERYQISCGIAVVINNTVFETNLNLSDR

VGSDVDAASLFKGFQKLGFNTERLDNASQEDLEKKFDEIKEDKKRLEKTDCLVITLMTHG

NEEEVFLTDGSIKMKTFLSYFNAENCRELHMKPKIFIIQASRGTGLGGGVEMKVVKDKET

HMKDPNEADAGGIYNEYDQVYGKQIRIPNAADFLVVYSTSFGYTAFRTSDVGSPFVHHLS

EELMNISKDENFYRVLTRVNKKVGMEYKPSHKISGIVTQMPCFSSHLTKELFFNLKGMAD

GSKLDVNNKATSTTSSTTKETERNTGQGIINKSHLDQKMYTIRTGRAVVVNNTEFDSGLG

LADRTGSDVDASSLFQRFMELGFEGELLSDATFKDLEEKFQENDKEELQKTDCLIVAVLS

HGDEDSVYMTDTPVKIKAVMDSFNAQNCPELIQQPKIFIFQCSRGPALRDSANFKVGAEK

KENSIPYGLETVRIPNEADFLAVYSTSLVMNYTGLLSKVPGLLLVLSLPTFHSEGQPLPD

SLLSNIFHIHAYNFIFCYESINRTLQKLEKERQNPAFNNMTCLLQDTRLYVWC

>ADZ24783_1 ADZ24783.1 caspase 3/7-4 [Mytilus galloprovincialis]

MNEMASQDTLDAKPDLIEETPTKLPVTKGTEPKTNEGTIYQSQLDQERYPIKSGIAVVIN

NTEFDARLGLSDRSGSDVDASSLFQRFTELGFESDLMNDASRDDLVEKLQEIMQDKEQLS

KTDCLIVALLTHGNEDSVYMTDRSIKMKSIMNCFNAENCPQLILKPKIFIFQCSRKPDSR

QGEHITVQDTLKNKVTIPDAVDTVRIPNESDFLEVFSTSIGDRSLHNTESSSPFLRHLIE

ELHNIQEDDDIYKVLTKVNSKVMTFEPGHTASKDMRQMPYFVSHLTKDLYLKQTN

>VDI74420_1 VDI74420.1 Hypothetical predicted protein [Mytilus galloprovincialis]

MKSIMNCFNAENCPELILKPKIFIFQCSRKPDSRQGEHITVQDTLKNKVTIPDAVDTVRI

PNESDFLEVFSTSIGDRSLYNTESSSPFLRHLIEELHNIQEDDDIYKVLTKVNSKVMTFE

PGHTGSKDMRQMPYFVSHLTKDLYLKQTN

>VDI75808_1 VDI75808.1 Hypothetical predicted protein [Mytilus galloprovincialis]

MAINVSKTKAICIGSKQKLSVTSNENINLALNGQQIIESTCEKVLALDTSLIPETWTGPH

GDPRGQRPRTEPAFTAAITKEEFDSKYYKIKSGHVLVLNNKEFHVPNADLGNRDGSDIDA

STISQRFSELGFEYELETDLSKSATVEWFEKVKRQLITKPVDCFVLVILSHGTANGVYAT

DQLMTTEEIVENFNATNCPPLRFKPKILIIQSCRGTNLSRGLNVPIDPNQADSSADLSEY

YDWLGIQTVRLPNEADFLFSYATIPGRAAFRNSELGTPFIRHLSKAIEEMEENEDFYSVL

TSVNKMVGMIYKPNDGVAAYSDIVQMPCFVSHLTKSLRLKQSG

>VDI01848_1 VDI01848.1 caspase 7 [Mytilus galloprovincialis]

MADKNDTTDAKNMEDTVDTFPFSFGKKSVPVNIHMSREEFEADIYPMNRKNRGFAVIINN

KTFDQGVGLGARNGTDVDASALANRFAELGFDVELEDDVTTEEMMQIMKKYAYMDHSEND

CFACAILSHGEEGIVYGKNGKVEIKKLLEPFKGNLCETLAGKPKIFFIQACRGTEFDEGV

EMNVADAKGEMDVDRQISTHKIPAEADFLVAYSVVPGYYAWRNSIDGSWFVQALSTVLEK

YGNQLDLVRILTRVNKLVAYKFQSNTSNPYMSRKKQIPCITSMLTKELYFFPKQ

>VDI01850_1 VDI01850.1 caspase 7 [Mytilus galloprovincialis]

MEDTVDTFPFSFGKKSVPVNIHMSREEFEADIYPMNRKNRGFAVIINNKTFDQGVGLGAR

NGTDVDASALANRFAELGFDVELEDDVTTEEMMQIMKKYAYMDHSENDCFACAILSHGEE

GIVYGKNGKVEIKKLLEPFKGNLCETLAGKPKIFFIQACRGTEFDEGVEMNVADAKGEMD

VDRQISTHKIPAEADFLVAYSVVPGYYAWRNSIDGSWFVQALSTVLEKYGNQLDLVRILT

RVNKLVAYKFQSNTSNPYMSRKKQIPCITSMLTKELYFFPKQ

>VDI01849_1 VDI01849.1 caspase 7 [Mytilus galloprovincialis]

MEGTQTETDKNDTTDAKNMEDTVDTFPFSFGKKSVPVNIHMSREEFEADIYPMNRKNRGF

AVIINNKTFDQGVGLGARNGTDVDASALANRFAELGFDVELEDDVTTEEMMQIMKKYAYM

DHSENDCFACAILSHGEEGIVYGKNGKVEIKKLLEPFKGNLCETLAGKPKIFFIQACRGT

EFDEGVEMNVADAKGEMDVDRQISTHKIPAEADFLVAYSVVPGYYAWRNSIDGSWFVQAL

STVLEKYGNQLDLVRILTRVNKLVAYKFQSNTSNPYMSRKKQIPCITSMLTKELYFFPKQ

>VDI04987_1 VDI04987.1 caspase 7 [Mytilus galloprovincialis]

MNNCGERRSLRPGAPSEPNAYNYNYPKRGLFIIINNKNFHPSTGQRCREGTGADAERLEE

RFQDLGFEVRRYDDATASSMTKIMVEAANYDHSECDCFGCAILSHGIEGRIYATDGMLSL

DVLTGPFRGDRCSSLVGKPKLFFVQACRTAPMNHSFDSMSDKSGLSEEYEPALSSIRKVP

VEADFLFAYSTVPGHYSWRNNQDGSWFIQALCIVIENYGTKMELVHMLTQVNRIVAYEFE

SCKDEEFTDKMKMMPCIVSMLTRYVYFRPKKPDIRD

>VDH92082_1 VDH92082.1 caspase 8 [Mytilus galloprovincialis]

MANIRLRQILVEICSDLGSDDLEKLKFLLRDVTGSAKLEMATMAIDIFSAIEKNKHVALH

DGSYLAECFLLIGRKDLVRKLELDPRRVEVGMETDPRLLPFRILLYKISEDLSDKEFEQL

KFHSSKTLGIKRGQMQKVKHVFALFVLLEKDLLISPTSINVLKDMLKALHRGDLLELVVE

YQARYPHLIDGESSTDSTDRNGATQPLESGAPVVDQNPSLPSRPQNISNGRQQNPTASPV

LVYEMNAFPRGLCIIINNNKFHSQSGFGSRGGSDEDARMLDETFLKLGFIVTIYKDLTAK

DMFAVMHGLSTLDHSKYNAVVVCILSHGGEHVVYGVDGKDIAIRYLTSFFRSSKCKSLAN

KPKLFFIQACQGNERQTGINQTDLSIQADGPSDLEEGQSFPETDAEDVGRSNNSKIIPDE

SDFLIGYATVTGYTSFRSRTRGSFYIRKLTENLNQYANSQDLISILTRVNHDVSTENIPA

GNNSEISKQVPMPQFTLRKQLFFRPENLPRDGTVPRSAAGLSIISQSIQAVRTIAQPTDA

TSQSGEVTSGSDATSQSSEVQNVISRSDNLRTVTQPSDANM

>VDI67414_1 VDI67414.1 caspase 8 [Mytilus galloprovincialis]

MASTCLTKNILVDICSDLDSDDMANLKFLMKDVAGLHKLMPATIALDLFSAIENNKNVAL

HNGRFLAECFELMGRTDLVRRIGLNPTQVEQQMGQTKYSVRPFRILLYKISEDLASDDVE

LLASFCLRVFELKRQEKETIKSAFDLFVLLEKRGIIEVVHTNPLKEMMKSLENRGDLVDL

VDEYQVIVPDRSGPTCSRTIPDYPGGYDISQSTRHLPQPRPYIGPSVPFYEMTSTPRGIC

VIINNNKFIKMKSREGTDVDQARLEATFKKLGFVIKVYRDLTVKDMAGVMKDISIMDHSK

YNALVVCVLSHGGENFIYRMDDKGGKRGNSRRRQLVHGDGDRPIQGNFVYGVDDKPIPIR

YLTMYFQSSKCKTLANKPKLFFIQACQGEDMQTGVLLSSDLQMDGPSVDESIDMDVTPVF

ENAEETDAAPHASDNQKVIPDESDFLLGYATVFGYVSYRSRSQGSFYVRSLTENLDKHAD

TLDLQALMVKVNHDVSEENIDIVKGEQAKKQVPMTQITLRKLLFFSPDNLK

>AIS73193_1 AIS73193.1 caspase-8 [Mytilus galloprovincialis]

MMALTCLTKDILVDICSDLDSDDMDNLKFLMKDVAGLHKLMPATNALDLFSAIENNTHVA

FHNGRFLAECFELMGRTDLVGRLGLNPTQIEKEMGQHNYSVRPFRILLYKISEDLASDDV

ELLASFSLRVFELTQQEKATIKSALDLFVLLEKRGIIEIDDTNPLKEMVKSLENRGDLVD

LVDKYQDRPIPPGPPVPIYEMTSKPRGICVIINNKKFIKMKAREGTDVDQARLEATFKKL

WFKIAVYRDLTVKNMADVMKDISIMDHSKFNALVVCVLSHGGENFIYRMDDRGGKTGNSQ

RQLVHGDGDRLIQGNFVYGVDDKPIPIRYLTMYFQSSKCKTLANKPKLFFIQACQGEKME

PAVLLSSDLQMDGPSVDDSIDLDVAPVSENTEEKDAAPHASDNQKVIPDESDFLLGYATV

FGYVSYRSRSQGSFYVKSLTENLDNHADTLDLQALMVKVTHDVSEENIGKQAKKQVPMTQ

ITLRRLLLFSPHNLK

>AIS73193Mytilus_galloprovincialisDED

PRGICVIINNKKFIAREGTDVDQARLEATFKKLWFKIAVYRDLTVKNMADVMKDISIMDH

SKFNALVVCVLSHGGENFVYGVDDKPIPIRYLTMYFQSSKCKTLANKPKLFFIQACQGEK

MEPAVLIPDESDFLLGYATVFGYVSYRSRSQGSFYVKSLTENLDNHADTLDLQALMVKVT

HDVSEENIQAKKQVPMTQITLRRLLLF

>VDI58512_1 VDI58512.1 caspase 8 [Mytilus galloprovincialis]

METADLKTALFTLSSVIPRSKHDKIKSICDVFIQLERSGDLSPLNVDVLYKLLKMMDRGD

LITKVRAYQGAQPTGGNTEIGYGPVEEGQVPRTVEIAMEPEYQGEGQIPQANTVEQQRPS

QATERSVQFTGNQLPLHLEEFQNSTAIVNRSINDIRGVNQSQPGTVDIQGQGQNKHSRTD

PYMHEYMRKRLKMEDRENDRSRVVPSLQENPVNLGSQQEQPSPGNHPSLRGVGNQTPTEP

SPFFRTGISSEVPVNHLVHRNDVITEEQSPGDSGLTAGTVSLSDRSSQEISSGDIPLSIT

ESKLVELSEVIARRDDCTLLASALMIDYDQTTQIMQANPTNVLRVHGFLKHWFIEQRRRI

GTSPTADGLLDAFRKLGFTDAIEEYVRLSREPESEFVQGSENLAALSEGSGNQEENFNVS

SGEARVSQGNEDTDFRAATPPINQSNSGAQVGPSLNPLINQSMRQQNRQDEVIVRQMEAV

AIDDSAMPFYKMNARPRGLAIIINNQHFHKIRNDPQSKEMPARTGTEKDAEKLDYIWRKL

DFEVRLFNDVDSTRMTQIMVETGFEDHSKYDCLVVSILTHGVLGHVYGADGRIVRINQLT

SCFSGRRCPSLAGKPKLFFIQACQGREKQEGHAIETDAGGSIPEEDRPRELIPDEADFLL

GYATVPGYVSYRSRSQGSWYINKLSQNLDKYAYQHDLLSILIKVNEEVGRATANIDNGAY

KQIPAPMFTLRKRIIFR

>VDI63351_1 VDI63351.1 caspase 8 [Mytilus galloprovincialis]

MNAIPIGIAVIINNEIFDGEFEDRTGTHIDAEKLNKLWTDLKFEVDYHYNLTAAQMIQTA

CDIARLDHTKYDSLVFCVLTHGANGVLYGKDENFVHTQQIIDLYLSSRCPTLAGKPKLFF

IQACQGTNTQEGLQLSTDTTNIYPDNYHREIIPNEADFLVSFSTVRGCVSYRCPQNGTWY

INSLVENLRKMCKKRDLIAIMTKVNKDVKEKSFQDGSNTYIQIPIYNSTLCKDVFLD

>VDH91938_1 VDH91938.1 caspase 9 [Mytilus galloprovincialis]DED

MVDRGTVVDIDNELEAFDVQSIKFLVKNFIHHVQLKKCFSLLDVFTALEVVKHITQNNWK

EFLSECLFMIGKRNIIHILGLNSSEIEERIQRKEGFLIPFRTALYNIAEDLDSTEIEKLK

QEAINMVPNIIPALRKVTSMYDFLDILEKRLLISHHSSDIFLVMLERINRSDLGIFIRDF

SGGFYHMTRPYSGMCVIINNKTFSKESKLLPRRGTEFDEERLSQTFTKLKFKTCIYRDLS

AEEIVEKITELAEVDHSKYGACVVCILSHGNETAVFGSYGHSVGINHLTSLLSPRNCQSL

TGKPKLLFIQACRGIRDQTIQNNNIGQIDDVSKTSEEEQHDIGDTNSDIQYIPPPSDPPL

GEPRELGDINSDIQYLPPPPDTALRGPHDIADINGDTQCLPPQPDPPLGEPHSLQGEFDF

LLGYATTPGYVSYRDCKQGTVYVRTLTEELMAYSDRLDLLSILTIVTNKLSDMNIDGGAG

KVHKQCPMPQFSLTKKLYFFPENFV

>VDH91689_1 VDH91689.1 caspase 8 [Mytilus galloprovincialis]DED

MIELSNNGLKNFSIRRLERIDESARIFELFNILEQRKDISSNDTTFLTTLFEEIHRKDLF

NHLEPFTKGGLKPIGIYPVNTESPGLCIILNNEKFEDTEKFHERLGSQSDVECITKLFDN

LGYKIIVKSNLRSGDIVDFVHRLKTEDHGVYTSFVFFILSHGGPRMIYGVDGLPVQISKL

TEEFTASSCKTLAGKPKVFFIQACQGKKEQLIQTRASPRCLREVRSDIVPMPPVQDEPLD

DVMHPNPSDFLLGFSTAPGYISYRDSRSGSFYIQSLCRQIQEKCKSHHLLDILTEVNSQV

SKENILLDGGHRALTVPTQETSLTAKFYFYPELVIKHV

>VDI58510_1 VDI58510.1 caspase 8 [Mytilus galloprovincialis]

MTCGDLRGREMLDYVNTVSKLDHSDGSSLVVCILSHGSLRTVCGVDGFPIPLRDITEKFT

ASNCKSLAGKPKLFFIQACQGKKEQILPYHRRQQFSRESELQGTRQVESAKNDIIVMDTD

ENETVADVIPEESDFLLAFSTAPGWSSYRDPDKGSYYIQKLCEQIQKECMRNHLQDILTV

VNNTISKCDIPIEDGQIVKTMPSYFTSLTRKLYFFPENSR

>VDI52580_1 VDI52580.1 caspase 7 [Mytilus galloprovincialis]

MGKNFSKPNTKRNKKGRTEQRLSSIHENDHYGFVSQNQNSFQPGYAGFSYFPVREDKRCN

LNQTTLRRYASTSNISTAPTTSDYVSQRGMNRSSQASNEHYSFDSDRSTQAETNIKLNHF

ETDARPTMRRTQSWLAIPDIYQMKEEFYNMNHARRGDCVIFNIKHVGTEEVRAGSEQDVQ

RLTESFSFLGFEVIEKVDLTKKEIIREIHSIADKNHYDNDCFVCVILSHGREGYVHTTDG

EVSLSTLFDPIKNCKSLLGKPKLFFIQACRGNTIDFGNLVQDGSPRGSETLQMLSSESDV

LIAYATTPDYVSWRDKKMGSIYIEVLCELLRLEGTETKLTELLQKVSNYTAKCFAVSRLC

QVPCYTSSLSKSLVFNLKKRVVMVMYNDGTYSNVSNTNAQINATHSNA

>VDI42160_1 VDI42160.1 blast:Caspase-8 [Mytilus galloprovincialis]CARD

MNDEDRRKLQTSRSFLVKNIENIEDINEELYTREVFTDGMKADVEVQDTKEKKVRKFLDI

LPRRGPEAFDIFLDVLNQTKNKHVYDRLMTGSPGATYQRSYIAQPVNLPTIGGDNQLPET

WPDERSILEASQCKIKPHDPSLMQHSISGRAYQMKRNPRGRAVIINNKIFHGPVQKIDGK

NRVMLAMREGTDEDVQGLHKLFTDLHFHVKIHKDKNAQKIVDILREEASDVGFHRNADCF

ILIILTHGKQNAIYGSDGKTVEIRDIKDMFNARNFPCMENKPKLFFIQACRGEEYDGGRN

KTVPDASSTDCAMSGNVVNLALERTIKDFERISVNNANDTDATVEKSVATDSHRLVAMAT

TMDMVSFRNRILGSWFMQAVIYVFRKCSHNEDILSMLSDVNHLVSLGRAKDRENRDAVAV

SEFESTLSKRFYFFPQQCETVPIQASTS

>VDI06563_1 VDI06563.1 caspase 2 [Mytilus galloprovincialis]CARD

MDWKKAFAKNRVQIVKNLANPNNVADSLFSDRVFTEEMRDEVQQEPETEKKNRMILDTLH

RRGQRSVKSLYNAFQETGNDHLADLLLPYAKIIEQKENFNDPKAWPPEKPEQTTMEEHGV

FKIKDLKSPLLHDYNREDVYRLCGETRGKVFIMNNKFSKTSWEREGSDVDVTNLTDLFQQ

LHFEVVTKTDTPAKEMFEFLIKERNKIENWKAIECIVLIVMSHGTSGYIYGSDRKTIKLS

DLTDVFDSEHCLGLDDKPRLVFVQACRGVTDQDKIAELCVKVENQKIQPKSCDELDSETF

HEEKPTQIKHPSADFFIVYATPEGTSAYRNIESGSWFLNAVVWTFKYHAKREELQHLLIR

VNRLVAKGKGDAGYDQKLTVSEVKSNLRKKFYFFPGVYGDSPQFFKE

>VDI22398_1 VDI22398.1 caspase 2 [Mytilus galloprovincialis]CARD

MAVDPKKALIKNRVAIVRDLDNGDDVADYLYANKLFTEDMRDSVLQEPEKDKKNRLILDT

LGRRGQKAAKGLHDAFLDTENSDLAKLLAPYIRVIESEENRYRPKEWPPNSEEQDKMQKD

SVRRIKDLKSPSYIHEYTDENVYEIRNKTRGKVFIINNVEFQTKLSTRQGSEKDAENMKS

LFTELHFDVVQEDNLTGQDMLKFLKEERNKVDWNAMDCVILVVMSHGHTTGIFGTDGQEI

ELKKITELFESSQCPGLNEKPRLVFVQACRGENKESDHIGTGVQVSGNDSVTEGVHAMSL

IGKDQIDAVAEMPNEDLKEKKTVHSAADFLIAYATPEGTPAYRHIHVGSWFLNAIVWTFK

YHAHQEEIQHMLLKVNELVSQGRTMAGNLTVSEIKNNLRKKFYFFPGVYDDPPKFIDG

>VDI33080_1 VDI33080.1 caspase 7 [Mytilus galloprovincialis]CARD

MDKNHRKALTQCRSKIVSDLSPDNEFWDKIAEQEIFTPLMLEYIQAKNPRPEKVRHLLDD

LLRRGPDAFGKFLYCLDNSGHGHLADVIRHKEQMILNPNNNPILQEARFTGNIRGTSTGF

QTTAPNNHGDSHLLHSVSAQTINAHIPMGQFPHSQSNPNIASSGTLVTFPSDESASSGTP

VTSPSDASSRSTTSYPSTSLSGQNSSNNDISMENEHDGLTSQLPGTSSQYDNRQQTSSLE

STSETGSHQATERTKKYKMESTPRGFVLIINNIHFHNHKHRPGAQQDRDKLKAMFEDFGF

QVDVKEDQTAREIENLLSEFSLLQDLHRVDSLIVVLMSHGNDEVILGTDCRRLNVIRLIK

KFNADYCPAMMGKPKMFIVNACRGDDVDCNPILGSTQPCFDKTETDSGPTSLDRKQPANS

DLLIAYSTFPGYVSFRDEEEGSWFIQCLEEVFRESASEEHVMDMLTQVNDKVASKSESMM

GYNQTPAPSTTLRYKWFLNPLTV

>VDI78201_1 VDI78201.1 caspase 7 [Mytilus galloprovincialis]CARD

MIMDKIHKKALNQCRRSLVRDIILESEFCDLIKEKEIFTDGMFEEILANTSRRKRVIQLL

DDLTRRGPDAYNKFLYCLENSGHSNLADAIKEKEQKIRYPNQQPVHGAQHLQHSVSAQQL

QSPDLLSNMNSSQSAPNPLSNMNTGQSAPNPLSNMNSSQSAPNLASSGSIVASPSEEVQD

ITNRKEVQGNSYFQRETGYNEDVRMDVADHINPGFSNPLPGAESSRSSSSDGLGSYNIRT

DPKGIVLIINNINFVSNNENRHGAEIDQERLTEMLSNFGFDIRFHIDKTAQEIEETLKYF

SKDEQLLEVDSLIVVLMSHGNNEIILGTDGGSKNIIDLITLFNSNNCTSMTGKPKMFIVN

ACRGDGRDKIFGKTQPVDVHYTVHTDGPISRKPANSDFLIAYSTFPGYVSYRDDQDGSPF

VKCLVEVFTEMAHLVHVSDMLTEVNNRMAESHNQIPNPTSTLRLKWHLIDSHIVRTNLTD

DNRPLATLLAVLSIRPSGIPEAGYGVFAERSFPKGTRFGPYGGKITENIDNWSYAFNVYD

ADDSVKYYVDAYYKNNSNWLRYANCARTVHEENLYVLQFNDEVYYVAYECITKGTELLVW

YGPDYGEDLGLKRKPEDYQERFHEQDIRYVIENTITENVHKLYTHCYKFFDQAPFIKPPG

ENGTFDKYCFHDGSYQVCLYSECNVNPCTNQTDCKDCEGCCLHQGKYFKATEKFKHVDTI

NTCVCNEDGHFGCTIYAQFPEYICS

>OPL32628_1 OPL32628.1 caspase 8, partial [Mytilus galloprovincialis]

MAINKTTFVTEDLTCIIQGSTDDVEKLKSFFEDKFKFEIETLSDLKKEVLQKELTEFRDE

YLNSSRHAKKYYCFVCVIMSHGNEKGIKTDDGQPITMDEITSYFKNKETPQFIGKPKIFL

VQACRGSATQPVVTPDDWPEDTMSVPHDADILIAYATTPGYTCFRTKYSDIGAWFIHVLL

EEIEESYESDHLEDMLISVRHKLAIDEEWRQGDFCQMPCTWTTLTKRLFFR

>VDI38397_1 VDI38397.1 caspase 7 [Mytilus galloprovincialis]

MPVPKEALDKHAYLTTQLSNYIDSSIDKDQYEQLLNRLKSVLTPKDRNNAKTIMHILESM

EKKSFIAPGKYADLKKILSEFDERIARDVIEPIEEEINDILAKNSKSTKIQPELRGGGDI

AVASSYTDEQPPNKKRKTSQIAIPDSAKYVYANSGQKGLMLILNFSRDREGSTDDVEKLK

SFFEDKFKFEIETLSDLKKEVLQKELTEFRDEYLNSSRHAKKYYCFVCVIMSHGNEKGIM

TDDRQPITMDEITSYFKNKETPHFIGKPKIFLVQACRGSATQPVVTPDDWPEDTMSVPHD

ADILIAYATTPGYKCFRTKYSDIGAWFIHVLLEEIEESYESDHLEDMLISVRHRLAIDEE

WRRSDFCQMPCTWTTLTKRLFFR

>VDI80363_1 VDI80363.1 Hypothetical predicted protein [Mytilus galloprovincialis]

MIYFIIWLVINIVTAYDFMNLTCPHPGQWKLRARAKNCSSIQNYVCLFDAVQLTFEENCI

GPRDERIGNKLVLNIGNFNRKPCSSTHYQPLTFFSFGNTNCSHLKSPCSEEGQVIADNGT

SYSDRSCRCDYRQGYAFVNDTSVHCSCMPSKEECTCFQKSCRENEVLSPDYKCVKEIEQP

GNYTCSQLLDKNSIESGTADVNPMYDFSNRYYDTDTDYFVEPLEEIVNCLKGETVTLSCK

VNKERTTGIWFKDSEKLNGESPDIRLIAEGKFHKLEIVNVRIEHRGYYTLYIKKIASSSL

LLVIGKNFRVIDRHDKYEKKPGQRGLALIVNFMFHGETSERKGSDVDCKNLTIFFKEELK

YQVECTTRDMTKLQLMKYFQEIARTYLTEKSKDYHSFICVIMSHGNEDGILTTNGLITVE

EIKKPFKNKSGGKFNGKPKLFLIQACRGTEEQEKVVVSDCFGTKIHTGHVGRYEEFPGNR

QRRSSVETSVIRPGILDQPSASSEVFYTDAHSNSSKNVPTDADIIVAYASTPGAFFIRAF

LSVAKSKYNEEHIEEILRYVRNEIATNPEYAPQSGSLAGKCQMAQVETTSTKKFFL

>VDI00435_1 VDI00435.1 caspase 7 [Mytilus galloprovincialis]

MEMYCRRKTEDHEQNMGNNRKDGQKTKHKLRTFIAALHADAIPGNDTNVNFLNTPTDEAL

SELSQHLGIGTLQLGIELGLNLAEIEQSCYKFQRNLQELNKDILLKWKNKSRVKTTRCLM

MALQMVKNGGTTYLDKLVKNSSMFERITYPKQMFDKPMDPHGLLLEDKCDSHIEKIYEKK

AGQTGLAVIVNYSHKRHASEADVERLETFFVNIVKYHVEIVEDKAKSTLMQTFESIRKNE

MTSKAADKYHCFVLVMMCHGEKDTLHIITTKGVVENIKVQEVIDQFKLTDFSGKPKFFIF

QACRHQPGTRTDEADSLESQNLKGKLHQDADLLVSYATTPGYKAYRELYTTEEGSWFIRE

MINVFKERYEETDVINIMMAVRRNVEKCRTSEGGIQMTQDLLTLRKKFFL

>VDH97760_1 VDH97760.1 Hypothetical predicted protein [Mytilus galloprovincialis]

MHSCKPKSDTKRRQSRSPMTLNTTQITLPDGKTTPRGTGNLLNQRLSPLHVSSSSPRNLT

QPPDIIIHSPHNERKEISKEHLAVKSVRRKRTNSRTRRRKRSSDSNRSAHVNNDFNLLDD

SASLRRLSVSDNESNYSFESDSRLSSPNSILNVGRAFTDHSLSKCISVLDRRGQYYFMNH

KRRGYFAIFNFQVCGDKIRQGSEKDTERLDACFTKLGFQTEVLQNCTTAEMDRELKRLMA

IDQTDCDCFGMAILSHGENGRISTRDGSISLENLIRKVKKCESLLGKPKLIFVQSCRLDN

NLTTDASIVQSIPKLTIESDLLVSYATTKGNVAYRSPDLGSPYIQILTEILEAKGDKESL

FDLLTEVNAKLQERVQSLGASCEQVSIFETTLKKKLQFDKIVAVYTKYSDGRIEYENKLE

DNIQEQTHVKSTQDLRN

>VDH97448_1 VDH97448.1 caspase 2 [Mytilus galloprovincialis]CARD

MDERSRTILRKGRTILIKGLCDIPGICDKLFAYGVLTEGMMSEITSEHTREQQARALLDI

LPRRGQTAYDYFHQALVDTHSQDLADVLKPELASRPELSKAKTLEETPQRYDRTTLQHAA

SVGSPIIDPPSLFSVGSIDIHSTVTVPPFTPETVEPMKIKMLKCNRQQIISNNPDCYSMD

GNPRGKLIIINIMSFDNHGIPSRDKQAHDLSATSLSLLFKQLKFKIDIKTDINKKELLDL

ITTEQQSRETYDCFAMVILSHGLGDLIFMKDGEPLKMQNIMDQFDTENCPKLKGKPKLFF

IHACPSDSGEQDNTEMSNIFPLIHEDHHTLTKADIFLARATKYDPIVSNTESKGFWFIQA

FVHIMKYRGHHDHLLDIMTKVNGLAREDDQLVESMNTLTRSLYFFPD

>VDI05140_1 VDI05140.1 Hypothetical predicted protein [Mytilus galloprovincialis]

MNVSDGSSYNFDQKERGIALLINNRNFKTTNDRKGDEIDQQNMIKLFGDLRFKTCSYEDQ

TADGMRGIATKIAQDSIHEDSDCFVCVVASHGREKQTKTDTHGPYLEREQEIMGVDGNEI

RTSDFVGIFSDDKCKGLRGKPKFFFIQACRTAGVLKMDEGCPLRPLETLKLIQETHGNKF

MNSGEEVPMVTMAPCPEDMLIMFASVSGNLAMRNPKDGSWMFQELYNCLEKNRKDLGKRN

FLEILTYVLARLSKKETSAKQFLKQFSPGCFIHCLTKDVFMKEKV

>VDI05141_1 VDI05141.1 Hypothetical predicted protein [Mytilus galloprovincialis]

MDSSDSITYNFGQAERGIALLIHNKNFESKYGTRNGDELDRRHLKKIFKELGFKILSFED

QKADEMLKIATDIAKESVHEKSDCLVCVVSSHGNEEQTKSDTHQSTLEREQEIMGVDGKG

IKTSDLVDIFNDDNCKGLSGKPKFFFIQACRVSGIRKMDEGYPKQNMTPMEGSHKQDKVD

STTEVTDIAFDHEKGSADAIYEEIDTSDTDDDEDLTYLKAKYQKKDTQIHPALSLIKYKV

LRSEGTHTEEVDSKGSEPDEQVPVITLVPCPENTLLMFAALSGNYAVRSSGKGGWMLNEL

YKCLAKYAKDGKKLERINFLEILTKVLAGISEKTFNPPKDSKENHLSGHFSPACFTHCLT

KEVYFKKKSSLRKIKQVISKSKKKEIRAVLPRRVKKSFDYRVFDQLGMQSEAVTTESEVG

DELVHGAAGGSDSMSKVDFEDLKERLCIYMDSEESDTDEETKELGRKLKEVKQKEKVMKN

KKRKVELRQEIEKRMKELEELEELELKGASKPKKEKSGKKVEQSKKNKGKSKETDKEKGK

ETDREVKVKIKIKTKVQLKIKVMNFLV

>VDH98847_1 VDH98847.1 caspase 7 [Mytilus galloprovincialis]

MDTKIKECAYNFGYAKRGIALLIHNEEYDPNSDFNDRLGDNIDFKHMLEIFKKLGFDVRP

YRNLTSTDIIEKAEEVSKMKDLHKESDCFVCVIAAHGEEMEIQTAEGTKEHVIFGTNGKP

VRTSEIVGMFDERNCKELADKPKFFFIQACRTGQVIGNRLDIGHDVSKVDQMVRDEIFKT

TQQNSSNQIIDVECPADTLLMFASLSSNSAVRTPSTGGWLLASLYNQIKEYIKNGKLWSI

EFTQVLNSTLQEMATKFYKPSDETSKWYGAFSPGCFTHTLSADVLFNKKL

>VDI16491_1 VDI16491.1 caspase 7 [Mytilus galloprovincialis]

MDVSNKGFSYDFNNERRGIALLIHNEVFDKFSGYKDRPGDSGDFIRMQEIFTKLGFEVDA

KRNLTGRKIFDTIQTVSQSKSHEKSDCFVCVIASHGEEMQIETKNNVIIKEHVIVGTDGQ

CVKTRDIVEMFNEDYCEGLKDKPKFFFIQACRTSNTTVLGMDSGHILTITERKQPPGGVD

DDEEMHVDSKITDNVDANYISTNKDNLDTRHMSTADTYLSRRQLVHQVTDVECPEDTLLM

FASLSNNFAVRNSSSGGWLLTSLYSQIKEHIETGKIGSTEFTQILNGALKEMTTKNFKPS

DTTSRLYGAFSPGCFTHCLSRDVLFLEKQYR

>VDI73650_1 VDI73650.1 caspase 7 [Mytilus galloprovincialis]

METSESASKNNESTPDSAKKKIMQLIDRFKKTTRLSQSQTDQAKRQQLPSVKNDLFNDIY

DFNHPKRGVAIFIINSSFHNQSKRPFAGEDIKKMRELFELLDFDVVLLENKTSVDLWKSL

QDINLKVEENCDCLACVISSHGAEISITSSIDTSTQPRHHVLFTADGIIPTDRILQMFDD

DHCPKLQGKPRMFFIQACRCILNTENDTESEIDEGFAFDLVKEQVGDDMIPVKVVKKKRS

FFRRRQKETLADTKGRPEEEEINLTNCKSIEPVDFCLIPCYEDYLVMFSSSSEKIAWSDS

GKGGWLTFCLYEVFKHLLDTEYSVSLLHILTDVCGKMAKEMETRTPSIPSYDRSKSAATI

YHMLTKDIFLKPMVHKLVPLPGPVLESSV

>VDI73651_1 VDI73651.1 Hypothetical predicted protein [Mytilus galloprovincialis]

MSTDSSDSDDTEDTQFTEIDSSPDGIKLRQKILTVIKRIKINRKSGTRLRNGKTVKGRQI

EAETNSTEDKEFQPALESNDYNLSWRKPGLAVFIINSRFEHQPYRPFAETDCHYMVKLFK

HLGFEVRLLKDLKGQDLWRELIAIQTSITIEYDCFVCVISSHGAELPLSRFWDSQRKLRE

HVIYTRDGSIPTDKILNIFNDEGCKNLQGKPRIFFIQACRCVMGADEDLDVDKGVDIEVT

RTRIHEQDYPVFHFKRRSKYNFDNTKDEVKKASKSDENENCDFILDETLLTTSKADTRGR

PEDHHRKVSSGRFHSAEPASFCFIPCHEDFLVMFSSASERIAWSDSGKGGWLMYCIYKVF

KGLLNAEFKTDLLYILVGVCDKMARGLAAATPSDPLRNRTKSAATIYHMLTKDIYLQPKH

VMKQSVCSGQFEGK

>ADZ24778Mytilus_galloprovincialisCARD

KKGKFLSVSNVQCTTCRCCVDFDKTCISQLFTHMQYDSDNVRGVSGQDVKVFLEEHLNKE

NTVFDSLVIVFFSGGDKGKIYDKGGNEIDKEEIFTMIKKSTAFKGKPKIVILRTYDFEEK

TTPPDKPNVDDLFVVPKTKKGPWVVGDGMNGSYFIQALIHVFKNMAHEKSFMEMMEEVNT

CIMNAMVPTVAEVMILEHSVQKSLF

>VDI18073_1 VDI18073.1 Hypothetical predicted protein [Mytilus galloprovincialis]

MIVAAWLWGIMLGVAAVSGLSKAVYRQKKLQCGPMYPYVNIPKDSILFFANFTINILLPI

VIMGITYRKIFTILHQSAIFRRNSATSTEDIFSDDKRVIITLILVLGVFIVCWLPYVVYI

FLALMLEDKNDIPSYINPLAYCSGFTNSICNPVIYAFRFKDFRKGYKQILQRSSCMSTRD

LRMIELRESRRPSQNLICVGTNVVIIILSDVCLINLVFLPESIRPLEKHMQGMSLSEKKG

WVLIIQNTDFGHGRALPESADIDTLQRLFTDYNYNVRVRKNVTGTALIEHVKYYADKEKS

KCFVCFISSHGNLEYVECVVKKDEDERTRLVKIENIFDKANTRRLANLPKVFFIDACRSE

QPGRQLRNPMIPSDNFILGMSCLGRKKSMANRSIGDYYFHHIAKVFEESFPLGLQGNGYV

RDIDYLLKKVNRDVAEYAEKKGYSQRPIYSNSMNRHLFLQAK

>XP_033764175_1 XP_033764175.1 caspase-2-like [Pecten maximus]

MRTIYECDKVTEKCYQRNDSLDGMTKRPFKPNQDSHDHPGVKRLRTEETVILPARDGSGG

SICVRDSSYDSRAIYKPLPEIFMVDPCSQRNARVPPIQSIGEFDDPDIDLTDGPVSVNVA

LCQRQFYLSHFKKSYAMRRIPRGQALIINVNEVEGKLPRRGTNIDRDNLHNLLTQLHFCV

TVYNDVDKLTADEIGKKLQQFSLLEDHVQADCCVVCLLSHGEEGYLFGTDGNKLKLDDIF

ALFDNNSCKSLIGKPKIFFIQACRGGALDEGVEWKQADEIDGSHPSMRQLPTMSDMLICY

PTQSGYYAWRNRERGSWYIEALVQILMKYAKNEDMCTMLNRVNQLVSRKISRCPQIEMDQ

MSQMSEYKSTLRKPHLFFFPGIGSV

>XP_033747025_1 XP_033747025.1 caspase-3-like [Pecten maximus]

MGDKGDTKKDMPDANPNLVSDTDARYGHGSGGIFSKGQSRKPVPRTTILTEAELNSPLYK

MDHARRGKAVIINNKEFKGNMGLGDRKGTDVDASVMYRRLQELDFDVTLEHNLSADKMQQ

ILINLSLEDHNDCDCLFVVVLSHGEDGVVFGTEGSVEVKKLFEPFKGNNCPSLAGKPKIF

AIQACRGSQFDDGLGVTVADAEGVMEVDRTVRTQSIPAEADFLVAYSVVPGYYSWRNSTN

GSWFIQALSLALDDEKLKDLDFLRLLTVVNRKVAYHFQSQTSDPRMAMKKQIPCITSMLT

KEIYFNPK

>XP_033734463_1 XP_033734463.1 caspase-3-like isoform X2 [Pecten maximus]

MGRTVTMHTSTDSLTPTSTFLSPHNSVDCPGPPYILRRPSGDRRSVRPGAPSEPNEYNFN

YHKRGLFLIINNKNFHPSTGQVTREGTNVDAERLEERFQDLGFEVRRYNDVASSKITKIM

YDASEYDHSDCDCFGCAILSHGIEGRIYATDGILPLEMLIVPFKGERCPTLVGKPKLFFL

QACRGISNEASLDLTDSVSDRSGYSEEFESCMHPLRKIPVEADFLFSYSTVPGCYSWRNN

QDGSWFIQALCIVLENYGPKMELLHMLTQVNRIVTYEFESCTDEEFTENIKQMPCIVSML

TRYVYFRPKKPDIRD

>XP_033734454_1 XP_033734454.1 caspase-3-like isoform X1 [Pecten maximus]

MSRLTDMDTISPFDHMSPKKKSHGSTIDIHVEYDRKDSATSGTTSASSPRMSPCSVESGS

PYLLRKVSGDRRSVRPGAPSEPNEYNFNYHKRGLFLIINNKNFHPSTGQVTREGTNVDAE

RLEERFQDLGFEVRRYNDVASSKITKIMYDASEYDHSDCDCFGCAILSHGIEGRIYATDG

ILPLEMLIVPFKGERCPTLVGKPKLFFLQACRGISNEASLDLTDSVSDRSGYSEEFESCM

HPLRKIPVEADFLFSYSTVPGCYSWRNNQDGSWFIQALCIVLENYGPKMELLHMLTQVNR

IVTYEFESCTDEEFTENIKQMPCIVSMLTRYVYFRPKKPDIRD

>XP_033736656_1 XP_033736656.1 caspase-3-like [Pecten maximus]

MPDRVGSDQDASTMYQRFSDLGFEPQLLNDCTHDEMMEKLKEAASDNHSLCSSFACVVLS

HGDQDGIWATDKAPVLKTEDLMATVNSQNCKSLIGKPKIFIIQACRGLQSDPGVDVNFDD

RVMSDAASESEEYLAHLSQQENTYQNIRIPSDVDFLLVQATSPGRTAIRNNQTGSPFIQH

LSEAMSNMEEEDDFIAVLTKVNRNMAYNFSTGRMGGVKQMPCFTSMLTKKLLLKKGC

>XP_033736655_1 XP_033736655.1 caspase-3-like [Pecten maximus]

MSMQEASDTVPYGPNEGPPMPQPGQDHRNQNDGQEDPVLQSHDLYATEYKTCSEPGVAVV

INNKTFEEPDLDERVGTDRDASNLYNRFNDFGYDTKHLIDKSKHDLLEELKNIAGMDHTE

HSAFLCAVLTHGNKDGLWAKDSLIKISELTKIFDAENCPSLVNKPKVFIIQACRGNEEGK

PIVIKVQNTEDPVVDDALPMEQDENMEVQEDARDPVSVDILTIPAIMDFLIIHATVEGEK

AWRNTATGTPFIRNFCDELETLLGNRGEDDIYTLLNRVNLAVALKFQAQSGGKQMPCFVS

TLTKTFKLKAPQ

>XP_033728057_1 XP_033728057.1 uncharacterized protein LOC117317363 [Pecten maximus]DEDDED

MIDMETDSVEDQLGDIRQFSQHKLLLEVDSELDSGDIENMKFLCQDVIPLSKLQKIDRGI

DLFVQLEHANKISPKKGDMEMLVEILFRVRRLDLVKKLGQNSKEVRRKYGGSDRAVSTNL

SPLRNLLYELLEELTHDEFVTTKFVLQSEFGKGTMSKINSMMALFTFLEREQELEIEDID

ILYRIMDIVKRTDLKQKIQEYEKYPYKKGRPGRPVLPVTGLGPKPTKTPVRPVPSSGVQV

SASDVEMMDTTNVQPPAPTGTGLLDSHLYSISVELAKETRWRALCGELNINIEDLDELTA

TADSPAGKVYLMLMYWRESVVNDSPNAGRMLYLALYQSNFQILALNLALKNLFPGVVMEP

IPNKPEDNTQTGTTEQVNPPTQPVPPPAQGSDSSNEASPNIAHVAPLPQTGPNPDTQEQA

NIAIHSEEAQISIINPNPPTTNQQVQQSGQTVVPADGRMSGAVELQRQAVNETEQQRQDN

DLVRQLQRTHVDNELDLPSYKMDAKPRGVCVIINNEEFYKIPRDDTSKEMPNRTGTHRDA

ERLTYIFQKLDFLVEMHNNQTDIEIAQLLSGVGYRDHNNYDCFVCCILTHGASGHIFGSN

GKMLRINILTGFFRSQVCPSLAGKPKLFFIQACQGRDKQGGYDLEPDAAVEIHHPRKPDL

GVDINGREMIPDEADFILGYATVPGYVSYRSRSQGSWFINKLVDMLDKYAYRYDLLSVLV

KVNEEVGNAIANVDGGRYKQIPAPLVTLRKKLYFR

>XP_033724604_1 XP_033724604.1 caspase-2-like [Pecten maximus]CARD

MDEVDRRKLTSNRSYLVKNIYKTMNVVDKLMEDRIITDGMKQDIEAERTNDDRNRRLLDI

IPRRGPQAFEAFLNALVNTDNDFAAEQLSPGAGSIYKVSMTSSEKGMVSPEEEKPQNQHS

EIILPKKWPTEQDCVTDFIVKKCSHSEERKWQSDYYYKMRSKPRGRVLMFNNFVAVAHRV

KEDGSWKEKLQMREGTEKDEQSLRRLFEQLHFDVEVKKDRKVEQMRSDIAEEKAHPFHKD

ADCFILVFLSHGSTEGIYGTDGNVLTTDEVKSSFNNKNFGSMTNKPKLIFIQACRGDKYD

KGANVADNMAKDMSDARKSMSALNLNDPKLENVDEADSKVEKSPSNADMFIANATTQDNV

SFRNTKFGSWFIQALVYILRNYACQTDLLDMLTLVNKLVSEGRALDRDRNISVAVSVFES

TLRHKLHFFPGLTEVVTHSPQD

>XP_033724734_1 XP_033724734.1 caspase-3-like isoform X1 [Pecten maximus]CARD

MMEAIHKKALQKTRTRLVRDLDPCSELQDLLVSEHIFTPIMLEYIQAEKTRTDKARRLLD

DLVRRGPKAYTTFLKCLRESGHGDLANAVEDKEYELRGLPVPFRPDHQEGQSAAMMSATT

TVYTREHGPVTNPVHTEIQPPQSSFMQTQEYSAQQCQPSEESNPCGSMFTPSGASITQQV

ANPVHGISHVITGRETGGQRTEDVMEVTDGEVEQPPPQLPADPVLSGYQSVPVDRRYKME

SNPRGLCFIINNKNYTQMSNRSGTDVNRDQLMDLFQHLNFGVDIRNDLSGMEMKRQLAEL

TQYDGLTRVDSLVICLLCHGDNENMYGTDGVAVNLLTDVFKIFGPQTCPALIGKPKLFVL

NSCRGEARDVGNLNYTTITNPTGDNPQTDITHRLPRNATDVRPMTHLSLPERNPSYKDLY

IAYSTFPGHVSYRDTQDGSWFIQKLVEVFKEEAAINEIDIMMREVNRRVSMEYDPNSLEG

QMPAPSNTLTRQWFFNPST

>XP_033724726_1 XP_033724726.1 uncharacterized protein LOC117314744 isoform X1 [Pecten maximus]CARD

MEKIHKDALRKTRIRLVNDLHPDAEMWDLLVTEGIFTPIMMEYIQAEKTRADKVRKLLDD

LVRRGPKAYKKFLRCLRDSGHEDLANTVEDNEHILRGLPVPFRQGQEDLNRNTQRPPEPP

RPSSVTSQHFINIEPIPAQFSEESSATHYQPSAEWQTGPASLPCSLMTPSGARILPTAKI

PGNSSSQPGSHIATTSQPGSYIATTSQPGSYIATTSQPGSHIATTSQPGISIPNKSQTEG

HIATTSQPEHIATTSQPEHIATTSQPEHIATTSQSGGHIATGNTAGQSGDDDVVMETADA

PAPGNEVGTAYQRASAERPTYRMESKPRGYCLIINNKDYTLDSLMDPRVGTNVNREELQS

LFTDLGFQVDAKDNLTGEAMKNTLKEFAAYQGLPAVDSLIVAFLCHGNSEKMYGVDGKPI

YVQNDVLKVFSPRECPVMTGKPKLFLMNSCRGDMRDVGYPANSTVGGTVPFDAAQTDTKP

LSARPPNDLQLTPNFQDLYLAYSTFHGHVSYRNREEGSWYIQKFVEVFREQAAFCDINAI

MTQVNRRVASMYDPQSLEVQVPAPCNTLTKDWYFNPPTNF

>XP_033742269_1 XP_033742269.1 caspase-3-like [Pecten maximus]CARD

MADDEWKQDIRKNWTVIIKNLANPDELLEYLHHNITPPLLNEEQMRKIKSMTSKKEKTRK

LMSTLMYGDDRVLPAFVQALRATKNEALADKLGQCENKLTQERRQKLHLGPKVWPPSIDE

LEQMHEANAEQCDMADVLRGEYTHQEEVYTMTAERRGLAVIINNTKFKDPTKDRIGSDED

RDKLSRVLCQLHFDIRIYNNVTGADILRIAEESSHWGHHQESDCFIFVVLSHGSSEGVHG

TDNGIAHFDEIEANFNSNNCPHLTGKPKVFLFQACRITKGTKDEKNQSGTQEFDSMEPSS

EPEASNEADYDDDDVDTDIDKAFENLPDDCDQVQEDSSMDEEPGKQIPRSHVSSDFFIAY

ATPEGFRAYRQEKRGSWFIQAIVWVLKYNAHKDHLLNLITKVNNYVSTAMYGRHKGVQVS

EMRGSPQKRLYFFPGIYNDPPEKNDYTGKCINY

>XP_033740501_1 XP_033740501.1 caspase-10-like isoform X1 [Pecten maximus]DEDDED

MERVYYGQRITVETLHTYHRLFIKKYPRIQMDSKKAERKIRLETVGHQLSPDETAMLKFL

CHGVVSTKRLHECANGSALLFEIATQGNDEPVVEVLARVGRKDLAKTFSEGCEHKNALLQ

HISNLDDLGNTCLNEFRVLLVHICNELTDEEDLQKLKDITTNIPKRLLAKAVTILDFFSL

LMKGDHLQPQTLDTALLDPMRHLRITAWEKHLEKISGFIERLALREAEEREGSRVTLAAA

EYVPKEAEAEQTSEDSNESSEETQETIPKIAETEQIGEETQENMYDVRAKEAAEAEQTRD

SEESKETPEVSQENVAETENSDTDEQKQDNLGEARVSGLDYINLIGYCASNQHQINSWPS

QSSMKESGTKPKRQRVYQRVENVRLEEYRMTRIPRGKAVIFNIEKFNNSTDPMLRDRTGS

SVDADGIERIFRRLHFDVERFNDHDCSFNKIDKELKTLAYDTDHQDNDCLAIFLLTHGRN

GQLLDANCKSFDLTPLRKYFYASNCKTLAGKPKMFFIQACQGPQRQGIVYKRIEEDVVAL

KPSEVPVTPVSTSGDDYTNQSPDEADFLLCNSTPPGYVSYRDTENGTWFISTFIETLERD

CNSFDINDILLRMNRILGEKTGNLEGNPCAQIASITSSLRRKVFLRTMPNT

>XP_033740498_1 XP_033740498.1 caspase-10-like isoform X1 [Pecten maximus]DEDDED

MLHFHHKGTVGCRMDSKKAERRTRLESIGDELSLDNICILKFLSHGVVPARKLQECTKGS

TLLLEIGAEGNDVHLVEGLARAGRKDLASRYSEGCQNEGEALYSQMMKNFDDLEKTCLNE

FRVLLVYIANELTDSEDLQKLKDVTTGVPKRALAKAATVLDFFNLLMEEKNLLPQTLDTA

LLNPMDRLRVTTWKKHSEQISGFKERIALRESGEREGSRVKPTAETIPKVAESKQTSGDR

KECLEEAQETEPKLAENEQTGERTEETATKVNAVEAKQTDELAERSEKAQETVSKVTETE

HTREKTRENIQEAGGQGVERTEHPEISGENEERPDVSQENVTETEHSNTGGQKQENQDQD

PEKRQDGATSEQPHIEDGRRNTLNPDTDSGAKPKRHPVNRRMKNIRLKEYRMTCKPRGKA

VIFNIEKFSSLYSHPTDQKPRDRIGSSVDADGIQRIFKQLHFDVERFDDCLLHMLKDRLW

KLAYETDHDHFDCIAIFILTHGSNGQLLDANCKSFDFEPLRKYFYASNCKTLAGKPKMFF

IQACQGSVTQGRYNHLEKDTIVVEPTTLSSTPSSTPSSSPPSTPSSSPPSTPISFAESQG

NQSPDEADFLLCHSTSPGYVSYRHTEKGSWFISTFIETLERDCNRVDLLEILTRANCILG

GKTGTVKGKPCAQIANIITSLRGKVFLRTIPDT

>XP_033736246_1 XP_033736246.1 caspase-7-like [Pecten maximus]

MSDQSDPSKPRHPVATGENGDYVFHPGVRDSSKLPKRSEEDLMREVYNIRGQTESSRQAL

IINNLTKITGDKDSNLMESNLQKSGFSVKCKTRLTQEKLEECLKTVADDMADDTDCFVCV

VLALGGNGFIKGWNSNVSQTKSLTTEKLCEPFQGNNCKPLVLKPKIFIIMSWDTEPDSRI

QIDGDADQSEVFKVPTESDVLVYVCHMPGKFKQGYDGDGSWFIKTLADVMQKNRERCKRR

DFLRLLTIVNREMANKYKGTKLKRSECIPTVTSTLTKLIYI

>XP_033762013_1 XP_033762013.1 caspase-7-like [Pecten maximus]

MASAALQYQPLSGPLPASLISFSRSSVDQDLSLQNNTNRIYSPPLRYTDSSCIQVNNVVT

DKPHNDPVTDASLSESMDDLETDGLDNQENLSCNLYLDDEHKSNNLKSDIENDRINFVDE

SQETAVKSVPGESCDQWPQENGCAFKVKAVDMDRRQPTKETVEKTIIRTHQDFVSAEYRH

DHLYRGRCLIVSNSEFRDKKNRPKRDGSLRDARTLRNQFLALGFHPIRTKDGSDEMIHQN

LSPEDLLQILSEVTDPSLNDHSDYDCFACVILSYGLEGAVVCPGETEDVYLEINSILEHF

TPQKCPSLAMKPKLFFIQGCTPYEIIQNQAMAKTPSSTTWCPMMKTVIPSMADCLVQFSG

VDGCLGWGRKRNRGLSYYVQALCQTLKKFCILKLEQGQKSQEMTSLLLRINRAFTHILED

SDIDMDETYCVPLTYSTLTKQMYFYPKSPQDDLETG

>XP_033724674_1 XP_033724674.1 cell death protein 3-like [Pecten maximus]CARD

MEDYHLELLRKARPIIVKNMADIGAVCDHLIAFDILTESMAEEILSKSSNASKARALLHV

LVMRGPKAFDLFHSTMIITKNHTIADILKPPVSPQKLKKSVSLPSPECEDFPGEPVQLKY

STIDRRARKHPLSRQQGVDDAKTEYGEYEIETSQKKGTESKEVFGALAQVCEESNLSSGI

LVSGSDQHVSHFASEDVSQEADIHREMDDIQRYRDRSENVGHSFDRWTSIPTDSKQITDP

FTSSQVQTEQGNQVHVASKDDFIQHLVETSSSSQLVDEDEDEMEDEGRNWPATRKHSRPL

VHLTFRQPAIDRQKFIKDITSNEVYHIDPKCQRRLVIFNNSVFPGLPPRTTGNIDATSLD

LLFKKMSYSTKTYSDCSCERIMDVLGEELNSNTLHHMETFILILLSHGKGEKIYGSEGNM

VDIKDITKLFDNEHCPALKGKPKLFFISACEGQQNKSGGEACVDSSLGNVGYDSLSQDML

NSLVSMEMDDGRTESSDMFIARATTQCDATFQQVSFGSQFIQALVHVIKEHSFIDDVVTL

MDKMNQLVQRPSGSPLVEYSNTLPKKFYFFHNSKHSLTLTI

>XP_033749873_1 XP_033749873.1 uncharacterized protein LOC117334389 [Pecten maximus]

MSNREEEDEEKHGATAAQPPHSTTDGLLSSCFPPILGKKKCSSAIDDDGEVYSKANTKSE

TTEGQTLDAEKPEDVSKQHKGGSQANVIAEGSQKHQGGSQATASAEGSQKHQGVSQANVT

TEGSQKHQGGSQATVTAQLAHQQGGAQNVIDDDDEEECFYPINQNCPGVAILIGHSKFDT

GEKERKGAHMDLKNTFEVFERLGFLINFFMDEEVMVLQHKLKELAMSERVKKADCLVCVV

STHGSEQEVGKGDRDPRLPSNVKVFEHFLSMKNGVIKTRELIEIFDDSHCKALRGKPKLF

FIQACRSRTDKSLLQSMDPGIQVDIVGPVLRPLPTNTTDFMSGARSALDNQGSQPLSDQR

SQSQVDARAYMAPVLERDGEMYYEHTPPFPLFEDIVEVVPPVCPNNFLIMFATPSEMVAW

SEDDSGGWLLSALYLEFIKHIRNGTRIALLPTLTDVSRHVAYKYASRMEGTDYHKMKSVP

CLEHKLTRDVYLTPSKKTSY

>XP_033734730_1 XP_033734730.1 uncharacterized protein LOC117323554 [Pecten maximus]

MIAYDFNHFRRGVAVLIVNTEFCNVKDNRPRAEKDIDYLTALFEVLDFQIKVLKNKTTRD

LMDSLIDIRSQLEKDSDCFACLISTHGGEFPVQGSKNLHQHVLHTFDGSIPTDDIISTFN

DNGCAAMRGKPKLFFIQACRGRMDVDDTELVDPGVDVDIITYNSQPPLAGLREESRHVSQ

HQGQNGTHDCANTHGDGLEKEEGHHGNQLTTQEMVGQEVDLQITGKDHGVTDGDDSKTTT

KADAKGRSDDNPARMYGPGYIPFIDGEEDQTYANEEEQRRHREAWCRYQEKVQKDQEQKR

RERFLELQLKREMEDADFCSIPCYSHCLVMFSSAPERLAWGDDHTGGWLPYCMYNVVHWF

HRYDFRTDLLQVLTEVNNEMAIYLQTNMPSKPEWHQSKSAACIYHMLTKDIYFRLKWSKK

AALVTAV

>XP_033733480_1 XP_033733480.1 uncharacterized protein LOC117322653 [Pecten maximus]

MIGLKTHGNEPKPKKKPEVPPVDECPKHGDDQPDDRYDFNHSQRGVAVLIVNTEFRNAKK

NLPHAEKDIGYLTSLFEVLSFQIKVLENKTTQDLKDSLIEIQSKLEEDSDCFACLISTHG

SECPVQDSKNLRQHVLHTFDGSIPTDDIISTFNDNECAAMRGKPKLFFIQACRGRMDVDD

AELVDPGVDVDIITYNSQPPLAGLREESRHVSQHQGRNGTHDCANTHGDGLEKEEGHHGN

QLTTQEMVGQGVDLQITGKDHGVTDGDDSKTTTTADAKGRSDDNPARMYGPGYIPFIDGE

EDQTYANEEEQRRNREAWCRYREKVQKDQEQKRRERILELQPKREMEDADFCSIPCYSHC

LVMFSSAPGRVTFSDDHTGGWLPYCMYNLVHRFQSYDFRTDLLQVLTEVNNEMAMNMKTH

FPKKPEWHQSKSVACIYHMLTKDIYFRLE

>XP_033734732_1 XP_033734732.1 uncharacterized protein LOC117323556 isoform X2 [Pecten maximus]

MLLSEEVINYYNDIPPQNDPDYSRVASLSESPASSPTIPRENARSPNDSTSNPQGSSGDA

FFGKVFDDQHSSQSTSTTPEHVYDFSYRSRGVAVLIVNVTFRDKIMSPDRIYAKHDIEVF

EKILKALNFTIIKLIDHSTADLRRRLEEIRSSLTSDCDCFLCLISSHGLETRISDDKRKR

QHTTFTYDRWIPTDSIIRPFTENQCRCLSGKPKLFFIQACRTVHRSADADSIDRGVITTE

AVCQPPGHHGDVRDNPVRLEGTGNTAIDNLKTDQDDLNSSSQDLSQSVREKVNLDEIPFA

DETDDSDDPDFSTADNSRASYYPPRPRQSDDVDFCTIPCYPHCLVMFSSAPERFAWSNDF

KGGWLPYCMYKVLYKMAQRHFRIDLLQALTEVNETMAKSLETNSPSDKVLHQVKSAACIY

HMLTKDIYFTPKYAPVPVVDRVQVSAV

>XP_033725096_1 XP_033725096.1 uncharacterized protein LOC117315065 [Pecten maximus]

MEDEDMQYDFSKQKYPKRGLVYIIVNERYSAEGYPDRHGAYTDYRNMKNLFKPLNLKLMT

YYDKTGKQIKDIIQNASKRPDLADRPCFICVISSHGEQWPDLDRKERKKTMEHVMDHVIV

GVDGMGVLTKDLVGFFSKDKCFDMVGKPKLFFIQACRETFYSNGGSKAVLYSTDRSKAVL

YSTDRSKAVLYSTEAFDRGVAVQYVVSAKLTKHRRMRQRLSLNTKYDSADSKPNQSEVPL

DALDHIIDPTGNGDSDDDDQLSDGYDTATTDEDSDADSDRETGHEEIKKRDRSVNTENEE

VISKDDVKIEIRHIQATTPSDQNKDLNVVPKEEEKTEIQQSERMVPSGVPLDQDDAGASG

KRRVISDIQEAEPMEVTAIPCHDDMLVMFASPPGKIAWRSTSEGSWLLANLYNVFKQYIR

SRKISERDFLSILIEVSQDVSLKETNTVETEFKNMKMVPTIYHRLTKEVVMCFGKETGLL

KSLKRRFQSLSHNDDLK

>XP_033761469_1 XP_033761469.1 LOW QUALITY PROTEIN: caspase-3-like [Pecten maximus]

MATRITKEMKSRHTRLLNDLANHFDPALNGHEFDQLKNSCRVHIQPLALKNSNNAIELLK

NLERNNTISVGKYDELKKIIKEFDFTVLDVITEAEDDLTQLGLQAPGAASSPVQQPQSDS

NPPSSRKTNLNVPGANTGELESQHPSPGNTDKPSAPKKKRDDEPDDYVYKTNGKTGLLVI

LNFTQKREGALEDERNLKGFFKDSLQWEVETRQDLKYDRLASVSAGITERLSDHETADKY

HSLVVAVMSHGNEDGIFPHDHLEEVEEKRVYKFDTIFNFFKNDEIKAFAGKPKVFLVQAC

RGSQIQGEVIQDDHDPIDIGELPLPASVRISIPVDADILIMWATTPGFKAFRSVISGSHF

LREVVHQFEQNYGTDHVEAMLMKVRHTFACTEKYRWKHVGTDKDGRKKTSVSVQMPCTWT

TLTKMLYLK

**Supplementary File 4: Caspase repertoire of platyhelminths species which are included in analysis of Figure 3.**

>TNN06772.1 Caspase-2 [Schistosoma japonicum]CARD

MEREHRDVIRKLRVYFVKNVSNLEEILDFLESEMIITENQRHSVLSGSTVAEKIRKFLDI

IVRCGPLAYDKFKEALLKTNQKGICDVINETEELKKSVGCNISSECTPGNKDNYIMTPDR

LSSHPLTPYTVTSFPRGYVLIINIEEYSRESGVSSRLGSSHDVTKLQLLFEDFMYSVAVI

RNVNGENLIKVVREFARKPEHNNVHAAVLIILAHGLEHHIIASDGIHVSIDELVSCFTNK

RCPLLAGKPKFILIQACRGEERNHNDSKMGDFLDSLPSNSREISLDPSSWLSLPHMSDCV

IAYTTLPGFVSWRSETEGSWYVKVFVEVTRMHGHRLHMLELLTEVNNRLVSEASNYGIKQ

ISQPVTTLTKPYYLSTLKA

>TNN08539.1 Caspase-8, partial [Schistosoma japonicum]ANK

MQSNEQNEDVVMWTKTLHQSINSFNIKQLEDVLIKLNTTLGLEFTKEVLNYSVDGANSVY

VAVEAGFEKAVTLLLDTGMVNVTQRNYQNNWTTLHLACYLGLATIVKSLLNFNASVNSVQ

INELYQLACIQTIENTKSSNQGTALHLAVYSKNSETVNQLLHDVDKISKCYENQCLQNER

KYKETINFKGKLFDEFCTLSDKNGNTALHCAVMNKLYNIVDTICQYHSKSIRLPNKHGYT

CIYLAKNILKDVRMLSLLIKYSTSTTPSNSHDHHHHHRHFGQHSFKQVNSINSSRVSSNN

PLNRKEIKKSSSNLIKTTELIKSNKKNIYKKLKKCKHSLNSLTIEQLNKNINYNYTTNNI

TQNTNNRLSMYSYHNDDYYSMRTLSTKMSYLSIQSKNKVHFQPLNGLSGNRIDLNERELN

RNLCQNIEQVNNETIQCVQNRNNSIENSILQSNSSDEPCLAKCKNATAISTTQKNNSISK

TEKIDQPTYRQKKGIGDIDTCQLTTDDNYNINNKIENDSNGNNEDDSDDYYKMNSTRKGI

CLIINNMMFWHSDFQNRSGCDVDEKSLEKVFESLNFLVKLLRNLSAGEIQAQLEHLGKKT

NHNEYDCLIICLMSHGTIGRIYGVDGNSLSIHELTSLFTSDNCPTLAGKPKLFFIQACRG

DDYQKGYVIQRQQSIETIQTNQENDINQ

>AAX26691.2 SJCHGC07329 protein, partial [Schistosoma japonicum]

IINIEEYSRESGVSSRLGSSHDVTKLQLLFEDFMYSVAVIRNVNGENLIKVVREFARKPE

HNNVHAAVLIILAHGLEHHIIASDGIHVSIDELVSCFTNKRCPLLAGKPKFILIQACRGE

ERNHNDSKMGDFLDSLPSNSREISLDPSSWLSLPHMSDCVIAYTTLPGFVSWRSETEGSW

YVKVFVEVTRMHGHRLHMLELLTEVNNRLVSEASNYGIKQISQPVTTLTKPYYLSTLKA

>AAW26244.1 SJCHGC00800 protein [Schistosoma japonicum]

MGESNPTDEKHVNKGSYLSVSTDDLHNPYFSYPVVSPDRPGQFYRGECLLINQRDFDPST

NQSKRNGTDIDADRIERVFRSLNYKVTRILNITKKALDQTLIEASQKDHSSYDSFVFVML

THGDKDVVYASDDEVQTSYIMSFFRGDRCPSLIGKPKLFFFQACRGVSFDKGVSTMVADA

GDDVIVRKLPVEADILVSYSTVPGYFAWRNSTKGSWYIQELCNALETDIKNGNHTDIMTL

LTVVARVVAYQYRSSTRQVETDNMKQMTSTFSTLTRLFHITGPPKEIK

>CAX71159.1 caspase 3, apoptosis-related cysteine peptidase [Schistosoma japonicum]

MEEKVDGVGFLGTLSNPTGGKPVNKGSYLSVSTDDLHNPYFSYPVVSPDRPGKFYRGECL

LINQRDFDPSTNHSRRNGTDIDADRIERVFRSLNYKVTRILNITKKALGQTLVEASQKDH

SSYDSFVFVMLTHGENNIVYTSDDEVQTSYIMSFFRGDRCPSLIGKPKLFFFQACRGVSF

DKGVSTMVADAGDDVIVHKLPVEADILVSYSTVPGYFAWRNSTTGSWYIQELCNALETDI

KNGNHTDIMTLLTVVARVVAYQYRSSTSQVEADNMKQMTSTVSTLTRLFHITGQPKEIK

>TNN08040.1 Caspase-3 [Schistosoma japonicum]

MEEKVDGVGFLGTLSNPTGGKPVNKGSYLSVSTDDLHNPYFSYPVVSPDRPGKFYRGECL

LINQRDFDPSTNHSRRNGTDIDADRIERVFRSLNYKVTRILNITKKALGQTLVEASQKDH

SSYDSFVFVMLTHGENNIVYTSDDEVQTSYIMSFFRGDRCPSLIGKPKLFFFQACRGVSF

DKGVSTMVADAGDDVIVHKLPVEADILVSYSTVPGYFAWRNMTRGSWYIQELCNALETDI

KNGNHTDIMTLLTVVARVVAYQYRSSTGEAETDNMKQMTSTFSTLTRLFHITGPPKEIK

>KAH8863223.1 Caspase-3 [Schistosoma japonicum]

MEEKVDGVGFLGTLSNPTGGKPVNKGSYLSVSTDDLHNPYFSYPVVSPDRPGKFYRGECL

LINQRDFDPSTNHSRRNGTDIDADRIERVFRSLNYKVTRILNITKKALGQTLVEASQKDH

SSYDSFVFVMLTHGENNIVYTSDDEVQTSYIMSFFRGDRCPSLIGKPKLFFFQACRGVSF

DKGVSTMVADAGDDVIVHKLPVEADILVSYSTVPGYFAWRNSTTGSWYIQELCNALETDI

KNGNHTDIMTLLTVVARVVAYQYRSSTGQVEADNMKQMTSTVSTLTRLFHITGPPKEIK

>KAH8863222.1 Caspase-3 [Schistosoma japonicum]

MEEKVDGVGFLGTFYPVVSPDRPGKFYRGECLLINQRDFDPSTNHSRRNGTDIDADRIER

VFRSLNYKVTRILNITKKALGQTLVEASQKDHSSYDSFVFVMLTHGENNIVYTSDDEVQT

SYIMSFFRGDRCPSLIGKPKLFFFQACRGVSFDKGVSTMVADAGDDVIVHKLPVEADILV

SYSTVPGYFAWRNSTTGSWYIQELCNALETDIKNGNHTDIMTLLTVVARVVAYQYRSSTG

QVEADNMKQMTSTVSTLTRLFHITGPPKEIK

>AAX25947.2 SJCHGC04214 protein, partial [Schistosoma japonicum]

PNPGGGKPVDEGLYLSVSTDDLHNPYLSYAVVSPDCPGEFYRGECLLINQRDFDPSIDQS

PRDGTDLDADLIERVFRSLNYKVTRILNITKKALDKTLVNASQIDHSSYDSFVFVMLTHG

EYNVVYATDDEVQTSYIMSFFRGDRCPSLIGKPKLFFFQACQGVSYDKGVSTMVADAGDN

IIVHKLPVEADILVSYSTVPGYIAWRNMTSGSWYIQELCNALETDIKNGNHTDLITLLTV

VARVVAYQYQSSTGEPETDI

>KAH8878343.1 Caspase-7 [Schistosoma japonicum]

MICFRDLSVTVENNSLSQARSSLHYSNSQYRPTNKTQISPSKLSVSCASLLPVFSLPSEK

SSSFVYRTNGAERGLCVIFSVDKFDPILCLPRRTGSLVDVQNIKRAFLSLDFKVIVHSNP

SSKTLFSVIQTISTQNLRDHDCFACVILSHGDEGGLVYSTDGSIPVERIIAPFRGDQCSD

LRGKPKLFFIQACRGMSLDDGVFVCDGPSSNSKPTDTATAMRRIPIEADLFIAYAVQPGY

YAFRNSMNGSWFIRSLSDALLRYGNTLDLLSIMTRVNYDVAYEYESTTANPALCGKKQMP

SFVSTLTKHVIFPPKRSKNLSKLM

>TNN10203.1 Caspase-7 isoform 1 [Schistosoma japonicum]

MQASGFPQAEKTLMVDGVDNCNTDLSVTVQNNSLSQARSSLHYSNSQYRPTSKTQISPSK

LSVSCASLLPVFSLPSEKSSSFVYRTNGAERGLCVIFSVDKFDPILCLPRRTGSLVDVQN

IKRAFLSLDFKVIVHSNPSSKTLFSVIQTISTQNLRDHDCFACVILSHGDEGGLIYSTDG

SIPVERIIAPFRGDQCSDLRGKPKLFFIQACRGMSLDDGVFVCDGPSSNSKPTDTATAMR

RIPIEADLFIAYAVQPGYYAFRNSMNGSWFIRSLSDALLRYGNTLDLLSIMTRVNYDVAY

EYESTTANPALCGKKQMPSFVSTLTKHVIFPPKRSKNLSKLM

>CAX76580.1 caspase 3, apoptosis-related cysteine peptidase [Schistosoma japonicum]

MGESNPTGGKLVNKGSYLTVSTDDLHNPYFSYPVVSPDRPGEFYRGECLLINQRDFDPSV

GQSPRDGTDIDADSIERVFRSLNYKVTRILNITKKALDQTLVEASQKDHSSYDSFVFIML

THGGNNIVYTSDDKVQTSYIMSFFRGDRCPSLIRKPKLFFFQVSRGASYDKGVSTMVADA

GDDVIVHKLPVDADILISYSTVPGYITWCNTTKGSWYIQELCNALETDIKNGNHTDIMTL

LTVVARVVAYQYRSSTRQVETDNMKQMTSTFSTLTRLFHITGPPKEIK

>AAX25741.2 SJCHGC03058 protein, partial [Schistosoma japonicum]

NPTGGKLVNKGSYLTVSTDDLHNPYFSYPVVSPDRPGEFYRGECLLINQRDFDPSVGQSP

RDGTDIDADRIERVFRSLNYKVTRILNITKKALDQTLVEASQKDHSSYDSFVFIMLTHGG

NNIVYTSDDKVQTSYIMSFFRGDRCPSLIRKPKLFFFQVSRGASYDKGVSTMVADAGDDV

IVHKLPVDADILISYSTVPGYFAWRNSTTGSWYIQETMYALETDIKNGNHTDIMT

>CAX76587.1 caspase 3, apoptosis-related cysteine peptidase [Schistosoma japonicum]

MGESNPTGGKLVNKGSYLTVSTDDLHNPYFSYPVVSPDRPGEFYRGECLLINQRDFDPSV

GQSPRDGTDIDADRIERVFRSLNYKVTRILNITKKALDQTLVEASQKDHSSYDSFVFIML

THGGNNIVYTSDDKVQTSYIMSFFRGDRCPSLIRKPKLFFFQVSRGASYDKGVSTMVADA

GDDVIVHKLPVDADILISYSTVPGYITWCNTTKGSWYIQELCNALETDIKNGNHTDIMTL

LTVVARVVAYQYRSSTRQVETDNMKQMTSTFSTLTRLFHITGPPKEIK

>CAX76583.1 caspase 3, apoptosis-related cysteine peptidase [Schistosoma japonicum]

MGESNPTGGKLVNKGSYLTVSTDDLHNPYFSYPVVSPDRPGEFYRGECLLINQRDFDPSV

GQSPRDGTDIDADSIERVFRSLNYKVTRILNITKKALDQTLVEASQKDHLSYDSFVFIML

THGGNNIVYTSDDKVQTSYIMSFFRGDRCPSLIRKPKLFFFQVSRGASYDKGVSTMVADA

GDDVIVHKLPVDADILISYSTVPGYITCCNTTKGSWYIQELCNALETDIKNGNHTDIMTL

LTVVARVVAYQYRSSTRQVETDNMKQMTSTFSTLTRLFHITGPPKEIK

>CAX76582.1 caspase 3, apoptosis-related cysteine peptidase [Schistosoma japonicum]

MLTHGENDVVYASDDKVQTSYIMSFFRGDRCPSLIGKPKLFFFQACRGVSYDKGVSTMVA

DAGDDVIVHKLPVEADILVSYSTVPGYFAWRNSTKGSWYIQELCNALETDIKNGNHTDIL

TLLTVVARVVAYQYRSSTREVEIDNMKQMTSTVSTLTRLFYLTGPPKEIK

>KAH8878344.1 Caspase-7 [Schistosoma japonicum]

MAHAYTYFQDYVISVSTQNLRDHDCFACVILSHGDEGGLVYSTDGSIPVERIIAPFRGDQ

CSDLRGKPKLFFIQACRGMSLDDGVFVCDGPSSNSKPTDTATAMRRIPIEADLFIAYAVQ

PGYYAFRNSMNGSWFIRSLSDALLRYGNTLDLLSIMTRVNYDVAYEYESTTANPALCGKK

QMPSFVSTLTKHVIFPPKRSKNLSKLM

>KAH8863219.1 Caspase [Schistosoma japonicum]

MLTHGDKDVVYASDDKVQTSYIMSFFRGDRCPSLIGKPKLFFFQACRGVSYDKGVSTMVA

DAGDDVIVHKLPVEADILVSYSTVPGYFAWRNSTKGSWYIQELCNALETDIKNGNHTDIM

TLLTVVARVVAYQYRSSTREVETDNMKQMTSTFSTLTRLFHITGPPKEIK

>KAH9283631.1 Caspase-3 [Echinococcus granulosus]

MWRKGETLNPLCYNDLQFSDRSPCDKDTTEASPPVDSALVAPTDKMKNIFPEKMFERSLF

SCKLMPLESQSDTPAYKRQAVESSAHHRAATSFVYKTRAPRRGTCLVISVDTFKAALGLP

SRPGADVDLKKLEDTFLFLDFDVKTYQNPSATQILSIVETESSANHADADCFACVILTHG

DDAGSIYGTDGPVPLDQVIHPFRGNTCPSLAGKPKMFFIQACRGAMLDSGTLVSTDSSGG

PGDSDTLGGLRRIPIEADLLVAYAVQPGKFTGYYAFRNSVHGSWFIQALNNCLRKYGRIL

DLMSIMTRVNYEVAFEFESLASTAAYSGKKQVPSIVSTLTKDVFFPPKTTAPSTCTSSLP

PSRTP

>XP_024351257.1 Caspase-3 [Echinococcus granulosus]

MSAKVLALKSPCDKDTTEASPPVDSALVAPTDKMKNIFPEKMFERSLFSCKLMPLESQSD

TPAYKRQAVESSAHHRAATSFVYKTRAPRRGTCLVISVDTFKAALGLPSRPGADVDLKKL

EDTFLFLDFDVKTYQNPSATQILSIVETESSANHADADCFACVILTHGDDAGSIYGTDGP

VPLDQVIHPFRGNTCPSLAGKPKMFFIQACRGAMLDSGTLVSTDSSGGPGDSDTLGGLRR

IPIEADLLVAYAVQPGNLEITRVISNSTSFSLGYYAFRNSVHGSWFIQALNNCLRKYGRI

LDLMSIMTRVNYEVAFEFESLASTAAYSGKKQVPSIVSTLTKDASIFYVATVVKVMSAVP

NDLRELRGDLWVRCVGVQRSILLGKKQVLSIRAIRTHRDDGGSIYGLDGTIPLDQIIHHF

RGNPRLFLVGKLKRVFQIPKFFDKIKYEWLISPFGDQAREESSPYNLFAYPIQGHSRRTG

KQAKRVMRNFARYQDVESRVLGLAIAAPLET

>XP_024351559.1 Caspase-3 [Echinococcus granulosus]

MDRCDGFLFLTASSSTGGKASPTPRGTTSPCQASPTLSKSVKPAEVTTGIGSGDRRRYLS

EIELCDPDLGYPRVANPTNRSNPRGVCLLINQRDFDSAKTGQERRDGTDVDADNIERTFI

RCGYAVNRATNLTLRKMEFLLDDVRSQNHSKYDSFACVILSHGAEGIVYASDGTINVDRL

IGYFRSDRCPTLAGKPKMFFIQACRGCKFDKGVTLSTDASSDSVLVTKLPIEADIFVANS

TFPGYYAWRNSHAGSWFIQELCKVIKAAQESGRHHDVASLLTVVARKVALLYESNTGQSD

SHGSKQMVSVNSTLTRKAFIV

>KAH9280007.1 Caspase-3 [Echinococcus granulosus]

MENKVLSSKPTADIPVKDDAEDASFIFSSTGGKASPTPRGTTSPCQASPTLSKSVKPAEV

TTGIGSGDRRRYLSEIELCDPDLGYPRVANPTNRSNPRGVCLLINQRDFDSAKTGQERRD

GTDVDADNIERTFIRCGYAVNRATNLTLRKMEFLLDDVRSQNHSKYDSFACVILSHGAEG

IVYASDGTINVDRLIGYFRSDRCPTLAGKPKMFFIQACRGCKFDKGVTLSTDASSDSVLV

TKLPIEADIFVANSTFPGYYAWRNSHAGSWFIQELCKVIKAAQESGRHHDVASLLTVVAR

KVALLYESNTGQSDSHGSKQMVSVNSTLTRKAFIV

>KAH9278879.1 Caspase-8 [Echinococcus granulosus] ANK

MPQTTSKNINRLPSSSSTSTSATDTEMLLQLLRRELLLALESMDGAVFRMKLQWILENAG

EFVDLVFMSGLDFEGKTALHIAVANRFEEAVKLLVSTAKVDLLATAGDNGWSALHTAVDL

GDAKLTQLLLPTHNTTLTPHHVLSSLKQATASPYAVTALHMAVRRNSVPCAVMLLHTANA

LLNDHSSGWTSSEEELAVDEIQQKHHFHGQAGKGSRKLNKLKDRNGDTPLHWAVKTNNYA

MARLLCSYGAHPRHSHNCQGQTPFALAKYVLRNTKMTKLLASFSPTSSTNTSVTDNFEDD

NEETRIKEDEAIMPLKPPSPTESTLRRVVKASEVDGSCQKLQQTLIRVEVPTPTPGTVTA

VTAMPVKKSPSGEASNGGSILGRPYLHREVKSTIPQGSLDMKVDSDTLEACRLRSNSLKV

GRREMDMLRQKLPGLLSRPLLKRAVVHRPRPVSLKARRKASLMDPVEYYPMASPRRGICV

VINNMTYWHPKFQNRGGCDKDEKRVEKVFCGLGFLVKVLRNLSSGEMRSELAFIGNKTDH

SAYDAFIAVIMTHGGLGELYGVNGDAFPVHQLTLDFTAERCPSLAGKPKLFFIQACRGDE

YQLGYALPATGITEDAVTSNPGSALPNPLQIGDTQVEGEAVEEATDDVEEKPAIVAAKLP

RKQRLVPNYADFLLSYATLAGFKAQRDPQKGSIYIQTLCHHLEMHGRSRSLLDIVTSVHR

EVSEKVFREVESPDQEGTVFQQTPEVRHTLTRRVQFV

>XP_024352296.1 Caspase-2 [Echinococcus granulosus] CARD

MNSEQRQRLSRMQVDFVKGLRDIEEITDYLLCHEILSRTQAEKILSSGPIIADQVRTLLN

VLVRCGPDAYDTFVSALSETNQSSLAEKMANLNTNPPPVPSVLPSAKVETHVSEQPTQSL

AAYPVNSTPRGHILLVNISNFEVGSGLSNRVGSSEDVAALKSLFTDLGYYVEVLLDPTAT

ALESNLSAFIKSPSHYNVDTGGLIVMSHGVQDYIYTADGKLFAINDILEAFTNKSFPALA

GKPKFILFQACRGEEKDRGYIYEPDNASFVASPLESVDAGVHKTLWKCLPYMSDYIIAYS

TLPGFVSWRSEKAGSWFIQILVDVFRKYASSLHVLDLLTEVNRRLVEESQEREFKQITQQ

SHTLTRPFYLAGTQSQPR

>KAH9285259.1 Caspase-2 [Echinococcus granulosus] CARD

MGRCYFLTPGMNSEQRQRLSRMQVDFVKGLRDIEEITDYLLCHEILSRTQAEKILSSGPI

IADQVRTLLNVLVRCGPDAYDTFVSALSETNQSSLAEKMANLNTNPPPVPSVLPSAKVET

HVSEQPTQSLAAYPVNSTPRGHILLVNISNFEVGSGLSNRVGSSEDVAALKSLFTDLGYY

VEVLLDPTATALESNLSAFIKSPSHYNVDTGGLIVMSHGVQDYIYTADGKLFAINDILEA

FTNKSFPALAGKPKFILFQACRGEEKDRGYIYEPDNASFVASPLESVDAGVHKTLWKCLP

YMSDYIIAYSTLPGFVSWRSEKAGSWFIQILVDVFRKYASSLHVLDLLTEVNRRLVEESQ

EREFKQITQQSHTLTRPFYLAGTQSQPR

>VZI03839.1 unnamed protein product [Spirometra erinaceieuropaei]

MWEDKLYSQQCFKDLRLRSRNSPDPNESDEILPVDSAHVPPILRPSNASFVCGEDLRLPP

QKVPITSTIPPHASKLNSPIPQILPTRSPASFVYGTRAPRRGTCLIISVDTFKPALCLPG

RPGAEVDLRKLQQTFSMLDFDVKMYCNPTAANMTAIVEAESKANHADADTFVMVLLSHGD

DGGLIYGTDGAVYLENLIKPFRGDSCPDLAGKPKMFFVQACRGSQLDPGTVVETDGFYPG

GEGGHLGSLRRIPVEADLLVAYAVQPGYFAFRNSLYGSWFIQALTNCLQRYGRTLDLMSI

LTRVNHEVAYEFESMASNPEYSGKKQVSSIVSTLTKDVFFPPKKPSSRS

>VZI43280.1 unnamed protein product [Spirometra erinaceieuropaei]

MSETADENDAGLFARPMKVSGFGAACDRERRGVVLSKADLCDPTLQYFRLSAPHNRNNPR

GICLLINQRDFDREKTGQERRDGTDVDADAIERTFIKLGYSVNRATNLTLRKMQMLLSNV

SCQDHSHFDSFVCVVLSHGSDGIIYASDGYLPADNLLSFFRGDRCPSLVGKPKMFFIQAG

SNIVFIQIHEACSPLSLSRACRGSAFDRGVSLASDAGAEDVLVCKLPSEADILVANSTVP

GYFAWRNSNTGSWFIQELCTVLGADAASSGEHFDIVTLLTVVSRKVALLYESNTGQPGSH

GMKQMVSVSSTLTRRAFLTG

>VZI50365.1 unnamed protein product [Spirometra erinaceieuropaei]

MVLLSHGDDGGLIYGTDGAVYLENLIKPFRGDSCPDLAGKPKMFFVQACRGSQLDPGTVV

ETDGFYPGGEGGHLGSLRRIPVEADLLVAYAVQPGYFAFRNSLYGSWFIQALTNCLQRYG

RTLDLMSILTRVNHEVAYEFESMASNPEYSGKKQVSSIVSTLTKDVFFPPKKPSSRS

>VZI32601.1 unnamed protein product [Spirometra erinaceieuropaei]ANK

MKSTARESRLGPYSNSVETEATLSLMRRGLVQSLESLDVELFQRKLEALLEGDFADMVLQ

SGLGLEGKTALHLAVECRFQEALKLLLATDKVNLLAISGETGWTALHTAVHLEDPTAIRI

LLRYPIETLDAAGSTGDETVEHCCEDRLLRARELLWCHSTNPIGATALHMAVRRNNAECV

RVLLEDANWLHSRLAEFRPSEYGASDLEGSCELCYIQDRNGDSALHWAVSNRSHEIARLL

CSYGAQIHRPINQKGHEPLTVCKDVLGDDEMTAILTPFRRADPAVDTSGHSVHQPCIVVA

DSASNQTSSRLRRAGVTTGTSKSLTEGGQRTTADQQADPVSQRRPQEQPMISGEDRGGCD

KDESRIEKVFRDLGFLVKLLRNLSSGEMKAQLKHLGTATDHSTYDCFVSVIMAHGGVGEI

YGVDGSPFLLHEITAYFTADKCPTLAGKPKLFFIQACRGEEYQLGYAPPSRVSAAEPPQP

TPGLPTRATTEAPRDTSSNPFQARSSFVEAVPVDPPSMPADRSTANPDSEAVAREQEPAV

LAAKLPRRQRLVPTYADFLISCATKAGFKAQRDPKEGSIYIQSLCRHLEQYGRERSLLDI

ITGVHREVSERVFVAASSSEAGSGPIFLQTPEVRHTLTRRVQFV

>ACU88130.1 caspase-7 [Schistosoma mansoni]

MVDDVDSSNKNLSVVEEHSSTQIRSSLSCDSPQNKLNNKTWSNSLKCSVSCSSLLPVFSI

PSEKSPSSYVYRTDGAERGLCVVFSVDNFDPILCLPPRNGSLVDVQNIKRAFANLDFRVI

VHSNPTSESLFSLIRTISTQNLRDHDCFACVILSHGDEGGLIYATDGSIPVDRIIAPFRG

DQCLDLRGKPKLFFIQACRGMALDDGVFVCDGPLSNSKSADAATTIRRIPVEADLFIAYA

VQPGYYAFRNSVNGSWFIRALSDVLLRYGNTLDLLSIMTRVNYDVAFEYESTAANPSFCG

KKQMPSFVSTLTKHVIFPPKKPRNPSKLM

>ACU88129.1 caspase-3 [Schistosoma mansoni]

MGEVVDGVHLMTTKPNQGRGQMADKDGYLSVSMADLHNPHLSYPVVSPNRSGKFYRGECI

LINQRDFHPSTNQSRRDGTDVDADRVERVFKSLNYKVTRILNITKSVLHQTLLEASQADH

SSYDSFIFVMLSHGDNNIIYANDGEVLTSYIMAFFRGDRCPSLIAKPKLFFFQACRGAAF

DKGVSTMVTDAGEDLIVHKLPIEADILVAYSTVPGFFAWRNSSSGSWFIQELCNALESDM

KNANHSDIMSLLTVVARVVAYQYRSNTGQIETDNMKQMTSTVSTLTRLFYINGLKS

>XP_018647858.1 subfamily C14A unassigned peptidase (C14 family) [Schistosoma mansoni]ANK

MQNNGHDEDIVIWTKSLHKSINSLNIHQLEDILVKLNTTKGLEFAKNVINHSINGENSVY

IAVEAGFDQAVKQLLNTDMVNIIQRNPQNNWTSLHLACYLGLATIVKSLLNCNTSNKFIR

INELYQLICMQTAETESTSQGTALHLAVYSKNIETVNQLLHDVDLISKCYESQCIENGKQ

PKEAMNSKGKLFDEFCVLTDKNGNTALHCAVMDKLYDIVNTICKYHNKSIRLPNKHGYTC

IYLAKNVLKDVKMVSLLVKYSTSSITSGNPGHHSFKQNNSIKSARVSCMNQLNKKDNKKS

HSNLIKKSENNGNYKKLKKFKYPLNHLTIDQLNKNLNSKYIITTNNNKKKWYAYSYEDEE

NHSVRTLSTKISYLSMQQKSKKSYRPFSGLSGNRVDLNEREFNRNSRQNIEQSHIENIYC

GQNRNNSIQYNGSNEPCLATCKNDIPFATTLQNNLILQPENIDQLAHQKKKESEDKNNRS

GCDIDEKSLENVFGSFNFLVKLLRNLSAGEIQAQLEHLGKKTNHHEYDCLVICLMSHGTI

GRIYGVDGNSLSIHELTSIFTADNCPSLAGKPKLFFIQACRGEDYQKGYVIQKQSNEITQ

TNNDITNQEDNLIPTYADFLLSYATVAGFRAQRDPTGGTVYIQALCKRLHNGLQNESILD

IVTAVHREVSKQLYQETSFQLMKTNGELFLQIPEVRHTLTKKVCFKR

>XP_018646351.1 subfamily C14A unassigned peptidase (C14 family) [Schistosoma mansoni]CARD

MEKEHRDVIRKLRVFLVKNISNLEEILDILESSLIITENQRHSILSGVTLAEKIRRFLDV

IVRCGPLAYDRFKQALLETNQKSIYDLLNETEELKVVTGHGMSSECLSDAKDESIMIPDR

VSSHPLIPYTVTSFPRGYVLIINIEDYSRESGVPNRHGSSHDASKLQILFEDFMYSVAVI

KNLEGEKLKKVVREFACKPEHSNVHAAVLIILAHGLEHHIIASDGIHVSIDELVGCFTNK

RCPLLAGKPKLILIQACRGEERNHNGLVKRDFQDSLLSNSCEVSLDPSSWLSLPHMSDCV

IVYSTLPGFVSWRSEIEGSWYVKVFVEVTRLHGHRLHILELLTEVNNRLVSEASNYGIKQ

ISQPVTTLTKPYYLSTLKT

>VDK22955.1 unnamed protein product [Taenia asiatica]

MLLPLTTSSSTGSKASPTLRGTASPCQASPTLPKSVKPAEVTTGTGSSDGRRYLSEVELC

DPDLGYPRVANPNNRSNPRGVCLLINQRDFDPAKTGQERRDGTDVDADNIERTFIRCGYA

VNRATNLTLRKMEFLLDDVRSQNHSKYDSFACVILSHGAEGIVYASDGTINVDRLIGYFR

SDRCPTLAGKPKMFFIQACRGCKFDKGVTLSTDASSDCVLVTKLPIEADVFVANSTFPGY

YAWRNSQAGSWFIQELCKVIKAAQESGRHHDVASLLTVVARKVALLYESNTGQSDSHGSK

QMVSINSTLTRKAFIV

>VDK23326.1 unnamed protein product [Taenia asiatica] ANK

MPQTTLRNIKRLPSSSSTSTSATDAGILLQLLRRELLLALESMDGAVFKMKLQWILENAG

EFVDLVLMSGLDLEGKTALHIAVGNRFEEAVKLLVSTGKVDLLATAGDSGWSALHTAVDL

GDVKLTQALLPIHNAVITPHRILSSLKQPTASPYAVTALHMAVRRNSASCAVVLLHTANA

LLDGHLGGWTSSEEEEDLASVDAIQQKNHFHGQAGKGSQKLIKLKDRNGDTPLHWAVKTN

NYAMARLLCSYGARPRHSHNCQGQTPFALAKYVLRNTKMTKLLASFSPTFNTNTTDNFEN

DNNETESREYEEILPFESPSPGNSTLQRVPKASEVDGSCRKLERTLVAVEMPTPPTGTVI

ATAIPAKESPSGEASNGGSISARPYLHREVKSTIPQVCHLLPGPLDLKVESDTLETSRPR

SYSLKVERRELDMLRQRVPSLLPRPQLKRAVVHRPRPVSLKARRKASLMDPVEYYPMASS

RRGICVVINNMTYWHPKFQNRGGCDKDEKRVEKVFCGLGFLVKVLRNLSSGEMRSELAFI

GNKTDHSAYDAFVAVIMTHGGLGELYGVNGDAFPVHQLTLDFTAERCPSLAGKPKLFFIQ

ACRGDDYQFGYALPASGSVEDAAPTNLGSALSNPLENGITQVEGETVEEATEDVEEKPPI

VAAKLPRKQRLVPNFADFLLSYATLAGFKAQRDPQQGSIYIQTLCHHLEVYGRSRSLLDI

VTSVHREVSEKVFREAESPDQEGTVFQQTPEVRHTLTRRVQFV

>VDK23513.1 unnamed protein product [Taenia asiatica] CARD

MNAEQRQRLSRMQVDFVKGLRDIEEITDYLLCHEILSRTQAEKILSSGPVVADQVRTLLN

VLVRCGPNAYDTFVSALSETNQSSFAEKMANLSISPRPVHPVSPPSKTQSHVTEHPTQPI

AAYPVNSAPHGYILLVNISNFEAGSGLSNRIGSSEDVVALKSLFTDLGYFVEVLLDPTAT

VLESSLSAFIKSPSHYEVDAGGLIVMSHGVQDYIYTADGKLFAINDILEAFTNKSFPALA

GKPKFILFQACRGEEKDRGCIFEPDDASFVASPLESVDAGVHKTLWKCLPYMSDYIIAYS

TLPGFVSWRSEKAGSWFIQILVDVFRKYASSLHVLDLLTEVNRRLVEESQEREFKQITQQ

SHTLTRPFYLAGIHSQSR

>VDK35885.1 unnamed protein product, partial [Taenia asiatica]

FSPCDNDTTEASPPVDSVQIAPAVKVKKSSSEKMFESSLFSCKLMPPESQSDTPSYKRQA

VESAVHHRTAASFVYKTRAPRRGTCLIISVDTFKAALRLPNRPGAGVDLRKLEDTFHFLD

FDVKTYQNPSAAQILSIVEAESSANHADADYFACVILTHGDDGGSIYGTDGPISLDQLIH

PFRGNTCPSLAGKPKMFFIQVRSLFHS

>VDL94269Schistocephalus_solidus

MCSGAHGLAAXAENSGFASSSSDPHQADEILPVDSAHVPQILRSSKTSDVCVGENPKYSP

PKILMSSAAPTQALKLNSPVPQVLPRRAPSSFIYGTRALRRGTCLIISVDTFKPALCLPG

RPGSEVDLQKLQQTFAMLDFDVKIYCNPTAANMTAIVEAESKANHADSDTFVMVLLSHGD

DGGLIYGTDGAVYLENLIKAFRGDSCPDLAGKPKLFFVQTKGATCVDV

>VDM00376Schistocephalus_solidusCARD

MEKQHSDTLRKHRVQLVSDVHSVEEILDHLFGENIISDNQRCLILAATTPAQKIRLLLDT

LVRCGPNAFSSFCSALRATKQFHILELLETSLPTEESVNIRLLSNKSSSTGIMRADEPQG

EDIMQENYSGPRKLSHYIISPYPVCTTPLGYILIVNISEFTPVCGLKPRLGSRKDTEELA

LLFQDIGYSVEVLEDPDTPTLLDRLAAFAAKPEHASVDAGGMVLMSHGLRDYIYTSEGHL

VSVEKLLSYFTNKALPGLAGKPKLLILQACRGRAQDFGTVNTDYVVTSNIFKPADGTDAP

ISNSSWNSLPYMSDCVIAYSTLPGFVSWRSEEEGSWFIQVLLDVFRRLSDKLHVIDLLAE

VNRIIVEQSQDQMCKQISQPATNLRRPFFLSTRDSLNFQAMVK

>VDL97252Schistocephalus_solidusANK

MMSTFSMQLKLLLATDKVNLLAIAGETGWTALHAAVHQEDPVAIRILLRYPVETLEATTT

TTVEAETAEDQRCEERLLRARELLWCHCDNPIGATALHMAVRRNNLECVRVLLEDANWLH

SRLAEYRPSEYGVSELEGSCEICCIQDRSGDTALHWAVSNGSHEIAWLLCSYGAQIHKPT

NHKGHEPLTICKDVLRDHEMTAILTPFKSSDPPPNAELIFKNHRPSPVLADSASNTSVRL

NASSRLRRTAVLPDPMKLPTEGGAQTIAGGPSDQVSQRRPPEQPAMFAEFCRTRPYLDRS

TKDNKQVAEFRRQATALNAVRAQKPQQLAGLPALSVEKHQGMAPRLSGLSAINLERKRPS

VCLSVPGMTCPAPLLVADQVPLQPPPPKPITLRARRRASLMDPNEYYPMASPRRGICLIM

NSMTYWHPKFQDRGGCDKDEIRIEKVFRDLGFLVKLLRNLSSGEMKAQLKHLGTATDHST

YDCFVTIIMAHGGMGEIYGVDGSPFPLHEITSYFTADKCSTLAGKPKLFFIQACRGEEYQ

LGYAPPSRNSMVGCLQPVPGLLRRAPPQSPSLTVSNPFEARSSFVEAAPVPSPSVSAQTV

TANPDAETVSREEEPAVLAANLPRRQRLVPTYADFLISCATKAGFKAQRDPKEGSIYVQS

LCRHLEQYGRLRSLLDIITGVHREIGHGNDDDDGDEDDDDNDDDGDDEDAEDDDDSGWKG

HRSDLTNLEPLNSLAKALPDASDESLWGILV

>CAH8647555.1 unnamed protein product [Schistosoma haematobium]

MASKDGYLSVSMADLHNPHLSYPIVSPNRSGKFYRGECILINQRDFHPSTNQSRRDGTDV

DADRVERVFKSLNYKVTRILNITKSVLHQTLLEASQTDHSSYDSFIFVMLSHGDNNIVYA

NDGEVLTSYIMAFFRGDRCPSLIAKPKLFFFQACRGAAFDKGVSTMVTDAGEDLIVHKLP

IEADILVAYSTVPGFFAWRNSSSGSWFIQELCNALESDMKSANHSDIMSLLTVVARVVAY

QYRSNTGHIETDNMKQMTSSVSTLTRLFYINGPPS

>XP_051070475.1 Caspase-3 [Schistosoma haematobium]

MGEIVDGVRLVTTVPKQERGQMASKDGYLSVSMADLHNPYLSYPIVSPNRSGKFYRGECI

LINQRDFHPSTNQSRRDGTDVDADRVERVFKSLNYKVTRILNITKSVLHQTLLEASQTDH

SSYDSFIFVMLSHGDNNIVYANDGEVLTSYIMAFFRGDRCPSLIAKPKLFFFQACRGAAF

DKGVSTMVTDAGEDLIVHKLPIEADILVAYSTVPGFFAWRNSSSGSWFIQELCNALESDM

KSANHSDIMSLLTVVARVVAYQYRSNTGHIETDNMKQMTSSVSTLTRLFYINGPPS

>CAH8647525.1 unnamed protein product [Schistosoma haematobium]

MGEIVDGVRLVTTVVKNVLYFERRPKQERGQMASKDGYLSVSMADLHNPHLSYPIVSPNR

SGKFYRGECILINQRDFHPSTNQSRRDGTDVDADRVERVFKSLNYKVTRILNITKSVLHQ

TLLEASQTDHSSYDSFIFVMLSHGDNNIVYANDGEVLTSYIMAFFRGDRCPSLIAKPKLF

FFQACRGAAFDKGVSTMVTDAGEDLIVHKLPIEADILVAYSTVPGFFAWRNSSSGSWFIQ

ELCNALESDMKSANHSDIMSLLTVVARVVAYQYRSNTGHIETDNMKQMTSSVSTLTRLFY

INGPPS

>CAH8555065.1 unnamed protein product [Schistosoma haematobium]

MVDDVDSSNTNLSLVEENSSTQIRSSLSCDSPQNKPNNKTWSSSLKCSVSCTSLLPVFSI

PSEKSPSSYVYRTDGAERGLCVVFSVDKFDPILCLPPRNGSLVDIQNIKRAFTSLDFRVI

VHSNPTSESLFSLIRTISTQNLRDHDCFACVILSHGDEGGLIYATDGSIPVDRIIAPFRG

DQCLDLRGKPKLFFIQACRGMALDDGVFVCDGPLSNSKSADAATTTRRIPVEADLFIAYS

VQPGYYAFRNSVNGSWFIRALSDVLLRYGNTLDLLSIMTRVNYEVAFEYESTAANPSFCC

KKQMPSFVSTLTKHVIFPPKKHRNTSKLL

>CAH8464935.1 unnamed protein product [Schistosoma haematobium]CARD

MEKEHRDVIRKLRVFLVKNISNLEEILDILESALIITENQRHSILSGTTLAEKIRRFLDI

IVRCGPSAYDKFKQALLETNQKSIYDRLNETEELKVLAGHRISSERLSDTRDESIMIPDR

VSSHPLTPYTVTSFPRGYVLIINIEDYSCESGVPNRHGSSHDASKLQILFEDFMYSVAVI

RNLEGEKLKKVVREFACKPEHSNVHAAVLIILAHGLEHHIIASDGTHVSIDELVSCFTNK

KCPLLAGKPKLILIQACRGEERNHNALVKRDFLDSLPSNFCEVSLDPSSWLSLPHMSDCV

IVYSTLPGFVSWRSEIEGSWYVKVFVEVTRLHGHRLHILELLTEVNNRLVSEASNYGIKQ

ISQPVTTLTKPYYLSTLNT

>XP_051073460.1 Caspase-8 [Schistosoma haematobium]

MLICDEKIVQFNVSEKYCSQNRSGCDTDEKSLENVFGSFNFLVKLLRNLSAGEIQAQLEH

LGKKTDHHEYDCLVICLMSHGTIGRIYGVDGNSLSIHELTSIFTADNCPSLAGKPKLFFI

QACRGEDYQKGYVIQKQSNETTQTKNDLTDQEDVDNFIQHQSSSLNLRLSVSDAKFIGFK

LDNLLHNSIPLQLNNVQLIPTYADFLLSYATVAGFRAQRDPTGGTVYIQALCKRLYKGLQ

DESILDIVTAVHREVSKQLYQETSCQLMKTNGELFLQVPEVRHTLTKKVCFKR

>XP_051073461.1 Caspase-8, variant 2 [Schistosoma haematobium]

MEMDALTTATTTTTTTTTTTTTTTTNSNNNNNNNNNDSDDYYKMNSTRRGICLIINNMMF

WHSNFQNRSGCDTDEKSLENVFGSFNFLVKLLRNLSAGEIQAQLEHLGKKTDHHEYDCLV

ICLMSHGTIGRIYGVDGNSLSIHELTSIFTADNCPSLAGKPKLFFIQACRGEDYQKGYVI

QKQSNETTQTKNDLTDQEDVDNFIQHQSSSLNLRLSVSDAKFIGFKLDNLLHNSIPLQLN

NVQLIPTYADFLLSYATVAGFRAQRDPTGGTVYIQALCKRLYKGLQDESILDIVTAVHRE

VSKQLYQETSCQLMKTNGELFLQVPEVRHTLTKKVCFKR

>CAH8442799.1 unnamed protein product [Schistosoma haematobium]ANK

MQNNEHDEDIVIWTKSLHQSINSLNIHQLEDILVKLKTTKGLEFTKNVINHLIYGENSVY

VAVEAGFDQAVIQLLNTDMVNIIQRNLQNNWTSLHLACHLGLATIVKSLLNCNTSNKFIR

TNELYQLICMQTTEIESASQGTALHLAVYSKNIETVNQLLHDVDLISKCYENQCIKNRKQ

PKEELNSKGKLFDEFCVLTDKNGNTALHCAVMDKLYDIVNTICKYHNKSIRLPNKHGYTC

IYLAKNILKDMKMLSLLIQYSNSSITSYNSGQYLFKQNNFIKNTKISNLNQLNIKKSYSN

LIRKSENNNNNKKKKLKKFNKPLNHLTIDQLNKNLNLKYGFTSNKKYYSYSYNDEDYHSV

RTLSTKFSYISMKQNNKNYYNRPFSGLSGNRVDLNERELNRNSRLNIEQNYIEDISDVQN

KNNSMQYNGSNEPCIAACKNNTAFATTLQDNPILQPENIDQLAHQQKKESEDKNICQLIT

NNISNITNKMEMDALTTATTTTTTTTTTTTTTTTTTTTTNNNNNNNNNNNNNDSDDYYKM

NSTRRGICLIINNMMFWHSDFQNRSGCDTDEKSLENVFGSFNFLVKLLRNLSAGEIQAQL

EHLGKKTDHHEYDCLVICLMSHGTIGRIYGVDGNSLSIHELTSIFTADNCPSLAGKPKLF

FIQACRGEDYQKGYVIQKQSNETTQTKNDLTDQEDVDNFIQHQSSSLNLRLSVSDAKFIG

FKLDNLLHNSIPLQLNNVQLIPTYADFLLSYATVAGFRAQRDPTGGTVYIQALCKRLYKG

LQDESILDIVTAVHREVSKQLYQETSCQLMKTNGELFLQVPEVRHTLTKKVCFKR

>CDS42948.1 caspase [Echinococcus multilocularis]

MWRKSETLNPLCYNDLQFSDRSPCDKDTTEAPPPVDSALVPPTDKMKNIFPEKMFERSLF

SCKLMPLESQSDTPAYKRQAVESAVHHRTATSFVYKTRAPRRGTCLVISVDTFKAALGLP

SRPGADVDLRKLEDTFLFLDFNVKTYQNPSATQILSIVETESSANHADADCFACVILTHG

DDAGSIYGTDGPVPLDQVIHPFRGNTCPSLAGKPKMFFIQACRGAMLDSGTLVSTDSSGG

PGDSDTLGGLRRIPIEADLLVAYAVQPGYYAFRNSVHGSWFIQALNNCLRKYGRILDLMS

IMTRVNYEVAFEFESLASTAAYSGKKQVPSIVSTLTKDVFFPPKTTAPSTCTSSLPPSRT

L

>CDI97360.1 caspase 2 [Echinococcus multilocularis]

MNSEQRQRLSRMQVDFVKGLRDIEEITDYLLCHEILSRTQAEKILSSGPIIADQVRTLLN

VLVRCGPDAYDTFVSALSETNQSSLAEKMANLNANPPPVPSVLPSAKVETHVSEQPTQSL

AAYPVNSTPRGHILLVNISNFEVGSGLSNRVGSSEDVAALKSLFTDLGYYVEVLLDPTAT

ALESNLSAFIKSPSHYNVDTGGLIIMSHGVQDYIYTADGKLFAINDILEAFTNKSFPALA

GKPKFILFQACRGEEKDRGYIYEPDNASFVASPLESVDAGVHKTLWKCLPYMSDYIIAYS

TLPGFVSWRSEKAGSWFIQILVDVFRKYASSLHVLDLLTEVNRRLVEESQEREFKQITQQ

LHTLTRPFYLAGTQSQPR

>CDS37385.1 caspase 3 apoptosis cysteine peptidase [Echinococcus multilocularis]

MENKVSSSKPTSGISVKDDAEDASFIFSSTGGKASPTPRGTTSPRQASPTLSKCVKPAEV

TTGIGSGDGRRYLSEIELCDPDLGYPRVANPNNRSNPRGVCLLINQRDFDSAKTGQERRD

GTDVDADNIERTFIRCGYAVNRATNLTLRKMEFLLDDVRSQNHSKYDSFACVILSHGAEG

IVYASDGTINVDRLIGYFRSDRCPTLAGKPKMFFIQACRGCKFDKGITLSTDASSDSVLV

TKLPIEADIFVANSTFPGYYAWRNSHAGSWFIQELCKVIKAAQESGQHHDLASLLTVVAR

KVALLYESNTGQSDSHGSKQMVSVNSTLTRKAFIV

>CDS36952.1 caspase 8 [Echinococcus multilocularis]

MPQTTSKNINRLPSSSSTSASATDTEMLLQLLRRELLLALESMDGAVFRMKLQWILENAG

EFVDLVFMSGLDFEGKTALHIAVANRFEEAVKFLVSTGKVDLLATAGDNGWSALHTAVDL

GDAKLTQLLLPTHSTTLTPHRVLSSLKQSTASPYAVTALHMAVRRNSVPCAVMLLHTANA

LPNDHSSGWTSSEEELAVDEIQQKHHFHGQAGKGSRKLNKLKDRNGDTPLHWAVKTNNYA

MARLLCSYGAHPRHSHNCQGQTPFALAKYVLRNTKMTKLLASFSPTSSTNTSVTDNFEDD

NEETRIKEDEAFMPLKPPSPTESTLRRVAKASEVDSSCRKLQQTLIRVEVPTPTPGTVTA

VTAMPVKKSPSGEASNGESTLGRPYLHREVKSTIPQGPLDMKVDSDTLEACRLRSNSLKV

GQREMDMLRQKLPGLLSRPLLKRAVVHRPRPVSLKARRKASLMDPVEYYPMASPRRGICV

VINNMTYWHPKFQNRGGCDKDEKRVEKVFCGLGFLVKVLRNLSSGEMRSELAFIGNKTDH

SAYDAFIAVIMTHGGLGELYGVNGDAFPVHQLTLDFTAERCPSLAGKPKLFFIQACRGDE

YQLGYALPATGNTEDAVTSNPGSALPNPLQIGVTQVEGEAVEEATDDVEEKPAIVAAKLP

RKQRLVPNYADFLLSYATLAGFKAQRDPQQGSIYIQTLCHHLEMHGRSRSLLDIVTSVHR

EVSEKVFREVESPDQEGTVFQQTPEVRHTLTRRVQFV

>VDL19533.1 unnamed protein product [Hymenolepis diminuta]CARD

MNNEQRQRLRRMQVDFVKNLHDVEDIIDYLLSMEILSRTQSEQILLSGPTIPDKVRSLLS

TLVRCGPNAYDAFIDALRKTNQFSLADKMESLVVDSPSFTTYSSSCHVRNPQKKPIGSTV

NQYLTAYPVNSNPRGYILLVNIANFEPGCGLSNRFGSSEDVADLKSLFSGMGYVVEVLLD

PNSTKLESALISFIKNPVHYSVDAVGLIIMSHGIQDYIYTSDGKLFPINDILESFTNKSF

PAMAGKPKFILFQACRGDEKDRGYIFEPDAAILNQQPSHLENVDGTITVNKTNWKCLPSM

SDYIIAYSTLPGFVSWRSEKEGSWFIQILVEVFQKFASTLHVLDLLTEVNRRLVEESQDR

EFKQITQQQHTLTRPFYLSTVNFKP

>VDL63263.1 unnamed protein product [Hymenolepis diminuta]

MRREKEAINPLVYNDLEFTDRSVCDQDVSEELPPVDSAQIGPAAKIQKISSGCMYENSLF

SCKLMSFDSNSPHSSAPFKRQAVEPTSSHRSAASFVYKTRAPRRGTCLIISVDTFKPALC

LPSRPGADVDLVKLEDTFHFLDFDVKKYQNPTAANILSIIEAESSASHEDADCFACVILT

HGDDSGSIYGTDGPVHLDQLIHPFRGNTCPSLAGKPKLFFIQACRGTLLDSGTTVSTDSS

SGTSDSGTLGSLRRIPIEADLLIAYAVQPGMCVSFHLTAI

>VDL11723.1 unnamed protein product [Hymenolepis diminuta]

MPEPRIENVSRRLSSVNAESAMLLQMLRRELLLALESKDPIVFQIKLNWILENSREHVDL

VLRSGLDSEGKTALHIAVETRFIEAVKFLLSTGKVDVLATAGEGGWTALHTAVDLGDANL

TKILLPTFPNIYLPPHQVLAALRQPTASSCAVTALHMAVRRNSASCAVVLLQTANALVHG

HSGGWTTSEEEDDSAFVERVQQHRDFHGQVADGSRKLNKLKDLNGDTPLHWAVKTNNYAM

ARLLCSYGARPRHFYNNQGQTPLALAKYVLQNAKMTKLLTSFSSDSSPKTMHSMKGENEN

PPGGIDDIKNSPEDAIIPMELPPHRAVRDSGVDGTCARLQRTSIGVEMMNSSQTESIEQY

EQKQTLCGISNNAGISSRPYLHRQVKTACSQVPTDRRFDSDSLEPTARQRSNSLKVGRRE

LEMFQQKLPSLFPRQNLKRAVTHYPRPISLKARRKASLMDPADYYPMASPQRGICVVINN

MTYWHPKFQNRGGCDKDERRVEKVFRDLGFMVKVLRNLSSGEMQSELAYVGRKTDHSAYD

AFVAVIMTHGGLGELYGVNGDAFPVHRLTLDFTAEICPSLAGKPKLFFIQACRGDEYQLG

YAVPVNGTSEDEISMCQVSFSPNPLRFTTSKLESNEPEYPEEEPAIAAAKLPRKQRLIPS

FADFLLSYATLAGFKAQRDPQQGSIYIQTLCHYLELYGKSRSLLDIVTSVHREVSEKVFR

EAESTDRDGGTIFQQTPEVRHTLTRRVKFV

>VDL62354.1 unnamed protein product, partial [Hymenolepis diminuta]

METQVSSQTVPSEGDEKDTSFLFPTRNVPAPCPSPPILTKAMGSPETSMSVDSWRYITES

ELSDPNLAYPRVLTPNNRSNPRGVCLLINQRDFDQAKTGQERRDGTDVDADIIERTFIKC

GYSVNRATNLTLRKMEYLLDDVRNQNHSKYDSFVCVILSHGCEGMVFASDGTINVDRLIS

YFRSDRCPTLAGKPKMFFI

>CDS28794.1 caspase [Hymenolepis microstoma]

MWQKKEAINPLAYNDLEFTDRRCDQCVSEQLPPVDSAQIAPAAKIQRISSDSMYENSLFS

CKLMAFDPNSHSSTFFKCQSVEPTLSHRSPASFVYKTRAPRRGTCLIISVDTFKPALCLP

KRPGSDVDLVKLEDTFHFLDFDVKTYQNPTAAKLLSIVEAESSASHEDADCFACVILTHG

DDGGSIYATDGPVHLDQLIHPFRGNTCPSLAGKPKLFFIQACRGTLLDSGTTVSTDSSCG

GSDSGTLGSLRRIPIEADLLIAYAVQPGYYAFRNSVYGSWFIQALNNGLRKYGRILDIMS

ILTRVNYEVAFEFESLASTAEYTGKKQVPSVVSTLTKDVFFPPKSPPSAPGPSTSYSCPT

TP

>CDS30656.1 caspase 2 [Hymenolepis microstoma]CARD

MNNEQRQRLSRMQVDFVKNLHDVEDILDYLLSLEILSRAQSEQILLSGPTIPDKVRSLLN

TLVRCGPNAYDAFIDALLKTSQCSLVDKMKSLTVDSPSFTTYSSPSLVRNPQPKPFESAA

NQYLTAYPVNSTPRGYILLVNIANFEPGCGLSNRLGSSEDVADLESMLTDLGYSVQVLLN

LSSTQLEAALSSFVKNPVHYSVDAGGLIVMSHGVQDYIYTSDGKLFSINDILEAFTNKSF

PAMAGKPKLILFQACRGEEKDRGYIFEPDAVALNQHSSHLESVDGTIVVNKTNWKCLPSM

SDYIIAYSTLPGFVSWRSETKGSWFIQTLVGVFRKFASTLHVLDLLTEVNRRLVEESQDR

EFKQITQQQHTLTRPFYLSTVNLKL

>CDS31953.1 caspase 3 apoptosis cysteine peptidase [Hymenolepis microstoma]

METQLSPSQTMSSHGDEGDASFQFPTHNVPTICPNPPTLSKSIGSPEASLNVDSWRYIAE

SELSDPDLAYPRVLNPNNRSNPRGVCLLINQRDFDQAKTGQERRDGTDIDADIVERTFIK

CGYAVNRATNLTLRKMEYLLDDVRNQNHSKYDSFVCVLLSHGCEGMVFASDGTINVDRLI

SYFRSDRCPTLAGKPKMFFIQACRGSKFDRGFALSTDASSEGMLITKMPIEADVFVANST

FPGYYAWRNSHTGSWFIQELCKVIKADQDLGKSHDVAALLTVVARKVAILYESNTGQADS

HASKQMVSVNSTLTRKAYLF

>CDS29941.1 caspase 8 [Hymenolepis microstoma]

MPEPKTRNASRRHSSVNAESEMMLQMLRRELLLALESKDPAVFKMKLTWILEHAREYADL

VLRSGFDSEGKTALHIAVEIQFIEAVKLLVSTGKCDLLATTRERGWTALHTAVDLDDANL

TRILLPTFPNIYLPPHQVLTVLRQPTESSCAATALHMAVRRNSASCAIVLLQTANALVHG

HSGGWTTSEDEDDFVFAEKVQQRRDFHGQAEDGSRKLNKLKDANGDTPLHWAVKTNNYTM

ARLLCSYGARPRHSHNNQGQTPLALAKYVLRNDKMTKLLTSFALTSPKPTYSMGGENENL

PDGIENIQNSPEDAIIPMEFPFHRAVRNSGVDGTCTRLQRTSICVEMMNSTPTEGGEQCG

KEPPSCEMSNNASISGRPYFHRQVKAACSQATTDMRFNSDSLDPKVRHRSNSLKVGRREL

EILRQQIPSLFPRQHLKRAVTYHPRLSSLRSRRKISLMDPADYYPMASPQRGICVVINNM

TYWHPKFQNRGGCDKDERRAEKVFCDLGFMVKVFRNLSSGEMQSELAYVGRKTDHSSYDA

FVAVIMTHGGLGELYGVNGDAFPVHRLTLDFTAEKCPSLAGKPKLFFIQACRGDEYQLGY

AVPVNGASEGELSMHQVSSSLNPLQFRPSKLNSDEPDDLEEEPAIAAAKLPKKQRLVPSF

ADFLLSYATLAGFKAQRDPQQGSIYIQTLCHYLELYGKSRSLLDIVTSVHREVSEKVFRE

AESTDRDGGTIFQQTPEVRHTLTRRVQFV

>VDD79867.1 unnamed protein product [Mesocestoides corti]

MAVNVRQCSSDVFIENAHFPGKPELHQSQEFTVFHELPTVDPLLSHHSTASFVYKTRAPR

RGTCLIISVDTFKPALCLPCRPGADVDLKNLEDTFQSLDFDVKSYQNPTAAMITAIIEAE

STASHADADCFACVILTHGDEGGSVYGTDGPIHLDQLIHPFRGDTCPGLAGKPKMFFIQA

CRGTLLDSGTIVSTDSSFGSGDSGTLGSLRRIPVEADLLVAYAVQPGKSTNSIYGSWFIQ

ALNNCLRKYGRILDMMSILTRVNYEVAYEFESLATTAEYSGKKQVPSIVSTLTKDIFFPP

KK

>VDD79553.1 unnamed protein product [Mesocestoides corti]

MNPEHVQHLSRMQVDLVRDLNNIEAITDYLVSEQILSRTQAEQILSQISTSDKTRSLLNT

IVRCGPDAYQAFVSALSKTDQLSLVEKLATVSVNAQSLPPPPQLKARINDIGRRPQSLAV

YPVTSNPRGYILLVNVSKFKVGCGLNERVGSLEDVSALQALFSDIGYCVEVLLDPDFRSL

DSSLKAFIRSPNHNKVDAGGLIVMSHGLQDYIYTSDGKLFAINDILNVFTNKSLPALAGK

PKFILFQACRGEEKDRGCMLTSDVTRGIPSPIEFVDAPTKNASWKCLPYMSDCVIAYSTL

PGFVSWRCEKQGSWFIQVLVDVFRKYASTFHVLELLTEVNRRMVEESQHREFKQITQPSN

TLTRAFYLSTSHFSSI

>VDD75766.1 unnamed protein product [Mesocestoides corti]

METEIPPSQLMADMSLKGDDGDARFISSSSPGSSSPTLCGIAASRASPTLSKSAASKEEA

GNKRYLSEEELNDFDLAYPLVPNPHHRKNPRGICLLINQRDFDLAKTGQERRDGTDVDAD

MVERTFIRCGYAVERATNLTLRKMELLLDDVSSQNHSIYDSFVCVVLSHGSDGVIYAADG

IIPADKFVSYFRSDRCPSLAGKPKIFFIQACRGSRFDPGVTMSTDAGEGGVLITKLPVEA

DIFVANSTAPGKFRYLQPICYFAWRNSQTGSWFIQELCAVIKAAKESGKSHDVASLLTVV

ARKVALLYESNTGQRNSHGMKQMVTVNSTLTRKALLV

>VDD77147.1 unnamed protein product [Mesocestoides corti]

MARLLCSYGARPRQSHNSQGQTPLVLSKYVLRNSKMTKLLSSFSPTSSPQSKRSTVSTSE

DIDAIQNTEEDAVILPVLPESSLNLTAKSSTVDSGAIATRLRRTGVVAAMATKVAALSPR

NDAIETMSTSSLPLSSVVDNETIGVSPRPYLHREAKVAFTEVDLAVDPDTLEPPRPRANS

LKVCRRELEVLRRKLPAFLTRSSLQKAMVHQPRPVPLRARRKASTMDPSEYYPMASPKRG

ICVIINNMTYWHPQFRNRGGCDKDEKRVEKVFGDLGFLVKVLRNLSSGEMRSELKFIGRK

TDHTAYDCFVAVVMAHGGLGELYGVNGDAFPVHQLTFDFTAEKCPSLAGKPKLFFIQACR

GDEYQLGYALPVQSNAEEPPQVVTNPPRVESASNPLYISSGKDDEPAIAAAKLPRKQRLV

PSFADFLLSYATLAGFKAQRDAKQGSMYIQTLCHHLEVYGRVRSLLDIVTSVHREVSERV

FREADSPEDEGTIFQQTPEVRHTLTRRVQRGRSSSSSDSRSNDGLQNGISKKERLEEQRK

KEEKKAEEERLTRQQWAESKLLEEEVARRLERYLEAELASLLKEREAEFNAEVERRVAIE

KEALERRRKEEAERKAREEEEMRLRKEEEERQLKELNERLTRQREEELSELRRKEEEIRQ

KQYEEELKKLEEKQRLEAEEQRRIKKEQEMILNRKRVRPKLSFSLKK

>AGY56134.1 caspase-8 [Schmidtea mediterranea]

MCLSNFHNFYKYDLNDDDTECLKFLLSDFLRNWDIFQNDSGRRIQDFVFLLLINGFSENE

TLNIIIESLFYIPRLDILYECSVPIDSYKTSPFGKLICSKRLFLFKIFNEFEERDFKTLL

NNFCRAYCKSFNLIINEQSILKKLNMILKLCIKDPDIFNHFQLYFLNYGNNRTEKLVESF

KDFEELQISPSEQLWEKITEDEKHCLDENYYLALEKGDEVSTPNFVYSAEGRVKMLKYKH

DEFYPYPESDRVKSVCLIINIKNYEHYPMKGSERDVANIYKTFGKLHFTIQKYENLNKNE

MISVAKTYSKIEASKSGSMLVCFILAHGCKDCIYDINEELVSIREFSLIFNAKNCEAYKN

KPKLFFIQACQGDRFHEGIPESGIQSSQISLTQENLHLLKPPIKQYKNSDIIPEEADFIY

GMSTVADYVAFRSTKKGSWYISELCKRVNKEAQTNDIISILTNVIHDVSMKTHMNACQLP

SVMFTSRKKCFLTKQII

>AEX93483.1 caspase 2 [Schmidtea mediterranea]

MHPNHKNLLKVNNFYLLNHLEISNDPNPISILNALGILKENHVADINSQSSNIKKVDKFLSILPKRGSKA

FDKFMEALVQSQQEYIVDKLNMCSGVPVSEVPVPFVDTTDNLIDNRSLNKTNRLMLLMNNYNFGNDEHNR

KGSEKDVEQLKSFFTEIGYKLFDDKSYEDLTSTEMNEIIKEFSLYIAKNSINSAVIIIMSHGKNKRISGI

DGHFVDLQHVYKILSCGTEQIFKLIIFSCCRGETAALASMLVMESISHSDSLSINDEIQKDRYDEFAWQV

GNYIPKNAVVLYSTQENYQSHRETTGTVFIQALLSIFKTHYKTKPLREMVMMIRSEMKRLYPKYDQWPDI

KENYFANFYL

>PAA50311[Macrostomum_lignano]

MKQSQDKAMIKLLVERKSNMAPQPGTSRQAAARATRIDNAAQKQQSMAGRDLPNQGRFFM

TESHSNKPDRPNLTPSRQLWQQHPQTRHQQSVIPEEPTKHPQAAQHPQYHQQKQQQQQQQ

QHRLSNANYSRQRDTKSAGVQATPAESDKEDDDEEDAYSMDSKIRGLCIIVNNQDYFNSD

DFKERAGTDLDERCLERLFKSLRFHVKIFKNLDAKDIYWNLSNYAKSDHSGYDCYVCCIL

SHGTLGGIYGVDGRVQKLSDLTVMLSSTNCPSLAGKPKLFFIQACRGRCLQRGAHHFTGD

RDPAMIPMANHETDALSADPDQEYILPDHGDFLIGYATVAQHIAQRHREQGSVYVQTLCD

VIERGRHKEDLLSLLTKVKYRVSKHVFHNEDGSVFVQMPEVKETLRKKIYFK

>PAA88208[Macrostomum_lignano]

MSSPVESAKRDLARALRSEEPHRVKAILRSPVADLAWLDKPLDRDGSCALHIGSYCGQEV

GVHALLEFGCDPNVTNQNGETPLHVAAEQRFEHMIDLLVDAEGIRLDAARKLDRRTALHI

AASHGCCDGVRRLLEAGAGIDVRDRDGNTPVFLAVQNNKHRVVHELVSQGANLKLKNNGL

QTCLAFAMKQSQDKAMIKLLVERKSTMAPQPGTSRQAAVRATRIDNAAQKQQSMAGRDLP

NQGRFFMTESHSNKPDRPNLTPSRQLWQQHPQTRHQQSVIPEEPTKHPQAAQHPQYHQQK

QQQQQQQQHRLSNANYSRQRDTKSAGVQATPAESDKEDDDEEDAYSMDSKIRGLCIIVNN

QDYFNSDDFKERAGTDLDERRLERLFKSLRFHVKIFKNLDAKDIYWNLSNYAKSDHSGYD

CYVCCILSHGTLGGIYGVDGRVQKLSDLTVMLSSTNCPSLAGKPKLFFIQACRGRCLQRG

AHHFTGDRDPAMIPMANHETDALSADPDQEYILPDHGDFLIGYATVAQHIAQRHREQGSV

YVQTLCDVIERGRHKEDLLSLLTKVKYRVSKHVFHNEDGSVFVQMPEVKETLRKKIYFK

>PAA85491[Macrostomum_lignano]

MSTREQNLPETLERPEVSPVVLDDLENNFPEIGKVEETAGVLPLIGINFDNFEARILEKH

EADRFETAYDTRGNPRGLCLIINNKSFAGKLEDRTGTDVDRDNLSNLFNKLLYKVEIVNN

QTAEGMKETCRRFILDPEHERYSSCVLAVLSHGTNGKILGSDSKPVDEYKDILSLFNHSD

CKLTGKPKFVIIQACRIELDRTDKGTQHPMVDQEVSDGPVVEGDENFDLVDHLRRLNYGD

AQRVENRLTPDFADLLIAHSTLPGYFSWRHTKQGSWFIQAIVQVFSIHFHSEHVADMMVQ

VNAVISKAFGRNIRCQMPSPTNHLRKKFYFFPSK

>PAA49678[Macrostomum_lignano]

MSTREQNLPETLEQPEVSPVVLDDDLENNFPEIDKVEETDGLLPVGINFDNFEARILEKH

EADRFETAYDTRGNPRGLCLIINNKSFDGKLDDRTGTDVDRDNLSNLFNKLLYKVEIVNN

QTAEEMKEACRSFILNPEHEQYTSCVLAVLSHGTNGKILGRDSKPVDEYKDILSLFNHSD

CKLTGKPKFVIIQACRIELDRTDKDTQHPMVGQEVSDGPVVEGDENFDLVDHLQRLNYGD

AQKMENRLPPDYADLLIAHSTLPGFISWRNTKQGSWFIQAIVQIFSRNFHSEHVADMMVQ

VNLLVSKAFRGENKKNKNQACQMPSPTNHLRKKFYFFPSK

>PAA61954[Macrostomum_lignano]

IMSDQAAVEQPNKGDATDAGSSIFKRSSQVQPAFPQPLPTAALEDIDAEYQMGNPSRGFC

LIFNHVNFLSSTHQPTREGSDVDASSIENSFRRLGFDVRTFRDLTASNLLDVLREFRLKN

HSNNDCFAVVLLSHGYEGHFYAKDRDLPIDFVVRQFKPDACKTLAGKPKLFFIQACRGDN

FDRGVEVTDGEDDTLDALPPAGVGVGSGAGISGPFRVPLESDFFFFYSTPPGYFSWRNSH

RGSWFGQQLTALLDRHGDRMELQHLMLMVQRLVSSAYASDTGDPKSTGMKQMPVTVSQLR

KLLYFRPKQH

>PAA87421[Macrostomum_lignano]

AMTAPLPNTPLDPALEYSMDHKKRGFFIIINQVKFESRTGQKERTGSDLDAAKLEETFKI

LGFNVLPFRNLKLADLRRLLAEYGQKDHSDNDCFGLAMMSHGEQDIIYCTDGSLKVDELT

KPFRADNCASLIGKPKLFFFQACRGFTFDDGVEKTDADLPEDEDLEEERVQRLPIEADNF

FFYSTPPGYFSWKNALYGSWFIQELCRVLDQWGTRHEFSQLMVSVQREVASNRHESLTDR

AATSRKKQMPVSVSQLTKLLYFRPKLPPQLPRVAPSSES

>PAA86298[Macrostomum_lignano]

MTAPLPNTPLDPALEYSMDHKKRGFFIIINQVKFESRTGQKERTGSDLDAAKLEETFKIL

GFNVLPFRNLKLADLRRLLAEYGQKDHSDNDCFGLAMMSHGEQDIIYCTDGSLKVDELTK

PFRADNCASLIGKPKLFFFQACRGFTFDDGVEKTDADLPEDEDLEEERVQRLPIEADNFF

FYSTPPGYFSWKNALYGSWFIQELCRVLDQWGTRHEFSQLMVSVQREVASNRHESLTDRA

ATSRKKQMPVSVSQLTKLLYFRPKLPPQLPRVAPSSES

>PAA76232[Macrostomum_lignano]

QECTMSDSVSQQDADQVDAGLFRRGGGRRDPPPPLEPPTGGNPAPKLAEYQMNHPKRGLC

LIFNQVTFDPRTGQSTRIGSDVDADNLERSFKQLGFSIMPFRDLSCEKIVSVIKETARKD

HSNYDCVAVVLLSHGENGAIYGSDGKIKMEKLFRLLRPDYCPTLAGKPKLFFIQACRGSE

YDKGCEVADGDNGTGRDEEDGGKLSRIPLESDFLYFYSTPPGYYSWRNDIQGSWFIQQLC

HVLQRYGDSDELGHLLLMVQQAVAGNYQSNTNDDSTSGMKEIPVSVSTLRKLVYFRRK

>PAA65253[Macrostomum_lignano]

IMSDQAAVEQPNKDDATDAGPSIFKRSSQVQPAFPQPLPTAALEDIDVEYKMDHPSRGFC

LIFNHVNFLSSTHQPTREGSDVDASNIENSFRRLGFDVRTFRDLTASSLLDVLREFRLKN

HSNNDCFAVVLLSHGYEGHFYTKDRDLPIDFVVRQFKPDACKTLAGKPKLFFIQACRGDN

FDRGVEVTDGEGDTLDALPPAGVGVGSGAGISGPLRVPLESDFFFFYSTPPGYFSWRNSQ

RGSWFGQQLTALLDRHGDRMELQHLMLMVQRLVSSAYASDTGEPESTGMKQMPVTVSQLR

KLLYFRPKQH

>PAA78189[Macrostomum_lignano]

IMSDQAAVEQPNNGDATDAGSSIFKRSSQVQPAFPQPLPTAALEDIDVEYKMDHPSRGFC

LIFNHVDFDSTRQPTREGSDVDASNIENSFRRLGFDVRTFRDLTASSLLDVLREFRLKNH

SNNDCFAVVLLSHGYEGHFYAKDRDLPIDFVVRQFKPDACKTLAGKPKLFFIQACRGDNF

DRGVEVTDGEDDTLDALPPAGVGVGSGAGISGPFRVPLESDFFFFYSTPPGYFSWRNSHR

GSWFGQQLTALLDRHGDRMELQHLMLMVQRLVSSAYASDTGDPKSTGMKQMPVTVSQLRK

LLYFRPKQH

>PAA75167[Macrostomum_lignano]

LLFSTIIVTDCVKSRQYLQEQRGPECSVLDFLAVKKLERLSEMSERSVSLELLAVDGRRF

RCRPDPRHFAPVQGRFGRIFRVRISAAETSESTADETAAVYCADKAGRKLLSCFSILAQL

KHDNIVRLIAAARDADSAASVCVLMEWVGAGSLRDLIDRSGCLTEPLIGAYAKQIGSALN

YLHSRSILHRNVKCRSALITESGHTKLAGFTACCLLEEDDPDASLDVRQFGCFLVEMATG

RTGCKEAPESSTNQLRSLCEACAEQQRPRMEQLLQLEFFQTMQSLKPLLSPSPPTPTQAL

RPVKSDRCLLGGRAEVGASAPEVLLPQRRLGESDATEELDDDDEAGARGLDKYRQTDDED

EDYFELEGTDFEELIEEDENEQDEEDAQACQIDRERLLGRGRFGSVFRADLHGMPAAAYR

TTPGCPEETLKQFRASCLILGKLRHDCIVRLVKVLSDSSGCLTVLTELVEGRNLRSILTQ

RKAGFREGDLRSYCRQIASALLYLHEQRPALVHRNVKCKSVLLQPNSSLKLVGFSLCMRQ

QRTARPREWGSPCYLAPEVLHSAGAEFTTKSDVWAFGCLVFELATGRPPNAEQSRNFDRL

HASICVGSMPSLPIDSSAALGELYRLCTQRRRKRRPGVQHLLALPFFHRSSRPLLERRNS

SCQRRKPSLVRQRTTSDSQVVEELQLQQQQQQSKQQQEQQQEQQKQQQQPSLRFPLVTPR

LQQLKPKQQQQDLEAFESVKDDVLKGRSEKLHRSSSEGGPVMQRAQTAAKQQQQQPQPEP

LFEPINSLPSSNIYPHCQRPHGYCLLINVERFANPSEFPTRNGSLVDVERVSRLLQRLGY

RVRSIADPTAAGICKAAMRIALKNHENYDSFVCFLMSHGTNNQIVGSDGQYISLGRIASF

FDSDVCAGLAGKPKLFFIQACRGERQQRNFATTRAVAVGASADQVATAITDDDDEAEEPN

TGGVLFAPKGVDFFFGYATSEKYIAYRDSQSGSAYVRVLCRVVEAYGAKEDLLQLHTRVT

ALISRNPQENQFGSKLFQVPDMRSKLTKQFYFLLGSLKKVSSDLTS

>PAA90669[Macrostomum_lignano]

AMTAPLPNTPLDPALEYSMDHKKRGFFIIINQVKFEPRTGQKERTGSDLDAAKLEETFKI

LGFNVLPFRNLKLADLRRLLAEYGQKDHSDNDCFGLAMMSHGEQDIIYCTDGPLKVDELT

KPFRADNCASLIGKPKLFFFQACRGFTFDDGVEKTDADLPEDEDLEEERVQRLPIEADNF

FFYSSPPGYFSWKNALYGSWFIQELCRVLDQWGTRHEFSQLMVSVQREVASNRHESLTDR

AATSRKKQMPVSVSQLTKLLYFRPKLPPQLPRVAPSSES

>PAA54824[Macrostomum_lignano]

MSERSVSLELLAVDGRRFRCRPDPRHFAPVQGRFGRIFRVRISAAETSESTADETAAVYC

ADKAGRKLLSCFSILAQLKHDNIVRLIAAARDADSAASVCVLMEWVGAGSLRDLIDRSGC

LTEPLIGAHGKQIGSALHYLHSRRILHRNVKCRSALITESGHTKLAGFTACCLLEEDELD

ASLDVRQFGCFLVEMATGRTGCKEAPESSTNQLRSLCEACAEQQRPRMEQLLQLEFFQTM

QSLKPLLSPSPPTPTQALRPVKSDRCLLGGRAEVGASAPEVLLQQRRLGESDATEELDDD

DEAGARGLDKYRKTDDEDEDYFELEGTDFEELIEEDENEQDEEDAQACQIDRERLLGRGR

FGSVFRADLHGMPAAAYRTTPGCPEETLKQFRASCLILRKLRHDCIVRLVKVLSDSSGCL

TVLTELVEGRNLRSILTQRKAGFREGDLRSYCRQIASALLYLHEQRPALVHRNVKCKSVL

LQPNSSLKLVGFSLCMRQQRTARPREWGSPCYLAPEVLHSAGAEFTTKSDVWAFGCLVFE

LASGRPPNAEQSRNFDRLHASICVGSMPSLPIDSSAALGELYRLCTQRRRKRRPGVQHLL

ALPFFHRSNRPLLERRNSSCQRRKPSLVRQRTTSDSQVVEELQLQQQQQQSKQQQEQQQE

QQKQQKQPSLRFPLVTPRLQQLKPKQQQQQQQDLEAFESVKDDVWKGRSEKLHRSSSEGG

PVMQRAQTAAKQQQQQPQSEPLFEPINSLPSSNIYPHCQRPHGYCLLINVERFANPSEFP

TRNGSLVDVERVSRLLQRLGYRVRSIADPTAAGICKAAMRIALKNHENYDSFVCFLMSHG

TNNQIVGSDGQYISLGRIASFFDSDVCAGLAGKPKLFFIQACRGERQQRNFATTRAVAVG

ASADQVATAITDDDDEAEEPNTGGVLFAPKGVDFFFGYATSEKYIAYRDSQSGSAYIRVL

CRVVEAYGAKEDLLQLHTRVTALISRNPQENQFGSKLFQVPDMRSKLTKQFYFLLGSLKK

VYSDLTS

>CDS17893[Echinococcus_granulosus]

MWRKGETLNPLCYNDLQFSDRSPCDKDTTEASPPVDSALVAPTDKMKNIFPEKMFERSLF

SCKLMPLESQSDTPAYKRQAVESSAHHRAATSFVYKTRAPRRGTCLVISVDTFKAALGLP

SRPGADVDLKKLEDTFLFLDFDVKTYQNPSATQILSIVETESSANHADADCFACVILTHG

DDAGSIYGTDGPVPLDQVIHPFRGNTCPSLAGKPKMFFIQACRGAMLDSGTLVSTDSSGG

PGDSDTLGGLRRIPIEADLLVAYAVQPGKFTGYYAFRNSVHGSWFIQALNNCLRKYGRIL

DLMSIMTRVNYEVAFEFESLASTAAYSGKKQVPSIVSTLTKDVFFPPKTTAPSTCTSSLP

PSRTP

>CDS23854[Echinococcus_granulosus]

MPQTTSKNINRLPSSSSTSTSATDTEMLLQLLRRELLLALESMDGAVFRMKLQWILENAG

EFVDLVFMSGLDFEGKTALHIAVANRFEEAVKLLVSTAKVDLLATAGDNGWSALHTAVDL

GDAKLTQLLLPTHNTTLTPHHVLSSLKQATASPYAVTALHMAVRRNSVPCAVMLLHTANA

LLNDHSSGWTSSEEELAVDEIQQKHHFHGQAGKGSRKLNKLKDRNGDTPLHWAVKTNNYA

MARLLCSYGAHPRHSHNCQGQTPFALAKYVLRNTKMTKLLASFSPTSSTNTSVTDNFEDD

NEETRIKEDEAIMPLKPPSPTESTLRRVVKASEVDGSCQKLQQTLIRVEVPTPTPGTVTA

VTAMPVKKSPSGEASNGGSILGRPYLHREVKSTIPQGSLDMKVDSDTLEACRLRSNSLKV

GRREMDMLRQKLPGLLSRPLLKRAVVHRPRPVSLKARRKASLMDPVEYYPMASPRRGICV

VINNMTYWHPKFQNRGGCDKDEKRVEKVFCGLGFLVKVLRNLSSGEMRSELAFIGNKTDH

SAYDAFIAVIMTHGGLGELYGVNGDAFPVHQLTLDFTAERCPSLAGKPKLFFIQACRGDE

YQLGYALPATGITEDAVTSNPGSALPNPLQIGDTQVEGEAVEEATDDVEEKPAIVAAKLP

RKQRLVPNYADFLLSYATLAGFKAQRDPQKGSIYIQTLCHHLEMHGRSRSLLDIVTSVHR

EVSEKVFREVESPDQEGTVFQQTPEVRHTLTRRVQFV

>XP_024351559[Echinococcus_granulosus]

MDRCDGFLFLTASSSTGGKASPTPRGTTSPCQASPTLSKSVKPAEVTTGIGSGDRRRYLS

EIELCDPDLGYPRVANPTNRSNPRGVCLLINQRDFDSAKTGQERRDGTDVDADNIERTFI

RCGYAVNRATNLTLRKMEFLLDDVRSQNHSKYDSFACVILSHGAEGIVYASDGTINVDRL

IGYFRSDRCPTLAGKPKMFFIQACRGCKFDKGVTLSTDASSDSVLVTKLPIEADIFVANS

TFPGYYAWRNSHAGSWFIQELCKVIKAAQESGRHHDVASLLTVVARKVALLYESNTGQSD

SHGSKQMVSVNSTLTRKAFIV

>CDS19333[Echinococcus_granulosus]

MENKVLSSKPTADIPVKDDAEDASFIFSSTGGKASPTPRGTTSPCQASPTLSKSVKPAEV

TTGIGSGDRRRYLSEIELCDPDLGYPRVANPTNRSNPRGVCLLINQRDFDSAKTGQERRD

GTDVDADNIERTFIRCGYAVNRATNLTLRKMEFLLDDVRSQNHSKYDSFACVILSHGAEG

IVYASDGTINVDRLIGYFRSDRCPTLAGKPKMFFIQACRGCKFDKGVTLSTDASSDSVLV

TKLPIEADIFVANSTFPGYYAWRNSHAGSWFIQELCKVIKAAQESGRHHDVASLLTVVAR

KVALLYESNTGQSDSHGSKQMVSVNSTLTRKAFIV

>PAA59014[Macrostomum_lignano]

LMDDIHRNALLKARCVLVSQLVADNKLWGLLLERKVFTKPMLERILGQGTPHERRQQLMD

DLPRRGPDAFGQFLGILRDTRQQDLADYLDGMDRHLRHGHPLPSEPRPQPRPPSGPSQPA

QPSQPQDDPSRRPSVDRSVSRIGVTNQPNIPDDVLSVQEVLEHHEDGLEELNFHTCGIID

STELMPQPVTQQHLTEKSKNDWEFTYQLRSNPKGLLLIISNRDFVKKSKRTGTEVDCENL

TRVFKNLGYELYRNQVFNDKTAAEMKSLCRDFILDPRHESLDSCVLCILSHGDQGGVIYG

RDDLPATEDELFEHFNSDRCKLHNKPKLVIIQACRGSMKDKGHIFDQTDSASGDKTDQTT

VQKIIHSLQEENYSDTMDLEKRVPLYADIVFAHSTTPGFVSWRHELRGSWFIQTIAKTFA

AYAHEEHVFDMLTTVNRIVTETFGGRLGYVSMPHPISRLRKKFYFFPK

>RTG87993.1 caspase 2 [Schistosoma bovis]CARD

MEKEHRDVIRKLRVFLVKNISNLEEILDILESALIITENQRHSILSGTTLAEKIRRFLDI

IVRCGPSAYDKFKQALLETNQKSIYDRLNETEELKVLAGHGISSERLSDTRDESIMIPDR

VSSHPLTPYTVTSFPRGYVLIINIEDYSCESGVPNRHGSSHDASKLQILFEDFMYSVAVI

KNLEGEKLKKVVREFACKPEHSNVHAAVLIILAHGLEHHIIASDGTHVSIDELVSCFTNK

KCPLLAGKPKLILIQACRGEERNHNALVKRDFLDSLPSNFCEVSLDPSSWLSLPHMSDCV

IVYSTLPGFVSWRSEIEGSWRNYKENLEVTIYNNLVRSNV

>CAH8479610.1 unnamed protein product [Schistosoma bovis]

MQYNGSNEPCIAACKNNTAFATTLQDNPILQPENIDQLAHQQKKESEDKNICQLITNNIS

NITNKMEMDAITTATTTTTTTTTTTTTTTNNNNNNNNNNNNNNNEDDSDDYYKMNSTRRG

ICLIINNMMFWHSDFQNRSGCDIDEKSLENVFGSFNFLVKLLRNLSAGEIQAQLEHLGKK

TDHHEYDCLVICLMSHGTIGRIYGVDGNSLSIHELTSIFTADNCPSLAGKPKLFFIQACR

GEDYQKGYVIQKQSNETTQTKNDLTDQEDVDVDNFIQHQSSSLNLRLSVSDAKFIGFKLD

NLLHNSIPLHQLNNVQLIPTYADFLLSYATVAGFRAQRDPTGGTVYIQALCKRLYKGLQN

ESILDIVTAVHREVSKQLYQETSSQLMKTNGELFLQIPEVRHTLTKKVCFKR

>RTG88383.1 caspase 8, partial [Schistosoma bovis]ANK

SINPQGITMQNNEHDEDIVVWTKSLHQSINSLNIHQLEDILVKLKTTKGLEFTKNVINHL

IDGENSVYVAVEAGFDQAVIQLLNTDMVNIIQRNLQNNWTSLHLACHLGLATIVKSLLNC

NTSNKFIRANELYQLICMQTTEIESASQGTALHLAVYSKNIETVNQLLHDVDLISKCYEN

QCIKNRKQPKEELNSKGKLFDEFCVLTDKNGNTALHCAVMDKLYDIVNTICKYHNKSIRL

PNKHGYTCIYLAKNILKDMKMLSLLIQYSNSSITSYNSGQYLFKQNNFIKNTKISNFNQL

NIKKSYSNLIRKSENNNNNKKKKLKKFNKPLNHLTIDQLNKNLNLNYGFTNNKKYYSYSY

NDEDYHSVRTLSTKFSYISMKQNNKNYYNRPFSGLSGNRVDLNERELNRNSRLNIEQNYI

EDIYDVQNKNNSMQYNGSSEPCIAACKNNTAFATTLQDNPILQPENIDQLAHQQKKESED

KNICQLITNNISNITNKMEMDALTTATTTTTTTTTTTTTTTTTNNNNNNNNNNNNNNNED

DSDDYYKMNSTRRGICLIINNMMFWHSDFQNRSGCDIDEKSLENVFGSFKFLVKLLRNLS

AGEIQAQLEHLGKKTDHHEYDCLVICLMSHGTIGRIYGVDGNSLSIHELTSIFTADNCPS

LAGKPKLFFIQACRGEDYQKGYVIQKQSNETTQTKNDLTDQEDVDVDNFIQHQSSSLNLR

LSVSDAKFIGFKLDNLLHNSIPLHQLNNVQLIPTYADFLLSYATVAGFRAQRDPTGGTVY

IQALCKRLYKGLQNESILDIVTAVHREVSKQLYQETSSQLMKTNGELFLQIPEVRHTLTK

KVCFKR

>RTG86123.1 caspase 3, partial [Schistosoma bovis]

VKNVLYFERRPKQERGQMASKDGYLSVSMADLHNPHLSYPIVSPNRSGKFYRGECILINQ

RDFHPSTNQSRRDGTDVDADRVERVFKSLNYKVTRILNITKSVLHQTLLEASQTDHSSYD

SFIFVMLSHGDNNIVYANDGEVLTSYIMAFFRGDRCPSLIAKPKLFFFQACRGAAFDKGV

STMVTDAGEDLIVHKLPIEADILVAYSTVPGFFAWRNSSSGSWFIQELCNALESDMKSAN

HSDIMSLLTVVARVVAYQYRSNTGQIETDNMKQMTSSVSTLTRLFYINGPPS

>CAH8632254.1 unnamed protein product [Schistosoma bovis]

MGEIVDGVRLMTTVPKQERGQMASKDGYLSVSMADLHNPHLSYPIVSPNRSGKFYRGECI

LINQRDFHPSTNQSRRDGTDVDADRVERVFKSLNYKVTRILNITKSVLHQTLLEASQTDH

SSYDSFIFVMLSHGDNNIVYANDGEVLTSYIMAFFRGDRCPSLIAKPKLFFFQACRGAAF

DKGVSTMVTDAGEDLIVHKLPIEADILVAYSTVPGFFAWRNSSSGSWFIQELCNALESDM

KSANHSDIMSLLTVVARVVAYQYRSNTGQIETDNMKQMTSSVSTLTRLFYINGPPS

>CAH8637253.1 unnamed protein product [Schistosoma bovis]

MGEIVDGVRLMTTVVKNVLYFERRPKQERGQMASKDGYLSVSMADLHNPHLSYPIVSPNR

SGKFYRGECILINQRDFHPSTNQSRRDGTDVDADRVERVFKSLNYKVTRILNITKSVLHQ

TLLEASQTDHSSYDSFIFVMLSHGDNNIVYANDGEVLTSYIMAFFRGDRCPSLIAKPKLF

FFQACRGAAFDKGVSTMVTDAGEDLIVHKLPIEADILVAYSTVPGFFAWRNSSSGSWFIQ

ELCNALESDMKSANHSDIMSLLTVVARVVAYQYRSNTGQIETDNMKQMTSSVSTLTRLFY

INGPPS

>CAH8545811.1 unnamed protein product [Schistosoma bovis]

MVDDVDSSNTNLSLVEEHSSTQIRSSLSCDSPQNKPNNKTWSSSLKCSVSCTSLLPVFSI

PSEKSPSSYVYRTDGAERGLCVVFSVDKFDPILCLPPRNGSLVDIQNIKRAFTSLDFRVI

VHSNPTSESLFSLIRTISTQNLRDHDCFACVILSHGDEGGLIYATDGSIPVDRIIAPFRG

DQCLDLRGKPKLFFIQACRGMALDDGVFVCDGPLSNSKSADAATTTRRIPVEADLFIAYS

VQPGYYAFRNSVNGSWFIRSLSDVLLRYGNTLDLLSIMTRVNYEVAFEYESTAANPSFCG

KKQMPSFVSTLTKHVIFPPKKHRNTSKLL

>VDM32914.1 unnamed protein product [Hydatigera taeniaeformis]CARD

MNAEHRQRLGRMQVDFVKGLRDIDEITDYLLCHEILSRTQAEKILSSGPVIADQVRTLLSVLVRCGPDAYDTFVSALLET

NQSSFVEKMTNLDISPRPVPPISPPSRTDAHVTQQSTQSIAAYPVNSTPRGYILLINISKFEVGSGLSNRIGSSEDVVAL

KSLFTDLGYCVEILLDPTASTLESSLSTFIKSPCHYGVDAGGLIVMSHGVQDYIYTADGKLFAINDILEGFTNRSFPALA

GKPKFILFQACRGEEKDRGYTFNPDNASFIPSPLESVDAGIHRTLWKCLPYMSDYIIAYSTLPGFVSWRSEKAGSWFIQI

LVDVFRKYANTLHVLDLLTEVNRRLVEESQEREFKQITQQSHTLTRPFYLTGTHSQSR

>VDM18528.1 unnamed protein product [Hydatigera taeniaeformis]

MCSSLTASSSAGGKSSPTLRRTASPCHASPPLQKSVKPAEVMTGTGSSDGRRYLSEVELCDPDLGYPRVANPNNRSNPRG

VCLLINQRDFDLAKTGQERRDGTDVDADNIERTFIRCGYAVNRATNLTLRKMEFLLDDVRSQNHTKYDSFACVILSHGAE

GIVYASDGTINVDRLIGYFRSDRCPTLAGKPKMFFIQACRGCKFDKGVALSTDANSDYVLVTKLPIEADVFVANSTFPGY

YAWRNSQAGSWFVQELCKVIKAAQESGRHHDVASLLTVVARKVALLYESNTGQSDSHGSKQMVSINSTLTRKAFIV

>VDM32254.1 unnamed protein product [Hydatigera taeniaeformis]

MLTSIRLDNTVVFDYGSIEVKLLSVLLQNRGGCDKDEKLVEKVFCGLGFLVKVLRNLSSGEMRSELSFIGNKTDHSAYDA

FVAVIMTHGGLGELYGVNGDAFPVHQLTLDFTAERCPSLAGKPKLFFIQACRGDEYQFGYALPAAGSTDDVAATNSGPAF

SNLLQVGVTQVEGEVAEEAAEDVEEKLPIVAAKLPRKQRLVPNFADFLLSYATLAGFKAQRDPQQGSIYIQTLCHHLELH

GRSRSLLDIVTSVHREVSEKVFREAESPNQEGTVFQQTPEVRHTLTRRVQFV

>QDF63030.1 caspase-7 [Dugesia japonica]

MDTVDAKESDFYSTDKKKLCYIINNEEFDRTLNLSSRSGTQRDFEKLKTLFKKLNFVVENHKNCTAGEMRKNLEEISKRK

DLKDYSMFVCFILSHGDEGIIYGRDAFVPIDSMLNLFKGDKCKGLVGKPKLFFIQACRGDRFDKGVVADSVGSNNLDKIP

IEADVLIGYSTVPGYFSWRNSSNGSWYIQALCSEIEKNAETEDVVHILVRVNASVAANFSSNTGDIKSHGMIQIPSFTCM

LRKKVLLVDQN

>QAV56480.1 caspase-7 [Dugesia japonica]

MDTVDAKESDFYSTDKKKLCYIINNEEFDRGLNLSPRSGTQRDFEKLKTLFKKLNFVVENHKNCTAYEMRKNLEEISKRK

DLKDYSMFVCFILSHGDEGIIYGRDAHVSIDTMLNYFKGDKCKGLVGKPKLFFIQACRGDKFDKGVLADSVGSNNLDKIP

IEADVLIGYSTVPGYFSWRNSSNGSWYIQALCSEIEKNAKTEDVVHILVRVNASVAGNFSSNTGDTKSHGMIQIPSFTCM

LRKKVLLLDQN

>QDF63029.1 caspase-3B [Dugesia japonica]

MDNENDTIEFIDTDDLDTVDGQENNAYDMSQKKLCYIISNEEFQPHLNLSTRSGTDKDFKQLRQVFKTLKFEVEPFKNCT

CRQINENLKQLSQRKDLDGVSMMVCIILSHGDEGIIYGTDGIVELDRLFGYFKGDKCKGLIGKPKIFFIQACRGQKYDKG

VSADSIISNYIEKIPIEADFLIGYSTVPGYFSWRNGASGSWYIQALCKELIKCSKTHELVQLLVKVNNNVATNFKSKTNQ

KETDGMVQISSFVCMLRKSLWFS

>QDF63028.1 caspase-3A [Dugesia japonica]

MEDIEKTNFDDVDGTIRECYDVSKIKLCYIINNEEFYPKLNLPTRSGTDEDFKSLKQVFTKLGFKVENYKNFKKDEICSN

LRELSERKELKDVSILVCIILTHGADGIIYGIDGPIELDRIFGYFKGDKCRKLIGKPKVFLFQACRGEKFDKGVVADGFF

QTSLEKIPVEADFLIGYSTVAGYFSWRNGRNGSWYIQALCRELERSAQSMELIQLLVNVNNYVATNYFSMTKNDKTDAMV

QIPSFVCMLRKPLWLVDKN

>QDF63032.1 caspase-3C [Dugesia japonica]

MNFNNSNSRELYPMNGKKLCYIINNELFHKKSGLTRRNGTNIDFKNIKITFEKLNFEIQWNDNCRSLEMIQNLQKISLKS

DLKDYSIFVCVILSHGDEGIIYGFEDSVEINKLLAIFTEENCEALRDKPKIFLIQACKGDKCDSGAYADSIQSNNINKMF

IEPNFFIGYSTVPGYFSWRNQVVGSWFISSLCDVINSNENREDLVQLLVRVNNRVSKYRSNVNRNDMNNKLQISSFVCML

LKDVYL

>BAP74679.1 caspase-8 [Dugesia japonica]DED

MCKINFHNFYQYDLSTDDAECIQFLLSDFLLHWDIFQMDNGGRIQDFVFLLIRNGLTENETLNVIIESLYFVPRLDILYE

CSISYDSYISNPVGKLICNKRLLFFNLFNELEEKDFKSILHILWKEYVKDEPWEPKCSILQNLDKLLKMSRENDTLFDEI

KSYFSNFGNNRTKKVVDVYEENRECTICDSEQLWQKVSNDNKQSNNDKIIIAMEKGDEFSNPSFAINDDEGRMKLINYKA

DESYLVPERDSIKSFCLIFDIRDYENKPRKGSEKDILKIRKTFEKLHFTIQPYKNVTKSEMFDIAKTYANIDAKSSGSML

VCFILAHGSKDCIFDINEEKVNIRQFSLCFSAKNCEAYRNKPKVFFIQACQGDRFHEGIPETSIDLEESSQCSYLQTGNQ

PIKIEEYKKSDMIPEEADFIYGMATVSDYVAFRSTKKGSWYITELCKRINEEVKTNDVIGILTNVNGDVSRRNYKNKCQL

PSVIFTTRKKCYLTEQTSK

>UKF18831.1 caspase-8A [Dugesia japonica]

MGSYDMSKSKYCLIIGNITFIDKDSLPGVIDDANRLEKVFEKLGFEVRFGEKKKYKKNEMTTNLTASEIDRKFQNLSKDE

NLLNDISILVCIILSHGDAGIIYGSDEKAVKLDEIFGYFEGNNCKELRRKPKVFLIDACRGAKRATITSRGSSTLPLRST

PQLPNKADFLEGYATVKGYIALMTDEGFGWYMQALCNELEGLTESKDLLSILTIVNNKVATEYVKWKKYEIWLKDPKNQD

AVKKEIWKNDPKNEVGTMYKKWLDDTKIQSAINEELEGIVQMPIFKCTLRNAVHLQLKKEATTIGKNSITE

>QDF63031.1 caspase-6 [Dugesia japonica]

MNNFENILKNIYLNSTDNHICNYILCEKFEPEHSKLELIFELLKRNYPANEAFDMIICALYIINYQIHQRYCECPETIYE

KYKTKLTNEKVFYVRVYKNLPDEILITFLQRIFNISKVSITCEEYFEICSKATIDYDNKLLDSVNTFYEEIFGDKFQNSL

ELDGKPQNKRPRNSISKNNNNQQDENESIFYYAIKDKRDNLCLIFNASFNRRGSEQDVIEIKSTFENINGFKVLSNEITT

REKLLNDIKDSLKLVTEDSDLIVTFIMSHGEQGIVFDCFDESVIINDILELYSSKMVPQLSFKPKIFVFQTCRGNKKQEV

IHFNSPNVENIASDAPRKVSNDDIINGSDIFIVYSTIPGHSSMRDKYKGAYFIRKFCEVIKENSKKRNLDEICTLVNTSY

EKYAFGEQEITQLCEVRKVVMKFNIYLEKPDKLKMFLL

>ASL70034.1 caspase 8-like protein [Dugesia japonica]

MSYLNKSDFINLKYLIVSTGLVRNEDIVDLKDERDLIPKLINHLEKVPGIDNAAHILEELLNESFYYLKHYAYLHKNGVC

PLSYEASEWGINLTDESILLLRLYQNLSQHEFFFVLDNFMGVQKEIYLNLQMDFNNLNASDYLPKVYDWLHQQRNPLAEK

LEILIKDRFGDFETKPESVSLNPSAAGFQGVCLIINNENFQSMKYKKREGTQVDEDAIFDTFSKLNFNVLVYKDVKRENI

NKTLDHAKSLVEMKHRMFITFVLSHGEEGRFITSDDKEMKFEELWGHFRGSQFKPLIGKPKLFFFVCCQGLRYNQTVCAD

EITEDSETIEFDSISSPSGHPDIADFFFGYATYKSLRSFRSTLNGTFYVKFLCASINENPKKSIHDIHIEVQRSLAKSKI

PQVNSFQMAYYENLLREHCCFAI

>VDN15196.1 unnamed protein product [Dibothriocephalus latus]CARD

MEKQHVEALRKHRVELVRDVHSVEEILDHLFGEGIISDNQQCLILAGTTPAQKVRLLLDTLVRCGPKAFTSFCSALKATK

QFHLLELLETSQPTEESAEYRPPPDKQPASTGIMQADEAHGTSIVQQNYSGPRKPPSQYVISPYPVCSRPLGYILIVNIV

EFAPACRLKPRLGSRKDAEGLVRLFRDIGYSVEVLENPDTPTLLDSLEAYAAKPEHASVDAGGMVLMSHGLRDHIYTSEG

HLVSVEKLLSYFTNKALPGLAGKPKLLILQACRGRAQDFGTSNTDYVVRRDTFKPADAPVSNSSWGSLPYMSDCVIAYST

LPGFVSWRSEDEGSWFIQVLLDVFRRLGNKLHVIDLLAEVNRIIVEQSQDQVCKQISQPATNLRRPFFLCTRDAS
